# First description of a *Yersinia pseudotuberculosis* clonal outbreak in France, confirmed using a new core genome multilocus sequence typing method

**DOI:** 10.1101/2022.03.23.485572

**Authors:** Cyril Savin, Anne-Sophie Le Guern, Fanny Chereau, Julien Guglielmini, Guillaume Heuzé, Christian Demeure, Javier Pizarro-Cerdá

**Affiliations:** Institut Pasteur, Université de Paris Cité, Yersinia Research Unit, F-75015, Paris, France; Institut Pasteur, Université de Paris Cité, Yersinia National Reference Laboratory, F-75015, Paris, France; Institut Pasteur, Université de Paris Cité, WHO Collaborating Research & Reference Centre for Yersinia FRA-140, F-75015, Paris, France; French National Public Health Agency, Department of Infectious Diseases, F-94410, Saint-Maurice, France; Institut Pasteur, Université de Paris Cité, Hub de Bioinformatique et Biostatistique, F-75015, Paris, France; French National Public Health Agency, Corse, F-20000, Ajaccio, France

**Keywords:** Yersinia pseudotuberculosis, outbreak, epidemiological investigation, cgMLST, enteric yersiniosis

## Abstract

*Yersinia pseudotuberculosis* is an enteric pathogen causing mild enteritis that can lead to mesenteric adenitis and septicemia in elderly patients. Most cases are sporadic, but outbreaks have already been described in different countries. We report for the first time a *Y. pseudotuberculosis* clonal outbreak in France, that occurred in 2020. An epidemiological investigation pointed towards the consumption of tomatoes as the likely source of contamination. The *Yersinia* National Reference Laboratory (YNRL) developed a new cgMLST scheme with 1,921 genes specific to *Y. pseudotuberculosis* that identified the clustering of isolates associated to the outbreak and allowed to perform molecular typing in real time. In addition, this method allowed to retrospectively identify isolates belonging to this cluster from earlier in 2020. This method, which does not require specific bioinformatic skills, is now used systematically at the YNRL and proves to display an excellent discriminatory power and is available to the scientific community.

## Introduction

The *Yersinia* genus encompasses 26 different species. Two of them are enteropathogenic for humans: *Yersinia enterocolitica* and *Yersinia pseudotuberculosis* (1). Occurring predominantly in children, this latter species can cause mild enteritis characterized by fever, abdominal pain, and sometimes diarrhea that can lead to mesenteric adenitis (2). *Y. pseudotuberculosis* (*Yptb*) can cause an invasive infection, leading to bacteremia in elderly patients or in individuals presenting underlying medical disorders (diabetes, cirrhosis, iron overload) (3). Most of the *Yptb* associated-cases are sporadic, but some outbreaks have been reported in different parts of the world, including Japan (4), Canada (5), Europe (6), Russia (7) and more recently in New Zealand (8). The reservoir of *Yptb* is mostly wild mammals (particularly rodents, lagomorphs, wild boars) and birds. The pathogen can enter the food chain and outbreaks caused by consumption of contaminated iceberg lettuce (9), carrots (10) or raw milk (11) have been described.

In case of outbreak suspicion, epidemiological investigations are of key importance to establish a link between patients and to identify a common exposure. Molecular investigation methods allowing the establishment of a genetic link between bacterial isolates have been essential in many outbreak investigations to confirm a genetic link between clinical isolates or between clinical and environmental isolates. Pulsed-field gel electrophoresis (PFGE) used to be the gold standard technique (9). However, this method is time-consuming and labor-intensive. Its lack of reproducibility and resolution led to its replacement by multilocus variable-number tandem repeat analysis (MLVA) which has a better discriminatory power, but is still time-consuming (12). Recent advances in sequencing methods have made whole-genome sequencing a rapid and affordable approach, available to surveillance laboratories. This has led to the development of core-genome Single Nucleotide Polymorphism (cgSNP) analyses to determine the genetic distance between bacterial isolates, with an excellent discriminatory power (8, 13). Nevertheless, cgSNP analyses require advanced bioinformatic skills and is not yet standardized between laboratories.

In France, the surveillance of enteric yersiniosis is conducted by the *Yersinia* National Reference Laboratory (YNRL) and Santé publique France (SpF), the national public health agency. The routine procedure at the YNRL includes whole-genome sequencing of all bacterial isolates, followed by a bioinformatic analysis using a 500-gene core genome multilocus sequence typing (cgMLST) dedicated to the *Yersinia* genus (14), which allows to identify the species and eventually the lineage. Due to its relative low number of genes, this technique is not used to detect clusters. Therefore, the YNRL developed a new and easy-to-use public tool for their identification. This method is as powerful as other bioinformatic tools based on whole-genome sequencing such as cgSNP analysis and does not require specific bioinformatic skills. It is based on a cgMLST of 1,921 genes shared by most *Yptb* strains. We hereby describe this tool and highlight its usefulness in the investigation of the first described *Yptb* outbreak in France.

## Methods

### *Y. pseudotuberculosis* isolates and taxonomic assignment

*Yersinia* isolates, together with some clinical and demographic data, are regularly sent to the French YNRL for enteric yersiniosis by medical laboratories for complete characterization. Isolation and taxonomic assignments are performed as described by Savin *et al*. (14) based on a 500-gene cgMLST scheme designed to identify all the species of the *Yersinia* genus, as well as the lineage. A total of 359 isolates of *Yptb* received at the YNRL between 1991 and 2020 for complete characterization were genotypically assigned. In addition, genomic data of 9 isolates of *Yptb* lineage 16 with a clinical origin (1969 to 1990) were extracted from our database for comparison of their genetic relatedness.

### A novel cgMLST as a tool for identification of *Y. pseudotuberculosis* clusters

From our 1,346 *Yersinia* reference genomes dataset, constituted for the phylogenetic analysis of the *Yersinia* genus (14), 485 genomes representative of the *Yptb* species diversity were selected. Selection of the genes was performed as described previously (14) and led to the selection of 1,421 *Yptb* core genes to which the 500 genes of the *Yersinia* spp cgMLST scheme (14) were added, resulting in 1,921 core genes deemed suitable for cgMLST analysis and molecular investigation (Table S1).

A database was created for *Yptb* in the Institut Pasteur’s MLST and whole-genome MLST resource (https://bigsdb.pasteur.fr), which uses the BIGSdb software tool (15, 16). All *de novo* assembled genomes were uploaded into the isolates database, and the reference alleles of the 1,921 cgMLST loci were defined in the linked database of reference sequences and profiles (‘seqdef’). Within BIGSdb, a scan of the genome sequence was performed for each isolate using parameters (min 80□% identity, min 80□% alignment, blastn word size of 20 nt) to check for the presence of each core gene and to determine its allele number. The BIGSdb –

*Yptb* database of cgMLST profiles is accessible at https://bigsdb.pasteur.fr/yersinia/. A comparison of the allelic profiles can be performed either with the ‘Genome comparator’ plugin or by the construction of a minimum spanning tree (MST) with GrapeTree (17) using the corresponding BIGSdb plugin.

Since 2018 at the YNRL, each isolate identified as *Yptb* is also submitted routinely to this new cgMLST to evaluate its genetic distance with other isolates from the database. When a cluster of isolates (≤ 5 allelic difference) is identified, the YNRL alerts SpF who determines whether an epidemiological investigation is required.

### Core-genome SNP analysis

Genome sequences of all the isolates and paired-end quality-filtered FASTQ files were obtained as described by Savin *et al*. (14). Variant calling was performed using the IP32953 reference strain (accession number: NC_006155) with Snippy version 4.6.0 and core-SNPs were extracted using snp-sites (https://github.com/tseemann/snippy). A comparison of the isolates using the core-SNPs was performed by the construction of a MST with GrapeTree (17).

### Discriminatory power determination

The discriminatory power of the molecular typing method was determined using the Simpson’s index of diversity (ID). It calculates the probability of a technique to attribute the same profile to epidemiologically unrelated isolates. The higher the index is, the better the discriminatory power is (18).

### Epidemiological, trace-back, and environmental investigations

The YNRL alerts SpF of any unusual signal of *Yptb* from the microbiological surveillance, including clusters, to determine whether an epidemiological investigation is required. For this outbreak, all patients corresponding to the outbreak case definition were contacted by SpF and queried about their previous exposition to animals, visits in natural areas (sea, lake, forest, river, farms), drinking water supply and food consumption (dairy products, meats, fresh vegetables), using a standard trawling questionnaire. The questionnaire covered the 10 days before the onset of the symptoms. Places of travel (holiday period) were also recorded. Historical records of tap water quality were verified by the regional Health Agency (ARS de Corse) and trace-back investigation of suspected foods were performed by the French Directorate General for Food (DGAl) and the General Directorate for Competition Policy, Consumer Affairs and Fraud Control (DGCCRF).

## Results

### Historical diversity of *Y. pseudotuberculosis* isolates in France

According to the French YNRL database, among the 324 *Yptb* isolates received between 1991 and 2019, seventeen lineages currently circulate in France: isolates from lineages 15 and 10 are the most frequent (76 and 70 isolates respectively), followed by lineages 17, 5, 7, 2 and 16 (Figure S1). Even if the number of isolates has been quite stable over time (11.2 ±5.2 per year since 1991), very few strains were reported in 1997 and 2002 (2 isolates each year) while a peak was observed in 2005 (28 isolates) (Figure S1).

### Suspicion of a *Y. pseudotuberculosis* outbreak in France during 2020

In 2020, enteric yersiniosis surveillance by the YNRL led to the identification of 35 isolates of *Yptb*. Their geographical distribution indicates that, for almost all the lineages, isolates originated from different French departments (Figure 1.A). The 20 ones identified in the first semester belonged to 9 different lineages, lineage 10 being the most frequently isolated (5 isolates) whereas 3 specimens of each lineage 15 and 16 were found (Figure 1.B). At the end of July, 3 additional lineage 16 isolates were identified. The isolation of these specimens by a single medical laboratory in Porto-Vecchio (Corse-du-Sud) led the YNRL to alert SpF (see below) of a potential outbreak concerning *Yptb* lineage 16 (Figure S2). In August 2020, 4 more lineage 16 isolates were identified, from the same laboratory, together with 3 other isolates (lineage 10 and 15). In September 2020, 2 lineage 16 isolates were identified but they originated from a laboratory located in Lyon (Rhône). Afterwards, no more isolates from lineage 16 were reported in 2020. Interestingly, in addition to the 8 lineage 16 isolates identified during the summer, 3 lineage 16 specimens were found, 2 by the same laboratory in Corse-du-Sud, and one in Dijon (Côte d’Or) at the beginning of the year. Whereas no *Yptb* were identified in October and November, 4 isolates belonging to 3 different lineages (7, 10 and 15) appeared in December (Figure S2).

**Figure 1:**
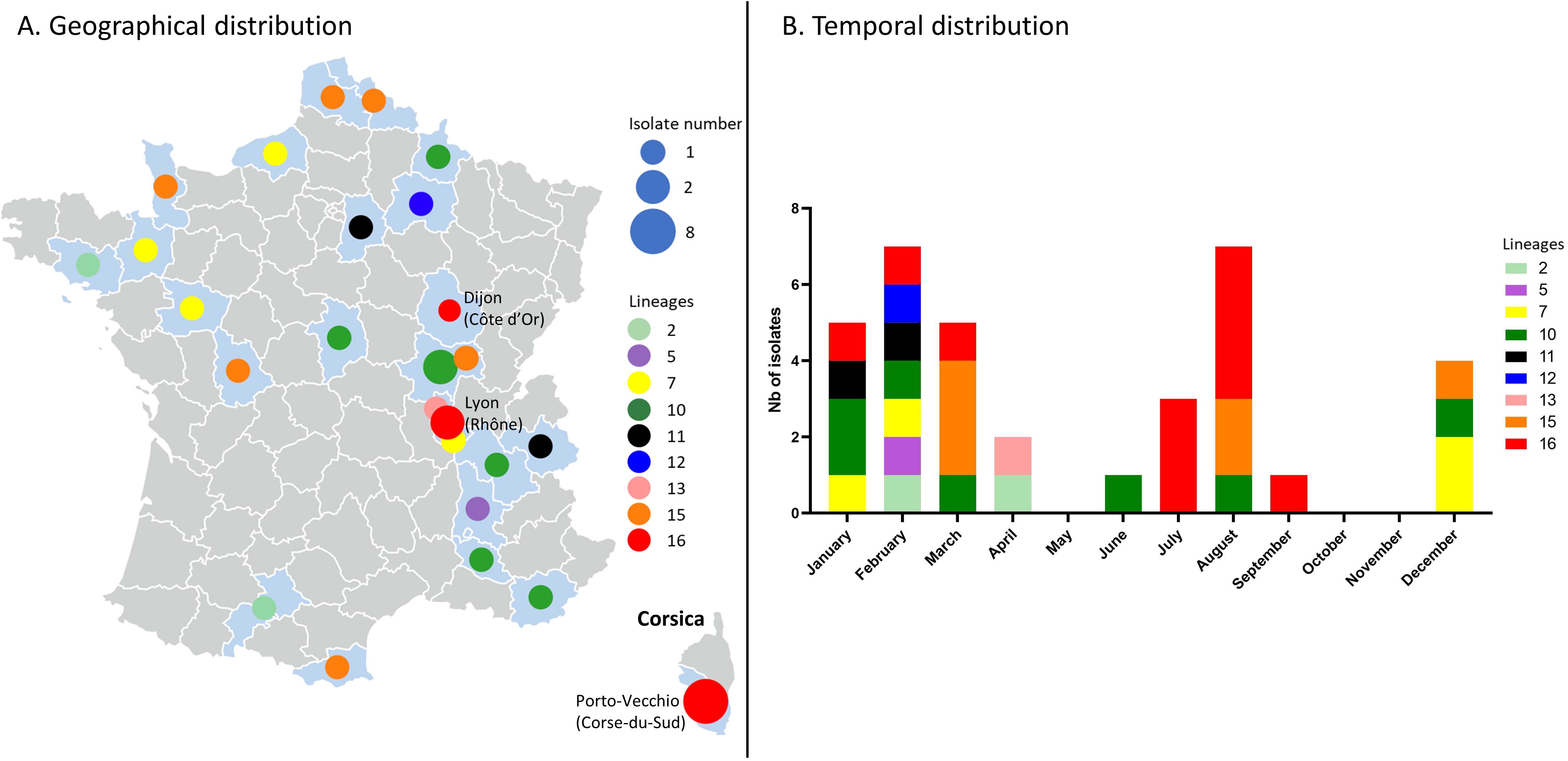
Geographical and temporal distribution of the 35 *Y. pseudotuberculosis* isolated in France in2020. (A) Map of France with the departments. Size of the circle depends on the number of isolates. Colors of the circles depends on the isolates’ lineages. (B) Number of strains per month. Colors of the squares depends on the isolates’ lineages.

### Epidemiological, trace back and environmental investigations

On August 12, 2020, the YNRL informed SpF of the identification of 3 patients infected by *Yptb* lineage 16 as determined by a cgMLST 500 genes, isolated in the same week (week 30) in a single medical laboratory in Corsica (Figure S2). By comparison, 0 to 2 isolates belonging to lineage 16 had been isolated per year in France since 1991 (Figure S1). Moreover, no *Yptb* had been isolated the previous years by the medical laboratory, while already using the same identification method. This unusual temporal and geographical group of cases, combined with the potential for invasive infections by *Yptb*, instigated an epidemiological investigation led by SpF and local public health authorities, to identify a potential common source of contamination and to implement control measures.

Cases were defined as any patient with identification of a *Yptb* lineage 16 isolate in the YNRL national database, from any type of specimen sampled from July 1^st^ in metropolitan France. In total, 8 cases were identified with sampling dates between July 23^rd^ and September 1^st^. The 8 *Yptb* specimens were recovered from stool samples in a laboratory in Porto-Vecchio (Corse-du-Sud department) for 6 patients and in a laboratory in Lyon (Rhône department) for 2 patients. The median age of the patients was 25 years old, with 4 patients between 5 to 15, 3 patients between 30 to 60 and 1 patient older than 90. The patients sex ratio was 1.7 (three women and five men). The 8 cases were interviewed: the onset of the symptoms covered a period from July 10^th^ to August 26^th^. Most patients have managed their symptoms at home, one 10-year-old child was admitted to the hospital during one night for observation. Two patients were residents of Corsica and 6 were in holidays in Corsica during the incubation period. Moreover, 7 of them were located (residency or holidays) in a 10km radius area in Northeast of Corsica (Haute-Corse department).

Food queries pointed towards the consumption of tomatoes from the same grocery store in Northeast of Corsica (6 cases). No other common consumption of food nor leisure activity was identified. Seven of the cases resided in an area supplied by the same water distribution network. No contamination episodes of the water distribution network covering the area were identified in the historical records (15 campaigns in 2019).

Food investigation established that the suspected tomatoes originated from a local production unit, based in the same geographical area. On-site inspection did not identify any non-conformity potentially leading to contamination of the tomatoes during production, harvest, storage, packaging, or transport, nor any problem with traceability. Three other companies commercialized tomatoes from the suspected batch, but no trace back could be performed as no sales records were kept. The tomatoes were not rinsed before distribution and the water used for irrigation came from the public distribution network for agricultural use. No verification of the quality of this water system was conducted.

### A new *Y. pseudotuberculosis* cgMLST confirmed the cluster of isolates

As our newly developed 1,921 genes cgMLST scheme specific to the species *Yptb* is used in routine at the YNRL, identification of the cluster of isolates associated to this pseudotuberculosis outbreak was possible. This new typing method was also applied to the *Yptb* strains isolated in France in 2020 and their genetic relatedness was determined (Table S2 and Figure 2).

**Figure 2:**
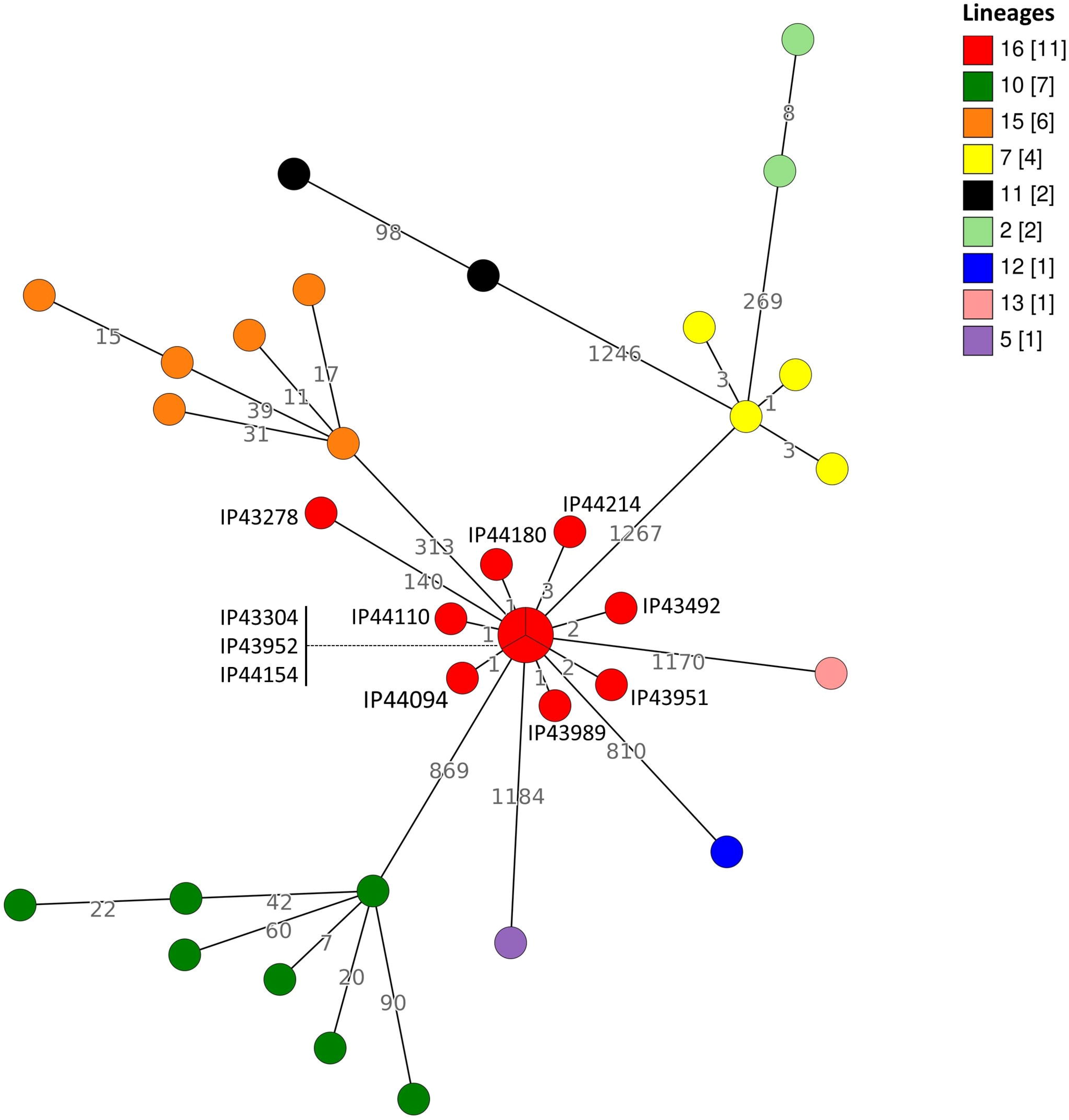
Minimum spanning tree obtained using the allelic profiles of the cgMLST (1921 genes) on the 35 *Y. pseudotuberculosis* isolates in France in 2020. The branch lengths are based on a logarithmic scale. Numbers close to the branches reveals the alleles differences. Colors of the circles depends on the isolates’ lineages.

The observed distance between isolates within each lineage (0 to 140 alleles) is much lower than distances observed between isolates from different lineages (269 to 1267 alleles), confirming that the lineages are well demarcated from each other using this novel cgMLST (Figure 2).

Whereas the distances between isolates from lineages 10 and 15 are higher, lineages 7 and 16 display isolates more closely related to each other, suggesting more clonality. Lineage 7 isolates (4 specimens) have 1 to 3 allele differences and may be considered belonging to the same cluster. However, no interviews of these patients were conducted to identify a potential common exposure and their distant isolation dates (Figure 1 and 2) weaken the hypothesis of a common source of contamination.

Interestingly, among the 11 lineage 16 isolates, 10 of them showed close genetic relatedness (between 0 and 3 differences) and may be considered as belonging to the same cluster. Whereas isolates IP43304 and IP43492 were recovered at the beginning of 2020, the 8 other specimens were isolated within 38 days (from 28^th^ of July to 3^rd^ of September 2020) and correspond to the cases investigated during the summer (see above). The close genetic relatedness of the 8 isolates, together with their close geographical and temporal isolation, confirmed a cluster of cases due to a *Yptb* lineage 16 infection. Interestingly, the 2 isolates IP43304 and IP43492 were also recovered from the laboratory in Porto-Vecchio in February and March 2020. No interviews of these two patients from early 2020 on the exposures were conducted, given the distance to onset of symptoms. The high allele difference number between isolate IP43278 and the other isolates (≥140) from lineage 16 (Figure 2) excludes IP43278 from the cluster.

### Performance comparison between the novel *Y. pseudotuberculosis* cgMLST and classical cgSNP analysis

Taking advantage of our novel, easy-to-use *Yptb* cgMLST scheme, we compared the performance of both methods on 39 lineage-16 clinical isolates from our French database (1969 – 2020). Minimum spanning trees (MST) were reconstructed with both cgMLST data and cgSNP analyses (Figure 3). cgMLST-based MST allows to determine the genetic distance in terms of allelic distance. Among the 39 studied isolates, 37 cgMLST profiles were identified. Only 3 isolates belonged to the same cgMLST profile, and they corresponded to specimens from the 2020 Corsica outbreak (Figure 3.A). Pairwise allelic distances between the 39 isolates (table S3) confirm that 3 isolates have the same cgMLST profile (IP43304, IP43952 and IP44154) whereas other isolates have allelic distances between 1 and 157 (table S3).

**Figure 3:**
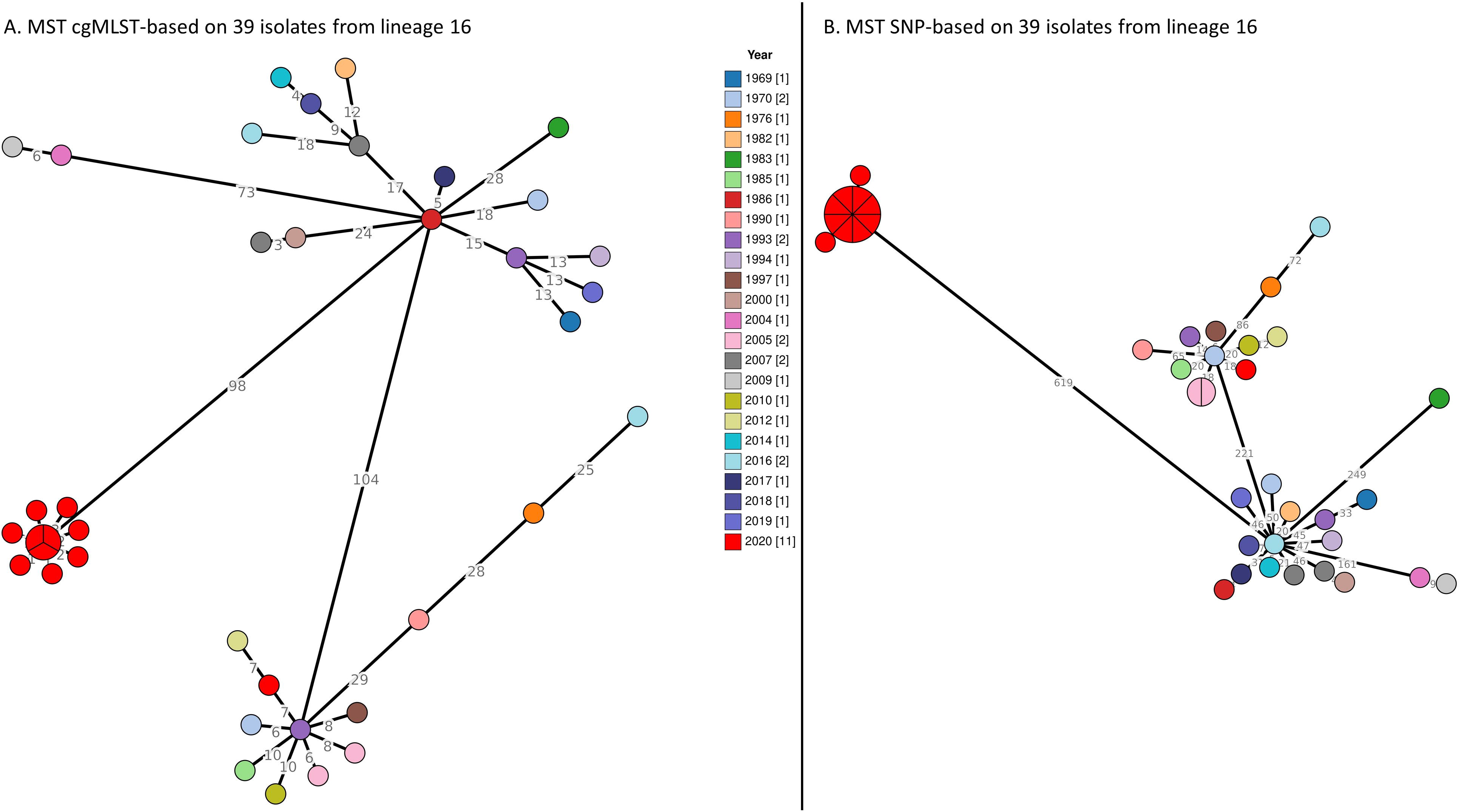
Minimum spanning tree reconstructed on the 39 *Y. pseudotuberculosis* belonging to the lineage 16 isolated in France, 1969-2020. (A) MST cgMLST-based (B) MST SNP-based. The branch lengths are based on a linear scale. Numbers close to the branches reveals the alleles differences (A) or SNP differences (B). Colors of the circles depends on the isolates’ lineages.

The cgSNP-based MST (Figure 3.B) allows to determine the genetic distance in terms of point mutations (SNPs). On this tree, we can observe 31 different SNP profiles: 2 isolates from 2005 have the same SNP profile and 8 isolates (IP43304, IP43951, IP43952, IP43989, IP44094, IP44110, IP44154 and IP44180) from the 2020 outbreak have the same profile. Pairwise SNP comparisons (Table S3) confirm the null distance between those isolates, whereas other isolates have between 1 and 802 SNP distances.

This lower number of profiles obtained with the cgSNP-based MST indicates that some of the isolates with different cgMLST profiles have merged into a single cgSNP profile. Simpson’s index of diversity estimation for cgMLST is 0.996, whereas for cgSNP analysis is 0.96. Our cgMLST proved to have a better discriminatory power than cgSNP.

## Discussion

We report here for the first time a *Yptb* clonal outbreak in France, with 8 cases identified during the summer 2020. All cases had been exposed in the same area in Corsica and consumption of local tomatoes was the suspected source of contamination. A new cgMLST confirmed that the 8 cases belonged to the same cluster. Moreover, two earlier cases (February and March 2020, both detected in Corsica) were also identified as belonging to the same cluster (although they could not be interviewed on their exposures and no common exposure with the summer cases could be identified).

The number of cases reported during the outbreak is low. However, incidence of pseudotuberculosis is also low in France, with an average number of 11 isolates per year (Figure S1). This incidence is probably underestimated: all symptomatic patients do not visit their doctor and they rarely prescribe stool examinations in case of diarrhea with no complications. In addition, notification of *Yptb* or transmission of the isolates to the YNRL are not mandatory. Furthermore, detection and isolation of *Yptb* in medical laboratories is difficult: a slower growth rate as compared to other enterobacteria and the presence of competitive microbiota renders *Yersinia* spp. isolation complex (19, 20). Growth of some *Yptb* strains is impaired on semi-selective CIN agar (21). The recent implementation of panel-based testing systems (i.e. multiplex PCR) targeting enterobacteria could alleviate this issue, leading to stool culture only when a PCR positive signal is obtained (22, 23). However, some PCR kits target only *Y. enterocolitica* specific chromosomal genes, reinforcing the low identification rate of *Yptb*.

Epidemiological investigation pointed tomatoes as the suspected source of contamination source for the summer cases. However, as no sampling of tomatoes was performed, this suspected source could not be confirmed. Contaminated vegetables were also suspected in previously reported *Yptb* outbreaks, with iceberg lettuce in 1998 (9) and carrots in 2006 (10) in Finland were confirmed as sources of contamination. As wildlife is considered the *Yptb* reservoir, feces of carrier animals may contaminate environmental water, soil, and grass (24). Contamination of vegetables in the fields can be direct (feces) or indirect (irrigation with contaminated water). Wild boars and pigs are recognized as reservoirs of *Yptb* (25): as Corsica hosts a large population of wild boars and allows the wandering of pigs, it is possible to hypothesize a contamination of vegetables in the fields from this reservoir. Wild rodents may also contaminate vegetables in the fields or during storage. Carrier animals may contaminate their environment as long as they host the pathogen, possibly leading to several episodes or sources of contamination (26).

An increase in *Yptb* cases had been observed and investigated in 2005 in France (Figure S1 and 1). However, the epidemiological investigation identified a high genetic diversity in the isolates as well as the absence of a geographically defined cluster. This increase in clinical cases has been linked to an increase in prevalence in rodent reservoirs (3).

Identification of outbreak-related isolates and trace-back investigations to identify a potential source of contamination were difficult when techniques such as PFGE (9) or MLVA (12) were used. Depending on the pathogen, the discriminatory power of PFGE may differ and not be optimal (27-29) and its lack of reproducibility between laboratories restricts its use to retrospective epidemiology (30, 31). PFGE has often been replaced by MLVA, which has proven to display a better discriminatory power (29, 32) but is still labor-intensive and time-consuming. Development of whole-genome-based typing methods alleviates these issues and allows rapid detection of clusters as well near-real time alerts of public health authorities. The confirmation of genetic relatedness of clinical and food samples remains a strong lever for recalling food products from the market. Rapid trace-back investigation strengthens the possibility to identify a common source of contamination and to remove it from the food chain (33).

Different cgMLST schemes have been developed for foodborne pathogen identification. They have proven to be useful in public health surveillance and have provided tools allowing international collaboration (13, 34, 35). Discriminatory power comparisons between cgMLST and cgSNP analysis have shown a very high discriminatory power for both methods, thus arguing for the use of whole-genome-based methods for epidemiological investigation (Figure 3) (13). The comparison confirmed the better performance and resolution power of our novel cgMLST specific to *Yptb*. cgSNP analysis has already been used in the investigation of an outbreak due to *Yptb* infection. This tool allowed the identification of a point-source contamination in the food chain (8). However, it requires advanced bioinformatic skills not widely available in National Reference laboratories worldwide. In this framework, we developed a new cgMLST for *Yptb* that proved to be more discriminant than cgSNP analysis (Figure 3 and Table S2). Here, allelic distance identification was very useful as it confirms that the 8 summer isolates belong to the same cluster with 0 to 3 alleles difference and suggests a persistent or recurrent contamination of the food chain as 2 isolates were identified in February and March 2020. Interestingly, lineage 16 specimens are absent in previous years samplings (Figure 3.A) suggesting that this clone emerged recently.

Our new cgMLST does not require additional laboratory manipulation and is usable in real-time after the identification of the bacterial species, as it only requires the genome assembly of the isolate. Furthermore, it relies on the simple comparison of allelic profiles and should help future international collaboration to determine whether a clone is circulating in several countries.

We report, for the first time in France, an outbreak of *Yptb* infections due to the same clone. Epidemiological and microbiological investigations established a link between the patients and identified the consumption of tomatoes from a unique grocery store as the suspected source of contamination. Our recently developed cgMLST (available for the community at https://bigsdb.pasteur.fr/yersinia/) exhibits an excellent discriminatory power and allows epidemiological investigation in real-time.

## Acknowledgements

This project received funding from Santé publique France (Saint-Maurice, France) and the LabEX Integrative Biology of Emerging Infectious Diseases (ANR LBX-62 IBEID). This work used the computational and storage services (MAESTRO cluster) provided by the IT department at the Institut Pasteur, Paris. We acknowledge the continuous support of Keith Jolley (Oxford University) for the development of the BIGSdb web application.

## Supplemental Material

Table S1: List of the 1,921 genes used for this cgMLST.

Table S2: Allelic profiles of the 35 isolates from 2020 in France.

Table S3: Pairwise distance matrix cgMLST-based and SNP-based obtained comparing the 39 *Y. pseudotuberculosis* isolates belonging to the lineage 16.

Figure S1: Repartition of the different *Y. pseudotuberculosis* lineages in France according to the year of isolation.

Figure S2: Timeline of the lineage 16 isolates during summer 2020. Number between brackets correspond to the isolation month.

## References

1. Le Guern AS, Savin C, Angermeier H, Bremont S, Clermont D, Muhle E, et al. Yersinia artesiana sp. nov., Yersinia proxima sp. nov., Yersinia alsatica sp. nov., Yersina vastinensis sp. nov., Yersinia thracica sp. nov. and Yersinia occitanica sp. nov., isolated from humans and animals. Int J Syst Evol Microbiol. 2020 Oct;70(10):5363–72.

2. Tertti R, Vuento R, Mikkola P, Granfors K, Makela AL, Toivanen A. Clinical manifestations of Yersinia pseudotuberculosis infection in children. Eur J Clin Microbiol Infect Dis. 1989 Jul;8(7):587–91.

3. Vincent P, Leclercq A, Martin L, Yersinia Surveillance N, Duez JM, Simonet M, et al. Sudden onset of pseudotuberculosis in humans, France, 2004-05. Emerg Infect Dis. 2008 Jul;14(7):1119–22.

4. Inoue M, Nakashima H, Ueba O, Ishida T, Date H, Kobashi S, et al. Community outbreak of Yersinia pseudotuberculosis. Microbiol Immunol. 1984;28(8):883–91.

5. Nowgesic E, Fyfe M, Hockin J, King A, Ng H, Paccagnella A, et al. Outbreak of Yersinia pseudotuberculosis in British Columbia--November 1998. Can Commun Dis Rep. 1999 Jun 1;25(11):97–100.

6. Jalava K, Hallanvuo S, Nakari UM, Ruutu P, Kela E, Heinasmaki T, et al. Multiple outbreaks of Yersinia pseudotuberculosis infections in Finland. J Clin Microbiol. 2004 Jun;42(6):2789–91.

7. Somova LM, Antonenko FF, Timchenko NF, Lyapun IN. Far Eastern Scarlet-Like Fever is a Special Clinical and Epidemic Manifestation of Yersinia pseudotuberculosis Infection in Russia. Pathogens. 2020 Jun 2;9(6).

8. Williamson DA, Baines SL, Carter GP, da Silva AG, Ren X, Sherwood J, et al. Genomic Insights into a Sustained National Outbreak of Yersinia pseudotuberculosis. Genome Biol Evol. 2016 Dec 1;8(12):3806–14.

9. Nuorti JP, Niskanen T, Hallanvuo S, Mikkola J, Kela E, Hatakka M, et al. A widespread outbreak of Yersinia pseudotuberculosis O:3 infection from iceberg lettuce. J Infect Dis. 2004 Mar 1;189(5):766–74.

10. Rimhanen-Finne R, Niskanen T, Hallanvuo S, Makary P, Haukka K, Pajunen S, et al. Yersinia pseudotuberculosis causing a large outbreak associated with carrots in Finland, 2006. Epidemiol Infect. 2009 Mar;137(3):342–7.

11. Parn T, Hallanvuo S, Salmenlinna S, Pihlajasaari A, Heikkinen S, Telkki-Nykanen H, et al. Outbreak of Yersinia pseudotuberculosis O:1 infection associated with raw milk consumption, Finland, spring 2014. Euro Surveill. 2015;20(40).

12. Halkilahti J, Haukka K, Siitonen A. Genotyping of outbreak-associated and sporadic Yersinia pseudotuberculosis strains by novel multilocus variable-number tandem repeat analysis (MLVA). J Microbiol Meth. 2013 Nov;95(2):245–50.

13. Vincent C, Usongo V, Berry C, Tremblay DM, Moineau S, Yousfi K, et al. Comparison of advanced whole genome sequence-based methods to distinguish strains of Salmonella enterica serovar Heidelberg involved in foodborne outbreaks in Quebec. Food Microbiol. 2018 Aug;73:99–110.

14. Savin C, Criscuolo A, Guglielmini J, Le Guern AS, Carniel E, Pizarro-Cerda J, et al. Genus-wide Yersinia core-genome multilocus sequence typing for species identification and strain characterization. Microb Genom. 2019 Oct;5(10).

15. Jolley KA, Bray JE, Maiden MCJ. Open-access bacterial population genomics: BIGSdb software, the http://PubMLST.org website and their applications. Wellcome Open Res. 2018;3:124.

16. Jolley KA, Maiden MC. BIGSdb: Scalable analysis of bacterial genome variation at the population level. BMC Bioinformatics. 2010 Dec 10;11:595.

17. Zhou Z, Alikhan NF, Sergeant MJ, Luhmann N, Vaz C, Francisco AP, et al. GrapeTree: visualization of core genomic relationships among 100,000 bacterial pathogens. Genome Res. 2018 Sep;28(9):1395–404.

18. Hunter PR, Gaston MA. Numerical index of the discriminatory ability of typing systems: an application of Simpson’s index of diversity. J Clin Microbiol. 1988 Nov;26(11):2465–6.

19. Schiemann DA, Olson SA. Antagonism by gram-negative bacteria to growth of Yersinia enterocolitica in mixed cultures. Appl Environ Microbiol. 1984 Sep;48(3):539–44.

20. Tan LK, Ooi PT, Carniel E, Thong KL. Evaluation of a modified Cefsulodin-Irgasan-Novobiocin agar for isolation of Yersinia spp. PLoS One. 2014;9(8):e106329.

21. Fukushima H, Gomyoda M. Growth of Yersinia pseudotuberculosis and Yersinia enterocolitica biotype 3B serotype O3 inhibited on cefsulodin-Irgasan-novobiocin agar. J Clin Microbiol. 1986 Jul;24(1):116–20.

22. Engberg J, Vejrum LK, Madsen TV, Nielsen XC. Verification of analytical bacterial spectrum of QIAstat-Dx(R) GI V2 and Novodiag(R) Bacterial GE+ V2-0 diagnostic panels. J Antimicrob Chemother. 2021 Sep 23;76(Supplement_3):iii50–iii7.

23. Schaumburg F, Frobose N, Kock R. A comparison of two multiplex-PCR assays for the diagnosis of traveller’s diarrhoea. BMC Infect Dis. 2021 Feb 16;21(1):181.

24. Mair NS. Yersiniosis in wildlife and its public health implications. J Wildl Dis. 1973 Jan;9(1):64–71.

25. Reinhardt M, Hammerl JA, Kunz K, Barac A, Nockler K, Hertwig S. Yersinia pseudotuberculosis Prevalence and Diversity in Wild Boars in Northeast Germany. Appl Environ Microbiol. 2018 Sep 15;84(18).

26. Backhans A, Fellstrom C, Lambertz ST. Occurrence of pathogenic Yersinia enterocolitica and Yersinia pseudotuberculosis in small wild rodents. Epidemiol Infect. 2011 Aug;139(8):1230–8.

27. Gilpin BJ, Robson B, Lin S, Hudson JA, Weaver L, Dufour M, et al. The limitations of pulsed-field gel electrophoresis for analysis of Yersinia enterocolitica isolates. Zoonoses Public Health. 2014 Sep;61(6):405–10.

28. Magistrali CF, Cucco L, Pezzotti G, Farneti S, Cambiotti V, Catania S, et al. Characterisation of Yersinia pseudotuberculosis isolated from animals with yersiniosis during 1996-2013 indicates the presence of pathogenic and Far Eastern strains in Italy. Vet Microbiol. 2015 Oct 22;180(1-2):161–6.

29. Sihvonen LM, Toivonen S, Haukka K, Kuusi M, Skurnik M, Siitonen A. Multilocus variable-number tandem-repeat analysis, pulsed-field gel electrophoresis, and antimicrobial susceptibility patterns in discrimination of sporadic and outbreak-related strains of Yersinia enterocolitica. BMC Microbiol. 2011 Feb 25;11:42.

30. Chiou CS, Wei HL, Yang LC. Comparison of pulsed-field gel electrophoresis and coagulase gene restriction profile analysis techniques in the molecular typing of Staphylococcus aureus. J Clin Microbiol. 2000 Jun;38(6):2186–90.

31. Zhou H, Liu W, Qin T, Liu C, Ren H. Defining and Evaluating a Core Genome Multilocus Sequence Typing Scheme for Whole-Genome Sequence-Based Typing of Klebsiella pneumoniae. Front Microbiol. 2017;8:371.

32. Hu Y, Li B, Jin D, Cui Z, Tao X, Zhang B, et al. Comparison of multiple-locus variable-number tandem-repeat analysis with pulsed-field gel electrophoresis typing of Acinetobacter baumannii in China. J Clin Microbiol. 2013 Apr;51(4):1263–8.

33. Jackson BR, Tarr C, Strain E, Jackson KA, Conrad A, Carleton H, et al. Implementation of Nationwide Real-time Whole-genome Sequencing to Enhance Listeriosis Outbreak Detection and Investigation. Clin Infect Dis. 2016 Aug 1;63(3):380–6.

34. Moura A, Criscuolo A, Pouseele H, Maury MM, Leclercq A, Tarr C, et al. Whole genome-based population biology and epidemiological surveillance of Listeria monocytogenes. Nat Microbiol. 2016 Oct 10;2:16185.

35. van den Beld MJC, Reubsaet FAG, Pijnacker R, Harpal A, Kuiling S, Heerkens EM, et al. A Multifactorial Approach for Surveillance of Shigella spp. and Entero-Invasive Escherichia coli Is Important for Detecting (Inter)national Clusters. Front Microbiol. 2020;11:564103.

