## Supplementary figures and images for "First description of a *Yersinia pseudotuberculosis* clonal outbreak in France, confirmed using a new core genome multilocus sequence typing method"

### Figure_S1

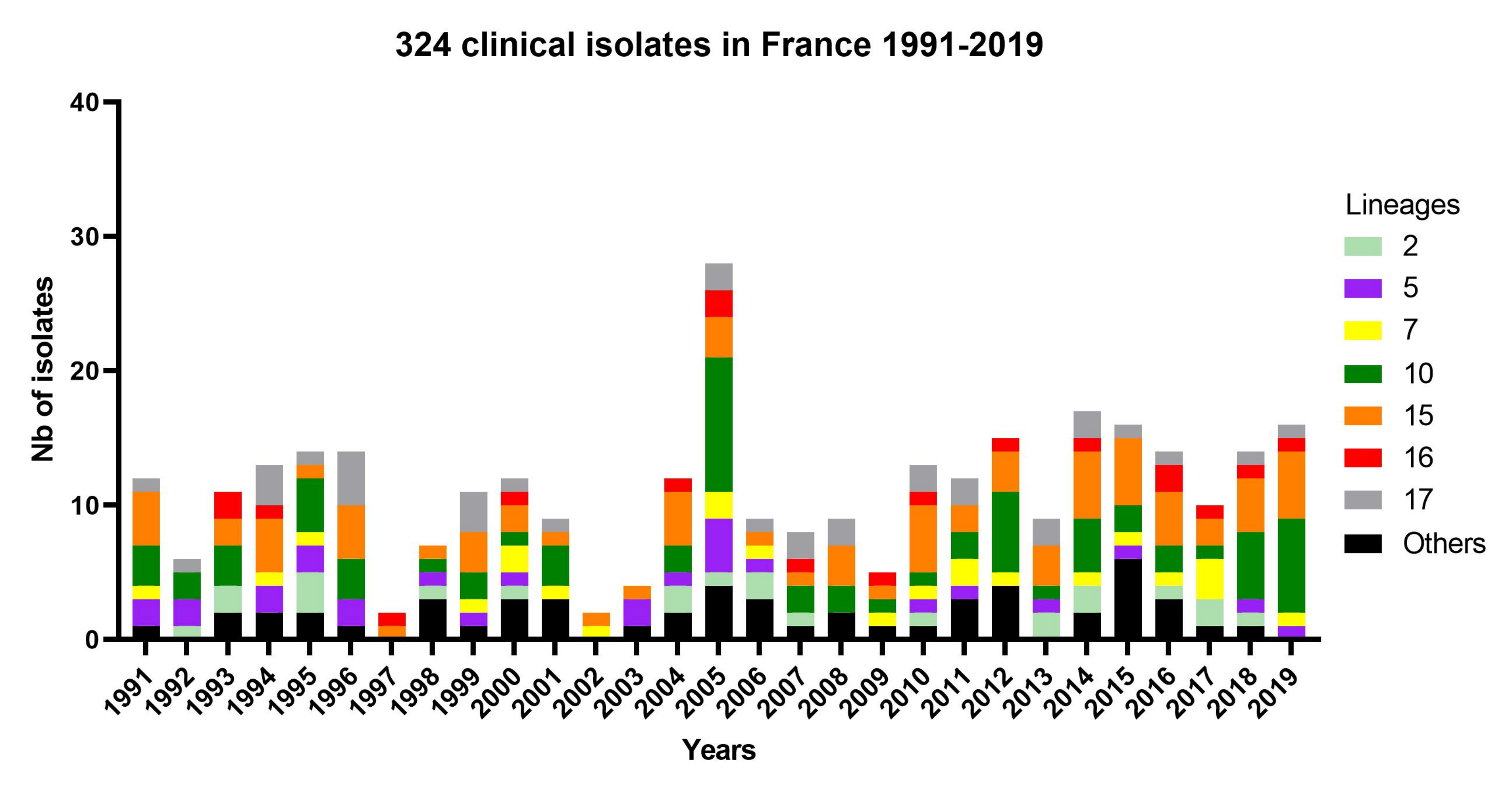

### Figure_S2

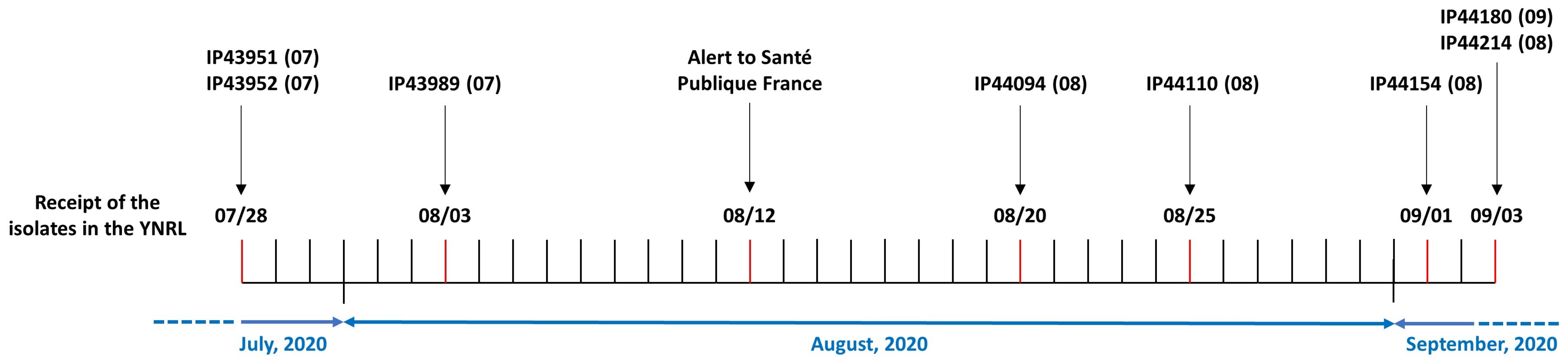
