## Supplementary material for "First description of a *Yersinia pseudotuberculosis* clonal outbreak in France, confirmed using a new core genome multilocus sequence typing method": Table_S1

| BIGSdb locus | gene | synonym | locus tag | old locus tag | alternative locus tag | product | protein id | Comment |
|---|---|---|---|---|---|---|---|---|
| yeps_YPTB0001 | mioC | | YPTB_RS00075 | YPTB0001 | | FMN-binding protein MioC | WP_002212258.1 | |
| yeps_YPTB0004 | viaA | | YPTB_RS00090 | YPTB0004 | | ATPase RavA stimulator ViaA | WP_002212255.1 | |
| yeps_YPTB0006 | kup | trkD | YPTB_RS00100 | YPTB0006 | | low affinity potassium transporter Kup | WP_011191440.1 | |
| yeps_YPTB0007 | rbsD | | YPTB_RS00105 | YPTB0007 | | D-ribose pyranase | WP_002212252.1 | |
| yeps_YPTB0008 | rbsK | | YPTB_RS00110 | YPTB0008 | | ribokinase | WP_011191441.1 | |
| yeps_YPTB0009 | | | YPTB_RS00115 | YPTB0009 | | hypothetical protein | WP_002215906.1 | |
| yeps_YPTB0020 | yihI | | YPTB_RS00195 | YPTB0020 | | Der GTPase-activating protein YihI | WP_002213158.1 | |
| yeps_YPTB0021 | hemN | | YPTB_RS00200 | YPTB0021 | | oxygen-independent coproporphyrinogen III oxidase | WP_002213155.1 | |
| yeps_YPTB0023 | glnL | | YPTB_RS00210 | YPTB0023 | | nitrogen regulation protein NR(II) | WP_002213151.1 | |
| yeps_YPTB0025 | typA | | YPTB_RS00225 | YPTB0025 | | ribosome-dependent GTPase TypA | WP_002217380.1 | |
| yeps_YPTB0026 | yihX | | YPTB_RS00230 | YPTB0026 | | glucose-1-phosphatase | WP_002209011.1 | |
| yeps_YPTB0027 | | | YPTB_RS00235 | YPTB0027 | | virulence factor BrkB family protein | WP_011191445.1 | |
| yeps_YPTB0029 | fabY | | YPTB_RS00245 | YPTB0029 | | fatty acid biosynthesis protein FabY | WP_002209008.1 | |
| yeps_YPTB0030 | | | YPTB_RS00250 | YPTB0030 | | hypothetical protein | WP_002209007.1 | |
| yeps_YPTB0031 | | | YPTB_RS00255 | YPTB0031 | | uracil-xanthine permease family protein | WP_011191446.1 | |
| yeps_YPTB0034 | trmH | spoU | YPTB_RS00270 | YPTB0034 | | tRNA (guanosine(18)-2'-O)-methyltransferase TrmH | WP_002209003.1 | |
| yeps_YPTB0035 | spoT | | YPTB_RS00275 | YPTB0035 | | bifunctional GTP diphosphokinase/guanosine-3',5'-bis pyrophosphate 3'-pyrophosphohydrolase | WP_002209002.1 | |
| yeps_YPTB0036 | rpoZ | | YPTB_RS00280 | YPTB0036 | | DNA-directed RNA polymerase subunit omega | WP_004392061.1 | |
| yeps_YPTB0038 | ligB | | YPTB_RS00290 | YPTB0038 | | NAD-dependent DNA ligase LigB | WP_011191447.1 | |
| yeps_YPTB0041 | rph | | YPTB_RS00305 | YPTB0041 | | ribonuclease PH | WP_002208997.1 | |
| yeps_YPTB0042 | pyrE | | YPTB_RS00310 | YPTB0042 | | orotate phosphoribosyltransferase | WP_002208996.1 | |
| yeps_YPTB0045 | coaBC | | YPTB_RS00325 | YPTB0045 | | bifunctional phosphopantothenoylcysteine decarboxylase/phosphopantothenate--cysteine ligase CoaBC | WP_002208993.1 | |
| yeps_YPTB0046 | radC | | YPTB_RS00330 | YPTB0046 | | DNA repair protein RadC | WP_002208992.1 | |
| yeps_YPTB0050 | coaD | | YPTB_RS00350 | YPTB0050 | | pantetheine-phosphate adenylyltransferase | WP_011191449.1 | |
| yeps_YPTB0051 | | | YPTB_RS00355 | YPTB0051 | | glycosyltransferase family 2 protein | WP_002208987.1 | |
| yeps_YPTB0054 | rfaF | waaF | YPTB_RS00370 | YPTB0054 | | ADP-heptose--LPS heptosyltransferase RfaF | WP_011191450.1 | |
| yeps_YPTB0056 | | | YPTB_RS00380 | YPTB0056 | | glycine C-acetyltransferase | WP_002208982.1 | |
| yeps_YPTB0057 | tdh | | YPTB_RS00385 | YPTB0057 | | L-threonine 3-dehydrogenase | WP_011191451.1 | |
| yeps_YPTB0058 | | | YPTB_RS00390 | YPTB0058 | | divergent polysaccharide deacetylase family protein | WP_011191452.1 | |
| yeps_YPTB0059 | envC | | YPTB_RS00395 | YPTB0059 | | murein hydrolase activator EnvC | WP_002208980.1 | |
| yeps_YPTB0060 | gpmM | | YPTB_RS00400 | YPTB0060 | | 2,3-bisphosphoglycerate-independent phosphoglycerate mutase | WP_011191453.1 | |
| yeps_YPTB0061 | | | YPTB_RS00405 | YPTB0061 | | rhodanese-like domain-containing protein | WP_002208978.1 | |
| yeps_YPTB0062 | grxC | | YPTB_RS00410 | YPTB0062 | | glutaredoxin 3 | WP_002208977.1 | |
| yeps_YPTB0063 | secB | prlG | YPTB_RS00415 | YPTB0063 | | protein-export chaperone SecB | WP_002208976.1 | |
| yeps_YPTB0066 | cysE | | YPTB_RS00430 | YPTB0066 | | serine O-acetyltransferase | WP_002208974.1 | |
| yeps_YPTB0067 | trmL | | YPTB_RS00435 | YPTB0067 | | tRNA (uridine(34)/cytosine(34)/5-carboxymethylaminomethyluridine(34)-2'-O)-methyltransferase TrmL | WP_011191455.1 | |
| yeps_YPTB0068 | ada | | YPTB_RS00440 | YPTB0068 | | bifunctional DNA-binding transcriptional regulator/O6-methylguanine-DNA methyltransferase Ada | WP_011191456.1 | |
| yeps_YPTB0069 | cpxA | | YPTB_RS00445 | YPTB0069 | | envelope stress sensor histidine kinase CpxA | WP_002208971.1 | |
| yeps_YPTB0071 | cpxP | | YPTB_RS00455 | YPTB0071 | | cell-envelope stress modulator CpxP | WP_011191457.1 | |
| yeps_YPTB0073 | fieF | | YPTB_RS00465 | YPTB0073 | | CDF family cation-efflux transporter FieF | WP_002208967.1 | |
| yeps_YPTB0075 | | | YPTB_RS00475 | YPTB0075 | | sulfate ABC transporter substrate-binding protein | WP_011191459.1 | |
| yeps_YPTB0076 | | | YPTB_RS00480 | YPTB0076 | | molybdate ABC transporter substrate-binding protein | WP_011191460.1 | |
| yeps_YPTB0078 | | | YPTB_RS00490 | YPTB0078 | | 4-carboxy-4-hydroxy-2-oxoadipate aldolase/oxaloacetate decarboxylase | WP_002208962.1 | |
| yeps_YPTB0079 | | | YPTB_RS00495 | YPTB0079 | | PIG-L family deacetylase | WP_002208961.1 | |
| yeps_YPTB0081 | tpiA | tpi | YPTB_RS00505 | YPTB0081 | | triose-phosphate isomerase | WP_002208959.1 | |
| yeps_YPTB0083 | | | YPTB_RS00515 | YPTB0083 | | DUF805 domain-containing protein | WP_002208957.1 | |
| yeps_YPTB0084 | fpr | mvrA | YPTB_RS00520 | YPTB0084 | | ferredoxin--NADP(+) reductase | WP_002208956.1 | |
| yeps_YPTB0085 | glpX | | YPTB_RS00525 | YPTB0085 | | class II fructose-bisphosphatase | WP_002223912.1 | |
| yeps_YPTB0087 | | | YPTB_RS00535 | YPTB0087 | | aquaporin family protein | WP_002208954.1 | |
| yeps_YPTB0088 | | | | YPTB0088 | | hypothetical protein | CAH19328.1 | |
| yeps_YPTB0090 | | | YPTB_RS00550 | YPTB0090 | | peptidase domain-containing ABC transporter | WP_011191463.1 | |
| yeps_YPTB0091 | | | YPTB_RS00555 | YPTB0091 | | HlyD family secretion protein | WP_002208949.1 | |
| yeps_YPTB0095 | rraA | | YPTB_RS00575 | YPTB0095 | | ribonuclease E activity regulator RraA | WP_002208945.1 | |
| yeps_YPTB0096 | | | YPTB_RS00580 | YPTB0096 | | 1,4-dihydroxy-2-naphthoate polyprenyltransferase | WP_011191464.1 | |
| yeps_YPTB0098 | hslV | | YPTB_RS00590 | YPTB0098 | | ATP-dependent protease subunit HslV | WP_002208942.1 | |
| yeps_YPTB0100 | cytR | | YPTB_RS00600 | YPTB0100 | | DNA-binding transcriptional regulator CytR | WP_002216730.1 | |
| yeps_YPTB0102 | rpmE | | YPTB_RS00615 | YPTB0102 | | 50S ribosomal protein L31 | WP_002216737.1 | |
| yeps_YPTB0106 | | | YPTB_RS00640 | YPTB0106 | | bifunctional aspartate kinase/homoserine dehydrogenase II | WP_002208934.1 | |
| yeps_YPTB0115 | | | YPTB_RS00695 | YPTB0115 | | type I secretion system permease/ATPase | WP_011191472.1 | |
| yeps_YPTB0119 | | | YPTB_RS00715 | YPTB0119 | | glutathione peroxidase | WP_002209479.1 | |
| yeps_YPTB0120 | oxyR | | YPTB_RS00720 | YPTB0120 | | DNA-binding transcriptional regulator OxyR | WP_011191476.1 | |
| yeps_YPTB0122 | fabR | | YPTB_RS00730 | YPTB0122 | | HTH-type transcriptional repressor FabR | WP_002209476.1 | |
| yeps_YPTB0123 | | | YPTB_RS00735 | YPTB0123 | | YijD family membrane protein | WP_002209475.1 | |
| yeps_YPTB0124 | trmA | | YPTB_RS00740 | YPTB0124 | | tRNA (uridine(54)-C5)-methyltransferase TrmA | WP_002209474.1 | |
| yeps_YPTB0128 | | | YPTB_RS00800 | YPTB0128 | | sugar ABC transporter ATP-binding protein | WP_011191480.1 | |
| yeps_YPTB0129 | | | YPTB_RS00805 | YPTB0129 | | ABC transporter permease | WP_002220992.1 | |
| yeps_YPTB0130 | yjfF | | YPTB_RS00810 | YPTB0130 | | sugar ABC transporter permease YjfF | WP_002212021.1 | |
| yeps_YPTB0131 | hdfR | | YPTB_RS00815 | YPTB0131 | | HTH-type transcriptional regulator HdfR | WP_002212020.1 | |
| yeps_YPTB0135 | ilvM | | YPTB_RS00835 | YPTB0135 | | acetolactate synthase 2 small subunit | WP_002212016.1 | |
| yeps_YPTB0136 | | | YPTB_RS00840 | YPTB0136 | | branched-chain amino acid transaminase | WP_002212015.1 | |
| yeps_YPTB0144 | ilvY | | YPTB_RS00880 | YPTB0144 | | HTH-type transcriptional activator IlvY | WP_002212008.1 | |
| yeps_YPTB0163 | rep | | YPTB_RS00995 | YPTB0163 | | DNA helicase Rep | WP_002211993.1 | |
| yeps_YPTB0165 | rhlB | mmrA | YPTB_RS01005 | YPTB0165 | | ATP-dependent RNA helicase RhlB | WP_002228177.1 | |
| yeps_YPTB0168 | wecA | | YPTB_RS01020 | YPTB0168 | | UDP-N-acetylglucosamine--undecaprenyl-phosphate N-acetylglucosaminephosphotransferase | WP_002211988.1 | |
| yeps_YPTB0169 | wzzE | wzz | YPTB_RS01025 | YPTB0169 | | ECA polysaccharide chain length modulation protein | WP_012304740.1 | |
| yeps_YPTB0172 | rffG | | YPTB_RS01040 | YPTB0172 | | dTDP-glucose 4,6-dehydratase | WP_172601537.1 | |
| yeps_YPTB0174 | rffC | wecD | YPTB_RS01050 | YPTB0174 | | dTDP-4-amino-4,6-dideoxy-D-galactose acyltransferase | WP_012304738.1 | |
| yeps_YPTB0175 | rffA | wecE | YPTB_RS01055 | YPTB0175 | | dTDP-4-amino-4,6-dideoxygalactose transaminase | WP_002211981.1 | |
| yeps_YPTB0176 | wzxE | wzx | YPTB_RS01060 | YPTB0176 | | lipid III flippase WzxE | WP_011191502.1 | |
| yeps_YPTB0178 | wzyE | | YPTB_RS01070 | YPTB0178 | | ECA oligosaccharide polymerase | WP_002211978.1 | |
| yeps_YPTB0183 | hemD | | YPTB_RS01115 | YPTB0183 | | uroporphyrinogen-III synthase | WP_011191507.1 | |
| yeps_YPTB0191 | | | YPTB_RS01160 | YPTB0191 | | DUF484 domain-containing protein | WP_002211472.1 | |
| yeps_YPTB0192 | xerC | | YPTB_RS01165 | YPTB0192 | | tyrosine recombinase XerC | WP_011191512.1 | |
| yeps_YPTB0194 | uvrD | mutU,pdeB,rad,recL | YPTB_RS01175 | YPTB0194 | | DNA helicase II | WP_002211475.1 | |
| yeps_YPTB0198 | corA | | YPTB_RS01195 | YPTB0198 | | magnesium/cobalt transporter CorA | WP_002211480.1 | |
| yeps_YPTB0200 | | | YPTB_RS01205 | YPTB0200 | | thioesterase family protein | WP_002211482.1 | |
| yeps_YPTB0201 | pldA | | YPTB_RS01210 | YPTB0201 | | phospholipase A | WP_002211483.1 | |
| yeps_YPTB0202 | recQ | | YPTB_RS01215 | YPTB0202 | | ATP-dependent DNA helicase RecQ | WP_002211484.1 | |
| yeps_YPTB0203 | rhtC | | YPTB_RS01220 | YPTB0203 | | threonine export protein RhtC | WP_002211485.1 | |
| yeps_YPTB0205 | pldB | | YPTB_RS01230 | YPTB0205 | | lysophospholipase L2 | WP_002211487.1 | |
| yeps_YPTB0208 | glpQ | | YPTB_RS01245 | YPTB0208 | | glycerophosphodiester phosphodiesterase | WP_002211490.1 | |
| yeps_YPTB0209 | glpA | | YPTB_RS01255 | YPTB0209 | | anaerobic glycerol-3-phosphate dehydrogenase subunit A | WP_002211491.1 | |
| yeps_YPTB0210 | glpB | | YPTB_RS01260 | YPTB0210 | | glycerol-3-phosphate dehydrogenase subunit GlpB | WP_011191516.1 | |
| yeps_YPTB0212 | | | YPTB_RS01270 | YPTB0212 | | DcrB family lipoprotein | WP_002211494.1 | |
| yeps_YPTB0213 | | | YPTB_RS01275 | YPTB0213 | | 7-cyano-7-deazaguanine/7-aminomethyl-7-deazaguanine transporter | WP_011191518.1 | |
| yeps_YPTB0214 | tusA | | YPTB_RS01280 | YPTB0214 | | sulfurtransferase TusA | WP_002215973.1 | |
| yeps_YPTB0217 | | | YPTB_RS01295 | YPTB0217 | | DUF1820 family protein | WP_002211499.1 | |
| yeps_YPTB0218 | | | YPTB_RS01300 | YPTB0218 | | DUF2500 domain-containing protein | WP_002211500.1 | |
| yeps_YPTB0220 | rsmD | | YPTB_RS01310 | YPTB0220 | | 16S rRNA (guanine(966)-N(2))-methyltransferase | WP_011191520.1 | |
| yeps_YPTB0223 | ftsX | ftsS | YPTB_RS01330 | YPTB0223 | | permease-like cell division protein FtsX | WP_002211505.1 | |
| yeps_YPTB0225 | panM | | YPTB_RS01340 | YPTB0225 | | aspartate 1-decarboxylase autocleavage activator PanM | WP_011191522.1 | |
| yeps_YPTB0226 | | | YPTB_RS01345 | YPTB0226 | | branched-chain amino acid ABC transporter substrate-binding protein | WP_011191523.1 | |
| yeps_YPTB0230 | livF | | YPTB_RS01365 | YPTB0230 | | high-affinity branched-chain amino acid ABC transporter ATP-binding protein LivF | WP_002211512.1 | |
| yeps_YPTB0235 | | | YPTB_RS01390 | YPTB0235 | | fimbria/pilus periplasmic chaperone | WP_011191529.1 | |
| yeps_YPTB0242 | ugpQ | | YPTB_RS01425 | YPTB0242 | | glycerophosphodiester phosphodiesterase | WP_011191534.1 | |
| yeps_YPTB0243 | | | YPTB_RS01430 | YPTB0243 | | AEC family transporter | WP_002224020.1 | |
| yeps_YPTB0246 | | | YPTB_RS01445 | YPTB0246 | | carboxylate/amino acid/amine transporter | WP_002215597.1 | |
| yeps_YPTB0248 | metE | | YPTB_RS01455 | YPTB0248 | | 5-methyltetrahydropteroyltriglutamate--homocysteine S-methyltransferase | WP_011191538.1 | |
| yeps_YPTB0249 | | | YPTB_RS01460 | YPTB0249 | | dienelactone hydrolase family protein | WP_012413425.1 | |
| yeps_YPTB0254 | rmuC | | YPTB_RS01485 | YPTB0254 | | DNA recombination protein RmuC | WP_002211532.1 | |
| yeps_YPTB0255 | ubiE | | YPTB_RS01490 | YPTB0255 | | bifunctional demethylmenaquinone methyltransferase/2-methoxy-6-polyprenyl-1,4-benzoquinol methylase UbiE | WP_002224024.1 | |
| yeps_YPTB0256 | | | YPTB_RS01495 | YPTB0256 | | SCP2 domain-containing protein | WP_002215928.1 | |
| yeps_YPTB0257 | ubiB | | YPTB_RS01500 | YPTB0257 | | ubiquinone biosynthesis regulatory protein kinase UbiB | WP_002211535.1 | |
| yeps_YPTB0258 | | | YPTB_RS01505 | YPTB0258 | | Sec-independent protein translocase subunit TatA | WP_002211536.1 | |
| yeps_YPTB0259 | tatB | mtta2 | YPTB_RS01510 | YPTB0259 | | Sec-independent protein translocase protein TatB | WP_011191542.1 | |
| yeps_YPTB0260 | tatC | mttB | YPTB_RS01515 | YPTB0260 | | Sec-independent protein translocase subunit TatC | WP_011191543.1 | |
| yeps_YPTB0268 | pepQ | | YPTB_RS01555 | YPTB0268 | | Xaa-Pro dipeptidase | WP_011191548.1 | |
| yeps_YPTB0269 | | | YPTB_RS01560 | YPTB0269 | | IMPACT family protein | WP_002231137.1 | |
| yeps_YPTB0273 | birA | bioR,dhbB | YPTB_RS01610 | YPTB0273 | | bifunctional biotin--[acetyl-CoA-carboxylase] ligase/biotin operon repressor BirA | WP_011191550.1 | |
| yeps_YPTB0274 | coaA | panK,rts | YPTB_RS01615 | YPTB0274 | | type I pantothenate kinase | WP_002212290.1 | |
| yeps_YPTB0276 | tuf | | YPTB_RS01645 | YPTB0276 | | elongation factor Tu | WP_002210669.1 | |
| yeps_YPTB0281 | rplJ | | YPTB_RS01670 | YPTB0281 | | 50S ribosomal protein L10 | WP_002210674.1 | |
| yeps_YPTB0283 | rpoB | | YPTB_RS01680 | YPTB0283 | | DNA-directed RNA polymerase subunit beta | WP_002210676.1 | |
| yeps_YPTB0286 | | | YPTB_RS01700 | YPTB0286 | | thiazole synthase | WP_002228257.1 | |
| yeps_YPTB0287 | thiS | | YPTB_RS01705 | YPTB0287 | | sulfur carrier protein ThiS | WP_002217275.1 | |
| yeps_YPTB0288 | | | YPTB_RS01710 | YPTB0288 | | HesA/MoeB/ThiF family protein | WP_002210680.1 | |
| yeps_YPTB0289 | thiE | | YPTB_RS01715 | YPTB0289 | | thiamine phosphate synthase | WP_012303440.1 | |
| yeps_YPTB0291 | rsd | | YPTB_RS01730 | YPTB0291 | | sigma D regulator | WP_011191553.1 | |
| yeps_YPTB0292 | nudC | | YPTB_RS01735 | YPTB0292 | | NAD(+) diphosphatase | WP_011191554.1 | |

| | | | | | | | |
|---|---|---|---|---|---|---|---|
| yeps_YPTB0294 | hemE | | YPTB_RS01745 | YPTB0294 | | uroporphyrinogen decarboxylase | WP_011191555.1 |
| yeps_YPTB0297 | hupA | | YPTB_RS01760 | YPTB0297 | | DNA-binding protein HU-alpha | WP_002210689.1 |
| yeps_YPTB0299 | purD | | YPTB_RS01770 | YPTB0299 | | phosphoribosylamine--glycine ligase | WP_011191558.1 |
| yeps_YPTB0310 | | | YPTB_RS01850 | YPTB0310 | | two component system response regulator | WP_011191565.1 |
| yeps_YPTB0311 | | | YPTB_RS01855 | YPTB0311 | | two component system sensor kinase | WP_011191566.1 |
| yeps_YPTB0315 | | | YPTB_RS01875 | YPTB0315 | | AraC family transcriptional regulator | WP_011171760.1 |
| yeps_YPTB0316 | sctF | | YPTB_RS01880 | YPTB0316 | | type III secretion system needle filament subunit SctF | WP_011191571.1 |
| yeps_YPTB0317 | | | YPTB_RS01885 | YPTB0317 | | EscG/YscG/SsaH family type III secretion system needle protein co-chaperone | WP_002209040.1 |
| yeps_YPTB0318 | sctI | | YPTB_RS01890 | YPTB0318 | | type III secretion system inner rod subunit SctI | WP_002209041.1 |
| yeps_YPTB0319 | ssaJ | | YPTB_RS01895 | YPTB0319 | | EscJ/YscJ/HrcJ family type III secretion inner membrane ring protein SsaJ | WP_011191572.1 |
| yeps_YPTB0321 | sctL | | YPTB_RS01905 | YPTB0321 | | type III secretion system stator protein SctL | WP_011191573.1 |
| yeps_YPTB0325 | | | | YPTB0325 | | conserved hypothetical protein | CAH19565.1 |
| yeps_YPTB0326 | | | YPTB_RS01930 | YPTB0326 | | YscQ/HrcQ family type III secretion apparatus protein | WP_012413433.1 |
| yeps_YPTB0328 | | | YPTB_RS01940 | YPTB0328 | | EscS/YscS/HrcS family type III secretion system export apparatus protein | WP_002209051.1 |
| yeps_YPTB0335 | metC | | YPTB_RS01975 | YPTB0335 | | cystathionine beta-lyase | WP_002215044.1 |
| yeps_YPTB0336 | | | YPTB_RS01980 | YPTB0336 | | heme ABC transporter ATP-binding protein | WP_002209058.1 |
| yeps_YPTB0338 | | | YPTB_RS01990 | YPTB0338 | | hemin ABC transporter substrate-binding protein | WP_011191579.1 |
| yeps_YPTB0339 | | | YPTB_RS01995 | YPTB0339 | | hemin-degrading factor | WP_002209061.1 |
| yeps_YPTB0340 | | | YPTB_RS02000 | YPTB0340 | | TonB-dependent hemoglobin/transferrin/lactoferrin family receptor | WP_011191580.1 |
| yeps_YPTB0341 | | | YPTB_RS23515 | YPTB0341 | | hemin uptake protein HemP | WP_011191581.1 |
| yeps_YPTB0342 | | | YPTB_RS02010 | YPTB0342 | | SDR family oxidoreductase | WP_011191582.1 |
| yeps_YPTB0347 | | | YPTB_RS02035 | YPTB0347 | | citrate lyase subunit beta | WP_002209069.1 |
| yeps_YPTB0348 | | | YPTB_RS02040 | YPTB0348 | | cysteine protease StiP family protein | WP_002209070.1 |
| yeps_YPTB0349 | | | YPTB_RS02045 | YPTB0349 | | hypothetical protein | WP_002209071.1 |
| yeps_YPTB0350 | | | YPTB_RS02050 | YPTB0350 | | phosphoribosyltransferase domain-containing protein | WP_002209072.1 |
| yeps_YPTB0352 | | | YPTB_RS02060 | YPTB0352 | | tellurium resistance TerZ family protein | WP_011191586.1 |
| yeps_YPTB0354 | | | YPTB_RS02070 | YPTB0354 | | tellurite resistance TerB family protein | WP_002209076.1 |
| yeps_YPTB0355 | | | YPTB_RS02075 | YPTB0355 | | TerC/Alx family metal homeostasis membrane protein | WP_011191588.1 |
| yeps_YPTB0356 | | | YPTB_RS02080 | YPTB0356 | | TerD family protein | WP_002209078.1 |
| yeps_YPTB0367 | ubiA | cyr | YPTB_RS02135 | YPTB0367 | | 4-hydroxybenzoate octaprenyltransferase | WP_002209088.1 |
| yeps_YPTB0369 | | | YPTB_RS02145 | YPTB0369 | | diacylglycerol kinase | WP_002209089.1 |
| yeps_YPTB0371 | zur | | YPTB_RS02155 | YPTB0371 | | zinc uptake transcriptional repressor Zur | WP_002214647.1 |
| yeps_YPTB0378 | | | YPTB_RS02190 | YPTB0378 | | MmcQ/YjbR family DNA-binding protein | WP_002209098.1 |
| yeps_YPTB0379 | uvrA | dinE | YPTB_RS02195 | YPTB0379 | | excinuclease ABC subunit UvrA | WP_011191598.1 |
| yeps_YPTB0381 | rhaM | | YPTB_RS02205 | YPTB0381 | | L-rhamnose mutarotase | WP_002209101.1 |
| yeps_YPTB0382 | fucO | | YPTB_RS02210 | YPTB0382 | | lactaldehyde reductase | WP_011191599.1 |
| yeps_YPTB0383 | rhaD | | YPTB_RS02215 | YPTB0383 | | rhamnulose-1-phosphate aldolase | WP_011191600.1 |
| yeps_YPTB0384 | | | YPTB_RS02220 | YPTB0384 | | L-rhamnose isomerase | WP_011191601.1 |
| yeps_YPTB0385 | rhaB | | YPTB_RS02225 | YPTB0385 | | rhamnulokinase | WP_002209105.1 |
| yeps_YPTB0386 | rhaS | rhaC2 | YPTB_RS02240 | YPTB0386 | | HTH-type transcriptional activator RhaS | WP_011191602.1 |
| yeps_YPTB0387 | rhaR | rhaC1 | YPTB_RS02245 | YPTB0387 | | HTH-type transcriptional activator RhaR | WP_011191603.1 |
| yeps_YPTB0391 | | | YPTB_RS02260 | YPTB0391 | | subtilase family AB5 toxin binding subunit | WP_002209112.1 |
| yeps_YPTB0396 | aegA | | YPTB_RS02285 | YPTB0396 | | formate-dependent uric acid utilization protein AegA | WP_011191607.1 |
| yeps_YPTB0397 | | | YPTB_RS02290 | YPTB0397 | | 4Fe-4S binding protein | WP_002209119.1 |
| yeps_YPTB0400 | cutA | | YPTB_RS02305 | YPTB0400 | | divalent cation tolerance protein CutA | WP_002209122.1 |
| yeps_YPTB0401 | | | YPTB_RS02310 | YPTB0401 | | anaerobic C4-dicarboxylate transporter | WP_011191610.1 |
| yeps_YPTB0409 | sugE | | YPTB_RS02350 | YPTB0409 | | quaternary ammonium compound efflux SMR transporter SugE | WP_002228124.1 |
| yeps_YPTB0410 | frdD | | YPTB_RS02355 | YPTB0410 | | fumarate reductase subunit FrdD | WP_002209134.1 |
| yeps_YPTB0414 | epmA | | YPTB_RS02380 | YPTB0414 | | elongation factor P--(R)-beta-lysine ligase | WP_002209139.1 |
| yeps_YPTB0415 | mscM | | YPTB_RS02385 | YPTB0415 | | miniconductance mechanosensitive channel MscM | WP_032466065.1 |
| yeps_YPTB0416 | psd | | YPTB_RS02390 | YPTB0416 | | archaetidylserine decarboxylase | WP_002209141.1 |
| yeps_YPTB0417 | rsgA | | YPTB_RS02395 | YPTB0417 | | small ribosomal subunit biogenesis GTPase RsgA | WP_002209142.1 |
| yeps_YPTB0420 | nnr | | YPTB_RS02435 | YPTB0420 | | bifunctional ADP-dependent NAD(P)H-hydrate dehydratase/NAD(P)H-hydrate epimerase | WP_011191613.1 |
| yeps_YPTB0421 | tsaE | | YPTB_RS02440 | YPTB0421 | | tRNA (adenosine(37)-N6)-threonylcarbamoyltransferase complex ATPase subunit type 1 TsaE | WP_011191614.1 |
| yeps_YPTB0426 | hflX | | YPTB_RS02465 | YPTB0426 | | GTPase HflX | WP_002209152.1 |
| yeps_YPTB0428 | hflC | hflA | YPTB_RS02475 | YPTB0428 | | protease modulator HflC | WP_002209155.1 |
| yeps_YPTB0430 | | | YPTB_RS02485 | YPTB0430 | | adenylosuccinate synthase | WP_002209157.1 |
| yeps_YPTB0431 | nsrR | | YPTB_RS02490 | YPTB0431 | | nitric oxide-sensing transcriptional repressor NsrR | WP_002217229.1 |
| yeps_YPTB0435 | bsmA | | YPTB_RS02510 | YPTB0435 | | biofilm peroxide resistance protein BsmA | WP_002215294.1 |
| yeps_YPTB0436 | yjfP | | YPTB_RS02515 | YPTB0436 | | esterase | WP_011191618.1 |
| yeps_YPTB0437 | | | | YPTB0437 | | hypothetical protein | CAH19677.1 |
| yeps_YPTB0442 | | | YPTB_RS02545 | YPTB0442 | | DUF3757 domain-containing protein | WP_002210157.1 |
| yeps_YPTB0443 | | | YPTB_RS02550 | YPTB0443 | | OapA family protein | WP_002210158.1 |
| yeps_YPTB0445 | ytfE | | YPTB_RS02560 | YPTB0445 | | iron-sulfur cluster repair protein YtfE | WP_011191621.1 |
| yeps_YPTB0446 | | | YPTB_RS02565 | YPTB0446 | | bifunctional 2',3'-cyclic-nucleotide 2'-phosphodiesterase/3'-nucleotidase | WP_011191622.1 |
| yeps_YPTB0447 | cysQ | amtA | YPTB_RS02570 | YPTB0447 | | 3'(2'),5'-bisphosphate nucleotidase CysQ | WP_032466061.1 |
| yeps_YPTB0448 | | | YPTB_RS02575 | YPTB0448 | | YtfJ family protein | WP_002210162.1 |
| yeps_YPTB0449 | | | YPTB_RS02580 | YPTB0449 | | DUF1107 domain-containing protein | WP_002210163.1 |
| yeps_YPTB0451 | msrA | pms | YPTB_RS02590 | YPTB0451 | | peptide-methionine (S)-S-oxide reductase MsrA | WP_002210165.1 |
| yeps_YPTB0452 | | | YPTB_RS02595 | YPTB0452 | | autotransporter assembly complex protein TamA | WP_011191624.1 |
| yeps_YPTB0454 | | | YPTB_RS02605 | YPTB0454 | | gamma-glutamylcyclotransferase | WP_002210168.1 |
| yeps_YPTB0455 | ppa | | YPTB_RS02610 | YPTB0455 | | inorganic diphosphatase | WP_002210169.1 |
| yeps_YPTB0456 | fbp | fdp | YPTB_RS02615 | YPTB0456 | | class 1 fructose-bisphosphatase | WP_011191626.1 |
| yeps_YPTB0467 | cgtA | | YPTB_RS02675 | YPTB0467 | | Obg family GTPase CgtA | WP_002210181.1 |
| yeps_YPTB0468 | pmrB | | YPTB_RS02680 | YPTB0468 | | two-component system sensor histidine kinase PmrB | WP_011191629.1 |
| yeps_YPTB0469 | pmrA | | YPTB_RS02685 | YPTB0469 | | two-component system response regulator PmrA | WP_002210183.1 |
| yeps_YPTB0474 | ftsH | hflB,mrsC,tolZ | YPTB_RS02710 | YPTB0474 | | ATP-dependent zinc metalloprotease FtsH | WP_002228195.1 |
| yeps_YPTB0477 | secG | | YPTB_RS02725 | YPTB0477 | | preprotein translocase subunit SecG | WP_002210190.1 |
| yeps_YPTB0479 | nusA | | YPTB_RS02745 | YPTB0479 | | transcription termination factor NusA | WP_002209253.1 |
| yeps_YPTB0482 | truB | p35 | YPTB_RS02760 | YPTB0482 | | tRNA pseudouridine(55) synthase TruB | WP_002209256.1 |
| yeps_YPTB0491 | | | YPTB_RS02805 | YPTB0491 | | efflux RND transporter periplasmic adaptor subunit | WP_002209266.1 |
| yeps_YPTB0493 | | | YPTB_RS02815 | YPTB0493 | | efflux transporter outer membrane subunit | WP_011191634.1 |
| yeps_YPTB0495 | | | YPTB_RS02825 | YPTB0495 | | U32 family peptidase | WP_011191635.1 |
| yeps_YPTB0496 | | | YPTB_RS02835 | YPTB0496 | | SCP2 domain-containing protein | WP_002209272.1 |
| yeps_YPTB0497 | | | YPTB_RS02840 | YPTB0497 | | N-acetyltransferase | WP_011191636.1 |
| yeps_YPTB0498 | | | YPTB_RS02850 | YPTB0498 | | GIY-YIG nuclease family protein | WP_011191637.1 |
| yeps_YPTB0499 | | | YPTB_RS02855 | YPTB0499 | | heparinase II/III family protein | WP_011191638.1 |
| yeps_YPTB0502 | | | YPTB_RS02870 | YPTB0502 | | sugar ABC transporter permease | WP_011191641.1 |
| yeps_YPTB0503 | | | YPTB_RS02875 | YPTB0503 | | carbohydrate ABC transporter permease | WP_002209279.1 |
| yeps_YPTB0504 | ugpC | | YPTB_RS02880 | YPTB0504 | | sn-glycerol-3-phosphate ABC transporter ATP-binding protein UgpC | WP_002209280.1 |
| yeps_YPTB0505 | | | YPTB_RS02885 | YPTB0505 | | alginate lyase family protein | WP_002209281.1 |
| yeps_YPTB0507 | | | YPTB_RS02895 | YPTB0507 | | hypothetical protein | WP_002209283.1 |
| yeps_YPTB0508 | phnP | | YPTB_RS02900 | YPTB0508 | | phosphonate metabolism protein PhnP | WP_002209284.1 |
| yeps_YPTB0509 | phnN | | YPTB_RS02905 | YPTB0509 | | ribose 1,5-bisphosphokinase | WP_002209285.1 |
| yeps_YPTB0510 | phnM | | YPTB_RS02910 | YPTB0510 | | alpha-D-ribose 1-methylphosphonate 5-triphosphate diphosphatase | WP_011191643.1 |
| yeps_YPTB0511 | phnL | | YPTB_RS02915 | YPTB0511 | | phosphonate C-P lyase system protein PhnL | WP_011191644.1 |
| yeps_YPTB0512 | phnK | | YPTB_RS02920 | YPTB0512 | | phosphonate C-P lyase system protein PhnK | WP_011191645.1 |
| yeps_YPTB0513 | | | YPTB_RS02925 | YPTB0513 | | alpha-D-ribose 1-methylphosphonate 5-phosphate C-P-lyase PhnJ | WP_011191646.1 |
| yeps_YPTB0516 | phnG | | YPTB_RS24180 | YPTB0516 | | phosphonate C-P lyase system protein PhnG | WP_002209292.1 |
| yeps_YPTB0517 | phnF | | YPTB_RS24185 | YPTB0517 | | phosphonate metabolism transcriptional regulator PhnF | WP_011191649.1 |
| yeps_YPTB0518 | nrdG | yjgE | YPTB_RS02950 | YPTB0518 | | anaerobic ribonucleoside-triphosphate reductase-activating protein | WP_025470743.1 |
| yeps_YPTB0519 | nrdD | | YPTB_RS02955 | YPTB0519 | | anaerobic ribonucleoside-triphosphate reductase | WP_002209296.1 |
| yeps_YPTB0520 | | | YPTB_RS02960 | YPTB0520 | | ABC transporter ATP-binding protein | WP_002209297.1 |
| yeps_YPTB0521 | | | YPTB_RS02965 | YPTB0521 | | ABC transporter ATP-binding protein | WP_002209298.1 |
| yeps_YPTB0522 | | | YPTB_RS02970 | YPTB0522 | | ABC transporter permease | WP_002209299.1 |
| yeps_YPTB0526 | argF | | YPTB_RS02990 | YPTB0526 | | ornithine carbamoyltransferase | WP_011191651.1 |
| yeps_YPTB0529 | | | YPTB_RS03005 | YPTB0529 | | valine--tRNA ligase | WP_011191653.1 |
| yeps_YPTB0533 | lptG | | YPTB_RS03025 | YPTB0533 | | LPS export ABC transporter permease LptG | WP_002223173.1 |
| yeps_YPTB0541 | | | YPTB_RS03070 | YPTB0541 | | AraC family transcriptional regulator | WP_011191658.1 |
| yeps_YPTB0542 | | | YPTB_RS03075 | YPTB0542 | | PTS fructose-like transporter subunit IIB | WP_002209180.1 |
| yeps_YPTB0546 | | | YPTB_RS03095 | YPTB0546 | | DUF2501 domain-containing protein | WP_002209184.1 |
| yeps_YPTB0547 | lsrG | | YPTB_RS03100 | YPTB0547 | | (4S)-4-hydroxy-5-phosphonooxypentane-2,3-dione isomerase | WP_002209186.1 |
| yeps_YPTB0549 | lsrB | | YPTB_RS03110 | YPTB0549 | | autoinducer 2 ABC transporter substrate-binding protein LsrB | WP_011191662.1 |
| yeps_YPTB0550 | lsrD | | YPTB_RS03115 | YPTB0550 | | autoinducer 2 ABC transporter permease LsrD | WP_011191663.1 |
| yeps_YPTB0551 | lsrC | | YPTB_RS03120 | YPTB0551 | | autoinducer 2 ABC transporter permease LsrC | WP_011191664.1 |
| yeps_YPTB0552 | lsrA | | YPTB_RS03125 | YPTB0552 | | autoinducer 2 ABC transporter ATP-binding protein LsrA | WP_011191665.1 |
| yeps_YPTB0554 | lsrK | | YPTB_RS03140 | YPTB0554 | | autoinducer-2 kinase | WP_002230543.1 |
| yeps_YPTB0564 | | | YPTB_RS03185 | YPTB0564 | | LapA family protein | WP_002209198.1 |
| yeps_YPTB0565 | | | YPTB_RS03190 | YPTB0565 | | efflux RND transporter periplasmic adaptor subunit | WP_002209199.1 |
| yeps_YPTB0575 | prfC | miaD,tos | YPTB_RS03260 | YPTB0575 | | peptide chain release factor 3 | WP_011191681.1 |
| yeps_YPTB0581 | deoC | dra | YPTB_RS03290 | YPTB0581 | | deoxyribose-phosphate aldolase | WP_002216078.1 |
| yeps_YPTB0582 | deoA | tpp,ttg | YPTB_RS03295 | YPTB0582 | | thymidine phosphorylase | WP_011191687.1 |
| yeps_YPTB0585 | | | YPTB_RS03310 | YPTB0585 | | YtjB family periplasmic protein | WP_002209218.1 |
| yeps_YPTB0586 | serB | | YPTB_RS03315 | YPTB0586 | | phosphoserine phosphatase | WP_002209219.1 |
| yeps_YPTB0587 | radA | sms | YPTB_RS03320 | YPTB0587 | | DNA repair protein RadA | WP_002209220.1 |
| yeps_YPTB0588 | nadR | nadI,pnuA | YPTB_RS03325 | YPTB0588 | | multifunctional transcriptional regulator/nicotinamide-nucleotide adenylyltransferase/ribosylnicotinamide kinase NadR | WP_002209221.1 |
| yeps_YPTB0590 | ettA | | YPTB_RS03335 | YPTB0590 | | energy-dependent translational throttle protein EttA | WP_011191691.1 |
| yeps_YPTB0591 | | | YPTB_RS03340 | YPTB0591 | | OmpA family protein | WP_011191692.1 |
| yeps_YPTB0593 | | | YPTB_RS03350 | YPTB0593 | | YfiR family protein | WP_002209225.1 |
| yeps_YPTB0595 | sltY | | YPTB_RS03360 | YPTB0595 | | murein transglycosylase | WP_011191694.1 |

| | | | | | | | |
|---|---|---|---|---|---|---|---|
| yeps_YPTB0597 | yjjX | | YPTB_RS03370 | YPTB0597 | | inosine/xanthosine triphosphatase | WP_011191695.1 |
| yeps_YPTB0600 | creA | yjjD | YPTB_RS03385 | YPTB0600 | | protein CreA | WP_002209232.1 |
| yeps_YPTB0611 | dnaK | | YPTB_RS03440 | YPTB0611 | | molecular chaperone DnaK | WP_002209248.1 |
| yeps_YPTB0612 | dnaJ | groP,grpC | YPTB_RS03445 | YPTB0612 | | molecular chaperone DnaJ | WP_002209249.1 |
| yeps_YPTB0613 | nhaA | ant,antA | YPTB_RS03450 | YPTB0613 | | Na+/H+ antiporter NhaA | WP_011191700.1 |
| yeps_YPTB0615 | rpsT | sup(s20) | YPTB_RS03460 | YPTB0615 | | 30S ribosomal protein S20 | WP_002220715.1 |
| yeps_YPTB0617 | ileS | ilvS | YPTB_RS03475 | YPTB0617 | | isoleucine--tRNA ligase | WP_002210509.1 |
| yeps_YPTB0618 | lspA | | YPTB_RS03480 | YPTB0618 | | signal peptidase II | WP_002210508.1 |
| yeps_YPTB0619 | fkpB | slpA,yaaD | YPTB_RS03485 | YPTB0619 | | FKBP-type peptidyl-prolyl cis-trans isomerase | WP_002228112.1 |
| yeps_YPTB0623 | carA | arg,cap,pyrA | YPTB_RS03505 | YPTB0623 | | glutamine-hydrolyzing carbamoyl-phosphate synthase small subunit | WP_002224759.1 |
| yeps_YPTB0624 | carB | arg,cap,pyrA | YPTB_RS03510 | YPTB0624 | | carbamoyl-phosphate synthase large subunit | WP_011191702.1 |
| yeps_YPTB0625 | | | YPTB_RS03515 | YPTB0625 | | LysE family translocator | WP_011191703.1 |
| yeps_YPTB0626 | | | YPTB_RS03520 | YPTB0626 | | threonine/serine exporter ThrE family protein | WP_011191704.1 |
| yeps_YPTB0627 | | | YPTB_RS03525 | YPTB0627 | | threonine/serine exporter | WP_032466075.1 |
| yeps_YPTB0631 | apaH | cfcB | YPTB_RS03545 | YPTB0631 | | bis(5'-nucleosyl)-tetraphosphatase (symmetrical) ApaH | WP_011191708.1 |
| yeps_YPTB0634 | pdxA | | YPTB_RS03560 | YPTB0634 | | 4-hydroxythreonine-4-phosphate dehydrogenase PdxA | WP_011191711.1 |
| yeps_YPTB0635 | surA | | YPTB_RS03565 | YPTB0635 | | peptidylprolyl isomerase SurA | WP_002210488.1 |
| yeps_YPTB0637 | djlA | yabH | YPTB_RS03575 | YPTB0637 | | co-chaperone DjlA | WP_011191713.1 |
| yeps_YPTB0641 | tssB | | YPTB_RS03595 | YPTB0641 | | type VI secretion system contractile sheath small subunit | WP_002210481.1 |
| yeps_YPTB0643 | | | YPTB_RS03605 | YPTB0643 | | Hcp family type VI secretion system effector | WP_002210479.1 |
| yeps_YPTB0644 | | | YPTB_RS03610 | YPTB0644 | | type VI secretion system baseplate subunit TssE | WP_002210478.1 |
| yeps_YPTB0645 | tssF | | YPTB_RS03615 | YPTB0645 | | type VI secretion system baseplate subunit TssF | WP_011191714.1 |
| yeps_YPTB0647 | tssH | | YPTB_RS03625 | YPTB0647 | | type VI secretion system ATPase TssH | WP_002210475.1 |
| yeps_YPTB0649 | | | YPTB_RS03635 | YPTB0649 | | pentapeptide repeat-containing protein | WP_011191716.1 |
| yeps_YPTB0651 | | | YPTB_RS03645 | YPTB0651 | | DUF3540 domain-containing protein | WP_002210473.1 |
| yeps_YPTB0652 | | | YPTB_RS03650 | YPTB0652 | | DUF4150 domain-containing protein | WP_002214737.1 |
| yeps_YPTB0653 | tssJ | | YPTB_RS03655 | YPTB0653 | | type VI secretion system lipoprotein TssJ | WP_011191717.1 |
| yeps_YPTB0654 | tssK | | YPTB_RS03660 | YPTB0654 | | type VI secretion system baseplate subunit TssK | WP_002210470.1 |
| yeps_YPTB0660 | | | YPTB_RS03690 | YPTB0660 | | DedA family protein | WP_002210465.1 |
| yeps_YPTB0661 | thiQ | sfuC,yabJ | YPTB_RS03695 | YPTB0661 | | thiamine ABC transporter ATP-binding protein ThiQ | WP_011191721.1 |
| yeps_YPTB0662 | thiP | sfuB,yabK | YPTB_RS03700 | YPTB0662 | | thiamine/thiamine pyrophosphate ABC transporter permease ThiP | WP_011191722.1 |
| yeps_YPTB0668 | | | YPTB_RS03735 | YPTB0668 | | sugar efflux transporter | WP_011191726.1 |
| yeps_YPTB0669 | leuD | | YPTB_RS03740 | YPTB0669 | | 3-isopropylmalate dehydratase small subunit | WP_002210456.1 |
| yeps_YPTB0670 | leuC | | YPTB_RS03745 | YPTB0670 | | 3-isopropylmalate dehydratase large subunit | WP_011191727.1 |
| yeps_YPTB0680 | rsmH | | YPTB_RS03815 | YPTB0680 | | 16S rRNA (cytosine(1402)-N(4))-methyltransferase RsmH | WP_002210442.1 |
| yeps_YPTB0681 | ftsL | mraR,yabD | YPTB_RS03820 | YPTB0681 | | cell division protein FtsL | WP_002210441.1 |
| yeps_YPTB0682 | | | YPTB_RS03825 | YPTB0682 | | peptidoglycan glycosyltransferase FtsI | WP_011191731.1 |
| yeps_YPTB0683 | murE | | YPTB_RS03830 | YPTB0683 | | UDP-N-acetylmuramoyl-L-alanyl-D-glutamate--2,6-diaminopimelate ligase | WP_012105673.1 |
| yeps_YPTB0684 | murF | mra | YPTB_RS03835 | YPTB0684 | | UDP-N-acetylmuramoyl-tripeptide--D-alanyl-D-alanine ligase | WP_011191733.1 |
| yeps_YPTB0686 | murD | | YPTB_RS03845 | YPTB0686 | | UDP-N-acetylmuramoyl-L-alanine--D-glutamate ligase | WP_011191734.1 |
| yeps_YPTB0687 | ftsW | | YPTB_RS03850 | YPTB0687 | | cell division protein FtsW | WP_002210435.1 |
| yeps_YPTB0688 | murG | | YPTB_RS03855 | YPTB0688 | | undecaprenyldiphospho-muramoylpentapeptide beta-N-acetylglucosaminyltransferase | WP_011191735.1 |
| yeps_YPTB0689 | murC | | YPTB_RS03860 | YPTB0689 | | UDP-N-acetylmuramate--L-alanine ligase | WP_002216457.1 |
| yeps_YPTB0694 | lpxC | asmB,envA | YPTB_RS03885 | YPTB0694 | | UDP-3-O-acyl-N-acetylglucosamine deacetylase | WP_002228285.1 |
| yeps_YPTB0695 | | | YPTB_RS03890 | YPTB0695 | | DUF721 domain-containing protein | WP_002210428.1 |
| yeps_YPTB0696 | secM | | YPTB_RS03895 | YPTB0696 | | secA translation cis-regulator SecM | WP_011191738.1 |
| yeps_YPTB0706 | gspE | pulE | YPTB_RS03955 | YPTB0706 | | type II secretion system protein GspE | WP_011191740.1 |
| yeps_YPTB0708 | nadC | | YPTB_RS03965 | YPTB0708 | | carboxylating nicotinate-nucleotide diphosphorylase | WP_002209324.1 |
| yeps_YPTB0710 | ampE | | YPTB_RS03975 | YPTB0710 | | beta-lactamase regulator AmpE | WP_002209326.1 |
| yeps_YPTB0717 | | | YPTB_RS04010 | YPTB0717 | | protein YacL | WP_002209334.1 |
| yeps_YPTB0718 | yddG | | YPTB_RS04015 | YPTB0718 | | aromatic amino acid DMT transporter YddG | WP_011191744.1 |
| yeps_YPTB0720 | speE | | YPTB_RS04025 | YPTB0720 | | polyamine aminopropyltransferase | WP_011191746.1 |
| yeps_YPTB0721 | | | YPTB_RS04030 | YPTB0721 | | YacC family pilotin-like protein | WP_002228215.1 |
| yeps_YPTB0722 | cueO | | YPTB_RS04035 | YPTB0722 | | multicopper oxidase CueO | WP_011191747.1 |
| yeps_YPTB0724 | can | | YPTB_RS04045 | YPTB0724 | | carbonate dehydratase | WP_002209342.1 |
| yeps_YPTB0725 | | | YPTB_RS04050 | YPTB0725 | | ABC transporter ATP-binding protein | WP_011191749.1 |
| yeps_YPTB0727 | | | YPTB_RS04060 | YPTB0727 | | polysaccharide deacetylase family protein | WP_012413481.1 |
| yeps_YPTB0728 | panD | | YPTB_RS04065 | YPTB0728 | | aspartate 1-decarboxylase | WP_011191751.1 |
| yeps_YPTB0729 | panC | | YPTB_RS04070 | YPTB0729 | | pantoate--beta-alanine ligase | WP_011191752.1 |
| yeps_YPTB0730 | panB | | YPTB_RS04075 | YPTB0730 | | 3-methyl-2-oxobutanoate hydroxymethyltransferase | WP_012304571.1 |
| yeps_YPTB0738 | mrcB | pbpF,ponB | YPTB_RS04115 | YPTB0738 | | bifunctional glycosyl transferase/transpeptidase | WP_011191759.1 |
| yeps_YPTB0739 | fhuC | | YPTB_RS04120 | YPTB0739 | | Fe3+-hydroxamate ABC transporter ATP-binding protein FhuC | WP_002209359.1 |
| yeps_YPTB0745 | | | YPTB_RS04150 | YPTB0745 | | TRIC cation channel family protein | WP_002209366.1 |
| yeps_YPTB0746 | btuF | | YPTB_RS04155 | YPTB0746 | | vitamin B12 ABC transporter substrate-binding protein BtuF | WP_011191763.1 |
| yeps_YPTB0748 | dgt | optA | YPTB_RS04165 | YPTB0748 | | dGTPase | WP_011191765.1 |
| yeps_YPTB0749 | degP | | YPTB_RS04170 | YPTB0749 | | serine endoprotease DegP | WP_011191766.1 |
| yeps_YPTB0753 | mazG | | YPTB_RS04190 | YPTB0753 | | nucleoside triphosphate pyrophosphohydrolase | WP_011191769.1 |
| yeps_YPTB0754 | pyrG | | YPTB_RS04195 | YPTB0754 | | CTP synthase (glutamine hydrolyzing) | WP_002209376.1 |
| yeps_YPTB0755 | eno | | YPTB_RS04200 | YPTB0755 | | phosphopyruvate hydratase | WP_011191770.1 |
| yeps_YPTB0757 | queE | | YPTB_RS04210 | YPTB0757 | | 7-carboxy-7-deazaguanine synthase QueE | WP_002209379.1 |
| yeps_YPTB0758 | queD | | YPTB_RS04215 | YPTB0758 | | 6-carboxytetrahydropterin synthase QueD | WP_002209380.1 |
| yeps_YPTB0760 | cysI | | YPTB_RS04225 | YPTB0760 | | assimilatory sulfite reductase (NADPH) hemoprotein subunit | WP_011191773.1 |
| yeps_YPTB0763 | | | YPTB_RS04240 | YPTB0763 | | type II toxin-antitoxin system HicA family toxin | WP_002209384.1 |
| yeps_YPTB0766 | cysN | | YPTB_RS04260 | YPTB0766 | | sulfate adenylyltransferase subunit CysN | WP_002209387.1 |
| yeps_YPTB0767 | cysC | | YPTB_RS04265 | YPTB0767 | | adenylyl-sulfate kinase | WP_002209388.1 |
| yeps_YPTB0769 | ftsB | | YPTB_RS04275 | YPTB0769 | | cell division protein FtsB | WP_002209390.1 |
| yeps_YPTB0771 | ispF | mecS,ygbB | YPTB_RS04285 | YPTB0771 | | 2-C-methyl-D-erythritol 2,4-cyclodiphosphate synthase | WP_002209392.1 |
| yeps_YPTB0772 | truD | | YPTB_RS04290 | YPTB0772 | | tRNA pseudouridine(13) synthase TruD | WP_011191777.1 |
| yeps_YPTB0773 | surE | ygbC | YPTB_RS04295 | YPTB0773 | | 5'/3'-nucleotidase SurE | WP_011191778.1 |
| yeps_YPTB0775 | nlpD | | YPTB_RS04305 | YPTB0775 | | murein hydrolase activator NlpD | WP_012304557.1 |
| yeps_YPTB0779 | rpiB | yjcA | YPTB_RS04330 | YPTB0779 | | ribose 5-phosphate isomerase B | WP_002209402.1 |
| yeps_YPTB0780 | | | YPTB_RS04335 | YPTB0780 | | erythritol/L-threitol dehydrogenase | WP_002209403.1 |
| yeps_YPTB0781 | | | YPTB_RS04340 | YPTB0781 | | D-threitol dehydrogenase | WP_002209404.1 |
| yeps_YPTB0782 | | | YPTB_RS04345 | YPTB0782 | | dihydroxyacetone kinase subunit DhaK | WP_002228229.1 |
| yeps_YPTB0783 | dhaL | | YPTB_RS04350 | YPTB0783 | | dihydroxyacetone kinase subunit L | WP_011191782.1 |
| yeps_YPTB0792 | | | YPTB_RS04395 | YPTB0792 | | TonB-dependent receptor | WP_002223222.1 |
| yeps_YPTB0794 | map | pepM(S.t.) | YPTB_RS04410 | YPTB0794 | | type I methionyl aminopeptidase | WP_002209416.1 |
| yeps_YPTB0795 | | | YPTB_RS04415 | YPTB0795 | | ParD-like family protein | WP_002209417.1 |
| yeps_YPTB0798 | | | YPTB_RS04430 | YPTB0798 | | L-ribulose-5-phosphate 3-epimerase | WP_011191789.1 |
| yeps_YPTB0800 | | | YPTB_RS04440 | YPTB0800 | | ABC transporter permease | WP_002209421.1 |
| yeps_YPTB0802 | | | YPTB_RS04450 | YPTB0802 | | substrate-binding domain-containing protein | WP_041175489.1 |
| yeps_YPTB0803 | | | YPTB_RS04455 | YPTB0803 | | DeoR/GlpR family DNA-binding transcription regulator | WP_011191793.1 |
| yeps_YPTB0806 | dmsB | | YPTB_RS04470 | YPTB0806 | | dimethylsulfoxide reductase subunit B | WP_011191795.1 |
| yeps_YPTB0807 | | | YPTB_RS04475 | YPTB0807 | | dimethyl sulfoxide reductase anchor subunit | WP_011191796.1 |
| yeps_YPTB0811 | katG | | YPTB_RS04495 | YPTB0811 | | catalase/peroxidase HPI | WP_002209433.1 |
| yeps_YPTB0812 | | | YPTB_RS04500 | YPTB0812 | | D-ribose ABC transporter substrate-binding protein | WP_011191800.1 |
| yeps_YPTB0813 | | | YPTB_RS04505 | YPTB0813 | | DUF2291 family protein | WP_002209435.1 |
| yeps_YPTB0814 | | | YPTB_RS04510 | YPTB0814 | | sugar ABC transporter ATP-binding protein | WP_011191801.1 |
| yeps_YPTB0816 | | | YPTB_RS04520 | YPTB0816 | | transketolase | WP_002209438.1 |
| yeps_YPTB0819 | | | YPTB_RS04535 | YPTB0819 | | L-fucose/L-arabinose isomerase family protein | WP_011191804.1 |
| yeps_YPTB0820 | | | YPTB_RS04540 | YPTB0820 | | sugar-binding transcriptional regulator | WP_002209443.1 |
| yeps_YPTB0821 | | | YPTB_RS04545 | YPTB0821 | | MFS transporter | WP_011191805.1 |
| yeps_YPTB0822 | pncC | | YPTB_RS04550 | YPTB0822 | | nicotinamide-nucleotide amidase | WP_002209445.1 |
| yeps_YPTB0825 | alaS | act,ala-act,lovB | YPTB_RS04565 | YPTB0825 | | alanine--tRNA ligase | WP_011191808.1 |
| yeps_YPTB0828 | | | YPTB_RS04600 | YPTB0828 | | DedA family protein | WP_002209451.1 |
| yeps_YPTB0829 | gshA | gsh-I | YPTB_RS04605 | YPTB0829 | | glutamate--cysteine ligase | WP_172957065.1 |
| yeps_YPTB0832 | | | YPTB_RS04620 | YPTB0832 | | inner membrane protein YpjD | WP_002209455.1 |
| yeps_YPTB0836 | trmD | | YPTB_RS04640 | YPTB0836 | | tRNA (guanosine(37)-N1)-methyltransferase TrmD | WP_002222284.1 |
| yeps_YPTB0850 | pssA | pss | YPTB_RS04745 | YPTB0850 | | CDP-diacylglycerol--serine O-phosphatidyltransferase | WP_002208751.1 |
| yeps_YPTB0853 | trxC | yfiG | YPTB_RS04760 | YPTB0853 | | thioredoxin TrxC | WP_011191817.1 |
| yeps_YPTB0856 | emrA | | YPTB_RS04775 | YPTB0856 | | multidrug efflux MFS transporter periplasmic adaptor subunit EmrA | WP_002208745.1 |
| yeps_YPTB0860 | | | YPTB_RS04795 | YPTB0860 | | MFS transporter | WP_011191821.1 |
| yeps_YPTB0862 | | | YPTB_RS04805 | YPTB0862 | | AtzE family amidohydrolase | WP_011191823.1 |
| yeps_YPTB0863 | hpxX | | YPTB_RS04810 | YPTB0863 | | oxalurate catabolism protein HpxX | WP_002208738.1 |
| yeps_YPTB0866 | | | YPTB_RS04825 | YPTB0866 | | transporter substrate-binding domain-containing protein | WP_011191825.1 |
| yeps_YPTB0867 | | | YPTB_RS04830 | YPTB0867 | | amino acid ABC transporter permease | WP_002208734.1 |
| yeps_YPTB0868 | | | YPTB_RS04835 | YPTB0868 | | amino acid ABC transporter permease | WP_011191826.1 |
| yeps_YPTB0869 | | | YPTB_RS04840 | YPTB0869 | | amino acid ABC transporter ATP-binding protein | WP_002208732.1 |
| yeps_YPTB0881 | | | YPTB_RS04900 | YPTB0881 | | hypothetical protein | WP_011191838.1 |
| yeps_YPTB0883 | fadE | | YPTB_RS04910 | YPTB0883 | | acyl-CoA dehydrogenase FadE | WP_011191839.1 |
| yeps_YPTB0884 | lpcA | gmhA,tfrA,yafI | YPTB_RS04915 | YPTB0884 | | D-sedoheptulose 7-phosphate isomerase | WP_002208720.1 |
| yeps_YPTB0885 | | | YPTB_RS04920 | YPTB0885 | | class II glutamine amidotransferase | WP_002208719.1 |
| yeps_YPTB0886 | | | YPTB_RS04925 | YPTB0886 | | murein L,D-transpeptidase | WP_011191840.1 |
| yeps_YPTB0888 | | | YPTB_RS04935 | YPTB0888 | | NADH:ubiquinone reductase (Na(+)-transporting) subunit B | WP_002208716.1 |
| yeps_YPTB0891 | nqrE | nqr5 | YPTB_RS04950 | YPTB0891 | | NADH:ubiquinone reductase (Na(+)-transporting) subunit E | WP_002208713.1 |
| yeps_YPTB0893 | | | YPTB_RS04960 | YPTB0893 | | FAD:protein FMN transferase | WP_002227821.1 |
| yeps_YPTB0912 | ppnP | | YPTB_RS05065 | YPTB0912 | | pyrimidine/purine nucleoside phosphorylase | WP_002208692.1 |
| yeps_YPTB0914 | mak | | YPTB_RS05075 | YPTB0914 | | fructokinase | WP_011191852.1 |
| yeps_YPTB0916 | sbcD | yajA | YPTB_RS05085 | YPTB0916 | | exonuclease subunit SbcD | WP_011191854.1 |
| yeps_YPTB0917 | phoB | phoRc | YPTB_RS05095 | YPTB0917 | | phosphate response regulator transcription factor PhoB | WP_002208685.1 |

| | | | | | | | |
|---|---|---|---|---|---|---|---|
| yeps_YPTB0918 | phoR | | YPTB_RS05100 | YPTB0918 | | phosphate regulon sensor histidine kinase PhoR | WP_002208684.1 |
| yeps_YPTB0923 | | | YPTB_RS05125 | YPTB0923 | | SDR family oxidoreductase | WP_002208677.1 |
| yeps_YPTB0927 | queA | | YPTB_RS05150 | YPTB0927 | | tRNA preQ1(34) S-adenosylmethionine ribosyltransferase-isomerase QueA | WP_011191859.1 |
| yeps_YPTB0929 | yajC | | YPTB_RS05160 | YPTB0929 | | preprotein translocase subunit YajC | WP_002208671.1 |
| yeps_YPTB0935 | ribE | ribC | YPTB_RS05195 | YPTB0935 | | 6,7-dimethyl-8-ribityllumazine synthase | WP_002208666.1 |
| yeps_YPTB0937 | thiL | | YPTB_RS05205 | YPTB0937 | | thiamine-phosphate kinase | WP_011191861.1 |
| yeps_YPTB0938 | pgpA | yajN | YPTB_RS05210 | YPTB0938 | | phosphatidylglycerophosphatase A | WP_002208663.1 |
| yeps_YPTB0939 | dxs | | YPTB_RS05215 | YPTB0939 | | 1-deoxy-D-xylulose-5-phosphate synthase | WP_011191862.1 |
| yeps_YPTB0940 | ispA | | YPTB_RS05220 | YPTB0940 | | (2E,6E)-farnesyl diphosphate synthase | WP_011191863.1 |
| yeps_YPTB0941 | xseB | yajE | YPTB_RS05225 | YPTB0941 | | exodeoxyribonuclease VII small subunit | WP_002208660.1 |
| yeps_YPTB0942 | | | | YPTB0942 | | hypothetical protein | CAH20182.1 |
| yeps_YPTB0943 | thiI | nuvA,yajJ,yajK | YPTB_RS05235 | YPTB0943 | | tRNA 4-thiouridine(8) synthase ThiI | WP_011191864.1 |
| yeps_YPTB0944 | yajL | | YPTB_RS05240 | YPTB0944 | | protein deglycase YajL | WP_002208657.1 |
| yeps_YPTB0945 | panE | abpA | YPTB_RS05245 | YPTB0945 | | 2-dehydropantoate 2-reductase | WP_002208656.1 |
| yeps_YPTB0946 | | | YPTB_RS05250 | YPTB0946 | | YajQ family cyclic di-GMP-binding protein | WP_002208655.1 |
| yeps_YPTB0949 | | | YPTB_RS05265 | YPTB0949 | | cytochrome o ubiquinol oxidase subunit IV | WP_002208652.1 |
| yeps_YPTB0954 | ampG | | YPTB_RS05290 | YPTB0954 | | muropeptide MFS transporter AmpG | WP_002223277.1 |
| yeps_YPTB0956 | bolA | | YPTB_RS05300 | YPTB0956 | | transcriptional regulator BolA | WP_002208644.1 |
| yeps_YPTB0957 | | | | YPTB0957 | | hypothetical protein | CAH20197.1 |
| yeps_YPTB0965 | | | YPTB_RS05345 | YPTB0965 | | acyl-CoA thioesterase | WP_002208636.1 |
| yeps_YPTB0967 | | | YPTB_RS05355 | YPTB0967 | | SgrR family transcriptional regulator | WP_011191867.1 |
| yeps_YPTB0968 | cof | | YPTB_RS05360 | YPTB0968 | | HMP-PP phosphatase | WP_011191868.1 |
| yeps_YPTB0973 | glnK | ybaI | YPTB_RS05385 | YPTB0973 | | P-II family nitrogen regulator | WP_002208627.1 |
| yeps_YPTB0974 | amtB | ybaG | YPTB_RS05390 | YPTB0974 | | ammonium transporter AmtB | WP_002228344.1 |
| yeps_YPTB0975 | tesB | | YPTB_RS05395 | YPTB0975 | | acyl-CoA thioesterase II | WP_002208625.1 |
| yeps_YPTB0976 | | | YPTB_RS05400 | YPTB0976 | | YbaY family lipoprotein | WP_011191872.1 |
| yeps_YPTB0986 | | | YPTB_RS05450 | YPTB0986 | | DsrE/DsrF/TusD sulfur relay family protein | WP_002208613.1 |
| yeps_YPTB0992 | dnaX | dnaZ,dnaZX | YPTB_RS05480 | YPTB0992 | | DNA polymerase III subunit gamma/tau | WP_002208605.1 |
| yeps_YPTB1024 | | | YPTB_RS05650 | YPTB1024 | | NfeD family protein | WP_002208577.1 |
| yeps_YPTB1026 | | | YPTB_RS05660 | YPTB1026 | | co-chaperone YbbN | WP_011191894.1 |
| yeps_YPTB1028 | tesA | apeA,pldC | YPTB_RS05670 | YPTB1028 | | multifunctional acyl-CoA thioesterase I/protease I/lysophospholipase L1 | WP_002208573.1 |
| yeps_YPTB1030 | | | YPTB_RS05680 | YPTB1030 | | ABC transporter permease | WP_011191896.1 |
| yeps_YPTB1031 | purK | | YPTB_RS05685 | YPTB1031 | | 5-(carboxyamino)imidazole ribonucleotide synthase | WP_002208570.1 |
| yeps_YPTB1032 | purE | | YPTB_RS05690 | YPTB1032 | | 5-(carboxyamino)imidazole ribonucleotide mutase | WP_002208569.1 |
| yeps_YPTB1035 | cysS | | YPTB_RS05705 | YPTB1035 | | cysteine--tRNA ligase | WP_011191897.1 |
| yeps_YPTB1036 | ybcJ | | YPTB_RS05710 | YPTB1036 | | ribosome-associated protein YbcJ | WP_002209775.1 |
| yeps_YPTB1037 | folD | ads | YPTB_RS05715 | YPTB1037 | | bifunctional methylenetetrahydrofolate dehydrogenase/methenyltetrahydrofolate cyclohydrolase FolD | WP_002209774.1 |
| yeps_YPTB1038 | | | YPTB_RS05725 | YPTB1038 | | AlpA family phage regulatory protein | WP_002215270.1 |
| yeps_YPTB1040 | | | YPTB_RS05735 | YPTB1040 | | hypothetical protein | WP_011191899.1 |
| yeps_YPTB1041 | | | YPTB_RS05740 | YPTB1041 | | hypothetical protein | WP_002215266.1 |
| yeps_YPTB1043 | | | YPTB_RS05745 | YPTB1043 | | hypothetical protein | WP_041175443.1 |
| yeps_YPTB1044 | | | YPTB_RS05755 | YPTB1044 | | CDP-alcohol phosphatidyltransferase family protein | WP_002209770.1 |
| yeps_YPTB1045 | | | YPTB_RS05760 | YPTB1045 | | phosphatidate cytidylyltransferase | WP_011191901.1 |
| yeps_YPTB1047 | | | YPTB_RS05770 | YPTB1047 | | bifunctional alpha/beta hydrolase/class I SAM-dependent methyltransferase | WP_011191902.1 |
| yeps_YPTB1048 | | | YPTB_RS05775 | YPTB1048 | | phosphatase PAP2/dual specificity phosphatase family protein | WP_011191903.1 |
| yeps_YPTB1050 | | | YPTB_RS05785 | YPTB1050 | | hypothetical protein | WP_002215253.1 |
| yeps_YPTB1051 | | | YPTB_RS05790 | YPTB1051 | | LysR family transcriptional regulator | WP_011191905.1 |
| yeps_YPTB1053 | | | YPTB_RS05800 | YPTB1053 | | aldo/keto reductase | WP_011191906.1 |
| yeps_YPTB1054 | | | YPTB_RS05815 | YPTB1054 | | hypothetical protein | WP_011191907.1 |
| yeps_YPTB1056 | | | YPTB_RS05825 | YPTB1056 | | GDYXXLXY domain-containing protein | WP_011191909.1 |
| yeps_YPTB1057 | | | YPTB_RS05830 | YPTB1057 | | DUF4401 domain-containing protein | WP_011191910.1 |
| yeps_YPTB1068 | pbpC | | YPTB_RS05890 | YPTB1068 | | penicillin-binding protein 1C | WP_002223314.1 |
| yeps_YPTB1071 | | | YPTB_RS05905 | YPTB1071 | | CoA-acylating methylmalonate-semialdehyde dehydrogenase | WP_011191918.1 |
| yeps_YPTB1073 | iolG | | YPTB_RS05915 | YPTB1073 | | inositol 2-dehydrogenase | WP_011191920.1 |
| yeps_YPTB1074 | | | YPTB_RS05920 | YPTB1074 | | substrate-binding domain-containing protein | WP_002210300.1 |
| yeps_YPTB1075 | | | YPTB_RS05925 | YPTB1075 | | sugar ABC transporter ATP-binding protein | WP_011191921.1 |
| yeps_YPTB1076 | | | YPTB_RS05930 | YPTB1076 | | ABC transporter permease | WP_002210302.1 |
| yeps_YPTB1082 | | | YPTB_RS05960 | YPTB1082 | | ABC transporter ATP-binding protein/permease | WP_002214723.1 |
| yeps_YPTB1083 | | | YPTB_RS05965 | YPTB1083 | | antibiotic biosynthesis monooxygenase | WP_002210309.1 |
| yeps_YPTB1086 | | | YPTB_RS05985 | YPTB1086 | | LuxR family transcriptional regulator | WP_002210313.1 |
| yeps_YPTB1087 | | | YPTB_RS05990 | YPTB1087 | | cold shock domain-containing protein | WP_012304395.1 |
| yeps_YPTB1089 | crcB | | YPTB_RS06000 | YPTB1089 | | fluoride efflux transporter CrcB | WP_011191930.1 |
| yeps_YPTB1090 | tatA | mttaI | YPTB_RS06010 | YPTB1090 | | Sec-independent protein translocase subunit TatA | WP_002210319.1 |
| yeps_YPTB1092 | lipB | | YPTB_RS06020 | YPTB1092 | | lipoyl(octanoyl) transferase LipB | WP_011228201.1 |
| yeps_YPTB1093 | | | YPTB_RS06025 | YPTB1093 | | DUF493 family protein YbeD | WP_002210322.1 |
| yeps_YPTB1094 | dacA | | YPTB_RS06030 | YPTB1094 | | D-alanyl-D-alanine carboxypeptidase DacA | WP_011191931.1 |
| yeps_YPTB1095 | rlpA | | YPTB_RS06035 | YPTB1095 | | endolytic peptidoglycan transglycosylase RlpA | WP_002210324.1 |
| yeps_YPTB1099 | rsfS | | YPTB_RS06055 | YPTB1099 | | ribosome silencing factor | WP_002210329.1 |
| yeps_YPTB1100 | nadD | | YPTB_RS06060 | YPTB1100 | | nicotinate-nucleotide adenylyltransferase | WP_002210330.1 |
| yeps_YPTB1101 | holA | | YPTB_RS06065 | YPTB1101 | | DNA polymerase III subunit delta | WP_011191933.1 |
| yeps_YPTB1102 | lptE | | YPTB_RS06070 | YPTB1102 | | LPS assembly lipoprotein LptE | WP_002210332.1 |
| yeps_YPTB1107 | | | YPTB_RS06095 | YPTB1107 | | amino acid ABC transporter permease | WP_002223337.1 |
| yeps_YPTB1109 | lnt | | YPTB_RS06105 | YPTB1109 | | apolipoprotein N-acyltransferase | WP_002210341.1 |
| yeps_YPTB1111 | ybeY | | YPTB_RS06115 | YPTB1111 | | rRNA maturation RNase YbeY | WP_011191934.1 |
| yeps_YPTB1117 | | | YPTB_RS06185 | YPTB1117 | | ROK family transcriptional regulator | WP_002224863.1 |
| yeps_YPTB1118 | nagA | | YPTB_RS06190 | YPTB1118 | | N-acetylglucosamine-6-phosphate deacetylase | WP_002210351.1 |
| yeps_YPTB1123 | | | YPTB_RS06215 | YPTB1123 | | beta-N-acetylhexosaminidase | WP_011191940.1 |
| yeps_YPTB1124 | fur | | YPTB_RS06220 | YPTB1124 | | ferric iron uptake transcriptional regulator | WP_011191941.1 |
| yeps_YPTB1126 | | | | YPTB1126 | | hypothetical | CAH20366.1 |
| yeps_YPTB1136 | | | YPTB_RS06285 | YPTB1136 | | type II toxin-antitoxin system RatA family toxin | WP_002210715.1 |
| yeps_YPTB1137 | | | YPTB_RS06290 | YPTB1137 | | RnfH family protein | WP_002210716.1 |
| yeps_YPTB1138 | bamE | | YPTB_RS06295 | YPTB1138 | | outer membrane protein assembly factor BamE | WP_002210717.1 |
| yeps_YPTB1140 | nadK | | YPTB_RS06305 | YPTB1140 | | NAD(+) kinase | WP_011191944.1 |
| yeps_YPTB1142 | | | YPTB_RS06315 | YPTB1142 | | citrate synthase | WP_002210721.1 |
| yeps_YPTB1143 | sdhC | cybA | YPTB_RS06320 | YPTB1143 | | succinate dehydrogenase cytochrome b556 subunit | WP_002210722.1 |
| yeps_YPTB1152 | cydB | | YPTB_RS06365 | YPTB1152 | | cytochrome d ubiquinol oxidase subunit II | WP_002210731.1 |
| yeps_YPTB1153 | cydX | | YPTB_RS06370 | YPTB1153 | | cytochrome bd-I oxidase subunit CydX | WP_002210732.1 |
| yeps_YPTB1154 | ybgE | | YPTB_RS06375 | YPTB1154 | | cyd operon protein YbgE | WP_011191945.1 |
| yeps_YPTB1156 | tolQ | | YPTB_RS06385 | YPTB1156 | | Tol-Pal system protein TolQ | WP_002210735.1 |
| yeps_YPTB1160 | pal | excC | YPTB_RS06405 | YPTB1160 | | peptidoglycan-associated lipoprotein Pal | WP_011191946.1 |
| yeps_YPTB1167 | | | YPTB_RS06475 | YPTB1167 | | PsiF family protein | WP_011191948.1 |
| yeps_YPTB1168 | galM | | YPTB_RS06480 | YPTB1168 | | galactose-1-epimerase | WP_011191949.1 |
| yeps_YPTB1170 | galT | galB | YPTB_RS06490 | YPTB1170 | | galactose-1-phosphate uridylyltransferase | WP_002210749.1 |
| yeps_YPTB1171 | galE | galD | YPTB_RS06495 | YPTB1171 | | UDP-glucose 4-epimerase GalE | WP_011191951.1 |
| yeps_YPTB1173 | modF | phrA | YPTB_RS06510 | YPTB1173 | | molybdate ABC transporter ATP-binding protein ModF | WP_011191952.1 |
| yeps_YPTB1174 | modE | modR | YPTB_RS06515 | YPTB1174 | | molybdenum-dependent transcriptional regulator | WP_002210754.1 |
| yeps_YPTB1175 | | | YPTB_RS06520 | YPTB1175 | | AcrZ family multidrug efflux pump-associated protein | WP_002217291.1 |
| yeps_YPTB1176 | modA | | YPTB_RS06525 | YPTB1176 | | molybdate ABC transporter substrate-binding protein | WP_002210756.1 |
| yeps_YPTB1178 | modC | chlD,narD | YPTB_RS06535 | YPTB1178 | | molybdenum ABC transporter ATP-binding protein ModC | WP_011191953.1 |
| yeps_YPTB1179 | | | YPTB_RS06540 | YPTB1179 | | pyridoxal phosphatase | WP_011191954.1 |
| yeps_YPTB1180 | pgl | | YPTB_RS06545 | YPTB1180 | | 6-phosphogluconolactonase | WP_011191955.1 |
| yeps_YPTB1182 | bioB | | YPTB_RS06555 | YPTB1182 | | biotin synthase BioB | WP_002210762.1 |
| yeps_YPTB1183 | bioF | | YPTB_RS06560 | YPTB1183 | | 8-amino-7-oxononanoate synthase | WP_011191957.1 |
| yeps_YPTB1184 | bioC | | YPTB_RS06565 | YPTB1184 | | malonyl-ACP O-methyltransferase BioC | WP_011191958.1 |
| yeps_YPTB1185 | bioD | | YPTB_RS06570 | YPTB1185 | | dethiobiotin synthase | WP_002216554.1 |
| yeps_YPTB1186 | | | YPTB_RS06575 | YPTB1186 | | ABC transporter ATP-binding protein | WP_002210766.1 |
| yeps_YPTB1187 | uvrB | | YPTB_RS06580 | YPTB1187 | | excinuclease ABC subunit B | WP_011191959.1 |
| yeps_YPTB1189 | yvcK | | YPTB_RS06590 | YPTB1189 | | uridine diphosphate-N-acetylglucosamine-binding protein YvcK | WP_002210769.1 |
| yeps_YPTB1193 | moaE | chlA5 | YPTB_RS06615 | YPTB1193 | | molybdopterin synthase catalytic subunit MoaE | WP_002210774.1 |
| yeps_YPTB1197 | betI | | YPTB_RS06635 | YPTB1197 | | transcriptional regulator BetI | WP_002218278.1 |
| yeps_YPTB1199 | | | YPTB_RS06645 | YPTB1199 | | LysR family transcriptional regulator | WP_002214199.1 |
| yeps_YPTB1201 | xapA | pndA | YPTB_RS06655 | YPTB1201 | | xanthosine phosphorylase | WP_011191966.1 |
| yeps_YPTB1202 | | | YPTB_RS06660 | YPTB1202 | | nucleoside permease | WP_011191967.1 |
| yeps_YPTB1203 | | | YPTB_RS06665 | YPTB1203 | | zinc resistance sensor/chaperone ZraP | WP_011191968.1 |
| yeps_YPTB1204 | | | YPTB_RS06670 | YPTB1204 | | PAS domain S-box protein | WP_011191969.1 |
| yeps_YPTB1205 | | | YPTB_RS06675 | YPTB1205 | | sigma 54-interacting transcriptional regulator | WP_011191970.1 |
| yeps_YPTB1207 | | | YPTB_RS06685 | YPTB1207 | | LysR family transcriptional regulator | WP_002220168.1 |
| yeps_YPTB1208 | | | YPTB_RS06695 | YPTB1208 | | EthD family reductase | WP_002220163.1 |
| yeps_YPTB1211 | | | YPTB_RS06710 | YPTB1211 | | ATP-binding cassette domain-containing protein | WP_011191972.1 |
| yeps_YPTB1220 | yajD | | YPTB_RS06760 | YPTB1220 | | HNH nuclease YajD | WP_002214185.1 |
| yeps_YPTB1221 | | | YPTB_RS06765 | YPTB1221 | | suppressor of fused domain protein | WP_011191976.1 |
| yeps_YPTB1222 | | | YPTB_RS06770 | YPTB1222 | | GlsB/YeaQ/YmgE family stress response membrane protein | WP_002210786.1 |
| yeps_YPTB1229 | urtB | | YPTB_RS06805 | YPTB1229 | | urea ABC transporter permease subunit UrtB | WP_002210793.1 |
| yeps_YPTB1231 | urtD | | YPTB_RS06815 | YPTB1231 | | urea ABC transporter ATP-binding protein UrtD | WP_011191979.1 |
| yeps_YPTB1250 | | | YPTB_RS06905 | YPTB1250 | | nicotinamide mononucleotide deamidase-related protein YfaY | WP_011191993.1 |
| yeps_YPTB1251 | eco | eti | YPTB_RS06910 | YPTB1251 | | serine protease inhibitor ecotin | WP_002210815.1 |
| yeps_YPTB1252 | | | YPTB_RS06915 | YPTB1252 | | 2Fe-2S ferredoxin-like protein | WP_002210816.1 |
| yeps_YPTB1255 | | | YPTB_RS06930 | YPTB1255 | | bifunctional 3-demethylubiquinone 3-O-methyltransferase/2-octaprenyl-6-hydroxy phenol methylase | WP_002210820.1 |
| yeps_YPTB1257 | rcsC | hmwC | YPTB_RS06945 | YPTB1257 | | two-component system sensor histidine kinase RcsC | WP_011171779.1 |
| yeps_YPTB1258 | rcsB | | YPTB_RS06950 | YPTB1258 | | transcriptional regulator RcsB | WP_002210824.1 |

| | | | | | | | |
|---|---|---|---|---|---|---|---|
| yeps_YPTB1259 | rcsD | | YPTB_RS06955 | YPTB1259 | | phosphotransferase RcsD | WP_011191996.1 |
| yeps_YPTB1263 | | | YPTB_RS06995 | YPTB1263 | | MFS transporter | WP_011191998.1 |
| yeps_YPTB1268 | | | YPTB_RS07020 | YPTB1268 | | LexA family transcriptional regulator | WP_002208864.1 |
| yeps_YPTB1270 | | | YPTB_RS07030 | YPTB1270 | | four-carbon acid sugar kinase family protein | WP_012304340.1 |
| yeps_YPTB1272 | | | YPTB_RS07040 | YPTB1272 | | DeoR/GlpR family DNA-binding transcription regulator | WP_002208861.1 |
| yeps_YPTB1273 | | | YPTB_RS07045 | YPTB1273 | | hydroxypyruvate isomerase family protein | WP_002208860.1 |
| yeps_YPTB1274 | | | YPTB_RS07050 | YPTB1274 | | hypothetical protein | WP_011192001.1 |
| yeps_YPTB1276 | | | YPTB_RS07060 | YPTB1276 | | DUF2635 domain-containing protein | WP_002208857.1 |
| yeps_YPTB1279 | | | YPTB_RS07075 | YPTB1279 | | phage tail assembly protein | WP_002208854.1 |
| yeps_YPTB1283 | | | YPTB_RS07100 | YPTB1283 | | phage baseplate assembly protein | WP_002215460.1 |
| yeps_YPTB1285 | | | YPTB_RS07110 | YPTB1285 | | baseplate J/gp47 family protein | WP_011192005.1 |
| yeps_YPTB1286 | | | YPTB_RS22575 | YPTB1286 | | YmfQ family protein | WP_011192006.1 |
| yeps_YPTB1289 | | | YPTB_RS07135 | YPTB1289 | | MurR/RpiR family transcriptional regulator | WP_002208843.1 |
| yeps_YPTB1290 | | | YPTB_RS07140 | YPTB1290 | | 6-phospho-beta-glucosidase | WP_002208842.1 |
| yeps_YPTB1300 | | | YPTB_RS07205 | YPTB1300 | | DEAD/DEAH box helicase | WP_011192012.1 |
| yeps_YPTB1301 | rsuA | | YPTB_RS07210 | YPTB1301 | | 16S rRNA pseudouridine(516) synthase RsuA | WP_011192013.1 |
| yeps_YPTB1304 | yejF | | YPTB_RS07225 | YPTB1304 | | microcin C ABC transporter ATP-binding protein YejF | WP_011192015.1 |
| yeps_YPTB1305 | | | YPTB_RS07230 | YPTB1305 | | ABC transporter permease subunit | WP_002353954.1 |
| yeps_YPTB1306 | | | YPTB_RS07235 | YPTB1306 | | microcin C ABC transporter permease YejB | WP_002208826.1 |
| yeps_YPTB1307 | | | YPTB_RS07240 | YPTB1307 | | extracellular solute-binding protein | WP_011192016.1 |
| yeps_YPTB1310 | | | YPTB_RS07255 | YPTB1310 | | phosphatase PAP2 family protein | WP_011192018.1 |
| yeps_YPTB1311 | | | YPTB_RS07260 | YPTB1311 | | GTP-binding protein | WP_011192019.1 |
| yeps_YPTB1312 | | | | YPTB1312 | | putative membrane protein | CAH20552.1 |
| yeps_YPTB1313 | | | YPTB_RS07270 | YPTB1313 | | GntR family transcriptional regulator | WP_002208816.1 |
| yeps_YPTB1314 | | | YPTB_RS07275 | YPTB1314 | | mannitol dehydrogenase family protein | WP_002224659.1 |
| yeps_YPTB1315 | uxuA | | YPTB_RS07280 | YPTB1315 | | mannonate dehydratase | WP_011192020.1 |
| yeps_YPTB1317 | mtr | | YPTB_RS07290 | YPTB1317 | | tryptophan permease | WP_002208812.1 |
| yeps_YPTB1318 | | | YPTB_RS07295 | YPTB1318 | | YkgJ family cysteine cluster protein | WP_011192021.1 |
| yeps_YPTB1329 | fruB | fpr,fruF | YPTB_RS07350 | YPTB1329 | | fused PTS fructose transporter subunit IIA/HPr protein | WP_002208799.1 |
| yeps_YPTB1342 | | | YPTB_RS07415 | YPTB1342 | | iron ABC transporter permease | WP_002215451.1 |
| yeps_YPTB1343 | | | YPTB_RS07420 | YPTB1343 | | ABC transporter ATP-binding protein | WP_011192030.1 |
| yeps_YPTB1347 | | | YPTB_RS07445 | YPTB1347 | | isopenicillin N synthase family oxygenase | WP_011192033.1 |
| yeps_YPTB1348 | | | YPTB_RS07450 | YPTB1348 | | MetQ/NlpA family ABC transporter substrate-binding protein | WP_011192034.1 |
| yeps_YPTB1355 | ybjG | | YPTB_RS07490 | YPTB1355 | | undecaprenyl-diphosphate phosphatase | WP_002208768.1 |
| yeps_YPTB1356 | | | YPTB_RS07495 | YPTB1356 | | phosphatase PAP2 family protein | WP_024063536.1 |
| yeps_YPTB1359 | | | YPTB_RS07510 | YPTB1359 | | YbjC family protein | WP_002216036.1 |
| yeps_YPTB1360 | | | YPTB_RS07520 | YPTB1360 | | YbjN domain-containing protein | WP_002208763.1 |
| yeps_YPTB1365 | | | YPTB_RS07545 | YPTB1365 | | YbjO family protein | WP_011192038.1 |
| yeps_YPTB1366 | rlmC | | YPTB_RS07550 | YPTB1366 | | 23S rRNA (uracil(747)-C(5))-methyltransferase RlmC | WP_011192039.1 |
| yeps_YPTB1369 | | | YPTB_RS07570 | YPTB1369 | | ABC transporter substrate-binding protein | WP_012413601.1 |
| yeps_YPTB1370 | | | YPTB_RS07575 | YPTB1370 | | iron ABC transporter permease | WP_197684172.1 |
| yeps_YPTB1371 | | | YPTB_RS07580 | YPTB1371 | | ABC transporter ATP-binding protein | WP_002211375.1 |
| yeps_YPTB1372 | | | YPTB_RS07585 | YPTB1372 | | nicotianamine synthase | WP_012304311.1 |
| yeps_YPTB1374 | | | YPTB_RS07595 | YPTB1374 | | DMT family transporter | WP_002223483.1 |
| yeps_YPTB1375 | artM | | YPTB_RS07605 | YPTB1375 | | arginine ABC transporter permease ArtM | WP_002211370.1 |
| yeps_YPTB1378 | artP | | YPTB_RS07620 | YPTB1378 | | arginine ABC transporter ATP-binding protein ArtP | WP_011192044.1 |
| yeps_YPTB1379 | | | YPTB_RS07625 | YPTB1379 | | chorismate mutase | WP_002211366.1 |
| yeps_YPTB1380 | | | YPTB_RS07630 | YPTB1380 | | lipoprotein | WP_002211365.1 |
| yeps_YPTB1382 | | | YPTB_RS07640 | YPTB1382 | | DUF2867 domain-containing protein | WP_011192045.1 |
| yeps_YPTB1385 | hcr | | YPTB_RS07655 | YPTB1385 | | NADH oxidoreductase | WP_011192047.1 |
| yeps_YPTB1388 | | | YPTB_RS07670 | YPTB1388 | | ATP-dependent endonuclease | WP_002211356.1 |
| yeps_YPTB1389 | | | YPTB_RS07675 | YPTB1389 | | VirK/YbjX family protein | WP_002211355.1 |
| yeps_YPTB1392 | cspD | cspH | YPTB_RS07690 | YPTB1392 | | cold shock-like protein CspD | WP_002211350.1 |
| yeps_YPTB1393 | clpS | | YPTB_RS07695 | YPTB1393 | | ATP-dependent Clp protease adapter ClpS | WP_002211349.1 |
| yeps_YPTB1396 | aat | | YPTB_RS07710 | YPTB1396 | | leucyl/phenylalanyl-tRNA--protein transferase | WP_002211346.1 |
| yeps_YPTB1402 | lolA | lplA | YPTB_RS07745 | YPTB1402 | | outer membrane lipoprotein chaperone LolA | WP_002211338.1 |
| yeps_YPTB1404 | serS | | YPTB_RS07755 | YPTB1404 | | serine--tRNA ligase | WP_002211336.1 |
| yeps_YPTB1415 | aroA | | YPTB_RS07810 | YPTB1415 | | 3-phosphoshikimate 1-carboxyvinyltransferase | WP_011192056.1 |
| yeps_YPTB1419 | | | YPTB_RS07830 | YPTB1419 | | ComEC family protein | WP_011192058.1 |
| yeps_YPTB1420 | msbA | | YPTB_RS07835 | YPTB1420 | | lipid A ABC transporter ATP-binding protein/permease MsbA | WP_002211320.1 |
| yeps_YPTB1421 | lpxK | | YPTB_RS07840 | YPTB1421 | | tetraacyldisaccharide 4'-kinase | WP_002211319.1 |
| yeps_YPTB1423 | | | YPTB_RS07855 | YPTB1423 | | cold-shock protein | WP_002211317.1 |
| yeps_YPTB1424 | | | YPTB_RS07860 | YPTB1424 | | hypothetical protein | WP_002211315.1 |
| yeps_YPTB1425 | kdsB | | YPTB_RS07865 | YPTB1425 | | 3-deoxy-manno-octulosonate cytidylyltransferase | WP_002211314.1 |
| yeps_YPTB1426 | | | YPTB_RS07870 | YPTB1426 | | YcbJ family phosphotransferase | WP_011192060.1 |
| yeps_YPTB1427 | cmoM | | YPTB_RS07880 | YPTB1427 | | tRNA uridine 5-oxyacetic acid(34) methyltransferase CmoM | WP_002211311.1 |
| yeps_YPTB1429 | mukE | kicA | YPTB_RS07890 | YPTB1429 | | chromosome partition protein MukE | WP_002211309.1 |
| yeps_YPTB1437 | pncB | | YPTB_RS07930 | YPTB1437 | | nicotinate phosphoribosyltransferase | WP_002228013.1 |
| yeps_YPTB1438 | pepN | | YPTB_RS07935 | YPTB1438 | | aminopeptidase N | WP_011192063.1 |
| yeps_YPTB1440 | | | YPTB_RS07945 | YPTB1440 | | cell division protein ZapC | WP_011192064.1 |
| yeps_YPTB1443 | | | YPTB_RS07960 | YPTB1443 | | ABC transporter ATP-binding protein | WP_002211292.1 |
| yeps_YPTB1450 | fabA | | YPTB_RS08000 | YPTB1450 | | bifunctional 3-hydroxydecanoyl-ACP dehydratase/trans-2-decenoyl-ACP isomerase | WP_002220006.1 |
| yeps_YPTB1451 | | | YPTB_RS08005 | YPTB1451 | | Lon protease family protein | WP_002224683.1 |
| yeps_YPTB1456 | yccS | | YPTB_RS08030 | YPTB1456 | | TIGR01666 family membrane protein | WP_011192070.1 |
| yeps_YPTB1457 | | | YPTB_RS08035 | YPTB1457 | | YccF domain-containing protein | WP_011192071.1 |
| yeps_YPTB1459 | | | YPTB_RS08045 | YPTB1459 | | methylglyoxal synthase | WP_002213060.1 |
| yeps_YPTB1462 | hspQ | | YPTB_RS08060 | YPTB1462 | | heat shock protein HspQ | WP_002213054.1 |
| yeps_YPTB1464 | yccX | | YPTB_RS08070 | YPTB1464 | | acylphosphatase | WP_002213049.1 |
| yeps_YPTB1465 | tusE | | YPTB_RS08075 | YPTB1465 | | sulfurtransferase TusE | WP_002213046.1 |
| yeps_YPTB1478 | | | YPTB_RS08150 | YPTB1478 | | SDR family oxidoreductase | WP_002213024.1 |
| yeps_YPTB1479 | | | YPTB_RS08155 | YPTB1479 | | lipocalin-like domain-containing protein | WP_002228570.1 |
| yeps_YPTB1480 | | | YPTB_RS08160 | YPTB1480 | | phosphopantetheine-binding protein | WP_002213017.1 |
| yeps_YPTB1481 | | | YPTB_RS08165 | YPTB1481 | | ACP S-malonyltransferase | WP_002213016.1 |
| yeps_YPTB1517 | | | YPTB_RS08355 | YPTB1517 | | S-(hydroxymethyl)glutathione dehydrogenase/class III alcohol dehydrogenase | WP_002224699.1 |
| yeps_YPTB1518 | | | YPTB_RS08360 | YPTB1518 | | LysR family transcriptional regulator | WP_002211958.1 |
| yeps_YPTB1520 | folE | | YPTB_RS08370 | YPTB1520 | | GTP cyclohydrolase I FolE | WP_002211960.1 |
| yeps_YPTB1521 | | | YPTB_RS08375 | YPTB1521 | | DUF418 family protein | WP_012413631.1 |
| yeps_YPTB1522 | mglB | | YPTB_RS08380 | YPTB1522 | | galactose/glucose ABC transporter substrate-binding protein MglB | WP_011192106.1 |
| yeps_YPTB1526 | | | YPTB_RS08400 | YPTB1526 | | NAD-dependent malic enzyme | WP_002211968.1 |
| yeps_YPTB1527 | cdd | | YPTB_RS08405 | YPTB1527 | | cytidine deaminase | WP_011192106.1 |
| yeps_YPTB1528 | | | YPTB_RS08410 | YPTB1528 | | CidB/LrgB family autolysis modulator | WP_011192107.1 |
| yeps_YPTB1536 | udk | | YPTB_RS08450 | YPTB1536 | | uridine kinase | WP_002211872.1 |
| yeps_YPTB1538 | asmA | | YPTB_RS08460 | YPTB1538 | | outer membrane assembly protein AsmA | WP_002211874.1 |
| yeps_YPTB1542 | | | YPTB_RS08480 | YPTB1542 | | lysine N(6)-hydroxylase/L-ornithine N(5)-oxygenase family protein | WP_002211878.1 |
| yeps_YPTB1543 | | | YPTB_RS08485 | YPTB1543 | | acetyltransferase | WP_002214999.1 |
| yeps_YPTB1548 | | | YPTB_RS08510 | YPTB1548 | | iron-siderophore ABC transporter substrate-binding protein | WP_011192116.1 |
| yeps_YPTB1552 | galU | | YPTB_RS08530 | YPTB1552 | | UTP--glucose-1-phosphate uridylyltransferase GalU | WP_002211887.1 |
| yeps_YPTB1556 | hisF | | YPTB_RS08555 | YPTB1556 | | imidazole glycerol phosphate synthase subunit HisF | WP_011192119.1 |
| yeps_YPTB1560 | hisC | | YPTB_RS08575 | YPTB1560 | | histidinol-phosphate transaminase | WP_011192121.1 |
| yeps_YPTB1561 | hisD | | YPTB_RS08580 | YPTB1561 | | histidinol dehydrogenase | WP_011192122.1 |
| yeps_YPTB1562 | hisG | | YPTB_RS08585 | YPTB1562 | | ATP phosphoribosyltransferase | WP_002211896.1 |
| yeps_YPTB1563 | | | YPTB_RS08590 | YPTB1563 | | SDR family oxidoreductase | WP_002211898.1 |
| yeps_YPTB1574 | | | YPTB_RS08660 | YPTB1574 | | thiol-disulfide oxidoreductase DCC family protein | WP_012413636.1 |
| yeps_YPTB1575 | | | YPTB_RS08665 | YPTB1575 | | SDR family oxidoreductase | WP_002216393.1 |
| yeps_YPTB1577 | | | YPTB_RS08675 | YPTB1577 | | L-fuconate dehydratase | WP_002211917.1 |
| yeps_YPTB1579 | argG | | YPTB_RS08685 | YPTB1579 | | argininosuccinate synthase | WP_002211920.1 |
| yeps_YPTB1580 | | | YPTB_RS08690 | YPTB1580 | | EmmdR/YeeO family multidrug/toxin efflux MATE transporter | WP_002224723.1 |
| yeps_YPTB1604 | | | YPTB_RS08830 | YPTB1604 | | TRAP transporter large permease | WP_002213124.1 |
| yeps_YPTB1605 | | | YPTB_RS08835 | YPTB1605 | | TRAP transporter small permease | WP_002213122.1 |
| yeps_YPTB1609 | | | YPTB_RS08860 | YPTB1609 | | MFS transporter | WP_002230721.1 |
| yeps_YPTB1611 | mtfA | | YPTB_RS08875 | YPTB1611 | | DgsA anti-repressor MtfA | WP_002211042.1 |
| yeps_YPTB1612 | | | YPTB_RS08885 | YPTB1612 | | Ycil family protein | WP_002211043.1 |
| yeps_YPTB1613 | | | YPTB_RS08890 | YPTB1613 | | serine protein kinase RIO | WP_002211044.1 |
| yeps_YPTB1633 | | | YPTB_RS09005 | YPTB1633 | | PTS mannose/fructose/sorbose transporter subunit IIC | WP_002211068.1 |
| yeps_YPTB1634 | manX | gptB,ptsL | YPTB_RS09010 | YPTB1634 | | PTS mannose transporter subunit IIAB | WP_002211069.1 |
| yeps_YPTB1635 | | | YPTB_RS09015 | YPTB1635 | | TerC family protein | WP_002231099.1 |
| yeps_YPTB1641 | | | YPTB_RS09045 | YPTB1641 | | 5-carboxymethyl-2-hydroxymuconate Delta-isomerase | WP_002211073.1 |
| yeps_YPTB1642 | hpaH | hpcG | YPTB_RS09050 | YPTB1642 | | 2-oxo-hepta-3-ene-1,7-dioic acid hydratase | WP_011192151.1 |
| yeps_YPTB1645 | hpaB | | YPTB_RS09065 | YPTB1645 | | 4-hydroxyphenylacetate 3-monooxygenase, oxygenase component | WP_002211077.1 |
| yeps_YPTB1646 | hpaC | | YPTB_RS09070 | YPTB1646 | | 4-hydroxyphenylacetate 3-monooxygenase, reductase component | WP_002211078.1 |
| yeps_YPTB1648 | | | YPTB_RS09080 | YPTB1648 | | CoA pyrophosphatase | WP_002211080.1 |
| yeps_YPTB1649 | pabB | | YPTB_RS09085 | YPTB1649 | | aminodeoxychorismate synthase component 1 | WP_011192153.1 |
| yeps_YPTB1650 | | | YPTB_RS09090 | YPTB1650 | | YoaH family protein | WP_011192154.1 |
| yeps_YPTB1652 | dbpA | | YPTB_RS09100 | YPTB1652 | | ATP-dependent RNA helicase DbpA | WP_011192156.1 |
| yeps_YPTB1653 | | | YPTB_RS09105 | YPTB1653 | | YebG family protein | WP_002211085.1 |
| yeps_YPTB1654 | | | YPTB_RS09110 | YPTB1654 | | N(4)-acetylcytidine aminohydrolase | WP_002211086.1 |
| yeps_YPTB1655 | | | YPTB_RS09115 | YPTB1655 | | protein YebF | WP_012304226.1 |
| yeps_YPTB1669 | | | YPTB_RS09195 | YPTB1669 | | flagellar synthesis protein FlgN | WP_002211107.1 |
| yeps_YPTB1670 | flgM | | YPTB_RS09200 | YPTB1670 | | anti-sigma-28 factor FlgM | WP_002211108.1 |
| yeps_YPTB1674 | flgD | lfgD | YPTB_RS09220 | YPTB1674 | | flagellar hook assembly protein FlgD | WP_011192164.1 |

| | | | | | | | |
|---|---|---|---|---|---|---|---|
| yeps_YPTB1676 | | | YPTB_RS09230 | YPTB1676 | | flagellar basal body rod protein FlgF | WP_011192165.1 |
| yeps_YPTB1677 | flgG | | YPTB_RS09235 | YPTB1677 | | flagellar basal-body rod protein FlgG | WP_011192166.1 |
| yeps_YPTB1678 | flgH | flaFVIII,flaY | YPTB_RS09240 | YPTB1678 | | flagellar basal body L-ring protein FlgH | WP_002211116.1 |
| yeps_YPTB1679 | | | YPTB_RS09245 | YPTB1679 | | flagellar basal body P-ring protein FlgI | WP_002211117.1 |
| yeps_YPTB1682 | flgL | flaT,flaU | YPTB_RS09260 | YPTB1682 | | flagellar hook-associated protein FlgL | WP_002211120.1 |
| yeps_YPTB1683 | | | YPTB_RS09265 | YPTB1683 | | sugar-binding transcriptional regulator | WP_041175448.1 |
| yeps_YPTB1684 | | | YPTB_RS09270 | YPTB1684 | | bifunctional aldolase/short-chain dehydrogenase | WP_011192168.1 |
| yeps_YPTB1686 | | | YPTB_RS09280 | YPTB1686 | | substrate-binding domain-containing protein | WP_002211125.1 |
| yeps_YPTB1687 | | | YPTB_RS09285 | YPTB1687 | | sugar ABC transporter ATP-binding protein | WP_011192170.1 |
| yeps_YPTB1688 | | | YPTB_RS09290 | YPTB1688 | | ABC transporter permease | WP_011192171.1 |
| yeps_YPTB1691 | fliR | mopE | YPTB_RS09305 | YPTB1691 | | flagellar type III secretion system protein FliR | WP_011192174.1 |
| yeps_YPTB1693 | fliP | mopC | YPTB_RS09315 | YPTB1693 | | flagellar type III secretion system pore protein FliP | WP_002220494.1 |
| yeps_YPTB1696 | fliM | CheC2,flaAII,flaQII | YPTB_RS09330 | YPTB1696 | | flagellar motor switch protein FliM | WP_011192177.1 |
| yeps_YPTB1697 | fliL | cheC1,flaAI,flaQI | YPTB_RS09335 | YPTB1697 | | flagellar basal body-associated protein FliL | WP_002227961.1 |
| yeps_YPTB1699 | fliJ | flaO,flaS | YPTB_RS09345 | YPTB1699 | | flagella biosynthesis chaperone FliJ | WP_002211139.1 |
| yeps_YPTB1701 | fliH | flaAII.3,flaBIII | YPTB_RS09355 | YPTB1701 | | flagellar assembly protein FliH | WP_011192180.1 |
| yeps_YPTB1703 | fliF | flaAII.1,flaBI | YPTB_RS09365 | YPTB1703 | | flagellar M-ring protein FliF | WP_011192181.1 |
| yeps_YPTB1704 | fliE | flaAI,flaN | YPTB_RS09370 | YPTB1704 | | flagellar hook-basal body complex protein FliE | WP_011192182.1 |
| yeps_YPTB1705 | | | YPTB_RS09375 | YPTB1705 | | hypothetical protein | WP_002215990.1 |
| yeps_YPTB1708 | | | YPTB_RS09390 | YPTB1708 | | DNA-3-methyladenine glycosylase | WP_002211145.1 |
| yeps_YPTB1709 | | | YPTB_RS09395 | YPTB1709 | | metal-dependent phosphohydrolase | WP_011192184.1 |
| yeps_YPTB1710 | | | YPTB_RS09400 | YPTB1710 | | AraC family transcriptional regulator | WP_002211147.1 |
| yeps_YPTB1711 | fliT | | YPTB_RS09410 | YPTB1711 | | flagella biosynthesis regulatory protein FliT | WP_011192185.1 |
| yeps_YPTB1712 | fliS | lafC | YPTB_RS09415 | YPTB1712 | | flagellar export chaperone FliS | WP_011192186.1 |
| yeps_YPTB1713 | fliD | flaV,flbC | YPTB_RS09420 | YPTB1713 | | flagellar filament capping protein FliD | WP_011192187.1 |
| yeps_YPTB1714 | | | YPTB_RS09425 | YPTB1714 | | FliC/FljB family flagellin | WP_002211152.1 |
| yeps_YPTB1716 | fliZ | | YPTB_RS09435 | YPTB1716 | | flagella biosynthesis regulatory protein FliZ | WP_002211154.1 |
| yeps_YPTB1718 | tcyJ | | YPTB_RS09445 | YPTB1718 | | cystine ABC transporter substrate-binding protein | WP_002211157.1 |
| yeps_YPTB1719 | tcyL | | YPTB_RS09450 | YPTB1719 | | cystine ABC transporter permease | WP_011192190.1 |
| yeps_YPTB1722 | | | YPTB_RS09465 | YPTB1722 | | DUF6516 family protein | WP_012413668.1 |
| yeps_YPTB1725 | | | YPTB_RS09485 | YPTB1725 | | FTR1 family protein | WP_002403646.1 |
| yeps_YPTB1728 | wrbA | | YPTB_RS09500 | YPTB1728 | | NAD(P)H:quinone oxidoreductase | WP_002211168.1 |
| yeps_YPTB1729 | | | YPTB_RS09505 | YPTB1729 | | N-acetylneuraminate epimerase | WP_002211169.1 |
| yeps_YPTB1912 | | | YPTB_RS10495 | YPTB1912 | | Gfo/Idh/MocA family oxidoreductase | WP_011192351.1 |
| yeps_YPTB1915 | | | YPTB_RS10510 | YPTB1915 | | carbohydrate ABC transporter permease | WP_002213186.1 |
| yeps_YPTB1916 | ugpC | | YPTB_RS10515 | YPTB1916 | | sn-glycerol-3-phosphate ABC transporter ATP-binding protein UgpC | WP_011192353.1 |
| yeps_YPTB1917 | | | YPTB_RS10530 | YPTB1917 | | molecular chaperone | WP_002227954.1 |
| yeps_YPTB1919 | | | YPTB_RS10540 | YPTB1919 | | fimbrial biogenesis outer membrane usher protein | WP_032466356.1 |
| yeps_YPTB1920 | | | YPTB_RS10545 | YPTB1920 | | molecular chaperone | WP_011192355.1 |
| yeps_YPTB1926 | ripC | | YPTB_RS10575 | YPTB1926 | | itaconate degradation C-C-lyase RipC | WP_002212068.1 |
| yeps_YPTB1927 | ripR | | YPTB_RS10580 | YPTB1927 | | itaconate degradation transcriptional regulator RipR | WP_012105184.1 |
| yeps_YPTB1928 | | | YPTB_RS10585 | YPTB1928 | | carboxypeptidase-like regulatory domain-containing protein | WP_011192359.1 |
| yeps_YPTB1929 | | | YPTB_RS10590 | YPTB1929 | | hypothetical protein | WP_011192360.1 |
| yeps_YPTB1931 | | | YPTB_RS10600 | YPTB1931 | | amidohydrolase | WP_011192361.1 |
| yeps_YPTB1933 | nqrE | nqr5 | YPTB_RS10610 | YPTB1933 | | NADH:ubiquinone reductase (Na(+)-transporting) subunit E | WP_002212061.1 |
| yeps_YPTB1944 | | | YPTB_RS10670 | YPTB1944 | | ABC transporter ATP-binding protein | WP_011192364.1 |
| yeps_YPTB1946 | | | YPTB_RS10680 | YPTB1946 | | cytochrome c | WP_002227951.1 |
| yeps_YPTB1950 | pgaB | | YPTB_RS10700 | YPTB1950 | | poly-beta-1,6-N-acetyl-D-glucosamine N-deacetylase PgaB | WP_002212043.1 |
| yeps_YPTB1951 | | | YPTB_RS10705 | YPTB1951 | | poly-beta-1,6 N-acetyl-D-glucosamine synthase | WP_002224456.1 |
| yeps_YPTB1952 | pgaD | | YPTB_RS10710 | YPTB1952 | | poly-beta-1,6-N-acetyl-D-glucosamine biosynthesis protein PgaD | WP_011192367.1 |
| yeps_YPTB1954 | | | YPTB_RS10720 | YPTB1954 | | hypothetical protein | WP_002212038.1 |
| yeps_YPTB1957 | narX | narR | YPTB_RS10735 | YPTB1957 | | nitrate/nitrite two-component system sensor histidine kinase NarX | WP_011192369.1 |
| yeps_YPTB1965 | hutG | | YPTB_RS10780 | YPTB1965 | | N-formylglutamate deformylase | WP_011192373.1 |
| yeps_YPTB1968 | | | YPTB_RS10795 | YPTB1968 | | formimidoylglutamate deiminase | WP_011192376.1 |
| yeps_YPTB1969 | | | YPTB_RS10800 | YPTB1969 | | HutD family protein | WP_011192377.1 |
| yeps_YPTB1972 | | | YPTB_RS10815 | YPTB1972 | | NAD(P)-dependent oxidoreductase | WP_011192380.1 |
| yeps_YPTB1974 | | | YPTB_RS10825 | YPTB1974 | | coenzyme F390 synthetase | WP_011192382.1 |
| yeps_YPTB1975 | | | YPTB_RS10830 | YPTB1975 | | NAD(P)-dependent oxidoreductase | WP_012413710.1 |
| yeps_YPTB1977 | | | YPTB_RS10840 | YPTB1977 | | hypothetical protein | WP_002211270.1 |
| yeps_YPTB1978 | | | YPTB_RS10845 | YPTB1978 | | glycosyltransferase | WP_011192385.1 |
| yeps_YPTB1983 | | | YPTB_RS10875 | YPTB1983 | | RpiB/LacA/LacB family sugar-phosphate isomerase | WP_002211263.1 |
| yeps_YPTB1984 | | | YPTB_RS10880 | YPTB1984 | | class I SAM-dependent methyltransferase | WP_002211262.1 |
| yeps_YPTB1985 | | | YPTB_RS10885 | YPTB1985 | | SDR family oxidoreductase | WP_011192390.1 |
| yeps_YPTB1986 | | | YPTB_RS10890 | YPTB1986 | | hypothetical protein | WP_002211260.1 |
| yeps_YPTB1987 | | | YPTB_RS10895 | YPTB1987 | | hypothetical protein | WP_002211259.1 |
| yeps_YPTB1990 | | | YPTB_RS10910 | YPTB1990 | | cupin domain-containing protein | WP_002211256.1 |
| yeps_YPTB1991 | | | YPTB_RS10915 | YPTB1991 | | carboxymuconolactone decarboxylase family protein | WP_011192392.1 |
| yeps_YPTB1992 | | | YPTB_RS10920 | YPTB1992 | | heavy metal sensor histidine kinase | WP_002211254.1 |
| yeps_YPTB1993 | | | YPTB_RS10925 | YPTB1993 | | heavy metal response regulator transcription factor | WP_011192393.1 |
| yeps_YPTB1998 | ychF | | YPTB_RS10950 | YPTB1998 | | redox-regulated ATPase YchF | WP_002211245.1 |
| yeps_YPTB2000 | ychH | | YPTB_RS10960 | YPTB2000 | | stress-induced protein YchH | WP_002211242.1 |
| yeps_YPTB2003 | lolB | | YPTB_RS10985 | YPTB2003 | | lipoprotein insertase outer membrane protein LolB | WP_011192397.1 |
| yeps_YPTB2004 | hemA | | YPTB_RS10990 | YPTB2004 | | glutamyl-tRNA reductase | WP_002211237.1 |
| yeps_YPTB2006 | prmC | | YPTB_RS11000 | YPTB2006 | | peptide chain release factor N(5)-glutamine methyltransferase | WP_011192399.1 |
| yeps_YPTB2014 | | | YPTB_RS11050 | YPTB2014 | | ABC transporter permease | WP_011192400.1 |
| yeps_YPTB2015 | | | YPTB_RS11055 | YPTB2015 | | ABC transporter permease | WP_002211226.1 |
| yeps_YPTB2020 | | | YPTB_RS11085 | YPTB2020 | | molecular chaperone | WP_002211220.1 |
| yeps_YPTB2021 | | | YPTB_RS11090 | YPTB2021 | | lipoprotein | WP_002211218.1 |
| yeps_YPTB2029 | argS | | YPTB_RS11130 | YPTB2029 | | arginine--tRNA ligase | WP_011192408.1 |
| yeps_YPTB2033 | cmoA | | YPTB_RS11150 | YPTB2033 | | carboxy-S-adenosyl-L-methionine synthase CmoA | WP_002211207.1 |
| yeps_YPTB2034 | | | YPTB_RS11155 | YPTB2034 | | MAPEG family protein | WP_002211206.1 |
| yeps_YPTB2037 | nudB | ntpA | YPTB_RS11170 | YPTB2037 | | dihydroneopterin triphosphate diphosphatase | WP_002211203.1 |
| yeps_YPTB2039 | ruvC | | YPTB_RS11180 | YPTB2039 | | crossover junction endodeoxyribonuclease RuvC | WP_002211201.1 |
| yeps_YPTB2041 | ruvB | | YPTB_RS11190 | YPTB2041 | | Holliday junction branch migration DNA helicase RuvB | WP_002211198.1 |
| yeps_YPTB2043 | znuC | | YPTB_RS11200 | YPTB2043 | | zinc ABC transporter ATP-binding protein ZnuC | WP_011192413.1 |
| yeps_YPTB2044 | znuA | | YPTB_RS11205 | YPTB2044 | | zinc ABC transporter substrate-binding protein ZnuA | WP_011192414.1 |
| yeps_YPTB2048 | | | YPTB_RS11225 | YPTB2048 | | MurR/RpiR family transcriptional regulator | WP_002211192.1 |
| yeps_YPTB2049 | zwf | | YPTB_RS11230 | YPTB2049 | | glucose-6-phosphate dehydrogenase | WP_011906215.1 |
| yeps_YPTB2054 | | | YPTB_RS11260 | YPTB2054 | | ATP-dependent DNA helicase | WP_011192419.1 |
| yeps_YPTB2055 | tsaB | | YPTB_RS11265 | YPTB2055 | | tRNA (adenosine(37)-N6)-threonylcarbamoyltransferase complex dimerization subunit type 1 TsaB | WP_011192420.1 |
| yeps_YPTB2062 | | | YPTB_RS11300 | YPTB2062 | | YcgL domain-containing protein | WP_002211743.1 |
| yeps_YPTB2063 | | | YPTB_RS11305 | YPTB2063 | | lytic murein transglycosylase | WP_011192421.1 |
| yeps_YPTB2065 | | | YPTB_RS11315 | YPTB2065 | | YcgN family cysteine cluster protein | WP_002211739.1 |
| yeps_YPTB2076 | | | YPTB_RS11370 | YPTB2076 | | pirin family protein | WP_011192427.1 |
| yeps_YPTB2085 | | | YPTB_RS11415 | YPTB2085 | | YeaC family protein | WP_002217933.1 |
| yeps_YPTB2087 | ansA | | YPTB_RS11425 | YPTB2087 | | asparaginase | WP_011192430.1 |
| yeps_YPTB2088 | sppA | | YPTB_RS11430 | YPTB2088 | | signal peptide peptidase SppA | WP_011192431.1 |
| yeps_YPTB2091 | | | YPTB_RS11445 | YPTB2091 | | DNA topoisomerase III | WP_041175458.1 |
| yeps_YPTB2119 | | | YPTB_RS11590 | YPTB2119 | | septation protein A | WP_002210640.1 |
| yeps_YPTB2124 | | | YPTB_RS11615 | YPTB2124 | | BON domain-containing protein | WP_002210636.1 |
| yeps_YPTB2127 | trpCF | | YPTB_RS11630 | YPTB2127 | | bifunctional indole-3-glycerol-phosphate synthase TrpC/phosphoribosylanthranilate isomerase TrpF | WP_011192443.1 |
| yeps_YPTB2130 | | | YPTB_RS11645 | YPTB2130 | | anthranilate synthase component 1 | WP_002228454.1 |
| yeps_YPTB2136 | cobO | | YPTB_RS11675 | YPTB2136 | | cob(I)yrinic acid a,c-diamide adenosyltransferase | WP_002210626.1 |
| yeps_YPTB2138 | sohB | | YPTB_RS11685 | YPTB2138 | | protease SohB | WP_002210624.1 |
| yeps_YPTB2153 | | | YPTB_RS11765 | YPTB2153 | | DUF2164 domain-containing protein | WP_002210609.1 |
| yeps_YPTB2154 | fsa | | YPTB_RS11770 | YPTB2154 | | fructose-6-phosphate aldolase | WP_002215192.1 |
| yeps_YPTB2155 | | | YPTB_RS11775 | YPTB2155 | | YbdD/YjiX family protein | WP_002210607.1 |
| yeps_YPTB2157 | | | YPTB_RS11785 | YPTB2157 | | exoribonuclease II | WP_011192452.1 |
| yeps_YPTB2158 | | | YPTB_RS11790 | YPTB2158 | | MFS transporter | WP_011192453.1 |
| yeps_YPTB2159 | | | YPTB_RS11795 | YPTB2159 | | SDR family oxidoreductase | WP_011192454.1 |
| yeps_YPTB2161 | | | YPTB_RS11805 | YPTB2161 | | electron transport complex subunit E | WP_011192455.1 |
| yeps_YPTB2162 | rsxG | | YPTB_RS11810 | YPTB2162 | | electron transport complex subunit RsxG | WP_002210600.1 |
| yeps_YPTB2168 | | | YPTB_RS11840 | YPTB2168 | | DUF2569 domain-containing protein | WP_002210594.1 |
| yeps_YPTB2170 | | | YPTB_RS11850 | YPTB2170 | | HlyD family type I secretion periplasmic adaptor subunit | WP_011192461.1 |
| yeps_YPTB2173 | | | YPTB_RS11865 | YPTB2173 | | ribulokinase | WP_011192464.1 |
| yeps_YPTB2175 | araG | | YPTB_RS11875 | YPTB2175 | | L-arabinose ABC transporter ATP-binding protein AraG | WP_002210588.1 |
| yeps_YPTB2176 | araH | | YPTB_RS11880 | YPTB2176 | | L-arabinose ABC transporter permease AraH | WP_164491550.1 |
| yeps_YPTB2177 | araC | | YPTB_RS11885 | YPTB2177 | | arabinose operon transcriptional regulator AraC | WP_011192467.1 |
| yeps_YPTB2178 | | | YPTB_RS11890 | YPTB2178 | | oxidoreductase | WP_002210585.1 |
| yeps_YPTB2179 | | | YPTB_RS11895 | YPTB2179 | | bile acid:sodium symporter | WP_011192468.1 |
| yeps_YPTB2184 | | | YPTB_RS11925 | YPTB2184 | | YdgA family protein | WP_011192473.1 |
| yeps_YPTB2200 | | | YPTB_RS12010 | YPTB2200 | | pyridoxal phosphate-dependent aminotransferase | WP_011192488.1 |
| yeps_YPTB2206 | | | YPTB_RS12040 | YPTB2206 | | sugar ABC transporter ATP-binding protein | WP_011192492.1 |
| yeps_YPTB2209 | | | YPTB_RS12055 | YPTB2209 | | ABC transporter permease | WP_002211034.1 |
| yeps_YPTB2210 | | | YPTB_RS12060 | YPTB2210 | | ABC transporter ATP-binding protein | WP_011192495.1 |
| yeps_YPTB2211 | | | YPTB_RS12065 | YPTB2211 | | AMP nucleosidase | WP_002223601.1 |
| yeps_YPTB2217 | ilvN | | YPTB_RS12095 | YPTB2217 | | acetolactate synthase small subunit | WP_002230908.1 |
| yeps_YPTB2218 | yjjG | | YPTB_RS12100 | YPTB2218 | | pyrimidine 5'-nucleotidase | WP_002211025.1 |
| yeps_YPTB2221 | ogt | | YPTB_RS12120 | YPTB2221 | | methylated-DNA--[protein]-cysteine S-methyltransferase | WP_002211023.1 |
| yeps_YPTB2222 | | | YPTB_RS12125 | YPTB2222 | | FNR family transcription factor | WP_011192502.1 |

| | | | | | | | |
|---|---|---|---|---|---|---|---|
| yeps_YPTB2225 | pntA | | YPTB_RS12140 | YPTB2225 | | Re/Si-specific NAD(P)(+) transhydrogenase subunit alpha | WP_011192503.1 |
| yeps_YPTB2226 | | | | YPTB2226 | | hypothetical protein | CAH21464.1 |
| yeps_YPTB2227 | | | YPTB_RS12150 | YPTB2227 | | DUF1471 family protein YdgH | WP_002211018.1 |
| yeps_YPTB2228 | | | YPTB_RS12155 | YPTB2228 | | amino acid permease | WP_002211017.1 |
| yeps_YPTB2229 | | | YPTB_RS12160 | YPTB2229 | | hypothetical protein | WP_002211016.1 |
| yeps_YPTB2230 | rstA | urpT | YPTB_RS12165 | YPTB2230 | | two-component system response regulator RstA | WP_002211015.1 |
| yeps_YPTB2231 | rstB | uspT | YPTB_RS12170 | YPTB2231 | | two-component system sensor histidine kinase RstB | WP_011192505.1 |
| yeps_YPTB2232 | | | YPTB_RS12175 | YPTB2232 | | carboxypeptidase M32 | WP_011192506.1 |
| yeps_YPTB2239 | | | YPTB_RS12215 | YPTB2239 | | helix-turn-helix domain-containing protein | WP_002211007.1 |
| yeps_YPTB2241 | hrpA | | YPTB_RS12225 | YPTB2241 | | ATP-dependent RNA helicase HrpA | WP_012413748.1 |
| yeps_YPTB2242 | azoR | | YPTB_RS12230 | YPTB2242 | | FMN-dependent NADH-azoreductase | WP_002211004.1 |
| yeps_YPTB2246 | | | YPTB_RS12250 | YPTB2246 | | YnbE family lipoprotein | WP_002211000.1 |
| yeps_YPTB2248 | | | YPTB_RS12260 | YPTB2248 | | 2-hydroxyacid dehydrogenase | WP_002210998.1 |
| yeps_YPTB2251 | | | YPTB_RS12275 | YPTB2251 | | MgtC/SapB family protein | WP_011192514.1 |
| yeps_YPTB2252 | | | YPTB_RS12280 | YPTB2252 | | multidrug efflux SMR transporter | WP_002210994.1 |
| yeps_YPTB2254 | ttcA | | YPTB_RS12290 | YPTB2254 | | tRNA 2-thiocytidine(32) synthetase TtcA | WP_011192516.1 |
| yeps_YPTB2258 | | | YPTB_RS12310 | YPTB2258 | | ABC transporter substrate-binding protein | WP_002210987.1 |
| yeps_YPTB2262 | tyrR | | YPTB_RS12330 | YPTB2262 | | transcriptional regulator TyrR | WP_011192519.1 |
| yeps_YPTB2264 | | | YPTB_RS12340 | YPTB2264 | | RES family NAD+ phosphorylase | WP_011192520.1 |
| yeps_YPTB2265 | | | YPTB_RS12345 | YPTB2265 | | YcjF family protein | WP_002210980.1 |
| yeps_YPTB2267 | pspD | | YPTB_RS12355 | YPTB2267 | | phage shock protein PspD | WP_002216375.1 |
| yeps_YPTB2268 | pspC | | YPTB_RS12360 | YPTB2268 | | envelope stress response membrane protein PspC | WP_002210977.1 |
| yeps_YPTB2269 | pspB | | YPTB_RS12365 | YPTB2269 | | envelope stress response membrane protein PspB | WP_002210976.1 |
| yeps_YPTB2270 | pspA | | YPTB_RS12370 | YPTB2270 | | phage shock protein PspA | WP_011192521.1 |
| yeps_YPTB2276 | sapF | | YPTB_RS12405 | YPTB2276 | | peptide ABC transporter ATP-binding protein SapF | WP_002210969.1 |
| yeps_YPTB2279 | | | YPTB_RS12420 | YPTB2279 | | YIP1 family protein | WP_002210965.1 |
| yeps_YPTB2280 | | | YPTB_RS12425 | YPTB2280 | | DUF3811 domain-containing protein | WP_002210964.1 |
| yeps_YPTB2281 | gstA | | YPTB_RS12430 | YPTB2281 | | glutathione transferase GstA | WP_002210962.1 |
| yeps_YPTB2282 | pdxY | | YPTB_RS12435 | YPTB2282 | | pyridoxal kinase PdxY | WP_011192523.1 |
| yeps_YPTB2284 | pdxH | | YPTB_RS12445 | YPTB2284 | | pyridoxamine 5'-phosphate oxidase | WP_002210959.1 |
| yeps_YPTB2285 | | | YPTB_RS12450 | YPTB2285 | | lipoprotein | WP_002218323.1 |
| yeps_YPTB2292 | | | YPTB_RS12485 | YPTB2292 | | TetR/AcrR family transcriptional regulator | WP_002210951.1 |
| yeps_YPTB2293 | | | YPTB_RS12490 | YPTB2293 | | alkene reductase | WP_011192525.1 |
| yeps_YPTB2295 | gloA | | YPTB_RS12500 | YPTB2295 | | lactoylglutathione lyase | WP_011192527.1 |
| yeps_YPTB2298 | | | YPTB_RS12520 | YPTB2298 | | C40 family peptidase | WP_012105035.1 |
| yeps_YPTB2299 | sodB | | YPTB_RS12525 | YPTB2299 | | superoxide dismutase [Fe] | WP_011192530.1 |
| yeps_YPTB2300 | purR | | YPTB_RS12530 | YPTB2300 | | HTH-type transcriptional repressor PurR | WP_002210943.1 |
| yeps_YPTB2303 | cfa | cdfA | YPTB_RS12545 | YPTB2303 | | cyclopropane fatty acyl phospholipid synthase | WP_002210940.1 |
| yeps_YPTB2304 | | | YPTB_RS12550 | YPTB2304 | | riboflavin synthase | WP_011192532.1 |
| yeps_YPTB2309 | sufE | | YPTB_RS12590 | YPTB2309 | | cysteine desulfuration protein SufE | WP_002211804.1 |
| yeps_YPTB2310 | sufS | | YPTB_RS12595 | YPTB2310 | | cysteine desulfurase SufS | WP_011192533.1 |
| yeps_YPTB2313 | sufB | | YPTB_RS12610 | YPTB2313 | | Fe-S cluster assembly protein SufB | WP_011192536.1 |
| yeps_YPTB2314 | sufA | | YPTB_RS12615 | YPTB2314 | | Fe-S cluster assembly scaffold SufA | WP_002211809.1 |
| yeps_YPTB2382 | | | YPTB_RS12960 | YPTB2382 | | spore coat U domain-containing protein | WP_002216613.1 |
| yeps_YPTB2387 | | | YPTB_RS12985 | YPTB2387 | | YebV family protein | WP_002210859.1 |
| yeps_YPTB2388 | | | YPTB_RS12990 | YPTB2388 | | ASCH domain-containing protein | WP_011192562.1 |
| yeps_YPTB2393 | | | YPTB_RS13020 | YPTB2393 | | hypothetical protein | WP_002210869.1 |
| yeps_YPTB2395 | | | YPTB_RS13030 | YPTB2395 | | N-acetylmuramoyl-L-alanine amidase | WP_011192565.1 |
| yeps_YPTB2398 | | | YPTB_RS13050 | YPTB2398 | | chemotaxis response regulator protein-glutamate methylesterase | WP_011192566.1 |
| yeps_YPTB2399 | cheR | cheX | YPTB_RS13055 | YPTB2399 | | protein-glutamate O-methyltransferase CheR | WP_002214065.1 |
| yeps_YPTB2401 | | | YPTB_RS13065 | YPTB2401 | | methyl-accepting chemotaxis protein | WP_011192568.1 |
| yeps_YPTB2413 | dsrB | | YPTB_RS24100 | YPTB2413 | | protein DsrB | WP_002210892.1 |
| yeps_YPTB2418 | | | YPTB_RS13155 | YPTB2418 | | Lrp/AsnC family transcriptional regulator | WP_002210900.1 |
| yeps_YPTB2419 | | | YPTB_RS13160 | YPTB2419 | | Kdo hydroxylase family protein | WP_012413770.1 |
| yeps_YPTB2420 | ydfZ | | YPTB_RS13165 | YPTB2420 | | putative selenium delivery protein YdfZ | WP_002210902.1 |
| yeps_YPTB2421 | | | YPTB_RS13170 | YPTB2421 | | AppA family phytase/histidine-type acid phosphatase | WP_011192581.1 |
| yeps_YPTB2425 | | | YPTB_RS13195 | YPTB2425 | | hypothetical protein | WP_002210908.1 |
| yeps_YPTB2430 | mnmA | | YPTB_RS13220 | YPTB2430 | | tRNA 2-thiouridine(34) synthase MnmA | WP_002210913.1 |
| yeps_YPTB2435 | phoQ | | YPTB_RS13245 | YPTB2435 | | two-component system sensor histidine kinase PhoQ | WP_002210918.1 |
| yeps_YPTB2438 | cobB | | YPTB_RS13260 | YPTB2438 | | NAD-dependent protein deacylase | WP_002210921.1 |
| yeps_YPTB2439 | nagK | | YPTB_RS13265 | YPTB2439 | | N-acetylglucosamine kinase | WP_011192586.1 |
| yeps_YPTB2440 | lolE | | YPTB_RS13270 | YPTB2440 | | lipoprotein-releasing ABC transporter permease subunit LolE | WP_011192587.1 |
| yeps_YPTB2441 | lolD | | YPTB_RS13275 | YPTB2441 | | lipoprotein-releasing ABC transporter ATP-binding protein LolD | WP_011192588.1 |
| yeps_YPTB2449 | nagZ | | YPTB_RS13315 | YPTB2449 | | beta-N-acetylhexosaminidase | WP_032466495.1 |
| yeps_YPTB2451 | lpoB | | YPTB_RS13325 | YPTB2451 | | penicillin-binding protein activator LpoB | WP_011192593.1 |
| yeps_YPTB2452 | | | YPTB_RS13330 | YPTB2452 | | YcfL family protein | WP_002213088.1 |
| yeps_YPTB2453 | hinT | | YPTB_RS13335 | YPTB2453 | | purine nucleoside phosphoramidase | WP_002213087.1 |
| yeps_YPTB2463 | ptsG | glcA,umG | YPTB_RS13390 | YPTB2463 | | PTS glucose transporter subunit IIBC | WP_011192602.1 |
| yeps_YPTB2464 | | | YPTB_RS13395 | YPTB2464 | | metal-dependent hydrolase | WP_002213084.1 |
| yeps_YPTB2465 | holB | | YPTB_RS13400 | YPTB2465 | | DNA polymerase III subunit delta' | WP_011192603.1 |
| yeps_YPTB2466 | | | YPTB_RS13405 | YPTB2466 | | dTMP kinase | WP_011192604.1 |
| yeps_YPTB2469 | fabF | fabJ | YPTB_RS13420 | YPTB2469 | | beta-ketoacyl-ACP synthase II | WP_002213079.1 |
| yeps_YPTB2472 | fabD | tfpA | YPTB_RS13435 | YPTB2472 | | ACP S-malonyltransferase | WP_002210934.1 |
| yeps_YPTB2473 | | | YPTB_RS13440 | YPTB2473 | | ketoacyl-ACP synthase III | WP_002210933.1 |
| yeps_YPTB2476 | yceD | | YPTB_RS13455 | YPTB2476 | | 23S rRNA accumulation protein YceD | WP_002210930.1 |
| yeps_YPTB2480 | | | YPTB_RS22940 | YPTB2480 | | IS3 family transposase | CAH21718.1 |
| yeps_YPTB2481 | | | YPTB_RS13475 | YPTB2481 | | antibiotic biosynthesis monooxygenase | WP_002213107.1 |
| yeps_YPTB2482 | pyrC | | YPTB_RS13480 | YPTB2482 | | dihydroorotase | WP_011192610.1 |
| yeps_YPTB2483 | dinI | | YPTB_RS13485 | YPTB2483 | | DNA damage-inducible protein I | WP_002213110.1 |
| yeps_YPTB2484 | bssS | | YPTB_RS13495 | YPTB2484 | | biofilm formation regulator BssS | WP_011192611.1 |
| yeps_YPTB2488 | | | YPTB_RS13515 | YPTB2488 | | rhodanese-related sulfurtransferase | WP_002211854.1 |
| yeps_YPTB2489 | | | | YPTB2489 | | putative membrane protein | CAH21727.1 |
| yeps_YPTB2503 | fae | | YPTB_RS13600 | YPTB2503 | | formaldehyde-activating enzyme | WP_011192622.1 |
| yeps_YPTB2504 | | | YPTB_RS13605 | YPTB2504 | | aldo/keto reductase | WP_011192623.1 |
| yeps_YPTB2505 | cas6f | | YPTB_RS13610 | YPTB2505 | | type I-F CRISPR-associated endoribonuclease Cas6/Csy4 | WP_002211866.1 |
| yeps_YPTB2506 | csy3 | | YPTB_RS13615 | YPTB2506 | | type I-F CRISPR-associated protein Csy3 | WP_011192624.1 |
| yeps_YPTB2507 | csy2 | | YPTB_RS13620 | YPTB2507 | | type I-F CRISPR-associated protein Csy2 | WP_011192625.1 |
| yeps_YPTB2508 | csy1 | | YPTB_RS13625 | YPTB2508 | | type I-F CRISPR-associated protein Csy1 | WP_011192626.1 |
| yeps_YPTB2513 | | | YPTB_RS13650 | YPTB2513 | | hypothetical protein | WP_002230953.1 |
| yeps_YPTB2514 | | | YPTB_RS13655 | YPTB2514 | | DUF1861 family protein | WP_011192631.1 |
| yeps_YPTB2516 | | | YPTB_RS13665 | YPTB2516 | | carbohydrate ABC transporter permease | WP_002210197.1 |
| yeps_YPTB2517 | | | YPTB_RS13670 | YPTB2517 | | sugar ABC transporter permease | WP_011192633.1 |
| yeps_YPTB2518 | | | YPTB_RS13675 | YPTB2518 | | ABC transporter substrate-binding protein | WP_011192634.1 |
| yeps_YPTB2519 | | | YPTB_RS13680 | YPTB2519 | | LacI family DNA-binding transcriptional regulator | WP_002210200.1 |
| yeps_YPTB2520 | | | YPTB_RS13685 | YPTB2520 | | phosphomannomutase/phosphoglucomutase | WP_002210201.1 |
| yeps_YPTB2535 | | | YPTB_RS13765 | YPTB2535 | | sugar ABC transporter permease | WP_002210221.1 |
| yeps_YPTB2536 | | | YPTB_RS13770 | YPTB2536 | | sugar ABC transporter ATP-binding protein | WP_011192642.1 |
| yeps_YPTB2537 | | | YPTB_RS13775 | YPTB2537 | | substrate-binding domain-containing protein | WP_011192643.1 |
| yeps_YPTB2541 | | | YPTB_RS13795 | YPTB2541 | | cation diffusion facilitator family transporter | WP_011192647.1 |
| yeps_YPTB2542 | ompX | omp4 | YPTB_RS13800 | YPTB2542 | | outer membrane protein OmpX | WP_002210229.1 |
| yeps_YPTB2543 | rhtA | | YPTB_RS13805 | YPTB2543 | | threonine/homoserine exporter RhtA | WP_002213809.1 |
| yeps_YPTB2544 | | | YPTB_RS13810 | YPTB2544 | | GNAT family N-acetyltransferase | WP_002210231.1 |
| yeps_YPTB2546 | dps | | YPTB_RS13820 | YPTB2546 | | DNA starvation/stationary phase protection protein Dps | WP_002210233.1 |
| yeps_YPTB2547 | | | YPTB_RS13825 | YPTB2547 | | YdcF family protein | WP_002210234.1 |
| yeps_YPTB2548 | glnH | | YPTB_RS13835 | YPTB2548 | | glutamine ABC transporter substrate-binding protein GlnH | WP_011192648.1 |
| yeps_YPTB2550 | glnQ | | YPTB_RS13845 | YPTB2550 | | glutamine ABC transporter ATP-binding protein GlnQ | WP_011192649.1 |
| yeps_YPTB2556 | menE | | YPTB_RS13870 | YPTB2556 | | o-succinylbenzoate--CoA ligase | WP_011192653.1 |
| yeps_YPTB2559 | menH | | YPTB_RS13885 | YPTB2559 | | 2-succinyl-6-hydroxy-2,4-cyclohexadiene-1-carboxylate synthase | WP_011192655.1 |
| yeps_YPTB2560 | menD | | YPTB_RS13890 | YPTB2560 | | 2-succinyl-5-enolpyruvyl-6-hydroxy-3-cyclohexene-1-carboxylic-acid synthase | WP_011192656.1 |
| yeps_YPTB2561 | menF | | YPTB_RS13895 | YPTB2561 | | isochorismate synthase MenF | WP_011192657.1 |
| yeps_YPTB2572 | | | YPTB_RS13960 | YPTB2572 | | gluconokinase | WP_002210265.1 |
| yeps_YPTB2575 | nuoN | | YPTB_RS13975 | YPTB2575 | | NADH-quinone oxidoreductase subunit NuoN | WP_002210268.1 |
| yeps_YPTB2576 | nuoM | | YPTB_RS13980 | YPTB2576 | | NADH-quinone oxidoreductase subunit M | WP_011192666.1 |
| yeps_YPTB2579 | nuoJ | | YPTB_RS13995 | YPTB2579 | | NADH-quinone oxidoreductase subunit J | WP_002210272.1 |
| yeps_YPTB2582 | nuoG | | YPTB_RS14010 | YPTB2582 | | NADH-quinone oxidoreductase subunit NuoG | WP_011192668.1 |
| yeps_YPTB2583 | nuoF | | YPTB_RS14015 | YPTB2583 | | NADH-quinone oxidoreductase subunit NuoF | WP_011192669.1 |
| yeps_YPTB2584 | nuoE | | YPTB_RS14020 | YPTB2584 | | NADH-quinone oxidoreductase subunit NuoE | WP_172601542.1 |
| yeps_YPTB2593 | | | YPTB_RS14070 | YPTB2593 | | sugar phosphatase | WP_011192675.1 |
| yeps_YPTB2596 | | | | YPTB2596 | | hypothetical protein | CAH21834.1 |
| yeps_YPTB2603 | yfcD | | YPTB_RS14120 | YPTB2603 | | NUDIX hydrolase YfcD | WP_002209742.1 |
| yeps_YPTB2605 | | | YPTB_RS14130 | YPTB2605 | | EAL domain-containing protein | WP_012303928.1 |
| yeps_YPTB2606 | | | YPTB_RS14135 | YPTB2606 | | TIGR01777 family oxidoreductase | WP_002209739.1 |
| yeps_YPTB2611 | | | YPTB_RS14160 | YPTB2611 | | UbiX family flavin prenyltransferase | WP_002209734.1 |
| yeps_YPTB2616 | accD | dedB,usg | YPTB_RS14185 | YPTB2616 | | acetyl-CoA carboxylase, carboxyltransferase subunit beta | WP_002209729.1 |
| yeps_YPTB2621 | | | YPTB_RS14210 | YPTB2621 | | helix-turn-helix transcriptional regulator | WP_011192684.1 |
| yeps_YPTB2628 | | | YPTB_RS14250 | YPTB2628 | | YfcL family protein | WP_002209715.1 |
| yeps_YPTB2631 | mepA | | YPTB_RS14265 | YPTB2631 | | penicillin-insensitive murein endopeptidase | WP_002225365.1 |
| yeps_YPTB2633 | prmB | | YPTB_RS14275 | YPTB2633 | | 50S ribosomal protein L3 N(5)-glutamine methyltransferase | WP_002209710.1 |
| yeps_YPTB2634 | smrB | | YPTB_RS14280 | YPTB2634 | | endonuclease SmrB | WP_002227846.1 |
| yeps_YPTB2635 | sixA | | YPTB_RS14285 | YPTB2635 | | phosphohistidine phosphatase SixA | WP_002209707.1 |

| | | | | | | | |
|---|---|---|---|---|---|---|---|
| yeps_YPTB2638 | | | YPTB_RS14300 | YPTB2638 | | YfcZ/YiiS family protein | WP_002227845.1 |
| yeps_YPTB2641 | ccmI | | YPTB_RS14315 | YPTB2641 | | c-type cytochrome biogenesis protein CcmI | WP_011192696.1 |
| yeps_YPTB2642 | | | YPTB_RS14320 | YPTB2642 | | cytochrome c-type biogenesis protein CcmH | WP_002214855.1 |
| yeps_YPTB2643 | | | YPTB_RS14325 | YPTB2643 | | DsbE family thiol:disulfide interchange protein | WP_002209699.1 |
| yeps_YPTB2644 | | | YPTB_RS14330 | YPTB2644 | | heme lyase CcmF/NrfE family subunit | WP_011192697.1 |
| yeps_YPTB2645 | ccmE | | YPTB_RS14335 | YPTB2645 | | cytochrome c maturation protein CcmE | WP_002209697.1 |
| yeps_YPTB2647 | | | YPTB_RS14345 | YPTB2647 | | heme ABC transporter permease | WP_011192699.1 |
| yeps_YPTB2648 | ccmB | | YPTB_RS14350 | YPTB2648 | | heme exporter protein CcmB | WP_002209694.1 |
| yeps_YPTB2649 | ccmA | | YPTB_RS14355 | YPTB2649 | | cytochrome c biogenesis heme-transporting ATPase CcmA | WP_011192700.1 |
| yeps_YPTB2650 | | | YPTB_RS14360 | YPTB2650 | | hypothetical protein | WP_011192701.1 |
| yeps_YPTB2651 | | | YPTB_RS14370 | YPTB2651 | | LemA family protein | WP_002209691.1 |
| yeps_YPTB2654 | | | YPTB_RS14385 | YPTB2654 | | hypothetical protein | WP_002209687.1 |
| yeps_YPTB2655 | | | YPTB_RS14390 | YPTB2655 | | type VI secretion system baseplate subunit TssE | WP_002209686.1 |
| yeps_YPTB2661 | tssL | | YPTB_RS14420 | YPTB2661 | | type VI secretion system protein TssL, long form | WP_011192706.1 |
| yeps_YPTB2662 | tssK | | YPTB_RS14425 | YPTB2662 | | type VI secretion system baseplate subunit TssK | WP_002211571.1 |
| yeps_YPTB2673 | tssG | | YPTB_RS14485 | YPTB2673 | | type VI secretion system baseplate subunit TssG | WP_011192712.1 |
| yeps_YPTB2674 | | | YPTB_RS14490 | YPTB2674 | | ImpA family type VI secretion system protein | WP_011192713.1 |
| yeps_YPTB2675 | | | YPTB_RS14495 | YPTB2675 | | fimbrial protein | WP_011192714.1 |
| yeps_YPTB2676 | | | YPTB_RS14500 | YPTB2676 | | glycosyl transferase | WP_011192715.1 |
| yeps_YPTB2678 | | | YPTB_RS14510 | YPTB2678 | | OmpA family protein | WP_011192717.1 |
| yeps_YPTB2679 | | | YPTB_RS14515 | YPTB2679 | | DcrB-related protein | WP_002215321.1 |
| yeps_YPTB2689 | dmsB | | YPTB_RS14565 | YPTB2689 | | dimethylsulfoxide reductase subunit B | WP_002211603.1 |
| yeps_YPTB2693 | | | YPTB_RS14585 | YPTB2693 | | DUF799 domain-containing protein | WP_002211607.1 |
| yeps_YPTB2697 | | | YPTB_RS14605 | YPTB2697 | | cytochrome b | WP_002211611.1 |
| yeps_YPTB2698 | alaC | | YPTB_RS14610 | YPTB2698 | | alanine transaminase | WP_002231034.1 |
| yeps_YPTB2700 | glk | | YPTB_RS14620 | YPTB2700 | | glucokinase | WP_002211615.1 |
| yeps_YPTB2701 | | | YPTB_RS14625 | YPTB2701 | | multidrug/biocide efflux PACE transporter | WP_011192730.1 |
| yeps_YPTB2703 | | | YPTB_RS14635 | YPTB2703 | | aldo/keto reductase | WP_011192731.1 |
| yeps_YPTB2704 | | | YPTB_RS14640 | YPTB2704 | | DUF2502 domain-containing protein | WP_002216909.1 |
| yeps_YPTB2711 | ligA | dnaL,lig,lop,pdeC | YPTB_RS14705 | YPTB2711 | | NAD-dependent DNA ligase LigA | WP_011192735.1 |
| yeps_YPTB2713 | cysZ | | YPTB_RS14715 | YPTB2713 | | sulfate transporter CysZ | WP_002231042.1 |
| yeps_YPTB2717 | crr | gsr,iex,tgs,treD | YPTB_RS14735 | YPTB2717 | | PTS glucose transporter subunit IIA | WP_002208491.1 |
| yeps_YPTB2718 | | | YPTB_RS14740 | YPTB2718 | | HAMP domain-containing protein | WP_011192737.1 |
| yeps_YPTB2719 | | | YPTB_RS14745 | YPTB2719 | | response regulator transcription factor | WP_002208494.1 |
| yeps_YPTB2721 | | | YPTB_RS14755 | YPTB2721 | | MacB family efflux pump subunit | WP_011192738.1 |
| yeps_YPTB2737 | | | YPTB_RS14835 | YPTB2737 | | MurR/RpiR family transcriptional regulator | WP_011192747.1 |
| yeps_YPTB2739 | | | YPTB_RS14850 | YPTB2739 | | N-acetylmannosamine kinase | WP_002208516.1 |
| yeps_YPTB2741 | | | YPTB_RS14860 | YPTB2741 | | N-acetylmannosamine-6-phosphate 2-epimerase | WP_071819112.1 |
| yeps_YPTB2742 | | | YPTB_RS14865 | YPTB2742 | | dihydrodipicolinate synthase family protein | WP_002208518.1 |
| yeps_YPTB2743 | | | YPTB_RS14870 | YPTB2743 | | Dyp-type peroxidase | WP_011192751.1 |
| yeps_YPTB2744 | | | YPTB_RS14875 | YPTB2744 | | RpoE-regulated lipoprotein | WP_002208521.1 |
| yeps_YPTB2755 | | | YPTB_RS14935 | YPTB2755 | | YaiI/YqxD family protein | WP_002208527.1 |
| yeps_YPTB2756 | maeB | | YPTB_RS14940 | YPTB2756 | | NADP-dependent oxaloacetate-decarboxylating malate dehydrogenase | WP_011192757.1 |
| yeps_YPTB2757 | nudK | | YPTB_RS14945 | YPTB2757 | | GDP-mannose pyrophosphatase NudK | WP_002208529.1 |
| yeps_YPTB2758 | napC | | YPTB_RS14950 | YPTB2758 | | cytochrome c-type protein NapC | WP_002208531.1 |
| yeps_YPTB2759 | napB | | YPTB_RS14955 | YPTB2759 | | nitrate reductase cytochrome c-type subunit | WP_002214527.1 |
| yeps_YPTB2764 | | | YPTB_RS14980 | YPTB2764 | | helix-turn-helix domain-containing protein | WP_002208538.1 |
| yeps_YPTB2765 | acrD | | YPTB_RS14985 | YPTB2765 | | multidrug efflux RND transporter permease AcrD | WP_011192758.1 |
| yeps_YPTB2766 | | | YPTB_RS14990 | YPTB2766 | | hypothetical protein | WP_002208540.1 |
| yeps_YPTB2773 | | | YPTB_RS15025 | YPTB2773 | | tetratricopeptide repeat protein | WP_002208547.1 |
| yeps_YPTB2774 | | | YPTB_RS15030 | YPTB2774 | | ArsC family reductase | WP_002208548.1 |
| yeps_YPTB2776 | | | YPTB_RS15040 | YPTB2776 | | M15 family metallopeptidase | WP_002208550.1 |
| yeps_YPTB2777 | | | YPTB_RS15045 | YPTB2777 | | YpfN family protein | WP_002208551.1 |
| yeps_YPTB2782 | bamC | | YPTB_RS15070 | YPTB2782 | | outer membrane protein assembly factor BamC | WP_011192764.1 |
| yeps_YPTB2783 | dapA | | YPTB_RS15075 | YPTB2783 | | 4-hydroxy-tetrahydrodipicolinate synthase | WP_032466692.1 |
| yeps_YPTB2784 | | | YPTB_RS15080 | YPTB2784 | | glycine cleavage system transcriptional repressor | WP_002227068.1 |
| yeps_YPTB2786 | | | | YPTB2786 | | hypothetical protein | CAH22024.1 |
| yeps_YPTB2791 | arsC | | YPTB_RS15115 | YPTB2791 | | arsenate reductase (glutaredoxin) | WP_002208565.1 |
| yeps_YPTB2792 | hda | | YPTB_RS15120 | YPTB2792 | | DnaA inactivator Hda | WP_002228401.1 |
| yeps_YPTB2793 | uraA | | YPTB_RS15125 | YPTB2793 | | uracil permease | WP_011192769.1 |
| yeps_YPTB2795 | purM | | YPTB_RS15135 | YPTB2795 | | phosphoribosylformylglycinamidine cyclo-ligase | WP_011192770.1 |
| yeps_YPTB2796 | purN | | YPTB_RS15140 | YPTB2796 | | phosphoribosylglycinamide formyltransferase | WP_011192771.1 |
| yeps_YPTB2800 | pstA | phoT | YPTB_RS15160 | YPTB2800 | | phosphate ABC transporter permease PstA | WP_011192773.1 |
| yeps_YPTB2801 | | | YPTB_RS15165 | YPTB2801 | | ABC transporter permease subunit | WP_011192774.1 |
| yeps_YPTB2802 | ppk1 | | YPTB_RS15170 | YPTB2802 | | polyphosphate kinase 1 | WP_002209782.1 |
| yeps_YPTB2803 | ppx | | YPTB_RS15175 | YPTB2803 | | exopolyphosphatase | WP_002209783.1 |
| yeps_YPTB2804 | | | YPTB_RS15180 | YPTB2804 | | DUF2633 family protein | WP_012303868.1 |
| yeps_YPTB2805 | mgtE | | YPTB_RS15185 | YPTB2805 | | magnesium transporter | WP_002209784.1 |
| yeps_YPTB2808 | | | YPTB_RS15200 | YPTB2808 | | ABC transporter permease | WP_011192776.1 |
| yeps_YPTB2812 | | | YPTB_RS15220 | YPTB2812 | | ABC transporter ATP-binding protein | WP_011192780.1 |
| yeps_YPTB2814 | | | YPTB_RS15230 | YPTB2814 | | MdtB/MuxB family multidrug efflux RND transporter permease subunit | WP_011192782.1 |
| yeps_YPTB2815 | mdtC | | YPTB_RS15235 | YPTB2815 | | multidrug efflux RND transporter permease subunit MdtC | WP_011192783.1 |
| yeps_YPTB2816 | | | YPTB_RS15240 | YPTB2816 | | MFS transporter | WP_011192784.1 |
| yeps_YPTB2818 | baeR | | YPTB_RS15250 | YPTB2818 | | two-component system response regulator BaeR | WP_011192786.1 |
| yeps_YPTB2821 | yegS | | YPTB_RS15265 | YPTB2821 | | lipid kinase YegS | WP_011192787.1 |
| yeps_YPTB2822 | | | YPTB_RS15270 | YPTB2822 | | inhibitor of vertebrate lysozyme family protein | WP_002209800.1 |
| yeps_YPTB2824 | thiD | | YPTB_RS15280 | YPTB2824 | | bifunctional hydroxymethylpyrimidine kinase/phosphomethylpyrimidine kinase | WP_011192789.1 |
| yeps_YPTB2833 | guaB | guaR | YPTB_RS15335 | YPTB2833 | | IMP dehydrogenase | WP_002227862.1 |
| yeps_YPTB2835 | | | YPTB_RS15345 | YPTB2835 | | zinc ribbon domain-containing protein | WP_011192798.1 |
| yeps_YPTB2839 | | | YPTB_RS15365 | YPTB2839 | | YfgM family protein | WP_002209815.1 |
| yeps_YPTB2840 | hisS | | YPTB_RS15370 | YPTB2840 | | histidine--tRNA ligase | WP_002209816.1 |
| yeps_YPTB2841 | ispG | | YPTB_RS15375 | YPTB2841 | | flavodoxin-dependent (E)-4-hydroxy-3-methylbut-2-enyl-diphosphate synthase | WP_002209817.1 |
| yeps_YPTB2845 | ndk | | YPTB_RS15395 | YPTB2845 | | nucleoside-diphosphate kinase | WP_011192804.1 |
| yeps_YPTB2852 | pepB | | YPTB_RS15430 | YPTB2852 | | aminopeptidase PepB | WP_011192810.1 |
| yeps_YPTB2856 | hscB | | YPTB_RS15450 | YPTB2856 | | co-chaperone HscB | WP_011192812.1 |
| yeps_YPTB2858 | iscU | | YPTB_RS15460 | YPTB2858 | | Fe-S cluster assembly scaffold IscU | WP_011192813.1 |
| yeps_YPTB2860 | iscR | | YPTB_RS15470 | YPTB2860 | | Fe-S cluster assembly transcriptional regulator IscR | WP_002222202.1 |
| yeps_YPTB2862 | suhB | | YPTB_RS15480 | YPTB2862 | | inositol-1-monophosphatase | WP_002213305.1 |
| yeps_YPTB2863 | | | YPTB_RS15485 | YPTB2863 | | nickel/cobalt transporter | WP_002213308.1 |
| yeps_YPTB2864 | | | YPTB_RS15490 | YPTB2864 | | DUF1007 family protein | WP_011192815.1 |
| yeps_YPTB2870 | hmpA | | YPTB_RS15520 | YPTB2870 | | NO-inducible flavohemoprotein | WP_002211553.1 |
| yeps_YPTB2874 | glrR | | YPTB_RS15540 | YPTB2874 | | two-component system response regulator GlrR | WP_002211556.1 |
| yeps_YPTB2880 | mltF | | YPTB_RS15570 | YPTB2880 | | membrane-bound lytic murein transglycosylase MltF | WP_024062994.1 |
| yeps_YPTB2882 | yfhb | | YPTB_RS15580 | YPTB2882 | | phosphatidylglycerophosphatase C | WP_002211564.1 |
| yeps_YPTB2885 | fdx | | | YPTB2885 | | putative [4Fe-4S] ferredoxin | CAH22123.1 |
| yeps_YPTB2888 | recO | | YPTB_RS15610 | YPTB2888 | | DNA repair protein RecO | WP_002209680.1 |
| yeps_YPTB2891 | lepB | | YPTB_RS15625 | YPTB2891 | | signal peptidase I | WP_011192827.1 |
| yeps_YPTB2892 | lepA | | YPTB_RS15630 | YPTB2892 | | translation elongation factor 4 | WP_002209677.1 |
| yeps_YPTB2896 | rseA | mclA | YPTB_RS15650 | YPTB2896 | | anti-sigma-E factor RseA | WP_002209673.1 |
| yeps_YPTB2901 | | | YPTB_RS15675 | YPTB2901 | | ankyrin repeat domain-containing protein | WP_002209667.1 |
| yeps_YPTB2903 | grcA | | YPTB_RS15690 | YPTB2903 | | autonomous glycyl radical cofactor GrcA | WP_002209664.1 |
| yeps_YPTB2904 | ung | | YPTB_RS15695 | YPTB2904 | | uracil-DNA glycosylase | WP_002209663.1 |
| yeps_YPTB2906 | | | YPTB_RS15705 | YPTB2906 | | DUF979 domain-containing protein | WP_011192829.1 |
| yeps_YPTB2908 | pxpA | | YPTB_RS15715 | YPTB2908 | | 5-oxoprolinase subunit PxpA | WP_002209659.1 |
| yeps_YPTB2910 | pxpB | | YPTB_RS15725 | YPTB2910 | | 5-oxoprolinase subunit PxpB | WP_011192830.1 |
| yeps_YPTB2911 | | | YPTB_RS15730 | YPTB2911 | | type 2 GTP cyclohydrolase I | WP_025471060.1 |
| yeps_YPTB2912 | | | YPTB_RS15735 | YPTB2912 | | 3',5'-cyclic-nucleotide phosphodiesterase | WP_002209656.1 |
| yeps_YPTB2913 | phrB | phr | YPTB_RS15740 | YPTB2913 | | deoxyribodipyrimidine photo-lyase | WP_011192831.1 |
| yeps_YPTB2917 | kdpB | | YPTB_RS15770 | YPTB2917 | | potassium-transporting ATPase subunit KdpB | WP_011192833.1 |
| yeps_YPTB2918 | kdpC | | YPTB_RS15775 | YPTB2918 | | potassium-transporting ATPase subunit KdpC | WP_072083175.1 |
| yeps_YPTB2922 | | | YPTB_RS15795 | YPTB2922 | | hypothetical protein | WP_002209646.1 |
| yeps_YPTB2923 | pgm | | YPTB_RS15805 | YPTB2923 | | phosphoglucomutase (alpha-D-glucose-1,6-bisphosphate-dependent) | WP_041175466.1 |
| yeps_YPTB2924 | seqA | | YPTB_RS15810 | YPTB2924 | | replication initiation negative regulator SeqA | WP_011192836.1 |
| yeps_YPTB2925 | | | YPTB_RS15820 | YPTB2925 | | hypothetical protein | WP_002212246.1 |
| yeps_YPTB2926 | | | | YPTB2926 | | hypothetical protein | CAH22164.1 |
| yeps_YPTB2927 | chbG | | YPTB_RS15830 | YPTB2927 | | chitin disaccharide deacetylase | WP_002212244.1 |
| yeps_YPTB2928 | chbR | | YPTB_RS15835 | YPTB2928 | | transcriptional regulator ChbR | WP_002212243.1 |
| yeps_YPTB2936 | | | YPTB_RS15885 | YPTB2936 | | HoxN/HupN/NixA family nickel/cobalt transporter | WP_011192840.1 |
| yeps_YPTB2937 | yut | | YPTB_RS15890 | YPTB2937 | | urea transporter | WP_011192841.1 |
| yeps_YPTB2938 | | | YPTB_RS15895 | YPTB2938 | | urease accessory protein UreD | WP_011192842.1 |
| yeps_YPTB2943 | | | YPTB_RS15920 | YPTB2943 | | urease subunit beta | WP_002222230.1 |
| yeps_YPTB2964 | dnaQ | mutD | YPTB_RS16030 | YPTB2964 | | DNA polymerase III subunit epsilon | WP_002210700.1 |
| yeps_YPTB2969 | | | YPTB_RS16055 | YPTB2969 | | endonuclease/exonuclease/phosphatase family protein | WP_002220059.1 |
| yeps_YPTB2976 | rcsF | | YPTB_RS16120 | YPTB2976 | | Rcs stress response system protein RcsF | WP_002212158.1 |
| yeps_YPTB2979 | nlpE | | YPTB_RS16135 | YPTB2979 | | envelope stress response activation lipoprotein NlpE | WP_002212155.1 |
| yeps_YPTB2980 | arfB | | YPTB_RS16140 | YPTB2980 | | aminoacyl-tRNA hydrolase | WP_002212154.1 |
| yeps_YPTB2981 | | | YPTB_RS16145 | YPTB2981 | | YaeQ family protein | WP_002212153.1 |
| yeps_YPTB2983 | rof | | YPTB_RS16155 | YPTB2983 | | Rho-binding antiterminator | WP_002218374.1 |
| yeps_YPTB2984 | | | YPTB_RS16160 | YPTB2984 | | cytochrome c | WP_002212150.1 |
| yeps_YPTB2985 | tilS | | YPTB_RS16165 | YPTB2985 | | tRNA lysidine(34) synthetase TilS | WP_011192857.1 |

| | | | | | | | |
|---|---|---|---|---|---|---|---|
| yeps_YPTB2986 | | | YPTB_RS16170 | YPTB2986 | | VOC family protein | WP_002212148.1 |
| yeps_YPTB2988 | dnaE | polC | YPTB_RS16180 | YPTB2988 | | DNA polymerase III subunit alpha | WP_012104742.1 |
| yeps_YPTB2989 | rnhB | | YPTB_RS16185 | YPTB2989 | | ribonuclease HII | WP_002212145.1 |
| yeps_YPTB2991 | lpxA | | YPTB_RS16195 | YPTB2991 | | acyl-ACP--UDP-N-acetylglucosamine O-acyltransferase | WP_002212143.1 |
| yeps_YPTB2995 | bamA | | YPTB_RS16215 | YPTB2995 | | outer membrane protein assembly factor BamA | WP_002212139.1 |
| yeps_YPTB2996 | rseP | | YPTB_RS16220 | YPTB2996 | | sigma E protease regulator RseP | WP_002212138.1 |
| yeps_YPTB2999 | ispC | | YPTB_RS16235 | YPTB2999 | | 1-deoxy-D-xylulose-5-phosphate reductoisomerase | WP_011192861.1 |
| yeps_YPTB3000 | frr | rrf | YPTB_RS16240 | YPTB3000 | | ribosome recycling factor | WP_002212134.1 |
| yeps_YPTB3001 | pyrH | smbA | YPTB_RS16245 | YPTB3001 | | UMP kinase | WP_002212133.1 |
| yeps_YPTB3002 | tsf | | YPTB_RS16250 | YPTB3002 | | translation elongation factor Ts | WP_002212132.1 |
| yeps_YPTB3007 | | | YPTB_RS16275 | YPTB3007 | | DUF3461 family protein | WP_002212127.1 |
| yeps_YPTB3008 | | | YPTB_RS16280 | YPTB3008 | | flavodoxin | WP_011192865.1 |
| yeps_YPTB3009 | truC | | YPTB_RS16285 | YPTB3009 | | tRNA pseudouridine(65) synthase TruC | WP_002212125.1 |
| yeps_YPTB3016 | | | YPTB_RS16325 | YPTB3016 | | DUF423 domain-containing protein | WP_002212118.1 |
| yeps_YPTB3019 | csdE | | YPTB_RS16340 | YPTB3019 | | cysteine desulfurase sulfur acceptor subunit CsdE | WP_002212115.1 |
| yeps_YPTB3027 | recC | | YPTB_RS16395 | YPTB3027 | | exodeoxyribonuclease V subunit gamma | WP_011192871.1 |
| yeps_YPTB3028 | | | YPTB_RS16400 | YPTB3028 | | peptidase | WP_012104731.1 |
| yeps_YPTB3029 | | | YPTB_RS16405 | YPTB3029 | | YgdB family protein | WP_002211629.1 |
| yeps_YPTB3030 | | | YPTB_RS16410 | YPTB3030 | | prepilin peptidase-dependent protein | WP_002211630.1 |
| yeps_YPTB3031 | | | YPTB_RS16415 | YPTB3031 | | prepilin peptidase-dependent protein | WP_002228638.1 |
| yeps_YPTB3044 | galR | | YPTB_RS16490 | YPTB3044 | | HTH-type transcriptional regulator GalR | WP_002209845.1 |
| yeps_YPTB3045 | lysA | | YPTB_RS16495 | YPTB3045 | | diaminopimelate decarboxylase | WP_011192878.1 |
| yeps_YPTB3046 | | | YPTB_RS16500 | YPTB3046 | | LysR family transcriptional regulator | WP_011192879.1 |
| yeps_YPTB3048 | | | YPTB_RS16510 | YPTB3048 | | LysR family transcriptional regulator | WP_011192880.1 |
| yeps_YPTB3049 | | | YPTB_RS16515 | YPTB3049 | | MBL fold metallo-hydrolase | WP_002209850.1 |
| yeps_YPTB3076 | kbaZ | | YPTB_RS16650 | YPTB3076 | | tagatose-bisphosphate aldolase subunit KbaZ | WP_011192897.1 |
| yeps_YPTB3077 | | | YPTB_RS16655 | YPTB3077 | | SIS domain-containing protein | WP_011192898.1 |
| yeps_YPTB3078 | | | YPTB_RS16660 | YPTB3078 | | PTS system mannose/fructose/N-acetylgalactosamine-transporter subunit IIB | WP_002209878.1 |
| yeps_YPTB3079 | | | YPTB_RS16665 | YPTB3079 | | PTS mannose/fructose/sorbose/N-acetylgalactosamine transporter subunit IIC | WP_002209879.1 |
| yeps_YPTB3080 | | | YPTB_RS16670 | YPTB3080 | | PTS system mannose/fructose/sorbose family transporter subunit IID | WP_002209880.1 |
| yeps_YPTB3081 | | | YPTB_RS16675 | YPTB3081 | | PTS sugar transporter subunit IIA | WP_002209881.1 |
| yeps_YPTB3082 | nagA | | YPTB_RS16680 | YPTB3082 | | N-acetylglucosamine-6-phosphate deacetylase | WP_002209882.1 |
| yeps_YPTB3083 | kduD | | YPTB_RS16685 | YPTB3083 | | 2-dehydro-3-deoxy-D-gluconate 5-dehydrogenase KduD | WP_002230627.1 |
| yeps_YPTB3087 | | | YPTB_RS16705 | YPTB3087 | | DUF2264 domain-containing protein | WP_011192901.1 |
| yeps_YPTB3088 | | | YPTB_RS16710 | YPTB3088 | | tagatose bisphosphate family class II aldolase | WP_011192902.1 |
| yeps_YPTB3089 | | | YPTB_RS16715 | YPTB3089 | | DJ-1/PfpI family protein | WP_011192903.1 |
| yeps_YPTB3097 | | | YPTB_RS16755 | YPTB3097 | | beta-galactosidase | WP_002209897.1 |
| yeps_YPTB3098 | | | YPTB_RS16760 | YPTB3098 | | arabinogalactan endo-beta-1,4-galactanase | WP_002209898.1 |
| yeps_YPTB3099 | | | YPTB_RS16765 | YPTB3099 | | sugar ABC transporter permease | WP_002209899.1 |
| yeps_YPTB3100 | | | YPTB_RS16770 | YPTB3100 | | sugar ABC transporter permease | WP_002209900.1 |
| yeps_YPTB3101 | | | YPTB_RS16775 | YPTB3101 | | extracellular solute-binding protein | WP_002209901.1 |
| yeps_YPTB3103 | | | YPTB_RS16785 | YPTB3103 | | sugar ABC transporter ATP-binding protein | WP_011192908.1 |
| yeps_YPTB3117 | | | YPTB_RS16865 | YPTB3117 | | Fic family protein | WP_002209918.1 |
| yeps_YPTB3118 | | | YPTB_RS16870 | YPTB3118 | | hypothetical protein | WP_011192915.1 |
| yeps_YPTB3166 | dsbC | | YPTB_RS17155 | YPTB3166 | | bifunctional protein-disulfide isomerase/oxidoreductase DsbC | WP_002209932.1 |
| yeps_YPTB3168 | fldB | | YPTB_RS17165 | YPTB3168 | | flavodoxin FldB | WP_011192956.1 |
| yeps_YPTB3171 | creB | blrA | YPTB_RS17180 | YPTB3171 | | two-component system response regulator CreB | WP_002209937.1 |
| yeps_YPTB3172 | | | YPTB_RS17185 | YPTB3172 | | protein YgfX | WP_002215144.1 |
| yeps_YPTB3173 | sdhE | | YPTB_RS17190 | YPTB3173 | | FAD assembly factor SdhE | WP_002209939.1 |
| yeps_YPTB3174 | ygfZ | | YPTB_RS17195 | YPTB3174 | | tRNA-modifying protein YgfZ | WP_011192958.1 |
| yeps_YPTB3175 | | | YPTB_RS17200 | YPTB3175 | | DUF2165 family protein | WP_002209941.1 |
| yeps_YPTB3176 | | | YPTB_RS17205 | YPTB3176 | | hemolysin III family protein | WP_011192959.1 |
| yeps_YPTB3184 | ubiH | | YPTB_RS17250 | YPTB3184 | | 2-octaprenyl-6-methoxyphenyl hydroxylase | WP_011192963.1 |
| yeps_YPTB3185 | pepP | | YPTB_RS17255 | YPTB3185 | | Xaa-Pro aminopeptidase | WP_011192964.1 |
| yeps_YPTB3186 | | | YPTB_RS17260 | YPTB3186 | | YecA family protein | WP_002209953.1 |
| yeps_YPTB3187 | zapA | | YPTB_RS17265 | YPTB3187 | | cell division protein ZapA | WP_002209954.1 |
| yeps_YPTB3190 | rpiA | | YPTB_RS17280 | YPTB3190 | | ribose-5-phosphate isomerase RpiA | WP_002209957.1 |
| yeps_YPTB3191 | | | YPTB_RS17285 | YPTB3191 | | LysR family transcriptional regulator ArgP | WP_002209958.1 |
| yeps_YPTB3192 | | | YPTB_RS17290 | YPTB3192 | | oxidative stress defense protein | WP_011192966.1 |
| yeps_YPTB3203 | metK | | YPTB_RS17350 | YPTB3203 | | methionine adenosyltransferase | WP_002209971.1 |
| yeps_YPTB3206 | rsmE | | YPTB_RS17370 | YPTB3206 | | 16S rRNA (uracil(1498)-N(3))-methyltransferase | WP_002209975.1 |
| yeps_YPTB3207 | gshB | gsh-II | YPTB_RS17375 | YPTB3207 | | glutathione synthase | WP_002209976.1 |
| yeps_YPTB3208 | | | YPTB_RS17380 | YPTB3208 | | YqgE/AlgH family protein | WP_011192971.1 |
| yeps_YPTB3209 | ruvX | | YPTB_RS17385 | YPTB3209 | | Holliday junction resolvase RuvX | WP_011192972.1 |
| yeps_YPTB3211 | aguA | | YPTB_RS17395 | YPTB3211 | | agmatine deiminase | WP_002209980.1 |
| yeps_YPTB3212 | | | YPTB_RS17400 | YPTB3212 | | type IV pilus twitching motility protein PilT | WP_002209981.1 |
| yeps_YPTB3213 | | | YPTB_RS17405 | YPTB3213 | | YggS family pyridoxal phosphate-dependent enzyme | WP_011192973.1 |
| yeps_YPTB3214 | | | YPTB_RS17410 | YPTB3214 | | pyrroline-5-carboxylate reductase | WP_011192974.1 |
| yeps_YPTB3216 | | | YPTB_RS17420 | YPTB3216 | | DUF167 family protein YggU | WP_011192976.1 |
| yeps_YPTB3221 | glsB | | YPTB_RS17445 | YPTB3221 | | glutaminase B | WP_011192980.1 |
| yeps_YPTB3223 | trmB | | YPTB_RS17455 | YPTB3223 | | tRNA (guanosine(46)-N7)-methyltransferase TrmB | WP_011192981.1 |
| yeps_YPTB3231 | | | YPTB_RS17500 | YPTB3231 | | substrate-binding domain-containing protein | WP_011192985.1 |
| yeps_YPTB3258 | | | YPTB_RS17640 | YPTB3258 | | acyl-homoserine-lactone synthase | WP_002215903.1 |
| yeps_YPTB3261 | | | YPTB_RS17655 | YPTB3261 | | intradiol ring-cleavage dioxygenase | WP_011192997.1 |
| yeps_YPTB3262 | | | YPTB_RS17660 | YPTB3262 | | MFS transporter | WP_011192998.1 |
| yeps_YPTB3263 | | | YPTB_RS17665 | YPTB3263 | | aerobactin synthase IucA | WP_011192999.1 |
| yeps_YPTB3264 | | | YPTB_RS17670 | YPTB3264 | | acetyltransferase | WP_011193000.1 |
| yeps_YPTB3265 | iucC | | YPTB_RS17675 | YPTB3265 | | IucA/IucC family siderophore biosynthesis protein | WP_011193001.1 |
| yeps_YPTB3266 | | | YPTB_RS17680 | YPTB3266 | | lysine N(6)-hydroxylase/L-ornithine N(5)-oxygenase family protein | WP_011193002.1 |
| yeps_YPTB3267 | | | YPTB_RS17685 | YPTB3267 | | TonB-dependent siderophore receptor | WP_011193003.1 |
| yeps_YPTB3269 | | | YPTB_RS17695 | YPTB3269 | | cellulase family glycosylhydrolase | WP_012303626.1 |
| yeps_YPTB3270 | | | YPTB_RS17700 | YPTB3270 | | LacI family transcriptional regulator | WP_032465892.1 |
| yeps_YPTB3290 | | | YPTB_RS17800 | YPTB3290 | | ABC transporter ATP-binding protein/permease | WP_011193020.1 |
| yeps_YPTB3291 | | | YPTB_RS17805 | YPTB3291 | | ABC transporter ATP-binding protein/permease | WP_011193021.1 |
| yeps_YPTB3294 | | | YPTB_RS17820 | YPTB3294 | | Gfo/Idh/MocA family oxidoreductase | WP_011193024.1 |
| yeps_YPTB3295 | | | YPTB_RS17825 | YPTB3295 | | NAD-dependent epimerase/dehydratase family protein | WP_011193025.1 |
| yeps_YPTB3355 | | | YPTB_RS18135 | YPTB3355 | | flagellar biosynthetic protein FliR | WP_002213766.1 |
| yeps_YPTB3382 | exbD | | YPTB_RS18275 | YPTB3382 | | TonB system transport protein ExbD | WP_012414002.1 |
| yeps_YPTB3384 | metC | | YPTB_RS18285 | YPTB3384 | | cystathionine beta-lyase | WP_002212172.1 |
| yeps_YPTB3385 | | | YPTB_RS18290 | YPTB3385 | | DedA family protein | WP_002212173.1 |
| yeps_YPTB3386 | | | YPTB_RS18295 | YPTB3386 | | AraC family transcriptional regulator | WP_002213786.1 |
| yeps_YPTB3387 | yqhD | | YPTB_RS18300 | YPTB3387 | | alcohol dehydrogenase | WP_002212174.1 |
| yeps_YPTB3388 | dkgA | | YPTB_RS18310 | YPTB3388 | | 2,5-didehydrogluconate reductase DkgA | WP_011193092.1 |
| yeps_YPTB3390 | ftsP | | YPTB_RS18320 | YPTB3390 | | cell division protein FtsP | WP_002212176.1 |
| yeps_YPTB3391 | | | YPTB_RS18325 | YPTB3391 | | 1-acylglycerol-3-phosphate O-acyltransferase | WP_002212177.1 |
| yeps_YPTB3392 | parC | | YPTB_RS18330 | YPTB3392 | | DNA topoisomerase IV subunit A | WP_002212178.1 |
| yeps_YPTB3393 | | | YPTB_RS18340 | YPTB3393 | | NAD(P)H-dependent oxidoreductase | WP_002212179.1 |
| yeps_YPTB3394 | | | YPTB_RS18345 | YPTB3394 | | LysR family transcriptional regulator | WP_011193094.1 |
| yeps_YPTB3399 | nudF | | YPTB_RS18370 | YPTB3399 | | ADP-ribose diphosphatase | WP_002212184.1 |
| yeps_YPTB3402 | | | YPTB_RS18390 | YPTB3402 | | glutathionylspermidine synthase family protein | WP_002212188.1 |
| yeps_YPTB3403 | ygiD | | YPTB_RS18395 | YPTB3403 | | 4,5-DOPA dioxygenase extradiol | WP_002212189.1 |
| yeps_YPTB3404 | ribB | htrP | YPTB_RS18400 | YPTB3404 | | 3,4-dihydroxy-2-butanone-4-phosphate synthase | WP_002212190.1 |
| yeps_YPTB3408 | glnE | | YPTB_RS18425 | YPTB3408 | | bifunctional [glutamate--ammonia ligase]-adenylyl-L-tyrosine phosphorylase/[glutamate--ammonia-ligase] adenylyltransferase | WP_002212194.1 |
| yeps_YPTB3409 | | | YPTB_RS18430 | YPTB3409 | | inorganic triphosphatase | WP_002212195.1 |
| yeps_YPTB3410 | | | YPTB_RS18435 | YPTB3410 | | SH3 domain-containing protein | WP_011193100.1 |
| yeps_YPTB3411 | | | YPTB_RS18440 | YPTB3411 | | multifunctional CCA addition/repair protein | WP_011193101.1 |
| yeps_YPTB3412 | bacA | | YPTB_RS18445 | YPTB3412 | | undecaprenyl-diphosphate phosphatase | WP_002212198.1 |
| yeps_YPTB3414 | plsY | | YPTB_RS18455 | YPTB3414 | | glycerol-3-phosphate 1-O-acyltransferase PlsY | WP_002217581.1 |
| yeps_YPTB3415 | tsaD | | YPTB_RS18460 | YPTB3415 | | tRNA (adenosine(37)-N6)-threonylcarbamoyltransferase complex transferase subunit TsaD | WP_002212201.1 |
| yeps_YPTB3417 | dnaG | dnaP,parB | YPTB_RS18470 | YPTB3417 | | DNA primase | WP_002210358.1 |
| yeps_YPTB3434 | | | YPTB_RS18585 | YPTB3434 | | hypothetical protein | WP_002210374.1 |
| yeps_YPTB3439 | | | YPTB_RS18610 | YPTB3439 | | glycoside hydrolase family 3 C-terminal domain-containing protein | WP_011193110.1 |
| yeps_YPTB3440 | | | YPTB_RS18615 | YPTB3440 | | hypothetical protein | WP_011193111.1 |
| yeps_YPTB3441 | | | YPTB_RS18620 | YPTB3441 | | carbohydrate ABC transporter permease | WP_011193112.1 |
| yeps_YPTB3442 | | | YPTB_RS18625 | YPTB3442 | | sugar ABC transporter permease | WP_011193113.1 |
| yeps_YPTB3443 | | | YPTB_RS18630 | YPTB3443 | | extracellular solute-binding protein | WP_011193114.1 |
| yeps_YPTB3444 | | | YPTB_RS18635 | YPTB3444 | | LacI family transcriptional regulator | WP_011193115.1 |
| yeps_YPTB3445 | | | YPTB_RS18640 | YPTB3445 | | hypothetical protein | WP_011193116.1 |
| yeps_YPTB3446 | | | YPTB_RS18645 | YPTB3446 | | ABC transporter ATP-binding protein | WP_002353714.1 |
| yeps_YPTB3447 | | | YPTB_RS18650 | YPTB3447 | | membrane protein | WP_011193117.1 |
| yeps_YPTB3448 | | | YPTB_RS18655 | YPTB3448 | | transporter | WP_011193118.1 |
| yeps_YPTB3468 | | | YPTB_RS18775 | YPTB3468 | | HdeD family acid-resistance protein | WP_002210400.1 |
| yeps_YPTB3469 | | | YPTB_RS18780 | YPTB3469 | | NADPH-dependent 2,4-dienoyl-CoA reductase | WP_011193129.1 |
| yeps_YPTB3472 | | | YPTB_RS18795 | YPTB3472 | | M48 family metallopeptidase | WP_002210404.1 |
| yeps_YPTB3473 | | | YPTB_RS18800 | YPTB3473 | | Gfo/Idh/MocA family oxidoreductase | WP_002210405.1 |
| yeps_YPTB3475 | | | YPTB_RS18810 | YPTB3475 | | YgjV family protein | WP_002210407.1 |
| yeps_YPTB3477 | | | YPTB_RS18820 | YPTB3477 | | tagaturonate reductase | WP_011193132.1 |
| yeps_YPTB3480 | exuR | | YPTB_RS18840 | YPTB3480 | | transcriptional regulator ExuR | WP_002210412.1 |
| yeps_YPTB3481 | | | YPTB_RS18845 | YPTB3481 | | hypothetical protein | WP_012303498.1 |
| yeps_YPTB3482 | | | YPTB_RS18850 | YPTB3482 | | DedA family protein | WP_002210414.1 |

| | | | | | | | |
|---|---|---|---|---|---|---|---|
| yeps_YPTB3483 | mzrA | | YPTB_RS18855 | YPTB3483 | | EnvZ/OmpR regulon moderator MzrA | WP_002210415.1 |
| yeps_YPTB3484 | | | YPTB_RS18860 | YPTB3484 | | DUF1090 domain-containing protein | WP_002210416.1 |
| yeps_YPTB3486 | | | YPTB_RS18875 | YPTB3486 | | phage holin family protein | WP_002210419.1 |
| yeps_YPTB3487 | | | YPTB_RS18880 | YPTB3487 | | YqjK-like family protein | WP_011193134.1 |
| yeps_YPTB3489 | | | YPTB_RS18890 | YPTB3489 | | glutathione S-transferase family protein | WP_011193135.1 |
| yeps_YPTB3490 | | | YPTB_RS18895 | YPTB3490 | | LysR family transcriptional regulator | WP_011193136.1 |
| yeps_YPTB3494 | | | YPTB_RS18915 | YPTB3494 | | YraN family protein | WP_002210147.1 |
| yeps_YPTB3495 | diaA | | YPTB_RS18920 | YPTB3495 | | DnaA initiator-associating protein DiaA | WP_011193139.1 |
| yeps_YPTB3496 | yraP | | YPTB_RS18925 | YPTB3496 | | divisome-associated lipoprotein YraP | WP_011193140.1 |
| yeps_YPTB3497 | mtgA | | YPTB_RS18930 | YPTB3497 | | monofunctional biosynthetic peptidoglycan transglycosylase | WP_011193141.1 |
| yeps_YPTB3499 | | | | YPTB3499 | | hypothetical protein | CAH22737.1 |
| yeps_YPTB3500 | arcB | | YPTB_RS18945 | YPTB3500 | | aerobic respiration two-component sensor histidine kinase ArcB | WP_002210142.1 |
| yeps_YPTB3501 | | | YPTB_RS18955 | YPTB3501 | | TIGR01212 family radical SAM protein | WP_002210140.1 |
| yeps_YPTB3505 | sspB | | YPTB_RS18975 | YPTB3505 | | ClpXP protease specificity-enhancing factor | WP_002216203.1 |
| yeps_YPTB3509 | | | YPTB_RS19000 | YPTB3509 | | cell division protein ZapE | WP_002210131.1 |
| yeps_YPTB3511 | degQ | hhoA | YPTB_RS19010 | YPTB3511 | | serine endoprotease DegQ | WP_002218066.1 |
| yeps_YPTB3512 | degS | hhoB,htrH | YPTB_RS19015 | YPTB3512 | | outer membrane-stress sensor serine endopeptidase DegS | WP_002210128.1 |
| yeps_YPTB3513 | murA | murZ | YPTB_RS19020 | YPTB3513 | | UDP-N-acetylglucosamine 1-carboxyvinyltransferase | WP_002210127.1 |
| yeps_YPTB3514 | ibaG | | YPTB_RS19025 | YPTB3514 | | BolA family iron metabolism protein IbaG | WP_012104414.1 |
| yeps_YPTB3516 | mlaC | | YPTB_RS19035 | YPTB3516 | | phospholipid-binding protein MlaC | WP_002210124.1 |
| yeps_YPTB3517 | mlaD | | YPTB_RS19040 | YPTB3517 | | outer membrane lipid asymmetry maintenance protein MlaD | WP_011193146.1 |
| yeps_YPTB3519 | mlaF | | YPTB_RS19050 | YPTB3519 | | phospholipid ABC transporter ATP-binding protein MlaF | WP_002210121.1 |
| yeps_YPTB3520 | | | YPTB_RS19055 | YPTB3520 | | calcium/sodium antiporter | WP_002210120.1 |
| yeps_YPTB3521 | kdsD | | YPTB_RS19060 | YPTB3521 | | arabinose-5-phosphate isomerase KdsD | WP_002210119.1 |
| yeps_YPTB3522 | kdsC | | YPTB_RS19065 | YPTB3522 | | 3-deoxy-manno-octulosonate-8-phosphatase KdsC | WP_002228203.1 |
| yeps_YPTB3529 | rapZ | | YPTB_RS19100 | YPTB3529 | | RNase adapter RapZ | WP_002210113.1 |
| yeps_YPTB3530 | npr | | YPTB_RS19105 | YPTB3530 | | PTS phosphocarrier protein NPr | WP_002210112.1 |
| yeps_YPTB3532 | | | YPTB_RS19115 | YPTB3532 | | aspartate carbamoyltransferase regulatory subunit | WP_012104412.1 |
| yeps_YPTB3541 | pmbA | tldE | YPTB_RS19160 | YPTB3541 | | metalloprotease PmbA | WP_011193152.1 |
| yeps_YPTB3542 | | | YPTB_RS19165 | YPTB3542 | | ribosome-associated protein | WP_011193153.1 |
| yeps_YPTB3543 | | | YPTB_RS19170 | YPTB3543 | | ribonuclease inhibitor | WP_002210098.1 |
| yeps_YPTB3544 | | | YPTB_RS19175 | YPTB3544 | | ribonuclease Ba | WP_002214288.1 |
| yeps_YPTB3546 | aaeB | | YPTB_RS19185 | YPTB3546 | | p-hydroxybenzoic acid efflux pump subunit AaeB | WP_002210095.1 |
| yeps_YPTB3566 | csrD | | YPTB_RS19285 | YPTB3566 | | RNase E specificity factor CsrD | WP_011193164.1 |
| yeps_YPTB3567 | | | YPTB_RS19290 | YPTB3567 | | oxidoreductase | WP_002210074.1 |
| yeps_YPTB3571 | accB | fabE | YPTB_RS19310 | YPTB3571 | | acetyl-CoA carboxylase biotin carboxyl carrier protein | WP_002230506.1 |
| yeps_YPTB3579 | | | YPTB_RS19350 | YPTB3579 | | GntR family transcriptional regulator | WP_011193167.1 |
| yeps_YPTB3580 | | | YPTB_RS19355 | YPTB3580 | | MFS transporter | WP_011193168.1 |
| yeps_YPTB3581 | | | YPTB_RS19360 | YPTB3581 | | carboxymuconolactone decarboxylase family protein | WP_011193169.1 |
| yeps_YPTB3582 | | | YPTB_RS19365 | YPTB3582 | | NAD(P)-dependent oxidoreductase | WP_011193170.1 |
| yeps_YPTB3589 | | | YPTB_RS19415 | YPTB3589 | | esterase-like activity of phytase family protein | WP_011193175.1 |
| yeps_YPTB3590 | yghU | | YPTB_RS19420 | YPTB3590 | | glutathione-dependent disulfide-bond oxidoreductase | WP_032466981.1 |
| yeps_YPTB3591 | | | YPTB_RS19425 | YPTB3591 | | SIS domain-containing protein | WP_011193177.1 |
| yeps_YPTB3592 | | | YPTB_RS19430 | YPTB3592 | | FGGY-family carbohydrate kinase | WP_002210047.1 |
| yeps_YPTB3593 | | | YPTB_RS19435 | YPTB3593 | | ABC transporter permease | WP_011193178.1 |
| yeps_YPTB3595 | | | YPTB_RS19445 | YPTB3595 | | sugar ABC transporter ATP-binding protein | WP_002210044.1 |
| yeps_YPTB3596 | | | YPTB_RS19450 | YPTB3596 | | autoinducer 2 ABC transporter substrate-binding protein | WP_002210043.1 |
| yeps_YPTB3597 | lpxP | | YPTB_RS19455 | YPTB3597 | | kdo(2)-lipid IV(A) palmitoleoyltransferase | WP_002210042.1 |
| yeps_YPTB3598 | | | YPTB_RS19460 | YPTB3598 | | hypothetical protein | WP_011193180.1 |
| yeps_YPTB3599 | | | YPTB_RS19465 | YPTB3599 | | Na+/H+ antiporter | WP_002215816.1 |
| yeps_YPTB3603 | ssuC | | YPTB_RS19485 | YPTB3603 | | aliphatic sulfonate ABC transporter permease SsuC | WP_002213853.1 |
| yeps_YPTB3604 | ssuD | | YPTB_RS19490 | YPTB3604 | | FMNH2-dependent alkanesulfonate monooxygenase | WP_011193183.1 |
| yeps_YPTB3607 | deoC | dra | YPTB_RS19505 | YPTB3607 | | deoxyribose-phosphate aldolase | WP_011193185.1 |
| yeps_YPTB3608 | | | YPTB_RS19510 | YPTB3608 | | ribokinase | WP_011193186.1 |
| yeps_YPTB3609 | rbsD | | YPTB_RS19515 | YPTB3609 | | D-ribose pyranase | WP_011193187.1 |
| yeps_YPTB3610 | | | YPTB_RS19520 | YPTB3610 | | helix-turn-helix domain-containing protein | WP_012303467.1 |
| yeps_YPTB3611 | | | YPTB_RS19525 | YPTB3611 | | Gfo/Idh/MocA family oxidoreductase | WP_002210025.1 |
| yeps_YPTB3626 | tssA | | YPTB_RS19610 | YPTB3626 | | type VI secretion system protein TssA | WP_002212110.1 |
| yeps_YPTB3628 | | | YPTB_RS19620 | YPTB3628 | | Fis family transcriptional regulator | WP_002212108.1 |
| yeps_YPTB3629 | tssH | | YPTB_RS19625 | YPTB3629 | | type VI secretion system ATPase TssH | WP_002212107.1 |
| yeps_YPTB3631 | tssK | | YPTB_RS19635 | YPTB3631 | | type VI secretion system baseplate subunit TssK | WP_002212105.1 |
| yeps_YPTB3632 | tssJ | | YPTB_RS19640 | YPTB3632 | | type VI secretion system lipoprotein TssJ | WP_002212104.1 |
| yeps_YPTB3633 | tagH | | YPTB_RS19645 | YPTB3633 | | type VI secretion system-associated FHA domain protein TagH | WP_011193200.1 |
| yeps_YPTB3635 | tssF | | YPTB_RS19655 | YPTB3635 | | type VI secretion system baseplate subunit TssF | WP_011193201.1 |
| yeps_YPTB3636 | tssE | | YPTB_RS19660 | YPTB3636 | | type VI secretion system baseplate subunit TssE | WP_002212100.1 |
| yeps_YPTB3637 | tssC | | YPTB_RS19665 | YPTB3637 | | type VI secretion system contractile sheath large subunit | WP_002230616.1 |
| yeps_YPTB3638 | tssB | | YPTB_RS19670 | YPTB3638 | | type VI secretion system contractile sheath small subunit | WP_011193202.1 |
| yeps_YPTB3639 | | | YPTB_RS19675 | YPTB3639 | | Hcp family type VI secretion system effector | WP_011193203.1 |
| yeps_YPTB3642 | | | YPTB_RS19690 | YPTB3642 | | maltoporin | WP_002212092.1 |
| yeps_YPTB3643 | malK | | YPTB_RS19695 | YPTB3643 | | maltose/maltodextrin ABC transporter ATP-binding protein MalK | WP_002212091.1 |
| yeps_YPTB3645 | malE | | YPTB_RS19705 | YPTB3645 | | maltose/maltodextrin ABC transporter substrate-binding protein MalE | WP_002212089.1 |
| yeps_YPTB3646 | malF | | YPTB_RS19710 | YPTB3646 | | maltose ABC transporter permease MalF | WP_011193206.1 |
| yeps_YPTB3647 | malG | | YPTB_RS19715 | YPTB3647 | | maltose ABC transporter permease MalG | WP_002212087.1 |
| yeps_YPTB3648 | psiE | | YPTB_RS19720 | YPTB3648 | | phosphate-starvation-inducible protein PsiE | WP_002212086.1 |
| yeps_YPTB3650 | lysC | apk | YPTB_RS19730 | YPTB3650 | | lysine-sensitive aspartokinase 3 | WP_011193207.1 |
| yeps_YPTB3653 | metH | | YPTB_RS19750 | YPTB3653 | | methionine synthase | WP_038400771.1 |
| yeps_YPTB3654 | iclR | | YPTB_RS19755 | YPTB3654 | | glyoxylate bypass operon transcriptional repressor IclR | WP_002230619.1 |
| yeps_YPTB3661 | tsaC | | YPTB_RS19830 | YPTB3661 | | L-threonylcarbamoyladenylate synthase type 1 TsaC | WP_002209025.1 |
| yeps_YPTB3662 | | | YPTB_RS19835 | YPTB3662 | | topoisomerase DNA-binding C4 zinc finger domain-containing protein | WP_012105865.1 |
| yeps_YPTB3666 | fmt | | YPTB_RS19855 | YPTB3666 | | methionyl-tRNA formyltransferase | WP_002209020.1 |
| yeps_YPTB3669 | mscL | | YPTB_RS19870 | YPTB3669 | | large-conductance mechanosensitive channel protein MscL | WP_002209017.1 |
| yeps_YPTB3670 | | | YPTB_RS19875 | YPTB3670 | | alternative ribosome-rescue factor A | WP_002209016.1 |
| yeps_YPTB3671 | zntR | | YPTB_RS19880 | YPTB3671 | | Zn(2+)-responsive transcriptional regulator | WP_002215702.1 |
| yeps_YPTB3672 | rplQ | | YPTB_RS19885 | YPTB3672 | | 50S ribosomal protein L17 | WP_002209014.1 |
| yeps_YPTB3673 | rpoA | | YPTB_RS19890 | YPTB3673 | | DNA-directed RNA polymerase subunit alpha | WP_002209013.1 |
| yeps_YPTB3703 | fusA | far,fus | YPTB_RS20040 | YPTB3703 | | elongation factor G | WP_002212325.1 |
| yeps_YPTB3707 | tusC | | YPTB_RS20060 | YPTB3707 | | sulfurtransferase complex subunit TusC | WP_002212321.1 |
| yeps_YPTB3708 | tusD | | YPTB_RS20065 | YPTB3708 | | sulfurtransferase complex subunit TusD | WP_002212320.1 |
| yeps_YPTB3713 | | | YPTB_RS20090 | YPTB3713 | | YheV family putative metal-binding protein | WP_002212315.1 |
| yeps_YPTB3715 | kefG | | YPTB_RS20100 | YPTB3715 | | glutathione-regulated potassium-efflux system ancillary protein KefG | WP_002215966.1 |
| yeps_YPTB3716 | | | YPTB_RS20105 | YPTB3716 | | ABC transporter ATP-binding protein | WP_002230378.1 |
| yeps_YPTB3719 | tauD | ssiD | YPTB_RS20120 | YPTB3719 | | taurine dioxygenase | WP_002212310.1 |
| yeps_YPTB3720 | tauC | ssiC | YPTB_RS20125 | YPTB3720 | | taurine ABC transporter permease TauC | WP_002212309.1 |
| yeps_YPTB3721 | tauB | ssiB | YPTB_RS20130 | YPTB3721 | | taurine ABC transporter ATP-binding subunit | WP_011193220.1 |
| yeps_YPTB3723 | | | YPTB_RS20140 | YPTB3723 | | LysE family translocator | WP_011193221.1 |
| yeps_YPTB3724 | | | YPTB_RS20145 | YPTB3724 | | hydrolase | WP_002212302.1 |
| yeps_YPTB3726 | | | YPTB_RS20155 | YPTB3726 | | phosphoribulokinase | WP_002212300.1 |
| yeps_YPTB3727 | | | YPTB_RS20160 | YPTB3727 | | DUF1240 domain-containing protein | WP_002228139.1 |
| yeps_YPTB3728 | | | YPTB_RS20165 | YPTB3728 | | OsmC family protein | WP_002212298.1 |
| yeps_YPTB3735 | | | YPTB_RS20200 | YPTB3735 | | 6-phospho-beta-glucosidase | WP_011193227.1 |
| yeps_YPTB3736 | | | YPTB_RS20205 | YPTB3736 | | LacI family transcriptional regulator | WP_002208881.1 |
| yeps_YPTB3737 | | | YPTB_RS20210 | YPTB3737 | | TonB-dependent copper receptor | WP_011193228.1 |
| yeps_YPTB3739 | | | YPTB_RS20220 | YPTB3739 | | cytosine deaminase | WP_011193230.1 |
| yeps_YPTB3740 | | | YPTB_RS20225 | YPTB3740 | | nitrite reductase large subunit NirB | WP_011193231.1 |
| yeps_YPTB3741 | nirD | | YPTB_RS20230 | YPTB3741 | | nitrite reductase small subunit NirD | WP_002208887.1 |
| yeps_YPTB3742 | nirC | | YPTB_RS20235 | YPTB3742 | | nitrite transporter NirC | WP_002208888.1 |
| yeps_YPTB3743 | cysG | | | YPTB3743 | | uroporphyrin-III C-methyltransferase / precorrin-2 oxidase /... | CAH22981.1 |
| yeps_YPTB3744 | trpS | | YPTB_RS20245 | YPTB3744 | | tryptophan--tRNA ligase | WP_002208891.1 |
| yeps_YPTB3745 | | | YPTB_RS20250 | YPTB3745 | | phosphoglycolate phosphatase | WP_011193233.1 |
| yeps_YPTB3747 | dam | | YPTB_RS20260 | YPTB3747 | | adenine-specific DNA-methyltransferase | WP_002208895.1 |
| yeps_YPTB3748 | | | YPTB_RS20265 | YPTB3748 | | SPOR domain-containing protein | WP_011193234.1 |
| yeps_YPTB3754 | | | YPTB_RS20295 | YPTB3754 | | PilN domain-containing protein | WP_011193236.1 |
| yeps_YPTB3755 | pilM | | YPTB_RS20300 | YPTB3755 | | pilus assembly protein PilM | WP_011193237.1 |
| yeps_YPTB3757 | nudE | | YPTB_RS20310 | YPTB3757 | | ADP compounds hydrolase NudE | WP_002208907.1 |
| yeps_YPTB3758 | | | YPTB_RS20320 | YPTB3758 | | intracellular growth attenuator family protein | WP_011193239.1 |
| yeps_YPTB3759 | yrfG | | YPTB_RS20325 | YPTB3759 | | GMP/IMP nucleotidase | WP_002208909.1 |
| yeps_YPTB3761 | hslO | | YPTB_RS20335 | YPTB3761 | | Hsp33 family molecular chaperone HslO | WP_002208911.1 |
| yeps_YPTB3765 | greB | | YPTB_RS20355 | YPTB3765 | | transcription elongation factor GreB | WP_002208915.1 |
| yeps_YPTB3769 | | | YPTB_RS20380 | YPTB3769 | | ferrous iron transporter C | WP_002208920.1 |
| yeps_YPTB3770 | | | YPTB_RS20385 | YPTB3770 | | DUF1471 domain-containing protein | WP_002208921.1 |
| yeps_YPTB3771 | bioH | | YPTB_RS20390 | YPTB3771 | | pimeloyl-ACP methyl ester esterase BioH | WP_011193243.1 |
| yeps_YPTB3772 | gntX | | YPTB_RS20395 | YPTB3772 | | DNA utilization protein GntX | WP_002208923.1 |
| yeps_YPTB3773 | nfuA | | YPTB_RS20400 | YPTB3773 | | Fe-S biogenesis protein NfuA | WP_002208924.1 |
| yeps_YPTB3774 | malQ | | YPTB_RS23610 | YPTB3774 | | 4-alpha-glucanotransferase | WP_002208925.1 |
| yeps_YPTB3775 | malP | | YPTB_RS23615 | YPTB3775 | | maltodextrin phosphorylase | WP_011193244.1 |
| yeps_YPTB3776 | malT | | YPTB_RS20415 | YPTB3776 | | HTH-type transcriptional regulator MalT | WP_002208926.1 |
| yeps_YPTB3777 | glpE | | YPTB_RS20420 | YPTB3777 | | thiosulfate sulfurtransferase GlpE | WP_002218928.1 |
| yeps_YPTB3778 | glpG | | YPTB_RS20425 | YPTB3778 | | rhomboid family intramembrane serine protease GlpG | WP_011193245.1 |
| yeps_YPTB3783 | glgP | glgY | YPTB_RS20460 | YPTB3783 | | glycogen phosphorylase | WP_011193248.1 |
| yeps_YPTB3784 | glgA | | YPTB_RS20465 | YPTB3784 | | glycogen synthase GlgA | WP_002209498.1 |

| | | | | | | | | |
|---|---|---|---|---|---|---|---|---|
| yeps_YPTB3785 | glgC | | YPTB_RS20470 | YPTB3785 | | glucose-1-phosphate adenylyltransferase | WP_011193249.1 | |
| yeps_YPTB3788 | | | YPTB_RS20485 | YPTB3788 | | sensor histidine kinase | WP_011193252.1 | |
| yeps_YPTB3796 | | | YPTB_RS20535 | YPTB3796 | | gluconokinase | WP_002209512.1 | |
| yeps_YPTB3797 | gntT | gntM,usgA | YPTB_RS20540 | YPTB3797 | | gluconate transporter | WP_011193259.1 | |
| yeps_YPTB3798 | gntR | | YPTB_RS20545 | YPTB3798 | | gluconate operon transcriptional repressor GntR | WP_011193260.1 | |
| yeps_YPTB3799 | | | YPTB_RS20550 | YPTB3799 | | pirin family protein | WP_002209515.1 | |
| yeps_YPTB3802 | | | YPTB_RS20565 | YPTB3802 | | adenosine kinase | WP_011193262.1 | |
| yeps_YPTB3803 | | | YPTB_RS20570 | YPTB3803 | | ketose 1,6-bisphosphate aldolase | WP_002209519.1 | |
| yeps_YPTB3804 | | | YPTB_RS20575 | YPTB3804 | | D-lyxose/D-mannose family sugar isomerase | WP_011193263.1 | |
| yeps_YPTB3805 | | | YPTB_RS20580 | YPTB3805 | | ABC transporter substrate-binding protein | WP_002209521.1 | |
| yeps_YPTB3806 | | | YPTB_RS20585 | YPTB3806 | | ribose ABC transporter permease | WP_002209522.1 | |
| yeps_YPTB3812 | uspA | | YPTB_RS20615 | YPTB3812 | | universal stress protein UspA | WP_002209528.1 | |
| yeps_YPTB3813 | gdhA | | YPTB_RS20620 | YPTB3813 | | NADP-specific glutamate dehydrogenase | WP_002209529.1 | |
| yeps_YPTB3814 | | | YPTB_RS20625 | YPTB3814 | | M10 family metallopeptidase C-terminal domain-containing protein | WP_012414110.1 | |
| yeps_YPTB3815 | rsmJ | | YPTB_RS20630 | YPTB3815 | | 16S rRNA (guanine(1516)-N(2))-methyltransferase RsmJ | WP_002215483.1 | |
| yeps_YPTB3818 | gorA | | YPTB_RS20645 | YPTB3818 | | glutathione-disulfide reductase | WP_032466815.1 | |
| yeps_YPTB3820 | | | YPTB_RS20655 | YPTB3820 | | GntP family permease | WP_011193273.1 | |
| yeps_YPTB3821 | | | YPTB_RS20660 | YPTB3821 | | glycerate kinase | WP_011193274.1 | |
| yeps_YPTB3829 | | | | YPTB3829 | | possible Tautomerase enzyme | CAH23067.1 | |
| yeps_YPTB3830 | | | YPTB_RS20705 | YPTB3830 | | sugar kinase | WP_011193283.1 | |
| yeps_YPTB3832 | | | YPTB_RS20715 | YPTB3832 | | dicarboxylate/amino acid:cation symporter | WP_002209553.1 | |
| yeps_YPTB3834 | | | YPTB_RS20725 | YPTB3834 | | pectate lyase | WP_002209555.1 | |
| yeps_YPTB3838 | | | YPTB_RS20745 | YPTB3838 | | ABC transporter ATP-binding protein | WP_161597821.1 | |
| yeps_YPTB3839 | dppD | | YPTB_RS20750 | YPTB3839 | | dipeptide ABC transporter ATP-binding protein | WP_002209562.1 | |
| yeps_YPTB3841 | dppB | | YPTB_RS20760 | YPTB3841 | | dipeptide ABC transporter permease DppB | WP_002228714.1 | |
| yeps_YPTB3842 | | | YPTB_RS20765 | YPTB3842 | | ABC transporter substrate-binding protein | WP_011193287.1 | |
| yeps_YPTB3846 | uhpB | | YPTB_RS20795 | YPTB3846 | | signal transduction histidine-protein kinase/phosphatase UhpB | WP_002230402.1 | |
| yeps_YPTB3848 | eptB | | YPTB_RS20805 | YPTB3848 | | kdo(2)-lipid A phosphoethanolamine 7''-transferase | WP_011193291.1 | |
| yeps_YPTB3850 | | | YPTB_RS20815 | YPTB3850 | | amino acid permease | WP_011193292.1 | |
| yeps_YPTB3856 | | | YPTB_RS20845 | YPTB3856 | | ATP-grasp domain-containing protein | WP_011193294.1 | |
| yeps_YPTB3857 | | | YPTB_RS20850 | YPTB3857 | | siderophore ABC transporter substrate-binding protein | WP_002209581.1 | |
| yeps_YPTB3858 | | | YPTB_RS20855 | YPTB3858 | | ABC transporter permease | WP_011193295.1 | |
| yeps_YPTB3859 | | | YPTB_RS20860 | YPTB3859 | | iron chelate uptake ABC transporter family permease subunit | WP_011193296.1 | |
| yeps_YPTB3888 | | | YPTB_RS21005 | YPTB3888 | | sugar ABC transporter permease | WP_002209589.1 | |
| yeps_YPTB3904 | ibpA | hslT,htpN | YPTB_RS21085 | YPTB3904 | | heat shock chaperone IbpA | WP_002209636.1 | |
| yeps_YPTB3905 | ibpB | hslS,htpE | YPTB_RS21090 | YPTB3905 | | heat shock chaperone IbpB | WP_002209635.1 | |
| yeps_YPTB3906 | | | YPTB_RS21095 | YPTB3906 | | putative transporter | WP_011193331.1 | |
| yeps_YPTB3910 | ghrB | | YPTB_RS21115 | YPTB3910 | | glyoxylate/hydroxypyruvate reductase GhrB | WP_002209630.1 | |
| yeps_YPTB3911 | | | YPTB_RS21120 | YPTB3911 | | transcriptional regulator | WP_011193334.1 | |
| yeps_YPTB3912 | | | YPTB_RS21125 | YPTB3912 | | OmpA family lipoprotein | WP_002209628.1 | |
| yeps_YPTB3913 | | | YPTB_RS21130 | YPTB3913 | | N-acetyltransferase | WP_011193335.1 | |
| yeps_YPTB3919 | | | YPTB_RS21160 | YPTB3919 | | mannitol-1-phosphate 5-dehydrogenase | WP_011193338.1 | |
| yeps_YPTB3922 | | | YPTB_RS21175 | YPTB3922 | | DM13 domain-containing protein | WP_002209617.1 | |
| yeps_YPTB3925 | sodA | | YPTB_RS21190 | YPTB3925 | | superoxide dismutase [Mn] | WP_002209614.1 | |
| yeps_YPTB3926 | fdhD | | YPTB_RS21195 | YPTB3926 | | formate dehydrogenase accessory sulfurtransferase FdhD | WP_011193341.1 | |
| yeps_YPTB3929 | fdoI | | YPTB_RS21215 | YPTB3929 | | formate dehydrogenase cytochrome b556 subunit | WP_011193343.1 | |
| yeps_YPTB3930 | fdhE | | YPTB_RS21220 | YPTB3930 | | formate dehydrogenase accessory protein FdhE | WP_002209609.1 | |
| yeps_YPTB3931 | | | YPTB_RS21225 | YPTB3931 | | L-seryl-tRNA(Sec) selenium transferase | WP_011193344.1 | |
| yeps_YPTB3941 | recF | uvrF | YPTB_RS21285 | YPTB3941 | | DNA replication/repair protein RecF | WP_002209643.1 | |
| yeps_YPTB3943 | dnaA | | YPTB_RS21300 | YPTB3943 | | chromosomal replication initiator protein DnaA | WP_002220732.1 | |
| yeps_YPTB3944 | | | | YPTB3944 | | hypothetical protein | CAH23182.1 | |
| yeps_YPTB3948 | yidC | | YPTB_RS21315 | YPTB3948 | | membrane protein insertase YidC | WP_002220739.1 | |
| yeps_YPTB3949 | mnmE | | YPTB_RS21320 | YPTB3949 | | tRNA uridine-5-carboxymethylaminomethyl(34) synthesis GTPase MnmE | WP_011193350.1 | |
| yeps_YPTB3967 | atpD | papB,uncD | YPTB_RS21410 | YPTB3967 | | F0F1 ATP synthase subunit beta | WP_002220753.1 | |
| yeps_YPTB3968 | atpG | papC,uncG | YPTB_RS21415 | YPTB3968 | | F0F1 ATP synthase subunit gamma | WP_002220756.1 | |
| yeps_YPTB3972 | atpE | papH,uncE | YPTB_RS21435 | YPTB3972 | | F0F1 ATP synthase subunit C | WP_000429386.1 | |
| x_1 | glnA | | YPTB_RS00215 | YPTB0024 | | glutamate--ammonia ligase | WP_002213150.1 | |
| x_2 | | | | | BZ17_946 | hypothetical protein | AJJ56340.1 | |
| x_3 | | | | | BZ17_887 | putative membrane protein | AJJ57145.1 | |
| x_4 | | | | | BZ17_159 | putative membrane protein | AJJ56946.1 | |
| x_5 | | | | | BZ17_111 | hypothetical protein | AJJ56799.1 | |
| x_6 | fdnG | | YPTB_RS21205 | YPTB3927 | | formate dehydrogenase-N subunit alpha | WP_011193342.1 | fragment, translation exception |
| yeps_x_1 | typA | bipA | YPTB_RS00225 | YPTB0025 | | ribosome-dependent GTPase TypA | WP_002217380.1 | fragment on the opposite strand |
| yeps_x_10 | emrB | | YPTB_RS04770 | YPTB0855 | | multidrug efflux MFS transporter permease subunit EmrB | WP_002208746.1 | fragment on the opposite strand |
| yeps_x_11 | | | YPTB_RS04895 | YPTB0880 | | filamentous hemagglutinin N-terminal domain-containing protein | WP_011191837.1 | fragment on the same strand but different frame |
| yeps_x_12 | | | YPTB_RS04895 | YPTB0880 | | filamentous hemagglutinin N-terminal domain-containing protein | WP_011191837.1 | fragment on the opposite strand |
| yeps_x_13 | | | YPTB_RS04895 | YPTB0880 | | filamentous hemagglutinin N-terminal domain-containing protein | WP_011191837.1 | fragment on the opposite strand |
| yeps_x_14 | aroL | | YPTB_RS05055 | YPTB0911 | | shikimate kinase AroL | WP_011191851.1 | |
| yeps_x_15 | | | | | BZ17_1504 | rhs element Vgr domain protein | AJJ53360.1 | |
| yeps_x_16 | | | YPTB_RS22525 | YPTB1081 | | hypothetical protein | WP_011191926.1 | fragment on the opposite strand |
| yeps_x_17 | | | | | BZ17_1459 | glycogen synthesis family protein | AJJ54910.1 | |
| yeps_x_18 | | | | | BZ17_1356 | hypothetical protein | AJJ56909.1 | |
| yeps_x_19 | | | | | BZ17_1338 | hypothetical protein | AJJ57070.1 | |
| yeps_x_2 | typA | bipA | YPTB_RS00225 | YPTB0025 | | ribosome-dependent GTPase TypA | WP_002217380.1 | fragment on the opposite strand |
| yeps_x_20 | | | | | BZ17_1234 | hypothetical protein | AJJ56635.1 | |
| yeps_x_21 | | | | | BZ17_1181 | hypothetical protein | AJJ53380.1 | |
| yeps_x_22 | | | YPTB_RS07475 | YPTB1352 | | HAAAP family serine/threonine permease | WP_011192035.1 | fragment on the opposite strand |
| yeps_x_23 | | | | | BZ17_1069 | putative membrane protein | AJJ54317.1 | |
| yeps_x_24 | | | YPTB_RS08855 | YPTB1608 | | alpha-galactosidase | WP_011192140.1 | fragment on the opposite strand |
| yeps_x_25 | mtfA | | YPTB_RS08875 | YPTB1611 | | DgsA anti-repressor MtfA | WP_002211042.1 | fragment on the opposite strand |
| yeps_x_26 | | | YPTB_RS09000 | YPTB1632 | | PTS mannose transporter subunit IID | WP_002211067.1 | fragment on the opposite strand |
| yeps_x_27 | | | | YPTB1706 | BZ17_795 | hypothetical protein | CAH20945.1 | fragment |
| yeps_x_28 | | | | | BZ17_562 | hypothetical protein | AJJ54038.1 | |
| yeps_x_29 | | | | | BZ17_511 | hypothetical protein | AJJ56644.1 | |
| yeps_x_3 | | | | | BZ17_2521 | hypothetical protein | AJJ56380.1 | |
| yeps_x_30 | | | | | BZ17_482 | hypothetical protein | AJJ56760.1 | |
| yeps_x_31 | | | | | | | | |
| yeps_x_32 | | | | | BZ17_324 | hypothetical protein | AJJ55948.1 | |
| yeps_x_33 | | | YPTB_RS12180 | YPTB2233 | | RHS repeat protein | CAH21471.1 | pseudogene ; fragment on the opposite strand |
| yeps_x_34 | | | | | BZ17_162 | hypothetical protein | AJJ54783.1 | |
| yeps_x_35 | | | | | BZ17_66 | hypothetical protein | AJJ55378.1 | |
| yeps_x_36 | | | | | BZ17_60 | putative membrane protein | AJJ55430.1 | |
| yeps_x_37 | | | | | BZ17_29 | putative membrane protein | AJJ53437.1 | |
| yeps_x_38 | | | | | | | | |
| yeps_x_39 | | | YPTB_RS13510 | YPTB2487 | | AI-2E family transporter | WP_002211853.1 | fragment on the opposite strand |
| yeps_x_4 | | | | | BZ17_2291 | hypothetical protein | AJJ55702.1 | |
| yeps_x_40 | | | | YPTB2564 | | hypothetical protein | CAH21802.1 | fragment on the opposite strand |
| yeps_x_41 | nuoM | | YPTB_RS13980 | YPTB2576 | | NADH-quinone oxidoreductase subunit M | WP_011192666.1 | fragment on the opposite strand |
| yeps_x_42 | | | | | BZ17_3718 | hypothetical protein | AJJ54599.1 | |
| yeps_x_43 | kdpD | | YPTB_RS15780 | YPTB2919 | | two-component system sensor histidine kinase KdpD | WP_002209649.1 | fragment on the opposite strand |
| yeps_x_44 | | | | YPTB3040 | | | | pseudogene |
| yeps_x_45 | | | YPTB_RS16755 | YPTB3097 | | beta-galactosidase | WP_002209897.1 | fragment on the opposite strand |
| yeps_x_46 | | | YPTB_RS17130 | YPTB3162 | | tyrosine-type recombinase/integrase | WP_011192954.1 | fragment on the opposite strand |
| yeps_x_47 | prfB | | YPTB_RS17145 | YPTB3164 | | peptide chain release factor 2 | WP_002228062.1 | fragment |
| yeps_x_48 | gcvP | | YPTB_RS17230 | YPTB3180 | | aminomethyl-transferring glycine dehydrogenase | WP_002209947.1 | fragment on the opposite strand |
| yeps_x_49 | | | | | BZ17_3408 | hypothetical protein | AJJ55544.1 | |
| yeps_x_5 | | | | | BZ17_1959 | hypothetical protein | AJJ56833.1 | |
| yeps_x_50 | | | YPTB_RS17680 | YPTB3266 | | lysine N(6)-hydroxylase/L-ornithine N(5)-oxygenase family protein | WP_011193002.1 | fragment on the opposite strand ; overlaps next cds |
| yeps_x_51 | | | | | BZ17_3208 | hypothetical protein | AJJ55852.1 | |
| yeps_x_52 | | | YPTB_RS18590 | YPTB3435 | | membrane protein | WP_002214224.1 | |
| yeps_x_53 | uxaC | | YPTB_RS18825 | YPTB3478 | | glucuronate isomerase | WP_002210410.1 | fragment on the opposite strand |
| yeps_x_54 | | | | | BZ17_3092 | hypothetical protein | AJJ56691.1 | |
| yeps_x_55 | murA | | YPTB_RS19020 | YPTB3513 | | UDP-N-acetylglucosamine 1-carboxyvinyltransferase | WP_002210127.1 | fragment on the opposite strand |
| yeps_x_56 | yhdP | | YPTB_RS19255 | YPTB3560 | | AsmA2 domain-containing protein YhdP | WP_011193163.1 | fragment on the opposite strand |
| yeps_x_57 | metH | | YPTB_RS19750 | YPTB3653 | | methionine synthase | WP_038400771.1 | fragment on the opposite strand |
| yeps_x_58 | mrcA | | YPTB_RS20305 | YPTB3756 | | peptidoglycan glycosyltransferase/peptidoglycan DD-transpeptidase MrcA | WP_011193238.1 | fragment on the opposite strand |
| yeps_x_59 | malT | | YPTB_RS20415 | YPTB3776 | | HTH-type transcriptional regulator MalT | WP_002208926.1 | fragment on the opposite strand |
| yeps_x_6 | | | YPTB_RS03685 | YPTB0659 | | DNA polymerase II | WP_011191720.1 | fragment on the opposite strand |
| yeps_x_60 | | | | | BZ17_2753 | hypothetical protein | AJJ56110.1 | |
| yeps_x_61 | | | | | BZ17_2741 | hypothetical protein | AJJ54297.1 | |
| yeps_x_62 | | | YPTB_RS21095 | YPTB3906 | | putative transporter | WP_011193331.1 | fragment on the opposite strand |
| yeps_x_63 | | | | | BZ17_2660 | hypothetical protein | AJJ53741.1 | |
| yeps_x_64 | fdnG | | YPTB_RS21205 | YPTB3927 | BZ17_2652 | formate dehydrogenase-N subunit alpha | WP_011193342.1 | fragment |
| yeps_x_65 | atpC | | YPTB_RS21405 | YPTB3966 | | F0F1 ATP synthase subunit epsilon | WP_002215546.1 | fragment on the opposite strand |
| yeps_x_66 | atpD | | YPTB_RS21410 | YPTB3967 | | F0F1 ATP synthase subunit beta | WP_002220753.1 | fragment on the opposite strand |
| yeps_x_7 | pyrG | | YPTB_RS04195 | YPTB0754 | | CTP synthase (glutamine hydrolyzing) | WP_002209376.1 | fragment on the opposite strand |
| yeps_x_8 | | | | | | | | |
| yeps_x_9 | emrB | | YPTB_RS04770 | YPTB0855 | | multidrug efflux MFS transporter permease subunit EmrB | WP_002208746.1 | fragment on the opposite strand |
| YPTB0002 | asnC | | YPTB_RS00080 | YPTB0002 | | transcriptional regulator AsnC | WP_002212257.1 | |
| YPTB0003 | | | YPTB_RS00085 | YPTB0003 | | aspartate--ammonia ligase | WP_002212256.1 | |

| | | | | | | | |
|---|---|---|---|---|---|---|---|
| YPTB0028 | dtd | | YPTB_RS00240 | YPTB0028 | | D-tyrosyl-tRNA(Tyr) deacylase | WP_002209009.1 |
| YPTB0033 | recG | | YPTB_RS00265 | YPTB0033 | | ATP-dependent DNA helicase RecG | WP_002209004.1 |
| YPTB0037 | gmk | spoR | YPTB_RS00285 | YPTB0037 | | guanylate kinase | WP_002209000.1 |
| YPTB0044 | dut | dnaS,sof | YPTB_RS00320 | YPTB0044 | | dUTP diphosphatase | WP_011566231.1 |
| YPTB0047 | rpmB | | YPTB_RS00335 | YPTB0047 | | 50S ribosomal protein L28 | WP_002208991.1 |
| YPTB0055 | rfaD | htrM | YPTB_RS00375 | YPTB0055 | | ADP-glyceromanno-heptose 6-epimerase | WP_002208983.1 |
| YPTB0070 | cpxR | | YPTB_RS00450 | YPTB0070 | | envelope stress response regulator transcription factor CpxR | WP_002208970.1 |
| YPTB0086 | glpK | | YPTB_RS00530 | YPTB0086 | | glycerol kinase GlpK | WP_002218876.1 |
| YPTB0089 | zapB | | YPTB_RS00545 | YPTB0089 | | septal ring assembly protein ZapB | WP_002208953.1 |
| YPTB0097 | hslU | | YPTB_RS00585 | YPTB0097 | | HslU--HslV peptidase ATPase subunit | WP_002208943.1 |
| YPTB0101 | priA | | YPTB_RS00610 | YPTB0101 | | primosomal protein N' | WP_011191465.1 |
| YPTB0104 | metJ | | YPTB_RS00630 | YPTB0104 | | met regulon transcriptional regulator MetJ | WP_004392248.1 |
| YPTB0121 | sthA | sth,udhA | YPTB_RS00725 | YPTB0121 | | Si-specific NAD(P)(+) transhydrogenase | WP_002209477.1 |
| YPTB0132 | | | YPTB_RS00820 | YPTB0132 | | DUF413 domain-containing protein | WP_002212019.1 |
| YPTB0134 | | | YPTB_RS00830 | YPTB0134 | | acetolactate synthase 2 catalytic subunit | WP_011191482.1 |
| YPTB0137 | ilvD | | YPTB_RS00845 | YPTB0137 | | dihydroxy-acid dehydratase | WP_011191483.1 |
| YPTB0166 | trxA | fipA,tsnC | YPTB_RS01010 | YPTB0166 | | thioredoxin TrxA | WP_002211990.1 |
| YPTB0167 | rho | | YPTB_RS01015 | YPTB0167 | | transcription termination factor Rho | WP_002211989.1 |
| YPTB0171 | wecC | | YPTB_RS01035 | YPTB0171 | | UDP-N-acetyl-D-mannosamine dehydrogenase | WP_011191499.1 |
| YPTB0177 | | | YPTB_RS01065 | YPTB0177 | | TDP-N-acetylfucosamine:lipid II N-acetylfucosaminyltransferase | WP_011191503.1 |
| YPTB0182 | hemX | | YPTB_RS01110 | YPTB0182 | | uroporphyrinogen-III C-methyltransferase | WP_011191506.1 |
| YPTB0190 | dapF | | YPTB_RS01155 | YPTB0190 | | diaminopimelate epimerase | WP_002211471.1 |
| YPTB0193 | yigB | | YPTB_RS01170 | YPTB0193 | | 5-amino-6-(5-phospho-D-ribitylamino)uracil phosphatase YigB | WP_011191513.1 |
| YPTB0219 | | | YPTB_RS01305 | YPTB0219 | | DUF1145 family protein | WP_002211501.1 |
| YPTB0222 | ftsE | | YPTB_RS01325 | YPTB0222 | | cell division ATP-binding protein FtsE | WP_004391337.1 |
| YPTB0224 | rpoH | fam,hin,htpR | YPTB_RS01335 | YPTB0224 | | RNA polymerase sigma factor RpoH | WP_002211506.1 |
| YPTB0227 | livH | | YPTB_RS01350 | YPTB0227 | | high-affinity branched-chain amino acid ABC transporter permease LivH | WP_002211509.1 |
| YPTB0228 | | | YPTB_RS01355 | YPTB0228 | | high-affinity branched-chain amino acid ABC transporter permease LivM | WP_002211510.1 |
| YPTB0229 | livG | | YPTB_RS01360 | YPTB0229 | | high-affinity branched-chain amino acid ABC transporter ATP-binding protein LivG | WP_011191524.1 |
| YPTB0240 | ugpE | | YPTB_RS01415 | YPTB0240 | | sn-glycerol-3-phosphate ABC transporter permease UgpE | WP_002211522.1 |
| YPTB0247 | metR | | YPTB_RS01450 | YPTB0247 | | HTH-type transcriptional regulator MetR | WP_011191537.1 |
| YPTB0250 | udp | | YPTB_RS01465 | YPTB0250 | | uridine phosphorylase | WP_002211528.1 |
| YPTB0261 | tatD | | YPTB_RS01520 | YPTB0261 | | 3'-5' ssDNA/RNA exonuclease TatD | WP_011191544.1 |
| YPTB0262 | hemB | | YPTB_RS01525 | YPTB0262 | | porphobilinogen synthase | WP_002211541.1 |
| YPTB0264 | ubiD | | YPTB_RS01535 | YPTB0264 | | 4-hydroxy-3-polyprenylbenzoate decarboxylase | WP_011191546.1 |
| YPTB0265 | fre | | YPTB_RS01540 | YPTB0265 | | NAD(P)H-flavin reductase | WP_002215918.1 |
| YPTB0266 | fadA | oldA | YPTB_RS01545 | YPTB0266 | | acetyl-CoA C-acyltransferase FadA | WP_002211545.1 |
| YPTB0267 | fadB | oldB | YPTB_RS01550 | YPTB0267 | | fatty acid oxidation complex subunit alpha FadB | WP_011191547.1 |
| YPTB0270 | trkH | | YPTB_RS01565 | YPTB0270 | | Trk system potassium transporter TrkH | WP_002211550.1 |
| YPTB0277 | secE | | YPTB_RS01650 | YPTB0277 | | preprotein translocase subunit SecE | WP_002210670.1 |
| YPTB0278 | nusG | | YPTB_RS01655 | YPTB0278 | | transcription termination/antitermination protein NusG | WP_002210671.1 |
| YPTB0279 | rplK | relC | YPTB_RS01660 | YPTB0279 | | 50S ribosomal protein L11 | WP_002210672.1 |
| YPTB0280 | rplA | | YPTB_RS01665 | YPTB0280 | | 50S ribosomal protein L1 | WP_002210673.1 |
| YPTB0282 | rplL | | YPTB_RS01675 | YPTB0282 | | 50S ribosomal protein L7/L12 | WP_002210675.1 |
| YPTB0296 | | | YPTB_RS01755 | YPTB0296 | | YjaG family protein | WP_002230622.1 |
| YPTB0337 | | | YPTB_RS01985 | YPTB0337 | | iron ABC transporter permease | WP_002209059.1 |
| YPTB0370 | lexA | exrA,spr,tsl,umuA | YPTB_RS02150 | YPTB0370 | | transcriptional repressor LexA | WP_002209090.1 |
| YPTB0373 | pspG | | YPTB_RS02165 | YPTB0373 | | envelope stress response protein PspG | WP_002209093.1 |
| YPTB0375 | dnaB | groP,grpA | YPTB_RS02175 | YPTB0375 | | replicative DNA helicase | WP_002209095.1 |
| YPTB0376 | alr | | YPTB_RS02180 | YPTB0376 | | alanine racemase | WP_011191596.1 |
| YPTB0377 | | | YPTB_RS02185 | YPTB0377 | | aspartate/tyrosine/aromatic aminotransferase | WP_011191597.1 |
| YPTB0402 | aspA | | YPTB_RS02315 | YPTB0402 | | aspartate ammonia-lyase | WP_002230464.1 |
| YPTB0404 | | | YPTB_RS02325 | YPTB0404 | | co-chaperone GroES | WP_002209127.1 |
| YPTB0408 | efp | | YPTB_RS02345 | YPTB0408 | | elongation factor P | WP_002209131.1 |
| YPTB0411 | frdC | | YPTB_RS02360 | YPTB0411 | | fumarate reductase subunit FrdC | WP_002209135.1 |
| YPTB0412 | | | YPTB_RS02365 | YPTB0412 | | succinate dehydrogenase/fumarate reductase iron-sulfur subunit | WP_002209136.1 |
| YPTB0424 | miaA | trpX | YPTB_RS02455 | YPTB0424 | | tRNA (adenosine(37)-N6)-dimethylallyltransferase MiaA | WP_002209149.1 |
| YPTB0425 | hfq | | YPTB_RS02460 | YPTB0425 | | RNA chaperone Hfq | WP_002209151.1 |
| YPTB0438 | rpsF | | YPTB_RS02525 | YPTB0438 | | 30S ribosomal protein S6 | WP_002210153.1 |
| YPTB0439 | priB | | YPTB_RS02530 | YPTB0439 | | primosomal replication protein N | WP_002210154.1 |
| YPTB0440 | rpsR | | YPTB_RS02535 | YPTB0440 | | 30S ribosomal protein S18 | WP_002210155.1 |
| YPTB0441 | rplI | | YPTB_RS02540 | YPTB0441 | | 50S ribosomal protein L9 | WP_002210156.1 |
| YPTB0444 | | | YPTB_RS02555 | YPTB0444 | | peptidylprolyl isomerase | WP_011191620.1 |
| YPTB0450 | | | YPTB_RS02585 | YPTB0450 | | hemolysin family protein | WP_002210164.1 |
| YPTB0463 | ispB | cel | YPTB_RS02655 | YPTB0463 | | octaprenyl diphosphate synthase | WP_011191628.1 |
| YPTB0464 | rplU | | YPTB_RS02660 | YPTB0464 | | 50S ribosomal protein L21 | WP_002210178.1 |
| YPTB0465 | rpmA | | YPTB_RS02665 | YPTB0465 | | 50S ribosomal protein L27 | WP_002210179.1 |
| YPTB0470 | dacB | | YPTB_RS02690 | YPTB0470 | | serine-type D-Ala-D-Ala carboxypeptidase | WP_002217314.1 |
| YPTB0471 | greA | | YPTB_RS02695 | YPTB0471 | | transcription elongation factor GreA | WP_002210184.1 |
| YPTB0472 | yhbY | | YPTB_RS02700 | YPTB0472 | | ribosome assembly RNA-binding protein YhbY | WP_002210185.1 |
| YPTB0473 | rlmE | | YPTB_RS02705 | YPTB0473 | | 23S rRNA (uridine(2552)-2'-O)-methyltransferase RlmE | WP_002228196.1 |
| YPTB0475 | folP | dhpS | YPTB_RS02715 | YPTB0475 | | dihydropteroate synthase | WP_011191630.1 |
| YPTB0476 | glmM | | YPTB_RS02720 | YPTB0476 | | phosphoglucosamine mutase | WP_002210189.1 |
| YPTB0478 | rimP | | YPTB_RS02740 | YPTB0478 | | ribosome maturation factor RimP | WP_002222054.1 |
| YPTB0481 | rbfA | p15B | YPTB_RS02755 | YPTB0481 | | 30S ribosome-binding factor RbfA | WP_002209255.1 |
| YPTB0483 | rpsO | | YPTB_RS02765 | YPTB0483 | | 30S ribosomal protein S15 | WP_002209257.1 |
| YPTB0485 | nlpI | | YPTB_RS02775 | YPTB0485 | | lipoprotein NlpI | WP_002209260.1 |
| YPTB0494 | | | YPTB_RS02820 | YPTB0494 | | U32 family peptidase | WP_002209269.1 |
| YPTB0523 | | | YPTB_RS02975 | YPTB0523 | | ABC transporter permease | WP_002209300.1 |
| YPTB0530 | | | YPTB_RS03010 | YPTB0530 | | DNA polymerase III subunit chi | WP_002209309.1 |
| YPTB0531 | pepA | ampA,carP,xerB | YPTB_RS03015 | YPTB0531 | | leucyl aminopeptidase | WP_002209310.1 |
| YPTB0532 | lptF | | YPTB_RS03020 | YPTB0532 | | LPS export ABC transporter permease LptF | WP_002209311.1 |
| YPTB0548 | lsrF | | YPTB_RS03105 | YPTB0548 | | 3-hydroxy-5-phosphonooxypentane-2,4-dione thiolase | WP_011191661.1 |
| YPTB0573 | | | YPTB_RS03250 | YPTB0573 | | DNA polymerase III subunit psi | WP_011191680.1 |
| YPTB0574 | rimI | | YPTB_RS03255 | YPTB0574 | | ribosomal protein S18-alanine N-acetyltransferase | WP_002216075.1 |
| YPTB0576 | osmY | | YPTB_RS03265 | YPTB0576 | | molecular chaperone OsmY | WP_011191682.1 |
| YPTB0577 | | | YPTB_RS03270 | YPTB0577 | | DUF1328 domain-containing protein | WP_011191683.1 |
| YPTB0583 | deoB | drm,thyR,tlr | YPTB_RS03300 | YPTB0583 | | phosphopentomutase | WP_011191688.1 |
| YPTB0598 | gpmB | | YPTB_RS03375 | YPTB0598 | | 2,3-diphosphoglycerate-dependent phosphoglycerate mutase GpmB | WP_002209230.1 |
| YPTB0599 | robA | | YPTB_RS03380 | YPTB0599 | | MDR efflux pump AcrAB transcriptional activator RobA | WP_002209231.1 |
| YPTB0603 | thrB | | YPTB_RS03400 | YPTB0603 | | homoserine kinase | WP_002209238.1 |
| YPTB0604 | thrC | | YPTB_RS03405 | YPTB0604 | | threonine synthase | WP_002209239.1 |
| YPTB0616 | ribF | | YPTB_RS03470 | YPTB0616 | | bifunctional riboflavin kinase/FAD synthetase | WP_002210510.1 |
| YPTB0620 | ispH | | YPTB_RS03490 | YPTB0620 | | 4-hydroxy-3-methylbut-2-enyl diphosphate reductase | WP_011191701.1 |
| YPTB0622 | dapB | | YPTB_RS03500 | YPTB0622 | | 4-hydroxy-tetrahydrodipicolinate reductase | WP_002210504.1 |
| YPTB0632 | apaG | corD | YPTB_RS03550 | YPTB0632 | | Co2+/Mg2+ efflux protein ApaG | WP_011191709.1 |
| YPTB0633 | rsmA | | YPTB_RS03555 | YPTB0633 | | 16S rRNA (adenine(1518)-N(6)/adenine(1519)-N(6))-dimethyltransferase RsmA | WP_011191710.1 |
| YPTB0671 | leuB | | YPTB_RS03750 | YPTB0671 | | 3-isopropylmalate dehydrogenase | WP_011191728.1 |
| YPTB0685 | mraY | murX | YPTB_RS03840 | YPTB0685 | | phospho-N-acetylmuramoyl-pentapeptide-transferase | WP_002210437.1 |
| YPTB0690 | | | YPTB_RS03865 | YPTB0690 | | D-alanine--D-alanine ligase | WP_002210432.1 |
| YPTB0692 | ftsA | divA | YPTB_RS03875 | YPTB0692 | | cell division protein FtsA | WP_002210431.1 |
| YPTB0693 | ftsZ | sfiB,sulB | YPTB_RS03880 | YPTB0693 | | cell division protein FtsZ | WP_002210430.1 |
| YPTB0701 | zapD | | YPTB_RS03925 | YPTB0701 | | cell division protein ZapD | WP_002209318.1 |
| YPTB0703 | | | YPTB_RS03940 | YPTB0703 | | GMP reductase | WP_002209320.1 |
| YPTB0712 | pdhR | aceC,genA,yacB | YPTB_RS03985 | YPTB0712 | | pyruvate dehydrogenase complex transcriptional repressor PdhR | WP_002216080.1 |
| YPTB0713 | aceE | | YPTB_RS03990 | YPTB0713 | | pyruvate dehydrogenase (acetyl-transferring), homodimeric type | WP_011191742.1 |
| YPTB0719 | speD | | YPTB_RS04020 | YPTB0719 | | adenosylmethionine decarboxylase | WP_011191745.1 |
| YPTB0723 | hpt | | YPTB_RS04040 | YPTB0723 | | hypoxanthine phosphoribosyltransferase | WP_011191748.1 |
| YPTB0726 | | | YPTB_RS04055 | YPTB0726 | | ABC transporter permease | WP_002209344.1 |
| YPTB0731 | folK | hppK | YPTB_RS04080 | YPTB0731 | | 2-amino-4-hydroxy-6-hydroxymethyldihydropteridine diphosphokinase | WP_002209351.1 |
| YPTB0734 | dksA | msmA | YPTB_RS04095 | YPTB0734 | | RNA polymerase-binding protein DksA | WP_002209354.1 |
| YPTB0744 | erpA | | YPTB_RS04145 | YPTB0744 | | iron-sulfur cluster insertion protein ErpA | WP_002209365.1 |
| YPTB0747 | mtnN | | YPTB_RS04160 | YPTB0747 | | 5'-methylthioadenosine/S-adenosylhomocysteine nucleosidase | WP_011191764.1 |
| YPTB0751 | relA | | YPTB_RS04180 | YPTB0751 | | GTP diphosphokinase | WP_002209373.1 |
| YPTB0765 | cysD | | YPTB_RS04255 | YPTB0765 | | sulfate adenylyltransferase subunit CysD | WP_002209386.1 |
| YPTB0768 | | | YPTB_RS04270 | YPTB0768 | | DUF3561 family protein | WP_002209389.1 |
| YPTB0774 | | | YPTB_RS04300 | YPTB0774 | | protein-L-isoaspartate(D-aspartate) O-methyltransferase | WP_002209395.1 |
| YPTB0833 | ffh | | YPTB_RS04625 | YPTB0833 | | signal recognition particle protein | WP_011191811.1 |
| YPTB0834 | rpsP | | YPTB_RS04630 | YPTB0834 | | 30S ribosomal protein S16 | WP_002209458.1 |
| YPTB0835 | rimM | yfjA | YPTB_RS04635 | YPTB0835 | | ribosome maturation factor RimM | WP_011191812.1 |
| YPTB0842 | tyrA | | YPTB_RS04670 | YPTB0842 | | bifunctional chorismate mutase/prephenate dehydrogenase | WP_002209465.1 |
| YPTB0851 | | | YPTB_RS04750 | YPTB0851 | | bifunctional acetate--CoA ligase family protein/GNAT family N-acetyltransferase | WP_011191815.1 |
| YPTB0857 | mprA | | YPTB_RS04780 | YPTB0857 | | transcriptional repressor MprA | WP_002208744.1 |
| YPTB0858 | ygaH | | YPTB_RS04785 | YPTB0858 | | L-valine transporter subunit YgaH | WP_011191819.1 |
| YPTB0873 | | | YPTB_RS04860 | YPTB0873 | | pyridoxal phosphate-dependent aminotransferase | WP_011191830.1 |
| YPTB0876 | | | YPTB_RS04875 | YPTB0876 | | acireductone dioxygenase | WP_011191833.1 |
| YPTB0887 | | | YPTB_RS04930 | YPTB0887 | | Na(+)-translocating NADH-quinone reductase subunit A | WP_002208717.1 |
| YPTB0889 | | | YPTB_RS04940 | YPTB0889 | | Na(+)-translocating NADH-quinone reductase subunit C | WP_011191841.1 |
| YPTB0890 | | | YPTB_RS04945 | YPTB0890 | | NADH:ubiquinone reductase (Na(+)-transporting) subunit D | WP_002208714.1 |
| YPTB0892 | nqrF | nqr6 | YPTB_RS04955 | YPTB0892 | | NADH:ubiquinone reductase (Na(+)-transporting) subunit F | WP_011191842.1 |

| | | | | | | | |
|---|---|---|---|---|---|---|---|
| YPTB0894 | nqrM | | YPTB_RS04965 | YPTB0894 | | (Na+)-NQR maturation NqrM | WP_002214684.1 |
| YPTB0903 | crl | | YPTB_RS05015 | YPTB0903 | | sigma factor-binding protein Crl | WP_002208702.1 |
| YPTB0920 | brnQ | hrbA | YPTB_RS05110 | YPTB0920 | | branched-chain amino acid transport system II carrier protein | WP_002208680.1 |
| YPTB0925 | | | YPTB_RS05140 | YPTB0925 | | peroxiredoxin C | WP_002208675.1 |
| YPTB0930 | secD | | YPTB_RS05165 | YPTB0930 | | protein translocase subunit SecD | WP_002223272.1 |
| YPTB0933 | nrdR | | YPTB_RS05185 | YPTB0933 | | transcriptional regulator NrdR | WP_002208668.1 |
| YPTB0934 | ribD | ribG,ybaE | YPTB_RS05190 | YPTB0934 | | bifunctional diaminohydroxyphosphoribosylaminopyrimidine deaminase/5-amino-6-(5-phosphoribosylamino)uracil reductase RibD | WP_002208667.1 |
| YPTB0936 | nusB | groNB,ssaD,ssyB | YPTB_RS05200 | YPTB0936 | | transcription antitermination factor NusB | WP_002208665.1 |
| YPTB0950 | | | YPTB_RS05270 | YPTB0950 | | cytochrome o ubiquinol oxidase subunit III | WP_002208651.1 |
| YPTB0951 | cyoB | | YPTB_RS05275 | YPTB0951 | | cytochrome o ubiquinol oxidase subunit I | WP_002208650.1 |
| YPTB0955 | | | YPTB_RS05295 | YPTB0955 | | lipoprotein | WP_002208646.1 |
| YPTB0958 | tig | | YPTB_RS05310 | YPTB0958 | | trigger factor | WP_002208643.1 |
| YPTB0959 | clpP | lopP | YPTB_RS05315 | YPTB0959 | | ATP-dependent Clp endopeptidase proteolytic subunit ClpP | WP_002208642.1 |
| YPTB0960 | clpX | lopC | YPTB_RS05320 | YPTB0960 | | ATP-dependent protease ATP-binding subunit ClpX | WP_011191865.1 |
| YPTB0962 | hupB | dpeA,hopD | YPTB_RS05330 | YPTB0962 | | DNA-binding protein HU-beta | WP_002208639.1 |
| YPTB0963 | ppiD | ybaU | YPTB_RS05335 | YPTB0963 | | peptidylprolyl isomerase | WP_011191866.1 |
| YPTB0966 | queC | | YPTB_RS05350 | YPTB0966 | | 7-cyano-7-deazaguanine synthase QueC | WP_002208635.1 |
| YPTB0970 | | | YPTB_RS05370 | YPTB0970 | | Lrp/AsnC family transcriptional regulator | WP_002208630.1 |
| YPTB0979 | tomB | | YPTB_RS05415 | YPTB0979 | | Hha toxicity modulator TomB | WP_002218472.1 |
| YPTB0981 | ykgO | | YPTB_RS05425 | YPTB0981 | | type B 50S ribosomal protein L36 | WP_002208618.1 |
| YPTB0990 | | | YPTB_RS22480 | YPTB0990 | | DUF454 family protein | WP_002228346.1 |
| YPTB0993 | | | YPTB_RS05485 | YPTB0993 | | YbaB/EbfC family nucleoid-associated protein | WP_002208604.1 |
| YPTB0994 | recR | | YPTB_RS05490 | YPTB0994 | | recombination mediator RecR | WP_002208603.1 |
| YPTB1025 | | | YPTB_RS05655 | YPTB1025 | | SPFH/Band 7/PHB domain protein | WP_002208576.1 |
| YPTB1027 | | | YPTB_RS05665 | YPTB1027 | | SDR family oxidoreductase | WP_002208574.1 |
| YPTB1029 | | | YPTB_RS05675 | YPTB1029 | | ABC transporter ATP-binding protein | WP_011191895.1 |
| YPTB1034 | ppiB | | YPTB_RS05700 | YPTB1034 | | peptidylprolyl isomerase B | WP_002208567.1 |
| YPTB1091 | lipA | lip | YPTB_RS06015 | YPTB1091 | | lipoyl synthase | WP_002210320.1 |
| YPTB1096 | mrdB | | YPTB_RS06040 | YPTB1096 | | peptidoglycan glycosyltransferase MrdB | WP_002210325.1 |
| YPTB1097 | mrdA | | YPTB_RS06045 | YPTB1097 | | peptidoglycan DD-transpeptidase MrdA | WP_011191932.1 |
| YPTB1098 | rlmH | | YPTB_RS06050 | YPTB1098 | | 23S rRNA (pseudouridine(1915)-N(3))-methyltransferase RlmH | WP_002210328.1 |
| YPTB1103 | leuS | | YPTB_RS06075 | YPTB1103 | | leucine--tRNA ligase | WP_002210333.1 |
| YPTB1104 | | | YPTB_RS06080 | YPTB1104 | | zinc ribbon-containing protein | WP_002210335.1 |
| YPTB1105 | | | YPTB_RS06085 | YPTB1105 | | amino acid ABC transporter ATP-binding protein | WP_002210336.1 |
| YPTB1106 | gltK | | YPTB_RS06090 | YPTB1106 | | glutamate/aspartate ABC transporter permease GltK | WP_002210337.1 |
| YPTB1110 | corC | | YPTB_RS06110 | YPTB1110 | | CNNM family magnesium/cobalt transport protein CorC | WP_002210342.1 |
| YPTB1116 | | | YPTB_RS06180 | YPTB1116 | | HAD-IIA family hydrolase | WP_002210349.1 |
| YPTB1119 | nagB | glmD | YPTB_RS06195 | YPTB1119 | | glucosamine-6-phosphate deaminase | WP_002210352.1 |
| YPTB1125 | fldA | | YPTB_RS06225 | YPTB1125 | | flavodoxin FldA | WP_002215386.1 |
| YPTB1135 | smpB | smqB | YPTB_RS06280 | YPTB1135 | | SsrA-binding protein SmpB | WP_002210714.1 |
| YPTB1141 | grpE | | YPTB_RS06310 | YPTB1141 | | nucleotide exchange factor GrpE | WP_002224622.1 |
| YPTB1144 | sdhD | | YPTB_RS06325 | YPTB1144 | | succinate dehydrogenase membrane anchor subunit | WP_002210723.1 |
| YPTB1145 | sdhA | | YPTB_RS06330 | YPTB1145 | | succinate dehydrogenase flavoprotein subunit | WP_002210724.1 |
| YPTB1146 | | | YPTB_RS06335 | YPTB1146 | | succinate dehydrogenase iron-sulfur subunit | WP_002210725.1 |
| YPTB1147 | sucA | | YPTB_RS06340 | YPTB1147 | | 2-oxoglutarate dehydrogenase E1 component | WP_002210726.1 |
| YPTB1149 | sucC | | YPTB_RS06350 | YPTB1149 | | ADP-forming succinate--CoA ligase subunit beta | WP_002210728.1 |
| YPTB1155 | ybgC | | YPTB_RS06380 | YPTB1155 | | tol-pal system-associated acyl-CoA thioesterase | WP_002210734.1 |
| YPTB1157 | tolR | | YPTB_RS06390 | YPTB1157 | | colicin uptake protein TolR | WP_002210736.1 |
| YPTB1159 | tolB | | YPTB_RS06400 | YPTB1159 | | Tol-Pal system beta propeller repeat protein TolB | WP_002210738.1 |
| YPTB1166 | gpmA | gpm | YPTB_RS06470 | YPTB1166 | | 2,3-diphosphoglycerate-dependent phosphoglycerate mutase | WP_002210746.1 |
| YPTB1169 | galK | galA | YPTB_RS06485 | YPTB1169 | | galactokinase | WP_011191950.1 |
| YPTB1172 | | | YPTB_RS06500 | YPTB1172 | | CPBP family intramembrane metalloprotease | WP_002210751.1 |
| YPTB1192 | moaD | chlA4,chlM | YPTB_RS06610 | YPTB1192 | | molybdopterin synthase sulfur carrier subunit | WP_002210773.1 |
| YPTB1209 | | | YPTB_RS06700 | YPTB1209 | | ABC transporter permease | WP_002220162.1 |
| YPTB1212 | hlyD | | YPTB_RS06715 | YPTB1212 | | secretion protein HlyD | WP_011191973.1 |
| YPTB1253 | nrdB | ftsB | YPTB_RS06920 | YPTB1253 | | ribonucleotide-diphosphate reductase subunit beta | WP_002210817.1 |
| YPTB1297 | | | YPTB_RS07190 | YPTB1297 | | YejL family protein | WP_002208836.1 |
| YPTB1309 | mepS | | YPTB_RS07250 | YPTB1309 | | bifunctional murein DD-endopeptidase/murein LD-carboxypeptidase | WP_002208822.1 |
| YPTB1316 | yeiP | | YPTB_RS07285 | YPTB1316 | | elongation factor P-like protein YeiP | WP_002228019.1 |
| YPTB1330 | fruK | fpk | YPTB_RS07355 | YPTB1330 | | 1-phosphofructokinase | WP_002208798.1 |
| YPTB1358 | | | YPTB_RS07505 | YPTB1358 | | GrxA family glutaredoxin | WP_002208765.1 |
| YPTB1361 | potF | | YPTB_RS07525 | YPTB1361 | | spermidine/putrescine ABC transporter substrate-binding protein PotF | WP_002208762.1 |
| YPTB1362 | potG | | YPTB_RS07530 | YPTB1362 | | putrescine ABC transporter ATP-binding subunit PotG | WP_002208760.1 |
| YPTB1376 | artQ | | YPTB_RS07610 | YPTB1376 | | arginine ABC transporter permease ArtQ | WP_002211369.1 |
| YPTB1377 | artJ | | YPTB_RS07615 | YPTB1377 | | arginine ABC transporter substrate-binding protein | WP_002211368.1 |
| YPTB1386 | hcp | | YPTB_RS07660 | YPTB1386 | | hydroxylamine reductase | WP_011192048.1 |
| YPTB1391 | macB | | YPTB_RS07685 | YPTB1391 | | macrolide ABC transporter ATP-binding protein/permease MacB | WP_002211351.1 |
| YPTB1394 | clpA | lopD | YPTB_RS07700 | YPTB1394 | | ATP-dependent Clp protease ATP-binding subunit ClpA | WP_011192049.1 |
| YPTB1395 | infA | | YPTB_RS07705 | YPTB1395 | | translation initiation factor IF-1 | WP_002211347.1 |
| YPTB1397 | cydC | mdrH | YPTB_RS07715 | YPTB1397 | | cysteine/glutathione ABC transporter ATP-binding protein/permease CydC | WP_002211344.1 |
| YPTB1403 | | | YPTB_RS07750 | YPTB1403 | | replication-associated recombination protein A | WP_002228009.1 |
| YPTB1408 | pflB | pfl,YPO1383 | YPTB_RS07775 | YPTB1408 | | formate C-acetyltransferase | WP_002211332.1 |
| YPTB1414 | serC | | YPTB_RS07805 | YPTB1414 | | 3-phosphoserine/phosphohydroxythreonine transaminase | WP_011192055.1 |
| YPTB1416 | cmk | | YPTB_RS07815 | YPTB1416 | | (d)CMP kinase | WP_002211324.1 |
| YPTB1417 | rpsA | ssyF | YPTB_RS07820 | YPTB1417 | | 30S ribosomal protein S1 | WP_011192057.1 |
| YPTB1428 | mukF | kicB | YPTB_RS07885 | YPTB1428 | | chromosome partition protein MukF | WP_002211310.1 |
| YPTB1436 | asnS | | YPTB_RS07925 | YPTB1436 | | asparagine--tRNA ligase | WP_002211301.1 |
| YPTB1439 | pyrD | | YPTB_RS07940 | YPTB1439 | | quinone-dependent dihydroorotate dehydrogenase | WP_002211296.1 |
| YPTB1442 | rlmKL | | YPTB_RS07955 | YPTB1442 | | bifunctional 23S rRNA (guanine(2069)-N(7))-methyltransferase RlmK/23S rRNA (guanine(2445)-N(2))-methyltransferase RlmL | WP_011192066.1 |
| YPTB1445 | pqiB | | YPTB_RS07970 | YPTB1445 | | intermembrane transport protein PqiB | WP_002211290.1 |
| YPTB1454 | sulA | sfiA | YPTB_RS08020 | YPTB1454 | | cell division inhibitor SulA | WP_011192069.1 |
| YPTB1524 | mglC | | YPTB_RS08390 | YPTB1524 | | galactose/methyl galactoside ABC transporter permease MglC | WP_002211965.1 |
| YPTB1531 | | | YPTB_RS08425 | YPTB1531 | | LVIVD repeat-containing protein | WP_011192108.1 |
| YPTB1535 | apbC | | YPTB_RS08445 | YPTB1535 | | iron-sulfur cluster carrier protein ApbC | WP_002211871.1 |
| YPTB1537 | dcd | dus,paxA | YPTB_RS08455 | YPTB1537 | | dCTP deaminase | WP_002211873.1 |
| YPTB1555 | | | YPTB_RS08550 | YPTB1555 | | bifunctional phosphoribosyl-AMP cyclohydrolase/phosphoribosyl-ATP diphosphatase HisIE | WP_002211889.1 |
| YPTB1557 | hisA | | YPTB_RS08560 | YPTB1557 | | 1-(5-phosphoribosyl)-5-[(5-phosphoribosylamino)methylideneamino]imidazole-4-carboxamide isomerase | WP_002211891.1 |
| YPTB1558 | hisH | | YPTB_RS08565 | YPTB1558 | | imidazole glycerol phosphate synthase subunit HisH | WP_002211892.1 |
| YPTB1559 | hisB | | YPTB_RS08570 | YPTB1559 | | bifunctional histidinol-phosphatase/imidazoleglycerol-phosphate dehydratase HisB | WP_011192120.1 |
| YPTB1615 | | | YPTB_RS08905 | YPTB1615 | | helix-turn-helix domain-containing protein | WP_002224436.1 |
| YPTB1647 | | | YPTB_RS09075 | YPTB1647 | | L-serine ammonia-lyase | WP_011192152.1 |
| YPTB1724 | putP | | YPTB_RS09480 | YPTB1724 | | sodium/proline symporter PutP | WP_002211164.1 |
| YPTB1795 | rdgC | yaiD | | YPTB1795 | | possible recombination associated protein RdgC | CAH21034.1 |
| YPTB1921 | | | YPTB_RS10550 | YPTB1921 | | type 1 fimbrial protein | WP_002213971.1 |
| YPTB1940 | | | YPTB_RS10650 | YPTB1940 | | iron transporter | WP_002212053.1 |
| YPTB2001 | prs | | YPTB_RS10970 | YPTB2001 | | ribose-phosphate diphosphokinase | WP_002211240.1 |
| YPTB2005 | prfA | sueB,uar | YPTB_RS10995 | YPTB2005 | | peptide chain release factor 1 | WP_011192398.1 |
| YPTB2007 | | | YPTB_RS11005 | YPTB2007 | | SirB2 family protein | WP_002211234.1 |
| YPTB2008 | | | YPTB_RS11010 | YPTB2008 | | invasion regulator SirB1 | WP_002211233.1 |
| YPTB2009 | kdsA | | YPTB_RS11015 | YPTB2009 | | 3-deoxy-8-phosphooctulonate synthase | WP_002211232.1 |
| YPTB2019 | | | YPTB_RS11080 | YPTB2019 | | phosphatase | WP_002211221.1 |
| YPTB2023 | rimJ | | YPTB_RS11100 | YPTB2023 | | ribosomal protein S5-alanine N-acetyltransferase | WP_002211216.1 |
| YPTB2032 | cmoB | | YPTB_RS11145 | YPTB2032 | | tRNA 5-methoxyuridine(34)/uridine 5-oxyacetic acid(34) synthase CmoB | WP_011192411.1 |
| YPTB2038 | | | YPTB_RS11175 | YPTB2038 | | YebC/PmpR family DNA-binding transcriptional regulator | WP_002211202.1 |
| YPTB2042 | znuB | | YPTB_RS11195 | YPTB2042 | | zinc ABC transporter permease subunit ZnuB | WP_002211197.1 |
| YPTB2045 | mepM | | YPTB_RS11210 | YPTB2045 | | murein DD-endopeptidase MepM | WP_002228437.1 |
| YPTB2047 | pyk | | YPTB_RS11220 | YPTB2047 | | pyruvate kinase | WP_002211193.1 |
| YPTB2053 | | | YPTB_RS11255 | YPTB2053 | | RidA family protein | WP_002211186.1 |
| YPTB2059 | minE | | YPTB_RS11285 | YPTB2059 | | cell division topological specificity factor MinE | WP_002211180.1 |
| YPTB2060 | minD | | YPTB_RS11290 | YPTB2060 | | septum site-determining protein MinD | WP_002211179.1 |
| YPTB2064 | | | YPTB_RS11310 | YPTB2064 | | fumarylacetoacetate hydrolase family protein | WP_002211740.1 |
| YPTB2083 | gapA | | YPTB_RS11405 | YPTB2083 | | glyceraldehyde-3-phosphate dehydrogenase | WP_002224141.1 |
| YPTB2084 | msrB | | YPTB_RS11410 | YPTB2084 | | peptide-methionine (R)-S-oxide reductase MsrB | WP_002211677.1 |
| YPTB2089 | | | YPTB_RS11435 | YPTB2089 | | NAD(P)H nitroreductase | WP_002211673.1 |
| YPTB2090 | selD | fdhB | YPTB_RS11440 | YPTB2090 | | selenide, water dikinase SelD | WP_011192432.1 |
| YPTB2116 | | | YPTB_RS11575 | YPTB2116 | | YciI family protein | WP_002210643.1 |
| YPTB2125 | trpA | | YPTB_RS11620 | YPTB2125 | | tryptophan synthase subunit alpha | WP_011192442.1 |
| YPTB2126 | trpB | | YPTB_RS11625 | YPTB2126 | | tryptophan synthase subunit beta | WP_002210633.1 |
| YPTB2137 | | | YPTB_RS11680 | YPTB2137 | | YciK family oxidoreductase | WP_002210625.1 |
| YPTB2139 | | | YPTB_RS11690 | YPTB2139 | | YciN family protein | WP_002210623.1 |
| YPTB2144 | ribA | | YPTB_RS11720 | YPTB2144 | | GTP cyclohydrolase II | WP_002227926.1 |
| YPTB2146 | | | YPTB_RS11730 | YPTB2146 | | LapA family protein | WP_002210616.1 |
| YPTB2147 | lapB | | YPTB_RS11735 | YPTB2147 | | lipopolysaccharide assembly protein LapB | WP_011192449.1 |
| YPTB2156 | | | YPTB_RS11780 | YPTB2156 | | carbon starvation protein A | WP_011192451.1 |
| YPTB2160 | nth | | YPTB_RS11800 | YPTB2160 | | endonuclease III | WP_002210602.1 |
| YPTB2167 | rsxA | | YPTB_RS11835 | YPTB2167 | | electron transport complex subunit RsxA | WP_002210595.1 |
| YPTB2261 | tpx | | YPTB_RS12325 | YPTB2261 | | thiol peroxidase | WP_002210984.1 |
| YPTB2263 | | | YPTB_RS12335 | YPTB2263 | | DUF2384 domain-containing protein | WP_002210982.1 |
| YPTB2266 | | | YPTB_RS12350 | YPTB2266 | | YcjX family protein | WP_002210979.1 |
| YPTB2273 | sapB | | YPTB_RS12390 | YPTB2273 | | putrescine ABC transporter permease SapB | WP_002210972.1 |

| | | | | | | | |
|---|---|---|---|---|---|---|---|
| YPTB2274 | sapC | | YPTB_RS12395 | YPTB2274 | | peptide ABC transporter permease SapC | WP_002210971.1 |
| YPTB2275 | sapD | | YPTB_RS12400 | YPTB2275 | | peptide ABC transporter ATP-binding protein SapD | WP_002210970.1 |
| YPTB2283 | tyrS | | YPTB_RS12440 | YPTB2283 | | tyrosine--tRNA ligase | WP_011192524.1 |
| YPTB2312 | sufC | | YPTB_RS12605 | YPTB2312 | | Fe-S cluster assembly ATPase SufC | WP_011192535.1 |
| YPTB2322 | | | YPTB_RS12660 | YPTB2322 | | lipoate--protein ligase A | WP_011192540.1 |
| YPTB2323 | | | YPTB_RS12665 | YPTB2323 | | lipoprotein | WP_002216102.1 |
| YPTB2324 | arnF | | YPTB_RS12670 | YPTB2324 | | 4-amino-4-deoxy-L-arabinose-phosphoundecaprenol flippase subunit ArnF | WP_002211819.1 |
| YPTB2336 | pheT | | YPTB_RS12730 | YPTB2336 | | phenylalanine--tRNA ligase subunit beta | WP_011192545.1 |
| YPTB2337 | pheS | | YPTB_RS12735 | YPTB2337 | | phenylalanine--tRNA ligase subunit alpha | WP_011192546.1 |
| YPTB2339 | rplT | pdzA | YPTB_RS12740 | YPTB2339 | | 50S ribosomal protein L20 | WP_002211833.1 |
| YPTB2340 | rpmI | | YPTB_RS12745 | YPTB2340 | | 50S ribosomal protein L35 | WP_002211834.1 |
| YPTB2341 | infC | fit | YPTB_RS22885 | YPTB2341 | | translation initiation factor IF-3 | WP_002227898.1 |
| YPTB2353 | | | YPTB_RS12815 | YPTB2353 | | YniB family protein | WP_002211847.1 |
| YPTB2354 | hxpB | | YPTB_RS12820 | YPTB2354 | | hexitol phosphatase HxpB | WP_002220277.1 |
| YPTB2389 | | | YPTB_RS12995 | YPTB2389 | | hypothetical protein | WP_002210862.1 |
| YPTB2396 | cheZ | | YPTB_RS13040 | YPTB2396 | | protein phosphatase CheZ | WP_002210872.1 |
| YPTB2397 | cheY | | YPTB_RS13045 | YPTB2397 | | chemotaxis response regulator CheY | WP_002210873.1 |
| YPTB2427 | icd | | YPTB_RS13205 | YPTB2427 | | NADP-dependent isocitrate dehydrogenase | WP_002210910.1 |
| YPTB2432 | purB | | YPTB_RS13230 | YPTB2432 | | adenylosuccinate lyase | WP_002230790.1 |
| YPTB2442 | lolC | | YPTB_RS13280 | YPTB2442 | | lipoprotein-releasing ABC transporter permease subunit LolC | WP_011192589.1 |
| YPTB2447 | | | YPTB_RS13305 | YPTB2447 | | NAD(P)/FAD-dependent oxidoreductase | WP_002213097.1 |
| YPTB2448 | | | YPTB_RS13310 | YPTB2448 | | alpha/beta hydrolase YcfP | WP_002213095.1 |
| YPTB2467 | mltG | | YPTB_RS13410 | YPTB2467 | | endolytic transglycosylase MltG | WP_011192605.1 |
| YPTB2470 | acpP | | YPTB_RS13425 | YPTB2470 | | acyl carrier protein | WP_002220787.1 |
| YPTB2471 | fabG | | YPTB_RS13430 | YPTB2471 | | 3-oxoacyl-ACP reductase FabG | WP_002210935.1 |
| YPTB2474 | plsX | | YPTB_RS13445 | YPTB2474 | | phosphate acyltransferase PlsX | WP_002210932.1 |
| YPTB2475 | rpmF | | YPTB_RS13450 | YPTB2475 | | 50S ribosomal protein L32 | WP_002210931.1 |
| YPTB2549 | glnP | | YPTB_RS13840 | YPTB2549 | | glutamine ABC transporter permease GlnP | WP_002210236.1 |
| YPTB2557 | menC | | YPTB_RS13875 | YPTB2557 | | o-succinylbenzoate synthase | WP_011192654.1 |
| YPTB2558 | menB | | YPTB_RS13880 | YPTB2558 | | 1,4-dihydroxy-2-naphthoyl-CoA synthase | WP_002210245.1 |
| YPTB2577 | nuoL | | YPTB_RS13985 | YPTB2577 | | NADH-quinone oxidoreductase subunit L | WP_011192667.1 |
| YPTB2578 | nuoK | | YPTB_RS13990 | YPTB2578 | | NADH-quinone oxidoreductase subunit NuoK | WP_002210271.1 |
| YPTB2580 | nuoI | | YPTB_RS14000 | YPTB2580 | | NADH-quinone oxidoreductase subunit NuoI | WP_002210273.1 |
| YPTB2581 | nuoH | | YPTB_RS14005 | YPTB2581 | | NADH-quinone oxidoreductase subunit NuoH | WP_002210274.1 |
| YPTB2585 | nuoC | | YPTB_RS14025 | YPTB2585 | | NADH-quinone oxidoreductase subunit C/D | WP_002210277.1 |
| YPTB2594 | | | YPTB_RS14075 | YPTB2594 | | YfbU family protein | WP_002210286.1 |
| YPTB2595 | | | YPTB_RS14080 | YPTB2595 | | DUF412 domain-containing protein | WP_011192676.1 |
| YPTB2597 | ackA | ack | YPTB_RS14090 | YPTB2597 | | acetate kinase | WP_011192678.1 |
| YPTB2600 | | | YPTB_RS14105 | YPTB2600 | | PTS sugar transporter subunit IIA | WP_011192680.1 |
| YPTB2601 | | | YPTB_RS14110 | YPTB2601 | | PTS sugar transporter subunit IIB | WP_002264679.1 |
| YPTB2602 | | | YPTB_RS14115 | YPTB2602 | | PTS ascorbate transporter subunit IIC | WP_002222248.1 |
| YPTB2609 | | | YPTB_RS14150 | YPTB2609 | | histidine ABC transporter permease HisQ | WP_002209736.1 |
| YPTB2612 | purF | | YPTB_RS14165 | YPTB2612 | | amidophosphoribosyltransferase | WP_002209733.1 |
| YPTB2613 | cvpA | dedE | YPTB_RS14170 | YPTB2613 | | colicin V production protein | WP_002209732.1 |
| YPTB2617 | | | YPTB_RS14190 | YPTB2617 | | DedA family protein | WP_002209728.1 |
| YPTB2630 | | | YPTB_RS14260 | YPTB2630 | | sulfite exporter TauE/SafE family protein | WP_002209713.1 |
| YPTB2632 | aroC | | YPTB_RS14270 | YPTB2632 | | chorismate synthase | WP_011192692.1 |
| YPTB2637 | fadI | flrD,fre,fsrC,ubiB | YPTB_RS14295 | YPTB2637 | | acetyl-CoA C-acyltransferase FadI | WP_002209704.1 |
| YPTB2640 | mlaA | | YPTB_RS14310 | YPTB2640 | | phospholipid-binding lipoprotein MlaA | WP_002209701.1 |
| YPTB2714 | cysK | cysZ | YPTB_RS14720 | YPTB2714 | | cysteine synthase A | WP_011192736.1 |
| YPTB2715 | ptsH | hpr | YPTB_RS14725 | YPTB2715 | | phosphocarrier protein Hpr | WP_002208488.1 |
| YPTB2716 | ptsI | | YPTB_RS14730 | YPTB2716 | | phosphoenolpyruvate-protein phosphotransferase PtsI | WP_002208490.1 |
| YPTB2761 | napD | | YPTB_RS14965 | YPTB2761 | | chaperone NapD | WP_002208535.1 |
| YPTB2775 | dapE | msgB | YPTB_RS15035 | YPTB2775 | | succinyl-diaminopimelate desuccinylase | WP_002208549.1 |
| YPTB2779 | | | YPTB_RS15055 | YPTB2779 | | DUF441 domain-containing protein | WP_002208553.1 |
| YPTB2781 | purC | | YPTB_RS15065 | YPTB2781 | | phosphoribosylaminoimidazolesuccinocarboxamide synthase | WP_002208555.1 |
| YPTB2785 | bcp | | YPTB_RS15085 | YPTB2785 | | thioredoxin-dependent thiol peroxidase | WP_011192766.1 |
| YPTB2789 | | | YPTB_RS15105 | YPTB2789 | | AI-2E family transporter | WP_002208563.1 |
| YPTB2794 | upp | uraP | YPTB_RS15130 | YPTB2794 | | uracil phosphoribosyltransferase | WP_002209776.1 |
| YPTB2797 | speG | | YPTB_RS15145 | YPTB2797 | | spermidine N1-acetyltransferase | WP_002209779.1 |
| YPTB2799 | pstB | phoT | YPTB_RS15155 | YPTB2799 | | phosphate ABC transporter ATP-binding protein PstB | WP_002209780.1 |
| YPTB2819 | | | YPTB_RS15255 | YPTB2819 | | YegP family protein | WP_002209797.1 |
| YPTB2838 | bamB | | YPTB_RS15360 | YPTB2838 | | outer membrane protein assembly factor BamB | WP_011192801.1 |
| YPTB2853 | iscX | | YPTB_RS15435 | YPTB2853 | | Fe-S cluster assembly protein IscX | WP_002209830.1 |
| YPTB2857 | iscA | | YPTB_RS15455 | YPTB2857 | | iron-sulfur cluster assembly protein IscA | WP_002209834.1 |
| YPTB2859 | | | YPTB_RS15465 | YPTB2859 | | IscS subfamily cysteine desulfurase | WP_011192814.1 |
| YPTB2869 | | | YPTB_RS15515 | YPTB2869 | | serine hydroxymethyltransferase | WP_002211552.1 |
| YPTB2872 | glnB | | YPTB_RS15530 | YPTB2872 | | nitrogen regulatory protein P-II | WP_002231018.1 |
| YPTB2873 | | | YPTB_RS15535 | YPTB2873 | | NAD+ synthase | WP_002211555.1 |
| YPTB2885 | | | YPTB_RS15595 | YPTB2885 | | YfhL family 4Fe-4S dicluster ferredoxin | WP_002211567.1 |
| YPTB2886 | | | YPTB_RS15600 | YPTB2886 | | holo-ACP synthase | WP_011192825.1 |
| YPTB2887 | pdxJ | | YPTB_RS15605 | YPTB2887 | | pyridoxine 5'-phosphate synthase | WP_011192826.1 |
| YPTB2889 | era | rbaA | YPTB_RS15615 | YPTB2889 | | GTPase Era | WP_002214829.1 |
| YPTB2890 | rnc | | YPTB_RS15620 | YPTB2890 | | ribonuclease III | WP_002209679.1 |
| YPTB2895 | rseB | | YPTB_RS15645 | YPTB2895 | | sigma-E factor regulatory protein RseB | WP_002209674.1 |
| YPTB2897 | rpoE | sigE | YPTB_RS15655 | YPTB2897 | | RNA polymerase sigma factor RpoE | WP_002209672.1 |
| YPTB2915 | | | YPTB_RS15750 | YPTB2915 | | YbfA family protein | WP_002209653.1 |
| YPTB2929 | chbA | | YPTB_RS15840 | YPTB2929 | | PTS N,N'-diacetylchitobiose transporter subunit IIA | WP_002212242.1 |
| YPTB2940 | | | YPTB_RS15905 | YPTB2940 | | urease accessory protein UreF | WP_011192843.1 |
| YPTB2942 | | | YPTB_RS15915 | YPTB2942 | | urease subunit alpha | WP_002212229.1 |
| YPTB2965 | rnhA | dasF,herA,rnh,sdrA | YPTB_RS16035 | YPTB2965 | | ribonuclease HI | WP_002210699.1 |
| YPTB2966 | | | YPTB_RS16040 | YPTB2966 | | class I SAM-dependent methyltransferase | WP_002210698.1 |
| YPTB2967 | gloB | | YPTB_RS16045 | YPTB2967 | | hydroxyacylglutathione hydrolase | WP_011192852.1 |
| YPTB2973 | metN | | YPTB_RS16105 | YPTB2973 | | methionine ABC transporter ATP-binding protein MetN | WP_011192854.1 |
| YPTB2974 | metI | | YPTB_RS16110 | YPTB2974 | | methionine ABC transporter permease MetI | WP_002212160.1 |
| YPTB2975 | | | YPTB_RS16115 | YPTB2975 | | MetQ/NlpA family lipoprotein | WP_011192855.1 |
| YPTB2977 | tsaA | | YPTB_RS16125 | YPTB2977 | | tRNA (N6-threonylcarbamoyladenosine(37)-N6)-methyltransferase TrmO | WP_002212157.1 |
| YPTB2978 | proS | drpA | YPTB_RS16130 | YPTB2978 | | proline--tRNA ligase | WP_011192856.1 |
| YPTB2987 | accA | | YPTB_RS16175 | YPTB2987 | | acetyl-CoA carboxylase carboxyl transferase subunit alpha | WP_002212147.1 |
| YPTB2993 | lpxD | firA,omsA | YPTB_RS16205 | YPTB2993 | | UDP-3-O-(3-hydroxymyristoyl)glucosamine N-acyltransferase | WP_002212141.1 |
| YPTB2994 | skp | | YPTB_RS16210 | YPTB2994 | | molecular chaperone Skp | WP_002212140.1 |
| YPTB2997 | cdsA | cds | YPTB_RS16225 | YPTB2997 | | phosphatidate cytidylyltransferase | WP_011192860.1 |
| YPTB2998 | ispU | | YPTB_RS16230 | YPTB2998 | | (2E,6E)-farnesyl-diphosphate-specific ditrans,polycis-undecaprenyl-diphosphate synthase | WP_002212136.1 |
| YPTB3003 | rpsB | | YPTB_RS16255 | YPTB3003 | | 30S ribosomal protein S2 | WP_002221800.1 |
| YPTB3012 | queF | | YPTB_RS16305 | YPTB3012 | | NADPH-dependent 7-cyano-7-deazaguanine reductase QueF | WP_011192866.1 |
| YPTB3017 | | | YPTB_RS16330 | YPTB3017 | | transcriptional regulator GcvA | WP_002212117.1 |
| YPTB3023 | argA | | YPTB_RS16375 | YPTB3023 | | amino-acid N-acetyltransferase | WP_002211624.1 |
| YPTB3033 | thyA | | YPTB_RS16425 | YPTB3033 | | thymidylate synthase | WP_011192873.1 |
| YPTB3034 | lgt | umpA | YPTB_RS16430 | YPTB3034 | | prolipoprotein diacylglyceryl transferase | WP_002211383.1 |
| YPTB3035 | ptsP | | YPTB_RS16435 | YPTB3035 | | phosphoenolpyruvate--protein phosphotransferase | WP_011192874.1 |
| YPTB3036 | rppH | | YPTB_RS16440 | YPTB3036 | | RNA pyrophosphohydrolase | WP_002211381.1 |
| YPTB3042 | aas | | YPTB_RS16480 | YPTB3042 | | bifunctional acyl-ACP--phospholipid O-acyltransferase/long-chain-fatty-acid--ACP ligase | WP_011192876.1 |
| YPTB3165 | recJ | | YPTB_RS17150 | YPTB3165 | | single-stranded-DNA-specific exonuclease RecJ | WP_002209931.1 |
| YPTB3167 | xerD | | YPTB_RS17160 | YPTB3167 | | site-specific tyrosine recombinase XerD | WP_002209933.1 |
| YPTB3181 | gcvH | | YPTB_RS17235 | YPTB3181 | | glycine cleavage system protein GcvH | WP_002209948.1 |
| YPTB3182 | gcvT | | YPTB_RS17240 | YPTB3182 | | glycine cleavage system aminomethyltransferase GcvT | WP_011192962.1 |
| YPTB3196 | pgk | | YPTB_RS17315 | YPTB3196 | | phosphoglycerate kinase | WP_002209963.1 |
| YPTB3197 | epd | gapB | YPTB_RS17320 | YPTB3197 | | erythrose-4-phosphate dehydrogenase | WP_002209964.1 |
| YPTB3198 | tkt | | YPTB_RS17325 | YPTB3198 | | transketolase | WP_011192968.1 |
| YPTB3205 | endA | nucM | YPTB_RS17360 | YPTB3205 | | deoxyribonuclease I | WP_002209973.1 |
| YPTB3210 | aguB | | YPTB_RS17390 | YPTB3210 | | N-carbamoylputrescine amidase | WP_002209979.1 |
| YPTB3215 | | | YPTB_RS17415 | YPTB3215 | | YggT family protein | WP_011192975.1 |
| YPTB3217 | | | YPTB_RS17425 | YPTB3217 | | XTP/dITP diphosphatase | WP_011192977.1 |
| YPTB3218 | hemW | | YPTB_RS17430 | YPTB3218 | | radical SAM family heme chaperone HemW | WP_002209987.1 |
| YPTB3222 | | | YPTB_RS17450 | YPTB3222 | | YggL family protein | WP_002209990.1 |
| YPTB3225 | | | YPTB_RS17465 | YPTB3225 | | oxidative damage protection protein | WP_002230648.1 |
| YPTB3230 | mglA | rbsA | | YPTB3230 | | ABC sugar transporter, fused ATP-binding domains | CAH22468.1 |
| YPTB3313 | slyB | pcpY | | YPTB3313 | | Putative outer membrane lipoprotein Pcp precursor | CAH22551.1 |
| YPTB3319 | motA | | | YPTB3319 | | putative flagellar motor transmembrane channel protein | CAH22557.1 |
| YPTB3320 | fliA | lafS | | YPTB3320 | | Putative RNA polymerase sigma factor for flagellar operon | CAH22558.1 |
| YPTB3339 | flgC | lfgC | | YPTB3339 | | Putative flagellar basal-body rod protein. | CAH22577.1 |
| YPTB3340 | flgB | lfgB | | YPTB3340 | | putative flagellar basal-body rod protein | CAH22578.1 |
| YPTB3347 | fliG | fla | | YPTB3347 | | puative flagellar motor switch protein | CAH22585.1 |
| YPTB3352 | fliN | mopA | | YPTB3352 | | Flagellar switch protein | CAH22590.1 |
| YPTB3354 | fliQ | mopD | | YPTB3354 | | Putative flagellar assembly/export protein, fliQ | CAH22592.1 |
| YPTB3357 | flhA | fhiA | | YPTB3357 | | Putative flagellar biosynthesis/export membrane protein flhA, fhiA. | CAH22595.1 |
| YPTB3395 | parE | nfxD | YPTB_RS18350 | YPTB3395 | | DNA topoisomerase IV subunit B | WP_011193095.1 |
| YPTB3396 | yqiA | | YPTB_RS18355 | YPTB3396 | | esterase YqiA | WP_011193096.1 |
| YPTB3398 | | | YPTB_RS18365 | YPTB3398 | | DUF1249 family protein | WP_002212183.1 |
| YPTB3407 | hldE | | YPTB_RS18420 | YPTB3407 | | bifunctional D-glycero-beta-D-manno-heptose-7-phosphate kinase/D-glycero-beta-D-manno-heptose 1-phosphate adenylyltransferase HldE | WP_011193099.1 |

| | | | | | | | |
|---|---|---|---|---|---|---|---|
| YPTB3413 | folB | | YPTB_RS18450 | YPTB3413 | | bifunctional dihydroneopterin aldolase/7,8-dihydroneopterin epimerase | WP_002212199.1 |
| YPTB3416 | rpsU | | YPTB_RS18465 | YPTB3416 | | 30S ribosomal protein S21 | WP_001144069.1 |
| YPTB3476 | | | YPTB_RS18815 | YPTB3476 | | altronate dehydratase family protein | WP_011193131.1 |
| YPTB3478 | uxaC | | YPTB_RS18825 | YPTB3478 | | glucuronate isomerase | WP_002210410.1 |
| YPTB3479 | | | YPTB_RS18835 | YPTB3479 | | MFS transporter | WP_002353699.1 |
| YPTB3485 | | | YPTB_RS18870 | YPTB3485 | | YqjD family protein | WP_002210418.1 |
| YPTB3488 | | | YPTB_RS18885 | YPTB3488 | | DoxX family protein | WP_002210421.1 |
| YPTB3491 | | | YPTB_RS18900 | YPTB3491 | | pirin family protein | WP_011193137.1 |
| YPTB3498 | elbB | elb2 | YPTB_RS18935 | YPTB3498 | | isoprenoid biosynthesis glyoxalase ElbB | WP_002210143.1 |
| YPTB3503 | | | YPTB_RS18965 | YPTB3503 | | glutamate synthase small subunit | WP_002210138.1 |
| YPTB3506 | sspA | pog,ssp | YPTB_RS18980 | YPTB3506 | | stringent starvation protein A | WP_002210135.1 |
| YPTB3508 | rplM | | YPTB_RS18995 | YPTB3508 | | 50S ribosomal protein L13 | WP_002210132.1 |
| YPTB3510 | | | YPTB_RS19005 | YPTB3510 | | DUF1043 family protein | WP_002210130.1 |
| YPTB3518 | mlaE | | YPTB_RS19045 | YPTB3518 | | lipid asymmetry maintenance ABC transporter permease subunit MlaE | WP_002210122.1 |
| YPTB3523 | lptC | | YPTB_RS19070 | YPTB3523 | | LPS export ABC transporter periplasmic protein LptC | WP_011193147.1 |
| YPTB3524 | lptA | | YPTB_RS19075 | YPTB3524 | | lipopolysaccharide ABC transporter substrate-binding protein LptA | WP_002210117.1 |
| YPTB3525 | lptB | | YPTB_RS19080 | YPTB3525 | | LPS export ABC transporter ATP-binding protein | WP_002210116.1 |
| YPTB3526 | rpoN | glnF,ntrA | YPTB_RS19085 | YPTB3526 | | RNA polymerase factor sigma-54 | WP_002210115.1 |
| YPTB3527 | hpf | | YPTB_RS19090 | YPTB3527 | | ribosome hibernation promoting factor | WP_004392031.1 |
| YPTB3528 | ptsN | rpoP | YPTB_RS19095 | YPTB3528 | | PTS IIA-like nitrogen regulatory protein PtsN | WP_002213952.1 |
| YPTB3531 | pyrB | | YPTB_RS19110 | YPTB3531 | | aspartate carbamoyltransferase | WP_011193148.1 |
| YPTB3533 | ridA | | YPTB_RS19120 | YPTB3533 | | 2-iminobutanoate/2-iminopropanoate deaminase | WP_011193150.1 |
| YPTB3536 | treB | | YPTB_RS19135 | YPTB3536 | | PTS trehalose transporter subunit IIBC | WP_011193151.1 |
| YPTB3538 | rnk | | YPTB_RS19145 | YPTB3538 | | nucleoside diphosphate kinase regulator | WP_002210103.1 |
| YPTB3547 | aaeA | | YPTB_RS19190 | YPTB3547 | | p-hydroxybenzoic acid efflux pump subunit AaeA | WP_002210094.1 |
| YPTB3548 | | | YPTB_RS19195 | YPTB3548 | | AaeX family protein | WP_002210093.1 |
| YPTB3549 | aaeR | | YPTB_RS19200 | YPTB3549 | | HTH-type transcriptional activator AaeR | WP_011193155.1 |
| YPTB3558 | tldD | | YPTB_RS19245 | YPTB3558 | | metalloprotease TldD | WP_002210082.1 |
| YPTB3561 | rng | | YPTB_RS19260 | YPTB3561 | | ribonuclease G | WP_002210079.1 |
| YPTB3563 | mreD | | YPTB_RS19270 | YPTB3563 | | rod shape-determining protein MreD | WP_002210077.1 |
| YPTB3565 | mreB | envB,rodY | YPTB_RS19280 | YPTB3565 | | rod shape-determining protein MreB | WP_002228205.1 |
| YPTB3570 | aroQ | | YPTB_RS19305 | YPTB3570 | | type II 3-dehydroquinate dehydratase | WP_002210071.1 |
| YPTB3574 | panF | | YPTB_RS19325 | YPTB3574 | | sodium/pantothenate symporter | WP_002210065.1 |
| YPTB3649 | pgi | | YPTB_RS19725 | YPTB3649 | | glucose-6-phosphate isomerase | WP_002212085.1 |
| YPTB3660 | aroE | | YPTB_RS19825 | YPTB3660 | | shikimate dehydrogenase | WP_002209026.1 |
| YPTB3663 | | | YPTB_RS19840 | YPTB3663 | | DUF494 family protein | WP_002209023.1 |
| YPTB3665 | def | fms | YPTB_RS19850 | YPTB3665 | | peptide deformylase | WP_002209021.1 |
| YPTB3668 | trkA | | YPTB_RS19865 | YPTB3668 | | Trk system potassium transporter TrkA | WP_002209018.1 |
| YPTB3674 | rpsD | ramA | YPTB_RS19895 | YPTB3674 | | 30S ribosomal protein S4 | WP_002218949.1 |
| YPTB3675 | rpsK | | YPTB_RS19900 | YPTB3675 | | 30S ribosomal protein S11 | WP_002218948.1 |
| YPTB3676 | rpsM | | YPTB_RS19905 | YPTB3676 | | 30S ribosomal protein S13 | WP_002213346.1 |
| YPTB3677 | rpmJ | | YPTB_RS19910 | YPTB3677 | | 50S ribosomal protein L36 | WP_002227352.1 |
| YPTB3678 | secY | prlA | YPTB_RS19915 | YPTB3678 | | preprotein translocase subunit SecY | WP_002213344.1 |
| YPTB3679 | rplO | | YPTB_RS19920 | YPTB3679 | | 50S ribosomal protein L15 | WP_002213341.1 |
| YPTB3680 | rpmD | | YPTB_RS19925 | YPTB3680 | | 50S ribosomal protein L30 | WP_002213339.1 |
| YPTB3681 | rpsE | | YPTB_RS19930 | YPTB3681 | | 30S ribosomal protein S5 | WP_002213337.1 |
| YPTB3682 | rplR | | YPTB_RS19935 | YPTB3682 | | 50S ribosomal protein L18 | WP_002213336.1 |
| YPTB3683 | rplF | | YPTB_RS19940 | YPTB3683 | | 50S ribosomal protein L6 | WP_002213334.1 |
| YPTB3684 | rpsH | | YPTB_RS19945 | YPTB3684 | | 30S ribosomal protein S8 | WP_002213332.1 |
| YPTB3685 | rpsN | | YPTB_RS19950 | YPTB3685 | | 30S ribosomal protein S14 | WP_002213330.1 |
| YPTB3686 | rplE | | YPTB_RS19955 | YPTB3686 | | 50S ribosomal protein L5 | WP_002213329.1 |
| YPTB3687 | rplX | | YPTB_RS19960 | YPTB3687 | | 50S ribosomal protein L24 | WP_002213327.1 |
| YPTB3688 | rplN | | YPTB_RS19965 | YPTB3688 | | 50S ribosomal protein L14 | WP_002213325.1 |
| YPTB3690 | rpmC | | YPTB_RS19975 | YPTB3690 | | 50S ribosomal protein L29 | WP_002218942.1 |
| YPTB3691 | rplP | | YPTB_RS19980 | YPTB3691 | | 50S ribosomal protein L16 | WP_002218940.1 |
| YPTB3692 | rpsC | | YPTB_RS19985 | YPTB3692 | | 30S ribosomal protein S3 | WP_002221644.1 |
| YPTB3693 | rplV | eryB | YPTB_RS19990 | YPTB3693 | | 50S ribosomal protein L22 | WP_002223844.1 |
| YPTB3694 | rpsS | | YPTB_RS19995 | YPTB3694 | | 30S ribosomal protein S19 | WP_002213430.1 |
| YPTB3695 | rplB | | YPTB_RS20000 | YPTB3695 | | 50S ribosomal protein L2 | WP_002213425.1 |
| YPTB3696 | rplW | | YPTB_RS20005 | YPTB3696 | | 50S ribosomal protein L23 | WP_002213423.1 |
| YPTB3697 | rplD | eryA | YPTB_RS20010 | YPTB3697 | | 50S ribosomal protein L4 | WP_002218934.1 |
| YPTB3698 | rplC | | YPTB_RS20015 | YPTB3698 | | 50S ribosomal protein L3 | WP_002218932.1 |
| YPTB3699 | rpsJ | | YPTB_RS20020 | YPTB3699 | | 30S ribosomal protein S10 | WP_001181005.1 |
| YPTB3704 | rpsG | | YPTB_RS20045 | YPTB3704 | | 30S ribosomal protein S7 | WP_002212324.1 |
| YPTB3705 | rpsL | | YPTB_RS20050 | YPTB3705 | | 30S ribosomal protein S12 | WP_002212323.1 |
| YPTB3706 | tusB | | YPTB_RS20055 | YPTB3706 | | sulfurtransferase complex subunit TusB | WP_002212322.1 |
| YPTB3709 | | | YPTB_RS20070 | YPTB3709 | | transcriptional regulator | WP_002212319.1 |
| YPTB3711 | | | YPTB_RS20080 | YPTB3711 | | protein SlyX | WP_002212317.1 |
| YPTB3714 | kefB | trkB | YPTB_RS20095 | YPTB3714 | | glutathione-regulated potassium-efflux system protein KefB | WP_002212314.1 |
| YPTB3725 | | | YPTB_RS20150 | YPTB3725 | | YheU family protein | WP_011193222.1 |
| YPTB3729 | crp | cap,csm | YPTB_RS20170 | YPTB3729 | | cAMP-activated global transcriptional regulator CRP | WP_002212297.1 |
| YPTB3732 | | | YPTB_RS20185 | YPTB3732 | | aminodeoxychorismate synthase component II | WP_011193225.1 |
| YPTB3734 | ppiA | rotA | YPTB_RS20195 | YPTB3734 | | peptidylprolyl isomerase A | WP_002208878.1 |
| YPTB3743 | cobA | | YPTB_RS20240 | YPTB3743 | | siroheme synthase CysG | WP_011193232.1 |
| YPTB3746 | rpe | | YPTB_RS20255 | YPTB3746 | | ribulose-phosphate 3-epimerase | WP_002215694.1 |
| YPTB3750 | aroK | | YPTB_RS20275 | YPTB3750 | | shikimate kinase AroK | WP_002208899.1 |
| YPTB3760 | hslR | | YPTB_RS20330 | YPTB3760 | | ribosome-associated heat shock protein Hsp15 | WP_002208910.1 |
| YPTB3764 | ompR | kmt,ompB | YPTB_RS20350 | YPTB3764 | | two-component system response regulator OmpR | WP_002208914.1 |
| YPTB3786 | glgX | | YPTB_RS20475 | YPTB3786 | | glycogen debranching protein GlgX | WP_011193250.1 |
| YPTB3790 | asd | usg-1 | | YPTB3790 | | aspartate semialdehyde dehydrogenase | CAH23028.1 |
| YPTB3791 | | | YPTB_RS20500 | YPTB3791 | | YhgN family NAAT transporter | WP_002209508.1 |
| YPTB3840 | dppC | | YPTB_RS20755 | YPTB3840 | | dipeptide ABC transporter permease DppC | WP_002209563.1 |
| YPTB3847 | uhpA | | YPTB_RS20800 | YPTB3847 | | transcriptional regulator UhpA | WP_002209570.1 |
| YPTB3914 | | | YPTB_RS21135 | YPTB3914 | | DNA-3-methyladenine glycosylase I | WP_002209626.1 |
| YPTB3915 | glyQ | | YPTB_RS21140 | YPTB3915 | | glycine--tRNA ligase subunit alpha | WP_002209624.1 |
| YPTB3916 | glyS | | YPTB_RS21145 | YPTB3916 | | glycine--tRNA ligase subunit beta | WP_002209623.1 |
| YPTB3920 | | | YPTB_RS21165 | YPTB3920 | | MltR family transcriptional regulator | WP_002209619.1 |
| YPTB3921 | | | YPTB_RS21170 | YPTB3921 | | YibL family ribosome-associated protein | WP_002209618.1 |
| YPTB3940 | gyrB | acrB,nalC,parA,pcbA | YPTB_RS21280 | YPTB3940 | | DNA topoisomerase (ATP-hydrolyzing) subunit B | WP_002209642.1 |
| YPTB3942 | dnaN | | YPTB_RS21295 | YPTB3942 | | DNA polymerase III subunit beta | WP_002209645.1 |
| YPTB3945 | rpmH | rimA,ssaF | YPTB_RS21305 | YPTB3945 | | 50S ribosomal protein L34 | WP_002220736.1 |
| YPTB3946 | rnpA | | YPTB_RS21310 | YPTB3946 | | ribonuclease P protein component | WP_002228153.1 |
| YPTB3947 | yidD | | YPTB_RS23380 | YPTB3947 | | membrane protein insertion efficiency factor YidD | WP_002228756.1 |
| YPTB3950 | | | YPTB_RS21325 | YPTB3950 | | trans-2-enoyl-CoA reductase family protein | WP_002215588.1 |
| YPTB3960 | pstB | phoT | YPTB_RS21375 | YPTB3960 | | phosphate ABC transporter ATP-binding protein PstB | WP_002215562.1 |
| YPTB3961 | pstA | phoT | YPTB_RS21380 | YPTB3961 | | phosphate ABC transporter permease PstA | WP_002215560.1 |
| YPTB3965 | glmU | | YPTB_RS21400 | YPTB3965 | | bifunctional UDP-N-acetylglucosamine diphosphorylase/glucosamine-1-phosphate N-acetyltransferase GlmU | WP_002215550.1 |
| YPTB3966 | atpC | papG,uncC | YPTB_RS21405 | YPTB3966 | | F0F1 ATP synthase subunit epsilon | WP_002215546.1 |
| YPTB3969 | atpA | papA,uncA | YPTB_RS21420 | YPTB3969 | | F0F1 ATP synthase subunit alpha | WP_002220758.1 |
| YPTB3970 | atpH | papE,uncH | YPTB_RS21425 | YPTB3970 | | F0F1 ATP synthase subunit delta | WP_002220760.1 |
| YPTB3971 | atpF | papF,uncF | YPTB_RS21430 | YPTB3971 | | F0F1 ATP synthase subunit B | WP_002220762.1 |
| YPTB3973 | atpB | papD,uncB | YPTB_RS21440 | YPTB3973 | | F0F1 ATP synthase subunit A | WP_002228150.1 |
