## Supplementary material for "First description of a *Yersinia pseudotuberculosis* clonal outbreak in France, confirmed using a new core genome multilocus sequence typing method": Table_S2

| Isolate | IP43168 | IP43186 | IP43211 | IP43237 | IP43278 | IP43304 | IP43305 | IP43330 | IP43355 | IP43377 | IP43382 | IP43410 | IP43453 | IP43476 | IP43492 | IP43500 | IP43579 | IP43611 | IP43692 | IP43716 | IP43951 | IP43952 | IP43989 | IP44039 | IP44042 | IP44063 | IP44094 | IP44110 | IP44154 | IP44180 | IP44214 | IP44616 | IP44623 | IP44632 | IP44652 |
|---|---|---|---|---|---|---|---|---|---|---|---|---|---|---|---|---|---|---|---|---|---|---|---|---|---|---|---|---|---|---|---|---|---|---|---|
| Lineage | 10 | 11 | 10 | 7 | 16 | 16 | 10 | 5 | 11 | 2 | 7 | 12 | 15 | 15 | 16 | 2 | 15 | 13 | 10 | 10 | 16 | 16 | 16 | 10 | 15 | 15 | 16 | 16 | 16 | 16 | 16 | 15 | 7 | 7 | 10 |
| yeps_YPTB0001 | 1 | 1 | 1 | 1 | 1 | 1 | 1 | 1 | 1 | 1 | 1 | 1 | 1 | 1 | 1 | 1 | 1 | 3 | 1 | 1 | 1 | 1 | 1 | 1 | 1 | 1 | 1 | 1 | 1 | 1 | 1 | 1 | 1 | 1 | 1 |
| yeps_YPTB0004 | 1 | 5 | 1 | 1 | 1 | 1 | 1 | 6 | 5 | 1 | 1 | 1 | 1 | 1 | 1 | 1 | 1 | 2 | 1 | 1 | 1 | 1 | 1 | 1 | 1 | 1 | 1 | 1 | 1 | 1 | 1 | 1 | 1 | 1 | 1 |
| yeps_YPTB0006 | 1 | 6 | 1 | 1 | 1 | 1 | 1 | 7 | 6 | 1 | 1 | 1 | 1 | 1 | 1 | 1 | 1 | 3 | 1 | 1 | 1 | 1 | 1 | 1 | 1 | 1 | 1 | 1 | 1 | 1 | 1 | 1 | 1 | 1 | 1 |
| yeps_YPTB0007 | 2 | 2 | 2 | 5 | 1 | 1 | 2 | 1 | 2 | 5 | 5 | 1 | 2 | 2 | 1 | 5 | 2 | 1 | 2 | 2 | 1 | 1 | 1 | 2 | 2 | 2 | 1 | 1 | 1 | 1 | 1 | 2 | 5 | 5 | 2 |
| yeps_YPTB0008 | 1 | 2 | 1 | 2 | 1 | 1 | 1 | 6 | 2 | 8 | 2 | 1 | 1 | 1 | 1 | 8 | 1 | 2 | 1 | 1 | 1 | 1 | 1 | 1 | 1 | 1 | 1 | 1 | 1 | 1 | 1 | 1 | 2 | 2 | 1 |
| yeps_YPTB0009 | 2 | 1 | 2 | 1 | 1 | 1 | 2 | 1 | 1 | 1 | 1 | 1 | 3 | 3 | 1 | 1 | 3 | 1 | 2 | 2 | 1 | 1 | 1 | 2 | 3 | 3 | 1 | 1 | 1 | 1 | 1 | 3 | 1 | 1 | 2 |
| yeps_YPTB0020 | 1 | 1 | 1 | 2 | 1 | 1 | 1 | 2 | 1 | 2 | 2 | 1 | 1 | 1 | 1 | 2 | 1 | 2 | 1 | 1 | 1 | 1 | 1 | 1 | 1 | 1 | 1 | 1 | 1 | 1 | 1 | 1 | 2 | 2 | 1 |
| yeps_YPTB0021 | 2 | 2 | 2 | 4 | 1 | 1 | 2 | 2 | 2 | 4 | 4 | 2 | 2 | 2 | 1 | 4 | 2 | 4 | 2 | 2 | 1 | 1 | 1 | 2 | 2 | 2 | 1 | 1 | 1 | 1 | 1 | 2 | 4 | 4 | 2 |
| yeps_YPTB0023 | 1 | 1 | 1 | 5 | 1 | 1 | 1 | 1 | 1 | 5 | 5 | 1 | 1 | 1 | 1 | 5 | 1 | 3 | 1 | 1 | 1 | 1 | 1 | 1 | 1 | 1 | 1 | 1 | 1 | 1 | 1 | 1 | 5 | 5 | 1 |
| yeps_YPTB0025 | 2 | 1 | 2 | 7 | 1 | 1 | 2 | 5 | 1 | 7 | 7 | 1 | 2 | 2 | 1 | 7 | 2 | 3 | 2 | 2 | 1 | 1 | 1 | 2 | 2 | 2 | 1 | 1 | 1 | 1 | 1 | 2 | 7 | 7 | 2 |
| yeps_YPTB0026 | 1 | 4 | 1 | 7 | 1 | 1 | 1 | 5 | 4 | 7 | 7 | 1 | 1 | 1 | 1 | 7 | 1 | 1 | 1 | 1 | 1 | 1 | 1 | 1 | 1 | 1 | 1 | 1 | 1 | 1 | 1 | 1 | 7 | 7 | 1 |
| yeps_YPTB0027 | 5 | 7 | 5 | 10 | 1 | 1 | 5 | 8 | 7 | 10 | 10 | 3 | 5 | 5 | 1 | 10 | 5 | 5 | 5 | 5 | 1 | 1 | 1 | 5 | 5 | 5 | 1 | 1 | 1 | 1 | 1 | 5 | 10 | 10 | 5 |
| yeps_YPTB0029 | 2 | 2 | 2 | 5 | 1 | 1 | 2 | 3 | 2 | 5 | 5 | 1 | 2 | 2 | 1 | 5 | 2 | 2 | 2 | 2 | 1 | 1 | 1 | 2 | 2 | 2 | 1 | 1 | 1 | 1 | 1 | 2 | 5 | 5 | 2 |
| yeps_YPTB0030 | 3 | 5 | 3 | 6 | 1 | 1 | 3 | 1 | 5 | 6 | 6 | 3 | 3 | 3 | 1 | 6 | 3 | 3 | 68 | 3 | 1 | 1 | 1 | 3 | 3 | 3 | 1 | 1 | 1 | 1 | 1 | 3 | 6 | 6 | 3 |
| yeps_YPTB0031 | 1 | 3 | 1 | 7 | 1 | 1 | 1 | 6 | 4 | 7 | 7 | 2 | 1 | 1 | 1 | 7 | 1 | 1 | 1 | 1 | 1 | 1 | 1 | 1 | 1 | 1 | 1 | 1 | 1 | 1 | 1 | 1 | 7 | 7 | 1 |
| yeps_YPTB0034 | 2 | 1 | 2 | 4 | 1 | 1 | 2 | 1 | 1 | 4 | 4 | 1 | 1 | 1 | 1 | 4 | 1 | 1 | 2 | 2 | 1 | 1 | 1 | 2 | 1 | 1 | 1 | 1 | 1 | 1 | 1 | 1 | 4 | 4 | 2 |
| yeps_YPTB0035 | 3 | 7 | 3 | 6 | 1 | 1 | 3 | 8 | 7 | 6 | 6 | 1 | 3 | 3 | 1 | 6 | 3 | 6 | 3 | 3 | 1 | 1 | 1 | 3 | 3 | 3 | 1 | 1 | 1 | 1 | 1 | 3 | 6 | 6 | 3 |
| yeps_YPTB0036 | 1 | 1 | 1 | 1 | 1 | 1 | 1 | 1 | 3 | 1 | 1 | 1 | 1 | 1 | 1 | 1 | 1 | 1 | 1 | 1 | 1 | 1 | 1 | 1 | 1 | 1 | 1 | 1 | 1 | 1 | 1 | 1 | 1 | 1 | 1 |
| yeps_YPTB0038 | 4 | 8 | 4 | 5 | 1 | 1 | 4 | 9 | 8 | 5 | 5 | 3 | 4 | 4 | 1 | 5 | 4 | 5 | 4 | 4 | 1 | 1 | 1 | 4 | 4 | 4 | 1 | 1 | 1 | 1 | 1 | 4 | 5 | 5 | 4 |
| yeps_YPTB0041 | 1 | 1 | 1 | 1 | 1 | 1 | 1 | 1 | 1 | 1 | 1 | 1 | 1 | 1 | 1 | 1 | 1 | 2 | 1 | 1 | 1 | 1 | 1 | 1 | 1 | 1 | 1 | 1 | 1 | 1 | 1 | 1 | 1 | 1 | 1 |
| yeps_YPTB0042 | 1 | 3 | 1 | 2 | 1 | 1 | 1 | 6 | 3 | 2 | 2 | 1 | 1 | 1 | 1 | 2 | 1 | 3 | 1 | 1 | 1 | 1 | 1 | 1 | 1 | 1 | 1 | 1 | 1 | 1 | 1 | 1 | 2 | 2 | 1 |
| yeps_YPTB0045 | 4 | 5 | 4 | 2 | 3 | 3 | 4 | 5 | 5 | 2 | 2 | 3 | 3 | 3 | 3 | 2 | 3 | 5 | 4 | 4 | 3 | 3 | 3 | 4 | 3 | 3 | 3 | 3 | 3 | 3 | 3 | 3 | 2 | 2 | 4 |
| yeps_YPTB0046 | 1 | 1 | 1 | 1 | 1 | 1 | 1 | 1 | 1 | 3 | 1 | 1 | 1 | 1 | 1 | 3 | 1 | 1 | 1 | 1 | 1 | 1 | 1 | 1 | 1 | 1 | 1 | 1 | 1 | 1 | 1 | 1 | 1 | 1 | 1 |
| yeps_YPTB0050 | 2 | 4 | 2 | 2 | 1 | 2 | 2 | 3 | 4 | 2 | 2 | 2 | 2 | 2 | 2 | 2 | 2 | 3 | 2 | 2 | 2 | 2 | 2 | 2 | 2 | 2 | 2 | 2 | 2 | 2 | 2 | 2 | 2 | 2 | 2 |
| yeps_YPTB0051 | 3 | 6 | 3 | 1 | 1 | 1 | 3 | 4 | 6 | 1 | 1 | 1 | 1 | 1 | 1 | 1 | 1 | 4 | 3 | 3 | 1 | 1 | 1 | 3 | 1 | 1 | 1 | 1 | 1 | 1 | 1 | 1 | 1 | 1 | 3 |
| yeps_YPTB0054 | 38 | 2 | 1 | 1 | 1 | 1 | 1 | 1 | 2 | 1 | 1 | 1 | 1 | 1 | 1 | 1 | 1 | 1 | 1 | 1 | 1 | 1 | 1 | 1 | 1 | 1 | 1 | 1 | 1 | 1 | 1 | 1 | 1 | 1 | 1 |
| yeps_YPTB0056 | 1 | 7 | 1 | 7 | 1 | 1 | 1 | 3 | 7 | 7 | 7 | 3 | 1 | 1 | 1 | 7 | 1 | 3 | 1 | 1 | 1 | 1 | 1 | 1 | 1 | 1 | 1 | 1 | 1 | 1 | 1 | 1 | 7 | 7 | 1 |
| yeps_YPTB0057 | 1 | 2 | 1 | 2 | 1 | 1 | 1 | 2 | 2 | 5 | 2 | 1 | 1 | 1 | 1 | 5 | 1 | 2 | 1 | 1 | 1 | 1 | 1 | 1 | 1 | 1 | 1 | 1 | 1 | 1 | 1 | 1 | 2 | 2 | 1 |
| yeps_YPTB0058 | 1 | 5 | 1 | 2 | 1 | 1 | 1 | 2 | 5 | 6 | 2 | 1 | 1 | 1 | 1 | 6 | 1 | 2 | 1 | 1 | 1 | 1 | 1 | 1 | 1 | 1 | 1 | 1 | 1 | 1 | 1 | 1 | 2 | 2 | 1 |
| yeps_YPTB0059 | 1 | 6 | 1 | 5 | 1 | 1 | 1 | 7 | 6 | 5 | 5 | 1 | 1 | 1 | 1 | 5 | 1 | 3 | 1 | 1 | 1 | 1 | 1 | 1 | 1 | 1 | 1 | 1 | 1 | 1 | 1 | 1 | 5 | 5 | 1 |
| yeps_YPTB0060 | 1 | 8 | 1 | 12 | 1 | 1 | 1 | 4 | 9 | 11 | 12 | 1 | 1 | 6 | 1 | 11 | 6 | 4 | 1 | 1 | 1 | 1 | 1 | 1 | 1 | 1 | 1 | 1 | 1 | 1 | 1 | 1 | 12 | 12 | 1 |
| yeps_YPTB0061 | 2 | 1 | 2 | 2 | 1 | 1 | 2 | 2 | 1 | 2 | 2 | 1 | 1 | 1 | 1 | 2 | 1 | 2 | 2 | 2 | 1 | 1 | 1 | 2 | 1 | 1 | 1 | 1 | 1 | 1 | 1 | 1 | 2 | 2 | 3 |
| yeps_YPTB0062 | 1 | 1 | 1 | 1 | 1 | 1 | 1 | 2 | 1 | 1 | 1 | 9 | 1 | 1 | 1 | 1 | 1 | 1 | 1 | 1 | 1 | 1 | 1 | 1 | 1 | 1 | 1 | 1 | 1 | 1 | 1 | 1 | 1 | 1 | 1 |
| yeps_YPTB0063 | 1 | 1 | 1 | 1 | 1 | 1 | 1 | 1 | 1 | 1 | 1 | 1 | 1 | 1 | 1 | 1 | 1 | 1 | 1 | 1 | 1 | 1 | 1 | 1 | 1 | 1 | 1 | 1 | 1 | 1 | 1 | 1 | 1 | 1 | 1 |
| yeps_YPTB0066 | 1 | 5 | 1 | 6 | 1 | 1 | 1 | 1 | 5 | 6 | 6 | 2 | 3 | 3 | 1 | 6 | 3 | 1 | 1 | 1 | 1 | 1 | 1 | 1 | 3 | 3 | 1 | 1 | 1 | 1 | 1 | 3 | 6 | 6 | 1 |
| yeps_YPTB0067 | 3 | 9 | 3 | 6 | 1 | 1 | 3 | 6 | 9 | 6 | 6 | 3 | 1 | 1 | 1 | 6 | 1 | 2 | 3 | 3 | 1 | 1 | 1 | 3 | 1 | 1 | 1 | 1 | 1 | 1 | 1 | 1 | 6 | 6 | 3 |
| yeps_YPTB0068 | 13 | 8 | 5 | 12 | 1 | 1 | 5 | 9 | 8 | 10 | 12 | 3 | 1 | 1 | 1 | 10 | 1 | 2 | 5 | 5 | 1 | 1 | 1 | 5 | 1 | 1 | 1 | 1 | 1 | 1 | 1 | 1 | 12 | 12 | 5 |
| yeps_YPTB0069 | 1 | 9 | 1 | 10 | 1 | 1 | 1 | 6 | 9 | 10 | 10 | 3 | 1 | 1 | 1 | 10 | 1 | 6 | 1 | 1 | 1 | 1 | 1 | 1 | 1 | 1 | 1 | 1 | 1 | 1 | 1 | 1 | 10 | 10 | 1 |
| yeps_YPTB0071 | 1 | 2 | 1 | 3 | 1 | 1 | 1 | 2 | 2 | 2 | 3 | 1 | 1 | 1 | 1 | 2 | 1 | 2 | 1 | 1 | 1 | 1 | 1 | 1 | 1 | 1 | 1 | 1 | 1 | 1 | 1 | 1 | 3 | 3 | 1 |
| yeps_YPTB0073 | 3 | 3 | 3 | 3 | 1 | 1 | 3 | 3 | 3 | 3 | 3 | 1 | 1 | 1 | 1 | 3 | 1 | 3 | 3 | 3 | 1 | 1 | 1 | 3 | 1 | 1 | 1 | 1 | 1 | 1 | 1 | 1 | 3 | 3 | 3 |
| yeps_YPTB0075 | 4 | 3 | 4 | 3 | 1 | 3 | 4 | 4 | 3 | 3 | 3 | 3 | 3 | 3 | 3 | 3 | 3 | 3 | 4 | 4 | 3 | 3 | 3 | 4 | 3 | 3 | 3 | 3 | 3 | 3 | 3 | 3 | 3 | 3 | 4 |
| yeps_YPTB0076 | 1 | 4 | 1 | 6 | 1 | 1 | 1 | 1 | 4 | 6 | 6 | 1 | 1 | 1 | 1 | 6 | 1 | 1 | 1 | 1 | 1 | 1 | 1 | 1 | 1 | 1 | 1 | 1 | 1 | 1 | 1 | 1 | 6 | 6 | 1 |
| yeps_YPTB0078 | 2 | 1 | 2 | 4 | 1 | 1 | 2 | 2 | 1 | 4 | 4 | 1 | 1 | 1 | 1 | 4 | 1 | 3 | 2 | 2 | 1 | 1 | 1 | 2 | 1 | 1 | 1 | 1 | 1 | 1 | 1 | 1 | 4 | 4 | 2 |
| yeps_YPTB0079 | 2 | 1 | 2 | 1 | 1 | 1 | 2 | 1 | 1 | 1 | 1 | 1 | 1 | 1 | 1 | 1 | 1 | 1 | 2 | 5 | 1 | 1 | 1 | 5 | 1 | 1 | 1 | 1 | 1 | 1 | 1 | 1 | 1 | 1 | 2 |
| yeps_YPTB0081 | 1 | 2 | 1 | 2 | 1 | 1 | 1 | 2 | 2 | 2 | 2 | 1 | 1 | 1 | 1 | 2 | 1 | 2 | 1 | 1 | 1 | 1 | 1 | 1 | 1 | 1 | 1 | 1 | 1 | 1 | 1 | 1 | 2 | 2 | 1 |
| yeps_YPTB0083 | 2 | 3 | 2 | 4 | 1 | 1 | 2 | 1 | 3 | 4 | 4 | 2 | 1 | 1 | 1 | 4 | 1 | 1 | 2 | 2 | 1 | 1 | 1 | 2 | 1 | 1 | 1 | 1 | 1 | 1 | 1 | 1 | 4 | 4 | 2 |
| yeps_YPTB0084 | 2 | 2 | 2 | 4 | 1 | 1 | 2 | 1 | 2 | 4 | 4 | 2 | 1 | 1 | 1 | 4 | 1 | 3 | 2 | 2 | 1 | 1 | 1 | 2 | 1 | 1 | 1 | 1 | 1 | 1 | 1 | 1 | 4 | 4 | 2 |
| yeps_YPTB0085 | 2 | 4 | 2 | 5 | 1 | 1 | 2 | 1 | 1 | 5 | 5 | 2 | 1 | 1 | 1 | 5 | 1 | 2 | 2 | 2 | 1 | 1 | 1 | 2 | 1 | 1 | 1 | 1 | 1 | 1 | 1 | 1 | 5 | 5 | 2 |
| yeps_YPTB0087 | 2 | 1 | 2 | 1 | 1 | 1 | 2 | 1 | 1 | 1 | 1 | 1 | 1 | 1 | 1 | 1 | 1 | 1 | 2 | 2 | 1 | 1 | 1 | 2 | 1 | 1 | 1 | 1 | 1 | 1 | 1 | 1 | 1 | 1 | 2 |
| yeps_YPTB0088 | 1 | 1 | 1 | 1 | 1 | 1 | 1 | 1 | 1 | 1 | 1 | 1 | 1 | 1 | 1 | 1 | 1 | 3 | 1 | 1 | 1 | 1 | 1 | 1 | 1 | 1 | 1 | 1 | 1 | 1 | 1 | 1 | 1 | 1 | 1 |
| yeps_YPTB0090 | 3 | 1 | 3 | 1 | 1 | 1 | 3 | 6 | 1 | 1 | 1 | 2 | 1 | 1 | 1 | 1 | 1 | 1 | 3 | 3 | 1 | 1 | 1 | 3 | 1 | 1 | 1 | 1 | 1 | 1 | 1 | 1 | 1 | 1 | 3 |
| yeps_YPTB0091 | 4 | 1 | 4 | 1 | 1 | 1 | 4 | 4 | 1 | 1 | 1 | 3 | 1 | 1 | 1 | 1 | 1 | 2 | 4 | 4 | 1 | 1 | 1 | 4 | 1 | 1 | 1 | 1 | 1 | 1 | 1 | 1 | 1 | 1 | 4 |
| yeps_YPTB0095 | 1 | 1 | 1 | 1 | 1 | 1 | 1 | 1 | 1 | 1 | 1 | 1 | 1 | 1 | 1 | 1 | 1 | 1 | 1 | 1 | 1 | 1 | 1 | 1 | 1 | 1 | 1 | 1 | 1 | 1 | 1 | 1 | 1 | 1 | 1 |
| yeps_YPTB0096 | 1 | 2 | 1 | 8 | 1 | 1 | 1 | 6 | 2 | 7 | 8 | 3 | 4 | 4 | 1 | 7 | 4 | 2 | 1 | 1 | 1 | 1 | 1 | 1 | 4 | 4 | 1 | 1 | 1 | 1 | 1 | 4 | 8 | 8 | 1 |
| yeps_YPTB0098 | 1 | 1 | 1 | 2 | 1 | 1 | 1 | 1 | 1 | 2 | 2 | 2 | 2 | 2 | 1 | 2 | 2 | 2 | 1 | 1 | 1 | 1 | 1 | 1 | 2 | 2 | 1 | 1 | 1 | 1 | 1 | 2 | 2 | 2 | 1 |
| yeps_YPTB0100 | 1 | 1 | 1 | 1 | 1 | 1 | 1 | 1 | 1 | 1 | 1 | 1 | 3 | 3 | 1 | 1 | 3 | 1 | 1 | 1 | 1 | 1 | 1 | 1 | 3 | 3 | 1 | 1 | 1 | 1 | 1 | 3 | 1 | 1 | 1 |
| yeps_YPTB0102 | 1 | 1 | 1 | 1 | 1 | 1 | 1 | 1 | 1 | 1 | 1 | 1 | 1 | 1 | 1 | 1 | 1 | 1 | 1 | 1 | 1 | 1 | 1 | 1 | 1 | 1 | 1 | 1 | 1 | 1 | 1 | 1 | 1 | 1 | 1 |
| yeps_YPTB0106 | 3 | 7 | 3 | 8 | 1 | 1 | 3 | 2 | 7 | 2 | 2 | 3 | 1 | 1 | 1 | 2 | 1 | 75 | 3 | 3 | 1 | 1 | 1 | 3 | 1 | 1 | 1 | 1 | 1 | 1 | 1 | 1 | 8 | 8 | 3 |
| yeps_YPTB0115 | 4 | 9 | 4 | 10 | 1 | 1 | 4 | 10 | 9 | 10 | 10 | 3 | 1 | 1 | 1 | 10 | 1 | 5 | 4 | 4 | 1 | 1 | 1 | 4 | 1 | 1 | 1 | 1 | 1 | 1 | 1 | 1 | 10 | 10 | 4 |
| yeps_YPTB0119 | 3 | 3 | 3 | 3 | 1 | 1 | 3 | 1 | 3 | 3 | 3 | 1 | 1 | 1 | 1 | 3 | 1 | 3 | 3 | 3 | 1 | 1 | 1 | 3 | 1 | 1 | 1 | 1 | 1 | 1 | 1 | 1 | 3 | 3 | 3 |
| yeps_YPTB0120 | 3 | 5 | 3 | 3 | 1 | 1 | 3 | 3 | 5 | 3 | 3 | 3 | 1 | 1 | 1 | 3 | 1 | 1 | 3 | 3 | 1 | 1 | 1 | 3 | 1 | 1 | 1 | 1 | 1 | 1 | 1 | 1 | 3 | 3 | 3 |
| yeps_YPTB0122 | 1 | 2 | 1 | 2 | 1 | 1 | 1 | 1 | 2 | 2 | 2 | 1 | 1 | 1 | 1 | 2 | 1 | 2 | 1 | 1 | 1 | 1 | 1 | 1 | 1 | 1 | 1 | 1 | 1 | 1 | 1 | 1 | 2 | 2 | 1 |
| yeps_YPTB0123 | 1 | 1 | 1 | 1 | 1 | 1 | 1 | 1 | 1 | 1 | 1 | 1 | 1 | 1 | 1 | 1 | 1 | 1 | 1 | 1 | 1 | 1 | 1 | 1 | 1 | 1 | 1 | 1 | 1 | 1 | 1 | 1 | 1 | 1 | 1 |
| yeps_YPTB0124 | 1 | 5 | 1 | 1 | 1 | 1 | 1 | 1 | 5 | 1 | 1 | 1 | 1 | 1 | 1 | 1 | 1 | 5 | 1 | 1 | 1 | 1 | 1 | 1 | 1 | 1 | 1 | 1 | 1 | 1 | 1 | 1 | 1 | 1 | 1 |
| yeps_YPTB0128 | 2 | 2 | 2 | 2 | 1 | 1 | 2 | 2 | 2 | 5 | 2 | 1 | 1 | 1 | 1 | 5 | 1 | 3 | 2 | 2 | 1 | 1 | 1 | 2 | 1 | 1 | 1 | 1 | 1 | 1 | 1 | 1 | 2 | 2 | 2 |
| yeps_YPTB0129 | 1 | 1 | 1 | 1 | 1 | 1 | 1 | 1 | 1 | 1 | 1 | 1 | 1 | 1 | 1 | 1 | 1 | 2 | 1 | 1 | 1 | 1 | 1 | 1 | 1 | 1 | 1 | 1 | 1 | 1 | 1 | 1 | 1 | 1 | 1 |
| yeps_YPTB0130 | 1 | 5 | 1 | 8 | 1 | 1 | 1 | 6 | 5 | 1 | 8 | 37 | 1 | 1 | 1 | 1 | 1 | 3 | 1 | 1 | 1 | 1 | 1 | 1 | 1 | 1 | 1 | 1 | 1 | 1 | 1 | 1 | 8 | 8 | 1 |
| yeps_YPTB0131 | 1 | 2 | 1 | 2 | 1 | 1 | 1 | 1 | 1 | 2 | 2 | 2 | 2 | 2 | 1 | 2 | 2 | 4 | 1 | 1 | 1 | 1 | 1 | 1 | 2 | 2 | 1 | 1 | 1 | 1 | 1 | 2 | 2 | 2 | 1 |
| yeps_YPTB0135 | 1 | 1 | 1 | 1 | 1 | 1 | 1 | 1 | 1 | 1 | 1 | 1 | 1 | 1 | 1 | 1 | 1 | 1 | 1 | 1 | 1 | 1 | 1 | 1 | 1 | 1 | 1 | 1 | 1 | 1 | 1 | 1 | 1 | 1 | 1 |
| yeps_YPTB0136 | 3 | 3 | 3 | 5 | 1 | 1 | 3 | 3 | 3 | 5 | 5 | 1 | 3 | 3 | 1 | 5 | 3 | 1 | 3 | 3 | 1 | 1 | 1 | 3 | 3 | 3 | 1 | 1 | 1 | 1 | 1 | 3 | 5 | 5 | 3 |
| yeps_YPTB0144 | 2 | 1 | 2 | 1 | 1 | 1 | 2 | 1 | 1 | 1 | 1 | 1 | 2 | 2 | 1 | 1 | 2 | 1 | 2 | 2 | 1 | 1 | 1 | 2 | 2 | 2 | 1 | 1 | 1 | 1 | 1 | 2 | 1 | 1 | 2 |
| yeps_YPTB0163 | 1 | 3 | 1 | 7 | 1 | 1 | 1 | 3 | 3 | 7 | 7 | 1 | 1 | 1 | 1 | 7 | 1 | 8 | 1 | 1 | 1 | 1 | 1 | 1 | 1 | 1 | 1 | 1 | 1 | 1 | 1 | 1 | 7 | 7 | 1 |
| yeps_YPTB0165 | 5 | 6 | 5 | 5 | 1 | 5 | 5 | 7 | 6 | 5 | 5 | 2 | 1 | 1 | 5 | 5 | 1 | 5 | 5 | 5 | 5 | 5 | 5 | 5 | 1 | 1 | 5 | 5 | 5 | 5 | 5 | 1 | 5 | 5 | 5 |
| yeps_YPTB0168 | 2 | 2 | 2 | 1 | 1 | 2 | 2 | 1 | 2 | 1 | 1 | 1 | 1 | 1 | 2 | 1 | 1 | 1 | 2 | 2 | 2 | 2 | 2 | 2 | 1 | 1 | 2 | 2 | 2 | 2 | 2 | 1 | 1 | 1 | 2 |
| yeps_YPTB0169 | 1 | 5 | 1 | 4 | 1 | 1 | 1 | 1 | 5 | 4 | 4 | 1 | 1 | 1 | 1 | 4 | 1 | 4 | 1 | 1 | 1 | 1 | 1 | 1 | 1 | 1 | 1 | 1 | 1 | 1 | 1 | 1 | 4 | 4 | 1 |
| yeps_YPTB0172 | 3 | 1 | 3 | 10 | 11 | 1 | 3 | 7 | 1 | 8 | 10 | 1 | 1 | 1 | 1 | 8 | 1 | 12 | 3 | 3 | 1 | 1 | 1 | 3 | 1 | 1 | 1 | 1 | 1 | 1 | 1 | 1 | 10 | 10 | 3 |
| yeps_YPTB0174 | 1 | 5 | 1 | 1 | 1 | 1 | 1 | 3 | 5 | 1 | 1 | 3 | 1 | 1 | 1 | 1 | 1 | 1 | 1 | 1 | 1 | 1 | 1 | 1 | 1 | 1 | 1 | 1 | 1 | 1 | 1 | 1 | 1 | 1 | 1 |
| yeps_YPTB0175 | 4 | 8 | 4 | 9 | 1 | 1 | 4 | 3 | 8 | 9 | 9 | 3 | 1 | 1 | 1 | 9 | 1 | 5 | 4 | 4 | 1 | 1 | 1 | 4 | 1 | 1 | 1 | 1 | 1 | 1 | 1 | 1 | 9 | 9 | 4 |
| yeps_YPTB0176 | 4 | 8 | 4 | 3 | 1 | 1 | 4 | 3 | 8 | 3 | 3 | 3 | 1 | 1 | 1 | 3 | 1 | 5 | 4 | 4 | 1 | 1 | 1 | 4 | 1 | 1 | 1 | 1 | 1 | 1 | 1 | 1 | 3 | 3 | 4 |
| yeps_YPTB0178 | 3 | 8 | 3 | 6 | 1 | 1 | 3 | 10 | 8 | 6 | 6 | 3 | 1 | 1 | 1 | 6 | 1 | 4 | 3 | 3 | 1 | 1 | 1 | 3 | 1 | 1 | 1 | 1 | 1 | 1 | 1 | 1 | 6 | 6 | 3 |
| yeps_YPTB0183 | 3 | 6 | 3 | 2 | 1 | 1 | 3 | 2 | 6 | 2 | 2 | 2 | 1 | 1 | 1 | 2 | 1 | 4 | 3 | 3 | 1 | 1 | 1 | 21 | 1 | 1 | 1 | 1 | 1 | 1 | 1 | 24 | 2 | 2 | 3 |
| yeps_YPTB0191 | 2 | 3 | 2 | 4 | 1 | 1 | 2 | 2 | 1 | 4 | 4 | 1 | 1 | 1 | 1 | 4 | 1 | 2 | 2 | 2 | 1 | 1 | 1 | 2 | 1 | 1 | 1 | 1 | 1 | 1 | 1 | 1 | 4 | 4 | 2 |
| yeps_YPTB0192 | 3 | 6 | 3 | 11 | 1 | 1 | 3 | 9 | 7 | 10 | 11 | 1 | 1 | 1 | 1 | 10 | 1 | 3 | 3 | 3 | 1 | 1 | 1 | 3 | 1 | 1 | 1 | 1 | 1 | 1 | 1 | 1 | 11 | 11 | 3 |
| yeps_YPTB0194 | 3 | 8 | 3 | 12 | 1 | 1 | 3 | 9 | 8 | 11 | 12 | 3 | 1 | 1 | 1 | 11 | 1 | 13 | 3 | 3 | 1 | 1 | 1 | 3 | 1 | 1 | 1 | 1 | 1 | 1 | 1 | 1 | 12 | 12 | 3 |
| yeps_YPTB0198 | 3 | 9 | 3 | 13 | 1 | 1 | 3 | 10 | 9 | 12 | 13 | 3 | 6 | 6 | 1 | 12 | 6 | 7 | 3 | 3 | 1 | 1 | 1 | 3 | 6 | 6 | 1 | 1 | 1 | 1 | 1 | 6 | 13 | 13 | 3 |
| yeps_YPTB0200 | 1 | 1 | 1 | 1 | 1 | 1 | 1 | 1 | 1 | 1 | 1 | 1 | 1 | 1 | 1 | 1 | 1 | 1 | 1 | 1 | 1 | 1 | 1 | 1 | 1 | 1 | 1 | 1 | 1 | 1 | 1 | 1 | 1 | 1 | 1 |
| yeps_YPTB0201 | 1 | 3 | 1 | 4 | 1 | 1 | 1 | 1 | 3 | 1 | 4 | 1 | 1 | 1 | 1 | 1 | 1 | 1 | 1 | 1 | 1 | 1 | 1 | 1 | 1 | 1 | 1 | 1 | 1 | 1 | 1 | 1 | 4 | 4 | 1 |
| yeps_YPTB0202 | 4 | 7 | 4 | 9 | 1 | 1 | 4 | 8 | 7 | 4 | 9 | 3 | 4 | 4 | 1 | 4 | 4 | 4 | 4 | 4 | 1 | 1 | 1 | 4 | 4 | 4 | 1 | 1 | 1 | 1 | 1 | 4 | 9 | 9 | 4 |
| yeps_YPTB0203 | 1 | 1 | 1 | 2 | 1 | 1 | 1 | 2 | 1 | 2 | 2 | 2 | 1 | 1 | 1 | 2 | 1 | 1 | 1 | 1 | 1 | 1 | 1 | 1 | 1 | 1 | 1 | 1 | 1 | 1 | 1 | 1 | 2 | 2 | 1 |
| yeps_YPTB0205 | 4 | 4 | 4 | 1 | 1 | 1 | 4 | 1 | 4 | 1 | 1 | 1 | 4 | 4 | 1 | 1 | 4 | 4 | 4 | 4 | 1 | 1 | 1 | 4 | 4 | 4 | 1 | 1 | 1 | 1 | 1 | 4 | 1 | 1 | 4 |
| yeps_YPTB0208 | 3 | 1 | 3 | 9 | 1 | 1 | 3 | 6 | 1 | 8 | 9 | 25 | 3 | 3 | 1 | 8 | 3 | 3 | 3 | 3 | 1 | 1 | 1 | 3 | 3 | 3 | 1 | 1 | 1 | 1 | 1 | 3 | 9 | 9 | 3 |
| yeps_YPTB0209 | 3 | 7 | 3 | 13 | 1 | 1 | 3 | 8 | 7 | 10 | 13 | 1 | 3 | 3 | 1 | 10 | 3 | 3 | 3 | 3 | 1 | 1 | 1 | 3 | 3 | 3 | 1 | 1 | 1 | 1 | 1 | 3 | 13 | 13 | 3 |
| yeps_YPTB0210 | 3 | 7 | 3 | 10 | 1 | 1 | 3 | 8 | 7 | 9 | 10 | 1 | 3 | 3 | 1 | 9 | 3 | 3 | 3 | 3 | 1 | 1 | 1 | 3 | 3 | 75 | 1 | 1 | 1 | 1 | 1 | 3 | 10 | 10 | 3 |
| yeps_YPTB0212 | 1 | 1 | 1 | 1 | 1 | 1 | 1 | 1 | 1 | 2 | 1 | 1 | 1 | 1 | 1 | 2 | 1 | 1 | 1 | 1 | 1 | 1 | 1 | 1 | 1 | 1 | 1 | 1 | 1 | 1 | 1 | 1 | 1 | 1 | 1 |
| yeps_YPTB0213 | 1 | 3 | 4 | 3 | 1 | 1 | 1 | 3 | 3 | 5 | 3 | 1 | 1 | 1 | 1 | 5 | 1 | 1 | 1 | 1 | 1 | 1 | 1 | 1 | 1 | 1 | 1 | 1 | 1 | 1 | 1 | 1 | 3 | 3 | 1 |
| yeps_YPTB0214 | 1 | 1 | 1 | 1 | 1 | 1 | 1 | 1 | 1 | 1 | 1 | 1 | 1 | 1 | 1 | 1 | 1 | 1 | 1 | 1 | 1 | 1 | 1 | 1 | 1 | 1 | 1 | 1 | 1 | 1 | 1 | 1 | 1 | 1 | 1 |
| yeps_YPTB0217 | 3 | 2 | 3 | 4 | 1 | 1 | 3 | 2 | 2 | 4 | 4 | 1 | 1 | 1 | 1 | 4 | 1 | 3 | 3 | 3 | 1 | 1 | 1 | 3 | 1 | 1 | 1 | 1 | 1 | 1 | 1 | 1 | 4 | 4 | 3 |
| yeps_YPTB0218 | 2 | 2 | 2 | 3 | 1 | 1 | 2 | 2 | 2 | 3 | 3 | 1 | 1 | 1 | 1 | 3 | 1 | 2 | 2 | 2 | 1 | 1 | 1 | 2 | 1 | 1 | 1 | 1 | 1 | 1 | 1 | 1 | 3 | 3 | 2 |
| yeps_YPTB0220 | 2 | 2 | 2 | 5 | 1 | 1 | 2 | 2 | 2 | 5 | 5 | 1 | 1 | 1 | 1 | 5 | 1 | 2 | 2 | 2 | 1 | 1 | 1 | 2 | 1 | 1 | 1 | 1 | 1 | 1 | 1 | 1 | 5 | 5 | 2 |
| yeps_YPTB0223 | 3 | 5 | 3 | 7 | 1 | 1 | 3 | 6 | 5 | 7 | 7 | 1 | 1 | 1 | 1 | 7 | 1 | 3 | 3 | 3 | 1 | 1 | 1 | 3 | 1 | 1 | 1 | 1 | 1 | 1 | 1 | 1 | 7 | 7 | 3 |
| yeps_YPTB0225 | 3 | 1 | 3 | 1 | 1 | 26 | 3 | 1 | 1 | 1 | 1 | 1 | 1 | 1 | 26 | 1 | 1 | 3 | 3 | 3 | 26 | 26 | 26 | 3 | 1 | 1 | 26 | 26 | 26 | 26 | 26 | 1 | 1 | 1 | 3 |
| yeps_YPTB0226 | 32 | 7 | 4 | 9 | 1 | 1 | 4 | 8 | 7 | 9 | 9 | 26 | 1 | 1 | 1 | 9 | 1 | 4 | 4 | 4 | 1 | 1 | 1 | 4 | 1 | 1 | 1 | 1 | 1 | 1 | 1 | 1 | 9 | 9 | 4 |
| yeps_YPTB0230 | 1 | 7 | 1 | 8 | 1 | 1 | 1 | 1 | 7 | 8 | 8 | 3 | 1 | 1 | 1 | 8 | 1 | 5 | 1 | 1 | 1 | 1 | 1 | 1 | 1 | 1 | 1 | 1 | 1 | 1 | 1 | 1 | 8 | 8 | 1 |
| yeps_YPTB0235 | 4 | 4 | 4 | 8 | 1 | 1 | 4 | 3 |  | 8 | 8 | 3 | 1 | 1 | 1 | 8 | 1 | 5 | 4 | 4 | 1 | 1 | 1 | 4 | 1 | 1 | 1 | 1 | 1 | 1 | 1 | 1 | 8 | 8 | 4 |
| yeps_YPTB0242 | 2 | 5 | 2 | 1 | 1 | 1 | 2 | 6 | 5 | 1 | 1 | 1 | 1 | 1 | 1 | 1 | 1 | 3 | 2 | 2 | 1 | 1 | 1 | 2 | 1 | 1 | 1 | 1 | 1 | 1 | 1 | 1 | 1 | 1 | 2 |
| yeps_YPTB0243 | 4 | 8 | 4 | 11 | 1 | 1 | 4 | 9 | 8 | 9 | 11 | 1 | 1 | 1 | 1 | 9 | 1 | 5 | 4 | 4 | 1 | 1 | 1 | 4 | 1 | 1 | 1 | 1 | 1 | 1 | 1 | 1 | 11 | 11 | 4 |
| yeps_YPTB0246 | 1 | 2 | 1 | 1 | 1 | 1 | 1 | 1 | 2 | 1 | 1 | 3 | 1 | 1 | 1 | 1 | 1 | 4 | 1 | 1 | 1 | 1 | 1 | 1 | 1 | 1 | 1 | 1 | 1 | 1 | 1 | 1 | 1 | 1 | 1 |
| yeps_YPTB0248 | 1 | 7 | 1 | 8 | 1 | 1 | 1 | 8 | 7 | 8 | 8 | 3 | 1 | 1 | 1 | 8 | 1 | 5 | 1 | 1 | 1 | 1 | 1 | 1 | 1 | 1 | 1 | 1 | 1 | 1 | 1 | 1 | 8 | 8 | 1 |
| yeps_YPTB0249 | 2 | 6 | 2 | 2 | 1 | 1 | 2 | 2 | 6 | 2 | 2 | 2 | 1 | 1 | 1 | 2 | 1 | 3 | 2 | 2 | 1 | 1 | 1 | 2 | 1 | 1 | 1 | 1 | 1 | 1 | 1 | 1 | 2 | 2 | 2 |
| yeps_YPTB0254 | 1 | 1 | 1 | 1 | 1 | 1 | 1 | 1 | 1 | 1 | 1 | 1 | 1 | 1 | 1 | 1 | 1 | 3 | 1 | 1 | 1 | 1 | 1 | 1 | 1 | 1 | 1 | 1 | 1 | 1 | 1 | 1 | 1 | 1 | 1 |
| yeps_YPTB0255 | 3 | 4 | 3 | 1 | 1 | 1 | 3 | 1 | 4 | 1 | 1 | 1 | 1 | 1 | 1 | 1 | 1 | 1 | 3 | 3 | 1 | 1 | 1 | 3 | 1 | 1 | 1 | 1 | 1 | 1 | 1 | 1 | 1 | 1 | 3 |
| yeps_YPTB0256 | 3 | 6 | 3 | 1 | 1 | 1 | 3 | 1 | 6 | 1 | 1 | 1 | 1 | 1 | 1 | 1 | 1 | 4 | 3 | 3 | 1 | 1 | 1 | 3 | 1 | 1 | 1 | 1 | 1 | 1 | 1 | 1 | 1 | 1 | 3 |
| yeps_YPTB0257 | 3 | 1 | 3 | 4 | 1 | 1 | 3 | 1 | 1 | 4 | 4 | 1 | 1 | 1 | 1 | 4 | 1 | 1 | 3 | 3 | 1 | 1 | 1 | 3 | 1 | 1 | 1 | 1 | 1 | 1 | 1 | 1 | 4 | 4 | 3 |
| yeps_YPTB0258 | 1 | 1 | 1 | 1 | 1 | 1 | 1 | 1 | 1 | 1 | 1 | 1 | 1 | 1 | 1 | 1 | 1 | 1 | 1 | 1 | 1 | 1 | 1 | 1 | 1 | 1 | 1 | 1 | 1 | 1 | 1 | 1 | 1 | 1 | 1 |
| yeps_YPTB0259 | 2 | 5 | 2 | 2 | 1 | 1 | 2 | 6 | 5 | 2 | 2 | 1 | 1 | 1 | 1 | 2 | 1 | 2 | 2 | 2 | 1 | 1 | 1 | 2 | 1 | 1 | 1 | 1 | 1 | 1 | 1 | 1 | 2 | 2 | 2 |
| yeps_YPTB0260 | 1 | 3 | 1 | 1 | 1 | 1 | 1 | 1 | 3 | 1 | 1 | 12 | 1 | 1 | 1 | 1 | 1 | 1 | 1 | 1 | 1 | 1 | 1 | 1 | 1 | 1 | 1 | 1 | 1 | 1 | 1 | 1 | 1 | 1 | 1 |
| yeps_YPTB0268 | 1 | 1 | 3 | 1 | 1 | 1 | 1 | 1 | 1 | 1 | 1 | 1 | 1 | 1 | 1 | 1 | 1 | 1 | 1 | 1 | 1 | 1 | 1 | 1 | 1 | 1 | 1 | 1 | 1 | 1 | 1 | 1 | 1 | 1 | 1 |
| yeps_YPTB0269 | 1 | 1 | 1 | 1 | 1 | 1 | 1 | 1 | 1 | 1 | 1 | 1 | 1 | 1 | 1 | 1 | 1 | 1 | 1 | 1 | 1 | 1 | 1 | 1 | 1 | 1 | 1 | 1 |  | 1 | 1 | 1 | 1 | 1 | 1 |
| yeps_YPTB0273 | 2 | 4 | 2 | 2 | 1 | 1 | 2 | 2 | 4 | 32 | 2 | 1 | 1 | 1 | 1 | 2 | 1 | 3 | 2 | 2 | 1 | 1 | 1 | 2 | 1 | 1 | 1 | 1 | 1 | 1 | 1 | 1 | 2 | 2 | 2 |
| yeps_YPTB0274 | 1 | 3 | 1 | 1 | 1 | 1 | 1 | 3 | 3 | 1 | 1 | 1 | 1 | 1 | 1 | 1 | 1 | 2 | 1 | 1 | 1 | 1 | 1 | 1 | 1 | 1 | 1 | 1 | 1 | 1 | 1 | 1 | 1 | 1 | 1 |
| yeps_YPTB0276 | 4 | 1 | 4 | 1 | 1 | 1 | 4 | 1 | 1 | 1 | 1 | 1 | 1 | 1 | 1 | 1 | 1 | 1 | 4 | 4 | 1 | 1 | 1 | 4 | 1 | 1 | 1 | 1 | 1 | 1 | 1 | 1 | 1 | 1 | 4 |
| yeps_YPTB0281 | 1 | 1 | 1 | 1 | 1 | 1 | 1 | 1 | 1 | 1 | 1 | 1 | 1 | 1 | 1 | 1 | 1 | 1 | 1 | 1 | 1 | 1 | 1 | 1 | 1 | 1 | 1 | 1 | 1 | 1 | 1 | 1 | 1 | 1 | 1 |
| yeps_YPTB0283 | 3 | 8 | 3 | 8 | 1 | 1 | 3 | 9 | 8 | 8 | 8 | 1 | 6 | 6 | 1 | 8 | 6 | 7 | 3 | 3 | 1 | 1 | 1 | 3 | 6 | 6 | 1 | 1 | 1 | 1 | 1 | 6 | 8 | 8 | 3 |
| yeps_YPTB0286 | 1 | 4 | 1 | 6 | 1 | 1 | 1 | 1 | 4 | 6 | 6 | 3 | 1 | 1 | 1 | 6 | 1 | 1 | 1 | 1 | 1 | 1 | 1 | 1 | 1 | 1 | 1 | 1 | 1 | 1 | 1 | 1 | 6 | 6 | 1 |
| yeps_YPTB0287 | 1 | 2 | 1 | 1 | 1 | 1 | 1 | 1 | 2 | 1 | 1 | 3 | 1 | 1 | 1 | 1 | 1 | 1 | 1 | 1 | 1 | 1 | 1 | 1 | 1 | 1 | 1 | 1 | 1 | 1 | 1 | 1 | 1 | 1 | 1 |
| yeps_YPTB0288 | 4 | 7 | 4 | 8 | 1 | 1 | 4 | 8 | 7 | 8 | 8 | 3 | 1 | 1 | 1 | 8 | 1 | 4 | 4 | 4 | 1 | 1 | 1 | 4 | 1 | 1 | 1 | 1 | 1 | 1 | 1 | 1 | 8 | 8 | 4 |
| yeps_YPTB0289 | 4 | 5 | 4 | 1 | 1 | 1 | 4 | 1 | 5 | 1 | 1 | 2 | 1 | 1 | 1 | 1 | 1 | 4 | 4 | 4 | 1 | 1 | 1 | 4 | 1 | 1 | 1 | 1 | 1 | 1 | 1 | 1 | 1 | 1 | 4 |
| yeps_YPTB0291 | 1 | 2 | 1 | 1 | 1 | 1 | 1 | 1 | 2 | 1 | 1 | 1 | 1 | 1 | 1 | 1 | 1 | 1 | 1 | 1 | 1 | 1 | 1 | 1 | 1 | 1 | 1 | 1 | 1 | 1 | 1 | 1 | 1 | 1 | 1 |
| yeps_YPTB0292 | 1 | 1 | 1 | 4 | 1 | 1 | 1 | 1 | 1 | 4 | 4 | 1 | 1 | 1 | 1 | 4 | 1 | 1 | 1 | 1 | 1 | 1 | 1 | 1 | 1 | 1 | 1 | 1 | 1 | 1 | 1 | 1 | 4 | 4 | 1 |
| yeps_YPTB0294 | 1 | 5 | 1 | 7 | 1 | 1 | 1 | 6 | 5 | 7 | 7 | 2 | 3 | 3 | 1 | 7 | 3 | 1 | 1 | 1 | 1 | 1 | 1 | 1 | 3 | 3 | 1 | 1 | 1 | 1 | 1 | 3 | 7 | 7 | 1 |
| yeps_YPTB0297 | 1 | 1 | 1 | 1 | 1 | 1 | 1 | 1 | 1 | 1 | 1 | 1 | 1 | 1 | 1 | 1 | 1 | 1 | 1 | 1 | 1 | 1 | 1 | 1 | 1 | 1 | 1 | 1 | 1 | 1 | 1 | 1 | 1 | 1 | 1 |
| yeps_YPTB0299 | 8 | 5 | 3 | 3 | 1 | 1 | 3 | 7 | 5 | 3 | 3 | 3 | 1 | 1 | 1 | 3 | 1 | 3 | 3 | 3 | 1 | 1 | 1 | 3 | 1 | 1 | 1 | 1 | 1 | 1 | 1 | 1 | 3 | 3 | 3 |
| yeps_YPTB0310 | 1 | 1 | 1 | 5 | 1 | 1 | 1 | 3 | 1 | 6 | 5 | 1 | 1 | 1 | 1 | 6 | 1 | 3 | 1 | 1 | 1 | 1 | 1 | 1 | 1 | 1 | 1 | 1 | 1 | 1 | 1 | 1 | 5 | 5 | 1 |
| yeps_YPTB0311 | 3 | 7 | 3 | 11 | 1 | 1 | 3 | 9 | 8 | 10 | 11 | 115 | 1 | 1 | 1 | 10 | 1 | 3 | 3 | 3 | 1 | 1 | 1 | 3 | 1 | 1 | 1 | 1 | 1 | 1 | 1 | 1 | 11 | 11 | 3 |
| yeps_YPTB0315 | 3 | 6 | 3 | 8 | 1 | 1 | 3 | 1 | 6 | 7 | 8 | 1 | 1 | 1 | 1 | 7 | 1 | 3 | 3 | 3 | 1 | 1 | 1 | 3 | 1 | 1 | 1 | 1 | 1 | 1 | 1 | 1 | 8 | 8 | 3 |
| yeps_YPTB0316 | 2 | 4 | 2 | 1 | 1 | 1 | 2 | 2 | 5 | 2 | 1 | 1 | 1 | 1 | 1 | 2 | 1 | 2 | 2 | 2 | 1 | 1 | 1 | 2 | 1 | 1 | 1 | 1 | 1 | 1 | 1 | 1 | 1 | 1 | 2 |
| yeps_YPTB0317 | 1 | 1 | 1 | 4 | 1 | 1 | 1 | 3 | 1 | 1 | 4 | 1 | 1 | 1 | 1 | 1 | 1 | 1 | 1 | 1 | 1 | 1 | 1 | 1 | 1 | 1 | 1 | 1 | 1 | 1 | 1 | 1 | 4 | 4 | 1 |
| yeps_YPTB0318 | 1 | 1 | 1 | 3 | 1 | 1 | 1 | 1 | 1 | 1 | 3 | 1 | 1 | 1 | 1 | 1 | 1 | 1 | 1 | 1 | 1 | 1 | 1 | 1 | 1 | 1 | 1 | 1 | 1 | 1 | 1 | 1 | 3 | 3 | 1 |
| yeps_YPTB0319 | 3 | 5 | 3 | 11 | 1 | 1 | 3 | 6 | 5 | 8 | 11 | 1 | 9 | 1 | 1 | 8 | 1 | 3 | 3 | 3 | 1 | 1 | 1 | 3 | 1 | 1 | 1 | 1 | 1 | 1 | 1 | 1 | 11 | 11 | 3 |
| yeps_YPTB0321 | 2 | 5 | 2 | 8 | 1 | 2 | 2 | 6 | 5 | 2 | 8 | 3 | 4 | 4 | 2 | 2 | 4 | 2 | 2 | 2 | 2 | 2 | 2 | 2 | 4 | 4 | 2 | 2 | 2 | 2 | 2 | 4 | 8 | 8 | 2 |
| yeps_YPTB0325 | 3 | 1 | 3 | 7 | 1 | 1 | 3 | 1 | 1 | 6 | 7 | 2 | 2 | 2 | 1 | 6 | 2 | 4 | 3 | 3 | 1 | 1 | 1 | 3 | 2 | 2 | 1 | 1 | 1 | 1 | 1 | 2 | 7 | 7 | 3 |
| yeps_YPTB0326 | 4 | 1 | 4 | 9 | 1 | 1 | 4 | 1 | 1 | 8 | 9 | 1 | 1 | 1 | 1 | 8 | 1 | 4 | 3 | 3 | 1 | 1 | 1 | 3 | 1 | 1 | 1 | 1 | 1 | 1 | 1 | 1 | 9 | 9 | 4 |
| yeps_YPTB0328 | 1 | 1 | 1 | 2 | 1 | 1 | 1 | 1 | 1 | 2 | 2 | 1 | 1 | 1 | 1 | 2 | 1 | 1 | 1 | 1 | 1 | 1 | 1 | 1 | 1 | 1 | 1 | 1 | 1 | 1 | 1 | 1 | 2 | 2 | 1 |
| yeps_YPTB0335 | 2 | 2 | 2 | 4 | 1 | 1 | 2 | 1 | 2 | 4 | 4 | 1 | 1 | 1 | 1 | 4 | 1 | 2 | 2 | 2 | 1 | 1 | 1 | 2 | 1 | 1 | 1 | 1 | 1 | 1 | 1 | 1 | 4 | 4 | 2 |
| yeps_YPTB0336 | 2 | 2 | 2 | 5 | 1 | 1 | 2 | 1 | 2 | 5 | 5 | 1 | 1 | 1 | 1 | 5 | 1 | 2 | 2 | 2 | 1 | 1 | 1 | 2 | 1 | 1 | 1 | 1 | 1 | 1 | 1 | 1 | 5 | 5 | 2 |
| yeps_YPTB0338 | 4 | 4 | 4 | 4 | 3 | 3 | 4 | 3 | 4 | 4 | 4 | 3 | 3 | 3 | 3 | 4 | 3 | 4 | 4 | 4 | 3 | 3 | 3 | 4 | 3 | 3 | 3 | 3 | 3 | 3 | 3 | 3 | 4 | 4 | 4 |
| yeps_YPTB0339 | 3 | 1 | 3 | 6 | 1 | 1 | 3 | 1 | 1 | 6 | 6 | 1 | 1 | 1 | 1 | 6 | 1 | 3 | 3 | 3 | 1 | 1 | 1 | 3 | 1 | 1 | 1 | 1 | 1 | 1 | 1 | 1 | 6 | 6 | 3 |
| yeps_YPTB0340 | 1 | 1 | 1 | 1 | 1 | 2 | 1 | 1 | 1 | 1 | 1 | 2 | 2 | 2 | 2 | 1 | 2 | 1 | 1 | 1 | 2 | 2 | 2 | 1 | 2 | 2 | 2 | 2 | 2 | 2 | 2 | 2 | 1 | 1 | 1 |
| yeps_YPTB0341 | 1 | 1 | 1 | 1 | 1 | 1 | 1 | 1 | 1 | 1 | 1 | 1 | 2 | 1 | 1 | 1 | 1 | 1 | 1 | 1 | 1 | 1 | 1 | 1 | 1 | 1 | 1 | 1 | 1 | 1 | 1 | 1 | 1 | 1 | 1 |
| yeps_YPTB0342 | 1 | 2 | 1 | 2 | 1 | 1 | 1 | 5 | 2 | 2 | 2 | 3 | 2 | 2 | 1 | 2 | 2 | 2 | 1 | 1 | 1 | 1 | 1 | 1 | 2 | 2 | 1 | 1 | 1 | 1 | 1 | 2 | 2 | 2 | 1 |
| yeps_YPTB0347 | 1 | 9 | 1 | 12 | 1 | 4 | 1 | 10 | 9 | 12 | 12 | 3 | 4 | 4 | 4 | 12 | 4 | 7 | 1 | 1 | 4 | 4 | 4 | 1 | 4 | 4 | 4 | 4 | 4 | 4 | 4 | 4 | 12 | 12 | 1 |
| yeps_YPTB0348 | 1 | 6 | 1 | 7 | 1 | 4 | 1 | 3 | 6 | 7 | 7 | 3 | 4 | 4 | 4 | 7 | 4 | 1 | 1 | 1 | 4 | 4 | 4 | 1 | 4 | 4 | 4 | 4 | 4 | 4 | 4 | 4 | 7 | 7 | 1 |
| yeps_YPTB0349 | 1 | 4 | 1 | 4 | 1 | 1 | 1 | 1 | 4 | 4 | 4 | 1 | 1 | 1 | 1 | 4 | 1 | 2 | 1 | 1 | 1 | 1 | 1 | 1 | 1 | 1 | 1 | 1 | 1 | 1 | 1 | 1 | 4 | 4 | 1 |
| yeps_YPTB0350 | 3 | 8 | 3 | 10 | 1 | 1 | 3 | 2 | 8 | 10 | 10 | 1 | 1 | 1 | 1 | 10 | 1 | 4 | 3 | 3 | 1 | 1 | 1 | 3 | 1 | 1 | 1 | 1 | 1 | 1 | 1 | 1 | 10 | 10 | 3 |
| yeps_YPTB0352 | 1 | 1 | 1 | 1 | 1 | 1 | 1 | 1 | 2 | 1 | 1 | 1 | 1 | 1 | 1 | 1 | 1 | 1 | 1 | 1 | 1 | 1 | 1 | 1 | 1 | 1 | 1 | 1 | 1 | 1 | 1 | 1 | 1 | 1 | 1 |
| yeps_YPTB0354 | 1 | 1 | 1 | 1 | 1 | 1 | 1 | 1 | 1 | 1 | 1 | 1 | 1 | 1 | 1 | 1 | 1 | 1 | 1 | 1 | 1 | 1 | 1 | 1 | 1 | 1 | 1 | 1 | 1 | 1 | 1 | 1 | 1 | 1 | 1 |
| yeps_YPTB0355 | 1 | 7 | 1 | 3 | 1 | 1 | 1 | 3 | 7 | 8 | 3 | 1 | 1 | 1 | 1 | 8 | 1 | 3 | 1 | 1 | 1 | 1 | 1 | 1 | 1 | 1 | 1 | 1 | 1 | 1 | 1 | 1 | 3 | 3 | 1 |
| yeps_YPTB0356 | 1 | 1 | 1 | 1 | 1 | 1 | 1 | 1 | 1 | 1 | 1 | 1 | 1 | 1 | 1 | 1 | 1 | 1 | 1 | 1 | 1 | 1 | 1 | 1 | 1 | 1 | 1 | 1 | 1 | 1 | 1 | 1 | 1 | 1 | 1 |
| yeps_YPTB0367 | 3 | 2 | 3 | 3 | 1 | 1 | 3 | 3 | 2 | 3 | 3 | 3 | 1 | 1 | 1 | 3 | 1 | 3 | 3 | 3 | 1 | 1 | 1 | 3 | 1 | 1 | 1 | 1 | 1 | 1 | 1 | 1 | 3 | 3 | 3 |
| yeps_YPTB0369 | 1 | 1 | 1 | 2 | 1 | 1 | 1 | 2 | 1 | 2 | 2 | 2 | 1 | 1 | 1 | 2 | 1 | 2 | 1 | 1 | 1 | 1 | 1 | 1 | 1 | 1 | 1 | 1 | 1 | 1 | 1 | 1 | 2 | 2 | 1 |
| yeps_YPTB0371 | 1 | 2 | 1 | 1 | 1 | 1 | 1 | 1 | 2 | 1 | 1 | 1 | 1 | 1 | 1 | 1 | 1 | 1 | 1 | 1 | 1 | 1 | 1 | 1 | 1 | 1 | 1 | 1 | 1 | 1 | 1 | 1 | 1 | 1 | 1 |
| yeps_YPTB0378 | 1 | 1 | 1 | 1 | 1 | 1 | 1 | 1 | 1 | 1 | 1 | 1 | 1 | 1 | 1 | 1 | 1 | 1 | 1 | 1 | 1 | 1 | 1 | 1 | 1 | 1 | 1 | 1 | 1 | 1 | 1 | 1 | 1 | 1 | 1 |
| yeps_YPTB0379 | 10 | 11 | 10 | 1 | 1 | 1 | 10 | 12 | 11 | 14 | 1 | 3 | 1 | 1 | 1 | 14 | 1 | 5 | 10 | 10 | 1 | 1 | 1 | 10 | 1 | 1 | 1 | 1 | 1 | 1 | 1 | 1 | 1 | 1 | 10 |
| yeps_YPTB0381 | 1 | 1 | 1 | 2 | 1 | 1 | 1 | 1 | 1 | 2 | 2 | 1 | 1 | 1 | 1 | 2 | 1 | 2 | 1 | 1 | 1 | 1 | 1 | 1 | 1 | 1 | 1 | 1 | 1 | 1 | 1 | 1 | 2 | 2 | 1 |

| | | | | | | | | | | | | | | | | | | | | | | | | | | | | | | | | | | | |
|---|---|---|---|---|---|---|---|---|---|---|---|---|---|---|---|---|---|---|---|---|---|---|---|---|---|---|---|---|---|---|---|---|---|---|---|
| yeps_YPTB0382 | 3 | 5 | 3 | 9 | 1 | 1 | 3 | 6 | 37 | 9 | 9 | 3 | 3 | 3 | 1 | 9 | 3 | 1 | 3 | 3 | 1 | 1 | 1 | 3 | 3 | 3 | 1 | 1 | 1 | 1 | 1 | 3 | 9 | 9 | 3 |
| yeps_YPTB0383 | 4 | 6 | 4 | 6 | 1 | 1 | 4 | 2 | 6 | 6 | 6 | 3 | 3 | 3 | 1 | 6 | 3 | 1 | 4 | 4 | 1 | 1 | 1 | 4 | 3 | 3 | 1 | 1 | 1 | 1 | 1 | 3 | 6 | 6 | 4 |
| yeps_YPTB0384 | 2 | 6 | 2 | 7 | 1 | 1 | 2 | 2 | 6 | 7 | 7 | 3 | 3 | 3 | 1 | 7 | 3 | 1 | 2 | 2 | 1 | 1 | 1 | 2 | 3 | 3 | 1 | 1 | 1 | 1 | 1 | 3 | 7 | 7 | 2 |
| yeps_YPTB0385 | 5 | 9 | 5 | 12 | 1 | 1 | 5 | 3 | 9 | 12 | 12 | 3 | 3 | 3 | 1 | 12 | 3 | 1 | 5 | 5 | 1 | 1 | 1 | 5 | 3 | 3 | 1 | 1 | 1 | 1 | 1 | 3 | 12 | 12 | 5 |
| yeps_YPTB0386 | 2 | 6 | 2 | 2 | 1 | 32 | 2 | 6 | 6 | 2 | 2 | 3 | 3 | 3 | 32 | 2 | 3 | 1 | 2 | 2 | 32 | 32 | 32 | 2 | 3 | 3 | 32 | 32 | 32 | 32 | 32 | 3 | 2 | 2 | 2 |
| yeps_YPTB0387 | 2 | 5 | 2 | 6 | 1 | 1 | 2 | 2 | 5 | 6 | 6 | 2 | 2 | 2 | 1 | 6 | 2 | 1 | 2 | 2 | 1 | 1 | 1 | 2 | 2 | 2 | 1 | 1 | 1 | 1 | 1 | 2 | 6 | 6 | 2 |
| yeps_YPTB0391 | 1 | 1 | 1 | 2 | 1 | 1 | 1 | 1 | 1 | 2 | 2 | 1 | 1 | 1 | 1 | 2 | 1 | 1 | 1 | 1 | 1 | 1 | 1 | 1 | 1 | 1 | 1 | 1 | 1 | 1 | 1 | 1 | 2 | 2 | 1 |
| yeps_YPTB0396 | 8 | 7 | 8 | 10 | 1 | 1 | 8 | 9 | 7 | 10 | 10 | 1 | 1 | 1 | 1 | 10 | 1 | 5 | 8 | 8 | 1 | 1 | 1 | 8 | 1 | 1 | 1 | 1 | 1 | 1 | 1 | 1 | 10 | 10 | 8 |
| yeps_YPTB0397 | 3 | 7 | 3 | 3 | 1 | 1 | 3 | 1 | 7 | 3 | 3 | 1 | 1 | 1 | 1 | 3 | 1 | 4 | 3 | 3 | 1 | 1 | 1 | 3 | 1 | 1 | 1 | 1 | 1 | 1 | 1 | 1 | 3 | 3 | 3 |
| yeps_YPTB0400 | 1 | 1 | 1 | 1 | 1 | 1 | 1 | 1 | 1 | 1 | 1 | 1 | 1 | 1 | 1 | 1 | 1 | 1 | 1 | 1 | 1 | 1 | 1 | 1 | 1 | 1 | 1 | 1 | 1 | 1 | 1 | 1 | 1 | 1 | 1 |
| yeps_YPTB0401 | 8 | 6 | 8 | 9 | 1 | 1 | 8 | 1 | 6 | 9 | 9 | 1 | 1 | 7 | 1 | 9 | 7 | 5 | 4 | 4 | 1 | 1 | 1 | 4 | 1 | 1 | 1 | 1 | 1 | 1 | 1 | 1 | 9 | 9 | 8 |
| yeps_YPTB0409 | 1 | 4 | 1 | 2 | 1 | 1 | 1 | 2 | 4 | 2 | 2 | 1 | 1 | 1 | 1 | 2 | 1 | 1 | 1 | 1 | 1 | 1 | 1 | 1 | 1 | 1 | 1 | 1 | 1 | 1 | 1 | 1 | 2 | 2 | 1 |
| yeps_YPTB0410 | 1 | 3 | 1 | 1 | 1 | 1 | 1 | 1 | 3 | 1 | 1 | 1 | 1 | 1 | 1 | 1 | 1 | 1 | 1 | 1 | 1 | 1 | 1 | 1 | 1 | 1 | 1 | 1 | 1 | 1 | 1 | 1 | 1 | 1 | 1 |
| yeps_YPTB0414 | 1 | 5 | 1 | 1 | 1 | 1 | 1 | 5 | 5 | 1 | 1 | 1 | 1 | 1 | 1 | 1 | 1 | 1 | 1 | 1 | 1 | 1 | 1 | 1 | 1 | 1 | 1 | 1 | 1 | 1 | 1 | 1 | 1 | 1 | 1 |
| yeps_YPTB0415 | 1 | 7 | 1 | 10 | 12 | 1 | 1 | 8 | 7 | 10 | 10 | 3 | 3 | 3 | 1 | 10 | 3 | 13 | 1 | 1 | 1 | 1 | 1 | 1 | 3 | 3 | 1 | 1 | 1 | 1 | 1 | 3 | 10 | 10 | 1 |
| yeps_YPTB0416 | 1 | 1 | 1 | 2 | 1 | 1 | 1 | 4 | 1 | 2 | 2 | 1 | 1 | 1 | 1 | 2 | 1 | 3 | 1 | 1 | 1 | 1 | 1 | 1 | 1 | 1 | 1 | 1 | 1 | 1 | 1 | 1 | 2 | 2 | 1 |
| yeps_YPTB0417 | 1 | 3 | 1 | 2 | 1 | 1 | 1 | 4 | 3 | 2 | 2 | 1 | 1 | 1 | 1 | 2 | 1 | 2 | 1 | 1 | 1 | 1 | 1 | 1 | 1 | 1 | 1 | 1 | 1 | 1 | 1 | 1 | 2 | 2 | 1 |
| yeps_YPTB0420 | 4 | 6 | 4 | 10 | 3 | 3 | 4 | 8 | 6 | 71 | 10 | 3 | 3 | 3 | 3 | 9 | 3 | 5 | 4 | 4 | 3 | 3 | 3 | 4 | 3 | 3 | 3 | 3 | 3 | 3 | 3 | 3 | 10 | 10 | 4 |
| yeps_YPTB0421 | 1 | 1 | 1 | 2 | 1 | 1 | 1 | 1 | 1 | 2 | 2 | 1 | 1 | 1 | 1 | 2 | 1 | 1 | 1 | 1 | 1 | 1 | 1 | 1 | 1 | 1 | 1 | 1 | 1 | 1 | 1 | 1 | 2 | 2 | 1 |
| yeps_YPTB0426 | 2 | 1 | 2 | 1 | 1 | 1 | 2 | 1 | 1 | 1 | 1 | 1 | 1 | 1 | 1 | 1 | 1 | 2 | 2 | 2 | 1 | 1 | 1 | 2 | 1 | 1 | 1 | 1 | 1 | 1 | 1 | 1 | 1 | 1 | 2 |
| yeps_YPTB0428 | 3 | 3 | 3 | 3 | 1 | 1 | 3 | 3 | 3 | 3 | 3 | 1 | 1 | 1 | 1 | 3 | 1 | 3 | 3 | 3 | 1 | 1 | 1 | 3 | 1 | 1 | 1 | 1 | 1 | 1 | 1 | 1 | 3 | 3 | 3 |
| yeps_YPTB0430 | 3 | 1 | 3 | 1 | 1 | 1 | 3 | 6 | 1 | 1 | 1 | 1 | 1 | 1 | 1 | 1 | 1 | 1 | 3 | 3 | 1 | 1 | 1 | 3 | 1 | 1 | 1 | 1 | 1 | 1 | 1 | 1 | 1 | 1 | 3 |
| yeps_YPTB0431 | 1 | 1 | 1 | 1 | 1 | 1 | 1 | 1 | 1 | 1 | 1 | 1 | 1 | 1 | 1 | 1 | 1 | 1 | 1 | 1 | 1 | 1 | 1 | 1 | 1 | 1 | 1 | 1 | 1 | 1 | 1 | 1 | 1 | 1 | 1 |
| yeps_YPTB0435 | 9 | 1 | 2 | 1 | 1 | 1 | 2 | 1 | 1 | 1 | 1 | 1 | 1 | 1 | 1 | 1 | 1 | 1 | 2 | 2 | 1 | 1 | 1 | 2 | 1 | 1 | 1 | 1 | 1 | 1 | 1 | 1 | 1 | 1 | 1 |
| yeps_YPTB0436 | 3 | 4 | 3 | 5 | 1 | 1 | 3 | 5 | 4 | 5 | 5 | 1 | 1 | 1 | 1 | 5 | 1 | 1 | 3 | 3 | 1 | 1 | 1 | 3 | 1 | 1 | 1 | 1 | 1 | 1 | 1 | 1 | 5 | 5 | 3 |
| yeps_YPTB0437 | 1 | 2 | 1 | 4 | 1 | 1 | 1 | 2 | 2 | 4 | 4 | 1 | 1 | 1 | 1 | 4 | 1 | 2 | 1 | 1 | 1 | 1 | 1 | 1 | 1 | 1 | 1 | 1 | 1 | 1 | 1 | 1 | 4 | 4 | 1 |
| yeps_YPTB0442 | 1 | 4 | 1 | 6 | 1 | 1 | 1 | 5 | 4 | 6 | 6 | 1 | 1 | 1 | 1 | 6 | 1 | 3 | 1 | 1 | 1 | 1 | 1 | 1 | 1 | 1 | 1 | 1 | 1 | 1 | 1 | 1 | 6 | 6 | 1 |
| yeps_YPTB0443 | 2 | 3 | 2 | 3 | 1 | 1 | 2 | 3 | 21 | 3 | 3 | 1 | 1 | 1 | 1 | 3 | 1 | 2 | 2 | 2 | 1 | 1 | 1 | 2 | 1 | 1 | 1 | 1 | 1 | 1 | 1 | 1 | 3 | 3 | 2 |
| yeps_YPTB0445 | 1 | 4 | 1 | 7 | 1 | 1 | 1 | 5 | 4 | 6 | 7 | 2 | 2 | 2 | 1 | 6 | 2 | 1 | 1 | 1 | 1 | 1 | 1 | 1 | 2 | 2 | 1 | 1 | 1 | 1 | 1 | 2 | 7 | 7 | 1 |
| yeps_YPTB0446 | 1 | 7 | 1 | 11 | 1 | 1 | 1 | 8 | 7 | 10 | 11 | 3 | 3 | 3 | 1 | 10 | 3 | 4 | 1 | 1 | 1 | 1 | 1 | 1 | 3 | 3 | 1 | 1 | 1 | 1 | 1 | 3 | 11 | 11 | 1 |
| yeps_YPTB0447 | 1 | 2 | 1 | 7 | 1 | 1 | 1 | 3 | 2 | 6 | 7 | 3 | 3 | 3 | 1 | 6 | 3 | 3 | 1 | 1 | 1 | 1 | 1 | 1 | 3 | 3 | 1 | 1 | 1 | 1 | 1 | 3 | 7 | 7 | 1 |
| yeps_YPTB0448 | 1 | 1 | 1 | 2 | 1 | 1 | 1 | 1 | 1 | 1 | 2 | 1 | 1 | 1 | 1 | 1 | 1 | 1 | 1 | 1 | 1 | 1 | 1 | 1 | 1 | 1 | 1 | 1 | 1 | 1 | 1 | 1 | 2 | 2 | 1 |
| yeps_YPTB0449 | 1 | 1 | 1 | 2 | 1 | 1 | 1 | 1 | 1 | 1 | 2 | 1 | 1 | 1 | 1 | 1 | 1 | 1 | 1 | 1 | 1 | 1 | 1 | 1 | 1 | 1 | 1 | 1 | 1 | 1 | 1 | 1 | 2 | 2 | 1 |
| yeps_YPTB0451 | 1 | 1 | 1 | 4 | 1 | 1 | 1 | 1 | 1 | 1 | 4 | 1 | 1 | 1 | 1 | 1 | 1 | 1 | 1 | 1 | 1 | 1 | 1 | 1 | 1 | 1 | 1 | 1 | 1 | 1 | 1 | 1 | 4 | 4 | 1 |
| yeps_YPTB0452 | 2 | 2 | 2 | 2 | 1 | 1 | 2 | 2 | 2 | 2 | 2 | 3 | 3 | 3 | 1 | 2 | 3 | 2 | 2 | 2 | 1 | 1 | 1 | 2 | 3 | 3 | 1 | 1 | 1 | 1 | 1 | 3 | 2 | 2 | 2 |
| yeps_YPTB0454 | 1 | 2 | 1 | 2 | 1 | 1 | 1 | 2 | 2 | 2 | 2 | 2 | 2 | 2 | 1 | 2 | 2 | 2 | 1 | 1 | 1 | 1 | 1 | 1 | 2 | 2 | 1 | 1 | 1 | 1 | 1 | 2 | 2 | 2 | 1 |
| yeps_YPTB0455 | 1 | 1 | 1 | 1 | 1 | 1 | 1 | 1 | 1 | 1 | 1 | 1 | 1 | 1 | 1 | 1 | 1 | 1 | 1 | 1 | 1 | 1 | 1 | 1 | 1 | 1 | 1 | 1 | 1 | 1 | 1 | 1 | 1 | 1 | 1 |
| yeps_YPTB0456 | 1 | 1 | 1 | 1 | 1 | 1 | 1 | 1 | 1 | 1 | 1 | 1 | 1 | 1 | 1 | 1 | 1 | 3 | 1 | 1 | 1 | 1 | 1 | 1 | 1 | 1 | 1 | 1 | 1 | 1 | 1 | 1 | 1 | 1 | 1 |
| yeps_YPTB0467 | 1 | 5 | 1 | 7 | 1 | 1 | 1 | 6 | 5 | 7 | 7 | 1 | 1 | 1 | 1 | 7 | 1 | 1 | 8 | 8 | 1 | 1 | 1 | 8 | 16 | 16 | 1 | 1 | 1 | 1 | 1 | 1 | 7 | 7 | 1 |
| yeps_YPTB0468 | 3 | 2 | 3 | 6 | 1 | 1 | 3 | 2 | 2 | 6 | 6 | 1 | 1 | 1 | 1 | 6 | 1 | 3 | 3 | 3 | 1 | 1 | 1 | 3 | 1 | 1 | 1 | 1 | 1 | 1 | 1 | 1 | 6 | 6 | 3 |
| yeps_YPTB0469 | 2 | 2 | 2 | 2 | 1 | 1 | 2 | 2 | 2 | 6 | 2 | 1 | 1 | 1 | 1 | 6 | 1 | 2 | 2 | 2 | 1 | 1 | 1 | 2 | 1 | 1 | 1 | 1 | 1 | 1 | 1 | 1 | 2 | 2 | 4 |
| yeps_YPTB0474 | 1 | 6 | 1 | 9 | 1 | 1 | 1 | 7 | 6 | 9 | 9 | 3 | 1 | 1 | 1 | 9 | 1 | 4 | 1 | 1 | 1 | 1 | 1 | 1 | 1 | 1 | 1 | 1 | 1 | 1 | 1 | 1 | 9 | 9 | 1 |
| yeps_YPTB0477 | 2 | 1 | 2 | 2 | 1 | 1 | 2 | 1 | 1 | 2 | 2 | 1 | 1 | 1 | 1 | 2 | 1 | 1 | 2 | 2 | 1 | 1 | 1 | 2 | 1 | 1 | 1 | 1 | 1 | 1 | 1 | 1 | 2 | 2 | 2 |
| yeps_YPTB0479 | 2 | 2 | 2 | 3 | 1 | 1 | 2 | 1 | 2 | 3 | 3 | 2 | 1 | 1 | 1 | 3 | 1 | 3 | 2 | 2 | 1 | 1 | 1 | 2 | 1 | 1 | 1 | 1 | 1 | 1 | 1 | 1 | 3 | 3 | 2 |
| yeps_YPTB0482 | 1 | 4 | 1 | 6 | 1 | 1 | 1 | 5 | 4 | 1 | 6 | 1 | 1 | 1 | 1 | 1 | 1 | 1 | 1 | 1 | 1 | 1 | 1 | 1 | 1 | 1 | 1 | 1 | 1 | 1 | 1 | 1 | 6 | 6 | 1 |
| yeps_YPTB0491 | 3 | 8 | 3 | 3 | 1 | 1 | 3 | 9 | 8 | 11 | 3 | 3 | 12 | 1 | 1 | 11 | 1 | 4 | 3 | 3 | 1 | 1 | 1 | 3 | 1 | 1 | 1 | 1 | 1 | 1 | 1 | 1 | 3 | 3 | 3 |
| yeps_YPTB0493 | 2 | | 2 | 2 | 1 | 1 | 2 | 5 | 2 | 2 | 2 | 2 | 1 | 1 | 1 | 2 | 1 | 3 | 2 | 2 | 1 | 1 | 1 | 2 | 1 | 1 | 1 | 1 | 1 | 1 | 62 | 1 | 2 | 2 | 2 |
| yeps_YPTB0495 | 3 | 1 | 3 | 8 | 1 | 1 | 3 | 1 | 1 | 8 | 8 | 3 | 1 | 1 | 1 | 8 | 1 | 4 | 3 | 3 | 1 | 1 | 1 | 3 | 1 | 1 | 1 | 1 | 1 | 1 | 1 | 1 | 8 | 8 | 3 |
| yeps_YPTB0496 | 1 | 4 | 1 | 5 | 1 | 1 | 1 | 1 | 4 | 1 | 5 | 1 | 1 | 1 | 1 | 1 | 1 | 2 | 1 | 1 | 1 | 1 | 1 | 1 | 1 | 1 | 1 | 1 | 1 | 1 | 1 | 1 | 5 | 5 | 1 |
| yeps_YPTB0497 | 1 | 4 | 6 | 4 | 1 | 1 | 1 | 1 | 4 | 5 | 4 | 1 | 1 | 1 | 1 | 5 | 1 | 2 | 1 | 1 | 1 | 1 | 1 | 1 | 1 | 1 | 1 | 1 | 1 | 1 | 1 | 1 | 4 | 4 | 1 |
| yeps_YPTB0498 | 2 | 1 | 2 | 4 | 1 | 1 | 2 | 2 | 1 | 4 | 4 | 2 | 1 | 1 | 1 | 4 | 1 | 3 | 2 | 2 | 1 | 1 | 1 | 2 | 1 | 1 | 1 | 1 | 1 | 1 | 1 | 1 | 4 | 4 | 2 |
| yeps_YPTB0499 | 5 | 9 | 5 | 14 | 1 | 1 | 5 | 11 | 9 | 18 | 14 | 3 | 1 | 1 | 1 | 13 | 1 | 6 | 5 | 5 | 1 | 1 | 1 | 5 | 1 | 1 | 1 | 1 | 1 | 1 | 1 | 1 | 14 | 14 | 5 |
| yeps_YPTB0502 | 6 | 7 | 6 | 11 | 1 | 1 | 6 | 1 | 7 | 10 | 11 | 3 | 1 | 1 | 1 | 10 | 16 | 5 | 4 | 4 | 1 | 1 | 1 | 4 | 1 | 1 | 1 | 1 | 1 | 1 | 1 | 1 | 11 | 11 | 6 |
| yeps_YPTB0503 | 4 | 8 | 4 | 8 | 1 | 1 | 4 | 9 | 8 | 8 | 8 | 3 | 1 | 1 | 1 | 8 | 1 | 5 | 4 | 4 | 1 | 1 | 1 | 4 | 1 | 1 | 1 | 1 | 1 | 1 | 1 | 1 | 8 | 8 | 4 |
| yeps_YPTB0504 | 1 | 6 | 1 | 9 | 1 | 1 | 1 | 7 | 6 | 9 | 9 | 3 | 1 | 1 | 1 | 9 | 1 | 4 | 1 | 1 | 1 | 1 | 1 | 1 | 1 | 1 | 1 | 1 | 1 | 1 | 1 | 1 | 9 | 9 | 1 |
| yeps_YPTB0505 | 1 | 4 | 1 | 4 | 1 | 1 | 1 | 3 | 4 | 4 | 4 | 3 | 1 | 1 | 1 | 4 | 1 | 4 | 1 | 1 | 1 | 1 | 1 | 1 | 1 | 1 | 1 | 1 | 1 | 1 | 1 | 1 | 4 | 4 | 1 |
| yeps_YPTB0507 | 1 | 6 | 1 | 9 | 1 | 1 | 1 | 7 | 6 | 9 | 9 | 3 | 1 | 1 | 1 | 9 | 1 | 4 | 1 | 1 | 1 | 1 | 1 | 1 | 1 | 1 | 1 | 1 | 1 | 1 | 1 | 1 | 9 | 9 | 1 |
| yeps_YPTB0508 | 1 | 1 | 1 | 8 | 1 | 1 | 1 | 5 | 1 | 7 | 8 | 1 | 1 | 1 | 1 | 7 | 1 | 2 | 1 | 1 | 1 | 1 | 1 | 1 | 1 | 1 | 1 | 1 | 1 | 1 | 1 | 1 | 8 | 8 | 1 |
| yeps_YPTB0509 | 1 | 4 | 1 | 7 | 1 | 1 | 1 | 5 | 4 | 7 | 7 | 1 | 1 | 1 | 1 | 7 | 1 | 2 | 1 | 1 | 1 | 1 | 1 | 1 | 1 | 1 | 1 | 1 | 1 | 1 | 1 | 1 | 7 | 7 | 1 |
| yeps_YPTB0510 | 1 | 6 | 12 | 10 | 1 | 1 | 1 | 7 | 6 | 9 | 10 | 1 | 1 | 1 | 1 | 9 | 1 | 11 | 1 | 1 | 1 | 1 | 1 | 1 | 1 | 1 | 1 | 1 | 1 | 1 | 1 | 1 | 10 | 10 | 1 |
| yeps_YPTB0511 | 3 | 7 | 3 | 6 | 1 | 1 | 3 | 3 | 7 | 6 | 6 | 3 | 1 | 5 | 1 | 6 | 5 | 4 | 3 | 3 | 1 | 1 | 1 | 3 | 1 | 1 | 1 | 1 | 1 | 1 | 1 | 1 | 6 | 6 | 3 |
| yeps_YPTB0512 | 3 | 4 | 3 | 7 | 1 | 1 | 3 | 3 | 4 | 7 | 7 | 3 | 1 | 1 | 1 | 7 | 1 | 4 | 3 | 3 | 1 | 1 | 1 | 3 | 1 | 1 | 1 | 1 | 1 | 1 | 1 | 1 | 7 | 7 | 3 |
| yeps_YPTB0513 | 9 | 8 | 9 | 13 | 1 | 1 | 9 | 11 | 10 | 13 | 13 | 3 | 1 | 1 | 1 | 13 | 1 | 5 | 9 | 9 | 1 | 1 | 1 | 9 | 1 | 1 | 1 | 1 | 1 | 1 | 1 | 1 | 13 | 13 | 9 |
| yeps_YPTB0516 | 4 | 1 | 4 | 1 | 1 | 1 | 4 | 8 | 1 | 10 | 1 | 3 | 1 | 1 | 1 | 10 | 1 | 5 | 4 | 4 | 1 | 1 | 1 | 4 | 1 | 1 | 1 | 1 | 1 | 1 | 1 | 1 | 1 | 1 | 4 |
| yeps_YPTB0517 | 4 | 8 | 4 | 11 | 1 | 1 | 4 | 3 | 8 | 9 | 11 | 3 | 10 | 4 | 1 | 9 | 4 | 5 | 4 | 4 | 1 | 1 | 1 | 4 | 4 | 4 | 1 | 1 | 1 | 1 | 1 | 4 | 11 | 11 | 4 |
| yeps_YPTB0518 | 1 | 1 | 1 | 1 | 1 | 1 | 1 | 1 | 1 | 1 | 1 | 2 | 1 | 1 | 1 | 1 | 1 | 2 | 1 | 1 | 1 | 1 | 1 | 1 | 1 | 1 | 1 | 1 | 1 | 1 | 1 | 1 | 1 | 1 | 1 |
| yeps_YPTB0519 | 1 | 6 | 1 | 9 | 1 | 1 | 1 | 1 | 6 | 8 | 9 | 3 | 1 | 1 | 1 | 8 | 1 | 4 | 1 | 1 | 1 | 1 | 1 | 1 | 1 | 1 | 1 | 1 | 1 | 1 | 1 | 1 | 9 | 9 | 1 |
| yeps_YPTB0520 | 9 | 7 | 5 | 6 | 1 | 1 | 5 | 8 | 7 | 6 | 6 | 3 | 1 | 1 | 1 | 6 | 1 | 3 | 5 | 5 | 1 | 1 | 1 | 5 | 1 | 1 | 1 | 1 | 1 | 1 | 1 | 1 | 6 | 6 | 5 |
| yeps_YPTB0521 | 3 | 6 | 3 | 8 | 9 | 1 | 3 | 7 | 6 | 8 | 8 | 2 | 1 | 1 | 1 | 8 | 1 | 1 | 3 | 3 | 1 | 1 | 1 | 3 | 1 | 1 | 1 | 1 | 1 | 1 | 1 | 1 | 8 | 8 | 3 |
| yeps_YPTB0522 | 1 | 6 | 1 | 9 | 1 | 1 | 1 | 10 | 6 | 8 | 9 | 2 | 1 | 1 | 1 | 8 | 1 | 3 | 1 | 1 | 1 | 1 | 1 | 1 | 1 | 1 | 1 | 1 | 1 | 1 | 1 | 1 | 9 | 9 | 1 |
| yeps_YPTB0526 | 3 | 1 | 3 | 6 | 1 | 1 | 3 | 3 | 1 | 6 | 6 | 1 | 1 | 1 | 1 | 6 | 1 | 1 | 3 | 3 | 1 | 1 | 1 | 3 | 1 | 1 | 1 | 1 | 1 | 1 | 1 | 1 | 6 | 6 | 3 |
| yeps_YPTB0529 | 11 | 10 | 11 | 14 | 1 | 1 | 11 | 2 | 88 | 14 | 14 | 3 | 4 | 4 | 1 | 14 | 4 | 6 | 11 | 11 | 1 | 1 | 1 | 11 | 4 | 4 | 1 | 1 | 1 | 1 | 1 | 4 | 14 | 14 | 11 |
| yeps_YPTB0533 | 3 | 6 | 3 | 3 | 1 | 1 | 3 | 8 | 6 | 3 | 3 | 3 | 1 | 1 | 1 | 3 | 1 | 4 | 3 | 3 | 1 | 1 | 1 | 3 | 1 | 1 | 1 | 1 | 1 | 1 | 1 | 1 | 3 | 3 | 3 |
| yeps_YPTB0541 | 3 | 2 | 3 | 2 | | 1 | 3 | 3 | 2 | 2 | 2 | 1 | 1 | 1 | 1 | 2 | 1 | 3 | 3 | 3 | 1 | 1 | 1 | 3 | 1 | 1 | 1 | 1 | 1 | 1 | 1 | 1 | 2 | 2 | 3 |
| yeps_YPTB0542 | 3 | 4 | 3 | 4 | 1 | 1 | 3 | 3 | 4 | 4 | 4 | 1 | 1 | 1 | 1 | 4 | 1 | 3 | 3 | 3 | 1 | 1 | 1 | 3 | 1 | 1 | 1 | 1 | 1 | 1 | 1 | 1 | 4 | 4 | 3 |
| yeps_YPTB0546 | 1 | 3 | 1 | 1 | 1 | 1 | 1 | 1 | 3 | 1 | 1 | 1 | 1 | 1 | 1 | 1 | 1 | 1 | 1 | 1 | 1 | 1 | 1 | 1 | 1 | 1 | 1 | 1 | 1 | 1 | 1 | 1 | 1 | 1 | 1 |
| yeps_YPTB0547 | 1 | 2 | 1 | 1 | 1 | 1 | 1 | 1 | 2 | 1 | 1 | 1 | 1 | 1 | 1 | 1 | 1 | 1 | 1 | 1 | 1 | 1 | 1 | 1 | 1 | 1 | 1 | 1 | 1 | 1 | 1 | 1 | 1 | 1 | 1 |
| yeps_YPTB0549 | 3 | 6 | 3 | 8 | 1 | 9 | 3 | 5 | 7 | 8 | 8 | 1 | 1 | 1 | 9 | 8 | 1 | 3 | 3 | 3 | 9 | 9 | 9 | 3 | 1 | 1 | 9 | 9 | 9 | 9 | 9 | 1 | 8 | 8 | 3 |
| yeps_YPTB0550 | 1 | 7 | 1 | 1 | 10 | 1 | 1 | 3 | 7 | 1 | 1 | 3 | 63 | 1 | 1 | 1 | 1 | 1 | 1 | 1 | 1 | 1 | 1 | 1 | 1 | 1 | 1 | 1 | 1 | 1 | 1 | 1 | 1 | 1 | 9 |
| yeps_YPTB0551 | 4 | 7 | 4 | 3 | 1 | 10 | 4 | 8 | 7 | 3 | 3 | 3 | 1 | 1 | 10 | 3 | 1 | 9 | 4 | 4 | 10 | 10 | 10 | 4 | 1 | 1 | 10 | 10 | 10 | 10 | 10 | 1 | 3 | 3 | 4 |
| yeps_YPTB0552 | 4 | 8 | 4 | 12 | 1 | 4 | 4 | 9 | 8 | 11 | 12 | 3 | 1 | 1 | 4 | 11 | 1 | 5 | 4 | 4 | 4 | 4 | 4 | 4 | 1 | 1 | 4 | 4 | 4 | 4 | 4 | 1 | 12 | 12 | 4 |
| yeps_YPTB0554 | 4 | 2 | 4 | 3 | 1 | 1 | 4 | 8 | 2 | 10 | 3 | 3 | 1 | 1 | 1 | 10 | 1 | 5 | 4 | 4 | 1 | 1 | 1 | 4 | 1 | 1 | 1 | 1 | 1 | 1 | 1 | 1 | 3 | 3 | 4 |
| yeps_YPTB0564 | 1 | 1 | 1 | 1 | 1 | 1 | 1 | 1 | 1 | 1 | 1 | 1 | 1 | 1 | 1 | 1 | 1 | 1 | 1 | 1 | 1 | 1 | 1 | 1 | 1 | 1 | 1 | 1 | 1 | 1 | 1 | 1 | 1 | 1 | 1 |
| yeps_YPTB0565 | 4 | 8 | 4 | 9 | 1 | 1 | 4 | 5 | 8 | 9 | 9 | 3 | 1 | 1 | 1 | 9 | 1 | 5 | 4 | 4 | 1 | 1 | 1 | 4 | 1 | 1 | 1 | 1 | 1 | 1 | 1 | 1 | 9 | 9 | 4 |
| yeps_YPTB0575 | 3 | 1 | 3 | 6 | 1 | 1 | 3 | 1 | 1 | 6 | 6 | 1 | 1 | 1 | 1 | 6 | 1 | 1 | 3 | 3 | 1 | 1 | 1 | 3 | 1 | 1 | 1 | 1 | 1 | 1 | 1 | 1 | 6 | 6 | 3 |
| yeps_YPTB0581 | 1 | 7 | 1 | 9 | 1 | 1 | 1 | 2 | 7 | 9 | 9 | 3 | 3 | 3 | 1 | 9 | 3 | 4 | 1 | 1 | 1 | 1 | 1 | 1 | 3 | 3 | 1 | 1 | 1 | 1 | 1 | 3 | 9 | 9 | 1 |
| yeps_YPTB0582 | 1 | 9 | 1 | 11 | 1 | 1 | 1 | 1 | 9 | 11 | 11 | 3 | 7 | 7 | 1 | 11 | 7 | 6 | 1 | 1 | 1 | 1 | 1 | 1 | 7 | 7 | 1 | 1 | 1 | 1 | 1 | 7 | 11 | 11 | 1 |
| yeps_YPTB0585 | 2 | 2 | 2 | 2 | 1 | 1 | 2 | 2 | 2 | 2 | 2 | 2 | 2 | 2 | 1 | 2 | 2 | 1 | 2 | 2 | 1 | 1 | 1 | 2 | 2 | 2 | 1 | 1 | 1 | 1 | 1 | 2 | 2 | 2 | 2 |
| yeps_YPTB0586 | 4 | 7 | 4 | 10 | 1 | 5 | 4 | 9 | 7 | 10 | 10 | 3 | 1 | 1 | 5 | 10 | 1 | 5 | 4 | 4 | 5 | 5 | 5 | 4 | 1 | 1 | 5 | 5 | 5 | 5 | 5 | 1 | 10 | 10 | 4 |
| yeps_YPTB0587 | 9 | 8 | 9 | 11 | 1 | 51 | 9 | 10 | 8 | 11 | 11 | 3 | 7 | 7 | 51 | 11 | 7 | 5 | 9 | 9 | 51 | 51 | 51 | 9 | 7 | 7 | 51 | 51 | 51 | 51 | 51 | 7 | 11 | 11 | 9 |
| yeps_YPTB0588 | 3 | 7 | 3 | 1 | 1 | 1 | 3 | 8 | 7 | 1 | 1 | 1 | 1 | 1 | 1 | 1 | 1 | 4 | 3 | 3 | 1 | 1 | 1 | 3 | 1 | 1 | 1 | 1 | 1 | 1 | 1 | 1 | 1 | 1 | 3 |
| yeps_YPTB0590 | 7 | 6 | 4 | 8 | 1 | 1 | 4 | 6 | 6 | 2 | 8 | 3 | 1 | 1 | 1 | 2 | 1 | 2 | 4 | 4 | 1 | 1 | 1 | 4 | 1 | 1 | 1 | 1 | 1 | 1 | 1 | 82 | 8 | 8 | 7 |
| yeps_YPTB0591 | 3 | 4 | 3 | 2 | 3 | 3 | 3 | 2 | 4 | 6 | 2 | 2 | 3 | 3 | 3 | 6 | 3 | 2 | 3 | 3 | 3 | 3 | 3 | 3 | 3 | 3 | 3 | 3 | 3 | 3 | 3 | 3 | 2 | 2 | 3 |
| yeps_YPTB0593 | 1 | 1 | 1 | 1 | 1 | 1 | 1 | 1 | 2 | 1 | 1 | 1 | 1 | 1 | 1 | 1 | 1 | 1 | 1 | 1 | 1 | 1 | 1 | 1 | 1 | 1 | 1 | 1 | 1 | 1 | 1 | 1 | 1 | 1 | 1 |
| yeps_YPTB0595 | 3 | 3 | 3 | 8 | 1 | 1 | 3 | 4 | 3 | 7 | 8 | 1 | 1 | 1 | 1 | 7 | 1 | 3 | 3 | 3 | 1 | 1 | 1 | 3 | 1 | 1 | 1 | 1 | 1 | 1 | 1 | 1 | 8 | 8 | 3 |
| yeps_YPTB0597 | 1 | 1 | 1 | 7 | 8 | 1 | 1 | 5 | 1 | 7 | 7 | 1 | 1 | 1 | 1 | 7 | 1 | 1 | 1 | 1 | 1 | 1 | 1 | 1 | 1 | 1 | 1 | 1 | 1 | 1 | 1 | 1 | 7 | 7 | 1 |
| yeps_YPTB0600 | 1 | 1 | 1 | 1 | 1 | 1 | 1 | 1 | 1 | 3 | 1 | 1 | 1 | 1 | 1 | 3 | 1 | 1 | 1 | 1 | 1 | 1 | 1 | 1 | 1 | 1 | 1 | 1 | 1 | 1 | 1 | 1 | 1 | 1 | 1 |
| yeps_YPTB0611 | 4 | 6 | 4 | 7 | 1 | 1 | 4 | 6 | 6 | 7 | 7 | 3 | 1 | 1 | 1 | 7 | 1 | 4 | 4 | 4 | 1 | 1 | 1 | 4 | 1 | 1 | 1 | 1 | 1 | 1 | 1 | 1 | 7 | 7 | 4 |
| yeps_YPTB0612 | 1 | 1 | 1 | 1 | 1 | 1 | 1 | 1 | 1 | 1 | 1 | 1 | 1 | 1 | 1 | 1 | 1 | 1 | 1 | 1 | 1 | 1 | 1 | 1 | 1 | 1 | 1 | 1 | 1 | 1 | 1 | 1 | 1 | 1 | 1 |
| yeps_YPTB0613 | 1 | 4 | 1 | 1 | 1 | 1 | 1 | 1 | 1 | 1 | 1 | 1 | 1 | 1 | 1 | 1 | 1 | 5 | 1 | 1 | 1 | 1 | 1 | 1 | 1 | 1 | 1 | 1 | 1 | 1 | 1 | 1 | 1 | 1 | 1 |
| yeps_YPTB0615 | 1 | 1 | 1 | 1 | 1 | 1 | 1 | 1 | 1 | 1 | 1 | 1 | 1 | 1 | 1 | 1 | 1 | 1 | 1 | 1 | 1 | 1 | 1 | 1 | 1 | 1 | 1 | 1 | 1 | 1 | 1 | 1 | 1 | 1 | 1 |
| yeps_YPTB0617 | 5 | 8 | 5 | 10 | 1 | 1 | 5 | 9 | 8 | 10 | 10 | 3 | 1 | 1 | 1 | 10 | 1 | 6 | 5 | 5 | 1 | 1 | 1 | 5 | 1 | 1 | 1 | 1 | 1 | 1 | 1 | 1 | 10 | 10 | 5 |
| yeps_YPTB0618 | 3 | 1 | 3 | 3 | 1 | 1 | 3 | 1 | 1 | 3 | 3 | 1 | 1 | 1 | 1 | 3 | 1 | 1 | 3 | 3 | 1 | 1 | 1 | 3 | 1 | 1 | 1 | 1 | 1 | 1 | 1 | 1 | 3 | 3 | 3 |
| yeps_YPTB0619 | 1 | 1 | 1 | 1 | 1 | 1 | 1 | 1 | 5 | 1 | 1 | 3 | 1 | 1 | 1 | 1 | 1 | 1 | 1 | 1 | 1 | 1 | 1 | 1 | 1 | 1 | 1 | 1 | 1 | 1 | 1 | 1 | 1 | 1 | 1 |
| yeps_YPTB0623 | 4 | 6 | 4 | 10 | 1 | 1 | 4 | 7 | 6 | 9 | 10 | 3 | 1 | 1 | 1 | 9 | 1 | 1 | 4 | 4 | 1 | 1 | 1 | 4 | 1 | 1 | 1 | 1 | 1 | 1 | 1 | 1 | 10 | 10 | 4 |
| yeps_YPTB0624 | 9 | 8 | 9 | 13 | 1 | 1 | 81 | 11 | 10 | 13 | 13 | 3 | 1 | 1 | 1 | 13 | 1 | 6 | 9 | 9 | 1 | 1 | 1 | 9 | 1 | 1 | 1 | 1 | 1 | 1 | 1 | 1 | 13 | 13 | 9 |
| yeps_YPTB0625 | 3 | 6 | 3 | 9 | 1 | 1 | 3 | 7 | 6 | 9 | 9 | 3 | 1 | 1 | 1 | 9 | 1 | 1 | 3 | 3 | 1 | 1 | 1 | 3 | 1 | 1 | 1 | 1 | 1 | 1 | 1 | 1 | 9 | 9 | 3 |
| yeps_YPTB0626 | 1 | 3 | 1 | 1 | 1 | 1 | 1 | 1 | 3 | 1 | 1 | 1 | 1 | 1 | 1 | 1 | 1 | 3 | 1 | 1 | 1 | 1 | 1 | 1 | 14 | 14 | 1 | 1 | 1 | 1 | 1 | 14 | 1 | 1 | 1 |
| yeps_YPTB0627 | 1 | 1 | 1 | 4 | 1 | 1 | 1 | 1 | 1 | 4 | 4 | 1 | 1 | 1 | 1 | 4 | 1 | 1 | 1 | 1 | 1 | 1 | 1 | 1 | 1 | 1 | 1 | 1 | 1 | 1 | 1 | 1 | 4 | 4 | 1 |
| yeps_YPTB0631 | 1 | 1 | 1 | 5 | 1 | 1 | 1 | 1 | 1 | 5 | 5 | 1 | 1 | 1 | 1 | 5 | 1 | 1 | 1 | 1 | 1 | 1 | 1 | 1 | 1 | 1 | 1 | 1 | 1 | 1 | 1 | 1 | 5 | 5 | 1 |
| yeps_YPTB0634 | 2 | 4 | 2 | 8 | 1 | 1 | 2 | 2 | 4 | 8 | 8 | 2 | 1 | 1 | 1 | 8 | 1 | 4 | 2 | 2 | 1 | 1 | 1 | 2 | 1 | 1 | 1 | 1 | 1 | 1 | 1 | 1 | 8 | 8 | 2 |
| yeps_YPTB0635 | 3 | 7 | 3 | 9 | 1 | 1 | 3 | 3 | 7 | 8 | 9 | 3 | 1 | 1 | 1 | 8 | 1 | 4 | 3 | 3 | 1 | 1 | 1 | 3 | 1 | 1 | 1 | 1 | 1 | 1 | 1 | 1 | 9 | 9 | 3 |
| yeps_YPTB0637 | 3 | 3 | 3 | 5 | 1 | 1 | 3 | 2 | 3 | 5 | 5 | 2 | 1 | 1 | 1 | 5 | 1 | 2 | 3 | 3 | 1 | 1 | 1 | 3 | 1 | 1 | 1 | 1 | 1 | 1 | 1 | 1 | 5 | 5 | 3 |
| yeps_YPTB0641 | 1 | 2 | 1 | 2 | 1 | 1 | 1 | 1 | 2 | 2 | 2 | 1 | 1 | 1 | 1 | 2 | 1 | 1 | 1 | 1 | 1 | 1 | 1 | 1 | 1 | 1 | 1 | 1 | 1 | 1 | 1 | 1 | 2 | 2 | 1 |
| yeps_YPTB0643 | 1 | 2 | 1 | 1 | 1 | 1 | 1 | 1 | 2 | 1 | 1 | 3 | 1 | 1 | 1 | 1 | 1 | 2 | 1 | 1 | 1 | 1 | 1 | 1 | 1 | 1 | 1 | 1 | 1 | 1 | 1 | 1 | 1 | 1 | 1 |
| yeps_YPTB0644 | 1 | 1 | 1 | 1 | 1 | 1 | 1 | 1 | 1 | 1 | 1 | 1 | 1 | 1 | 1 | 1 | 1 | 1 | 1 | 1 | 1 | 1 | 1 | 1 | 1 | 1 | 1 | 1 | 1 | 1 | 1 | 1 | 1 | 1 | 1 |
| yeps_YPTB0645 | 3 | 5 | 3 | 7 | 1 | 1 | 3 | 6 | 5 | 7 | 7 | 3 | 1 | 1 | 1 | 7 | 1 | 4 | 3 | 3 | 1 | 1 | 1 | 3 | 1 | 1 | 1 | 1 | 1 | 1 | 1 | 1 | 7 | 7 | 3 |
| yeps_YPTB0647 | 4 | 7 | 4 | 10 | 1 | 1 | 4 | 8 | 7 | 10 | 10 | 3 | 1 | 1 | 1 | 10 | 1 | 5 | 4 | 4 | 1 | 1 | 1 | 4 | 1 | 1 | 1 | 1 | 1 | 1 | 1 | 1 | 10 | 10 | 4 |
| yeps_YPTB0649 | 4 | 9 | 4 | 14 | 1 | 1 | 4 | 10 | 9 | 12 | 14 | 3 | 1 | 1 | 1 | 12 | 1 | 7 | 4 | 4 | 1 | 1 | 1 | 4 | 1 | 1 | 1 | 1 | 1 | 1 | 1 | 1 | 14 | 14 | 4 |
| yeps_YPTB0651 | 3 | 4 | 3 | 5 | 1 | 1 | 3 | 1 | 4 | 5 | 5 | 2 | 1 | 1 | 1 | 5 | 1 | 1 | 3 | 3 | 1 | 1 | 1 | 3 | 1 | 1 | 1 | 1 | 1 | 1 | 1 | 1 | 5 | 5 | 3 |
| yeps_YPTB0652 | 1 | 1 | 1 | 2 | 1 | 1 | 1 | 2 | 1 | 2 | 2 | 2 | 1 | 1 | 1 | 2 | 1 | 1 | 1 | 1 | 1 | 1 | 1 | 1 | 1 | 1 | 1 | 1 | 1 | 1 | 1 | 1 | 2 | 2 | 4 |
| yeps_YPTB0653 | 4 | 4 | 4 | 6 | 1 | 1 | 4 | 7 | 4 | 8 | 6 | 3 | 1 | 1 | 1 | 8 | 1 | 5 | 4 | 4 | 1 | 1 | 1 | 4 | 1 | 1 | 1 | 1 | 1 | 1 | 1 | 1 | 6 | 6 | 4 |
| yeps_YPTB0654 | 2 | 5 | 2 | 3 | 1 | 1 | 2 | 2 | 6 | 3 | 3 | 2 | 1 | 1 | 1 | 3 | 1 | 3 | 2 | 2 | 1 | 1 | 1 | 2 | 1 | 1 | 1 | 1 | 1 | 1 | 1 | 1 | 3 | 3 | 2 |
| yeps_YPTB0660 | 2 | 6 | 2 | 6 | 1 | 1 | 2 | 2 | 6 | 6 | 6 | 2 | 1 | 1 | 1 | 6 | 1 | 3 | 2 | 2 | 1 | 1 | 1 | 2 | 1 | 1 | 1 | 1 | 1 | 1 | 1 | 1 | 6 | 6 | 2 |
| yeps_YPTB0661 | 3 | 8 | 3 | 9 | 1 | 1 | 3 | 3 | 8 | 9 | 9 | 3 | 1 | 1 | 1 | 9 | 1 | 5 | 4 | 4 | 1 | 1 | 1 | 4 | 1 | 1 | 1 | 1 | 1 | 1 | 1 | 1 | 9 | 9 | 3 |
| yeps_YPTB0662 | 3 | 8 | 3 | 9 | 1 | 1 | 3 | 3 | 8 | 9 | 9 | 3 | 1 | 1 | 1 | 9 | 1 | 4 | 3 | 3 | 1 | 1 | 1 | 3 | 1 | 1 | 1 | 1 | 1 | 1 | 1 | 1 | 9 | 9 | 3 |
| yeps_YPTB0668 | 3 | 8 | 3 | 11 | 1 | 1 | 3 | 9 | 8 | 11 | 11 | 2 | 1 | 1 | 1 | 11 | 1 | 4 | 3 | 3 | 1 | 1 | 1 | 3 | 1 | 1 | 1 | 1 | 1 | 1 | 1 | 1 | 11 | 11 | 3 |
| yeps_YPTB0669 | 4 | 3 | 4 | 3 | 1 | 1 | 4 | 3 | 3 | 3 | 3 | 3 | 1 | 1 | 1 | 3 | 1 | 3 | 4 | 4 | 1 | 1 | 1 | 4 | 1 | 1 | 1 | 1 | 1 | 1 | 1 | 1 | 3 | 3 | 4 |
| yeps_YPTB0670 | 3 | 2 | 3 | 3 | 1 | 1 | 3 | 2 | 2 | 3 | 3 | 3 | 1 | 1 | 1 | 3 | 1 | 2 | 3 | 3 | 1 | 1 | 1 | 3 | 1 | 1 | 1 | 1 | 1 | 1 | 1 | 1 | 3 | 3 | 3 |
| yeps_YPTB0680 | 4 | 4 | 4 | 9 | 1 | 1 | 4 | 4 | 4 | 8 | 9 | 3 | 1 | 1 | 1 | 8 | 1 | 5 | 4 | 4 | 1 | 1 | 1 | 4 | 1 | 1 | 1 | 1 | 1 | 1 | 1 | 1 | 9 | 9 | 4 |
| yeps_YPTB0681 | 1 | 1 | 1 | 1 | 1 | 1 | 1 | 1 | 1 | 1 | 1 | 1 | 1 | 1 | 1 | 1 | 1 | 1 | 1 | 1 | 1 | 1 | 1 | 1 | 1 | 1 | 1 | 1 | 1 | 1 | 1 | 1 | 1 | 1 | 1 |
| yeps_YPTB0682 | 3 | 6 | 3 | 8 | 1 | 1 | 3 | 7 | 6 | 8 | 8 | 1 | 1 | 1 | 1 | 8 | 1 | 4 | 3 | 3 | 1 | 1 | 1 | 3 | 1 | 1 | 1 | 1 | 1 | 1 | 1 | 1 | 8 | 8 | 3 |
| yeps_YPTB0683 | 1 | 6 | 1 | 8 | 1 | 1 | 1 | 7 | 6 | 8 | 8 | 3 | 1 | 1 | 1 | 8 | 1 | 4 | 1 | 1 | 1 | 1 | 1 | 1 | 1 | 1 | 1 | 1 | 1 | 1 | 1 | 1 | 8 | 8 | 1 |
| yeps_YPTB0684 | 1 | 7 | 1 | 2 | 1 | 1 | 1 | 3 | 7 | 2 | 2 | 3 | 1 | 1 | 1 | 2 | 1 | 6 | 1 | 1 | 1 | 1 | 1 | 1 | 1 | 1 | 1 | 1 | 1 | 1 | 1 | 1 | 2 | 2 | 1 |
| yeps_YPTB0686 | 3 | 6 | 3 | 10 | 1 | 1 | 3 | 7 | 6 | 9 | 10 | 3 | 1 | 1 | 1 | 9 | 1 | 4 | 3 | 3 | 1 | 1 | 1 | 3 | 1 | 1 | 1 | 1 | 1 | 1 | 1 | 1 | 10 | 10 | 3 |
| yeps_YPTB0687 | 2 | 6 | 2 | 8 | 1 | 1 | 2 | 3 | 6 | 8 | 8 | 3 | 1 | 1 | 1 | 8 | 1 | 6 | 2 | 2 | 1 | 1 | 1 | 2 | 1 | 1 | 1 | 1 | 1 | 1 | 1 | 1 | 8 | 8 | 2 |
| yeps_YPTB0688 | 3 | 6 | 3 | 10 | 1 | 1 | 3 | 9 | 8 | 10 | 10 | 1 | 1 | 1 | 1 | 10 | 1 | 4 | 3 | 3 | 1 | 1 | 1 | 3 | 1 | 1 | 1 | 1 | 1 | 1 | 1 | 1 | 10 | 10 | 3 |
| yeps_YPTB0689 | 1 | 1 | 1 | 6 | 1 | 1 | 1 | 1 | 1 | 6 | 6 | 1 | 1 | 1 | 1 | 6 | 1 | 3 | 1 | 1 | 1 | 1 | 1 | 1 | 1 | 1 | 1 | 1 | 1 | 1 | 1 | 1 | 6 | 6 | 1 |
| yeps_YPTB0694 | 1 | 1 | 1 | 5 | 1 | 1 | 1 | 4 | 1 | 5 | 5 | 1 | 1 | 1 | 1 | 5 | 1 | 1 | 1 | 1 | 1 | 1 | 1 | 1 | 1 | 1 | 1 | 1 | 1 | 1 | 1 | 1 | 5 | 5 | 1 |
| yeps_YPTB0695 | 2 | 3 | 2 | 2 | 1 | 1 | 2 | 1 | 3 | 2 | 2 | 1 | 1 | 1 | 1 | 2 | 1 | 1 | 2 | 2 | 1 | 1 | 1 | 2 | 1 | 1 | 1 | 1 | 1 | 1 | 1 | 1 | 2 | 2 | 2 |
| yeps_YPTB0696 | 3 | 6 | 3 | 1 | 1 | 1 | 3 | 5 | 6 | 1 | 1 | 2 | 1 | 1 | 1 | 1 | 1 | 4 | 3 | 3 | 1 | 1 | 1 | 3 | 1 | 1 | 1 | 1 | 1 | 1 | 1 | 1 | 1 | 1 | 3 |
| yeps_YPTB0706 | 2 | 4 | 2 | 7 | 1 | 1 | 2 | 5 | 4 | 7 | 7 | 21 | 1 | 1 | 1 | 7 | 1 | 2 | 2 | 2 | 1 | 1 | 1 | 2 | 1 | 1 | 1 | 1 | 1 | 1 | 1 | 1 | 7 | 7 | 2 |
| yeps_YPTB0708 | 1 | 3 | 1 | 1 | 1 | 1 | 1 | 4 | 3 | 1 | 1 | 1 | 1 | 1 | 1 | 1 | 1 | 2 | 1 | 1 | 1 | 1 | 1 | 1 | 1 | 1 | 1 | 1 | 1 | 1 | 1 | 1 | 1 | 1 | 1 |
| yeps_YPTB0710 | 4 | 1 | 4 | 1 | 1 | 1 | 4 | 1 | 1 | 1 | 1 | 1 | 1 | 1 | 1 | 1 | 1 | 1 | 4 | 4 | 1 | 1 | 1 | 4 | 1 | 1 | 1 | 1 | 1 | 1 | 1 | 1 | 1 | 1 | 4 |
| yeps_YPTB0717 | 1 | 1 | 1 | 1 | 1 | 1 | 1 | 1 | 1 | 1 | 1 | 1 | 1 | 1 | 1 | 1 | 1 | 2 | 1 | 1 | 1 | 1 | 1 | 1 | 1 | 1 | 1 | 1 | 1 | 1 | 1 | 1 | 1 | 1 | 1 |
| yeps_YPTB0718 | 4 | 6 | 4 | 7 | 1 | 1 | 4 | 6 | 6 | 7 | 7 | 3 | 1 | 1 | 1 | 7 | 1 | 1 | 4 | 4 | 1 | 1 | 1 | 4 | 1 | 1 | 1 | 1 | 1 | 1 | 1 | 1 | 7 | 7 | 4 |
| yeps_YPTB0720 | 1 | 1 | 1 | 4 | 1 | 1 | 1 | 7 | 6 | 4 | 4 | 1 | 1 | 1 | 1 | 4 | 1 | 4 | 1 | 1 | 1 | 1 | 1 | 1 | 1 | 1 | 1 | 1 | 1 | 1 | 1 | 1 | 4 | 4 | 1 |
| yeps_YPTB0721 | 1 | 2 | 1 | 1 | 1 | 1 | 1 | 1 | 2 | 1 | 1 | 1 | 1 | 1 | 1 | 1 | 1 | 1 | 1 | 1 | 1 | 1 | 1 | 1 | 1 | 1 | 1 | 1 | 1 | 1 | 1 | 1 | 1 | 1 | 1 |
| yeps_YPTB0722 | 8 | 7 | 8 | 5 | 1 | 1 | 8 | 9 | 7 | 5 | 5 | 1 | 1 | 1 | 1 | 5 | 1 | 5 | 8 | 8 | 1 | 1 | 1 | 8 | 1 | 1 | 1 | 1 | 1 | 1 | 1 | 1 | 5 | 5 | 8 |
| yeps_YPTB0724 | 4 | 4 | 4 | 1 | 1 | 1 | 4 | 6 | 4 | 1 | 1 | 3 | 1 | 1 | 1 | 1 | 1 | 1 | 4 | 4 | 1 | 1 | 1 | 4 | 1 | 1 | 1 | 1 | 1 | 1 | 1 | 1 | 1 | 1 | 4 |
| yeps_YPTB0725 | 3 | 2 | 3 | 2 | 1 | 1 | 3 | 2 | 5 | 2 | 2 | 1 | 1 | 1 | 1 | 2 | 1 | 2 | 3 | 3 | 1 | 1 | 1 | 3 | 1 | 1 | 1 | 1 | 1 | 1 | 1 | 1 | 2 | 2 | 3 |
| yeps_YPTB0727 | 2 | 6 | 2 | 5 | 1 | 1 | 2 | 5 | 6 | 5 | 5 | 1 | 1 | 1 | 1 | 5 | 1 | 3 | 2 | 2 | 1 | 1 | 1 | 2 | 1 | 1 | 1 | 1 | 1 | 1 | 1 | 1 | 5 | 5 | 2 |
| yeps_YPTB0728 | 3 | 4 | 3 | 4 | 1 | 1 | 3 | 4 | 4 | 4 | 4 | 3 | 1 | 1 | 1 | 4 | 1 | 2 | 3 | 3 | 1 | 1 | 1 | 3 | 1 | 1 | 1 | 1 | 1 | 1 | 1 | 1 | 4 | 4 | 3 |
| yeps_YPTB0729 | 3 | 2 | 3 | 2 | 1 | 1 | 3 | 2 | 2 | 2 | 2 | 3 | 1 | 1 | 1 | 2 | 1 | 3 | 3 | 3 | 1 | 1 | 1 | 3 | 1 | 1 | 1 | 1 | 1 | 1 | 1 | 1 | 2 | 2 | 3 |
| yeps_YPTB0730 | 3 | 3 | 3 | 3 | 1 | 1 | 3 | 1 | 3 | 3 | 3 | 3 | 1 | 1 | 1 | 3 | 1 | 4 | 3 | 3 | 1 | 1 | 1 | 3 | 1 | 1 | 1 | 1 | 1 | 1 | 1 | 1 | 3 | 3 | 3 |
| yeps_YPTB0738 | 1 | 7 | 1 | 10 | 1 | 1 | 1 | 8 | 7 | 10 | 10 | 3 | 1 | 1 | 1 | 10 | 1 | 4 | 1 | 1 | 1 | 1 | 1 | 1 | 1 | 1 | 1 | 1 | 1 | 1 | 1 | 1 | 10 | 10 | 1 |
| yeps_YPTB0739 | 1 | 5 | 1 | 7 | 1 | 1 | 1 | 6 | 5 | 7 | 7 | 1 | 1 | 1 | 1 | 7 | 1 | 3 | 1 | 1 | 1 | 1 | 1 | 1 | 15 | 15 | 1 | 1 | 1 | 1 | 1 | 15 | 7 | 7 | 1 |
| yeps_YPTB0745 | 1 | 1 | 1 | 2 | 1 | 1 | 1 | 2 | 1 | 2 | 2 | 1 | 1 | 1 | 1 | 2 | 1 | 1 | 1 | 1 | 1 | 1 | 1 | 1 | 1 | 1 | 1 | 1 | 1 | 1 | 1 | 1 | 2 | 2 | 1 |
| yeps_YPTB0746 | 1 | 1 | 1 | 1 | 1 | 1 | 1 | 3 | 1 | 1 | 1 | 1 | 1 | 1 | 1 | 1 | 1 | 1 | 1 | 1 | 1 | 1 | 1 | 1 | 1 | 1 | 1 | 1 | 1 | 1 | 1 | 1 | 1 | 1 | 1 |
| yeps_YPTB0748 | 3 | 6 | 3 | 8 | 1 | 1 | 3 | 7 | 6 | 8 | 8 | 1 | 1 | 1 | 1 | 8 | 1 | 4 | 3 | 3 | 1 | 1 | 1 | 3 | 1 | 1 | 1 | 1 | 1 | 1 | 1 | 1 | 8 | 8 | 3 |
| yeps_YPTB0749 | 2 | 2 | 2 | 7 | 1 | 1 | 2 | 5 | 2 | 2 | 7 | 3 | 1 | 1 | 1 | 2 | 1 | 8 | 2 | 2 | 1 | 1 | 1 | 2 | 1 | 1 | 1 | 1 | 1 | 1 | 1 | 1 | 7 | 7 | 6 |
| yeps_YPTB0753 | 4 | 3 | 4 | 1 | 1 | 1 | 4 | 6 | 3 | 1 | 1 | 3 | 1 | 1 | 1 | 1 | 1 | 4 | 4 | 4 | 1 | 1 | 1 | 4 | 1 | 1 | 1 | 1 | 1 | 1 | 1 | 1 | 1 | 1 | 4 |
| yeps_YPTB0754 | 3 | 6 | 3 | 1 | 1 | 1 | 3 | 7 | 6 | 1 | 1 | 1 | 1 | 1 | 1 | 1 | 1 | 3 | 3 | 3 | 1 | 1 | 1 | 3 | 1 | 1 | 1 | 1 | 1 | 1 | 1 | 1 | 1 | 1 | 3 |
| yeps_YPTB0755 | 1 | 2 | 1 | 9 | 1 | 1 | 1 | 7 | 2 | 9 | 9 | 3 | 1 | 1 | 1 | 9 | 1 | 1 | 1 | 1 | 1 | 1 | 1 | 1 | 1 | 1 | 1 | 1 | 1 | 1 | 1 | 1 | 9 | 9 | 1 |
| yeps_YPTB0757 | 2 | 1 | 2 | 1 | 1 | 1 | 2 | 3 | 1 | 1 | 1 | 1 | 1 | 1 | 1 | 1 | 1 | 2 | 2 | 2 | 1 | 1 | 1 | 2 | 1 | 1 | 1 | 1 | 1 | 1 | 1 | 1 | 1 | 1 | 2 |
| yeps_YPTB0758 | 1 | 1 | 1 | 1 | 1 | 1 | 1 | 1 | 1 | 1 | 1 | 1 | 1 | 1 | 1 | 1 | 1 | 1 | 1 | 1 | 1 | 1 | 1 | 1 | 1 | 1 | 1 | 1 | 1 | 1 | 1 | 1 | 1 | 1 | 1 |

| | | | | | | | | | | | | | | | | | | | | | | | | | | | | | | | | | | | |
|---|---|---|---|---|---|---|---|---|---|---|---|---|---|---|---|---|---|---|---|---|---|---|---|---|---|---|---|---|---|---|---|---|---|---|---|
| yeps_YPTB0760 | 5 | 8 | 5 | 5 | 1 | 1 | 5 | 10 | 8 | 11 | 5 | 1 | 1 | 1 | 1 | 11 | 1 | 5 | 5 | 5 | 1 | 1 | 1 | 5 | 1 | 1 | 1 | 1 | 1 | 1 | 1 | 1 | 5 | 5 | 5 |
| yeps_YPTB0763 | 1 | 1 | 1 | 1 | 1 | 1 | 1 | 1 | 1 | 1 | 1 | 1 | 1 | 1 | 1 | 1 | 1 | 1 | 1 | 1 | 1 | 1 | 1 | 1 | 1 | 1 | 1 | 1 | 1 | 1 | 1 | 1 | 1 | 1 | 1 |
| yeps_YPTB0766 | 1 | 9 | 1 | 12 | 1 | 1 | 1 | 6 | 8 | 11 | 12 | 3 | 1 | 1 | 1 | 11 | 1 | 1 | 1 | 1 | 1 | 1 | 1 | 1 | 1 | 1 | 1 | 1 | 1 | 1 | 1 | 1 | 12 | 12 | 1 |
| yeps_YPTB0767 | 1 | 3 | 1 | 1 | 1 | 1 | 1 | 3 | 3 | 3 | 1 | 3 | 1 | 1 | 1 | 3 | 1 | 1 | 1 | 1 | 1 | 1 | 1 | 1 | 1 | 1 | 1 | 1 | 1 | 1 | 1 | 1 | 1 | 1 | 1 |
| yeps_YPTB0769 | 1 | 1 | 3 | 1 | 1 | 1 | 3 | 1 | 1 | 1 | 1 | 1 | 1 | 1 | 1 | 1 | 1 | 1 | 2 | 2 | 1 | 1 | 1 | 2 | 1 | 1 | 1 | 1 | 1 | 1 | 1 | 1 | 1 | 1 | 1 |
| yeps_YPTB0771 | 1 | 1 | 1 | 1 | 1 | 1 | 1 | 1 | 1 | 1 | 1 | 1 | 1 | 1 | 1 | 1 | 1 | 1 | 1 | 1 | 1 | 1 | 1 | 1 | 1 | 1 | 1 | 1 | 1 | 1 | 1 | 1 | 1 | 1 | 1 |
| yeps_YPTB0772 | 5 | 1 | 5 | 1 | 1 | 1 | 5 | 1 | 1 | 1 | 1 | 3 | 1 | 1 | 1 | 1 | 1 | 5 | 5 | 5 | 1 | 1 | 1 | 5 | 1 | 1 | 1 | 1 | 1 | 1 | 1 | 1 | 1 | 1 | 5 |
| yeps_YPTB0773 | 4 | 6 | 4 | 9 | 1 | 1 | 4 | 6 | 7 | 6 | 9 | 3 | 1 | 1 | 1 | 6 | 1 | 4 | 4 | 4 | 1 | 1 | 1 | 4 | 1 | 1 | 1 | 1 | 1 | 1 | 1 | 1 | 9 | 9 | 4 |
| yeps_YPTB0775 | 1 | 1 | 1 | 1 | 1 | 1 | 1 | 1 | 1 | 1 | 1 | 1 | 1 | 1 | 1 | 1 | 1 | 1 | 1 | 1 | 1 | 1 | 1 | 1 | 1 | 1 | 1 | 1 | 1 | 1 | 1 | 1 | 1 | 1 | 1 |
| yeps_YPTB0779 | 1 | 1 | 1 | 1 | 1 | 1 | 1 | 1 | 1 | 1 | 1 | 1 | 1 | 1 | 1 | 1 | 1 | 1 | 1 | 1 | 1 | 1 | 1 | 1 | 17 | 1 | 1 | 1 | 1 | 1 | 1 | 1 | 1 | 1 | 1 |
| yeps_YPTB0780 | 1 | 3 | 1 | 1 | 1 | 1 | 1 | 4 | 3 | 1 | 1 | 1 | 1 | 1 | 1 | 1 | 1 | 1 | 1 | 1 | 1 | 1 | 1 | 1 | 1 | 1 | 1 | 1 | 1 | 1 | 1 | 1 | 1 | 1 | 1 |
| yeps_YPTB0781 | 1 | 1 | 1 | 1 | 1 | 1 | 1 | 4 | 1 | 1 | 1 | 2 | 1 | 1 | 1 | 1 | 1 | 1 | 1 | 1 | 1 | 1 | 1 | 1 | 1 | 1 | 1 | 1 | 1 | 1 | 1 | 1 | 1 | 1 | 1 |
| yeps_YPTB0782 | 3 | 2 | 3 | 8 | 1 | 1 | 3 | 11 | 2 | 8 | 8 | 1 | 1 | 1 | 1 | 8 | 1 | 4 | 3 | 3 | 1 | 1 | 1 | 3 | 1 | 1 | 1 | 1 | 1 | 1 | 1 | 1 | 8 | 8 | 3 |
| yeps_YPTB0783 | 1 | 4 | 1 | 5 | 1 | 1 | 1 | 1 | 4 | 5 | 5 | 1 | 1 | 1 | 1 | 5 | 1 | 2 | 1 | 1 | 1 | 1 | 1 | 1 | 1 | 1 | 1 | 1 | 1 | 1 | 1 | 1 | 5 | 5 | 1 |
| yeps_YPTB0792 | 1 | 7 | 1 | 7 | 1 | 1 | 1 | 1 | 7 | 7 | 7 | 1 | 1 | 1 | 1 | 7 | 1 | 5 | 1 | 1 | 1 | 1 | 1 | 1 | 1 | 1 | 1 | 1 | 1 | 1 | 1 | 1 | 7 | 7 | 1 |
| yeps_YPTB0794 | 1 | 1 | 1 | 1 | 1 | 1 | 1 | 1 | 1 | 1 | 1 | 1 | 3 | 3 | 1 | 1 | 3 | 1 | 1 | 1 | 1 | 1 | 1 | 1 | 3 | 3 | 1 | 1 | 1 | 1 | 1 | 3 | 1 | 1 | 1 |
| yeps_YPTB0795 | 1 | 1 | 1 | 1 | 1 | 1 | 1 | 1 | 7 | 1 | 1 | 1 | 3 | 3 | 1 | 1 | 3 | 4 | 1 | 1 | 1 | 1 | 1 | 1 | 3 | 3 | 1 | 1 | 1 | 1 | 1 | 3 | 1 | 1 | 1 |
| yeps_YPTB0798 | 1 | 7 | 1 | 9 | 1 | 1 | 1 | 3 | 7 | 9 | 9 | 3 | 4 | 4 | 1 | 9 | 4 | 5 | 1 | 1 | 1 | 1 | 1 | 1 | 4 | 4 | 1 | 1 | 1 | 1 | 1 | 4 | 9 | 9 | 1 |
| yeps_YPTB0800 | 1 | 7 | 1 | 10 | 1 | 1 | 1 | 8 | 7 | 10 | 10 | 3 | 1 | 1 | 1 | 10 | 1 | 11 | 1 | 1 | 1 | 1 | 1 | 1 | 1 | 1 | 1 | 1 | 1 | 1 | 1 | 1 | 10 | 10 | 1 |
| yeps_YPTB0802 | 5 | 5 | 5 | 3 | 1 | 4 | 5 | 5 | 5 | 3 | 3 | 3 | 4 | 4 | 4 | 3 | 4 | 3 | 5 | 5 | 4 | 4 | 4 | 5 | 4 | 4 | 4 | 4 | 4 | 4 | 4 | 4 | 3 | 3 | 5 |
| yeps_YPTB0803 | 3 | 7 | 3 | 10 | 1 | 1 | 3 | 8 | 7 | | 10 | 3 | 1 | 1 | 1 | 10 | 1 | 4 | 3 | 3 | 1 | 1 | 1 | 3 | 1 | 1 | 1 | 1 | 1 | 1 | 1 | 1 | 10 | 10 | 3 |
| yeps_YPTB0806 | 1 | 1 | 1 | 5 | 1 | 1 | 1 | 4 | 1 | 5 | 5 | 3 | 3 | 3 | 1 | 5 | 3 | 4 | 1 | 1 | 1 | 1 | 1 | 1 | 3 | 3 | 1 | 1 | 1 | 1 | 1 | 3 | 5 | 5 | 1 |
| yeps_YPTB0807 | 5 | 9 | 5 | 9 | 1 | 1 | 5 | 13 | 9 | 11 | 9 | 3 | 4 | 4 | 1 | 11 | 4 | 6 | 5 | 5 | 1 | 1 | 1 | 5 | 4 | 4 | 1 | 1 | 1 | 1 | 1 | 4 | 9 | 9 | 5 |
| yeps_YPTB0811 | 1 | 7 | 1 | 12 | 1 | 1 | 1 | 8 | 7 | 10 | 12 | 3 | 3 | 3 | 1 | 10 | 3 | 4 | 1 | 1 | 1 | 1 | 1 | 1 | 3 | 3 | 1 | 1 | 1 | 1 | 1 | 3 | 12 | 12 | 1 |
| yeps_YPTB0812 | 1 | 7 | 1 | 11 | 1 | 1 | 1 | 9 | 7 | 11 | 11 | 3 | 3 | 3 | 1 | 11 | 3 | 4 | 1 | 1 | 1 | 1 | 1 | 1 | 3 | 3 | 1 | 1 | 1 | 1 | 1 | 3 | 11 | 11 | 1 |
| yeps_YPTB0813 | 1 | 1 | 1 | 1 | 1 | 1 | 1 | 4 | 1 | 1 | 1 | 1 | 1 | 1 | 1 | 1 | 1 | 1 | 1 | 1 | 1 | 1 | 1 | 1 | 1 | 1 | 1 | 1 | 1 | 1 | 1 | 1 | 1 | 1 | 1 |
| yeps_YPTB0814 | 10 | 8 | 4 | 8 | 1 | 1 | 4 | 8 | 8 | 8 | 8 | 3 | 3 | 3 | 1 | 8 | 3 | 4 | 4 | 4 | 1 | 1 | 1 | 4 | 3 | 3 | 1 | 1 | 1 | 1 | 1 | 3 | 8 | 8 | 4 |
| yeps_YPTB0816 | 3 | 5 | 3 | 7 | 1 | 1 | 3 | 5 | 5 | 5 | 7 | 1 | 1 | 1 | 1 | 5 | 1 | 4 | 3 | 3 | 1 | 1 | 1 | 3 | 1 | 1 | 1 | 1 | 1 | 1 | 1 | 1 | 7 | 7 | 3 |
| yeps_YPTB0819 | 4 | 6 | 4 | 9 | 1 | 1 | 4 | 7 | 6 | 9 | 9 | 3 | 1 | 1 | 1 | 9 | 1 | 4 | 4 | 4 | 1 | 1 | 1 | 4 | 1 | 1 | 1 | 1 | 1 | 1 | 1 | 1 | 9 | 9 | 4 |
| yeps_YPTB0820 | 3 | 3 | 3 | 6 | 1 | 1 | 3 | 2 | 5 | 6 | 6 | 3 | 1 | 1 | 1 | 6 | 1 | 3 | 3 | 3 | 1 | 1 | 1 | 3 | 1 | 1 | 1 | 1 | 1 | 1 | 1 | 1 | 6 | 6 | 3 |
| yeps_YPTB0821 | 3 | 5 | 3 | 5 | 1 | 1 | 3 | 7 | 5 | 5 | 5 | 1 | 1 | 1 | 1 | 5 | 1 | 3 | 3 | 3 | 1 | 1 | 1 | 3 | 1 | 1 | 1 | 1 | 1 | 1 | 1 | 1 | 5 | 5 | 3 |
| yeps_YPTB0822 | 1 | 3 | 1 | 3 | 1 | 1 | 1 | 1 | 3 | 3 | 3 | 1 | 1 | 1 | 1 | 3 | 1 | 1 | 1 | 1 | 1 | 1 | 1 | 1 | 1 | 1 | 1 | 1 | 1 | 1 | 1 | 1 | 3 | 3 | 1 |
| yeps_YPTB0825 | 14 | 8 | 4 | 12 | 1 | 1 | 4 | 10 | 8 | 12 | 12 | 3 | 13 | 6 | 1 | 12 | 6 | 4 | 4 | 4 | 1 | 1 | 1 | 4 | 1 | 1 | 1 | 1 | 1 | 1 | 1 | 1 | 12 | 12 | 4 |
| yeps_YPTB0828 | 1 | 1 | 2 | 1 | 1 | 1 | 2 | 1 | 1 | 1 | 1 | 1 | 1 | 1 | 1 | 1 | 1 | 1 | 2 | 2 | 1 | 1 | 1 | 2 | 1 | 1 | 1 | 1 | 1 | 1 | 1 | 1 | 1 | 1 | 2 |
| yeps_YPTB0829 | 1 | 7 | 1 | 13 | 1 | 1 | 1 | 9 | 7 | 11 | 13 | 3 | 1 | 1 | 1 | 11 | 1 | 1 | 1 | 1 | 1 | 1 | 1 | 1 | 1 | 1 | 1 | 1 | 1 | 1 | 1 | 1 | 13 | 13 | 12 |
| yeps_YPTB0832 | 1 | 1 | 1 | 7 | 1 | 1 | 1 | 6 | 1 | 7 | 7 | 1 | 1 | 1 | 1 | 7 | 1 | 4 | 1 | 1 | 1 | 1 | 1 | 1 | 1 | 1 | 1 | 1 | 1 | 1 | 1 | 1 | 7 | 7 | 1 |
| yeps_YPTB0836 | 1 | 1 | 1 | 2 | 1 | 1 | 1 | 1 | 1 | 2 | 2 | 1 | 1 | 1 | 1 | 2 | 1 | 2 | 1 | 1 | 1 | 1 | 1 | 1 | 1 | 1 | 1 | 1 | 1 | 1 | 1 | 1 | 2 | 2 | 1 |
| yeps_YPTB0850 | 3 | 2 | 3 | 2 | 1 | 1 | 3 | 4 | 2 | 2 | 2 | 3 | 3 | 3 | 1 | 2 | 3 | 2 | 3 | 3 | 1 | 1 | 1 | 3 | 3 | 3 | 1 | 1 | 1 | 1 | 1 | 3 | 2 | 2 | 3 |
| yeps_YPTB0853 | 1 | 6 | 1 | 4 | 1 | 1 | 1 | 7 | 6 | 4 | 4 | 3 | 4 | 4 | 1 | 4 | 4 | 3 | 1 | 1 | 1 | 1 | 1 | 1 | 4 | 4 | 1 | 1 | 1 | 1 | 1 | 4 | 4 | 4 | 1 |
| yeps_YPTB0856 | 3 | 6 | 3 | 1 | 1 | 1 | 3 | 7 | 6 | 1 | 1 | 3 | 1 | 1 | 1 | 1 | 1 | 3 | 3 | 3 | 1 | 1 | 1 | 3 | 1 | 1 | 1 | 1 | 1 | 1 | 1 | 1 | 1 | 1 | 3 |
| yeps_YPTB0860 | 1 | 6 | 1 | 9 | 1 | 1 | 1 | 8 | 6 | 9 | 9 | 1 | 1 | 1 | 1 | 9 | 1 | 3 | 1 | 1 | 1 | 1 | 1 | 1 | 1 | 1 | 1 | 1 | 1 | 1 | 1 | 1 | 9 | 9 | 10 |
| yeps_YPTB0862 | 4 | 8 | 4 | 11 | 1 | 1 | 4 | 9 | 8 | 11 | 11 | 3 | 1 | 1 | 1 | 11 | 1 | 5 | 4 | 4 | 1 | 1 | 1 | 4 | 1 | 1 | 1 | 1 | 1 | 1 | 1 | 1 | 11 | 11 | 4 |
| yeps_YPTB0863 | 1 | 3 | 1 | 4 | 1 | 1 | 1 | 1 | 3 | 4 | 4 | 1 | 1 | 1 | 1 | 4 | 1 | 2 | 1 | 1 | 1 | 1 | 1 | 1 | 1 | 1 | 1 | 1 | 1 | 1 | 1 | 1 | 4 | 4 | 1 |
| yeps_YPTB0866 | 4 | 8 | 4 | 9 | 1 | 1 | 4 | 4 | 8 | 9 | 9 | 3 | 1 | 1 | 1 | 9 | 1 | 5 | 4 | 4 | 1 | 1 | 1 | 4 | 1 | 1 | 1 | 1 | 1 | 1 | 1 | 1 | 9 | 9 | 4 |
| yeps_YPTB0867 | 1 | 3 | 1 | 4 | 1 | 1 | 1 | 1 | 3 | 4 | 4 | 3 | 7 | 1 | 1 | 4 | 1 | 4 | 1 | 1 | 1 | 1 | 1 | 1 | 1 | 1 | 1 | 1 | 1 | 1 | 1 | 1 | 4 | 4 | 1 |
| yeps_YPTB0868 | 1 | 5 | 1 | 4 | 1 | 1 | 1 | 6 | 5 | 4 | 4 | 3 | 1 | 1 | 1 | 4 | 1 | 4 | 1 | 1 | 1 | 1 | 1 | 1 | 1 | 1 | 1 | 1 | 1 | 1 | 1 | 1 | 4 | 4 | 1 |
| yeps_YPTB0869 | 1 | 4 | 1 | 6 | 1 | 1 | 1 | 5 | 4 | 6 | 6 | 3 | 1 | 1 | 1 | 6 | 1 | 1 | 1 | 1 | 1 | 1 | 1 | 1 | 1 | 1 | 1 | 1 | 1 | 1 | 1 | 1 | 6 | 6 | 1 |
| yeps_YPTB0881 | 4 | 8 | 4 | 12 | 1 | 1 | 4 | 9 | 8 | 11 | 12 | 1 | 1 | 5 | 1 | 11 | 5 | 4 | 4 | 4 | 1 | 1 | 1 | 4 | 1 | 1 | 1 | 1 | 1 | 1 | 1 | 1 | 12 | 12 | 4 |
| yeps_YPTB0883 | 4 | 1 | 4 | 10 | 1 | 1 | 4 | 8 | 52 | 10 | 10 | 3 | 1 | 1 | 1 | 10 | 1 | 11 | 4 | 4 | 1 | 1 | 1 | 4 | 1 | 1 | 1 | 1 | 1 | 1 | 1 | 1 | 10 | 10 | 4 |
| yeps_YPTB0884 | 2 | 1 | 2 | 1 | 1 | 1 | 2 | 2 | 1 | 1 | 1 | 2 | 1 | 1 | 1 | 1 | 1 | 1 | 2 | 2 | 1 | 1 | 1 | 2 | 1 | 1 | 1 | 1 | 1 | 1 | 1 | 1 | 1 | 1 | 2 |
| yeps_YPTB0885 | 3 | 4 | 3 | 5 | 1 | 1 | 3 | 1 | 4 | 5 | 5 | 1 | 1 | 1 | 1 | 5 | 1 | 1 | 3 | 3 | 1 | 1 | 1 | 3 | 1 | 1 | 1 | 1 | 1 | 1 | 1 | 1 | 5 | 5 | 3 |
| yeps_YPTB0886 | 3 | 6 | 3 | 8 | 1 | 1 | 3 | 7 | 6 | 8 | 8 | 1 | 1 | 1 | 1 | 8 | 1 | 4 | 3 | 3 | 1 | 1 | 1 | 3 | 1 | 1 | 1 | 1 | 1 | 1 | 1 | 1 | 8 | 8 | 3 |
| yeps_YPTB0888 | 1 | 1 | 1 | 1 | 1 | 1 | 1 | 1 | 1 | 1 | 1 | 1 | 1 | 1 | 1 | 1 | 1 | 1 | 1 | 1 | 1 | 1 | 1 | 1 | 1 | 1 | 1 | 1 | 1 | 1 | 1 | 1 | 1 | 1 | 1 |
| yeps_YPTB0891 | 1 | 1 | 1 | 1 | 1 | 1 | 1 | 1 | 1 | 1 | 1 | 1 | 1 | 1 | 1 | 1 | 1 | 1 | 1 | 1 | 1 | 1 | 1 | 1 | 1 | 1 | 1 | 1 | 1 | 1 | 1 | 1 | 1 | 1 | 1 |
| yeps_YPTB0893 | 4 | 8 | 4 | 8 | 1 | 1 | 4 | 4 | 8 | 8 | 8 | 3 | 1 | 1 | 1 | 8 | 1 | 4 | 4 | 4 | 1 | 1 | 1 | 4 | 1 | 1 | 1 | 1 | 1 | 1 | 1 | 1 | 8 | 8 | 4 |
| yeps_YPTB0912 | 1 | 1 | 1 | 1 | 1 | 1 | 1 | 1 | 1 | 1 | 1 | 1 | 2 | 2 | 1 | 1 | 2 | 1 | 1 | 1 | 1 | 1 | 1 | 1 | 2 | 2 | 1 | 1 | 1 | 1 | 1 | 2 | 1 | 1 | 1 |
| yeps_YPTB0914 | 10 | 3 | 3 | 3 | 1 | 1 | 3 | 31 | 3 | 3 | 3 | 1 | 3 | 3 | 183 | 3 | 3 | 3 | 3 | 3 | 1 | 1 | 1 | 28 | 3 | 3 | 1 | 1 | 1 | 1 | 50 | 28 | 180 | 3 | 3 |
| yeps_YPTB0916 | 1 | 3 | 1 | 7 | 1 | 1 | 1 | 1 | 3 | 7 | 7 | 1 | 1 | 1 | 1 | 7 | 1 | 8 | 1 | 1 | 1 | 1 | 1 | 1 | 1 | 1 | 1 | 1 | 1 | 1 | 1 | 1 | 7 | 7 | 1 |
| yeps_YPTB0917 | 1 | 1 | 1 | 1 | 1 | 1 | 1 | 1 | 1 | 1 | 1 | 1 | 1 | 1 | 1 | 1 | 1 | 1 | 1 | 1 | 1 | 1 | 1 | 1 | 1 | 1 | 1 | 1 | 1 | 1 | 1 | 1 | 1 | 1 | 1 |
| yeps_YPTB0918 | 3 | 7 | 3 | 4 | 1 | 1 | 3 | 3 | 7 | 4 | 4 | 1 | 1 | 1 | 1 | 4 | 1 | 4 | 3 | 3 | 1 | 1 | 1 | 3 | 1 | 1 | 1 | 1 | 1 | 1 | 1 | 1 | 4 | 4 | 3 |
| yeps_YPTB0923 | 3 | 5 | 3 | 5 | 1 | 1 | 3 | 1 | 15 | 5 | 5 | 1 | 1 | 1 | 1 | 5 | 1 | 1 | 3 | 3 | 1 | 1 | 1 | 3 | 1 | 1 | 1 | 1 | 1 | 1 | 1 | 1 | 5 | 5 | 3 |
| yeps_YPTB0927 | 1 | 7 | 1 | 9 | 1 | 1 | 1 | 8 | 7 | 9 | 9 | 3 | 1 | 1 | 1 | 9 | 1 | 5 | 1 | 1 | 1 | 1 | 1 | 1 | 1 | 1 | 1 | 1 | 1 | 1 | 1 | 1 | 9 | 9 | 1 |
| yeps_YPTB0929 | 1 | 1 | 1 | 1 | 1 | 1 | 1 | 1 | 1 | 1 | 1 | 1 | 1 | 1 | 1 | 1 | 1 | 1 | 1 | 1 | 1 | 1 | 1 | 1 | 1 | 1 | 1 | 1 | 1 | 1 | 1 | 1 | 1 | 1 | 1 |
| yeps_YPTB0935 | 1 | 1 | 1 | 1 | 1 | 1 | 1 | 1 | 1 | 1 | 1 | 1 | 1 | 1 | 1 | 1 | 1 | 1 | 1 | 1 | 1 | 1 | 1 | 1 | 1 | 1 | 1 | 1 | 1 | 1 | 1 | 1 | 1 | 1 | 1 |
| yeps_YPTB0937 | 3 | 5 | 3 | 5 | 1 | 1 | 3 | 6 | 5 | 5 | 5 | 3 | 1 | 1 | 1 | 5 | 1 | 3 | 3 | 3 | 1 | 1 | 1 | 3 | 1 | 1 | 1 | 1 | 1 | 1 | 1 | 1 | 5 | 5 | 3 |
| yeps_YPTB0938 | 3 | 1 | 3 | 6 | 1 | 1 | 3 | 1 | 1 | 6 | 6 | 2 | 1 | 1 | 1 | 6 | 1 | 4 | 3 | 3 | 1 | 1 | 1 | 3 | 1 | 1 | 1 | 1 | 1 | 1 | 1 | 1 | 6 | 6 | 3 |
| yeps_YPTB0939 | 4 | 1 | 4 | 7 | 1 | 1 | 4 | 7 | 1 | 7 | 7 | 3 | 1 | 1 | 1 | 7 | 1 | 4 | 4 | 4 | 1 | 1 | 1 | 4 | 1 | 1 | 1 | 1 | 1 | 1 | 1 | 1 | 7 | 7 | 4 |
| yeps_YPTB0940 | 3 | 5 | 3 | 7 | 1 | 1 | 3 | 6 | 5 | 7 | 7 | 2 | 1 | 1 | 1 | 7 | 1 | 4 | 3 | 3 | 1 | 1 | 1 | 3 | 1 | 1 | 1 | 1 | 1 | 1 | 1 | 1 | 7 | 7 | 3 |
| yeps_YPTB0941 | 1 | 1 | 1 | 1 | 1 | 1 | 1 | 1 | 1 | 1 | 1 | 1 | 1 | 1 | 1 | 1 | 1 | 1 | 1 | 1 | 1 | 1 | 1 | 1 | 1 | 1 | 1 | 1 | 1 | 1 | 1 | 1 | 1 | 1 | 1 |
| yeps_YPTB0942 | 1 | 1 | 1 | 1 | 1 | 1 | 1 | 1 | 1 | 1 | 1 | 1 | 1 | 1 | 1 | 1 | 1 | 1 | 1 | 1 | 1 | 1 | 1 | 1 | 1 | 1 | 1 | 1 | 1 | 1 | 1 | 1 | 1 | 1 | 1 |
| yeps_YPTB0943 | 8 | 6 | 4 | 7 | 1 | 1 | 4 | 7 | 6 | 7 | 7 | 3 | 1 | 1 | 1 | 7 | 1 | 5 | 4 | 4 | 1 | 1 | 1 | 4 | 1 | 1 | 1 | 1 | 1 | 1 | 1 | 1 | 7 | 7 | 4 |
| yeps_YPTB0944 | 3 | 3 | 3 | 3 | 1 | 1 | 3 | 3 | 3 | 3 | 3 | 3 | 1 | 1 | 1 | 3 | 1 | 3 | 3 | 3 | 1 | 1 | 1 | 3 | 1 | 1 | 1 | 1 | 1 | 1 | 1 | 1 | 3 | 3 | 3 |
| yeps_YPTB0945 | 4 | 2 | 4 | 2 | 1 | 1 | 4 | 2 | 2 | 2 | 2 | 3 | 1 | 1 | 1 | 2 | 1 | 2 | 4 | 4 | 1 | 1 | 1 | 4 | 1 | 1 | 1 | 1 | 1 | 1 | 1 | 1 | 2 | 2 | 4 |
| yeps_YPTB0946 | 1 | 2 | 1 | 2 | 1 | 1 | 1 | 2 | 2 | 3 | 2 | 1 | 1 | 1 | 1 | 3 | 1 | 2 | 1 | 1 | 1 | 1 | 1 | 1 | 1 | 1 | 1 | 1 | 1 | 1 | 1 | 1 | 2 | 2 | 1 |
| yeps_YPTB0949 | 2 | 3 | 2 | 2 | 1 | 1 | 2 | 2 | 2 | 2 | 2 | 2 | 1 | 1 | 1 | 2 | 1 | 2 | 2 | 2 | 1 | 1 | 1 | 2 | 1 | 1 | 1 | 1 | 1 | 1 | 1 | 1 | 2 | 2 | 2 |
| yeps_YPTB0954 | 1 | 4 | 1 | 5 | 1 | 1 | 1 | 1 | 4 | 5 | 5 | 1 | 1 | 1 | 1 | 5 | 1 | 2 | 1 | 1 | 1 | 1 | 1 | 1 | 1 | 1 | 1 | 1 | 1 | 1 | 1 | 1 | 5 | 5 | 1 |
| yeps_YPTB0956 | 1 | 1 | 1 | 1 | 1 | 1 | 1 | 1 | 1 | 2 | 1 | 1 | 1 | 1 | 1 | 2 | 1 | 1 | 1 | 1 | 1 | 1 | 1 | 1 | 1 | 1 | 1 | 1 | 1 | 1 | 1 | 1 | 1 | 1 | 1 |
| yeps_YPTB0957 | 1 | 1 | 1 | 1 | 1 | 1 | 1 | 1 | 1 | 1 | 1 | 1 | 1 | 1 | 1 | 1 | 1 | 1 | 1 | 1 | 1 | 1 | 1 | 1 | 1 | 1 | 1 | 1 | 1 | 1 | 1 | 1 | 1 | 1 | 1 |
| yeps_YPTB0965 | 1 | 2 | 1 | 1 | 1 | 1 | 1 | 3 | 2 | 4 | 1 | 1 | 1 | 1 | 1 | 4 | 1 | 1 | 1 | 1 | 1 | 1 | 1 | 1 | 1 | 1 | 1 | 1 | 1 | 1 | 1 | 1 | 1 | 1 | 1 |
| yeps_YPTB0967 | 4 | 8 | 4 | 11 | 1 | 1 | 4 | 9 | 8 | 11 | 11 | 3 | 1 | 1 | 1 | 11 | 1 | 4 | 4 | 4 | 1 | 1 | 1 | 4 | 1 | 1 | 1 | 1 | 1 | 1 | 1 | 1 | 11 | 11 | 4 |
| yeps_YPTB0968 | 5 | 7 | 5 | 8 | 5 | 5 | 26 | 3 | 7 | 8 | 8 | 3 | 5 | 5 | 5 | 8 | 5 | 5 | 5 | 5 | 5 | 5 | 5 | 5 | 5 | 5 | 5 | 5 | 5 | 5 | 5 | 5 | 8 | 8 | 5 |
| yeps_YPTB0973 | 1 | 1 | 1 | 1 | 1 | 1 | 1 | 1 | 1 | 1 | 1 | 1 | 1 | 1 | 1 | 1 | 1 | 1 | 1 | 1 | 1 | 1 | 1 | 1 | 1 | 1 | 1 | 1 | 1 | 1 | 1 | 1 | 1 | 1 | 1 |
| yeps_YPTB0974 | 2 | 1 | 2 | 5 | 1 | 1 | 2 | 4 | 1 | 5 | 5 | 1 | 1 | 1 | 1 | 5 | 1 | 2 | 2 | 2 | 1 | 1 | 1 | 2 | 1 | 1 | 1 | 1 | 1 | 1 | 1 | 1 | 5 | 5 | 2 |
| yeps_YPTB0975 | 3 | 3 | 3 | 3 | 1 | 1 | 3 | 4 | 3 | 3 | 3 | 3 | 1 | 1 | 1 | 3 | 1 | 3 | 3 | 3 | 1 | 1 | 1 | 3 | 1 | 1 | 1 | 1 | 1 | 1 | 1 | 1 | 3 | 3 | 3 |
| yeps_YPTB0976 | 1 | 4 | 1 | 1 | 1 | 1 | 1 | 6 | 4 | 7 | 1 | 1 | 1 | 1 | 1 | 7 | 1 | 1 | 1 | 1 | 1 | 1 | 1 | 1 | 1 | 1 | 1 | 1 | 1 | 1 | 1 | 1 | 1 | 1 | 1 |
| yeps_YPTB0986 | 1 | 1 | 1 | 5 | 1 | 1 | 1 | 3 | 1 | 5 | 5 | 1 | 1 | 1 | 1 | 5 | 1 | 1 | 1 | 1 | 1 | 1 | 1 | 1 | 1 | 1 | 1 | 1 | 1 | 1 | 1 | 1 | 5 | 5 | 1 |
| yeps_YPTB0992 | 3 | 7 | 5 | 9 | 1 | 1 | 5 | 3 | 7 | 9 | 9 | 3 | 4 | 4 | 1 | 9 | 4 | 3 | 5 | 5 | 1 | 1 | 1 | 5 | 4 | 4 | 1 | 1 | 1 | 1 | 1 | 4 | 9 | 9 | 3 |
| yeps_YPTB1024 | 2 | 6 | 2 | 2 | 1 | 1 | 2 | 2 | 6 | 2 | 2 | 2 | 1 | 1 | 1 | 2 | 1 | 2 | 2 | 2 | 1 | 1 | 1 | 2 | 1 | 1 | 1 | 1 | 1 | 1 | 1 | 1 | 2 | 2 | 2 |
| yeps_YPTB1026 | 3 | 3 | 3 | 2 | 1 | 1 | 3 | 4 | 3 | 2 | 2 | 1 | 1 | 1 | 1 | 2 | 1 | 3 | 3 | 3 | 1 | 1 | 1 | 3 | 1 | 1 | 1 | 1 | 1 | 1 | 1 | 1 | 2 | 2 | 3 |
| yeps_YPTB1028 | 2 | 2 | 2 | 2 | 1 | 1 | 2 | 2 | 2 | 4 | 2 | 1 | 1 | 1 | 1 | 4 | 1 | 2 | 2 | 2 | 1 | 1 | 1 | 2 | 1 | 1 | 1 | 1 | 1 | 1 | 1 | 1 | 2 | 2 | 2 |
| yeps_YPTB1030 | 4 | 8 | 4 | 14 | 1 | 1 | 4 | 13 | 8 | 12 | 14 | 3 | 1 | 1 | 1 | 12 | 1 | 4 | 95 | 4 | 1 | 1 | 1 | 4 | 1 | 1 | 1 | 1 | 1 | 1 | 1 | 1 | 14 | 14 | 4 |
| yeps_YPTB1031 | 5 | 8 | 5 | 13 | 1 | 1 | 5 | 10 | 8 | 12 | 13 | 3 | 1 | 1 | 1 | 12 | 1 | 5 | 5 | 5 | 1 | 1 | 1 | 5 | 1 | 1 | 1 | 1 | 1 | 1 | 1 | 1 | 13 | 13 | 5 |
| yeps_YPTB1032 | 4 | 3 | 4 | 3 | 1 | 1 | 4 | 5 | 3 | 3 | 3 | 3 | 1 | 1 | 1 | 3 | 1 | 4 | 4 | 4 | 1 | 1 | 1 | 4 | 1 | 1 | 1 | 1 | 1 | 1 | 1 | 1 | 3 | 3 | 4 |
| yeps_YPTB1035 | 11 | 7 | 4 | 10 | 1 | 1 | 4 | 8 | 7 | 10 | 10 | 3 | 1 | 1 | 1 | 10 | 1 | 12 | 4 | 4 | 1 | 1 | 1 | 4 | 1 | 1 | 1 | 1 | 1 | 1 | 1 | 1 | 10 | 10 | 4 |
| yeps_YPTB1036 | 1 | 1 | 1 | 1 | 1 | 1 | 1 | 1 | 1 | 1 | 1 | 1 | 1 | 1 | 1 | 1 | 1 | 1 | 1 | 1 | 1 | 1 | 1 | 1 | 1 | 1 | 1 | 1 | 1 | 1 | 1 | 1 | 1 | 1 | 1 |
| yeps_YPTB1037 | 1 | 1 | 1 | 2 | 1 | 1 | 1 | 1 | 1 | 2 | 2 | 1 | 1 | 1 | 1 | 2 | 1 | 2 | 1 | 1 | 1 | 1 | 1 | 1 | 1 | 1 | 1 | 1 | 1 | 1 | 1 | 1 | 2 | 2 | 1 |
| yeps_YPTB1038 | 1 | | 1 | 1 | 1 | 1 | 1 | 1 | 2 | 1 | 1 | 1 | 1 | 1 | 1 | 1 | 1 | 1 | 1 | 1 | 1 | 1 | 1 | 1 | 1 | 1 | 1 | 1 | 1 | 1 | 1 | 1 | 1 | 1 | 1 |
| yeps_YPTB1040 | 4 | | 4 | 7 | 1 | 1 | 4 | 8 | 7 | 10 | 7 | 3 | 1 | 1 | 1 | 10 | 1 | 5 | 4 | 4 | 1 | 1 | 1 | 4 | 1 | 1 | 1 | 1 | 1 | 1 | 1 | 1 | 7 | 7 | 4 |
| yeps_YPTB1041 | 1 | | 1 | 1 | 1 | 1 | 1 | 1 | 1 | 1 | 1 | 1 | 1 | 1 | 1 | 1 | 1 | 1 | 1 | 1 | 1 | 1 | 1 | 1 | 1 | 1 | 1 | 1 | 1 | 1 | 1 | 1 | 1 | 1 | 1 |
| yeps_YPTB1043 | 3 | | 3 | | 1 | 1 | 3 | 2 | 3 | 2 | | 3 | 1 | 1 | 1 | 2 | 1 | 3 | 3 | 3 | 1 | 1 | 1 | 3 | 1 | 1 | 1 | 1 | 1 | 1 | 1 | 1 | | | 3 |
| yeps_YPTB1044 | 1 | | 1 | 3 | 1 | 1 | 1 | 1 | 3 | 3 | 3 | 3 | 1 | 1 | 1 | 3 | 1 | 4 | 1 | 1 | 1 | 1 | 1 | 1 | 1 | 1 | 1 | 1 | 1 | 1 | 1 | 1 | 3 | 3 | 1 |
| yeps_YPTB1045 | 3 | | 3 | 8 | 1 | 1 | 3 | 3 | 6 | 8 | 8 | 1 | 1 | 1 | 1 | 8 | 1 | 3 | 3 | 3 | 1 | 1 | 1 | 3 | 1 | 1 | 1 | 1 | 1 | 1 | 1 | 1 | 8 | 8 | 3 |
| yeps_YPTB1047 | 3 | | 3 | 10 | 1 | 1 | 3 | 9 | 7 | 10 | 10 | 1 | 1 | 1 | 1 | 10 | 1 | 3 | 3 | 3 | 1 | 1 | 1 | 3 | 1 | 1 | 1 | 1 | 1 | 1 | 1 | 1 | 10 | 10 | 3 |
| yeps_YPTB1048 | 3 | 7 | 3 | 10 | 1 | 1 | 3 | 8 | 7 | 9 | 10 | 1 | 5 | 5 | 1 | 9 | 5 | 3 | 3 | 3 | 1 | 1 | 1 | 3 | 5 | 5 | 1 | 1 | 1 | 1 | 1 | 5 | 10 | 10 | 3 |
| yeps_YPTB1050 | 1 | 1 | 1 | 1 | 1 | 1 | 1 | 3 | 1 | 1 | 1 | 1 | 1 | 1 | 1 | 1 | 1 | 1 | 1 | 1 | 1 | 1 | 1 | 1 | 1 | 1 | 1 | 1 | 1 | 1 | 1 | 1 | 1 | 1 | 1 |
| yeps_YPTB1051 | 2 | 5 | | 4 | 1 | 1 | 2 | 4 | 5 | 4 | 4 | 3 | 1 | 1 | 1 | 4 | 1 | 2 | 2 | 2 | 1 | 1 | 1 | 2 | 10 | 10 | 1 | 1 | 1 | 1 | 1 | 10 | 4 | 4 | 2 |
| yeps_YPTB1053 | 4 | 7 | 4 | 7 | 1 | 1 | 4 | 1 | 7 | 7 | 7 | 3 | 1 | 1 | 1 | 7 | 1 | 4 | 4 | 4 | 1 | 1 | 1 | 4 | 1 | 1 | 1 | 1 | 1 | 1 | 1 | 1 | 7 | 7 | 4 |
| yeps_YPTB1054 | 5 | | 5 | 10 | 1 | 1 | 5 | 9 | 8 | 10 | 10 | 3 | 1 | 1 | 1 | 10 | 1 | 5 | 5 | 5 | 1 | 1 | 1 | 5 | 1 | 1 | 1 | 1 | 1 | 1 | 1 | 1 | 10 | 10 | 5 |
| yeps_YPTB1056 | 1 | 2 | 1 | 3 | 1 | 1 | 1 | 1 | 2 | 2 | 3 | 1 | 1 | 1 | 1 | 2 | 1 | 2 | 1 | 1 | 1 | 1 | 1 | 1 | 1 | 1 | 1 | 1 | 1 | 1 | 1 | 1 | 3 | 3 | 1 |
| yeps_YPTB1057 | 4 | 7 | 4 | 13 | 1 | 59 | 4 | 10 | 7 | 12 | 13 | 1 | 1 | 1 | 59 | 12 | 1 | 4 | 4 | 4 | 59 | 59 | 59 | 4 | 1 | 1 | 59 | 59 | 59 | 59 | 59 | 1 | 13 | 13 | 4 |
| yeps_YPTB1068 | 5 | 8 | 5 | 7 | 1 | 5 | 5 | 10 | 9 | 7 | 7 | 3 | 1 | 1 | 5 | 7 | 1 | 12 | 5 | 5 | 5 | 5 | 5 | 5 | 1 | 1 | 5 | 5 | 5 | 5 | 5 | 1 | 7 | 7 | 5 |
| yeps_YPTB1071 | 4 | 7 | 4 | 11 | 1 | 4 | 4 | 9 | 7 | 11 | 11 | 3 | 1 | 1 | 4 | 11 | 1 | 4 | 4 | 4 | 4 | 4 | 4 | 4 | 1 | 1 | 4 | 4 | 4 | 4 | 4 | 1 | 11 | 11 | 4 |
| yeps_YPTB1073 | 3 | 7 | 3 | 9 | 1 | 3 | 3 | 8 | 7 | 9 | 9 | 1 | 1 | 1 | 3 | 9 | 1 | 4 | 3 | 3 | 3 | 3 | 3 | 3 | 1 | 1 | 3 | 3 | 3 | 3 | 3 | 1 | 9 | 9 | 3 |
| yeps_YPTB1074 | 1 | 3 | 1 | 4 | 1 | 1 | 1 | 1 | 1 | 4 | 4 | 1 | 1 | 1 | 1 | 4 | 1 | 1 | 1 | 1 | 1 | 1 | 1 | 1 | 1 | 1 | 1 | 1 | 1 | 1 | 1 | 1 | 4 | 4 | 1 |
| yeps_YPTB1075 | 3 | 6 | 3 | 7 | 1 | 3 | 3 | 3 | 6 | 7 | 7 | 3 | 1 | 1 | 3 | 7 | 1 | 8 | 3 | 3 | 3 | 3 | 3 | 3 | 1 | 1 | 3 | 3 | 3 | 3 | 3 | 1 | 7 | 7 | 3 |
| yeps_YPTB1076 | 1 | 2 | 1 | 8 | 1 | 1 | 1 | 6 | 2 | 8 | 8 | 3 | 1 | 1 | 1 | 8 | 1 | 4 | 1 | 1 | 1 | 1 | 1 | 1 | 1 | 1 | 1 | 1 | 1 | 1 | 1 | 1 | 8 | 8 | 1 |
| yeps_YPTB1082 | 3 | 7 | 3 | 1 | 1 | 1 | 3 | 8 | 7 | 1 | 1 | 3 | 1 | 1 | 1 | 1 | 1 | 4 | 3 | 3 | 1 | 1 | 1 | 3 | 1 | 1 | 1 | 1 | 1 | 1 | 1 | 1 | 1 | 1 | 3 |
| yeps_YPTB1083 | 2 | 5 | 2 | 2 | 1 | 1 | 2 | 2 | 5 | 2 | 2 | 2 | 1 | 1 | 1 | 2 | 1 | 3 | 2 | 2 | 1 | 1 | 1 | 2 | 1 | 1 | 1 | 1 | 1 | 1 | 1 | 1 | 2 | 2 | 2 |
| yeps_YPTB1086 | 1 | 1 | 1 | 1 | 1 | 1 | 1 | 1 | 1 | 1 | 1 | 1 | 1 | 1 | 1 | 1 | 1 | 1 | 1 | 1 | 1 | 1 | 1 | 1 | 1 | 1 | 1 | 1 | 1 | 1 | 1 | 1 | 1 | 1 | 1 |
| yeps_YPTB1087 | 1 | 1 | 1 | 1 | 1 | 1 | 1 | 1 | 1 | 1 | 1 | 1 | 1 | 1 | 1 | 1 | 1 | 1 | 1 | 1 | 1 | 1 | 1 | 1 | 1 | 1 | 1 | 1 | 1 | 1 | 1 | 1 | 1 | 1 | 1 |
| yeps_YPTB1089 | 1 | 4 | 1 | 2 | 1 | 1 | 1 | 3 | 4 | 2 | 2 | 3 | 1 | 1 | 1 | 2 | 1 | 6 | 1 | 1 | 1 | 1 | 1 | 1 | 1 | 1 | 1 | 1 | 1 | 1 | 1 | 1 | 2 | 2 | 1 |
| yeps_YPTB1090 | 1 | 1 | 1 | 1 | 1 | 1 | 1 | 1 | 1 | 1 | 1 | 1 | 1 | 1 | 1 | 1 | 1 | 1 | 1 | 1 | 1 | 1 | 1 | 1 | 1 | 1 | 1 | 1 | 1 | 1 | 1 | 1 | 1 | 1 | 1 |
| yeps_YPTB1092 | 1 | 5 | 1 | 6 | 1 | 1 | 1 | 1 | 5 | 6 | 6 | 1 | 1 | 1 | 1 | 6 | 1 | 3 | 1 | 1 | 1 | 1 | 1 | 1 | 1 | 1 | 1 | 1 | 1 | 1 | 1 | 1 | 6 | 6 | 1 |
| yeps_YPTB1093 | 1 | 1 | 1 | 1 | 1 | 1 | 1 | 1 | 1 | 1 | 1 | 1 | 1 | 1 | 1 | 1 | 1 | 1 | 1 | 1 | 1 | 1 | 1 | 1 | 1 | 1 | 1 | 1 | 1 | 1 | 1 | 1 | 1 | 1 | 1 |
| yeps_YPTB1094 | 1 | 8 | 1 | 4 | 1 | 1 | 1 | 4 | 8 | 4 | 4 | 3 | 4 | 4 | 1 | 4 | 4 | 5 | 1 | 1 | 1 | 1 | 1 | 1 | 4 | 4 | 1 | 1 | 1 | 1 | 1 | 4 | 4 | 4 | 1 |
| yeps_YPTB1095 | 1 | 1 | 1 | 4 | 1 | 1 | 1 | 2 | 1 | 2 | 4 | 2 | 2 | 2 | 1 | 2 | 2 | 3 | 1 | 1 | 1 | 1 | 1 | 1 | 2 | 2 | 1 | 1 | 1 | 1 | 1 | 2 | 4 | 4 | 1 |
| yeps_YPTB1099 | 2 | 1 | 2 | 1 | 1 | 1 | 2 | 2 | 1 | 3 | 1 | 1 | 1 | 1 | 1 | 3 | 17 | 2 | 2 | 2 | 1 | 1 | 1 | 2 | 1 | 1 | 1 | 1 | 1 | 1 | 1 | 1 | 1 | 1 | 2 |
| yeps_YPTB1100 | 1 | 3 | 1 | 1 | 1 | 1 | 1 | 1 | 3 | 1 | 1 | 1 | 1 | 1 | 1 | 1 | 1 | 2 | 1 | 1 | 1 | 1 | 1 | 1 | 1 | 1 | 1 | 1 | 1 | 1 | 1 | 1 | 1 | 1 | 1 |
| yeps_YPTB1101 | 1 | 4 | 1 | 5 | 1 | 1 | 1 | 1 | 4 | 5 | 5 | 3 | 3 | 3 | 1 | 5 | 3 | 1 | 1 | 1 | 1 | 1 | 1 | 1 | 3 | 3 | 1 | 1 | 1 | 1 | 1 | 3 | 5 | 5 | 1 |
| yeps_YPTB1102 | 1 | 1 | 1 | 2 | 1 | 1 | 1 | 1 | 1 | 2 | 2 | 1 | 1 | 1 | 1 | 2 | 1 | 1 | 1 | 1 | 1 | 1 | 1 | 1 | 1 | 1 | 1 | 1 | 1 | 1 | 1 | 1 | 2 | 2 | 1 |
| yeps_YPTB1107 | 1 | 1 | 1 | 2 | 1 | 1 | 1 | 1 | 1 | 2 | 2 | 1 | 1 | 1 | 1 | 2 | 1 | 1 | 1 | 1 | 1 | 1 | 1 | 1 | 1 | 1 | 1 | 1 | 1 | 1 | 1 | 1 | 2 | 2 | 1 |
| yeps_YPTB1109 | 1 | 6 | 1 | 2 | 1 | 1 | 1 | 2 | 6 | 2 | 2 | 2 | 2 | 2 | 1 | 2 | 2 | 2 | 1 | 1 | 1 | 1 | 1 | 1 | 2 | 2 | 1 | 1 | 1 | 1 | 1 | 2 | 2 | 2 | 1 |
| yeps_YPTB1111 | 2 | 1 | 2 | 2 | 1 | 1 | 2 | 5 | 1 | 2 | 2 | 1 | 1 | 1 | 1 | 2 | 1 | 3 | 2 | 2 | 1 | 1 | 1 | 2 | 1 | 1 | 1 | 1 | 1 | 1 | 1 | 1 | 2 | 2 | 2 |
| yeps_YPTB1117 | 4 | 3 | 4 | 3 | 1 | 1 | 4 | 7 | 3 | 3 | 3 | 3 | 1 | 1 | 1 | 3 | 1 | 5 | 4 | 4 | 1 | 1 | 1 | 4 | 1 | 1 | 1 | 1 | 1 | 1 | 1 | 1 | 3 | 3 | 4 |
| yeps_YPTB1118 | 4 | 4 | 4 | 9 | 1 | 1 | 4 | 8 | 4 | 9 | 9 | 3 | 1 | 1 | 1 | 9 | 1 | 6 | 4 | 4 | 1 | 1 | 1 | 4 | 1 | 1 | 1 | 1 | 1 | 1 | 1 | 1 | 9 | 9 | 4 |
| yeps_YPTB1123 | 4 | 7 | 4 | 1 | 1 | 1 | 4 | 9 | 7 | 1 | 1 | 3 | 1 | 1 | 1 | 1 | 1 | 4 | 4 | 4 | 1 | 1 | 1 | 4 | 1 | 1 | 1 | 1 | 1 | 1 | 1 | 1 | 1 | 1 | 4 |
| yeps_YPTB1124 | 1 | 1 | 1 | 1 | 1 | 1 | 1 | 1 | 1 | 1 | 1 | 1 | 1 | 1 | 1 | 1 | 1 | 1 | 1 | 1 | 1 | 1 | 1 | 1 | 1 | 1 | 1 | 1 | 1 | 1 | 1 | 1 | 1 | 1 | 1 |
| yeps_YPTB1126 | 1 | 1 | 1 | 1 | 1 | 1 | 1 | 1 | 1 | 1 | 1 | 1 | 1 | 1 | 1 | 1 | 1 | 1 | 1 | 1 | 1 | 1 | 1 | 1 | 1 | 1 | 1 | 1 | 1 | 1 | 1 | 1 | 1 | 1 | 1 |
| yeps_YPTB1136 | 1 | 3 | 1 | 1 | 1 | 1 | 1 | 1 | 3 | 1 | 1 | 1 | 1 | 1 | 1 | 1 | 1 | 1 | 1 | 1 | 1 | 1 | 1 | 1 | 1 | 1 | 1 | 1 | 1 | 1 | 1 | 1 | 1 | 1 | 1 |
| yeps_YPTB1137 | 1 | 1 | 1 | 1 | 1 | 1 | 1 | 1 | 1 | 1 | 1 | 1 | 1 | 1 | 1 | 1 | 1 | 1 | 1 | 1 | 1 | 1 | 1 | 1 | 1 | 1 | 1 | 1 | 1 | 1 | 1 | 1 | 1 | 1 | 1 |
| yeps_YPTB1138 | 3 | 3 | 3 | 3 | 1 | 1 | 3 | 3 | 3 | 3 | 3 | 3 | 1 | 1 | 1 | 3 | 1 | 3 | 3 | 3 | 1 | 1 | 1 | 3 | 1 | 1 | 1 | 1 | 1 | 1 | 1 | 1 | 3 | 3 | 3 |
| yeps_YPTB1140 | 4 | 2 | 1 | 3 | 1 | 1 | 1 | 2 | 2 | 3 | 3 | 2 | 1 | 1 | 1 | 3 | 1 | 2 | 1 | 1 | 1 | 1 | 1 | 1 | 1 | 1 | 1 | 1 | 1 | 1 | 1 | 1 | 3 | 3 | 1 |
| yeps_YPTB1142 | 1 | 1 | 1 | 1 | 1 | 1 | 1 | 6 | 1 | 1 | 1 | 1 | 1 | 1 | 1 | 1 | 1 | 1 | 1 | 1 | 1 | 1 | 1 | 1 | 1 | 1 | 1 | 1 | 1 | 1 | 1 | 1 | 1 | 1 | 1 |
| yeps_YPTB1143 | 1 | 1 | 1 | 1 | 1 | 1 | 1 | 1 | 1 | 1 | 1 | 1 | 1 | 1 | 1 | 1 | 1 | 1 | 1 | 1 | 1 | 1 | 1 | 1 | 1 | 1 | 1 | 1 | 1 | 1 | 1 | 1 | 1 | 1 | 1 |
| yeps_YPTB1152 | 3 | 5 | 3 | 2 | 1 | 1 | 3 | 6 | 5 | 2 | 2 | 2 | 1 | 1 | 1 | 2 | 1 | 2 | 3 | 3 | 1 | 1 | 1 | 3 | 1 | 1 | 1 | 1 | 1 | 1 | 1 | 1 | 2 | 2 | 3 |
| yeps_YPTB1153 | 1 | 1 | 1 | 1 | 1 | 1 | 1 | 1 | 1 | 1 | 1 | 1 | 1 | 1 | 1 | 1 | 1 | 1 | 1 | 1 | 1 | 1 | 1 | 1 | 1 | 1 | 1 | 1 | 1 | 1 | 1 | 1 | 1 | 1 | 1 |
| yeps_YPTB1154 | 2 | 3 | 2 | 2 | 1 | 1 | 2 | 4 | 3 | 2 | 2 | 1 | 1 | 1 | 1 | 2 | 1 | 2 | 2 | 2 | 1 | 1 | 1 | 2 | 1 | 1 | 1 | 1 | 1 | 1 | 1 | 1 | 2 | 2 | 2 |
| yeps_YPTB1156 | 1 | 5 | 1 | 1 | 1 | 1 | 1 | 1 | 5 | 1 | 1 | 1 | 1 | 1 | 1 | 1 | 1 | 4 | 1 | 1 | 1 | 1 | 1 | 1 | 1 | 1 | 1 | 1 | 1 | 1 | 1 | 1 | 1 | 1 | 1 |
| yeps_YPTB1160 | 1 | 1 | 1 | 1 | 1 | 1 | 1 | 1 | 1 | 1 | 1 | 1 | 1 | 1 | 1 | 1 | 1 | 1 | 1 | 1 | 1 | 1 | 1 | 1 | 1 | 1 | 1 | 1 | 1 | 1 | 1 | 1 | 1 | 1 | 1 |
| yeps_YPTB1167 | 1 | 1 | 1 | 1 | 1 | 1 | 1 | 2 | 1 | 1 | 1 | 1 | 1 | 1 | 1 | 1 | 1 | 1 | 1 | 1 | 1 | 1 | 1 | 1 | 1 | 1 | 1 | 1 | 1 | 1 | 1 | 1 | 1 | 1 | 1 |
| yeps_YPTB1168 | 2 | 4 | 2 | 2 | 1 | 1 | 2 | 2 | 4 | 6 | 2 | 2 | 1 | 1 | 1 | 6 | 1 | 3 | 2 | 2 | 1 | 1 | 1 | 2 | 1 | 1 | 1 | 1 | 1 | 1 | 1 | 1 | 2 | 2 | 2 |
| yeps_YPTB1170 | 3 | 1 | 3 | 1 | 1 | 1 | 3 | 1 | 1 | 1 | 1 | 1 | 1 | 1 | 1 | 1 | 1 | 1 | 3 | 3 | 1 | 1 | 1 | 3 | 1 | 1 | 1 | 1 | 1 | 1 | 1 | 1 | 1 | 1 | 3 |
| yeps_YPTB1171 | 7 | 6 | 7 | 2 | 1 | 1 | 7 | 1 | 6 | 2 | 2 | 3 | 1 | 1 | 1 | 2 | 1 | 2 | 4 | 4 | 1 | 1 | 1 | 4 | 1 | 1 | 1 | 1 | 1 | 1 | 1 | 1 | 2 | 2 | 7 |
| yeps_YPTB1173 | 59 | 8 | 4 | 10 | 1 | 1 | 4 | 9 | 8 | 10 | 10 | 3 | 1 | 1 | 1 | 10 | 1 | 5 | 4 | 4 | 1 | 1 | 1 | 4 | 1 | 1 | 1 | 1 | 1 | 1 | 1 | 1 | 10 | 10 | 4 |
| yeps_YPTB1174 | 1 | 4 | 1 | 1 | 1 | 1 | 1 | 1 | 4 | 1 | 1 | 1 | 1 | 1 | 1 | 1 | 1 | 1 | 1 | 1 | 1 | 1 | 1 | 1 | 1 | 1 | 1 | 1 | 1 | 1 | 1 | 1 | 1 | 1 | 1 |
| yeps_YPTB1175 | 1 | 1 | 1 | 1 | 1 | 1 | 1 | 1 | 1 | 1 | 1 | 1 | 1 | 1 | 1 | 1 | 1 | 1 | 1 | 1 | 1 | 1 | 1 | 1 | 1 | 1 | 1 | 1 | 1 | 1 | 1 | 1 | 1 | 1 | 1 |
| yeps_YPTB1176 | 1 | 1 | 1 | 3 | 1 | 1 | 1 | 3 | 1 | 3 | 3 | 2 | 1 | 1 | 1 | 3 | 1 | 3 | 1 | 1 | 1 | 1 | 1 | 1 | 1 | 1 | 1 | 1 | 1 | 1 | 1 | 1 | 3 | 3 | 1 |
| yeps_YPTB1178 | 11 | 8 | 5 | 10 | 1 | 1 | 5 | 9 | 8 | 10 | 10 | 3 | 1 | 1 | 1 | 10 | 1 | 14 | 5 | 5 | 1 | 1 | 1 | 5 | 1 | 1 | 1 | 1 | 1 | 1 | 1 | 1 | 10 | 10 | 5 |
| yeps_YPTB1179 | 3 | 7 | 3 | 10 | 1 | 1 | 3 | 8 | 7 | 10 | 10 | 1 | 1 | 1 | 1 | 10 | 1 | 4 | 3 | 3 | 1 | 1 | 1 | 3 | 1 | 1 | 1 | 1 | 1 | 1 | 1 | 1 | 10 | 10 | 3 |
| yeps_YPTB1180 | 3 | 7 | 3 | 9 | 1 | 1 | 3 | 8 | 7 | 9 | 9 | 1 | 1 | 1 | 1 | 9 | 1 | 4 | 3 | 3 | 1 | 1 | 1 | 3 | 1 | 1 | 1 | 1 | 1 | 1 | 1 | 1 | 9 | 9 | 3 |
| yeps_YPTB1182 | 1 | 1 | 1 | 1 | 1 | 1 | 1 | 1 | 1 | 1 | 1 | 1 | 1 | 1 | 1 | 1 | 1 | 1 | 1 | 1 | 1 | 1 | 1 | 1 | 1 | 1 | 1 | 1 | 1 | 1 | 1 | 1 | 1 | 1 | 1 |
| yeps_YPTB1183 | 4 | 9 | 4 | 11 | 1 | 1 | 4 | 3 | 9 | 11 | 11 | 3 | 1 | 1 | 1 | 11 | 1 | 5 | 4 | 4 | 1 | 1 | 1 | 33 | 1 | 1 | 1 | 1 | 1 | 1 | 1 | 1 | 11 | 11 | 4 |
| yeps_YPTB1184 | 4 | 6 | 4 | 36 | 1 | 1 | 4 | 7 | 6 | 8 | 8 | 3 | 1 | 1 | 1 | 8 | 1 | 9 | 4 | 4 | 1 | 1 | 1 | 4 | 1 | 1 | 1 | 1 | 1 | 1 | 1 | 1 | 8 | 8 | 4 |
| yeps_YPTB1185 | 1 | 4 | 1 | 1 | 1 | 1 | 1 | 5 | 4 | 1 | 1 | 3 | 1 | 1 | 1 | 1 | 1 | 1 | 1 | 1 | 1 | 1 | 1 | 1 | 1 | 1 | 1 | 1 | 1 | 1 | 1 | 1 | 1 | 1 | 1 |
| yeps_YPTB1186 | 3 | 3 | 3 | 5 | 1 | 1 | 3 | 3 | 3 | 5 | 5 | 3 | 1 | 1 | 1 | 5 | 1 | 3 | 3 | 3 | 1 | 1 | 1 | 3 | 1 | 1 | 1 | 1 | 1 | 1 | 1 | 1 | 5 | 5 | 3 |
| yeps_YPTB1187 | 3 | 7 | 3 | 10 | 1 | 1 | 3 | 7 | 7 | 10 | 10 | 1 | 1 | 1 | | 10 | 1 | 4 | 3 | 3 | 1 | 1 | 1 | 3 | 1 | 1 | 1 | 1 | 1 | 1 | 1 | 1 | 10 | 10 | 3 |

| | | | | | | | | | | | | | | | | | | | | | | | | | | | | | | | | | | | |
|---|---|---|---|---|---|---|---|---|---|---|---|---|---|---|---|---|---|---|---|---|---|---|---|---|---|---|---|---|---|---|---|---|---|---|---|
| yeps_YPTB1189 | 3 | 5 | 3 | 6 | 1 | 1 | 3 | 6 | 5 | 6 | 6 | 2 | 1 | 1 | 1 | 6 | 1 | 3 | 3 | 3 | 1 | 1 | 1 | 3 | 1 | 1 | 1 | 1 | 1 | 1 | 1 | 1 | 6 | 6 | 3 |
| yeps_YPTB1193 | 1 | 1 | 1 | 1 | 1 | 1 | 1 | 1 | 1 | 1 | 1 | 2 | 1 | 1 | 1 | 1 | 1 | 1 | 1 | 1 | 1 | 1 | 1 | 1 | 1 | 1 | 1 | 1 | 1 | 1 | 1 | 1 | 1 | 1 | 1 |
| yeps_YPTB1197 | 1 | 1 | 1 | 2 | 1 | 1 | 1 | 1 | 1 | 2 | 2 | 1 | 1 | 1 | 1 | 2 | 1 | 3 | 1 | 1 | 1 | 1 | 1 | 1 | 1 | 1 | 1 | 1 | 1 | 1 | 1 | 1 | 2 | 2 | 1 |
| yeps_YPTB1199 | 1 | 5 | 1 | 6 | 1 | 1 | 1 | 1 | 5 | 6 | 6 | 1 | 1 | 1 | 1 | 6 | 1 | 1 | 1 | 1 | 1 | 1 | 1 | 1 | 1 | 1 | 1 | 1 | 1 | 1 | 1 | 1 | 6 | 6 | 1 |
| yeps_YPTB1201 | 1 | 5 | 1 | 6 | 1 | 1 | 1 | 4 | 5 | 6 | 6 | 3 | 4 | 4 | 1 | 6 | 4 | 1 | 1 | 1 | 1 | 1 | 1 | 1 | 4 | 4 | 1 | 1 | 1 | 1 | 1 | 4 | 6 | 6 | 1 |
| yeps_YPTB1202 | 5 | 7 | 5 | 9 | 1 | 1 | 5 | 8 | 7 | 9 | 9 | 3 | 4 | 4 | 1 | 9 | 4 | 1 | 5 | 5 | 1 | 1 | 1 | 5 | 4 | 4 | 1 | 1 | 1 | 1 | 1 | 4 | 9 | 9 | 5 |
| yeps_YPTB1203 | 2 | 4 | 2 | 3 | 1 | 1 | 2 | 3 | 4 | 3 | 3 | 2 | 3 | 3 | 1 | 3 | 3 | 1 | 2 | 2 | 1 | 1 | 1 | 2 | 3 | 3 | 1 | 1 | 1 | 1 | 1 | 3 | 3 | 3 | 2 |
| yeps_YPTB1204 | 8 | 7 | 8 | 10 | 1 | 1 | 32 | 9 | 7 | 10 | 10 | 3 | 4 | 4 | 1 | 10 | 4 | 1 | 8 | 8 | 1 | 1 | 1 | 8 | 4 | 4 | 1 | 1 | 1 | 1 | 1 | 4 | 10 | 10 | 8 |
| yeps_YPTB1205 | 5 | 7 | 5 | 10 | 1 | 1 | 5 | 3 | 7 | 9 | 10 | 3 | 5 | 5 | 1 | 9 | 5 | 1 | 5 | 5 | 1 | 1 | 1 | 5 | 5 | 5 | 1 | 1 | 1 | 1 | 1 | 5 | 10 | 10 | 5 |
| yeps_YPTB1207 | 1 | 1 | 1 | 4 | 1 | 1 | 1 | 1 | 1 | 4 | 4 | 1 | 1 | 1 | 1 | 4 | 1 | 1 | 1 | 1 | 1 | 1 | 1 | 1 | 1 | 1 | 1 | 1 | 1 | 1 | 1 | 1 | 4 | 4 | 1 |
| yeps_YPTB1208 | 2 | 2 | 2 | 3 | 1 | 1 | 2 | 2 | 2 | 3 | 3 | 2 | 1 | 1 | 1 | 3 | 1 | 1 | 2 | 2 | 1 | 1 | 1 | 2 | 1 | 1 | 1 | 1 | 1 | 1 | 1 | 1 | 3 | 3 | 2 |
| yeps_YPTB1211 | 2 | 5 | 2 | 2 | 1 | 1 | 2 | 6 | 5 | 2 | 2 | 3 | 4 | 4 | 1 | 2 | 4 | 1 | 2 | 2 | 1 | 1 | 1 | 2 | 4 | 4 | 1 | 1 | 1 | 1 | 1 | 4 | 2 | 2 | 2 |
| yeps_YPTB1220 | 3 | 3 | 3 | 1 | 1 | 1 | 3 | 2 | 3 | 2 | 1 | 2 | 2 | 2 | 1 | 2 | 2 | 1 | 3 | 3 | 1 | 1 | 1 | 3 | 2 | 2 | 1 | 1 | 1 | 1 | 1 | 2 | 1 | 1 | 3 |
| yeps_YPTB1221 | 2 | 2 | 2 | 2 | 1 | 1 | 2 | 1 | 2 | 6 | 2 | 3 | 4 | 4 | 1 | 6 | 4 | 1 | 2 | 2 | 1 | 1 | 1 | 2 | 4 | 4 | 1 | 1 | 1 | 1 | 1 | 4 | 2 | 2 | 2 |
| yeps_YPTB1222 | 1 | 1 | 1 | 1 | 1 | 1 | 1 | 2 | 1 | 1 | 1 | 1 | 1 | 1 | 1 | 1 | 1 | 1 | 1 | 1 | 1 | 1 | 1 | 1 | 1 | 1 | 1 | 1 | 1 | 1 | 1 | 1 | 1 | 1 | 1 |
| yeps_YPTB1229 | 4 | 8 | 4 | 12 | 1 | 1 | 4 | 10 | 8 | 12 | 12 | 4 | 1 | 1 | 1 | 12 | 1 | 5 | 4 | 4 | 1 | 1 | 1 | 4 | 1 | 1 | 1 | 1 | 1 | 1 | 1 | 1 | 12 | 12 | 4 |
| yeps_YPTB1231 | 1 | 1 | 1 | 4 | 1 | 1 | 1 | 2 | 1 | 4 | 4 | 1 | 1 | 1 | 1 | 4 | 1 | 4 | 1 | 1 | 1 | 1 | 1 | 1 | 1 | 1 | 1 | 1 | 1 | 1 | 1 | 1 | 4 | 4 | 1 |
| yeps_YPTB1250 | 1 | 6 | 1 | 7 | 1 | 1 | 1 | 3 | 6 | 7 | 7 | 3 | 1 | 1 | 1 | 7 | 1 | 1 | 1 | 1 | 1 | 1 | 1 | 1 | 1 | 1 | 1 | 1 | 1 | 1 | 1 | 1 | 7 | 7 | 1 |
| yeps_YPTB1251 | 1 | 1 | 1 | 1 | 1 | 1 | 1 | 1 | 1 | 1 | 1 | 1 | 1 | 1 | 1 | 1 | 1 | 1 | 1 | 1 | 1 | 1 | 1 | 1 | 1 | 1 | 1 | 1 | 1 | 1 | 1 | 1 | 1 | 1 | 1 |
| yeps_YPTB1252 | 1 | 1 | 1 | 1 | 1 | 1 | 1 | 1 | 1 | 1 | 1 | 1 | 1 | 1 | 1 | 1 | 1 | 1 | 1 | 1 | 1 | 1 | 1 | 1 | 1 | 1 | 1 | 1 | 1 | 1 | 1 | 1 | 1 | 1 | 1 |
| yeps_YPTB1255 | 1 | 1 | 1 | 1 | 1 | 1 | 1 | 4 | 1 | 1 | 1 | 1 | 1 | 1 | 1 | 1 | 1 | 1 | 5 | 5 | 1 | 1 | 1 | 5 | 1 | 1 | 1 | 1 | 1 | 1 | 1 | 1 | 1 | 1 | 1 |
| yeps_YPTB1257 | 5 | 9 | 5 | 10 | 1 | 1 | 5 | 4 | 9 | 10 | 10 | 3 | 1 | 1 | 1 | 10 | 1 | 6 | 5 | 5 | 1 | 1 | 1 | 5 | 1 | 1 | 1 | 1 | 1 | 1 | 1 | 1 | 10 | 10 | 5 |
| yeps_YPTB1258 | 1 | 1 | 1 | 1 | 1 | 1 | 1 | 1 | 1 | 1 | 1 | 1 | 1 | 1 | 1 | 1 | 1 | 1 | 1 | 1 | 1 | 1 | 1 | 1 | 1 | 1 | 1 | 1 | 1 | 1 | 1 | 1 | 1 | 1 | 1 |
| yeps_YPTB1259 | 1 | 5 | 1 | 8 | 1 | 1 | 1 | 1 | 5 | 6 | 8 | 1 | 1 | 1 | 1 | 6 | 1 | 1 | 1 | 1 | 1 | 1 | 1 | 1 | 1 | 1 | 1 | 1 | 1 | 1 | 1 | 1 | 8 | 8 | 1 |
| yeps_YPTB1263 | 5 | 8 | 5 | 11 | 1 | 1 | 5 | 9 | 8 | 11 | 11 | 3 | 1 | 1 | 1 | 11 | 1 | 1 | 5 | 5 | 1 | 1 | 1 | 5 | 1 | 1 | 1 | 1 | 1 | 1 | 1 | 1 | 11 | 11 | 5 |
| yeps_YPTB1268 | 1 | 1 | 1 | 1 | 1 | 1 | 1 | 1 | 1 | 1 | 1 | 1 | 1 | 1 | 1 | 1 | 1 | 1 | 1 | 1 | 1 | 1 | 1 | 1 | 1 | 1 | 1 | 1 | 1 | 1 | 1 | 1 | 1 | 1 | 1 |
| yeps_YPTB1270 | 4 | 8 | 4 | 7 | 1 | 1 | 4 | 9 | 8 | 7 | 7 | 3 | 1 | 1 | 1 | 7 | 1 | 5 | 4 | 4 | 1 | 1 | 1 | 4 | 1 | 1 | 1 | 1 | 1 | 1 | 1 | 20 | 7 | 7 | 4 |
| yeps_YPTB1272 | 2 | 1 | 2 | 1 | 1 | 1 | 2 | 3 | 1 | 1 | 1 | 1 | 1 | 1 | 1 | 1 | 1 | 2 | 2 | 2 | 1 | 1 | 1 | 2 | 1 | 1 | 1 | 1 | 1 | 1 | 1 | 1 | 1 | 1 | 2 |
| yeps_YPTB1273 | 2 | 1 | 2 | 4 | 1 | 1 | 2 | 2 | 1 | 3 | 4 | 1 | 1 | 1 | 1 | 3 | 1 | 1 | 2 | 2 | 1 | 1 | 1 | 2 | 1 | 1 | 1 | 1 | 1 | 1 | 1 | 1 | 4 | 4 | 2 |
| yeps_YPTB1274 | 1 | 2 | 1 | 4 | 1 | 1 | 1 | 2 | 2 | 4 | 4 | 1 | 1 | 1 | 1 | 4 | 1 | 2 | 1 | 1 | 1 | 1 | 1 | 1 | 1 | 1 | 1 | 1 | 1 | 1 | 1 | 1 | 4 | 4 | 1 |
| yeps_YPTB1276 | 1 | 1 | 1 | 1 | 1 | 1 | 1 | 1 | 1 | 1 | 1 | 1 | 1 | 1 | 1 | 1 | 1 | 4 | 1 | 1 | 1 | 1 | 1 | 1 | 1 | 1 | 1 | 1 | 1 | 1 | 1 | 1 | 1 | 1 | 1 |
| yeps_YPTB1279 | 1 | 1 | 1 | 1 | 1 | 1 | 1 | 1 | 1 | 1 | 1 | 1 | 1 | 1 | 1 | 1 | 1 | 3 | 1 | 1 | 1 | 1 | 1 | 1 | 1 | 1 | 1 | 1 | 1 | 1 | 1 | 1 | 1 | 1 | 1 |
| yeps_YPTB1283 | 3 | 3 | 3 | 4 | 1 | 1 | 3 | 3 | 3 | 4 | 4 | 3 | 3 | 3 | 1 | 4 | 3 | 1 | 3 | 3 | 1 | 1 | 1 | 3 | 3 | 3 | 1 | 1 | 1 | 1 | 1 | 3 | 4 | 4 | 3 |
| yeps_YPTB1285 | 3 | 8 | 5 | 8 | 1 | 1 | 5 | 11 | 8 | 8 | 8 | 3 | 3 | 3 | 1 | 8 | 3 | 1 | 5 | 5 | 1 | 1 | 1 | 5 | 3 | 3 | 1 | 1 | 1 | 1 | 1 | 3 | 8 | 8 | 5 |
| yeps_YPTB1286 | 1 | 6 | 1 | 8 | 1 | 1 | 1 | 3 | 6 | 8 | 8 | 1 | 1 | 1 | 1 | 8 | 1 | 1 | 1 | 1 | 1 | 1 | 1 | 1 | 1 | 1 | 1 | 1 | 1 | 1 | 1 | 1 | 8 | 8 | 1 |
| yeps_YPTB1289 | 1 | 4 | 1 | 8 | 1 | 1 | 1 | 4 | 4 | 7 | 8 | 1 | 4 | 4 | 1 | 7 | 4 | 2 | 1 | 1 | 1 | 1 | 1 | 1 | 4 | 4 | 1 | 1 | 1 | 1 | 1 | 4 | 8 | 8 | 1 |
| yeps_YPTB1290 | 1 | 1 | 1 | 1 | 1 | 1 | 1 | 6 | 1 | 1 | 1 | 1 | 1 | 1 | 1 | 1 | 1 | 1 | 1 | 1 | 1 | 1 | 1 | 1 | 1 | 1 | 1 | 1 | 1 | 1 | 1 | 1 | 1 | 1 | 1 |
| yeps_YPTB1300 | 3 | 7 | 3 | 11 | 1 | 1 | 3 | 9 | 7 | 11 | 11 | 1 | 1 | 1 | 1 | 11 | 1 | 2 | 3 | 3 | 1 | 1 | 1 | 3 | 1 | 1 | 1 | 1 | 1 | 1 | 1 | 1 | 11 | 11 | 3 |
| yeps_YPTB1301 | 3 | 2 | 3 | 7 | 1 | 1 | 3 | 2 | 2 | 7 | 7 | 1 | 1 | 1 | 1 | 7 | 1 | 4 | 3 | 3 | 1 | 1 | 1 | 3 | 1 | 1 | 1 | 1 | 1 | 1 | 1 | 1 | 7 | 7 | 3 |
| yeps_YPTB1304 | 5 | 58 | 5 | 8 | 1 | 1 | 5 | 10 | 9 | 8 | 8 | 3 | 1 | 1 | 1 | 8 | 1 | 6 | 5 | 5 | 1 | 1 | 1 | 5 | 1 | 1 | 1 | 1 | 1 | 1 | 1 | 1 | 8 | 8 | 5 |
| yeps_YPTB1305 | 1 | 3 | 1 | 2 | 1 | 1 | 1 | 1 | 3 | 2 | 2 | 1 | 1 | 1 | 1 | 2 | 1 | 1 | 1 | 1 | 1 | 1 | 1 | 1 | 1 | 1 | 1 | 1 | 1 | 1 | 1 | 1 | 2 | 2 | 1 |
| yeps_YPTB1306 | 1 | 5 | 1 | 6 | 1 | 1 | 1 | 1 | 5 | 6 | 6 | 1 | 1 | 1 | 1 | 6 | 1 | 3 | 1 | 1 | 1 | 1 | 1 | 1 | 1 | 1 | 1 | 1 | 1 | 1 | 1 | 1 | 6 | 6 | 1 |
| yeps_YPTB1307 | 4 | 8 | 4 | 12 | 1 | 1 | 4 | 9 | 8 | 11 | 12 | 3 | 1 | 1 | 1 | 11 | 1 | 7 | 4 | 4 | 1 | 1 | 1 | 4 | 1 | 1 | 1 | 1 | 1 | 1 | 1 | 1 | 12 | 12 | 4 |
| yeps_YPTB1310 | 1 | 3 | 1 | 2 | 1 | 1 | 1 | 5 | 3 | 2 | 2 | 3 | 1 | 1 | 1 | 2 | 1 | 2 | 1 | 1 | 1 | 1 | 1 | 1 | 1 | 1 | 1 | 1 | 1 | 1 | 1 | 1 | 2 | 2 | 1 |
| yeps_YPTB1311 | 1 | 8 | 1 | 2 | 1 | 1 | 1 | 9 | 8 | 11 | 2 | 3 | 3 | 3 | 1 | 11 | 3 | 4 | 1 | 1 | 1 | 1 | 1 | 1 | 3 | 3 | 1 | 1 | 1 | 1 | 1 | 3 | 2 | 2 | 1 |
| yeps_YPTB1312 | 1 | 1 | 1 | 1 | 1 | 1 | 1 | 2 | 1 | 1 | 1 | 1 | 1 | 1 | 1 | 1 | 1 | 1 | 1 | 1 | 1 | 1 | 1 | 1 | 1 | 1 | 1 | 1 | 1 | 1 | 1 | 1 | 1 | 1 | 1 |
| yeps_YPTB1313 | 1 | 3 | 1 | 2 | 1 | 1 | 1 | 2 | 3 | 2 | 2 | 2 | 2 | 2 | 1 | 2 | 2 | 2 | 1 | 1 | 1 | 1 | 1 | 1 | 2 | 2 | 1 | 1 | 1 | 1 | 1 | 2 | 2 | 2 | 1 |
| yeps_YPTB1314 | 1 | 6 | 1 | 1 | 1 | 1 | 1 | 7 | 6 | 8 | 1 | 3 | 3 | 3 | 1 | 8 | 3 | 4 | 1 | 1 | 1 | 1 | 1 | 1 | 3 | 3 | 1 | 1 | 1 | 1 | 1 | 3 | 1 | 1 | 1 |
| yeps_YPTB1315 | 1 | 7 | 1 | 9 | 1 | 1 | 1 | 3 | 7 | 9 | 9 | 3 | 3 | 3 | 1 | 9 | 3 | 4 | 1 | 1 | 1 | 1 | 1 | 1 | 3 | 3 | 1 | 1 | 1 | 1 | 1 | 3 | 9 | 9 | 1 |
| yeps_YPTB1317 | 1 | 1 | 1 | 1 | 1 | 1 | 1 | 6 | 1 | 1 | 1 | 1 | 1 | 1 | 1 | 1 | 1 | 1 | 1 | 1 | 1 | 1 | 1 | 1 | 1 | 1 | 1 | 1 | 1 | 1 | 1 | 1 | 1 | 1 | 1 |
| yeps_YPTB1318 | 2 | 2 | 2 | 2 | 1 | 1 | 2 | 1 | 2 | 2 | 2 | 1 | 1 | 1 | 1 | 2 | 1 | 1 | 2 | 2 | 1 | 1 | 1 | 2 | 1 | 1 | 1 | 1 | 1 | 1 | 1 | 1 | 2 | 2 | 2 |
| yeps_YPTB1329 | 3 | 3 | 3 | 9 | 1 | 1 | 3 | 1 | 3 | 9 | 9 | 3 | 1 | 1 | 1 | 9 | 1 | 5 | 3 | 3 | 1 | 1 | 1 | 3 | 1 | 1 | 1 | 1 | 1 | 1 | 1 | 1 | 9 | 9 | 3 |
| yeps_YPTB1342 | 1 | 6 | 1 | 5 | 1 | 1 | 1 | 1 | 6 | 5 | 5 | 1 | 1 | 1 | 1 | 5 | 1 | 3 | 1 | 1 | 1 | 1 | 1 | 1 | 1 | 1 | 1 | 1 | 1 | 1 | 1 | 1 | 5 | 5 | 1 |
| yeps_YPTB1343 | 1 | 3 | 1 | 2 | 1 | 1 | 1 | 4 | 3 | 2 | 2 | 1 | 1 | 1 | 1 | 2 | 1 | 2 | 1 | 1 | 1 | 1 | 1 | 1 | 1 | 1 | 1 | 1 | 1 | 1 | 1 | 1 | 2 | 2 | 1 |
| yeps_YPTB1347 | 1 | 7 | 1 | 10 | 1 | 1 | 1 | 9 | 7 | 10 | 10 | 4 | 3 | 3 | 1 | 10 | 3 | 5 | 1 | 1 | 1 | 1 | 1 | 1 | 3 | 3 | 1 | 1 | 1 | 1 | 1 | 58 | 10 | 10 | 1 |
| yeps_YPTB1348 | 1 | 1 | 1 | 4 | 1 | 1 | 1 | 1 | 1 | 4 | 4 | 2 | 1 | 1 | 1 | 4 | 1 | 1 | 1 | 1 | 1 | 1 | 1 | 1 | 1 | 1 | 1 | 1 | 1 | 1 | 1 | 1 | 4 | 4 | 1 |
| yeps_YPTB1355 | 4 | 4 | 4 | 4 | 1 | 1 | 4 | 4 | 4 | 4 | 4 | 3 | 1 | 1 | 1 | 4 | 1 | 4 | 4 | 4 | 1 | 1 | 1 | 4 | 1 | 1 | 1 | 1 | 1 | 1 | 1 | 1 | 4 | 4 | 4 |
| yeps_YPTB1356 | 4 | 5 | 4 | 7 | 1 | 1 | 4 | 8 | 5 | 7 | 7 | 3 | 1 | 1 | 1 | 7 | 1 | 5 | 4 | 4 | 1 | 1 | 1 | 4 | 1 | 1 | 1 | 1 | 1 | 1 | 1 | 1 | 7 | 7 | 4 |
| yeps_YPTB1359 | 1 | 1 | 1 | 1 | 1 | 1 | 1 | 1 | 1 | 1 | 1 | 1 | 1 | 1 | 1 | 1 | 1 | 1 | 1 | 1 | 1 | 1 | 1 | 1 | 1 | 1 | 1 | 1 | 1 | 1 | 1 | 1 | 1 | 1 | 1 |
| yeps_YPTB1360 | 1 | 1 | 1 | 1 | 1 | 1 | 1 | 1 | 1 | 1 | 1 | 1 | 1 | 1 | 1 | 1 | 1 | 1 | 1 | 1 | 1 | 1 | 1 | 1 | 1 | 1 | 1 | 1 | 1 | 1 | 1 | 1 | 1 | 1 | 1 |
| yeps_YPTB1365 | 2 | 1 | 2 | 1 | 1 | 1 | 2 | 4 | 1 | 1 | 1 | 3 | 1 | 1 | 1 | 1 | 1 | 2 | 2 | 2 | 1 | 1 | 1 | 2 | 1 | 1 | 1 | 1 | 1 | 1 | 1 | 1 | 1 | 1 | 2 |
| yeps_YPTB1366 | 10 | 6 | 3 | 2 | 1 | 1 | 3 | 8 | 6 | 2 | 2 | 2 | 1 | 1 | 1 | 2 | 1 | 3 | 3 | 3 | 1 | 1 | 1 | 3 | 1 | 1 | 1 | 1 | 1 | 1 | 1 | 1 | 2 | 2 | 3 |
| yeps_YPTB1369 | 4 | 8 | 4 | 12 | 1 | 1 | 4 | 1 | 8 | 11 | 12 | 3 | 1 | 1 | 1 | 11 | 1 | 4 | 4 | 4 | 1 | 1 | 1 | 4 | 1 | 1 | 1 | 1 | 1 | 1 | 1 | 1 | 12 | 12 | 4 |
| yeps_YPTB1370 | 5 | 9 | 5 | 11 | 1 | 1 | 5 | 10 | 9 | 11 | 11 | 3 | 7 | 7 | 1 | 11 | 7 | 5 | 5 | 5 | 1 | 1 | 1 | 5 | 7 | 7 | 1 | 1 | 1 | 1 | 1 | 7 | 11 | 11 | 5 |
| yeps_YPTB1371 | 2 | 1 | 2 | 5 | 1 | 1 | 2 | 4 | 1 | 5 | 5 | 1 | 1 | 1 | 1 | 5 | 1 | 2 | 2 | 2 | 1 | 1 | 1 | 2 | 1 | 1 | 1 | 1 | 1 | 1 | 1 | 1 | 5 | 5 | 2 |
| yeps_YPTB1372 | 1 | 4 | 1 | 1 | 1 | 1 | 1 | 1 | 4 | 1 | 1 | 1 | 1 | 1 | 1 | 1 | 1 | 1 | 1 | 1 | 1 | 1 | 1 | 1 | 1 | 1 | 1 | 1 | 1 | 1 | 1 | 1 | 1 | 1 | 1 |
| yeps_YPTB1374 | 4 | 6 | 4 | 7 | 1 | 1 | 4 | 7 | 6 | 7 | 7 | 3 | 1 | 1 | 1 | 7 | 1 | 4 | 4 | 4 | 1 | 1 | 1 | 4 | 1 | 1 | 1 | 1 | 1 | 1 | 1 | 1 | 7 | 7 | 4 |
| yeps_YPTB1375 | 4 | 6 | 4 | 2 | 1 | 1 | 4 | 7 | 6 | 2 | 2 | 1 | 1 | 1 | 1 | 2 | 1 | 4 | 4 | 4 | 1 | 1 | 1 | 4 | 1 | 1 | 1 | 1 | 1 | 1 | 1 | 1 | 2 | 2 | 4 |
| yeps_YPTB1378 | 3 | 4 | 3 | 4 | 1 | 1 | 1 | 3 | 4 | 4 | 4 | 4 | 1 | 1 | 1 | 4 | 1 | 4 | 3 | 3 | 1 | 1 | 1 | 3 | 1 | 1 | 1 | 1 | 1 | 1 | 1 | 1 | 4 | 4 | 3 |
| yeps_YPTB1379 | 1 | 4 | 1 | 6 | 1 | 1 | 1 | 5 | 4 | 6 | 6 | 1 | 1 | 1 | 1 | 6 | 1 | 1 | 1 | 1 | 1 | 1 | 1 | 1 | 25 | 1 | 1 | 1 | 1 | 1 | 1 | 1 | 6 | 6 | 1 |
| yeps_YPTB1380 | 1 | 1 | 1 | 1 | 1 | 20 | 1 | 3 | 1 | 1 | 1 | 1 | 1 | 1 | 20 | 1 | 1 | 1 | 1 | 1 | 20 | 20 | 20 | 1 | 1 | 1 | 20 | 20 | 20 | 20 | 20 | 1 | 1 | 1 | 1 |
| yeps_YPTB1382 | 3 | 4 | 3 | 8 | 1 | 1 | 3 | 7 | 4 | 8 | 8 | 3 | 1 | 1 | 1 | 8 | 1 | 4 | 3 | 3 | 1 | 1 | 1 | 3 | 1 | 1 | 1 | 1 | 1 | 1 | 1 | 1 | 8 | 8 | 3 |
| yeps_YPTB1385 | 3 | 2 | 3 | 5 | 1 | 1 | 3 | 1 | 2 | 5 | 5 | 1 | 1 | 1 | 1 | 5 | 1 | 2 | 3 | 3 | 1 | 1 | 1 | 3 | 1 | 1 | 1 | 1 | 1 | 1 | 1 | 1 | 5 | 5 | 3 |
| yeps_YPTB1388 | 1 | 1 | 1 | 1 | 1 | 1 | 1 | 8 | 1 | 1 | 1 | 1 | 1 | 1 | 1 | 1 | 1 | 1 | 1 | 1 | 1 | 1 | 1 | 1 | 1 | 1 | 1 | 1 | 1 | 1 | 1 | 1 | 1 | 1 | 1 |
| yeps_YPTB1389 | 1 | 5 | 1 | 6 | 1 | 1 | 1 | 1 | 5 | 6 | 6 | 1 | 1 | 1 | 1 | 6 | 1 | 1 | 1 | 1 | 1 | 1 | 1 | 1 | 1 | 1 | 1 | 1 | 1 | 1 | 1 | 1 | 6 | 6 | 1 |
| yeps_YPTB1392 | 1 | 1 | 1 | 1 | 1 | 1 | 1 | 1 | 1 | 1 | 1 | 1 | 1 | 1 | 1 | 1 | 1 | 1 | 1 | 1 | 1 | 1 | 1 | 1 | 1 | 1 | 1 | 1 | 1 | 1 | 1 | 1 | 1 | 1 | 1 |
| yeps_YPTB1393 | 1 | 1 | 1 | 1 | 1 | 1 | 1 | 1 | 1 | 1 | 1 | 1 | 1 | 1 | 1 | 1 | 1 | 1 | 1 | 1 | 1 | 1 | 1 | 1 | 1 | 1 | 1 | 1 | 1 | 1 | 1 | 1 | 1 | 1 | 1 |
| yeps_YPTB1396 | 3 | 1 | 3 | 2 | 1 | 1 | 3 | 3 | 1 | 2 | 2 | 1 | 1 | 1 | 1 | 2 | 1 | 4 | 3 | 3 | 1 | 1 | 1 | 3 | 1 | 1 | 1 | 1 | 1 | 1 | 1 | 1 | 2 | 2 | 3 |
| yeps_YPTB1402 | 1 | 2 | 1 | 1 | 1 | 1 | 1 | 1 | 2 | 3 | 1 | 1 | 1 | 1 | 1 | 3 | 1 | 1 | 1 | 1 | 1 | 1 | 1 | 1 | 1 | 1 | 1 | 1 | 1 | 1 | 1 | 1 | 1 | 1 | 1 |
| yeps_YPTB1404 | 1 | 3 | 1 | 30 | 1 | 1 | 1 | 1 | 3 | 1 | 1 | 2 | 1 | 1 | 1 | 1 | 1 | 1 | 1 | 1 | 1 | 1 | 1 | 1 | 1 | 1 | 1 | 1 | 1 | 1 | 1 | 1 | 1 | 1 | 1 |
| yeps_YPTB1415 | 1 | 5 | 1 | 3 | 1 | 1 | 1 | 1 | 5 | 6 | 3 | 2 | 3 | 3 | 1 | 6 | 3 | 3 | 1 | 1 | 1 | 1 | 1 | 1 | 3 | 3 | 1 | 1 | 1 | 1 | 1 | 3 | 3 | 3 | 1 |
| yeps_YPTB1419 | 1 | 7 | 1 | 9 | 1 | 1 | 33 | 8 | 7 | 5 | 9 | 29 | 5 | 5 | 1 | 5 | 5 | 5 | 1 | 1 | 1 | 1 | 1 | 1 | 5 | 5 | 1 | 1 | 1 | 1 | 1 | 5 | 9 | 9 | 1 |
| yeps_YPTB1420 | 1 | 1 | 1 | 6 | 1 | 1 | 1 | 1 | 1 | 6 | 6 | 1 | 3 | 3 | 1 | 6 | 3 | 4 | 1 | 1 | 1 | 1 | 1 | 1 | 3 | 3 | 1 | 1 | 1 | 1 | 1 | 3 | 6 | 6 | 1 |
| yeps_YPTB1421 | 1 | 1 | 1 | 3 | 1 | 1 | 1 | 1 | 1 | 3 | 3 | 1 | 1 | 1 | 1 | 3 | 1 | 2 | 1 | 1 | 1 | 1 | 1 | 1 | 1 | 1 | 1 | 1 | 1 | 1 | 1 | 1 | 3 | 3 | 1 |
| yeps_YPTB1423 | 1 | 1 | 1 | 1 | 1 | 1 | 1 | 1 | 1 | 1 | 1 | 1 | 1 | 1 | 1 | 1 | 1 | 2 | 1 | 1 | 1 | 1 | 1 | 1 | 1 | 1 | 1 | 1 | 1 | 1 | 1 | 1 | 1 | 1 | 1 |
| yeps_YPTB1424 | 1 | 1 | 1 | 1 | 1 | 1 | 1 | 1 | 1 | 1 | 1 | 1 | 1 | 1 | 1 | 1 | 1 | 1 | 1 | 1 | 1 | 1 | 1 | 1 | 1 | 1 | 1 | 1 | 1 | 1 | 1 | 1 | 1 | 1 | 1 |
| yeps_YPTB1425 | 1 | 4 | 1 | 1 | 1 | 1 | 1 | 1 | 5 | 1 | 1 | 1 | 1 | 1 | 1 | 1 | 1 | 2 | 1 | 1 | 1 | 1 | 1 | 1 | 1 | 1 | 1 | 1 | 1 | 1 | 1 | 1 | 1 | 1 | 1 |
| yeps_YPTB1426 | 1 | 4 | 1 | 7 | 1 | 1 | 1 | 4 | 4 | 7 | 7 | 1 | 1 | 1 | 1 | 7 | 1 | 3 | 1 | 1 | 1 | 1 | 1 | 1 | 1 | 1 | 1 | 1 | 1 | 1 | 1 | 1 | 7 | 7 | 1 |
| yeps_YPTB1427 | 1 | 3 | 1 | 4 | 1 | 1 | 1 | 1 | 3 | 4 | 4 | 1 | 1 | 1 | 1 | 4 | 1 | 1 | 1 | 1 | 1 | 1 | 1 | 1 | 1 | 1 | 1 | 1 | 1 | 1 | 1 | 1 | 4 | 4 | 5 |
| yeps_YPTB1429 | 1 | 1 | 1 | 1 | 1 | 1 | 1 | 1 | 1 | 1 | 1 | 1 | 1 | 1 | 1 | 1 | 1 | 1 | 1 | 1 | 1 | 1 | 1 | 1 | 1 | 1 | 1 | 1 | 1 | 1 | 1 | 1 | 1 | 1 | 1 |
| yeps_YPTB1437 | 2 | 2 | 3 | 2 | 1 | 1 | 3 | 5 | 2 | 2 | 2 | 1 | 1 | 1 | 1 | 2 | 1 | 4 | 3 | 3 | 1 | 1 | 1 | 3 | 1 | 1 | 1 | 1 | 1 | 1 | 1 | 1 | 2 | 2 | 2 |
| yeps_YPTB1438 | 4 | 2 | 4 | 13 | 9 | 9 | 43 | 11 | 2 | 13 | 13 | 3 | 3 | 3 | 9 | 13 | 3 | 15 | 4 | 4 | 9 | 9 | 9 | 4 | 3 | 3 | 9 | 9 | 9 | 9 | 9 | 3 | 13 | 13 | 4 |
| yeps_YPTB1440 | 1 | 1 | 1 | 1 | 1 | 1 | 1 | 1 | 1 | 1 | 1 | 1 | 1 | 1 | 1 | 1 | 1 | 1 | 1 | 1 | 1 | 1 | 1 | 1 | 1 | 1 | 1 | 1 | 1 | 1 | 1 | 1 | 1 | 1 | 1 |
| yeps_YPTB1443 | 1 | 8 | 1 | 10 | 1 | 1 | 1 | 9 | 8 | 10 | 10 | 3 | 4 | 4 | 1 | 10 | 4 | 5 | 1 | 1 | 1 | 1 | 1 | 1 | 4 | 4 | 1 | 1 | 1 | 1 | 1 | 4 | 10 | 10 | 1 |
| yeps_YPTB1450 | 1 | 1 | 1 | 1 | 1 | 1 | 1 | 1 | 1 | 1 | 1 | 1 | 1 | 18 | 1 | 1 | 1 | 1 | 1 | 1 | 1 | 1 | 1 | 1 | 1 | 1 | 1 | 1 | 1 | 1 | 1 | 1 | 1 | 1 | 1 |
| yeps_YPTB1451 | 1 | 3 | 1 | 1 | 1 | 1 | 1 | 4 | 3 | 1 | 1 | 1 | 1 | 44 | 1 | 1 | 1 | 2 | 1 | 1 | 1 | 1 | 1 | 1 | 1 | 1 | 1 | 1 | 1 | 1 | 1 | 1 | 1 | 1 | 1 |
| yeps_YPTB1456 | 49 | 7 | 1 | 10 | 1 | 1 | 1 | 8 | 7 | 9 | 10 | 3 | 3 | 3 | 1 | 9 | 3 | 4 | 1 | 1 | 1 | 1 | 1 | 1 | 3 | 3 | 1 | 1 | 1 | 1 | 1 | 3 | 10 | 10 | 1 |
| yeps_YPTB1457 | 4 | 2 | 4 | 2 | 1 | 4 | 4 | 2 | 2 | 2 | 2 | 2 | 3 | 3 | 4 | 2 | 3 | 2 | 4 | 4 | 4 | 4 | 4 | 4 | 3 | 3 | 4 | 4 | 4 | 4 | 4 | 3 | 2 | 2 | 4 |
| yeps_YPTB1459 | 1 | 2 | 1 | 1 | 1 | 1 | 1 | 1 | 2 | 1 | 1 | 1 | 1 | 1 | 1 | 1 | 1 | 1 | 1 | 1 | 1 | 1 | 1 | 1 | 1 | 1 | 1 | 1 | 1 | 1 | 1 | 1 | 1 | 1 | 1 |
| yeps_YPTB1462 | 1 | 1 | 1 | 1 | 1 | 1 | 1 | 1 | 1 | 1 | 1 | 1 | 1 | 1 | 1 | 1 | 1 | 1 | 1 | 1 | 1 | 1 | 1 | 1 | 1 | 1 | 1 | 1 | 1 | 1 | 1 | 1 | 1 | 1 | 1 |
| yeps_YPTB1464 | 1 | 1 | 1 | 1 | 1 | 1 | 1 | 1 | 1 | 1 | 1 | 1 | 1 | 1 | 1 | 1 | 1 | 1 | 1 | 1 | 1 | 1 | 1 | 1 | 1 | 1 | 1 | 1 | 1 | 1 | 1 | 1 | 1 | 1 | 1 |
| yeps_YPTB1465 | 1 | 1 | 1 | 1 | 1 | 1 | 1 | 1 | 1 | 1 | 1 | 1 | 1 | 1 | 1 | 1 | 1 | 2 | 1 | 1 | 1 | 1 | 1 | 1 | 1 | 1 | 1 | 1 | 1 | 1 | 1 | 1 | 1 | 1 | 1 |
| yeps_YPTB1478 | 1 | 1 | 1 | | 1 | 1 | 1 | 4 | 1 | 3 | 3 | 3 | 1 | 1 | 1 | 3 | 1 | 4 | 1 | 1 | 1 | 1 | 1 | 1 | 1 | 1 | 1 | 1 | 1 | 1 | 1 | 1 | 3 | 3 | 1 |
| yeps_YPTB1479 | 1 | 3 | 1 | | 1 | 1 | 1 | 1 | 3 | 1 | 1 | 1 | 1 | 1 | 1 | 1 | 1 | 1 | 1 | 1 | 1 | 1 | 1 | 1 | 1 | 1 | 1 | 1 | 1 | 1 | 1 | 1 | 1 | 1 | 1 |
| yeps_YPTB1480 | 1 | 1 | 1 | | 1 | 1 | 1 | 1 | 1 | 1 | 1 | 1 | 1 | 1 | 1 | 1 | 1 | 1 | 1 | 1 | 1 | 1 | 1 | 1 | 1 | 1 | 1 | 1 | 1 | 1 | 1 | 1 | 1 | 1 | 1 |
| yeps_YPTB1481 | 2 | 2 | 2 | | 1 | 1 | 2 | 3 | 2 | 2 | 2 | 1 | 1 | 1 | 1 | 2 | 1 | 3 | 2 | 2 | 1 | 1 | 1 | 2 | 1 | 1 | 1 | 1 | 1 | 1 | 1 | 1 | 2 | 2 | 2 |
| yeps_YPTB1517 | 4 | 7 | 4 | 8 | 1 | 1 | 4 | 6 | 7 | 8 | 8 | 1 | 1 | 1 | 1 | 8 | 1 | 4 | 4 | 4 | 1 | 1 | 1 | 4 | 1 | 1 | 1 | 1 | 1 | 1 | 1 | 1 | 8 | 8 | 4 |
| yeps_YPTB1518 | 4 | 6 | 4 | 1 | 1 | 1 | 4 | 4 | 6 | 1 | 1 | 1 | 3 | 1 | 1 | 1 | 1 | 4 | 4 | 4 | 1 | 1 | 1 | 4 | 18 | 18 | 1 | 1 | 1 | 1 | 1 | 18 | 1 | 1 | 4 |
| yeps_YPTB1520 | 2 | 2 | 2 | 4 | 1 | 1 | 2 | 2 | 2 | 4 | 4 | 1 | 1 | 1 | 1 | 4 | 1 | 2 | 2 | 2 | 1 | 1 | 1 | 2 | 1 | 1 | 1 | 1 | 1 | 1 | 1 | 1 | 4 | 4 | 2 |
| yeps_YPTB1521 | 4 | 8 | 4 | 9 | 1 | 1 | 4 | 8 | 8 | 9 | 9 | 3 | 1 | 1 | 1 | 9 | 1 | 4 | 4 | 4 | 1 | 1 | 1 | 4 | 1 | 1 | 1 | 1 | 1 | 1 | 1 | 1 | 9 | 9 | 4 |
| yeps_YPTB1522 | 1 | 1 | 1 | 5 | 1 | 1 | 1 | 1 | 1 | 5 | 5 | 1 | 1 | 1 | 1 | 5 | 1 | 1 | 1 | 1 | 1 | 1 | 1 | 1 | 1 | 1 | 1 | 1 | 1 | 1 | 1 | 1 | 5 | 5 | 1 |
| yeps_YPTB1526 | 2 | 2 | 2 | 6 | 1 | 1 | 2 | 5 | 4 | 6 | 6 | 1 | 1 | 1 | 1 | 6 | 1 | 2 | 2 | 2 | 1 | 1 | 1 | 2 | 1 | 1 | 1 | 1 | 1 | 1 | 1 | 1 | 6 | 6 | 2 |
| yeps_YPTB1527 | 2 | 2 | 2 | 4 | 1 | 1 | 2 | 2 | 2 | 4 | 4 | 3 | 1 | 1 | 1 | 4 | 1 | 2 | 2 | 2 | 1 | 1 | 1 | 2 | 1 | 1 | 1 | 1 | 1 | 1 | 1 | 1 | 4 | 4 | 2 |
| yeps_YPTB1528 | 3 | 4 | 3 | 6 | 1 | 1 | 3 | 2 | 4 | 6 | 6 | 2 | 2 | 2 | 1 | 6 | 2 | 3 | 3 | 3 | 1 | 1 | 1 | 3 | 2 | 2 | 1 | 1 | 1 | 1 | 1 | 2 | 6 | 6 | 3 |
| yeps_YPTB1536 | 2 | 1 | 2 | 1 | 1 | 1 | 2 | 1 | 1 | 1 | 1 | 1 | 1 | 1 | 1 | 1 | 1 | 1 | 2 | 2 | 1 | 1 | 1 | 2 | 1 | 1 | 1 | 1 | 1 | 1 | 1 | 1 | 1 | 1 | 2 |
| yeps_YPTB1538 | 1 | 2 | 1 | 2 | 1 | 1 | 1 | 5 | 2 | 2 | 2 | 1 | 1 | 1 | 1 | 2 | 1 | 3 | 1 | 1 | 1 | 1 | 1 | 1 | 1 | 1 | 1 | 1 | 1 | 1 | 1 | 1 | 2 | 2 | 1 |
| yeps_YPTB1542 | 3 | 6 | 3 | 9 | 1 | 1 | 3 | 7 | 6 | 9 | 9 | 1 | 1 | 1 | 1 | 9 | 1 | 3 | 3 | 3 | 1 | 1 | 1 | 3 | 1 | 1 | 1 | 1 | 1 | 1 | 1 | 1 | 9 | 9 | 3 |
| yeps_YPTB1543 | 1 | 1 | 1 | 1 | 1 | 1 | 1 | 1 | 1 | 1 | 1 | 3 | 1 | 1 | 1 | 1 | 1 | 1 | 1 | 1 | 1 | 1 | 1 | 1 | 1 | 1 | 1 | 1 | 1 | 1 | 1 | 1 | 1 | 1 | 1 |
| yeps_YPTB1548 | 3 | 6 | 3 | 7 | 1 | 1 | 3 | 7 | 6 | 7 | 7 | 2 | 1 | 1 | 1 | 7 | 1 | 3 | 3 | 3 | 1 | 1 | 1 | 3 | 1 | 1 | 1 | 1 | 1 | 1 | 1 | 1 | 7 | 7 | 3 |
| yeps_YPTB1552 | 4 | 6 | 4 | 7 | 1 | 1 | 4 | 7 | 6 | 7 | 7 | 3 | 1 | 1 | 1 | 7 | 1 | 4 | 4 | 4 | 1 | 1 | 1 | 4 | 1 | 1 | 1 | 1 | 1 | 1 | 1 | 1 | 7 | 7 | 4 |
| yeps_YPTB1556 | 10 | 7 | 3 | 9 | 1 | 1 | 3 | 8 | 7 | 9 | 9 | 1 | 1 | 1 | 1 | 9 | 1 | 4 | 3 | 3 | 1 | 1 | 1 | 3 | 1 | 1 | 1 | 1 | 1 | 1 | 1 | 1 | 9 | 9 | 3 |
| yeps_YPTB1560 | 2 | 8 | 9 | 10 | 1 | 1 | 9 | 1 | 8 | 10 | 10 | 3 | 1 | 6 | 1 | 10 | 6 | 4 | 2 | 2 | 1 | 1 | 1 | 2 | 1 | 1 | 1 | 1 | 1 | 1 | 1 | 1 | 10 | 10 | 2 |
| yeps_YPTB1561 | 4 | 8 | 4 | 1 | 1 | 1 | 4 | 1 | 8 | 1 | 1 | 3 | 1 | 1 | 1 | 1 | 1 | 10 | 4 | 4 | 1 | 1 | 1 | 4 | 1 | 1 | 1 | 1 | 1 | 1 | 1 | 1 | 1 | 1 | 4 |
| yeps_YPTB1562 | 4 | 6 | 4 | 6 | 1 | 1 | 4 | 1 | 6 | 6 | 6 | 3 | 1 | 1 | 1 | 6 | 1 | 3 | 4 | 4 | 1 | 1 | 1 | 4 | 1 | 1 | 1 | 1 | 1 | 1 | 1 | 1 | 6 | 6 | 4 |
| yeps_YPTB1563 | 3 | 4 | 3 | 3 | 1 | 1 | 3 | 4 | 4 | 3 | 3 | 3 | 1 | 1 | 1 | 3 | 1 | 3 | 3 | 3 | 1 | 1 | 1 | 3 | 1 | 1 | 1 | 1 | 1 | 1 | 1 | 1 | 3 | 3 | 3 |
| yeps_YPTB1574 | 3 | 4 | 3 | 3 | 1 | 1 | 3 | 5 | 4 | 3 | 3 | 1 | 1 | 1 | 1 | 3 | 1 | 2 | 3 | 3 | 1 | 1 | 1 | 3 | 1 | 1 | 1 | 1 | 1 | 1 | 1 | 1 | 3 | 3 | 3 |
| yeps_YPTB1575 | 1 | 4 | 1 | 7 | 1 | 1 | 1 | 5 | 4 | 7 | 7 | 1 | 1 | 1 | 1 | 7 | 1 | 3 | 1 | 1 | 1 | 1 | 1 | 1 | 1 | 1 | 1 | 1 | 1 | 1 | 1 | 1 | 7 | 7 | 1 |
| yeps_YPTB1577 | 3 | 6 | 3 | 8 | 1 | 1 | 3 | 7 | 6 | 8 | 8 | 1 | 1 | 1 | 1 | 8 | 1 | 3 | 3 | 3 | 1 | 1 | 1 | 3 | 1 | 1 | 1 | 1 | 1 | 1 | 1 | 1 | 8 | 8 | 3 |
| yeps_YPTB1579 | 4 | 7 | 4 | 1 | 1 | 1 | 4 | 8 | 7 | 1 | 1 | 3 | 1 | 1 | 1 | 1 | 1 | 11 | 4 | 4 | 1 | 1 | 1 | 4 | 1 | 1 | 1 | 1 | 1 | 1 | 1 | 1 | 1 | 1 | 4 |
| yeps_YPTB1580 | 9 | 6 | 5 | 8 | 1 | 1 | 5 | 7 | 6 | 8 | 8 | 3 | 1 | 1 | 1 | 8 | 1 | 6 | 5 | 5 | 1 | 1 | 1 | 5 | 1 | 1 | 1 | 1 | 1 | 1 | 1 | 1 | 8 | 8 | 5 |
| yeps_YPTB1604 | 3 | 4 | 3 | 4 | 1 | 1 | 3 | 4 | 4 | 4 | 4 | 4 | 1 | 1 | 1 | 4 | 1 | 4 | 3 | 3 | 1 | 1 | 1 | 3 | 1 | 1 | 1 | 1 | 1 | 1 | 1 | 1 | 4 | 4 | 3 |
| yeps_YPTB1605 | 1 | 1 | 23 | 1 | 1 | 1 | 1 | 1 | 1 | 1 | 1 | 1 | 1 | 1 | 1 | 1 | 1 | 1 | 1 | 1 | 1 | 1 | 1 | 1 | 1 | 1 | 1 | 1 | 1 | 1 | 1 | 1 | 1 | 1 | 1 |
| yeps_YPTB1609 | 4 | 7 | 4 | 5 | 1 | 1 | 4 | 4 | 7 | 5 | 5 | 1 | 1 | 1 | 1 | 5 | 1 | 5 | 4 | 4 | 1 | 1 | 1 | 4 | 1 | 1 | 1 | 1 | 1 | 1 | 1 | 1 | 5 | 5 | 4 |
| yeps_YPTB1611 | 1 | 1 | 1 | 1 | 1 | 1 | 1 | 1 | 4 | 1 | 1 | 1 | 1 | 1 | 1 | 1 | 1 | 3 | 1 | 1 | 1 | 1 | 1 | 1 | 1 | 1 | 1 | 1 | 1 | 1 | 1 | 1 | 1 | 1 | 1 |
| yeps_YPTB1612 | 3 | 1 | 3 | 1 | 1 | 1 | 3 | 4 | 1 | 1 | 1 | 1 | 1 | 1 | 1 | 1 | 1 | 3 | 3 | 3 | 1 | 1 | 1 | 3 | 1 | 1 | 1 | 1 | 1 | 1 | 1 | 1 | 1 | 1 | 3 |
| yeps_YPTB1613 | 1 | 4 | 1 | 1 | 1 | 1 | 1 | 1 | 4 | 1 | 1 | 1 | 1 | 1 | 1 | 1 | 1 | 2 | 1 | 1 | 1 | 1 | 1 | 1 | 1 | 1 | 1 | 1 | 1 | 1 | 1 | 1 | 1 | 1 | 1 |
| yeps_YPTB1633 | 1 | 1 | 1 | 1 | 1 | 1 | 1 | 3 | 1 | 1 | 1 | 1 | 1 | 1 | 1 | 1 | 1 | 1 | 1 | 1 | 1 | 1 | 1 | 1 | 1 | 1 | 1 | 1 | 1 | 1 | 1 | 1 | 1 | 1 | 1 |
| yeps_YPTB1634 | 1 | 3 | 1 | 4 | 1 | 1 | 1 | 1 | 3 | 4 | 4 | 1 | 1 | 1 | 1 | 4 | 1 | 1 | 1 | 1 | 1 | 1 | 1 | 1 | 1 | 1 | 1 | 1 | 1 | 1 | 1 | 1 | 4 | 4 | 1 |
| yeps_YPTB1635 | 4 | 8 | 4 | 11 | 1 | 1 | 4 | 1 | 8 | 11 | 11 | 1 | 1 | 1 | 1 | 11 | 1 | 4 | 4 | 4 | 1 | 1 | 1 | 4 | 1 | 1 | 1 | 1 | 1 | 1 | 1 | 1 | 11 | 11 | 4 |
| yeps_YPTB1641 | 2 | 4 | 2 | 3 | 1 | 1 | 2 | 3 | 4 | 3 | 3 | 2 | 1 | 1 | 1 | 3 | 1 | 2 | 2 | 2 | 1 | 1 | 1 | 2 | 1 | 1 | 1 | 1 | 1 | 1 | 1 | 1 | 3 | 3 | 2 |
| yeps_YPTB1642 | 1 | 1 | 1 | 2 | 1 | 1 | 1 | 1 | 1 | 2 | 2 | 1 | 1 | 1 | 1 | 2 | 1 | 1 | 1 | 1 | 1 | 1 | 1 | 1 | 1 | 1 | 1 | 1 | 1 | 1 | 1 | 1 | 2 | 2 | 1 |
| yeps_YPTB1645 | 2 | 6 | 2 | 8 | 1 | 1 | 2 | 9 | 6 | 8 | 8 | 3 | 1 | 1 | 1 | 8 | 1 | 5 | 2 | 2 | 1 | 1 | 1 | 2 | 1 | 1 | 1 | 1 | 1 | 1 | 1 | 1 | 8 | 8 | 2 |
| yeps_YPTB1646 | 1 | 1 | 1 | 1 | 1 | 1 | 1 | 1 | 1 | 1 | 1 | 1 | 1 | 1 | 1 | 1 | 1 | 1 | 1 | 1 | 1 | 1 | 1 | 1 | 1 | 1 | 1 | 1 | 1 | 1 | 1 | 1 | 1 | 1 | 1 |
| yeps_YPTB1648 | 1 | 1 | 1 | 1 | 1 | 1 | 1 | 4 | 1 | 1 | 1 | 1 | 1 | 1 | 1 | 1 | 1 | 2 | 1 | 1 | 1 | 1 | 1 | 1 | 1 | 1 | 1 | 1 | 1 | 1 | 1 | 1 | 1 | 1 | 1 |
| yeps_YPTB1649 | 1 | 4 | 1 | 1 | 1 | 1 | 1 | 5 | 4 | 1 | 1 | 1 | 1 | 1 | 1 | 1 | 1 | 2 | 1 | 1 | 1 | 1 | 1 | 1 | 1 | 1 | 1 | 1 | 1 | 1 | 1 | 1 | 1 | 1 | 1 |
| yeps_YPTB1650 | 1 | 2 | 1 | 2 | 1 | 1 | 1 | 2 | 2 | 2 | 2 | 2 | 1 | 1 | 1 | 2 | 1 | 2 | 1 | 1 | 1 | 1 | 1 | 1 | 1 | 1 | 1 | 1 | 1 | 1 | 1 | 1 | 2 | 2 | 1 |
| yeps_YPTB1652 | 1 | 6 | 1 | 4 | 1 | 1 | 1 | 4 | 6 | 4 | 4 | 2 | 1 | 1 | 1 | 4 | 1 | 1 | 1 | 1 | 1 | 1 | 1 | 1 | 1 | 1 | 1 | 1 | 1 | 1 | 1 | 1 | 4 | 4 | 1 |
| yeps_YPTB1653 | 1 | 1 | 1 | 1 | 1 | 1 | 1 | 1 | 1 | 1 | 1 | 1 | 1 | 1 | 1 | 1 | 1 | 2 | 1 | 1 | 1 | 1 | 1 | 1 | 1 | 1 | 1 | 1 | 1 | 1 | 1 | 1 | 1 | 1 | 1 |
| yeps_YPTB1654 | 1 | 1 | 1 | 1 | 1 | 1 | 1 | 1 | 1 | 1 | 1 | 1 | 1 | 1 | 1 | 1 | 1 | 1 | 1 | 1 | 1 | 1 | 1 | 1 | 1 | 1 | 1 | 1 | 1 | 1 | 1 | 1 | 1 | 1 | 1 |
| yeps_YPTB1655 | 1 | 1 | 1 | 1 | 1 | 1 | 1 | 1 | 1 | 1 | 1 | 1 | 1 | 1 | 1 | 1 | 1 | 1 | 1 | 1 | 1 | 1 | 1 | 1 | 1 | 1 | 1 | 1 | 1 | 1 | 1 | 1 | 1 | 1 | 1 |
| yeps_YPTB1669 | 1 | 1 | 1 | 2 | 1 | 1 | 1 | 1 | 1 | 2 | 2 | 1 | 1 | 1 | 1 | 2 | 1 | 2 | 1 | 1 | 1 | 1 | 1 | 1 | 1 | 1 | 1 | 1 | 1 | 1 | 1 | 1 | 2 | 2 | 1 |
| yeps_YPTB1670 | 1 | 1 | 1 | 2 | 1 | 1 | 1 | 1 | 1 | 2 | 2 | 14 | 1 | 1 | 1 | 2 | 1 | 2 | 1 | 1 | 1 | 1 | 1 | 1 | 1 | 1 | 1 | 1 | 1 | 1 | 1 | 1 | 2 | 2 | 1 |
| yeps_YPTB1674 | 1 | 1 | 1 | 1 | 1 | 1 | 1 | 1 | 1 | 1 | 1 | 1 | 1 | 1 | 1 | 1 | 1 | 1 | 1 | 1 | 1 | 1 | 1 | 1 | 1 | 1 | 1 | 1 | 1 | 1 | 1 | 1 | 1 | 1 | 1 |
| yeps_YPTB1676 | 1 | 6 | 1 | 7 | 1 | 1 | 1 | 6 | 6 | 1 | 7 | 1 | 1 | 1 | 1 | 1 | 1 | 1 | 1 | 1 | 1 | 1 | 1 | 1 | 1 | 1 | 1 | 1 | 1 | 1 | 1 | 1 | 7 | 7 | 1 |
| yeps_YPTB1677 | 1 | 1 | 1 | 1 | 1 | 1 | 1 | 6 | 1 | 1 | 1 | 3 | 1 | 1 | 1 | 1 | 1 | 2 | 1 | 1 | 1 | 1 | 1 | 1 | 1 | 1 | 1 | 1 | 1 | 1 | 1 | 1 | 1 | 1 | 1 |
| yeps_YPTB1678 | 1 | 1 | 1 | 1 | 1 | 1 | 1 | 1 | 1 | 1 | 1 | 1 | 1 | 1 | 1 | 1 | 1 | 4 | 1 | 1 | 1 | 1 | 1 | 1 | 1 | 1 | 1 | 1 | 1 | 1 | 1 | 1 | 1 | 1 | 1 |
| yeps_YPTB1679 | 1 | 7 | 1 | 8 | 1 | 1 | 1 | 1 | 7 | 8 | 8 | 3 | 1 | 1 | 1 | 8 | 1 | 4 | 1 | 1 | 1 | 1 | 1 | 1 | 1 | 1 | 1 | 1 | 1 | 1 | 1 | 1 | 8 | 8 | 1 |
| yeps_YPTB1682 | 1 | 7 | 1 | 10 | 1 | 1 | 1 | 1 | 8 | 10 | 10 | 3 | 1 | 1 | 1 | 10 | 1 | 4 | 1 | 1 | 1 | 1 | 1 | 1 | 1 | 1 | 1 | 1 | 1 | 1 | 1 | 1 | 10 | 10 | 1 |
| yeps_YPTB1683 | 4 | 7 | 4 | 2 | 4 | 4 | 4 | 9 | 7 | 2 | 2 | 3 | 4 | 4 | 4 | 2 | 4 | 5 | 4 | 4 | 4 | 4 | 4 | 4 | 4 | 4 | 4 | 4 | 4 | 4 | 4 | 4 | 2 | 2 | 4 |
| yeps_YPTB1684 | 59 | 9 | 1 | 13 | 89 | 1 | 1 | 10 | 9 | 12 | 13 | 3 | 1 | 1 | 1 | 12 | 1 | 5 | 1 | 1 | 1 | 1 | 1 | 1 | 1 | 1 | 1 | 1 | 1 | 1 | 1 | 1 | 13 | 13 | 1 |
| yeps_YPTB1686 | 1 | 5 | 1 | 2 | 1 | 1 | 1 | 2 | 5 | 2 | 2 | 3 | 1 | 1 | 1 | 2 | 1 | 2 | 1 | 1 | 1 | 1 | 1 | 1 | 1 | 1 | 1 | 1 | 1 | 1 | 1 | 1 | 2 | 2 | 1 |
| yeps_YPTB1687 | 1 | 8 | 1 | 13 | 1 | 1 | 1 | 2 | 9 | 12 | 13 | 3 | 1 | 1 | 1 | 12 | 1 | 5 | 1 | 1 | 1 | 1 | 1 | 1 | 1 | 1 | 1 | 1 | 1 | 1 | 1 | 1 | 13 | 13 | 1 |
| yeps_YPTB1688 | 1 | 7 | 1 | 9 | 1 | 1 | 1 | 3 | 1 | 9 | 9 | 3 | 1 | 1 | 1 | 9 | 1 | 4 | 1 | 1 | 1 | 1 | 1 | 1 | 1 | 1 | 1 | 1 | 1 | 1 | 1 | 1 | 9 | 9 | 1 |
| yeps_YPTB1691 | 3 | 6 | 3 | 10 | 1 | 1 | 3 | 7 | 6 | 9 | 10 | 1 | 1 | 1 | 1 | 9 | 1 | 2 | 3 | 3 | 1 | 1 | 1 | 3 | 1 | 1 | 1 | 1 | 1 | 1 | 1 | 1 | 10 | 10 | 3 |
| yeps_YPTB1693 | 3 | 5 | 3 | 4 | 1 | 1 | 3 | 6 | 5 | 7 | 4 | 1 | 1 | 1 | 1 | 7 | 1 | 4 | 3 | 3 | 1 | 1 | 1 | 3 | 1 | 1 | 1 | 1 | 1 | 1 | 1 | 1 | 4 | 4 | 3 |
| yeps_YPTB1696 | 5 | 9 | 5 | 3 | 1 | 1 | 5 | 10 | 9 | 3 | 3 | 3 | 1 | 1 | 1 | 3 | 1 | 6 | 5 | 5 | 1 | 1 | 1 | 5 | 1 | 1 | 1 | 1 | 1 | 1 | 1 | 1 | 3 | 3 | 5 |

| | | | | | | | | | | | | | | | | | | | | | | | | | | | | | | | | | | | |
|---|---|---|---|---|---|---|---|---|---|---|---|---|---|---|---|---|---|---|---|---|---|---|---|---|---|---|---|---|---|---|---|---|---|---|---|
| yeps_YPTB1697 | 1 | 2 | 1 | 1 | 1 | 1 | 1 | 1 | 2 | 1 | 1 | 2 | 1 | 1 | 1 | 1 | 1 | 1 | 1 | 1 | 1 | 1 | 1 | 1 | 1 | 1 | 1 | 1 | 1 | 1 | 1 | 1 | 1 | 1 | 1 |
| yeps_YPTB1699 | 1 | 2 | 1 | 2 | 1 | 1 | 1 | 2 | 2 | 2 | 2 | 3 | 1 | 1 | 1 | 2 | 1 | 2 | 1 | 1 | 1 | 1 | 1 | 1 | 1 | 1 | 1 | 1 | 1 | 1 | 1 | 1 | 2 | 2 | 1 |
| yeps_YPTB1701 | 1 | 3 | 1 | 5 | 1 | 1 | 1 | 2 | 3 | 5 | 5 | 3 | 1 | 1 | 1 | 5 | 1 | 4 | 1 | 1 | 1 | 1 | 1 | 1 | 1 | 1 | 1 | 1 | 1 | 1 | 1 | 1 | 5 | 5 | 1 |
| yeps_YPTB1703 | 4 | 6 | 4 | 8 | 1 | 1 | 4 | 5 | 6 | 8 | 8 | 3 | 1 | 1 | 1 | 8 | 1 | 2 | 4 | 4 | 1 | 1 | 1 | 4 | 1 | 1 | 1 | 1 | 1 | 1 | 1 | 1 | 8 | 8 | 4 |
| yeps_YPTB1704 | 2 | 3 | 2 | 3 | 1 | 1 | 2 | 3 | 3 | 3 | 3 | 1 | 1 | 1 | 1 | 3 | 1 | 2 | 2 | 2 | 1 | 1 | 1 | 2 | 1 | 1 | 1 | 1 | 1 | 1 | 1 | 1 | 3 | 3 | 2 |
| yeps_YPTB1705 | 1 | 1 | 1 | 1 | 1 | 1 | 1 | 1 | 1 | 1 | 1 | 1 | 1 | 1 | 1 | 1 | 1 | 1 | 1 | 1 | 1 | 1 | 1 | 1 | 1 | 1 | 1 | 1 | 1 | 1 | 1 | 1 | 1 | 1 | 1 |
| yeps_YPTB1708 | 1 | 1 | 1 | 1 | 1 | 1 | 1 | 1 | 1 | 6 | 1 | 2 | 1 | 1 | 1 | 6 | 1 | 1 | 1 | 1 | 1 | 1 | 1 | 1 | 1 | 1 | 1 | 1 | 1 | 1 | 1 | 1 | 1 | 1 | 1 |
| yeps_YPTB1709 | 3 | 1 | 3 | 9 | 1 | 1 | 3 | 7 | 6 | 8 | 9 | 1 | 1 | 1 | 1 | 8 | 1 | 3 | 3 | 3 | 1 | 1 | 1 | 3 | 1 | 1 | 1 | 1 | 1 | 1 | 1 | 1 | 9 | 9 | 3 |
| yeps_YPTB1710 | 1 | 1 | 1 | 1 | 1 | 1 | 1 | 1 | 1 | 1 | 1 | 1 | 1 | 1 | 1 | 1 | 1 | 1 | 1 | 1 | 1 | 1 | 1 | 1 | 1 | 1 | 1 | 1 | 1 | 1 | 1 | 1 | 1 | 1 | 1 |
| yeps_YPTB1711 | 1 | 1 | 1 | 2 | 1 | 1 | 1 | 2 | 1 | 2 | 2 | 1 | 1 | 1 | 1 | 2 | 1 | 1 | 1 | 1 | 1 | 1 | 1 | 1 | 1 | 25 | 1 | 1 | 1 | 1 | 1 | 1 | 2 | 2 | 1 |
| yeps_YPTB1712 | 1 | 1 | 1 | 1 | 1 | 1 | 1 | 1 | 1 | 1 | 1 | 1 | 1 | 1 | 1 | 1 | 1 | 1 | 1 | 1 | 1 | 1 | 1 | 1 | 1 | 1 | 1 | 1 | 1 | 1 | 1 | 1 | 1 | 1 | 1 |
| yeps_YPTB1713 | 1 | 5 | 1 | 6 | 1 | 1 | 1 | 1 | 5 | 6 | 6 | 1 | 1 | 1 | 1 | 6 | 1 | 1 | 1 | 1 | 1 | 1 | 1 | 1 | 1 | 1 | 1 | 1 | 1 | 1 | 1 | 1 | 6 | 6 | 1 |
| yeps_YPTB1714 | 23 | 5 | 1 | 7 | 1 | 1 | 1 | 5 | 5 | 7 | 7 | 1 | 1 | 1 | 1 | 7 | 1 | 1 | 1 | 1 | 1 | 1 | 1 | 1 | 1 | 1 | 1 | 1 | 1 | 1 | 1 | 1 | 7 | 7 | 1 |
| yeps_YPTB1716 | 2 | 4 | 2 | 2 | 1 | 1 | 2 | 5 | 4 | 2 | 2 | 1 | 1 | 1 | 1 | 2 | 1 | 2 | 2 | 2 | 1 | 1 | 1 | 2 | 1 | 1 | 1 | 1 | 1 | 1 | 1 | 1 | 2 | 2 | 2 |
| yeps_YPTB1718 | 1 | 2 | 1 | 1 | 1 | 1 | 1 | 3 | 2 | 1 | 1 | 1 | 1 | 1 | 1 | 1 | 1 | 1 | 1 | 1 | 1 | 1 | 1 | 1 | 1 | 1 | 1 | 1 | 1 | 1 | 1 | 1 | 1 | 1 | 1 |
| yeps_YPTB1719 | 6 | 7 | 6 | 1 | 1 | 1 | 6 | 8 | 7 | 1 | 1 | 1 | 1 | 1 | 1 | 1 | 1 | 1 | 6 | 6 | 1 | 1 | 1 | 6 | 1 | 1 | 1 | 1 | 1 | 1 | 1 | 1 | 1 | 1 | 6 |
| yeps_YPTB1722 | 1 | 2 | 1 | 2 | 1 | 1 | 1 | 2 | 2 | 2 | 2 | 1 | 1 | 1 | 1 | 2 | 1 | 3 | 1 | 1 | 1 | 1 | 1 | 1 | 1 | 1 | 1 | 1 | 1 | 1 | 1 | 1 | 2 | 2 | 1 |
| yeps_YPTB1725 | 8 | 6 | 8 | 9 | 1 | 1 | 8 | 6 | 6 | 9 | 9 | 1 | 1 | 1 | 1 | 9 | 1 | 4 | 3 | 3 | 1 | 1 | 1 | 3 | 1 | 1 | 1 | 1 | 1 | 1 | 1 | 1 | 9 | 9 | 8 |
| yeps_YPTB1728 | 2 | 4 | 2 | 5 | 1 | 1 | 2 | 3 | 4 | 5 | 5 | 1 | 1 | 1 | 1 | 5 | 1 | 3 | 2 | 2 | 1 | 1 | 1 | 2 | 1 | 1 | 1 | 1 | 1 | 1 | 1 | 1 | 5 | 5 | 2 |
| yeps_YPTB1729 | 3 | 7 | 3 | 13 | 1 | 1 | 3 | 10 | 7 | 12 | 13 | 1 | 1 | 1 | 1 | 12 | 1 | 4 | 3 | 3 | 1 | 1 | 1 | 3 | 1 | 1 | 1 | 1 | 1 | 1 | 1 | 1 | 13 | 13 | 3 |
| yeps_YPTB1912 | 4 | 8 | 4 | 12 | 1 | 1 | 4 | 9 | 8 | 11 | 12 | 3 | 1 | 1 | 1 | 11 | 1 | 5 | 4 | 4 | 1 | 1 | 1 | 4 | 1 | 1 | 1 | 1 | 1 | 1 | 1 | 1 | 12 | 12 | 4 |
| yeps_YPTB1915 | 5 | 5 | 5 | 2 | 1 | 1 | 5 | 1 | 5 | 6 | 2 | 3 | 1 | 1 | 1 | 6 | 1 | 1 | 5 | 5 | 1 | 1 | 1 | 5 | 1 | 1 | 1 | 1 | 1 | 1 | 1 | 1 | 2 | 2 | 5 |
| yeps_YPTB1916 | 4 | 2 | 4 | 3 | 1 | 1 | 4 | 1 | 2 | 5 | 3 | 3 | 1 | 1 | 1 | 5 | 1 | 4 | 4 | 4 | 1 | 1 | 1 | 4 | 1 | 1 | 1 | 1 | 1 | 1 | 1 | 1 | 3 | 3 | 4 |
| yeps_YPTB1917 | 3 | 2 | 3 | 5 | 1 | 1 | 3 | 1 | 2 | 5 | 5 | 1 | 1 | 1 | 1 | 5 | 1 | 3 | 3 | 3 | 1 | 1 | 1 | 3 | 1 | 1 | 1 | 1 | 1 | 1 | 1 | 1 | 5 | 5 | 3 |
| yeps_YPTB1919 | 5 | 10 | 5 | 13 | 3 | 3 | 5 | 12 | 10 | 13 | 13 | 3 | 3 | 7 | 3 | 13 | 7 | 5 | 5 | 5 | 3 | 3 | 3 | 5 | 3 | 3 | 3 | 3 | 3 | 3 | 3 | 3 | 13 | 13 | 5 |
| yeps_YPTB1920 | 1 | 1 | 1 | 1 | 1 | 1 | 1 | 3 | 1 | 1 | 1 | 1 | 1 | 1 | 1 | 1 | 1 | 1 | 1 | 1 | 1 | 1 | 1 | 1 | 1 | 1 | 1 | 1 | 1 | 1 | 1 | 1 | 1 | 1 | 1 |
| yeps_YPTB1926 | 3 | 4 | 3 | 6 | 1 | 1 | 3 | 4 | 4 | 5 | 6 | 1 | 1 | 1 | 1 | 5 | 1 | 3 | 3 | 3 | 1 | 1 | 1 | 3 | 17 | 17 | 1 | 1 | 1 | 1 | 1 | 17 | 6 | 6 | 3 |
| yeps_YPTB1927 | 2 | 4 | 2 | 7 | 1 | 1 | 2 | 2 | 4 | 7 | 7 | 1 | 1 | 1 | 1 | 7 | 1 | 2 | 2 | 2 | 1 | 1 | 1 | 2 | 1 | 1 | 1 | 1 | 1 | 1 | 1 | 1 | 7 | 7 | 2 |
| yeps_YPTB1928 | 2 | 3 | 2 | 6 | 1 | 1 | 2 | 4 | 3 | 6 | 6 | 1 | 1 | 1 | 1 | 6 | 1 | 1 | 2 | 2 | 1 | 1 | 1 | 2 | 1 | 1 | 1 | 1 | 1 | 1 | 1 | 1 | 6 | 6 | 2 |
| yeps_YPTB1929 | 3 | 5 | 3 | 8 | 1 | 1 | 3 | 6 | 5 | 8 | 8 | 1 | 1 | 1 | 1 | 8 | 1 | 1 | 3 | 3 | 1 | 1 | 1 | 3 | 1 | 1 | 1 | 1 | 1 | 1 | 1 | 1 | 8 | 8 | 3 |
| yeps_YPTB1931 | 3 | | 3 | 6 | 1 | 1 | 3 | 1 | 5 | 6 | 6 | 1 | 1 | 1 | 1 | 6 | 1 | 1 | 3 | 3 | 1 | 1 | 1 | 3 | 1 | 1 | 1 | 1 | 1 | 1 | 1 | 1 | 6 | 6 | 3 |
| yeps_YPTB1933 | 3 | 3 | 3 | 6 | 1 | 1 | 3 | 5 | 3 | 3 | 6 | 1 | 1 | 1 | 1 | 3 | 1 | 7 | 3 | 3 | 1 | 1 | 1 | 3 | 1 | 1 | 1 | 1 | 1 | 1 | 1 | 1 | 6 | 6 | 3 |
| yeps_YPTB1944 | 2 | 4 | 2 | 6 | 1 | 1 | 2 | 5 | 4 | 6 | 6 | 1 | 1 | 1 | 1 | 6 | 1 | 1 | 2 | 2 | 1 | 1 | 1 | 2 | 1 | 1 | 1 | 1 | 1 | 1 | 1 | 1 | 6 | 6 | 2 |
| yeps_YPTB1946 | 1 | 3 | 1 | 3 | 1 | 1 | 1 | 3 | 3 | 3 | 3 | 3 | 1 | 1 | 1 | 3 | 1 | 3 | 1 | 1 | 1 | 1 | 1 | 1 | 1 | 1 | 1 | 1 | 1 | 1 | 1 | 1 | 3 | 3 | 1 |
| yeps_YPTB1950 | 3 | 7 | 3 | 11 | 1 | 1 | 3 | 9 | 8 | 11 | 11 | 1 | 1 | 1 | 1 | 11 | 1 | 4 | 3 | 3 | 1 | 1 | 1 | 3 | 1 | 1 | 1 | 1 | 1 | 1 | 1 | 1 | 11 | 11 | 3 |
| yeps_YPTB1951 | 4 | 7 | 4 | 4 | 1 | 1 | 4 | 2 | 7 | 4 | 4 | 1 | 1 | 1 | 1 | 4 | 1 | 4 | 4 | 4 | 1 | 1 | 1 | 4 | 1 | 1 | 1 | 1 | 1 | 1 | 1 | 1 | 4 | 4 | 4 |
| yeps_YPTB1952 | 4 | 7 | 4 | 4 | 1 | 1 | 4 | 3 | 7 | 4 | 4 | 3 | 1 | 1 | 1 | 4 | 1 | 3 | 4 | 4 | 1 | 1 | 1 | 4 | 1 | 1 | 1 | 1 | 1 | 1 | 1 | 1 | 4 | 4 | 4 |
| yeps_YPTB1954 | 1 | 1 | 1 | 3 | 1 | 1 | 1 | 1 | 1 | 3 | 3 | 1 | 1 | 1 | 1 | 3 | 1 | 3 | 1 | 1 | 1 | 1 | 1 | 1 | 1 | 1 | 1 | 1 | 1 | 1 | 1 | 1 | 3 | 3 | 1 |
| yeps_YPTB1957 | 5 | 10 | 5 | 15 | 1 | 1 | 5 | 8 | 11 | 14 | 15 | 3 | 1 | 1 | 1 | 14 | 1 | 8 | 5 | 5 | 1 | 1 | 1 | 5 | 1 | 1 | 1 | 1 | 1 | 1 | 1 | 1 | 15 | 15 | 5 |
| yeps_YPTB1965 | 3 | 6 | 3 | 8 | 1 | 1 | 3 | 1 | 6 | 8 | 8 | 3 | 1 | 1 | 1 | 8 | 1 | 1 | 3 | 3 | 1 | 1 | 1 | 3 | 1 | 1 | 1 | 1 | 1 | 1 | 1 | 1 | 8 | 8 | 3 |
| yeps_YPTB1968 | 1 | 2 | 1 | 7 | 1 | 1 | 1 | 6 | 2 | 7 | 7 | 1 | 1 | 1 | 1 | 7 | 1 | 2 | 1 | 1 | 1 | 1 | 1 | 1 | 1 | 1 | 1 | 1 | 1 | 1 | 1 | 1 | 7 | 7 | 1 |
| yeps_YPTB1969 | 1 | 2 | 1 | 2 | 1 | 1 | 1 | 2 | 2 | 2 | 2 | 1 | 1 | 1 | 1 | 2 | 1 | 2 | 1 | 1 | 1 | 1 | 1 | 1 | 1 | 1 | 1 | 1 | 1 | 1 | 1 | 1 | 2 | 2 | 1 |
| yeps_YPTB1972 | 5 | 9 | 5 | 11 | 1 | 1 | 5 | 6 | 9 | 11 | 11 | 3 | 1 | 1 | 1 | 11 | 1 | 12 | 5 | 5 | 1 | 1 | 1 | 5 | 1 | 1 | 1 | 1 | 1 | 1 | 1 | 1 | 11 | 11 | 5 |
| yeps_YPTB1974 | 4 | 9 | 4 | 11 | 1 | 1 | 4 | 10 | 9 | 11 | 11 | 3 | 1 | 1 | 1 | 11 | 1 | 5 | 4 | 4 | 1 | 1 | 1 | 4 | 1 | 1 | 1 | 1 | 1 | 1 | 1 | 1 | 11 | 11 | 4 |
| yeps_YPTB1975 | 3 | 7 | 3 | 10 | 1 | 1 | 3 | 8 | 7 | 10 | 10 | 1 | 1 | 1 | 1 | 10 | 1 | 4 | 3 | 3 | 1 | 1 | 1 | 3 | 1 | 1 | 1 | 1 | 1 | 1 | 1 | 1 | 10 | 10 | 3 |
| yeps_YPTB1977 | 2 | 2 | 2 | 2 | 1 | 1 | 2 | 2 | 2 | 2 | 2 | 1 | 1 | 1 | 1 | 2 | 1 | 1 | 2 | 2 | 1 | 1 | 1 | 2 | 1 | 1 | 1 | 1 | 1 | 1 | 1 | 1 | 2 | 2 | 2 |
| yeps_YPTB1978 | 3 | 7 | 3 | 6 | 1 | 1 | 3 | 8 | 7 | 6 | 6 | 1 | 1 | 1 | 1 | 6 | 1 | 4 | 3 | 3 | 1 | 1 | 1 | 3 | 1 | 1 | 1 | 1 | 1 | 1 | 1 | 1 | 6 | 6 | 3 |
| yeps_YPTB1983 | 2 | 1 | 2 | 1 | 1 | 1 | 2 | 1 | 1 | 1 | 1 | 1 | 1 | 1 | 1 | 1 | 1 | 2 | 2 | 2 | | | | 2 | 1 | 1 | | | | | | 1 | 1 | 1 | 2 |
| yeps_YPTB1984 | 1 | 1 | 1 | 1 | 1 | 1 | 1 | 1 | 1 | 1 | 1 | 1 | 1 | 1 | 1 | 1 | 1 | 1 | 1 | 1 | | | | 1 | 1 | 1 | | | | | | 1 | 1 | 1 | 1 |
| yeps_YPTB1985 | 3 | 3 | 3 | 1 | 1 | 1 | 3 | 3 | 3 | 1 | 1 | 1 | 1 | 1 | 1 | 1 | 1 | 3 | 3 | 3 | | | | 3 | 1 | 1 | | | | | | 1 | 1 | 1 | 3 |
| yeps_YPTB1986 | 3 | 3 | 3 | 3 | 1 | 1 | 3 | 5 | 3 | 3 | 3 | 1 | 1 | 1 | 1 | 3 | 1 | 3 | 3 | 3 | | | | 3 | 1 | 1 | | | | | | 1 | 3 | 3 | 3 |
| yeps_YPTB1987 | 2 | 3 | 2 | 4 | 1 | 1 | 2 | 1 | 3 | 4 | 4 | 1 | 1 | 1 | 1 | 4 | 1 | 2 | 2 | 2 | | | | 2 | 1 | 1 | | | | | | 1 | 4 | 4 | 2 |
| yeps_YPTB1990 | 2 | 4 | 2 | 2 | 1 | 1 | 2 | 5 | 4 | 2 | 2 | 1 | 1 | 1 | 1 | 2 | 1 | 2 | 2 | 2 | 1 | 1 | 1 | 2 | 1 | 1 | 1 | 1 | 1 | 1 | 1 | 1 | 2 | 2 | 2 |
| yeps_YPTB1991 | | 2 | 2 | 4 | 1 | 1 | 2 | 3 | 2 | 4 | 4 | 1 | 1 | 1 | 1 | 4 | 1 | 2 | 2 | 2 | 1 | 1 | 1 | 2 | 1 | 1 | 1 | 1 | 1 | 1 | 1 | 1 | 4 | 4 | 2 |
| yeps_YPTB1992 | 3 | 1 | 3 | 10 | 1 | 1 | 3 | 7 | 1 | 10 | 10 | 1 | 1 | 5 | 1 | 10 | 5 | 3 | 3 | 3 | 1 | 1 | 1 | 3 | 1 | 1 | 1 | 1 | 1 | 1 | 1 | 1 | 10 | 10 | 3 |
| yeps_YPTB1993 | 3 | 6 | 3 | 8 | 1 | 1 | 3 | 7 | 6 | 8 | 8 | 1 | 1 | 1 | 1 | 8 | 1 | 3 | 3 | 3 | 1 | 1 | 1 | 3 | 1 | 1 | 1 | 1 | 1 | 1 | 1 | 1 | 8 | 8 | 3 |
| yeps_YPTB1998 | 3 | 3 | 3 | 7 | 3 | 3 | 3 | 6 | 3 | 7 | 7 | 3 | 3 | 3 | 3 | 7 | 3 | 3 | 3 | 3 | 3 | 3 | 3 | 3 | 3 | 3 | 3 | 3 | 3 | 3 | 3 | 3 | 7 | 7 | 3 |
| yeps_YPTB2000 | 3 | 2 | 3 | 2 | 1 | 1 | 3 | 2 | 2 | 2 | 2 | 1 | 1 | 1 | 1 | 2 | 1 | 2 | 3 | 3 | 1 | 1 | 1 | 3 | 1 | 1 | 1 | 1 | 1 | 1 | 1 | 1 | 2 | 2 | 3 |
| yeps_YPTB2003 | 4 | 7 | 4 | 1 | 1 | 1 | 4 | 3 | 7 | 1 | 1 | 3 | 1 | 1 | 1 | 1 | 1 | 5 | 4 | 4 | 1 | 1 | 1 | 4 | 1 | 1 | 1 | 1 | 1 | 1 | 1 | 1 | 1 | 1 | 4 |
| yeps_YPTB2004 | 3 | 6 | 3 | 5 | 1 | 1 | 3 | 3 | 6 | 5 | 5 | 3 | 1 | 1 | 1 | 5 | 1 | 5 | 3 | 3 | 1 | 1 | 1 | 3 | 1 | 1 | 1 | 1 | 1 | 1 | 1 | 1 | 5 | 5 | 3 |
| yeps_YPTB2006 | 1 | 5 | 1 | 1 | 1 | 1 | 1 | 1 | 5 | 1 | 1 | 1 | 1 | 1 | 1 | 1 | 1 | 1 | 1 | 1 | 1 | 1 | 1 | 1 | 1 | 1 | 1 | 1 | 1 | 1 | 1 | 1 | 1 | 1 | 1 |
| yeps_YPTB2014 | 1 | 6 | 1 | 8 | 1 | 1 | 1 | 9 | 8 | 11 | 8 | 3 | 3 | 3 | 1 | 11 | 3 | 12 | 1 | 1 | 1 | 1 | 1 | 1 | 3 | 3 | 1 | 1 | 1 | 1 | 1 | 3 | 8 | 8 | 1 |
| yeps_YPTB2015 | 1 | 6 | 1 | 6 | 1 | 1 | 1 | 7 | 6 | 6 | 6 | 3 | 3 | 3 | 1 | 6 | 3 | 4 | 1 | 1 | 1 | 1 | 1 | 1 | 3 | 3 | 1 | 1 | 1 | 1 | 1 | 3 | 6 | 6 | 1 |
| yeps_YPTB2020 | 1 | 3 | 1 | 4 | 1 | 1 | 1 | 3 | 3 | 4 | 4 | 3 | 3 | 3 | 1 | 4 | 3 | 3 | 1 | 1 | 1 | 1 | 1 | 1 | 3 | 3 | 1 | 1 | 1 | 1 | 1 | 3 | 4 | 4 | 1 |
| yeps_YPTB2021 | 5 | 1 | 1 | 4 | 1 | 1 | 1 | 1 | 1 | 4 | 4 | 1 | 1 | 1 | 1 | 4 | 1 | 1 | 1 | 1 | 1 | 1 | 1 | 1 | 1 | 1 | 1 | 1 | 1 | 1 | 1 | 1 | 4 | 4 | 1 |
| yeps_YPTB2029 | 4 | 8 | 4 | 11 | 1 | 3 | 4 | 10 | 9 | 11 | 11 | 3 | 3 | 3 | 3 | 11 | 3 | 6 | 4 | 4 | 3 | 3 | 3 | 4 | 3 | 3 | 3 | 3 | 3 | 3 | 3 | 3 | 11 | 11 | 4 |
| yeps_YPTB2033 | 3 | 4 | 3 | 2 | 1 | 1 | 3 | 3 | 4 | 2 | 2 | 1 | 1 | 1 | 1 | 2 | 1 | 2 | 3 | 3 | 1 | 1 | 1 | 3 | 1 | 1 | 1 | 1 | 1 | 1 | 1 | 1 | 2 | 2 | 3 |
| yeps_YPTB2034 | 2 | 2 | 2 | 2 | 1 | 1 | 2 | 2 | 2 | 2 | 2 | 1 | 1 | 1 | 1 | 2 | 1 | 2 | 2 | 2 | 1 | 1 | 1 | 2 | 1 | 1 | 1 | 1 | 1 | 1 | 1 | 1 | 2 | 2 | 2 |
| yeps_YPTB2037 | 1 | 1 | 1 | 2 | 1 | 1 | 1 | 1 | 1 | 2 | 2 | 1 | 1 | 1 | 1 | 2 | 1 | 2 | 1 | 1 | 1 | 1 | 1 | 1 | 1 | 1 | 1 | 1 | 1 | 1 | 1 | 1 | 2 | 2 | 1 |
| yeps_YPTB2039 | 1 | 1 | 1 | 1 | 1 | 1 | 1 | 1 | 1 | 1 | 1 | 1 | 1 | 1 | 1 | 1 | 1 | 1 | 1 | 1 | 1 | 1 | 1 | 1 | 1 | 1 | 1 | 1 | 1 | 1 | 1 | 1 | 1 | 1 | 1 |
| yeps_YPTB2041 | 3 | 6 | 3 | 9 | 1 | 1 | 3 | 7 | 6 | 8 | 9 | 1 | 1 | 1 | 1 | 8 | 1 | 4 | 3 | 3 | 1 | 1 | 1 | 3 | 1 | 1 | 1 | 1 | 1 | 1 | 1 | 1 | 9 | 9 | 3 |
| yeps_YPTB2043 | 3 | 4 | 3 | 4 | 1 | 1 | 3 | 4 | 4 | 4 | 4 | 1 | 1 | 1 | 1 | 4 | 1 | 4 | 3 | 3 | 1 | 1 | 1 | 3 | 1 | 1 | 1 | 1 | 1 | 1 | 1 | 31 | 4 | 4 | 3 |
| yeps_YPTB2044 | 1 | 4 | 1 | 2 | 1 | 1 | 1 | 5 | 4 | 2 | 2 | 1 | 1 | 1 | 1 | 2 | 1 | 2 | 1 | 1 | 1 | 1 | 1 | 1 | 1 | 1 | 1 | 1 | 1 | 1 | 1 | 1 | 2 | 2 | 1 |
| yeps_YPTB2048 | 1 | 4 | 1 | 1 | 1 | 1 | 1 | 1 | 4 | 1 | 1 | 1 | 1 | 1 | 1 | 1 | 1 | 3 | 1 | 1 | 1 | 1 | 1 | 1 | 1 | 1 | 1 | 1 | 1 | 1 | 1 | 1 | 1 | 1 | 1 |
| yeps_YPTB2049 | 1 | 3 | 1 | 2 | 1 | 1 | 1 | 1 | 3 | 2 | 2 | 1 | 1 | 1 | 1 | 2 | 1 | 1 | 1 | 1 | 1 | 1 | 1 | 1 | 1 | 1 | 1 | 1 | 1 | 1 | 1 | 1 | 2 | 2 | 1 |
| yeps_YPTB2054 | 9 | 7 | 3 | 7 | 1 | 1 | 3 | 3 | 38 | 7 | 7 | 1 | 1 | 1 | 1 | 7 | 1 | 3 | 3 | 3 | 1 | 1 | 1 | 3 | 1 | 1 | 1 | 1 | 1 | 1 | 1 | 1 | 7 | 7 | 3 |
| yeps_YPTB2055 | 3 | 4 | 3 | 4 | 1 | 1 | 3 | 3 | 4 | 4 | 4 | 15 | 1 | 1 | 1 | 4 | 1 | 3 | 3 | 3 | 1 | 1 | 1 | 3 | 1 | 1 | 1 | 1 | 1 | 1 | 1 | 1 | 4 | 4 | 3 |
| yeps_YPTB2062 | 1 | 1 | 1 | 1 | 1 | 1 | 1 | 1 | 1 | 1 | 1 | 1 | 1 | 1 | 1 | 1 | 1 | 1 | 1 | 1 | 1 | 1 | 1 | 1 | 1 | 1 | 1 | 1 | 1 | 1 | 1 | 1 | 1 | 1 | 1 |
| yeps_YPTB2063 | 7 | 4 | 6 | 4 | 1 | 1 | 6 | 8 | 4 | 4 | 4 | 1 | 1 | 1 | 1 | 4 | 1 | 4 | 6 | 6 | 1 | 1 | 1 | 6 | 1 | 1 | 1 | 1 | 1 | 1 | 1 | 1 | 4 | 4 | 7 |
| yeps_YPTB2065 | 1 | 1 | 1 | 1 | 1 | 1 | 1 | 1 | 1 | 1 | 1 | 1 | 1 | 1 | 1 | 1 | 1 | 1 | 1 | 1 | 1 | 1 | 1 | 1 | 1 | 1 | 1 | 1 | 1 | 1 | 1 | 1 | 1 | 1 | 1 |
| yeps_YPTB2076 | 4 | 7 | 4 | 10 | 1 | 1 | 4 | 8 | 7 | 10 | 10 | 3 | 1 | 1 | 1 | 10 | 1 | 2 | 4 | 4 | 1 | 1 | 1 | 4 | 1 | 1 | 1 | 1 | 1 | 1 | 1 | 1 | 10 | 10 | 4 |
| yeps_YPTB2085 | 2 | 3 | 2 | 1 | 1 | 1 | 2 | 2 | 3 | 1 | 1 | 1 | 1 | 1 | 1 | 1 | 1 | 2 | 2 | 2 | 1 | 1 | 1 | 2 | 1 | 1 | 1 | 1 | 1 | 1 | 1 | 1 | 1 | 1 | 2 |
| yeps_YPTB2087 | 3 | 7 | 3 | 7 | 1 | 1 | 3 | 8 | 7 | 7 | 7 | 1 | 1 | 1 | 1 | 7 | 1 | 4 | 3 | 3 | 1 | 1 | 1 | 3 | 1 | 1 | 1 | 1 | 1 | 1 | 1 | 1 | 7 | 7 | 3 |
| yeps_YPTB2088 | 2 | 2 | 2 | 2 | 1 | 1 | 2 | 5 | 2 | 2 | 2 | 1 | 1 | 1 | 1 | 2 | 1 | 2 | 2 | 2 | 1 | 1 | 1 | 2 | 1 | 1 | 1 | 1 | 1 | 1 | 1 | 1 | 2 | 2 | 2 |
| yeps_YPTB2091 | 4 | 7 | 4 | 12 | 8 | 8 | 4 | 10 | 7 | 12 | 12 | 3 | 4 | 4 | 8 | 12 | 4 | 2 | 4 | 4 | 8 | 8 | 8 | 4 | 4 | 4 | 8 | 8 | 8 | 8 | 8 | 4 | 12 | 12 | 4 |
| yeps_YPTB2119 | 2 | 1 | 2 | 1 | 1 | 1 | 2 | 2 | 1 | 1 | 1 | 1 | 1 | 1 | 1 | 1 | 1 | 1 | 2 | 2 | 1 | 1 | 1 | 2 | 1 | 1 | 1 | 1 | 1 | 1 | 1 | 1 | 1 | 1 | 2 |
| yeps_YPTB2124 | 1 | 1 | 1 | 1 | 1 | 1 | 1 | 1 | 1 | 1 | 1 | 1 | 1 | 1 | 1 | 1 | 1 | 1 | 1 | 1 | 1 | 1 | 1 | 1 | 1 | 1 | 1 | 1 | 1 | 1 | 1 | 1 | 1 | 1 | 1 |
| yeps_YPTB2127 | 1 | 7 | 1 | 9 | 1 | 1 | 1 | 1 | 7 | 6 | 9 | 3 | 1 | 1 | 1 | 6 | 1 | 1 | 1 | 1 | 1 | 1 | 1 | 1 | 24 | 24 | 1 | 1 | 1 | 1 | 1 | 24 | 9 | 9 | 1 |
| yeps_YPTB2130 | 1 | 3 | 1 | 8 | 1 | 1 | 1 | 7 | 3 | 8 | 8 | 1 | 1 | 1 | 1 | 8 | 1 | 3 | 1 | 1 | 1 | 1 | 1 | 1 | 1 | 1 | 1 | 1 | 1 | 1 | 1 | 1 | 8 | 8 | 1 |
| yeps_YPTB2136 | 1 | 1 | 1 | 1 | 1 | 1 | 1 | 1 | 1 | 4 | 1 | 1 | 1 | 1 | 1 | 4 | 1 | 2 | 1 | 1 | 1 | 1 | 1 | 1 | 1 | 1 | 1 | 1 | 1 | 1 | 1 | 1 | 1 | 1 | 1 |
| yeps_YPTB2138 | 1 | 1 | 1 | 1 | 1 | 1 | 1 | 1 | 1 | 1 | 1 | 1 | 1 | 1 | 1 | 1 | 1 | 38 | 1 | 1 | 1 | 1 | 1 | 1 | 1 | 1 | 1 | 1 | 1 | 1 | 1 | 1 | 1 | 1 | 1 |
| yeps_YPTB2153 | 1 | 3 | 1 | 1 | 1 | 1 | 1 | 1 | 3 | 2 | 1 | 1 | 1 | 1 | 1 | 2 | 1 | 1 | 1 | 1 | 1 | 1 | 1 | 1 | 1 | 1 | 1 | 1 | 1 | 1 | 1 | 1 | 1 | 1 | 1 |
| yeps_YPTB2154 | 1 | 1 | 1 | 4 | 1 | 1 | 1 | 1 | 1 | 3 | 4 | 1 | 1 | 1 | 1 | 3 | 1 | 1 | 1 | 1 | 1 | 1 | 1 | 1 | 1 | 1 | 1 | 1 | 1 | 1 | 1 | 1 | 4 | 4 | 1 |
| yeps_YPTB2155 | 4 | 5 | 3 | 5 | 1 | 1 | 3 | 5 | 5 | 5 | 5 | 1 | 1 | 1 | 1 | 5 | 1 | 4 | 3 | 3 | 1 | 1 | 1 | 3 | 1 | 1 | 1 | 1 | 1 | 1 | 1 | 1 | 5 | 5 | 3 |
| yeps_YPTB2157 | 4 | 7 | 4 | 10 | 1 | 1 | 4 | 9 | 7 | 10 | 10 | 1 | 1 | 1 | 1 | 10 | 1 | 5 | 4 | 4 | 1 | 1 | 1 | 4 | 1 | 1 | 1 | 1 | 1 | 1 | 1 | 1 | 10 | 10 | 4 |
| yeps_YPTB2158 | 1 | 5 | 1 | 5 | 1 | 1 | 1 | 7 | 5 | 5 | 5 | 3 | 1 | 1 | 1 | 5 | 1 | 4 | 1 | 1 | 1 | 1 | 1 | 1 | 1 | 1 | 1 | 1 | 1 | 1 | 1 | 1 | 5 | 5 | 1 |
| yeps_YPTB2159 | 4 | 7 | 4 | 10 | 1 | 1 | 4 | 8 | 7 | 9 | 10 | 3 | 1 | 1 | 1 | 9 | 1 | 5 | 4 | 4 | 1 | 1 | 1 | 4 | 1 | 1 | 1 | 1 | 1 | 1 | 1 | 1 | 10 | 10 | 4 |
| yeps_YPTB2161 | 8 | 7 | 8 | 12 | 1 | 1 | 8 | 10 | 7 | 11 | 12 | 1 | 1 | 1 | 1 | 11 | 1 | 4 | 8 | 8 | 1 | 1 | 1 | 8 | 1 | 1 | 1 | 1 | 1 | 1 | 1 | 1 | 12 | 12 | 8 |
| yeps_YPTB2162 | 4 | 4 | 4 | 3 | 1 | 1 | 4 | 5 | 4 | 3 | 3 | 3 | 1 | 1 | 1 | 3 | 1 | 1 | 4 | 4 | 1 | 1 | 1 | 4 | 1 | 1 | 1 | 1 | 1 | 1 | 1 | 1 | 3 | 3 | 4 |
| yeps_YPTB2168 | 1 | 1 | 1 | 1 | 1 | 1 | 1 | 4 | 1 | 1 | 1 | 1 | 1 | 1 | 1 | 1 | 1 | 1 | 1 | 1 | 1 | 1 | 1 | 1 | 1 | 1 | 1 | 1 | 1 | 1 | 1 | 1 | 1 | 1 | 1 |
| yeps_YPTB2170 | 5 | 4 | 5 | 6 | 1 | 34 | 5 | 6 | 4 | 7 | 6 | 1 | 1 | 1 | 34 | 7 | 1 | 1 | 5 | 5 | 34 | 34 | 34 | 5 | 1 | 1 | 34 | 34 | 34 | 34 | 34 | 1 | 6 | 6 | 5 |
| yeps_YPTB2173 | 3 | 6 | 3 | 1 | 1 | 1 | 3 | 3 | 6 | 1 | 1 | 1 | 1 | 1 | 1 | 1 | 1 | 1 | 3 | 3 | 1 | 1 | 1 | 3 | 1 | 1 | 1 | 1 | 1 | 1 | 1 | 1 | 1 | 1 | 3 |
| yeps_YPTB2175 | 1 | 6 | 1 | 7 | 1 | 1 | 1 | 1 | 6 | 7 | 7 | 3 | 1 | 1 | 1 | 7 | 1 | 2 | 1 | 1 | 1 | 1 | 1 | 1 | 1 | 1 | 1 | 1 | 1 | 1 | 1 | 1 | 7 | 7 | 1 |
| yeps_YPTB2176 | 3 | 1 | 3 | 8 | 1 | 48 | 3 | 1 | 1 | 7 | 8 | 1 | 1 | 1 | 48 | 7 | 1 | 6 | 3 | 3 | 48 | 48 | 48 | 3 | 1 | 1 | 48 | 48 | 48 | 48 | 48 | 1 | 8 | 8 | 3 |
| yeps_YPTB2177 | 3 | 3 | 3 | 1 | 1 | 1 | 3 | 4 | 3 | 1 | 1 | 3 | 1 | 1 | 1 | 1 | 1 | 2 | 3 | 3 | 1 | 1 | 1 | 3 | 1 | 1 | 1 | 1 | 1 | 1 | 1 | 1 | 1 | 1 | 3 |
| yeps_YPTB2178 | 1 | 4 | 1 | 7 | 1 | 1 | 1 | 5 | 4 | 6 | 7 | 1 | 1 | 1 | 1 | 6 | 1 | 3 | 58 | 1 | 1 | 1 | 1 | 1 | 1 | 1 | 1 | 1 | 1 | 1 | 1 | 1 | 7 | 7 | 1 |
| yeps_YPTB2179 | 1 | 2 | 1 | 7 | 1 | 1 | 1 | 5 | 2 | 7 | 7 | 1 | 1 | 1 | 1 | 7 | 1 | 4 | 1 | 1 | 1 | 1 | 1 | 1 | 1 | 1 | 1 | 1 | 1 | 1 | 1 | 1 | 7 | 7 | 1 |
| yeps_YPTB2184 | 1 | 6 | 1 | 9 | 1 | 1 | 1 | 7 | 6 | 9 | 9 | 3 | 1 | 1 | 1 | 9 | 1 | 4 | 1 | 1 | 1 | 1 | 1 | 1 | 1 | 1 | 1 | 1 | 1 | 1 | 1 | 1 | 9 | 9 | 1 |
| yeps_YPTB2200 | 3 | 2 | 3 | 6 | 1 | 1 | 3 | 5 | 2 | 6 | 6 | 1 | 1 | 1 | 1 | 6 | 1 | 4 | 3 | 3 | 1 | 1 | 1 | 3 | 1 | 1 | 1 | 1 | 1 | 1 | 1 | 1 | 6 | 6 | 3 |
| yeps_YPTB2206 | 1 | 2 | 1 | 1 | 1 | 1 | 1 | 1 | 2 | 1 | 1 | 3 | 3 | 3 | 1 | 1 | 3 | 5 | 1 | 1 | 1 | 1 | 1 | 1 | 3 | 3 | 1 | 1 | 1 | 1 | 1 | 3 | 1 | 1 | 1 |
| yeps_YPTB2209 | 1 | 2 | 1 | 5 | 1 | 1 | 1 | 4 | 2 | 5 | 5 | 2 | 1 | 1 | 1 | 5 | 1 | 2 | 1 | 1 | 1 | 1 | 1 | 1 | 1 | 1 | 1 | 1 | 1 | 1 | 1 | 1 | 5 | 5 | 1 |
| yeps_YPTB2210 | 1 | 4 | 1 | 4 | 1 | 1 | 1 | 7 | 4 | 4 | 4 | 2 | 3 | 3 | 1 | 4 | 3 | 4 | 1 | 1 | 1 | 1 | 1 | 1 | 3 | 3 | 1 | 1 | 1 | 1 | 1 | 3 | 4 | 4 | 1 |
| yeps_YPTB2211 | 1 | 4 | 1 | 9 | 1 | 1 | 1 | 6 | 4 | 8 | 9 | 2 | 3 | 3 | 1 | 8 | 3 | 4 | 1 | 1 | 1 | 1 | 1 | 1 | 3 | 3 | 1 | 1 | 1 | 1 | 1 | 3 | 9 | 9 | 1 |
| yeps_YPTB2217 | 1 | 1 | 1 | 2 | 1 | 1 | 1 | 1 | 1 | 2 | 2 | 3 | 3 | 3 | 1 | 2 | 3 | 1 | 1 | 1 | 1 | 1 | 1 | 1 | 3 | 3 | 1 | 1 | 1 | 1 | 1 | 3 | 2 | 2 | 1 |
| yeps_YPTB2218 | 1 | 3 | 1 | 6 | 1 | 1 | 1 | 5 | 3 | 6 | 6 | 3 | 3 | 3 | 1 | 6 | 3 | 1 | 1 | 1 | 1 | 1 | 1 | 1 | 3 | 3 | 1 | 1 | 1 | 1 | 1 | 3 | 6 | 6 | 1 |
| yeps_YPTB2221 | 1 | 4 | 1 | 1 | 1 | 4 | 1 | 1 | 4 | 1 | 1 | 3 | 4 | 4 | 4 | 1 | 4 | 1 | 1 | 1 | 4 | 4 | 4 | 1 | 4 | 4 | 4 | 4 | 4 | 4 | 4 | 4 | 1 | 1 | 5 |
| yeps_YPTB2222 | 1 | 2 | 1 | 1 | 1 | | 1 | 1 | 2 | 1 | 1 | 1 | 1 | 1 | | 1 | 1 | 1 | 1 | 1 | | | | 1 | 1 | 1 | | | | | | 1 | 1 | 1 | 1 |
| yeps_YPTB2225 | 1 | 8 | 1 | 11 | 1 | 4 | 1 | 9 | 8 | 11 | 11 | 3 | 4 | 4 | 4 | 11 | 4 | 5 | 1 | 1 | 4 | 4 | 4 | 1 | 4 | 4 | 4 | 4 | 4 | 4 | 4 | 4 | 11 | 11 | 1 |
| yeps_YPTB2226 | 1 | 3 | 1 | 5 | 1 | 3 | 1 | 3 | 3 | 4 | 5 | 1 | 3 | 3 | 3 | 4 | 3 | 3 | 1 | 1 | 3 | 3 | 3 | 1 | 3 | 3 | 3 | 3 | 3 | 3 | 3 | 3 | 5 | 5 | 1 |
| yeps_YPTB2227 | 1 | 1 | 1 | 2 | 1 | 3 | 1 | 3 | 1 | 2 | 2 | 1 | 3 | 3 | 3 | 2 | 3 | 1 | 1 | 1 | 3 | 3 | 3 | 1 | 3 | 3 | 3 | 3 | 3 | 3 | 3 | 3 | 2 | 2 | 1 |
| yeps_YPTB2228 | 1 | 8 | 9 | 4 | 1 | 4 | 9 | 4 | 8 | 4 | 4 | 3 | 4 | 4 | 4 | 4 | 4 | 5 | 9 | 9 | 4 | 4 | 4 | 9 | 4 | 4 | 4 | 4 | 4 | 4 | 4 | 4 | 4 | 4 | 1 |
| yeps_YPTB2229 | 1 | 1 | 1 | 1 | 1 | 1 | 1 | 1 | 1 | 1 | 1 | 3 | 1 | 1 | 1 | 1 | 1 | 1 | 1 | 1 | 1 | 1 | 1 | 1 | 1 | 1 | 1 | 1 | 1 | 1 | 1 | 1 | 1 | 1 | 1 |
| yeps_YPTB2230 | 1 | 2 | 1 | 6 | 1 | 3 | 1 | 2 | 2 | 5 | 6 | 2 | 3 | 3 | 3 | 5 | 3 | 2 | 1 | 1 | 3 | 3 | 3 | 1 | 3 | 3 | 3 | 3 | 3 | 3 | 3 | 3 | 6 | 6 | 1 |
| yeps_YPTB2231 | 1 | 4 | 1 | 4 | 1 | 4 | 1 | 9 | 8 | 10 | 4 | 3 | 6 | 6 | 4 | 10 | 6 | 5 | 1 | 1 | 4 | 4 | 4 | 1 | 6 | 6 | 4 | 4 | 4 | 4 | 4 | 6 | 4 | 4 | 1 |
| yeps_YPTB2232 | 1 | 7 | 1 | 9 | 1 | 6 | 1 | 7 | 7 | 9 | 9 | 3 | 6 | 6 | 6 | 9 | 6 | 5 | 1 | 1 | 6 | 6 | 6 | 1 | 6 | 6 | 6 | 6 | 6 | 6 | 6 | 6 | 9 | 9 | 1 |
| yeps_YPTB2239 | 3 | 1 | 3 | 1 | 1 | 1 | 3 | 5 | 1 | 1 | 1 | 3 | 1 | 1 | 1 | 1 | 1 | 6 | 3 | 3 | 1 | 1 | 1 | 3 | 1 | 1 | 1 | 1 | 1 | 1 | 1 | 1 | 1 | 1 | 3 |
| yeps_YPTB2241 | 5 | 8 | 5 | 12 | 13 | 1 | 5 | 10 | 9 | 12 | 12 | 3 | 1 | 1 | 1 | 12 | 1 | 5 | 5 | 5 | 1 | 1 | 1 | 5 | 1 | 1 | 1 | 1 | 1 | 1 | 1 | 1 | 12 | 12 | 5 |
| yeps_YPTB2242 | 1 | 1 | 1 | 1 | 1 | 1 | 1 | 1 | 1 | 1 | 1 | 1 | 1 | 1 | 1 | 1 | 1 | 1 | 1 | 1 | 1 | 1 | 1 | 1 | 1 | 1 | 1 | 1 | 1 | 1 | 1 | 11 | 1 | 1 | 1 |
| yeps_YPTB2246 | 1 | 1 | 1 | 2 | 1 | 1 | 1 | 1 | 1 | 2 | 2 | 1 | 1 | 1 | 1 | 2 | 1 | 1 | 1 | 1 | 1 | 1 | 1 | 1 | 1 | 1 | 1 | 1 | 1 | 1 | 1 | 1 | 2 | 2 | 1 |
| yeps_YPTB2248 | 3 | 3 | 3 | 3 | 1 | 1 | 3 | 5 | 3 | 3 | 3 | 1 | 1 | 1 | 1 | 3 | 1 | 3 | 3 | 3 | 1 | 1 | 1 | 3 | 1 | 1 | 1 | 1 | 1 | 1 | 1 | 1 | 3 | 3 | 3 |
| yeps_YPTB2251 | 1 | 3 | 1 | 4 | 1 | 1 | 1 | 1 | 3 | 4 | 4 | 1 | 1 | 1 | 1 | 4 | 1 | 3 | 1 | 1 | 1 | 1 | 1 | 1 | 1 | 1 | 1 | 1 | 1 | 1 | 1 | 1 | 4 | 4 | 1 |
| yeps_YPTB2252 | 1 | 1 | 1 | 1 | 1 | 1 | 1 | 1 | 1 | 1 | 1 | 1 | 1 | 1 | 1 | 1 | 1 | 1 | 1 | 1 | 1 | 1 | 1 | 1 | 1 | 1 | 1 | 1 | 1 | 1 | 1 | 1 | 1 | 1 | 1 |
| yeps_YPTB2254 | 1 | 3 | 1 | 4 | 1 | 1 | 1 | 1 | 3 | 4 | 4 | 1 | 1 | 1 | 1 | 4 | 1 | 2 | 1 | 1 | 1 | 1 | 1 | 1 | 1 | 1 | 1 | 1 | 1 | 1 | 1 | 1 | 4 | 4 | 1 |
| yeps_YPTB2258 | 3 | 3 | 3 | 7 | 1 | 1 | 3 | 1 | 3 | 7 | 7 | 1 | 1 | 1 | 1 | 7 | 1 | 3 | 3 | 3 | 1 | 1 | 1 | 3 | 1 | 1 | 1 | 1 | 1 | 1 | 1 | 1 | 7 | 7 | 3 |
| yeps_YPTB2262 | 1 | 6 | 1 | 2 | 1 | 1 | 1 | 8 | 6 | 2 | 2 | 3 | 3 | 3 | 1 | 2 | 3 | 4 | 1 | 1 | 1 | 1 | 1 | 1 | 3 | 3 | 1 | 1 | 1 | 1 | 1 | 3 | 2 | 2 | 1 |
| yeps_YPTB2264 | 1 | 1 | 1 | 1 | 1 | 1 | 1 | 2 | 1 | 1 | 1 | 1 | 1 | 1 | 1 | 1 | 1 | 3 | 1 | 1 | 1 | 1 | 1 | 1 | 1 | 1 | 1 | 1 | 1 | 1 | 1 | 1 | 1 | 1 | 1 |
| yeps_YPTB2265 | 1 | 5 | 1 | 7 | 1 | 1 | 1 | 6 | 5 | 7 | 7 | 3 | 4 | 4 | 1 | 7 | 4 | 2 | 1 | 1 | 1 | 1 | 1 | 1 | 4 | 4 | 1 | 1 | 1 | 1 | 1 | 4 | 7 | 7 | 1 |
| yeps_YPTB2267 | 1 | 2 | 1 | 1 | 1 | 1 | 1 | 1 | 2 | 1 | 1 | 1 | 1 | 1 | 1 | 1 | 1 | 1 | 1 | 1 | 1 | 1 | 1 | 1 | 1 | 1 | 1 | 1 | 1 | 1 | 1 | 1 | 1 | 1 | 1 |
| yeps_YPTB2268 | 1 | 1 | 1 | 1 | 1 | 1 | 1 | 1 | 1 | 1 | 1 | 2 | 1 | 1 | 1 | 1 | 1 | 1 | 1 | 1 | 1 | 1 | 1 | 1 | 1 | 1 | 1 | 1 | 1 | 1 | 1 | 1 | 1 | 1 | 1 |
| yeps_YPTB2269 | 1 | 1 | 1 | 1 | 1 | 1 | 1 | 1 | 1 | 1 | 1 | 1 | 1 | 1 | 1 | 1 | 1 | 1 | 1 | 1 | 1 | 1 | 1 | 1 | 1 | 1 | 1 | 1 | 1 | 1 | 1 | 1 | 1 | 1 | 1 |
| yeps_YPTB2270 | 1 | 7 | 1 | 8 | 1 | 1 | 1 | 1 | 7 | 8 | 8 | 3 | 4 | 4 | 1 | 8 | 4 | 1 | 1 | 1 | 1 | 1 | 1 | 1 | 4 | 4 | 1 | 1 | 1 | 1 | 1 | 4 | 8 | 8 | 1 |
| yeps_YPTB2276 | 4 | 6 | 4 | 2 | 1 | 4 | 4 | 7 | 6 | 2 | 2 | 3 | 1 | 1 | 4 | 2 | 1 | 5 | 4 | 4 | 4 | 4 | 4 | 4 | 1 | 1 | 4 | 4 | 4 | 4 | 4 | 1 | 2 | 2 | 4 |
| yeps_YPTB2279 | 3 | 5 | 3 | 1 | 1 | 3 | 3 | 2 | 5 | 1 | 1 | 1 | 1 | 1 | 3 | 1 | 1 | 2 | 3 | 3 | 3 | 3 | 3 | 3 | 1 | 1 | 3 | 3 | 3 | 3 | 3 | 1 | 1 | 1 | 3 |
| yeps_YPTB2280 | 1 | 1 | 1 | 1 | 1 | 1 | 1 | 1 | 1 | 1 | 1 | 1 | 1 | 1 | 1 | 1 | 1 | 1 | 2 | 2 | 1 | 1 | 1 | 2 | 1 | 1 | 1 | 1 | 1 | 1 | 1 | 1 | 1 | 1 | 1 |
| yeps_YPTB2281 | 1 | 1 | 1 | 7 | 1 | 1 | 1 | 1 | 1 | 7 | 7 | 1 | 1 | 1 | 1 | 7 | 1 | 5 | 1 | 1 | 1 | 1 | 1 | 1 | 1 | 1 | 1 | 1 | 1 | 1 | 1 | 1 | 7 | 7 | 1 |
| yeps_YPTB2282 | 1 | 1 | 1 | 1 | 1 | 1 | 1 | 4 | 1 | 1 | 1 | 1 | 1 | 1 | 1 | 1 | 1 | 1 | 1 | 1 | 1 | 1 | 1 | 1 | 1 | 1 | 1 | 1 | 1 | 1 | 1 | 1 | 1 | 1 | 1 |
| yeps_YPTB2284 | 1 | 1 | 1 | 4 | 1 | 1 | 1 | 1 | 1 | 3 | 4 | 1 | 1 | 1 | 1 | 3 | 1 | 1 | 1 | 1 | 1 | 1 | 1 | 1 | 1 | 1 | 1 | 1 | 1 | 1 | 1 | 1 | 4 | 4 | 1 |
| yeps_YPTB2285 | 6 | 3 | 3 | 3 | 1 | 3 | 3 | 4 | 3 | 5 | 3 | 1 | 1 | 1 | 3 | 5 | 1 | 3 | 3 | 3 | 3 | 3 | 3 | 3 | 1 | 1 | 3 | 3 | 3 | 3 | 3 | 1 | 3 | 3 | 3 |
| yeps_YPTB2292 | 3 | 3 | 3 | 8 | 1 | 1 | 3 | 6 | 3 | 8 | 8 | 3 | 1 | 1 | 1 | 8 | 1 | 4 | 3 | 3 | 1 | 1 | 1 | 3 | 1 | 1 | 1 | 1 | 1 | 1 | 1 | 1 | 8 | 8 | 3 |
| yeps_YPTB2293 | 3 | 3 | 3 | 3 | 1 | 1 | 3 | 6 | 3 | 3 | 3 | 3 | 1 | 1 | 1 | 3 | 1 | 4 | 3 | 3 | 1 | 1 | 1 | 3 | 1 | 1 | 1 | 1 | 1 | 1 | 1 | 1 | 3 | 3 | 3 |
| yeps_YPTB2295 | 1 | 1 | 1 | 1 | 1 | 1 | 1 | 1 | 1 | 1 | 1 | 1 | 1 | 1 | 1 | 1 | 1 | 2 | 1 | 1 | 1 | 1 | 1 | 1 | 1 | 1 | 1 | 1 | 1 | 1 | 1 | 1 | 1 | 1 | 1 |
| yeps_YPTB2298 | 3 | 1 | 3 | 6 | 1 | 1 | 3 | 3 | 1 | 6 | 6 | 3 | 1 | 1 | 1 | 6 | 1 | 1 | 3 | 3 | 1 | 1 | 1 | 3 | 1 | 1 | 1 | 1 | 1 | 1 | 1 | 1 | 6 | 6 | 3 |
| yeps_YPTB2299 | 3 | 7 | 3 | 5 | 1 | 1 | 3 | 1 | 7 | 5 | 5 | 3 | 1 | 1 | 1 | 5 | 1 | 4 | 3 | 3 | 1 | 1 | 1 | 3 | 1 | 1 | 1 | 1 | 1 | 1 | 1 | 1 | 5 | 5 | 3 |
| yeps_YPTB2300 | 4 | 8 | 4 | 10 | 1 | 1 | 4 | 9 | 8 | 10 | 10 | 3 | 1 | 1 | 1 | 10 | 1 | 5 | 4 | 4 | 1 | 1 | 1 | 4 | 1 | 1 | 1 | 1 | 1 | 1 | 1 | 1 | 10 | 10 | 4 |
| yeps_YPTB2303 | 1 | 2 | 1 | 8 | 1 | 1 | 1 | 4 | 2 | 8 | 8 | 3 | 1 | 1 | 1 | 8 | 1 | 5 | 1 | 1 | 1 | 1 | 1 | 1 | 1 | 1 | 1 | 1 | 1 | 1 | 1 | 1 | 8 | 8 | 1 |
| yeps_YPTB2304 | 4 | 7 | 4 | 4 | 1 | 1 | 4 | 1 | 7 | 4 | 4 | 1 | 1 | 1 | 1 | 4 | 1 | 1 | 4 | 4 | 1 | 1 | 1 | 4 | 1 | 1 | 1 | 1 | 1 | 1 | 1 | 1 | 4 | 4 | 4 |
| yeps_YPTB2309 | 3 | 6 | 3 | 1 | 1 | 1 | 3 | 7 | 6 | 1 | 1 | 1 | 1 | 1 | 1 | 1 | 1 | 1 | 3 | 3 | 1 | 1 | 1 | 3 | 1 | 1 | 1 | 1 | 1 | 1 | 1 | 1 | 1 | 1 | 3 |
| yeps_YPTB2310 | 8 | 7 | 8 | 6 | 1 | 1 | 8 | 9 | 7 | 6 | 6 | 1 | 1 | 1 | 1 | 6 | 1 | 4 | 8 | 8 | 1 | 1 | 1 | 8 | 1 | 1 | 1 | 1 | 1 | 1 | 1 | 1 | 6 | 6 | 8 |
| yeps_YPTB2313 | 4 | 4 | 4 | 4 | 1 | 1 | 4 | 6 | 4 | 4 | 4 | 1 | 1 | 1 | 1 | 4 | 1 | 4 | 4 | 4 | 1 | 1 | 1 | 4 | 1 | 1 | 1 | 1 | 1 | 1 | 1 | 1 | 4 | 4 | 4 |
| yeps_YPTB2314 | 1 | 1 | 1 | 2 | 1 | 1 | 1 | 1 | 1 | 2 | 2 | 1 | 1 | 1 | 1 | 2 | 1 | 1 | 1 | 1 | 1 | 1 | 1 | 1 | 1 | 1 | 1 | 1 | 1 | 1 | 1 | 1 | 2 | 2 | 1 |
| yeps_YPTB2382 | 3 | 3 | 3 | 4 | | 2 | 3 | 3 | 3 | 4 | 4 | 2 | 2 | 2 | 2 | 4 | 2 | 1 | 3 | 3 | 2 | 2 | 2 | 3 | 2 | 2 | 2 | 2 | 2 | 2 | 2 | 2 | 4 | 4 | 3 |
| yeps_YPTB2387 | 1 | 1 | 1 | 1 | 1 | 1 | 1 | 1 | 1 | 1 | 1 | 1 | 1 | 1 | 1 | 1 | 1 | 1 | 1 | 1 | 1 | 1 | 1 | 1 | 1 | 1 | 1 | 1 | 1 | 1 | 1 | 1 | 1 | 1 | 1 |
| yeps_YPTB2388 | 2 | 4 | 2 | 2 | 1 | 2 | 2 | 1 | 4 | 2 | 2 | 2 | 2 | 2 | 2 | 2 | 2 | 1 | 2 | 2 | 2 | 2 | 2 | 2 | 2 | 2 | 2 | 2 | 2 | 2 | 2 | 2 | 2 | 2 | 2 |
| yeps_YPTB2393 | 1 | 177 | 1 | 4 | 6 | 3 | 1 | 7 | 5 | 4 | 4 | 1 | 3 | 3 | 3 | 182 | 41 | 4 | 1 | 1 | 30 | 3 | 3 | 1 | 3 | 3 | 41 | 3 | 3 | 3 | 3 | 3 | 4 | 4 | 1 |
| yeps_YPTB2395 | 1 | 6 | 1 | 8 | 1 | 1 | 1 | 4 | 7 | 8 | 8 | 3 | 1 | 1 | 1 | 8 | 1 | 4 | 1 | 1 | 1 | 1 | 1 | 1 | 1 | 1 | 1 | 1 | 1 | 1 | 1 | 1 | 8 | 8 | 1 |
| yeps_YPTB2398 | 4 | 4 | 4 | 4 | 1 | 4 | 4 | 3 | 4 | 4 | 4 | 3 | 4 | 4 | 4 | 4 | 4 | 5 | 4 | 4 | 4 | 4 | 4 | 4 | 4 | 4 | 4 | 4 | 4 | 4 | 4 | 4 | 4 | 4 | 4 |

| | | | | | | | | | | | | | | | | | | | | | | | | | | | | | | | | | | | |
|---|---|---|---|---|---|---|---|---|---|---|---|---|---|---|---|---|---|---|---|---|---|---|---|---|---|---|---|---|---|---|---|---|---|---|---|
| yeps_YPTB2399 | 1 | 9 | 1 | 11 | 1 | 4 | 1 | 10 | 9 | 11 | 11 | 3 | 4 | 4 | 4 | 11 | 4 | 7 | 1 | 1 | 4 | 4 | 4 | 1 | 4 | 4 | 4 | 4 | 4 | 4 | 4 | 4 | 11 | 11 | 1 |
| yeps_YPTB2401 | 1 | 7 | 1 | 8 | 1 | 3 | 1 | 4 | 7 | 8 | 8 | 1 | 3 | 3 | 3 | 8 | 3 | 4 | 1 | 1 | 3 | 3 | 3 | 1 | 3 | 3 | 3 | 3 | 3 | 3 | 3 | 3 | 8 | 8 | 1 |
| yeps_YPTB2413 | 1 | 1 | 1 | 2 | 1 | 1 | 1 | 1 | 1 | 2 | 2 | 1 | 1 | 1 | 1 | 2 | 1 | 1 | 1 | 1 | 1 | 1 | 1 | 1 | 1 | 1 | 1 | 1 | 1 | 1 | 1 | 1 | 2 | 2 | 1 |
| yeps_YPTB2418 | 1 | 1 | 1 | 5 | 1 | 1 | 1 | 1 | 1 | 5 | 5 | 2 | 1 | 1 | 1 | 5 | 1 | 3 | 1 | 1 | 1 | 1 | 1 | 1 | 1 | 1 | 1 | 1 | 1 | 1 | 1 | 1 | 5 | 5 | 1 |
| yeps_YPTB2419 | 4 | 1 | 4 | 6 | 1 | 4 | 4 | 3 | 26 | 6 | 6 | 3 | 1 | 1 | 4 | 6 | 1 | 5 | 4 | 4 | 4 | 4 | 4 | 4 | 1 | 1 | 4 | 4 | 4 | 4 | 4 | 1 | 6 | 6 | 4 |
| yeps_YPTB2420 | 1 | 1 | 1 | 1 | 1 | 1 | 1 | 1 | 1 | 1 | 1 | 1 | 1 | 1 | 1 | 1 | 1 | 1 | 1 | 1 | 1 | 1 | 1 | 1 | 1 | 1 | 1 | 1 | 1 | 1 | 1 | 1 | 1 | 1 | 1 |
| yeps_YPTB2421 | 8 | 6 | 8 | 9 | 1 | 8 | 8 | 2 | 6 | 1 | 9 | 3 | 1 | 1 | 8 | 1 | 1 | 6 | 8 | 8 | 8 | 8 | 8 | 8 | 1 | 1 | 8 | 8 | 8 | 8 | 8 | 1 | 9 | 9 | 8 |
| yeps_YPTB2425 | 1 | 1 | 1 | 1 | 1 | 1 | 1 | 1 | 1 | 1 | 1 | 1 | 1 | 1 | 1 | 1 | 1 | 2 | 1 | 1 | 1 | 1 | 1 | 1 | 1 | 1 | 1 | 1 | 1 | 1 | 1 | 1 | 1 | 1 | 1 |
| yeps_YPTB2430 | 3 | 6 | 3 | 7 | 1 | 1 | 3 | 6 | 6 | 7 | 7 | 1 | 1 | 1 | 1 | 7 | 1 | 4 | 3 | 3 | 1 | 1 | 1 | 3 | 1 | 1 | 1 | 1 | 1 | 1 | 1 | 1 | 7 | 7 | 3 |
| yeps_YPTB2435 | 3 | 7 | 3 | 8 | 1 | 1 | 3 | 3 | 7 | 8 | 8 | 3 | 1 | 1 | 1 | 8 | 1 | 4 | 3 | 3 | 1 | 1 | 1 | 3 | 1 | 1 | 1 | 1 | 1 | 1 | 1 | 1 | 8 | 8 | 3 |
| yeps_YPTB2438 | 3 | 2 | 3 | 5 | 1 | 1 | 3 | 3 | 2 | 5 | 5 | 3 | 6 | 31 | 1 | 5 | 1 | 3 | 3 | 3 | 1 | 1 | 1 | 3 | 1 | 1 | 1 | 1 | 1 | 1 | 1 | 1 | 5 | 5 | 3 |
| yeps_YPTB2439 | 3 | 1 | 3 | 6 | 1 | 1 | 3 | 6 | 1 | 6 | 6 | 3 | 1 | 1 | 1 | 6 | 1 | 5 | 3 | 3 | 1 | 1 | 1 | 3 | 1 | 1 | 1 | 1 | 1 | 1 | 1 | 1 | 6 | 6 | 3 |
| yeps_YPTB2440 | 4 | 9 | 4 | 5 | 1 | 4 | 4 | 5 | 9 | 5 | 5 | 3 | 1 | 7 | 4 | 5 | 7 | 5 | 4 | 4 | 4 | 4 | 4 | 4 | 1 | 1 | 4 | 4 | 4 | 4 | 4 | 1 | 5 | 5 | 4 |
| yeps_YPTB2441 | 3 | 1 | 3 | 1 | 1 | 3 | 3 | 1 | 1 | 1 | 1 | 2 | 1 | 1 | 3 | 1 | 1 | 2 | 3 | 3 | 3 | 3 | 3 | 3 | 1 | 1 | 3 | 3 | 3 | 3 | 3 | 1 | 1 | 1 | 3 |
| yeps_YPTB2449 | 1 | 3 | 1 | 5 | 1 | 1 | 1 | 2 | 3 | 5 | 5 | 3 | 1 | 1 | 1 | 5 | 1 | 2 | 1 | 1 | 1 | 1 | 1 | 1 | 1 | 1 | 1 | 1 | 1 | 1 | 1 | 1 | 5 | 5 | 1 |
| yeps_YPTB2451 | 2 | 1 | 2 | 1 | 7 | 1 | 2 | 2 | 1 | 6 | 1 | 1 | 1 | 1 | 1 | 6 | 1 | 2 | 2 | 2 | 1 | 1 | 1 | 2 | 1 | 1 | 1 | 1 | 1 | 1 | 1 | 1 | 1 | 1 | 2 |
| yeps_YPTB2452 | 1 | 1 | 1 | 4 | 1 | 1 | 1 | 2 | 1 | 4 | 4 | 1 | 1 | 1 | 1 | 4 | 1 | 2 | 1 | 1 | 1 | 1 | 1 | 1 | 1 | 1 | 1 | 1 | 1 | 1 | 1 | 1 | 4 | 4 | 1 |
| yeps_YPTB2453 | 1 | 1 | 1 | 1 | 1 | 1 | 1 | 1 | 1 | 1 | 1 | 1 | 1 | 1 | 1 | 1 | 1 | 1 | 1 | 1 | 1 | 1 | 1 | 1 | 1 | 1 | 1 | 1 | 1 | 1 | 1 | 1 | 1 | 1 | 1 |
| yeps_YPTB2463 | 4 | 8 | 4 | 9 | 1 | 1 | 4 | 3 | 8 | 9 | 9 | 3 | 1 | 1 | 1 | 9 | 1 | 3 | 4 | 4 | 1 | 1 | 1 | 4 | 1 | 1 | 1 | 1 | 1 | 1 | 1 | 1 | 9 | 9 | 4 |
| yeps_YPTB2464 | 3 | 1 | 3 | 3 | 1 | 1 | 3 | 3 | 1 | 5 | 3 | 3 | 1 | 1 | 1 | 5 | 1 | 3 | | 3 | 1 | 1 | 1 | 3 | 1 | 1 | 1 | 1 | 1 | 1 | 1 | 1 | 3 | 3 | 3 |
| yeps_YPTB2465 | 4 | 4 | 4 | 7 | 1 | 1 | 4 | 4 | 4 | 7 | 7 | 3 | 1 | 1 | 1 | 7 | 1 | 5 | 4 | 4 | 1 | 1 | 1 | 4 | 1 | 1 | 1 | 1 | 1 | 1 | 1 | 1 | 7 | 7 | 4 |
| yeps_YPTB2466 | 3 | 6 | 3 | 4 | 1 | 1 | 3 | 4 | 6 | 4 | 4 | 3 | 1 | 1 | 1 | 4 | 1 | 4 | 3 | 3 | 1 | 1 | 1 | 3 | 1 | 1 | 1 | 1 | 1 | 1 | 1 | 1 | 4 | 4 | 3 |
| yeps_YPTB2469 | 4 | 5 | 4 | 3 | 1 | 1 | 4 | 3 | 25 | 3 | 3 | 3 | 1 | 1 | 1 | 3 | 1 | 3 | 4 | 4 | 1 | 1 | 1 | 4 | 1 | 1 | 1 | 1 | 1 | 1 | 1 | 1 | 3 | 3 | 4 |
| yeps_YPTB2472 | 4 | 8 | 4 | 1 | 1 | 1 | 4 | 9 | 8 | 1 | 1 | 3 | 1 | 1 | 1 | 1 | 1 | 5 | 4 | 4 | 1 | 1 | 1 | 4 | 1 | 1 | 1 | 1 | 1 | 1 | 1 | 1 | 1 | 1 | 4 |
| yeps_YPTB2473 | 2 | 1 | 2 | 2 | 1 | 1 | 2 | 1 | 1 | 2 | 2 | 1 | 1 | 1 | 1 | 2 | 1 | 3 | 2 | 2 | 1 | 1 | 1 | 2 | 1 | 1 | 1 | 1 | 1 | 1 | 1 | 1 | 2 | 2 | 2 |
| yeps_YPTB2476 | 1 | 1 | 1 | 1 | 1 | 1 | 1 | 3 | 1 | 1 | 1 | 1 | 1 | 1 | 1 | 1 | 1 | 1 | 1 | 1 | 1 | 1 | 1 | 1 | 1 | 1 | 1 | 1 | 1 | 1 | 1 | 1 | 1 | 1 | 1 |
| yeps_YPTB2480 | 1 | 1 | 1 | 1 | 1 | 1 | 1 | 1 | 1 | 1 | 1 | 1 | 1 | 1 | 1 | 1 | 1 | 1 | 1 | 1 | 1 | 1 | 1 | 1 | 1 | 1 | 1 | 1 | 1 | 1 | 1 | 1 | 1 | 1 | 1 |
| yeps_YPTB2481 | 1 | 1 | 1 | 2 | 1 | 1 | 1 | 1 | 1 | 2 | 2 | 1 | 1 | 1 | 1 | 2 | 1 | 2 | 1 | 1 | 1 | 1 | 1 | 1 | 1 | 1 | 1 | 1 | 1 | 1 | 1 | 1 | 2 | 2 | 1 |
| yeps_YPTB2482 | 3 | 6 | 3 | 8 | 1 | 1 | 3 | 7 | 6 | 8 | 8 | 1 | 1 | 1 | 1 | 8 | 1 | 4 | 3 | 3 | 1 | 1 | 1 | 3 | 1 | 1 | 1 | 1 | 1 | 1 | 1 | 1 | 8 | 8 | 3 |
| yeps_YPTB2483 | 1 | 1 | 1 | 1 | 1 | 1 | 1 | 1 | 1 | 1 | 1 | 1 | 1 | 1 | 1 | 1 | 1 | 1 | 1 | 1 | 1 | 1 | 1 | 1 | 1 | 1 | 1 | 1 | 1 | 1 | 1 | 1 | 1 | 1 | 1 |
| yeps_YPTB2484 | 1 | 1 | 1 | 1 | 1 | 1 | 1 | 1 | 1 | 1 | 1 | 1 | 1 | 1 | 1 | 1 | 1 | 1 | 1 | 1 | 1 | 1 | 1 | 1 | 1 | 1 | 1 | 1 | 1 | 1 | 1 | 1 | 1 | 1 | 1 |
| yeps_YPTB2488 | 4 | 6 | 4 | 3 | 1 | 1 | 4 | 3 | 6 | 3 | 3 | 3 | 1 | 1 | 1 | 3 | 1 | 3 | 4 | 4 | 1 | 1 | 1 | 4 | 1 | 1 | 1 | 1 | 1 | 1 | 1 | 1 | 3 | 3 | 4 |
| yeps_YPTB2489 | 1 | 3 | 1 | 1 | 1 | 1 | 1 | 1 | 3 | 1 | 1 | 1 | 1 | 1 | 1 | 1 | 1 | 4 | 1 | 1 | 1 | 1 | 1 | 1 | 1 | 1 | 1 | 1 | 1 | 1 | 1 | 1 | 1 | 1 | 1 |
| yeps_YPTB2503 | 5 | 4 | 5 | 1 | 1 | 1 | 5 | 5 | 4 | 1 | 1 | 1 | 1 | 1 | 1 | 1 | 1 | 1 | 5 | 5 | 1 | 1 | 1 | 5 | 1 | 1 | 1 | 1 | 1 | 1 | 1 | 1 | 1 | 1 | 5 |
| yeps_YPTB2504 | 4 | 5 | 4 | 8 | 1 | 1 | 4 | 7 | 5 | 8 | 8 | 3 | 1 | 1 | 1 | 8 | 1 | 3 | 4 | 4 | 1 | 1 | 1 | 4 | 1 | 1 | 1 | 1 | 1 | 1 | 1 | 1 | 8 | 8 | 4 |
| yeps_YPTB2505 | 3 | 4 | 3 | 6 | 1 | 1 | 3 | 5 | 4 | 6 | 6 | 1 | 1 | 1 | 1 | 6 | 1 | 3 | 3 | 3 | 1 | 1 | 1 | 3 | 1 | 1 | 1 | 1 | 1 | 1 | 1 | 1 | 6 | 6 | 3 |
| yeps_YPTB2506 | 4 | 6 | 4 | 8 | 1 | 1 | 4 | 7 | 6 | 8 | 8 | 1 | 1 | 1 | 1 | 8 | 1 | 5 | 4 | 4 | 1 | 1 | 1 | 4 | 1 | 1 | 1 | 1 | 1 | 1 | 1 | 1 | 8 | 8 | 4 |
| yeps_YPTB2507 | 3 | 6 | 3 | 3 | 1 | 1 | 3 | 7 | 6 | 3 | 3 | 1 | 1 | 1 | 1 | 3 | 1 | 4 | 3 | 3 | 1 | 1 | 1 | 3 | 1 | 1 | 1 | 1 | 1 | 1 | 1 | 1 | 3 | 3 | 3 |
| yeps_YPTB2508 | 4 | 7 | 4 | 11 | 1 | 1 | 4 | 8 | 7 | 10 | 11 | 1 | 1 | 1 | 1 | 10 | 1 | 5 | 4 | 4 | 1 | 1 | 1 | 4 | 1 | 1 | 1 | 1 | 1 | 1 | 1 | 1 | 11 | 11 | 4 |
| yeps_YPTB2513 | 3 | 6 | 3 | 9 | 1 | 1 | 3 | 7 | 6 | 8 | 9 | 1 | 1 | 1 | 1 | 8 | 1 | 4 | 3 | 3 | 1 | 1 | 1 | 3 | 1 | 1 | 1 | 1 | 1 | 1 | 1 | 1 | 9 | 9 | 3 |
| yeps_YPTB2514 | 1 | 5 | 1 | 10 | 1 | 1 | 1 | 6 | 5 | 8 | 10 | 1 | 1 | 1 | 1 | 8 | 1 | 3 | 1 | 1 | 1 | 1 | 1 | 1 | 1 | 1 | 1 | 1 | 1 | 1 | 1 | 1 | 10 | 10 | 1 |
| yeps_YPTB2516 | 1 | 3 | 1 | 3 | 1 | 1 | 1 | 1 | 3 | 3 | 3 | 1 | 1 | 1 | 1 | 3 | 1 | 3 | 1 | 1 | 1 | 1 | 1 | 1 | 1 | 1 | 1 | 1 | 1 | 1 | 1 | 1 | 3 | 3 | 1 |
| yeps_YPTB2517 | 3 | 3 | 3 | 4 | 1 | 1 | 3 | 5 | 3 | 4 | 4 | 1 | 1 | 1 | 1 | 4 | 1 | 4 | 3 | 3 | 1 | 1 | 1 | 3 | 1 | 1 | 1 | 1 | 1 | 1 | 1 | 1 | 4 | 4 | 3 |
| yeps_YPTB2518 | 3 | 4 | 3 | 4 | 1 | 1 | 3 | 4 | 4 | 4 | 4 | 1 | 1 | 1 | 1 | 4 | 1 | 4 | 3 | 3 | 1 | 1 | 1 | 3 | 1 | 1 | 1 | 1 | 1 | 1 | 1 | 1 | 4 | 4 | 3 |
| yeps_YPTB2519 | 3 | 1 | 3 | 2 | 1 | 1 | 3 | 6 | 1 | 2 | 2 | 1 | 1 | 1 | 1 | 2 | 1 | 2 | 3 | 3 | 1 | 1 | 1 | 3 | 1 | 1 | 1 | 1 | 1 | 1 | 1 | 1 | 2 | 2 | 3 |
| yeps_YPTB2520 | 1 | 5 | 1 | 6 | 1 | 1 | 1 | 1 | 5 | 6 | 6 | 1 | 1 | 1 | 1 | 6 | 1 | 3 | 1 | 1 | 1 | 1 | 1 | 1 | 1 | 1 | 1 | 1 | 1 | 1 | 1 | 1 | 6 | 6 | 1 |
| yeps_YPTB2535 | 1 | 6 | 1 | 8 | 1 | 1 | 1 | 7 | 6 | 8 | 8 | 1 | 1 | 34 | 1 | 8 | 1 | 2 | 1 | 1 | 1 | 1 | 1 | 1 | 1 | 1 | 1 | 1 | 1 | 1 | 1 | 1 | 8 | 8 | 1 |
| yeps_YPTB2536 | 1 | 7 | 1 | 10 | 1 | 1 | 1 | 8 | 7 | 10 | 10 | 1 | 1 | 1 | 1 | 10 | 1 | 3 | 1 | 1 | 1 | 1 | 1 | 1 | 1 | 1 | 1 | 1 | 1 | 1 | 1 | 1 | 10 | 10 | 1 |
| yeps_YPTB2537 | 50 | 8 | 3 | 5 | 1 | 3 | 3 | 9 | 8 | 5 | 5 | 1 | 1 | 1 | 3 | 5 | 1 | 4 | 3 | 3 | 3 | 3 | 3 | 3 | 1 | 1 | 3 | 3 | 3 | 3 | 3 | 1 | 5 | 5 | 3 |
| yeps_YPTB2541 | 3 | 6 | 3 | 8 | 1 | 3 | 3 | 6 | 6 | 8 | 8 | 3 | 3 | 3 | 3 | 8 | 3 | 1 | 3 | 3 | 3 | 3 | 3 | 3 | 3 | 3 | 3 | 3 | 3 | 3 | 3 | 3 | 8 | 8 | 3 |
| yeps_YPTB2542 | 1 | 1 | 1 | 3 | 1 | 1 | 1 | 5 | 1 | 3 | 3 | 1 | 1 | 1 | 1 | 3 | 1 | 3 | 1 | 1 | 1 | 1 | 1 | 1 | 1 | 1 | 1 | 1 | 1 | 1 | 1 | 1 | 3 | 3 | 1 |
| yeps_YPTB2543 | 1 | 6 | 1 | 8 | 1 | 1 | 1 | 6 | 6 | 8 | 8 | 3 | 3 | 3 | 1 | 8 | 3 | 4 | 1 | 1 | 1 | 1 | 1 | 1 | 3 | 3 | 1 | 1 | 1 | 1 | 1 | 3 | 8 | 8 | 1 |
| yeps_YPTB2544 | 1 | 1 | 1 | 8 | 1 | 1 | 1 | 1 | 1 | 8 | 8 | 3 | 3 | 3 | 1 | 8 | 3 | 4 | 1 | 37 | 1 | 1 | 1 | 1 | 3 | 3 | 1 | 1 | 1 | 1 | 1 | 3 | 8 | 8 | 1 |
| yeps_YPTB2546 | 1 | 1 | 1 | 1 | 1 | 1 | 1 | 1 | 1 | 1 | 1 | 1 | 1 | 1 | 1 | 1 | 1 | 2 | 1 | 1 | 1 | 1 | 1 | 1 | 1 | 1 | 1 | 1 | 1 | 1 | 1 | 1 | 1 | 1 | 1 |
| yeps_YPTB2547 | 1 | 6 | 1 | 8 | 1 | 1 | 1 | 1 | 6 | 8 | 8 | 1 | 1 | 1 | 1 | 8 | 1 | 9 | 1 | 1 | 1 | 1 | 1 | 1 | 1 | 1 | 1 | 1 | 1 | 1 | 1 | 1 | 8 | 8 | 1 |
| yeps_YPTB2548 | 1 | 1 | 1 | 7 | 1 | 1 | 1 | 1 | 1 | 7 | 7 | 1 | 1 | 1 | 1 | 7 | 1 | 3 | 1 | 1 | 1 | 1 | 1 | 1 | 1 | 1 | 1 | 1 | 1 | 1 | 1 | 1 | 7 | 7 | 1 |
| yeps_YPTB2550 | 1 | 6 | 1 | 8 | 1 | 1 | 1 | 7 | 6 | 8 | 8 | 1 | 1 | 1 | 1 | 8 | 1 | 3 | 1 | 1 | 1 | 1 | 1 | 1 | 1 | 1 | 1 | 1 | 1 | 1 | 1 | 1 | 8 | 8 | 1 |
| yeps_YPTB2556 | 4 | 9 | 4 | 12 | 1 | 4 | 4 | 10 | 9 | 12 | 12 | 3 | 3 | 3 | 4 | 12 | 3 | 5 | 4 | 4 | 4 | 4 | 4 | 4 | 3 | 3 | 4 | 4 | 4 | 4 | 4 | 3 | 12 | 12 | 4 |
| yeps_YPTB2559 | 5 | 9 | 5 | 5 | 1 | 5 | 5 | 10 | 9 | 5 | 5 | 3 | 12 | 7 | 5 | 5 | 7 | 5 | 5 | 5 | 5 | 5 | 5 | 5 | 4 | 4 | 5 | 5 | 5 | 5 | 5 | 4 | 5 | 5 | 5 |
| yeps_YPTB2560 | 5 | 8 | 5 | 12 | 1 | 5 | 5 | 10 | 9 | 12 | 12 | 3 | 4 | 4 | 5 | 12 | 4 | 1 | 5 | 5 | 5 | 5 | 5 | 5 | 4 | 4 | 5 | 5 | 5 | 5 | 5 | 89 | 12 | 12 | 5 |
| yeps_YPTB2561 | 5 | 3 | 5 | 3 | 1 | 5 | 5 | 8 | 3 | 3 | 3 | 3 | 4 | 4 | 5 | 3 | 4 | 1 | 5 | 5 | 5 | 5 | 5 | 5 | 4 | 4 | 5 | 5 | 5 | 5 | 5 | 4 | 3 | 3 | 5 |
| yeps_YPTB2572 | 1 | 4 | 1 | 7 | 1 | 1 | 1 | 5 | 4 | 7 | 7 | 1 | 1 | 1 | 1 | 7 | 1 | 1 | 1 | 1 | 1 | 1 | 1 | 1 | 1 | 1 | 1 | 1 | 1 | 1 | 1 | 1 | 7 | 7 | 1 |
| yeps_YPTB2575 | 4 | 7 | 4 | 10 | 1 | 1 | 4 | 8 | 7 | 10 | 10 | 1 | 1 | 1 | 1 | 10 | 1 | 4 | 30 | 4 | 1 | 1 | 1 | 4 | 1 | 1 | 1 | 1 | 1 | 1 | 1 | 1 | 10 | 10 | 4 |
| yeps_YPTB2576 | 4 | 7 | 4 | 11 | 3 | 3 | 4 | 9 | 7 | 11 | 11 | 30 | 3 | 3 | 3 | 11 | 3 | 4 | 4 | 4 | 3 | 3 | 3 | 4 | 3 | 3 | 3 | 3 | 3 | 3 | 3 | 3 | 11 | 11 | 4 |
| yeps_YPTB2579 | 3 | 2 | 3 | 5 | 1 | 1 | 3 | 2 | 2 | 5 | 5 | 1 | 1 | 1 | 1 | 5 | 1 | 3 | 3 | 3 | 1 | 1 | 1 | 3 | 1 | 1 | 1 | 1 | 1 | 1 | 1 | 1 | 5 | 5 | 3 |
| yeps_YPTB2582 | 4 | 7 | 4 | 10 | 1 | 79 | 4 | 18 | 7 | 121 | 10 | 1 | 1 | 1 | 79 | 10 | 1 | 12 | 4 | 11 | 79 | 79 | 79 | 11 | 1 | 1 | 79 | 79 | 79 | 79 | 79 | 1 | 10 | 10 | 4 |
| yeps_YPTB2583 | 13 | 7 | 3 | 12 | 1 | 52 | 3 | 8 | 7 | 10 | 12 | 1 | 5 | 5 | 52 | 10 | 5 | 3 | 3 | 3 | 52 | 52 | 52 | 3 | 5 | 5 | 52 | 52 | 52 | 52 | 52 | 5 | 12 | 12 | 3 |
| yeps_YPTB2584 | 1 | 4 | 1 | 6 | 1 | 1 | 1 | 5 | 4 | 6 | 6 | 1 | 1 | 1 | 1 | 6 | 1 | 1 | 1 | 1 | 1 | 1 | 1 | 1 | 1 | 1 | 1 | 1 | 1 | 1 | 1 | 1 | 6 | 6 | 1 |
| yeps_YPTB2593 | 1 | 1 | 1 | 6 | 1 | 1 | 1 | 5 | 1 | 6 | 6 | 3 | 1 | 1 | 1 | 6 | 1 | 1 | 1 | 1 | 1 | 1 | 1 | 1 | 1 | 1 | 1 | 1 | 1 | 1 | 1 | 1 | 6 | 6 | 1 |
| yeps_YPTB2596 | 1 | 2 | 1 | 1 | 1 | 1 | 1 | 1 | 2 | 1 | 1 | 1 | 1 | 1 | 1 | 1 | 1 | 1 | 1 | 1 | 1 | 1 | 1 | 1 | 1 | 1 | 1 | 1 | 1 | 1 | 1 | 1 | 1 | 1 | 1 |
| yeps_YPTB2603 | 3 | 3 | 3 | 6 | 1 | 1 | 3 | 5 | 3 | 6 | 6 | 1 | 3 | 3 | 1 | 6 | 3 | 2 | 3 | 3 | 1 | 1 | 1 | 3 | 3 | 3 | 1 | 1 | 1 | 1 | 1 | 3 | 6 | 6 | 3 |
| yeps_YPTB2605 | 1 | 1 | 1 | 1 | 1 | 1 | 1 | 2 | 1 | 1 | 1 | 1 | 1 | 1 | 1 | 1 | 1 | 1 | 1 | 1 | 1 | 1 | 1 | 1 | 1 | 1 | 1 | 1 | 1 | 1 | 1 | 1 | 1 | 1 | 1 |
| yeps_YPTB2606 | 4 | 4 | 4 | 7 | 1 | 1 | 4 | 4 | 4 | 7 | 7 | 3 | 4 | 4 | 1 | 7 | 4 | 4 | 4 | 4 | 1 | 1 | 1 | 4 | 4 | 4 | 1 | 1 | 1 | 1 | 1 | 4 | 7 | 7 | 4 |
| yeps_YPTB2611 | 1 | 2 | 1 | 1 | 1 | 1 | 1 | 1 | 2 | 1 | 1 | 1 | 1 | 1 | 1 | 1 | 1 | 1 | 1 | 1 | 1 | 1 | 1 | 1 | 1 | 1 | 1 | 1 | 1 | 1 | 1 | 1 | 1 | 1 | 1 |
| yeps_YPTB2616 | 2 | 1 | 2 | 2 | 1 | 1 | 2 | 2 | 1 | 2 | 2 | 1 | 1 | 1 | 1 | 2 | 1 | 1 | 2 | 2 | 1 | 1 | 1 | 2 | 1 | 1 | 1 | 1 | 1 | 1 | 1 | 1 | 2 | 2 | 2 |
| yeps_YPTB2621 | 1 | 1 | 1 | 1 | 1 | 1 | 1 | 1 | 1 | 1 | 1 | 1 | 1 | 1 | 1 | 1 | 1 | 1 | 1 | 1 | 1 | 1 | 1 | 1 | 1 | 1 | 1 | 1 | 1 | 1 | 1 | 1 | 1 | 1 | 1 |
| yeps_YPTB2628 | 1 | 1 | 1 | 1 | 1 | 1 | 1 | 1 | 1 | 1 | 1 | 1 | 1 | 1 | 1 | 1 | 1 | 1 | 1 | 1 | 1 | 1 | 1 | 1 | 1 | 1 | 1 | 1 | 1 | 1 | 1 | 1 | 1 | 1 | 1 |
| yeps_YPTB2631 | 1 | 7 | 1 | 8 | 1 | 1 | 1 | 8 | 7 | 8 | 8 | 1 | 5 | 5 | 1 | 8 | 5 | 6 | 1 | 1 | 1 | 1 | 1 | 1 | 5 | 5 | 1 | 1 | 1 | 1 | 1 | 5 | 8 | 8 | 1 |
| yeps_YPTB2633 | 2 | 4 | 2 | 6 | 1 | 1 | 2 | 2 | 4 | 6 | 6 | 3 | 1 | 1 | 1 | 6 | 1 | 2 | 2 | 2 | 1 | 1 | 1 | 2 | 1 | 1 | 1 | 1 | 1 | 1 | 1 | 1 | 6 | 6 | 2 |
| yeps_YPTB2634 | 1 | 4 | 1 | 1 | 1 | 1 | 1 | 1 | 4 | 1 | 1 | 1 | 1 | 1 | 1 | 1 | 1 | 1 | 1 | 1 | 1 | 1 | 1 | 1 | 1 | 1 | 1 | 1 | 1 | 1 | 1 | 1 | 1 | 1 | 1 |
| yeps_YPTB2635 | 1 | 1 | 1 | 3 | 1 | 1 | 1 | 1 | 1 | 3 | 3 | 1 | 1 | 1 | 1 | 3 | 1 | 1 | 1 | 1 | 1 | 1 | 1 | 1 | 1 | 1 | 1 | 1 | 1 | 1 | 1 | 1 | 3 | 3 | 1 |
| yeps_YPTB2638 | 1 | 1 | 1 | 1 | 1 | 1 | 1 | 1 | 1 | 1 | 1 | 1 | 1 | 1 | 1 | 1 | 1 | 1 | 1 | 1 | 1 | 1 | 1 | 1 | 1 | 1 | 1 | 1 | 1 | 1 | 1 | 1 | 1 | 1 | 1 |
| yeps_YPTB2641 | 9 | 8 | 4 | 5 | 1 | 1 | 4 | 3 | 5 | 5 | 5 | 3 | 1 | 1 | 1 | 5 | 1 | 5 | 4 | 4 | 1 | 1 | 1 | 4 | 1 | 1 | 1 | 1 | 1 | 1 | 1 | 1 | 5 | 5 | 4 |
| yeps_YPTB2642 | 1 | 1 | 1 | 1 | 1 | 1 | 1 | 1 | 1 | 1 | 1 | 3 | 1 | 1 | 1 | 1 | 1 | 1 | 1 | 1 | 1 | 1 | 1 | 1 | 1 | 1 | 1 | 1 | 1 | 1 | 1 | 1 | 1 | 1 | 1 |
| yeps_YPTB2643 | 1 | 1 | 1 | 1 | 1 | 1 | 1 | 3 | 1 | 1 | 1 | 1 | 1 | 1 | 1 | 1 | 1 | 1 | 1 | 1 | 1 | 1 | 1 | 1 | 1 | 1 | 1 | 1 | 1 | 1 | 1 | 1 | 1 | 1 | 1 |
| yeps_YPTB2644 | 1 | 6 | 1 | 1 | 1 | 1 | 1 | 8 | 6 | 1 | 1 | 1 | 1 | 1 | 1 | 1 | 1 | 4 | 1 | 1 | 1 | 1 | 1 | 1 | 1 | 1 | 1 | 1 | 1 | 1 | 1 | 1 | 1 | 1 | 1 |
| yeps_YPTB2645 | 1 | 1 | 1 | 7 | 1 | 1 | 1 | 5 | 1 | 1 | 7 | 2 | 1 | 1 | 1 | 1 | 1 | 1 | 1 | 1 | 1 | 1 | 1 | 1 | 1 | 1 | 1 | 1 | 1 | 1 | 1 | 1 | 7 | 7 | 1 |
| yeps_YPTB2647 | 4 | 6 | 4 | 4 | 1 | 1 | 4 | 3 | 6 | 4 | 4 | 3 | 1 | 1 | 1 | 4 | 1 | 5 | 4 | 4 | 1 | 1 | 1 | 4 | 1 | 1 | 1 | 1 | 1 | 1 | 1 | 1 | 4 | 4 | 4 |
| yeps_YPTB2648 | 3 | 3 | 3 | 3 | 1 | 1 | 3 | 4 | 3 | 3 | 3 | 3 | 1 | 1 | 1 | 3 | 1 | 3 | 3 | 3 | 1 | 1 | 1 | 3 | 1 | 1 | 1 | 1 | 1 | 1 | 1 | 1 | 3 | 3 | 3 |
| yeps_YPTB2649 | 3 | 7 | 3 | 3 | 1 | 1 | 3 | 8 | 7 | 3 | 3 | 3 | 1 | 1 | 1 | 3 | 1 | 3 | 3 | 3 | 1 | 1 | 1 | 3 | 1 | 1 | 1 | 1 | 1 | 1 | 1 | 1 | 3 | 3 | 3 |
| yeps_YPTB2650 | 3 | 5 | 3 | 6 | 1 | 1 | 3 | 4 | 5 | 6 | 6 | 1 | 1 | 1 | 1 | 6 | 1 | 4 | 3 | 3 | 1 | 1 | 1 | 3 | 1 | 1 | 1 | 1 | 1 | 1 | 1 | 1 | 6 | 6 | 3 |
| yeps_YPTB2651 | 4 | 6 | 4 | 1 | 1 | 1 | 4 | 7 | 6 | 1 | 1 | 3 | 1 | 1 | 1 | 1 | 1 | 1 | 4 | 4 | 1 | 1 | 1 | 4 | 1 | 1 | 1 | 1 | 1 | 1 | 1 | 1 | 1 | 1 | 4 |
| yeps_YPTB2654 | 2 | 2 | 2 | 2 | 1 | 1 | 2 | 2 | 2 | 2 | 2 | 1 | 1 | 1 | 1 | 2 | 1 | 2 | 2 | 2 | 1 | 1 | 1 | 2 | 1 | 1 | 1 | 1 | 1 | 1 | 1 | 1 | 2 | 2 | 2 |
| yeps_YPTB2655 | 2 | 2 | 2 | 4 | 1 | 1 | 2 | 3 | 2 | 4 | 4 | 1 | 1 | 1 | 1 | 4 | 1 | 2 | 2 | 2 | 1 | 1 | 1 | 2 | 1 | 1 | 1 | 1 | 1 | 1 | 1 | 1 | 4 | 4 | 2 |
| yeps_YPTB2661 | 1 | 7 | 1 | 8 | 1 | 1 | 1 | 4 | 7 | 8 | 8 | 3 | 1 | 1 | 1 | 8 | 1 | 4 | 1 | 1 | 1 | 1 | 1 | 1 | 1 | 1 | 1 | 1 | 1 | 1 | 1 | 1 | 8 | 8 | 1 |
| yeps_YPTB2662 | 1 | 6 | 1 | 7 | 1 | 1 | 1 | 3 | 6 | 7 | 7 | 3 | 1 | 1 | 1 | 7 | 1 | 4 | 1 | 1 | 1 | 1 | 1 | 1 | 1 | 1 | 1 | 1 | 1 | 1 | 1 | 1 | 7 | 7 | 1 |
| yeps_YPTB2673 | 3 | 1 | 3 | 10 | 1 | 1 | 3 | 2 | 1 | 9 | 10 | 3 | 1 | 1 | 1 | 9 | 1 | 7 | 4 | 4 | 1 | 1 | 1 | 4 | 1 | 1 | 1 | 1 | 1 | 1 | 1 | 1 | 10 | 10 | 3 |
| yeps_YPTB2674 | 2 | 5 | 2 | 3 | 1 | 1 | 2 | 1 | 5 | 7 | 3 | 2 | 1 | 1 | 1 | 7 | 1 | 3 | 2 | 2 | 1 | 1 | 1 | 2 | 1 | 1 | 1 | 1 | 1 | 1 | 1 | 1 | 3 | 3 | 2 |
| yeps_YPTB2675 | 1 | 1 | 1 | 5 | 1 | 1 | 1 | 1 | 1 | 5 | 5 | 1 | 1 | 1 | 1 | 5 | 1 | 2 | 1 | 1 | 1 | 1 | 1 | 1 | 1 | 1 | 1 | 1 | 1 | 1 | 1 | 1 | 5 | 5 | 1 |
| yeps_YPTB2676 | 4 | 8 | 9 | 12 | 1 | 1 | 9 | 1 | 8 | 11 | 12 | 3 | 1 | 1 | 1 | 11 | 1 | 5 | 4 | 4 | 1 | 1 | 1 | 4 | 1 | 1 | 1 | 1 | 1 | 1 | 1 | 1 | 12 | 12 | 4 |
| yeps_YPTB2678 | 4 | 3 | 4 | 10 | 1 | 1 | 4 | 9 | 3 | 10 | 10 | 3 | 1 | 1 | 1 | 10 | 1 | 5 | 4 | 4 | 1 | 1 | 1 | 4 | 1 | 1 | 1 | 1 | 1 | 1 | 1 | 1 | 10 | 10 | 11 |
| yeps_YPTB2679 | 1 | 1 | 1 | 2 | 1 | 1 | 1 | 1 | 1 | 1 | 2 | 2 | 1 | 1 | 1 | 1 | 1 | 2 | 1 | 1 | 1 | 1 | 1 | 1 | 1 | 1 | 1 | 1 | 1 | 1 | 1 | 1 | 2 | 2 | 1 |
| yeps_YPTB2689 | 2 | 2 | 2 | 2 | 1 | 1 | 2 | 2 | 2 | 2 | 2 | 1 | 1 | 1 | 1 | 2 | 1 | 1 | 2 | 2 | 1 | 1 | 1 | 2 | 1 | 1 | 1 | 1 | 1 | 1 | 1 | 1 | 2 | 2 | 2 |
| yeps_YPTB2693 | 3 | 5 | 3 | 1 | 1 | 1 | 3 | 5 | 5 | 1 | 1 | 3 | 1 | 1 | 1 | 1 | 1 | 4 | 3 | 3 | 1 | 1 | 1 | 3 | 1 | 1 | 1 | 1 | 1 | 1 | 1 | 1 | 1 | 1 | 3 |
| yeps_YPTB2697 | 1 | 1 | 1 | 1 | 1 | 1 | 1 | 1 | 1 | 1 | 1 | 1 | 1 | 1 | 1 | 1 | 1 | 1 | 1 | 1 | 1 | 1 | 1 | 1 | 1 | 1 | 1 | 1 | 1 | 1 | 1 | 1 | 1 | 1 | 1 |
| yeps_YPTB2698 | 7 | 4 | 3 | 5 | 1 | 1 | 3 | 6 | 4 | 5 | 5 | 1 | 1 | 1 | 1 | 5 | 1 | 4 | 3 | 3 | 1 | 1 | 1 | 3 | 1 | 1 | 1 | 1 | 1 | 1 | 1 | 1 | 5 | 5 | 3 |
| yeps_YPTB2700 | 3 | 7 | 3 | 7 | 1 | 7 | 3 | 3 | 7 | 7 | 7 | 3 | 3 | 5 | 7 | 7 | 5 | 3 | 3 | 3 | 7 | 7 | 7 | 3 | 3 | 3 | 7 | 7 | 7 | 7 | 7 | 3 | 7 | 7 | 3 |
| yeps_YPTB2701 | 4 | 5 | 4 | 5 | 1 | 1 | 4 | 3 | 5 | 5 | 5 | 3 | 4 | 4 | 1 | 5 | 4 | 3 | 4 | 4 | 1 | 1 | 1 | 4 | 4 | 4 | 1 | 1 | 1 | 1 | 1 | 4 | 5 | 5 | 4 |
| yeps_YPTB2703 | 1 | 2 | 1 | 4 | 1 | 1 | 1 | 3 | 2 | 4 | 4 | 1 | 1 | 1 | 1 | 4 | 1 | 1 | 1 | 1 | 1 | 1 | 1 | 1 | 1 | 1 | 1 | 1 | 1 | 1 | 1 | 1 | 4 | 4 | 1 |
| yeps_YPTB2704 | 2 | 2 | 2 | 2 | 1 | 1 | 2 | 2 | 2 | 2 | 2 | 2 | 2 | 2 | 1 | 2 | 2 | 1 | 2 | 2 | 1 | 1 | 1 | 2 | 2 | 2 | 1 | 1 | 1 | 1 | 1 | 2 | 2 | 2 | 2 |
| yeps_YPTB2711 | 1 | 7 | 1 | 11 | 1 | 1 | 1 | 8 | 7 | 11 | 11 | 3 | 1 | 1 | 1 | 10 | 1 | 4 | 1 | 1 | 1 | 1 | 1 | 1 | 1 | 1 | 1 | 1 | 1 | 1 | 1 | 1 | 11 | 11 | 1 |
| yeps_YPTB2713 | 1 | 3 | 1 | 4 | 1 | 1 | 1 | 1 | 3 | 4 | 4 | 1 | 1 | 1 | 1 | 4 | 1 | 1 | 1 | 1 | 1 | 1 | 1 | 1 | 1 | 1 | 1 | 1 | 1 | 1 | 1 | 1 | 4 | 4 | 1 |
| yeps_YPTB2717 | 1 | 1 | 1 | 1 | 1 | 1 | 1 | 1 | 1 | 1 | 1 | 1 | 1 | 1 | 1 | 1 | 1 | 1 | 1 | 1 | 1 | 1 | 1 | 1 | 1 | 1 | 1 | 1 | 1 | 1 | 1 | 1 | 1 | 1 | 1 |
| yeps_YPTB2718 | 1 | 6 | 1 | 7 | 1 | 1 | 1 | 8 | 7 | 7 | 7 | 2 | 1 | 1 | 1 | 7 | 1 | 3 | 1 | 1 | 1 | 1 | 1 | 1 | 1 | 1 | 1 | 1 | 1 | 1 | 1 | 1 | 7 | 7 | 1 |
| yeps_YPTB2719 | 1 | 1 | 1 | 1 | 1 | 1 | 1 | 1 | 1 | 1 | 1 | 1 | 1 | 1 | 1 | 1 | 1 | 4 | 1 | 1 | 1 | 1 | 1 | 1 | 1 | 1 | 1 | 1 | 1 | 1 | 1 | 1 | 1 | 1 | 1 |
| yeps_YPTB2721 | 1 | 7 | 1 | 10 | 1 | 1 | 1 | 8 | 7 | 10 | 10 | 1 | 5 | 5 | 1 | 10 | 5 | 3 | 1 | 1 | 1 | 1 | 1 | 1 | 5 | 5 | 1 | 1 | 1 | 1 | 1 | 5 | 10 | 10 | 1 |
| yeps_YPTB2737 | 1 | 5 | 1 | 6 | 1 | 1 | 1 | 1 | 5 | 6 | 6 | 3 | 1 | 1 | 1 | 6 | 1 | 4 | 1 | 1 | 1 | 1 | 1 | 1 | 1 | 1 | 1 | 1 | 1 | 1 | 1 | 1 | 6 | 6 | 1 |
| yeps_YPTB2739 | 1 | 46 | 1 | 8 | 1 | 1 | 1 | 1 | 6 | 7 | 8 | 1 | 1 | 1 | 1 | 7 | 1 | 3 | 1 | 1 | 1 | 1 | 1 | 1 | 1 | 1 | 1 | 1 | 1 | 1 | 1 | 1 | 8 | 8 | 1 |
| yeps_YPTB2741 | 1 | 5 | 1 | 8 | 1 | 1 | 1 | 7 | 6 | 8 | 8 | 3 | 1 | 1 | 1 | 8 | 1 | 4 | 1 | 1 | 1 | 1 | 1 | 1 | 1 | 1 | 1 | 1 | 1 | 1 | 1 | 1 | 8 | 8 | 1 |
| yeps_YPTB2742 | 1 | 3 | 1 | 1 | 1 | 1 | 1 | 5 | 3 | 1 | 1 | 3 | 1 | 1 | 1 | 1 | 1 | 1 | 1 | 1 | 1 | 1 | 1 | 1 | 1 | 1 | 1 | 1 | 1 | 1 | 1 | 1 | 1 | 1 | 1 |
| yeps_YPTB2743 | 1 | 7 | 1 | 1 | 1 | 1 | 1 | 9 | 7 | 1 | 1 | 3 | 1 | 1 | 1 | 1 | 1 | 4 | 1 | 1 | 1 | 1 | 1 | 1 | 1 | 1 | 1 | 1 | 1 | 1 | 1 | 1 | 1 | 1 | 1 |
| yeps_YPTB2744 | 1 | 5 | 1 | 1 | 1 | 1 | 1 | 6 | 1 | 1 | 1 | 2 | 1 | 1 | 1 | 1 | 1 | 1 | 1 | 1 | 1 | 1 | 1 | 1 | 1 | 1 | 1 | 1 | 1 | 1 | 1 | 1 | 1 | 1 | 1 |
| yeps_YPTB2755 | 1 | 2 | 1 | 1 | 1 | 1 | 1 | 1 | 2 | 1 | 1 | 1 | 1 | 1 | 1 | 1 | 1 | 1 | 1 | 1 | 1 | 1 | 1 | 1 | 1 | 1 | 1 | 1 | 1 | 1 | 1 | 1 | 1 | 1 | 1 |
| yeps_YPTB2756 | 1 | 8 | 1 | 11 | 1 | 1 | 1 | 12 | 8 | 11 | 11 | 3 | 7 | 7 | 1 | 11 | 7 | 5 | 1 | 1 | 1 | 1 | 1 | 1 | 7 | 7 | 1 | 1 | 1 | 1 | 1 | 7 | 11 | 11 | 1 |
| yeps_YPTB2757 | 1 | 8 | 1 | 9 | 1 | 1 | 1 | 7 | 8 | 9 | 9 | 2 | 2 | 2 | 1 | 9 | 6 | 4 | 1 | 1 | 1 | 1 | 1 | 1 | 2 | 2 | 1 | 1 | 1 | 1 | 1 | 2 | 9 | 9 | 1 |
| yeps_YPTB2758 | 3 | 1 | 3 | 3 | 1 | 1 | 3 | 3 | 1 | 3 | 3 | 3 | 5 | 5 | 1 | 3 | 5 | 1 | 3 | 3 | 1 | 1 | 1 | 3 | 5 | 5 | 1 | 1 | 1 | 1 | 1 | 5 | 3 | 3 | 3 |
| yeps_YPTB2759 | 4 | 1 | 4 | 1 | 1 | 1 | 4 | 1 | 1 | 1 | 1 | 1 | 1 | 1 | 1 | 1 | 1 | 1 | 4 | 4 | 1 | 1 | 1 | 4 | 1 | 1 | 1 | 1 | 1 | 1 | 1 | 1 | 1 | 1 | 4 |
| yeps_YPTB2764 | 1 | 1 | 1 | 1 | 1 | 1 | 1 | 1 | 1 | 1 | 1 | 2 | 1 | 1 | 1 | 1 | 1 | 1 | 1 | 1 | 1 | 1 | 1 | 1 | 1 | 1 | 1 | 1 | 1 | 1 | 1 | 1 | 1 | 1 | 1 |
| yeps_YPTB2765 | 5 | 9 | 5 | 13 | 1 | 1 | 5 | 11 | 9 | 13 | 13 | 4 | 6 | 6 | 1 | 13 | 6 | 6 | 5 | 5 | 1 | 1 | 1 | 5 | 6 | 6 | 1 | 1 | 1 | 1 | 1 | 6 | 13 | 13 | 5 |
| yeps_YPTB2766 | 1 | 1 | 1 | 1 | 1 | 1 | 1 | 1 | 1 | 1 | 1 | 1 | 1 | 1 | 1 | 1 | 1 | 3 | 1 | 1 | 1 | 1 | 1 | 1 | 1 | 1 | 1 | 1 | 1 | 1 | 1 | 1 | 1 | 1 | 1 |
| yeps_YPTB2773 | 1 | 1 | 1 | 1 | 1 | 1 | 1 | 1 | 1 | 1 | 1 | 1 | 1 | 1 | 1 | 1 | 1 | 1 | 1 | 1 | 1 | 1 | 1 | 1 | 1 | 1 | 1 | 1 | 1 | 1 | 1 | 1 | 1 | 1 | 1 |
| yeps_YPTB2774 | 1 | 3 | 1 | 5 | 1 | 1 | 1 | 1 | 3 | 5 | 5 | 1 | 1 | 1 | 1 | 5 | 1 | 3 | 1 | 1 | 1 | 1 | 1 | 1 | 1 | 1 | 1 | 1 | 1 | 1 | 1 | 1 | 5 | 5 | 1 |
| yeps_YPTB2776 | 1 | 1 | 1 | 1 | 1 | 1 | 1 | 1 | 1 | 1 | 1 | 1 | 1 | 1 | 1 | 1 | 1 | 1 | 1 | 1 | 1 | 1 | 1 | 1 | 1 | 1 | 1 | 1 | 1 | 1 | 1 | 1 | 1 | 1 | 1 |
| yeps_YPTB2777 | 1 | 1 | 1 | 1 | 1 | 1 | 1 | 2 | 1 | 1 | 1 | 1 | 1 | 1 | 1 | 1 | 1 | 1 | 1 | 1 | 1 | 1 | 1 | 1 | 1 | 1 | 1 | 1 | 1 | 1 | 1 | 1 | 1 | 1 | 1 |
| yeps_YPTB2782 | 3 | 8 | 3 | 10 | 1 | 1 | 3 | 9 | 8 | 10 | 10 | 3 | 7 | 7 | 1 | 10 | 7 | 7 | 3 | 3 | 1 | 1 | 1 | 3 | 7 | 7 | 1 | 1 | 1 | 1 | 1 | 7 | 10 | 10 | 3 |
| yeps_YPTB2783 | 1 | 3 | 1 | 2 | 1 | 1 | 1 | 4 | 3 | 2 | 2 | 1 | 3 | 3 | 1 | 2 | 3 | 2 | 1 | 1 | 1 | 1 | 1 | 1 | 3 | 3 | 1 | 1 | 1 | 1 | 1 | 3 | 2 | 2 | 1 |
| yeps_YPTB2784 | 1 | 1 | 1 | 2 | 1 | 1 | 1 | 1 | 1 | 2 | 2 | 1 | 1 | 1 | 1 | 2 | 1 | 1 | 1 | 1 | 1 | 1 | 1 | 1 | 1 | 1 | 1 | 1 | 1 | 1 | 1 | 1 | 2 | 2 | 1 |
| yeps_YPTB2786 | 1 | 2 | 1 | | 1 | 1 | 1 | 1 | 2 | 1 | 1 | 1 | 1 | 1 | 1 | 1 | 1 | 4 | 1 | | 1 | 1 | 1 | 1 | 1 | 1 | 1 | 1 | 1 | 1 | 1 | 1 | 1 | 1 | 1 |
| yeps_YPTB2791 | 1 | 1 | 1 | 1 | 1 | 1 | 1 | 3 | 1 | 1 | 1 | 1 | 1 | 1 | 1 | 1 | 1 | 1 | 1 | 1 | 1 | 1 | 1 | 1 | 1 | 1 | 1 | 1 | 1 | 1 | 1 | 1 | 1 | 1 | 1 |
| yeps_YPTB2792 | 4 | 5 | 4 | 5 | 1 | 1 | 4 | 2 | 5 | 5 | 5 | 3 | 2 | 2 | 1 | 5 | 2 | 1 | 4 | 4 | 1 | 1 | 1 | 4 | 2 | 2 | 1 | 1 | 1 | 1 | 1 | 2 | 5 | 5 | 4 |
| yeps_YPTB2793 | 3 | 4 | 3 | 4 | 1 | 1 | 3 | 3 | 4 | 4 | 4 | 3 | 3 | 3 | 1 | 4 | 3 | 1 | 24 | 3 | 1 | 1 | 1 | 3 | 3 | 3 | 1 | 1 | 1 | 1 | 1 | 3 | 4 | 4 | 3 |
| yeps_YPTB2795 | 5 | 8 | 5 | 4 | 1 | 1 | 5 | 1 | 8 | 4 | 4 | 3 | 4 | 4 | 1 | 4 | 4 | 1 | 5 | 5 | 1 | 1 | 1 | 5 | 4 | 4 | 1 | 1 | 1 | 1 | 1 | 4 | 4 | 4 | 5 |
| yeps_YPTB2796 | 5 | 9 | 5 | 13 | 1 | 1 | 5 | 10 | 9 | 13 | 13 | 3 | 4 | 4 | 1 | 13 | 4 | 6 | 5 | 5 | 1 | 1 | 1 | 5 | 4 | 4 | 1 | 1 | 1 | 1 | 1 | 4 | 13 | 13 | 5 |
| yeps_YPTB2800 | 2 | 6 | 2 | 10 | 1 | 1 | 2 | 8 | 77 | 10 | 10 | 2 | 3 | 3 | 1 | 10 | 3 | 1 | 2 | 2 | 1 | 1 | 1 | 2 | 3 | 3 | 1 | 1 | 1 | 1 | 1 | 3 | 10 | 10 | 2 |
| yeps_YPTB2801 | 5 | 8 | 15 | 14 | 1 | 1 | 5 | 12 | 8 | 14 | 14 | 3 | 4 | 4 | 1 | 14 | 4 | 1 | 5 | 5 | 1 | 1 | 1 | 5 | 4 | 4 | 1 | 1 | 1 | 1 | 1 | 4 | 14 | 14 | 5 |
| yeps_YPTB2802 | 4 | 7 | 4 | 10 | 1 | 1 | 4 | 8 | 7 | 10 | 10 | 3 | 1 | 1 | 1 | 10 | 1 | 5 | 4 | 4 | 1 | 1 | 1 | 4 | 1 | 1 | 1 | 1 | 1 | 1 | 1 | 1 | 10 | 10 | 4 |
| yeps_YPTB2803 | 4 | 8 | 4 | 12 | 1 | 1 | 4 | 10 | 8 | 12 | 12 | 3 | 1 | 1 | 1 | 12 | 1 | 5 | 4 | 4 | 1 | 1 | 1 | 4 | 1 | 1 | 1 | 1 | 1 | 1 | 1 | 1 | 12 | 12 | 4 |
| yeps_YPTB2804 | 1 | 1 | 1 | 1 | 1 | 1 | 1 | 1 | 1 | 1 | 1 | 1 | 1 | 1 | 1 | 1 | 1 | 1 | 1 | 1 | 1 | 1 | 1 | 1 | 1 | 1 | 1 | 1 | 1 | 1 | 1 | 1 | 1 | 1 | 1 |
| yeps_YPTB2805 | 4 | 10 | 4 | 5 | 1 | 1 | 4 | 11 | 10 | 13 | 5 | 3 | 7 | 7 | 1 | 13 | 7 | 5 | 4 | 4 | 1 | 1 | 1 | 4 | 7 | 7 | 1 | 1 | 1 | 1 | 1 | 7 | 5 | 5 | 4 |
| yeps_YPTB2808 | 3 | 8 | 3 | 12 | 14 | 1 | 3 | 10 | 8 | 12 | 12 | 3 | 1 | 1 | 1 | 12 | 1 | 4 | 3 | 3 | 1 | 1 | 1 | 3 | 1 | 33 | 1 | 1 | 1 | 1 | 1 | 1 | 12 | 12 | 3 |
| yeps_YPTB2812 | 12 | 7 | 4 | 10 | 1 | 1 | 4 | 8 | 7 | 10 | 65 | 3 | 1 | 1 | 1 | 10 | 1 | 5 | 4 | 4 | 1 | 1 | 1 | 4 | 1 | 1 | 1 | 1 | 1 | 1 | 1 | 1 | 10 | 10 | 4 |
| yeps_YPTB2814 | 5 | 10 | 5 | 13 | 1 | 1 | 5 | 11 | 10 | 13 | 13 | 3 | 4 | 4 | 1 | 13 | 4 | 8 | 5 | 5 | 1 | 1 | 1 | 5 | 4 | 4 | 1 | 1 | 1 | 1 | 1 | 4 | 13 | 13 | 5 |
| yeps_YPTB2815 | 5 | 10 | 5 | 15 | 1 | 1 | 5 | 11 | 10 | 14 | 15 | 3 | 4 | 4 | 1 | 14 | 4 | 1 | 5 | 5 | 1 | 1 | 1 | 5 | 4 | 4 | 1 | 1 | 1 | 1 | 1 | 4 | 15 | 15 | 5 |
| yeps_YPTB2816 | 5 | 9 | 5 | 9 | 1 | 1 | 5 | 5 | 9 | 10 | 9 | 3 | 5 | 5 | 1 | 10 | 5 | 6 | 5 | 5 | 1 | 1 | 1 | 5 | 5 | 5 | 1 | 1 | 1 | 1 | 1 | 5 | 9 | 9 | 5 |
| yeps_YPTB2818 | 4 | 4 | 4 | 4 | 1 | 1 | 4 | 3 | 4 | 4 | 4 | 3 | 4 | 4 | 1 | 4 | 4 | 1 | 4 | 4 | 1 | 1 | 1 | 4 | 18 | 4 | 1 | 1 | 1 | 1 | 1 | 4 | 4 | 4 | 4 |
| yeps_YPTB2821 | 4 | 5 | 4 | 5 | 1 | 1 | 4 | 2 | 5 | 5 | 5 | 3 | 3 | 3 | 1 | 5 | 3 | 1 | 4 | 4 | 1 | 1 | 1 | 4 | 3 | 3 | 1 | 1 | 1 | 1 | 1 | 3 | 5 | 5 | 4 |
| yeps_YPTB2822 | 1 | 1 | 1 | 1 | 1 | 1 | 1 | 5 | 1 | 1 | 1 | 1 | 1 | 1 | 1 | 1 | 1 | 1 | 1 | 1 | 1 | 1 | 1 | 1 | 1 | 1 | 1 | 1 | 1 | 1 | 1 | 1 | 1 | 1 | 1 |
| yeps_YPTB2824 | 3 | 5 | 3 | 5 | 1 | 1 | 3 | 6 | 5 | 5 | 5 | 2 | 1 | 1 | 1 | 5 | 1 | 2 | 3 | 3 | 1 | 1 | 1 | 3 | 1 | 1 | 1 | 1 | 1 | 1 | 1 | 1 | 5 | 5 | 3 |
| yeps_YPTB2833 | 4 | 7 | 4 | 7 | 1 | 1 | 4 | 8 | 7 | 10 | 7 | 3 | 11 | 1 | 1 | 10 | 1 | 4 | 4 | 91 | 1 | 1 | 1 | 4 | 1 | 1 | 1 | 1 | 1 | 1 | 1 | 1 | 7 | 7 | 4 |
| yeps_YPTB2835 | 2 | 1 | 2 | 1 | 1 | 1 | 2 | 2 | 1 | 1 | 1 | 1 | 1 | 1 | 1 | 1 | 1 | 2 | 2 | 2 | 1 | 1 | 1 | 2 | 1 | 1 | 1 | 1 | 1 | 1 | 1 | 1 | 1 | 1 | 2 |
| yeps_YPTB2839 | 1 | 3 | 1 | 1 | 1 | 1 | 1 | 1 | 3 | 1 | 1 | 1 | 1 | 1 | 1 | 1 | 1 | 1 | 1 | 1 | 1 | 1 | 1 | 1 | 1 | 1 | 1 | 1 | 1 | 1 | 1 | 1 | 1 | 1 | 1 |
| yeps_YPTB2840 | 3 | 1 | 3 | 1 | 1 | 1 | 3 | 6 | 1 | 1 | 1 | 1 | 8 | 1 | 1 | 1 | 1 | 3 | 3 | 3 | 1 | 1 | 1 | 3 | 1 | 1 | 1 | 1 | 1 | 1 | 1 | 1 | 1 | 1 | 3 |
| yeps_YPTB2841 | 3 | 3 | 3 | 3 | 1 | 1 | 3 | 3 | 3 | 3 | 3 | 1 | 1 | 1 | 1 | 3 | 1 | 3 | 3 | 3 | 1 | 1 | 1 | 3 | 1 | 1 | 1 | 1 | 1 | 1 | 1 | 1 | 3 | 3 | 3 |
| yeps_YPTB2845 | 1 | 1 | 1 | 1 | 1 | 1 | 1 | 6 | 1 | 1 | 1 | 3 | 1 | 1 | 1 | 1 | 1 | 1 | 1 | 1 | 1 | 1 | 1 | 1 | 1 | 1 | 1 | 1 | 1 | 1 | 1 | 1 | 1 | 1 | 1 |
| yeps_YPTB2852 | 1 | 6 | 1 | 9 | 1 | 1 | 1 | 7 | 6 | 9 | 9 | 3 | 1 | 1 | 1 | 9 | 1 | 1 | 1 | 1 | 1 | 1 | 1 | 1 | 1 | 1 | 1 | 1 | 1 | 1 | 1 | 1 | 9 | 9 | 1 |
| yeps_YPTB2856 | 1 | 2 | 1 | 1 | 1 | 1 | 1 | 2 | 2 | 1 | 1 | 2 | 1 | 1 | 1 | 1 | 1 | 2 | 1 | 1 | 1 | 1 | 1 | 1 | 1 | 1 | 1 | 1 | 1 | 1 | 1 | 1 | 1 | 1 | 1 |

| | | | | | | | | | | | | | | | | | | | | | | | | | | | | | | | | | | | |
|---|---|---|---|---|---|---|---|---|---|---|---|---|---|---|---|---|---|---|---|---|---|---|---|---|---|---|---|---|---|---|---|---|---|---|---|
| yeps_YPTB2858 | 1 | 3 | 1 | 3 | 1 | 1 | 1 | 5 | 3 | 3 | 3 | 2 | 1 | 1 | 1 | 3 | 1 | 3 | 1 | 1 | 1 | 1 | 1 | 1 | 1 | 1 | 1 | 1 | 1 | 1 | 1 | 1 | 3 | 3 | 1 |
| yeps_YPTB2860 | 1 | 1 | 1 | 1 | 1 | 1 | 1 | 1 | 1 | 1 | 1 | 1 | 1 | 1 | 1 | 1 | 1 | 1 | 1 | 1 | 1 | 1 | 1 | 1 | 1 | 1 | 1 | 1 | 1 | 1 | 1 | 1 | 1 | 1 | 1 |
| yeps_YPTB2862 | 1 | 1 | 1 | 1 | 1 | 1 | 1 | 3 | 1 | 1 | 1 | 1 | 1 | 1 | 1 | 1 | 1 | 2 | 1 | 1 | 1 | 1 | 1 | 1 | 1 | 1 | 1 | 1 | 1 | 1 | 1 | 1 | 1 | 1 | 1 |
| yeps_YPTB2863 | 1 | 1 | 1 | 1 | 1 | 1 | 16 | 3 | 1 | 4 | 1 | 1 | 1 | 1 | 1 | 4 | 1 | 1 | 1 | 1 | 1 | 1 | 1 | 1 | 1 | 1 | 1 | 1 | 1 | 1 | 1 | 1 | 1 | 1 | 1 |
| yeps_YPTB2864 | 2 | 2 | 2 | 2 | 2 | 22 | 2 | 2 | 2 | 2 | 2 | 2 | 2 | 2 | 22 | 2 | 2 | 2 | 2 | 2 | 22 | 22 | 22 | 2 | 2 | 2 | 22 | 22 | 22 | 22 | 22 | 2 | 2 | 2 | 2 |
| yeps_YPTB2870 | 4 | 1 | 4 | 9 | 1 | 1 | 4 | 1 | 1 | 1 | 9 | 3 | 1 | 1 | 1 | 1 | 1 | 5 | 4 | 4 | 1 | 1 | 1 | 4 | 1 | 1 | 1 | 1 | 1 | 1 | 1 | 1 | 9 | 9 | 4 |
| yeps_YPTB2874 | 4 | 7 | 4 | 7 | 1 | 1 | 4 | 8 | 7 | 7 | 7 | 3 | 1 | 1 | 1 | 7 | 1 | 4 | 4 | 4 | 1 | 1 | 1 | 4 | 1 | 1 | 1 | 1 | 1 | 1 | 1 | 1 | 7 | 7 | 4 |
| yeps_YPTB2880 | 3 | 7 | 3 | 4 | 1 | 1 | 3 | 8 | 7 | 4 | 4 | 1 | 1 | 1 | 1 | 4 | 1 | 4 | 3 | 3 | 1 | 1 | 1 | 3 | 1 | 1 | 1 | 1 | 1 | 1 | 1 | 1 | 4 | 4 | 3 |
| yeps_YPTB2882 | 2 | 6 | 2 | 3 | 1 | 1 | 2 | 7 | 6 | 3 | 3 | 1 | 1 | 1 | 1 | 3 | 1 | 3 | 2 | 2 | 1 | 1 | 1 | 2 | 1 | 1 | 1 | 1 | 1 | 1 | 1 | 1 | 3 | 3 | 2 |
| yeps_YPTB2885 | 1 | 6 | 1 | 8 | 1 | 1 | 7 | 1 | 6 | 8 | 8 | 3 | 1 | 1 | 1 | 8 | 1 | 1 | 1 | 7 | 1 | 1 | 1 | 7 | 1 | 1 | 1 | 1 | 1 | 1 | 1 | 1 | 8 | 8 | 1 |
| yeps_YPTB2888 | 4 | 3 | 4 | 3 | 1 | 1 | 4 | 3 | 3 | 3 | 3 | 3 | 1 | 1 | 1 | 3 | 1 | 3 | 4 | 4 | 1 | 1 | 1 | 4 | 1 | 1 | 1 | 1 | 1 | 1 | 1 | 1 | 3 | 3 | 4 |
| yeps_YPTB2891 | 3 | 5 | 3 | 5 | 1 | 1 | 3 | 2 | 5 | 5 | 5 | 2 | 2 | 2 | 1 | 5 | 2 | 3 | 3 | 3 | 1 | 1 | 1 | 3 | 2 | 2 | 1 | 1 | 1 | 1 | 1 | 2 | 5 | 5 | 3 |
| yeps_YPTB2892 | 1 | 5 | 1 | 6 | 1 | 1 | 1 | 4 | 5 | 6 | 6 | 3 | 3 | 3 | 1 | 6 | 3 | 4 | 1 | 1 | 1 | 1 | 1 | 1 | 3 | 3 | 1 | 1 | 1 | 1 | 1 | 3 | 6 | 6 | 1 |
| yeps_YPTB2896 | 1 | 2 | 1 | 5 | 1 | 1 | 1 | 4 | 2 | 1 | 5 | 1 | 1 | 1 | 1 | 1 | 1 | 1 | 1 | 1 | 1 | 1 | 1 | 1 | 1 | 1 | 1 | 1 | 1 | 1 | 1 | 1 | 5 | 5 | 1 |
| yeps_YPTB2901 | 3 | 6 | 3 | 3 | 1 | 1 | 3 | 7 | 6 | 3 | 3 | 3 | 3 | 3 | 1 | 3 | 3 | 3 | 3 | 3 | 1 | 1 | 1 | 3 | 3 | 3 | 1 | 1 | 1 | 1 | 1 | 3 | 3 | 3 | 3 |
| yeps_YPTB2903 | 1 | 1 | 1 | 1 | 1 | 1 | 1 | 1 | 1 | 1 | 1 | 1 | 1 | 1 | 1 | 1 | 1 | 1 | 1 | 1 | 1 | 1 | 1 | 1 | 1 | 1 | 1 | 1 | 1 | 1 | 1 | 1 | 1 | 1 | 1 |
| yeps_YPTB2904 | 1 | 4 | 1 | 3 | 1 | 1 | 1 | 4 | 4 | 3 | 3 | 3 | 4 | 4 | 1 | 3 | 4 | 4 | 1 | 1 | 1 | 1 | 1 | 1 | 4 | 4 | 1 | 1 | 1 | 1 | 1 | 4 | 3 | 3 | 1 |
| yeps_YPTB2906 | 1 | 9 | 1 | 11 | 1 | 1 | 1 | 4 | 40 | 10 | 11 | 3 | 4 | 4 | 1 | 10 | 4 | 5 | 1 | 1 | 1 | 1 | 1 | 1 | 4 | 4 | 1 | 1 | 1 | 1 | 1 | 4 | 11 | 11 | 1 |
| yeps_YPTB2908 | 1 | 1 | 1 | 1 | 1 | 1 | 1 | 1 | 1 | 5 | 1 | 1 | 1 | 1 | 1 | 5 | 1 | 3 | 1 | 1 | 1 | 1 | 1 | 1 | 1 | 1 | 1 | 1 | 1 | 1 | 1 | 1 | 1 | 1 | 1 |
| yeps_YPTB2910 | 4 | 1 | 4 | 1 | 1 | 1 | 4 | 1 | 1 | 7 | 1 | 3 | 3 | 3 | 1 | 7 | 3 | 1 | 4 | 4 | 1 | 1 | 1 | 4 | 3 | 3 | 1 | 1 | 1 | 1 | 1 | 3 | 1 | 1 | 4 |
| yeps_YPTB2911 | 1 | 5 | 1 | 3 | 1 | 1 | 1 | 1 | 5 | 3 | 3 | 1 | 1 | 1 | 1 | 3 | 1 | 3 | 44 | 1 | 1 | 1 | 1 | 1 | 1 | 1 | 1 | 1 | 1 | 1 | 1 | 1 | 3 | 3 | 1 |
| yeps_YPTB2912 | 8 | 1 | 1 | 1 | 1 | 1 | 1 | 1 | 1 | 1 | 1 | 3 | 3 | 3 | 1 | 1 | 3 | 4 | 1 | 1 | 1 | 1 | 1 | 1 | 3 | 3 | 1 | 1 | 1 | 1 | 1 | 3 | 1 | 1 | 1 |
| yeps_YPTB2913 | 1 | 7 | 1 | 10 | 1 | 1 | 1 | 8 | 7 | 10 | 10 | 3 | 3 | 3 | 1 | 10 | 3 | 4 | 1 | 1 | 1 | 1 | 1 | 1 | 3 | 3 | 1 | 1 | 1 | 1 | 1 | 3 | 10 | 10 | 1 |
| yeps_YPTB2917 | 1 | 10 | 1 | 14 | 1 | 1 | 1 | 11 | 10 | 13 | 14 | 3 | 7 | 7 | 1 | 13 | 7 | 5 | 1 | 1 | 1 | 1 | 1 | 1 | 7 | 7 | 1 | 1 | 1 | 1 | 1 | 7 | 14 | 14 | 1 |
| yeps_YPTB2918 | 1 | 5 | 1 | 8 | 1 | 1 | 1 | 6 | 5 | 8 | 8 | 3 | 4 | 4 | 1 | 8 | 4 | 2 | 1 | 1 | 1 | 1 | 1 | 1 | 4 | 4 | 1 | 1 | 1 | 1 | 1 | 4 | 8 | 8 | 1 |
| yeps_YPTB2922 | 1 | 1 | 1 | 2 | 1 | 1 | 1 | 1 | 1 | 2 | 2 | 1 | 1 | 1 | 1 | 2 | 1 | 1 | 1 | 1 | 1 | 1 | 1 | 1 | 1 | 1 | 1 | 1 | 1 | 1 | 1 | 1 | 2 | 2 | 1 |
| yeps_YPTB2923 | 3 | 5 | 3 | 11 | 1 | 3 | 3 | 8 | 5 | 10 | 11 | 3 | 4 | 4 | 3 | 10 | 4 | 5 | 3 | 3 | 3 | 3 | 3 | 3 | 4 | 4 | 3 | 3 | 3 | 3 | 3 | 4 | 11 | 11 | 3 |
| yeps_YPTB2924 | 1 | 5 | 1 | 1 | 1 | 1 | 1 | 2 | 5 | 1 | 1 | 3 | 4 | 4 | 1 | 1 | 4 | 1 | 1 | 1 | 1 | 1 | 1 | 1 | 4 | 4 | 1 | 1 | 1 | 1 | 1 | 4 | 1 | 1 | 1 |
| yeps_YPTB2925 | 1 | 6 | 1 | 3 | 1 | 1 | 1 | 2 | 6 | 3 | 3 | 3 | 1 | 1 | 1 | 3 | 1 | 3 | 1 | 1 | 1 | 1 | 1 | 1 | 1 | 1 | 1 | 1 | 1 | 1 | 1 | 1 | 3 | 3 | 1 |
| yeps_YPTB2926 | 1 | 4 | 1 | 1 | 1 | 1 | 1 | 1 | 4 | 1 | 1 | 1 | 1 | 1 | 1 | 1 | 1 | 1 | 1 | 1 | 1 | 1 | 1 | 1 | 1 | 1 | 1 | 1 | 1 | 1 | 1 | 1 | 1 | 1 | 1 |
| yeps_YPTB2927 | 1 | 6 | 1 | 8 | 1 | 1 | 1 | 7 | 6 | 8 | 8 | 1 | 1 | 1 | 1 | 8 | 1 | 3 | 1 | 1 | 1 | 1 | 1 | 1 | 1 | 1 | 1 | 1 | 1 | 1 | 1 | 1 | 8 | 8 | 1 |
| yeps_YPTB2928 | 1 | 5 | 1 | 5 | 1 | 1 | 1 | 6 | 5 | 5 | 5 | 1 | 1 | 1 | 1 | 5 | 1 | 3 | 1 | 1 | 1 | 1 | 1 | 1 | 1 | 1 | 1 | 1 | 1 | 1 | 1 | 1 | 5 | 5 | 1 |
| yeps_YPTB2936 | 1 | 5 | 1 | 7 | 1 | 1 | 1 | 3 | 5 | 7 | 7 | 1 | 1 | 1 | 1 | 7 | 1 | 8 | 1 | 1 | 1 | 1 | 1 | 1 | 1 | 1 | 1 | 1 | 1 | 1 | 1 | 1 | 7 | 7 | 1 |
| yeps_YPTB2937 | 1 | 6 | 1 | 7 | 1 | 1 | 1 | 7 | 6 | 7 | 7 | 3 | 1 | 1 | 1 | 7 | 1 | 1 | 1 | 1 | 1 | 1 | 1 | 1 | 1 | 1 | 1 | 1 | 1 | 1 | 1 | 1 | 7 | 7 | 1 |
| yeps_YPTB2938 | 1 | 5 | 1 | 8 | 1 | 1 | 1 | 6 | 5 | 8 | 8 | 1 | 1 | 1 | 1 | 8 | 1 | 3 | 1 | 1 | 1 | 1 | 1 | 1 | 1 | 1 | 1 | 1 | 1 | 1 | 1 | 1 | 8 | 8 | 1 |
| yeps_YPTB2943 | 1 | 5 | 1 | 6 | 1 | 1 | 1 | 2 | 5 | 6 | 6 | 2 | 1 | 1 | 1 | 6 | 1 | 1 | 1 | 1 | 1 | 1 | 1 | 1 | 1 | 1 | 1 | 1 | 1 | 1 | 1 | 1 | 6 | 6 | 1 |
| yeps_YPTB2964 | 1 | 3 | 1 | 2 | 1 | 1 | 1 | 4 | 3 | 2 | 2 | 1 | 1 | 1 | 1 | 2 | 1 | 1 | 1 | 1 | 1 | 1 | 1 | 1 | 1 | 1 | 1 | 1 | 1 | 1 | 1 | 1 | 2 | 2 | 1 |
| yeps_YPTB2969 | 1 | 3 | 1 | 1 | 1 | 1 | 1 | 1 | 3 | 1 | 1 | 1 | 1 | 1 | 1 | 1 | 1 | 1 | 1 | 1 | 1 | 1 | 1 | 1 | 1 | 1 | 1 | 1 | 1 | 1 | 1 | 1 | 1 | 1 | 1 |
| yeps_YPTB2976 | 1 | 4 | 1 | 3 | 1 | 1 | 1 | 1 | 4 | 5 | 3 | 3 | 1 | 1 | 1 | 5 | 1 | 1 | 1 | 1 | 1 | 1 | 1 | 1 | 1 | 1 | 1 | 1 | 1 | 1 | 1 | 1 | 3 | 3 | 1 |
| yeps_YPTB2979 | 1 | 1 | 1 | 1 | 1 | 1 | 1 | 1 | 1 | 1 | 1 | 3 | 1 | 1 | 1 | 1 | 1 | 1 | 1 | 1 | 1 | 1 | 1 | 1 | 1 | 1 | 1 | 1 | 1 | 1 | 1 | 1 | 1 | 1 | 1 |
| yeps_YPTB2980 | 1 | 4 | 1 | 1 | 1 | 1 | 1 | 1 | 4 | 1 | 1 | 1 | 1 | 1 | 1 | 1 | 1 | 1 | 1 | 1 | 1 | 1 | 1 | 1 | 1 | 1 | 1 | 1 | 1 | 1 | 1 | 1 | 1 | 1 | 1 |
| yeps_YPTB2981 | 1 | 1 | 1 | 3 | 1 | 1 | 1 | 1 | 1 | 3 | 3 | 1 | 1 | 1 | 1 | 3 | 1 | 1 | 1 | 1 | 1 | 1 | 1 | 1 | 1 | 1 | 1 | 1 | 1 | 1 | 1 | 1 | 3 | 3 | 1 |
| yeps_YPTB2983 | 1 | 1 | 1 | 1 | 1 | 1 | 1 | 1 | 1 | 1 | 1 | 1 | 1 | 1 | 1 | 1 | 1 | 1 | 1 | 1 | 1 | 1 | 1 | 1 | 1 | 1 | 1 | 1 | 1 | 1 | 1 | 1 | 1 | 1 | 1 |
| yeps_YPTB2984 | 1 | 2 | 1 | 1 | 1 | 1 | 1 | 1 | 2 | 1 | 1 | 1 | 1 | 1 | 1 | 1 | 1 | 2 | 1 | 1 | 1 | 1 | 1 | 1 | 1 | 1 | 1 | 1 | 1 | 1 | 1 | 1 | 1 | 1 | 1 |
| yeps_YPTB2985 | 4 | 3 | 4 | 7 | 1 | 1 | 4 | 7 | 3 | 7 | 7 | 3 | 1 | 1 | 1 | 7 | 1 | 5 | 4 | 4 | 1 | 1 | 1 | 4 | 1 | 1 | 1 | 1 | 1 | 1 | 1 | 1 | 7 | 7 | 4 |
| yeps_YPTB2986 | 2 | 2 | 2 | 1 | 1 | 1 | 2 | 1 | 2 | 1 | 1 | 2 | 1 | 1 | 1 | 1 | 1 | 3 | 2 | 2 | 1 | 1 | 1 | 2 | 1 | 1 | 1 | 1 | 1 | 1 | 1 | 1 | 1 | 1 | 2 |
| yeps_YPTB2988 | 3 | 7 | 3 | 12 | 1 | 1 | 3 | 7 | 7 | 10 | 12 | 1 | 1 | 1 | 1 | 10 | 1 | 4 | 3 | 3 | 1 | 1 | 1 | 3 | 1 | 1 | 1 | 1 | 1 | 1 | 1 | 1 | 12 | 12 | 3 |
| yeps_YPTB2989 | 1 | 1 | 1 | 4 | 1 | 1 | 1 | 1 | 1 | 4 | 4 | 1 | 1 | 1 | 1 | 4 | 1 | 2 | 1 | 1 | 1 | 1 | 1 | 1 | 1 | 1 | 1 | 1 | 1 | 1 | 1 | 1 | 4 | 4 | 1 |
| yeps_YPTB2991 | 1 | 6 | 1 | 1 | 1 | 1 | 1 | 7 | 6 | 1 | 1 | 1 | 1 | 1 | 1 | 1 | 1 | 3 | 1 | 1 | 1 | 1 | 1 | 1 | 1 | 1 | 1 | 1 | 1 | 1 | 1 | 1 | 1 | 1 | 1 |
| yeps_YPTB2995 | 1 | 2 | 1 | 2 | 1 | 1 | 1 | 1 | 2 | 2 | 2 | 1 | 1 | 1 | 1 | 2 | 1 | 2 | 1 | 1 | 1 | 1 | 1 | 1 | 1 | 1 | 1 | 1 | 1 | 1 | 1 | 1 | 2 | 2 | 1 |
| yeps_YPTB2996 | 1 | 1 | 1 | 2 | 1 | 1 | 1 | 2 | 1 | 2 | 2 | 1 | 1 | 1 | 1 | 2 | 1 | 5 | 1 | 1 | 1 | 1 | 1 | 1 | 1 | 1 | 1 | 1 | 1 | 1 | 1 | 1 | 2 | 2 | 1 |
| yeps_YPTB2999 | 6 | 5 | 6 | 2 | 1 | 1 | 6 | 5 | 5 | 2 | 2 | 1 | 1 | 1 | 1 | 2 | 1 | 7 | 6 | 6 | 1 | 1 | 1 | 6 | 1 | 1 | 1 | 1 | 1 | 1 | 1 | 1 | 2 | 2 | 6 |
| yeps_YPTB3000 | 1 | 1 | 1 | 1 | 1 | 1 | 1 | 3 | 1 | 1 | 1 | 1 | 1 | 1 | 1 | 1 | 1 | 1 | 1 | 1 | 1 | 1 | 1 | 1 | 1 | 1 | 1 | 1 | 1 | 1 | 1 | 1 | 1 | 1 | 1 |
| yeps_YPTB3001 | 1 | 1 | 1 | 1 | 1 | 1 | 1 | 1 | 1 | 1 | 1 | 1 | 1 | 1 | 1 | 1 | 1 | 1 | 1 | 1 | 1 | 1 | 1 | 1 | 1 | 1 | 1 | 1 | 1 | 1 | 1 | 1 | 1 | 1 | 1 |
| yeps_YPTB3002 | 1 | 1 | 1 | 1 | 1 | 1 | 1 | 1 | 1 | 1 | 1 | 1 | 1 | 1 | 1 | 1 | 1 | 2 | 1 | 1 | 1 | 1 | 1 | 1 | 1 | 1 | 1 | 1 | 1 | 1 | 1 | 1 | 1 | 1 | 1 |
| yeps_YPTB3007 | 2 | 1 | 2 | 1 | 1 | 1 | 2 | 2 | 1 | 1 | 1 | 1 | 1 | 1 | 1 | 1 | 1 | 1 | 2 | 2 | 1 | 1 | 1 | 2 | 1 | 1 | 1 | 1 | 1 | 1 | 1 | 1 | 1 | 1 | 2 |
| yeps_YPTB3008 | 5 | 7 | 5 | 8 | 1 | 4 | 5 | 6 | 7 | 8 | 8 | 3 | 4 | 4 | 4 | 8 | 4 | 6 | 5 | 5 | 4 | 4 | 4 | 5 | 4 | 4 | 4 | 4 | 4 | 4 | 4 | 4 | 8 | 8 | 5 |
| yeps_YPTB3009 | 2 | 2 | 2 | 5 | 1 | 1 | 2 | 4 | 2 | 2 | 5 | 2 | 1 | 1 | 1 | 2 | 1 | 2 | 2 | 2 | 1 | 1 | 1 | 2 | 1 | 1 | 1 | 1 | 1 | 1 | 1 | 1 | 5 | 5 | 2 |
| yeps_YPTB3016 | 1 | 1 | 1 | 1 | 1 | 1 | 1 | 1 | 1 | 1 | 1 | 1 | 1 | 1 | 1 | 1 | 1 | 1 | 1 | 1 | 1 | 1 | 1 | 1 | 1 | 1 | 1 | 1 | 1 | 1 | 1 | 1 | 1 | 1 | 1 |
| yeps_YPTB3019 | 1 | 3 | 1 | 3 | 1 | 1 | 1 | 5 | 3 | 3 | 3 | 3 | 3 | 3 | 1 | 3 | 3 | 3 | 1 | 1 | 1 | 1 | 1 | 1 | 3 | 3 | 1 | 1 | 1 | 1 | 1 | 3 | 3 | 3 | 1 |
| yeps_YPTB3027 | 2 | 8 | 2 | 10 | 1 | 1 | 2 | 2 | 8 | 10 | 10 | 1 | 1 | 1 | 1 | 10 | 1 | 14 | 2 | 2 | 1 | 1 | 1 | 2 | 1 | 1 | 1 | 1 | 1 | 1 | 1 | 1 | 10 | 10 | 2 |
| yeps_YPTB3028 | 1 | 1 | 1 | 1 | 1 | 1 | 1 | 1 | 1 | 1 | 1 | 1 | 1 | 1 | 1 | 1 | 1 | 1 | 1 | 1 | 1 | 1 | 1 | 1 | 1 | 1 | 1 | 1 | 1 | 1 | 1 | 1 | 1 | 1 | 1 |
| yeps_YPTB3029 | 1 | 1 | 1 | 5 | 1 | 1 | 1 | 1 | 1 | 1 | 5 | 1 | 1 | 1 | 1 | 1 | 1 | 1 | 1 | 1 | 1 | 1 | 1 | 1 | 1 | 1 | 1 | 1 | 1 | 1 | 1 | 1 | 5 | 5 | 1 |
| yeps_YPTB3030 | 1 | 5 | 1 | 1 | 1 | 1 | 1 | 2 | 5 | 1 | 1 | 1 | 1 | 1 | 1 | 1 | 1 | 1 | 1 | 1 | 1 | 1 | 1 | 1 | 1 | 1 | 1 | 1 | 1 | 1 | 1 | 1 | 1 | 1 | 1 |
| yeps_YPTB3031 | 5 | 3 | 1 | | 1 | 1 | 1 | 1 | 3 | 4 | | 1 | 1 | 1 | 1 | 4 | 1 | 1 | 1 | 1 | 1 | 1 | 1 | 1 | 1 | 1 | 1 | 1 | 1 | 1 | 1 | 1 | | | 1 |
| yeps_YPTB3044 | 3 | 4 | 3 | 6 | 1 | 1 | 3 | 5 | 4 | 6 | 6 | 1 | 1 | 1 | 1 | 6 | 1 | 2 | 3 | 3 | 1 | 1 | 1 | 3 | 1 | 1 | 1 | 1 | 1 | 1 | 1 | 1 | 6 | 6 | 3 |
| yeps_YPTB3045 | 3 | 7 | 3 | 9 | 1 | 1 | 3 | 3 | 7 | 3 | 9 | 1 | 1 | 1 | 1 | 3 | 1 | 4 | 3 | 3 | 1 | 1 | 1 | 3 | 1 | 1 | 1 | 1 | 1 | 1 | 1 | 1 | 9 | 9 | 3 |
| yeps_YPTB3046 | 3 | 6 | 3 | 9 | 3 | 3 | 3 | 7 | 6 | 8 | 9 | 3 | 3 | 3 | 3 | 8 | 3 | 4 | 3 | 3 | 3 | 3 | 3 | 3 | 3 | 3 | 3 | 3 | 3 | 3 | 3 | 3 | 9 | 9 | 3 |
| yeps_YPTB3048 | 1 | 4 | 1 | 5 | 1 | 1 | 1 | 2 | 4 | 2 | 5 | 1 | 1 | 1 | 1 | 2 | 1 | 3 | 1 | 1 | 1 | 1 | 1 | 1 | 1 | 1 | 1 | 1 | 1 | 1 | 1 | 1 | 5 | 5 | 1 |
| yeps_YPTB3049 | 1 | 4 | 1 | 2 | 1 | 1 | 1 | 3 | 4 | 2 | 2 | 1 | 1 | 1 | 1 | 2 | 1 | 3 | 1 | 1 | 1 | 1 | 1 | 1 | 1 | 1 | 1 | 1 | 1 | 1 | 1 | 1 | 2 | 2 | 1 |
| yeps_YPTB3076 | 4 | 6 | 4 | 8 | 1 | 1 | 4 | 7 | 6 | 8 | 8 | 3 | 4 | 4 | 1 | 8 | 4 | 4 | 4 | 4 | 1 | 1 | 1 | 4 | 4 | 4 | 1 | 1 | 1 | 1 | 1 | 4 | 8 | 8 | 4 |
| yeps_YPTB3077 | 5 | 8 | 5 | 10 | 1 | 1 | 5 | 2 | 8 | 10 | 10 | 3 | 7 | 7 | 1 | 10 | 7 | 5 | 5 | 5 | 1 | 1 | 1 | 5 | 7 | 7 | 1 | 1 | 1 | 1 | 1 | 7 | 10 | 10 | 5 |
| yeps_YPTB3078 | 2 | 3 | 2 | 2 | 1 | 1 | 2 | 2 | 3 | 2 | 2 | 2 | 1 | 1 | 1 | 2 | 1 | 2 | 2 | 2 | 1 | 1 | 1 | 2 | 1 | 1 | 1 | 1 | 1 | 1 | 1 | 1 | 2 | 2 | 2 |
| yeps_YPTB3079 | 5 | 8 | 5 | 5 | 1 | 1 | 5 | 7 | 8 | 9 | 5 | 3 | 1 | 1 | 1 | 9 | 1 | 5 | 5 | 5 | 1 | 1 | 1 | 5 | 1 | 1 | 1 | 1 | 1 | 1 | 1 | 1 | 5 | 5 | 5 |
| yeps_YPTB3080 | 4 | 7 | 4 | 4 | 1 | 1 | 4 | 8 | 7 | 10 | 4 | 3 | 3 | 3 | 1 | 10 | 3 | 4 | 4 | 4 | 1 | 1 | 1 | 4 | 3 | 3 | 1 | 1 | 1 | 1 | 1 | 3 | 4 | 4 | 4 |
| yeps_YPTB3081 | 3 | 3 | 3 | 7 | 1 | 1 | 3 | 3 | 3 | 6 | 7 | 1 | 1 | 1 | 1 | 6 | 1 | 3 | 3 | 3 | 1 | 1 | 1 | 3 | 1 | 1 | 1 | 1 | 1 | 1 | 1 | 1 | 7 | 7 | 3 |
| yeps_YPTB3082 | 6 | 9 | 6 | 11 | 1 | 1 | 6 | 6 | 9 | 10 | 11 | 3 | 3 | 3 | 1 | 10 | 3 | 6 | 6 | 6 | 1 | 1 | 1 | 6 | 3 | 3 | 1 | 1 | 1 | 1 | 1 | 3 | 11 | 11 | 6 |
| yeps_YPTB3083 | 1 | 2 | 1 | 6 | 1 | 1 | 1 | 1 | 2 | 5 | 6 | 2 | 1 | 1 | 1 | 5 | 1 | 1 | 1 | 1 | 1 | 1 | 1 | 1 | 1 | 1 | 1 | 1 | 1 | 1 | 1 | 1 | 6 | 6 | 1 |
| yeps_YPTB3087 | 4 | 2 | 4 | 8 | 1 | 1 | 4 | 7 | 2 | 8 | 8 | 3 | 1 | 1 | 1 | 8 | 1 | 4 | 4 | 4 | 1 | 1 | 1 | 4 | 1 | 1 | 1 | 1 | 1 | 1 | 1 | 1 | 8 | 8 | 4 |
| yeps_YPTB3088 | 3 | 5 | 3 | 5 | 1 | 1 | 3 | 3 | 5 | 6 | 5 | 1 | 1 | 1 | 1 | 6 | 1 | 3 | 3 | 3 | 1 | 1 | 1 | 3 | 1 | 1 | 1 | 1 | 1 | 1 | 1 | 1 | 5 | 5 | 3 |
| yeps_YPTB3089 | 1 | 5 | 1 | 4 | 1 | 32 | 1 | 1 | 5 | 6 | 4 | 1 | 1 | 1 | 32 | 6 | 1 | 1 | 1 | 1 | 32 | 32 | 32 | 1 | 1 | 1 | 32 | 32 | 32 | 32 | 32 | 1 | 4 | 4 | 1 |
| yeps_YPTB3097 | 4 | 8 | 4 | 8 | 1 | 1 | 4 | 8 | 8 | 8 | 8 | 3 | 3 | 3 | 1 | 8 | 3 | 5 | 4 | 4 | 1 | 1 | 1 | 4 | 3 | 3 | 1 | 1 | 1 | 1 | 1 | 3 | 8 | 8 | 4 |
| yeps_YPTB3098 | 1 | 5 | 1 | 7 | 1 | 1 | 1 | 1 | 5 | 2 | 7 | 2 | 2 | 2 | 1 | 2 | 2 | 1 | 3 | 3 | 1 | 1 | 1 | 3 | 2 | 2 | 1 | 1 | 1 | 1 | 1 | 2 | 7 | 7 | 6 |
| yeps_YPTB3099 | 1 | 1 | 6 | 2 | 1 | 1 | 1 | 4 | 1 | 2 | 2 | 1 | 2 | 2 | 1 | 2 | 2 | 5 | 1 | 1 | 1 | 1 | 1 | 1 | 2 | 2 | 1 | 1 | 1 | 1 | 1 | 2 | 2 | 2 | 1 |
| yeps_YPTB3100 | 1 | 4 | 1 | 4 | 1 | 1 | 1 | 5 | 4 | 4 | 4 | 3 | 3 | 3 | 1 | 4 | 3 | 3 | 1 | 1 | 1 | 1 | 1 | 1 | 3 | 3 | 1 | 1 | 1 | 1 | 1 | 3 | 4 | 4 | 1 |
| yeps_YPTB3101 | 1 | 6 | 1 | 5 | 1 | 1 | 1 | 4 | 6 | 5 | 5 | 2 | 2 | 2 | 1 | 5 | 2 | 4 | 1 | 10 | 1 | 1 | 1 | 10 | 2 | 2 | 1 | 1 | 1 | 1 | 1 | 2 | 5 | 5 | 1 |
| yeps_YPTB3103 | 5 | 7 | 5 | 11 | 1 | 1 | 5 | 9 | 7 | 11 | 11 | 3 | 3 | 3 | 1 | 11 | 3 | 5 | 5 | 5 | 1 | 1 | 1 | 5 | 3 | 3 | 1 | 1 | 1 | 1 | 1 | 3 | 11 | 11 | 5 |
| yeps_YPTB3117 | 3 | 4 | 3 | 6 | 1 | 1 | 3 | 5 | 4 | 6 | 6 | 1 | 1 | 1 | 1 | 6 | 1 | 3 | 3 | 3 | 1 | 1 | 1 | 3 | 1 | 1 | 1 | 1 | 1 | 1 | 1 | 1 | 6 | 6 | 3 |
| yeps_YPTB3118 | 2 | 2 | 2 | 1 | 1 | 1 | 2 | 2 | 2 | 1 | 1 | 1 | 2 | 2 | 1 | 1 | 2 | 2 | 2 | 2 | 1 | 1 | 1 | 2 | 11 | 11 | 1 | 1 | 1 | 1 | 1 | 2 | 1 | 1 | 2 |
| yeps_YPTB3166 | 1 | 6 | 1 | 1 | 1 | 1 | 1 | 1 | 6 | 7 | 1 | 1 | 5 | 5 | 1 | 7 | 5 | 1 | 1 | 1 | 1 | 1 | 1 | 1 | 5 | 5 | 1 | 1 | 1 | 1 | 1 | 5 | 1 | 1 | 1 |
| yeps_YPTB3168 | 2 | 3 | 4 | 3 | 1 | 1 | 2 | 3 | 3 | 3 | 3 | 2 | 3 | 3 | 1 | 3 | 3 | 1 | 2 | 2 | 1 | 1 | 1 | 2 | 3 | 3 | 1 | 1 | 1 | 1 | 1 | 3 | 3 | 3 | 2 |
| yeps_YPTB3171 | 3 | 1 | 3 | 1 | 1 | 1 | 3 | 1 | 1 | 1 | 1 | 1 | 1 | 1 | 1 | 1 | 1 | 1 | 3 | 3 | 1 | 1 | 1 | 3 | 1 | 1 | 1 | 1 | 1 | 1 | 1 | 1 | 1 | 1 | 3 |
| yeps_YPTB3172 | 1 | 1 | 1 | 1 | 1 | 1 | 1 | 1 | 1 | 1 | 1 | 3 | 3 | 3 | 1 | 1 | 3 | 1 | 1 | 1 | 1 | 1 | 1 | 1 | 3 | 3 | 1 | 1 | 1 | 1 | 1 | 3 | 1 | 1 | 1 |
| yeps_YPTB3173 | 1 | 1 | 1 | 1 | 1 | 1 | 1 | 1 | 1 | 1 | 1 | 1 | 1 | 1 | 1 | 1 | 1 | 1 | 1 | 1 | 1 | 1 | 1 | 1 | 1 | 1 | 1 | 1 | 1 | 1 | 1 | 1 | 1 | 1 | 1 |
| yeps_YPTB3174 | 5 | 5 | 5 | 7 | 1 | 1 | 5 | 1 | 5 | 7 | 7 | 3 | 5 | 5 | 1 | 7 | 5 | 1 | 5 | 5 | 1 | 1 | 1 | 5 | 5 | 5 | 1 | 1 | 1 | 1 | 1 | 5 | 7 | 7 | 5 |
| yeps_YPTB3175 | 2 | 3 | | 1 | 1 | 1 | 2 | 1 | 3 | 1 | 1 | 1 | 1 | 1 | 1 | 1 | 1 | 1 | 2 | 2 | 1 | 1 | 1 | 2 | 1 | 1 | 1 | 1 | 1 | 1 | 1 | 1 | 1 | 1 | 2 |
| yeps_YPTB3176 | 3 | 2 | 3 | 6 | 1 | 1 | 3 | 3 | 40 | 6 | 6 | 21 | 3 | 3 | 1 | 6 | 3 | 1 | 3 | 3 | 1 | 1 | 1 | 3 | 3 | 3 | 1 | 1 | 1 | 1 | 1 | 3 | 6 | 6 | 3 |
| yeps_YPTB3184 | 4 | 7 | 4 | 9 | 1 | 1 | 4 | 8 | 7 | 9 | 9 | 3 | 3 | 3 | 1 | 9 | 3 | 1 | 4 | 4 | 1 | 1 | 1 | 4 | 3 | 3 | 1 | 1 | 1 | 1 | 1 | 3 | 9 | 9 | 4 |
| yeps_YPTB3185 | 4 | 7 | 4 | 7 | 1 | 1 | 4 | 8 | 7 | 7 | 7 | 3 | 3 | 3 | 1 | 7 | 3 | 1 | 4 | 4 | 1 | 1 | 1 | 4 | 3 | 3 | 1 | 1 | 1 | 1 | 1 | 3 | 7 | 7 | 4 |
| yeps_YPTB3186 | 1 | 1 | 1 | 3 | 1 | 1 | 1 | 1 | 1 | 3 | 3 | 2 | 2 | 2 | 1 | 3 | 2 | 1 | 1 | 1 | 1 | 1 | 1 | 1 | 2 | 2 | 1 | 1 | 1 | 1 | 1 | 2 | 3 | 3 | 1 |
| yeps_YPTB3187 | 1 | 1 | 1 | 1 | 1 | 1 | 1 | 3 | 1 | 1 | 1 | 1 | 2 | 2 | 1 | 1 | 2 | 1 | 1 | 1 | 1 | 1 | 1 | 1 | 2 | 2 | 1 | 1 | 1 | 1 | 1 | 2 | 1 | 1 | 1 |
| yeps_YPTB3190 | 1 | 1 | 1 | 1 | 1 | 1 | 1 | 1 | 1 | 1 | 1 | 3 | 1 | 1 | 1 | 1 | 1 | 5 | 1 | 1 | 1 | 1 | 1 | 1 | 1 | 1 | 1 | 1 | 1 | 1 | 1 | 1 | 1 | 1 | 1 |
| yeps_YPTB3191 | 1 | 10 | 1 | 12 | 1 | 1 | 1 | 11 | 10 | 12 | 12 | 3 | 1 | 1 | 1 | 12 | 1 | 6 | 1 | 1 | 1 | 1 | 1 | 1 | 1 | 1 | 1 | 1 | 1 | 1 | 1 | 1 | 12 | 12 | 1 |
| yeps_YPTB3192 | 1 | 1 | 1 | 1 | 1 | 1 | 1 | 1 | 1 | 1 | 1 | 1 | 1 | 1 | 1 | 1 | 12 | 3 | 1 | 1 | 1 | 1 | 1 | 1 | 1 | 1 | 1 | 1 | 1 | 1 | 1 | 1 | 1 | 1 | 1 |
| yeps_YPTB3203 | 3 | 5 | 3 | 7 | 1 | 1 | 3 | 1 | 5 | 7 | 7 | 1 | 1 | 1 | 1 | 7 | 1 | 3 | 3 | 3 | 1 | 1 | 1 | 3 | 1 | 1 | 1 | 1 | 1 | 1 | 1 | 1 | 7 | 7 | 3 |
| yeps_YPTB3206 | 3 | 5 | 3 | 1 | 1 | 1 | 3 | 2 | 5 | 1 | 1 | 1 | 1 | 1 | 1 | 1 | 1 | 3 | 3 | 3 | 1 | 1 | 1 | 3 | 1 | 1 | 1 | 1 | 1 | 1 | 1 | 1 | 1 | 1 | 3 |
| yeps_YPTB3207 | 2 | 5 | 2 | 7 | 1 | 1 | 2 | 6 | 5 | 7 | 7 | 1 | 1 | 1 | 1 | 7 | 1 | 3 | 2 | 2 | 1 | 1 | 1 | 2 | 1 | 1 | 1 | 1 | 1 | 1 | 1 | 1 | 7 | 7 | 2 |
| yeps_YPTB3208 | 1 | 1 | 1 | 1 | 1 | 1 | 1 | 1 | 1 | 1 | 1 | 1 | 1 | 1 | 1 | 1 | 1 | 1 | 1 | 1 | 1 | 1 | 1 | 1 | 1 | 1 | 1 | 1 | 1 | 1 | 1 | 1 | 1 | 1 | 1 |
| yeps_YPTB3209 | 2 | 2 | 2 | 2 | 1 | 1 | 2 | 2 | 2 | 2 | 2 | 2 | 1 | 1 | 1 | 2 | 1 | 3 | 2 | 2 | 1 | 1 | 1 | 2 | 1 | 1 | 1 | 1 | 1 | 1 | 1 | 1 | 2 | 2 | 2 |
| yeps_YPTB3211 | 5 | 39 | 5 | 13 | 1 | 1 | 5 | 11 | 10 | 12 | 13 | 3 | 4 | 4 | 1 | 12 | 4 | 8 | 5 | 5 | 1 | 1 | 1 | 5 | 4 | 4 | 1 | 1 | 1 | 1 | 1 | 4 | 13 | 13 | 5 |
| yeps_YPTB3212 | 1 | 6 | 1 | 1 | 1 | 1 | 1 | 7 | 6 | 8 | 1 | 3 | 1 | 1 | 1 | 8 | 1 | 1 | 1 | 1 | 1 | 1 | 1 | 1 | 1 | 1 | 1 | 1 | 1 | 1 | 1 | 1 | 1 | 1 | 1 |
| yeps_YPTB3213 | 1 | 1 | 1 | 7 | 1 | 1 | 1 | 4 | 1 | 1 | 7 | 1 | 1 | 1 | 1 | 1 | 1 | 1 | 1 | 1 | 1 | 1 | 1 | 1 | 1 | 1 | 1 | 1 | 1 | 1 | 1 | 1 | 7 | 7 | 1 |
| yeps_YPTB3214 | 1 | 6 | 1 | 9 | 1 | 1 | 1 | 7 | 6 | 8 | 9 | 3 | 3 | 3 | 1 | 8 | 3 | 1 | 1 | 1 | 1 | 1 | 1 | 1 | 3 | 3 | 1 | 1 | 1 | 1 | 1 | 3 | 9 | 9 | 1 |
| yeps_YPTB3216 | 1 | 1 | 1 | 7 | 1 | 1 | 1 | 1 | 1 | 6 | 7 | 2 | 1 | 1 | 1 | 6 | 1 | 1 | 1 | 1 | 1 | 1 | 1 | 1 | 1 | 1 | 1 | 1 | 1 | 1 | 1 | 1 | 7 | 7 | 1 |
| yeps_YPTB3221 | 1 | 6 | 1 | 1 | 1 | 1 | 1 | 1 | 15 | 1 | 1 | 1 | 1 | 1 | 1 | 1 | 1 | 1 | 1 | 1 | 1 | 1 | 1 | 1 | 1 | 1 | 1 | 1 | 1 | 1 | 1 | 1 | 1 | 1 | 1 |
| yeps_YPTB3223 | 1 | 2 | 1 | 6 | 1 | 1 | 1 | 5 | 2 | 6 | 6 | 1 | 1 | 1 | 1 | 6 | 1 | 3 | 1 | 1 | 1 | 1 | 1 | 1 | 1 | 1 | 1 | 1 | 1 | 1 | 1 | 1 | 6 | 6 | 1 |
| yeps_YPTB3231 | 1 | 5 | 1 | 4 | 1 | 1 | 1 | 7 | 5 | 4 | 4 | 1 | 1 | 1 | 1 | 4 | 1 | 4 | 1 | 1 | 1 | 1 | 1 | 1 | 1 | 1 | 1 | 1 | 1 | 1 | 1 | 1 | 4 | 4 | 1 |
| yeps_YPTB3258 | 1 | 1 | 1 | 1 | 1 | 1 | 1 | 1 | 1 | 1 | 1 | 1 | 1 | 1 | 1 | 1 | 1 | 1 | 1 | 1 | 1 | 1 | 1 | 1 | 1 | 1 | 1 | 1 | 1 | 1 | 1 | 1 | 1 | 1 | 2 |
| yeps_YPTB3261 | 1 | 6 | 1 | 2 | 1 | 1 | 1 | 3 | 6 | 2 | 2 | 3 | 1 | 1 | 1 | 2 | 1 | 5 | 1 | 1 | 1 | 1 | 1 | 1 | 1 | 1 | 1 | 1 | 1 | 1 | 1 | 1 | 2 | 2 | 1 |
| yeps_YPTB3262 | 1 | 9 | 1 | 11 | 1 | 1 | 1 | 3 | 9 | 11 | 11 | 3 | 1 | 1 | 1 | 11 | 1 | 6 | 1 | 1 | 1 | 1 | 1 | 1 | 1 | 1 | 1 | 1 | 1 | 1 | 1 | 1 | 11 | 11 | 1 |
| yeps_YPTB3263 | 1 | 4 | 1 | 1 | 1 | 1 | 1 | 5 | 4 | 1 | 1 | 1 | 1 | 1 | 1 | 1 | 1 | 1 | 1 | 1 | 1 | 1 | 1 | 1 | 1 | 1 | 1 | 1 | 1 | 1 | 1 | 1 | 1 | 1 | 1 |
| yeps_YPTB3264 | 1 | 6 | 1 | 1 | 1 | 1 | 1 | 7 | 6 | 1 | 1 | 3 | 1 | 1 | 1 | 1 | 1 | 4 | 1 | 1 | 1 | 1 | 1 | 1 | 1 | 1 | 1 | 1 | 1 | 1 | 1 | 1 | 1 | 1 | 1 |
| yeps_YPTB3265 | 1 | 8 | 1 | 12 | 1 | 50 | 1 | 9 | 8 | 11 | 12 | 3 | 1 | 1 | 50 | 11 | 1 | 5 | 1 | 1 | 50 | 50 | 50 | 1 | 1 | 1 | 50 | 50 | 50 | 50 | 50 | 1 | 12 | 12 | 1 |
| yeps_YPTB3266 | 1 | 6 | 1 | 11 | 1 | 1 | 1 | 7 | 6 | 9 | 11 | 3 | 1 | 1 | 1 | 9 | 1 | 4 | 1 | 1 | 1 | 1 | 1 | 1 | 1 | 1 | 1 | 1 | 1 | 1 | 1 | 1 | 11 | 11 | 1 |
| yeps_YPTB3267 | 1 | 7 | 1 | 11 | 1 | 3 | 1 | 3 | 7 | 9 | 11 | 3 | 1 | 79 | 3 | 9 | 1 | 5 | 1 | 1 | 3 | 3 | 3 | 1 | 1 | 1 | 3 | 3 | 3 | 3 | 3 | 1 | 11 | 11 | 1 |
| yeps_YPTB3269 | 5 | 8 | 5 | 9 | 1 | 4 | 5 | 1 | 8 | 9 | 9 | 3 | 1 | 1 | 4 | 9 | 1 | 6 | 5 | 5 | 4 | 4 | 4 | 5 | 1 | 1 | 4 | 4 | 4 | 4 | 4 | 1 | 9 | 9 | 5 |
| yeps_YPTB3270 | 1 | 7 | 1 | 9 | 1 | 4 | 1 | 8 | 7 | 9 | 9 | 3 | 1 | 1 | 4 | 9 | 1 | 5 | 1 | 1 | 4 | 4 | 4 | 1 | 1 | 1 | 4 | 4 | 4 | 4 | 4 | 1 | 9 | 9 | 1 |
| yeps_YPTB3290 | 3 | 6 | 3 | 8 | 1 | 1 | 3 | 7 | 6 | 8 | 8 | 1 | 1 | 1 | 1 | 8 | 1 | 4 | 3 | 3 | 1 | 1 | 1 | 3 | 1 | 1 | 1 | 1 | 1 | 1 | 1 | 1 | 8 | 8 | 3 |
| yeps_YPTB3291 | 5 | 8 | 5 | 11 | 4 | 4 | 5 | 9 | 8 | 11 | 11 | 3 | 4 | 4 | 4 | 11 | 4 | 6 | 5 | 5 | 4 | 4 | 4 | 5 | 4 | 4 | 4 | 4 | 4 | 4 | 4 | 4 | 11 | 11 | 5 |
| yeps_YPTB3294 | 4 | 7 | 4 | 9 | 1 | 1 | 4 | 8 | 7 | 9 | 9 | 3 | 1 | 1 | 1 | 9 | 1 | 5 | 4 | 4 | 1 | 1 | 1 | 4 | 1 | 1 | 1 | 1 | 1 | 1 | 1 | 1 | 9 | 9 | 4 |
| yeps_YPTB3295 | 4 | 6 | 4 | 9 | 1 | 1 | 4 | 7 | 6 | 9 | 9 | 3 | 1 | 1 | 1 | 9 | 1 | 2 | 4 | 4 | 1 | 1 | 1 | 4 | 1 | 1 | 1 | 1 | 1 | 1 | 1 | 1 | 9 | 9 | 4 |
| yeps_YPTB3355 | 1 | 4 | 1 | 5 | 1 | 1 | 1 | 1 | 4 | 5 | 5 | 3 | 1 | 1 | 1 | 5 | 1 | 25 | 1 | 1 | 1 | 1 | 1 | 1 | 1 | 1 | 1 | 1 | 1 | 1 | 1 | 1 | 5 | 5 | 1 |
| yeps_YPTB3382 | 1 | 5 | 1 | 6 | 1 | 1 | 1 | 3 | 5 | 6 | 6 | 3 | 1 | 1 | 1 | 6 | 1 | 1 | 1 | 1 | 1 | 1 | 1 | 1 | 1 | 1 | 1 | 1 | 1 | 1 | 1 | 1 | 6 | 6 | 1 |
| yeps_YPTB3384 | 4 | 6 | 4 | 10 | 1 | 1 | 4 | 9 | 6 | 10 | 10 | 3 | 1 | 1 | 1 | 10 | 1 | 5 | 4 | 4 | 1 | 1 | 1 | 4 | 1 | 1 | 1 | 1 | 1 | 1 | 1 | 1 | 10 | 10 | 4 |
| yeps_YPTB3385 | 2 | 6 | 2 | 2 | 1 | 1 | 2 | 7 | 6 | 2 | 2 | 3 | 1 | 1 | 1 | 2 | 1 | 4 | 2 | 2 | 1 | 1 | 1 | 2 | 1 | 1 | 1 | 1 | 1 | 1 | 1 | 1 | 2 | 2 | 2 |
| yeps_YPTB3386 | 1 | 3 | 1 | 6 | 1 | 1 | 1 | 4 | 3 | 6 | 6 | 1 | 1 | 1 | 1 | 6 | 1 | 1 | 1 | 1 | 1 | 1 | 1 | 1 | 1 | 1 | 1 | 1 | 1 | 1 | 1 | 1 | 6 | 6 | 1 |
| yeps_YPTB3387 | 4 | 7 | 4 | 10 | 1 | 1 | 4 | 8 | 7 | 10 | 10 | 3 | 1 | 1 | 1 | 10 | 1 | 5 | 4 | 4 | 1 | 1 | 1 | 4 | 1 | 1 | 1 | 1 | 1 | 1 | 1 | 1 | 10 | 10 | 4 |
| yeps_YPTB3388 | 4 | 9 | 4 | 11 | 1 | 1 | 4 | 10 | 9 | 11 | 11 | 3 | 1 | 1 | 1 | 11 | 1 | 7 | 4 | 4 | 1 | 1 | 1 | 4 | 1 | 1 | 1 | 1 | 1 | 1 | 1 | 1 | 11 | 11 | 4 |
| yeps_YPTB3390 | 1 | 4 | 1 | 11 | 1 | 1 | 1 | 9 | 4 | 11 | 11 | 1 | 1 | 1 | 1 | 11 | 1 | 3 | 1 | 1 | 1 | 1 | 1 | 1 | 1 | 1 | 1 | 1 | 1 | 1 | 1 | 1 | 11 | 11 | 1 |
| yeps_YPTB3391 | 1 | 2 | 1 | 5 | 1 | 1 | 1 | 4 | 2 | 5 | 5 | 1 | 1 | 1 | 1 | 5 | 1 | 3 | 1 | 1 | 1 | 1 | 1 | 1 | 1 | 1 | 1 | 1 | 1 | 1 | 1 | 1 | 5 | 5 | 1 |
| yeps_YPTB3392 | 1 | 7 | 1 | 10 | 1 | 1 | 1 | 8 | 7 | 10 | 10 | 1 | 1 | 1 | 1 | 10 | 1 | 4 | 1 | 1 | 1 | 1 | 1 | 1 | 1 | 1 | 1 | 1 | 1 | 1 | 1 | 1 | 10 | 10 | 1 |
| yeps_YPTB3393 | 1 | 1 | 1 | 1 | 1 | 1 | 1 | 1 | 3 | 1 | 1 | 1 | 1 | 1 | 1 | 1 | 1 | 1 | 1 | 1 | 1 | 1 | 1 | 1 | 1 | 1 | 1 | 1 | 1 | 1 | 1 | 1 | 1 | 1 | 1 |
| yeps_YPTB3394 | 1 | 3 | 1 | 6 | 1 | 1 | 1 | 5 | 3 | 6 | 6 | 1 | 1 | 1 | 1 | 6 | 1 | 3 | 1 | 1 | 1 | 1 | 1 | 1 | 1 | 1 | 1 | 1 | 1 | 1 | 1 | 1 | 6 | 6 | 1 |
| yeps_YPTB3399 | 1 | 1 | 1 | 1 | 1 | 1 | 1 | 4 | 1 | 1 | 1 | 1 | 1 | 1 | 1 | 1 | 1 | 1 | 1 | 1 | 1 | 1 | 1 | 1 | 1 | 1 | 1 | 1 | 1 | 1 | 1 | 1 | 1 | 1 | 1 |
| yeps_YPTB3402 | 3 | 2 | 3 | 1 | 1 | 51 | 3 | 5 | 2 | 1 | 1 | 3 | 1 | 1 | 51 | 1 | 1 | 2 | 3 | 3 | 51 | 51 | 51 | 3 | 1 | 1 | 51 | 51 | 51 | 51 | 51 | 1 | 1 | 1 | 3 |
| yeps_YPTB3403 | 1 | 6 | 1 | 2 | 1 | 1 | 1 | 8 | 6 | 2 | 2 | 2 | 1 | 1 | 1 | 2 | 1 | 3 | 1 | 1 | 1 | 1 | 1 | 1 | 1 | 1 | 1 | 1 | 1 | 1 | 1 | 1 | 2 | 2 | 1 |
| yeps_YPTB3404 | 1 | 3 | 1 | 1 | 1 | 1 | 1 | 6 | 3 | 1 | 1 | 1 | 4 | 4 | 1 | 1 | 4 | 3 | 1 | 1 | 1 | 1 | 1 | 1 | 4 | 4 | 1 | 1 | 1 | 1 | 1 | 4 | 1 | 1 | 1 |
| yeps_YPTB3408 | 1 | 5 | 1 | 1 | 1 | 1 | 1 | 6 | 5 | 1 | 1 | 1 | 1 | 1 | 1 | 1 | 1 | 4 | 1 | 1 | 1 | 1 | 1 | 1 | 1 | 1 | 1 | 1 | 1 | 1 | 1 | 1 | 1 | 1 | 1 |
| yeps_YPTB3409 | 1 | 6 | 1 | 12 | 1 | 3 | 1 | 8 | 7 | 10 | 12 | 3 | 3 | 3 | 3 | 10 | 3 | 4 | 1 | 1 | 3 | 3 | 3 | 1 | 3 | 3 | 3 | 3 | 3 | 3 | 3 | 3 | 12 | 12 | 1 |
| yeps_YPTB3410 | 1 | 2 | 1 | 3 | 1 | 1 | 1 | 2 | 2 | 2 | 3 | 1 | 1 | 1 | 1 | 2 | 1 | 3 | 1 | 1 | 1 | 1 | 1 | 1 | 1 | 1 | 1 | 1 | 1 | 1 | 1 | 1 | 3 | 3 | 1 |
| yeps_YPTB3411 | 1 | 3 | 1 | 5 | 1 | 1 | 1 | 4 | 51 | 1 | 5 | 1 | 1 | 1 | 1 | 1 | 1 | 2 | 1 | 1 | 1 | 1 | 1 | 1 | 1 | 1 | 1 | 1 | 1 | 1 | 1 | 1 | 5 | 5 | 1 |
| yeps_YPTB3412 | 2 | 1 | 2 | 1 | 1 | 1 | 2 | 1 | 1 | 1 | 1 | 1 | 1 | 1 | 1 | 1 | 1 | 3 | 2 | 2 | 1 | 1 | 1 | 2 | 1 | 1 | 1 | 1 | 1 | 1 | 1 | 1 | 1 | 1 | 2 |
| yeps_YPTB3414 | 2 | 4 | 2 | 4 | 1 | 1 | 2 | 1 | 4 | 4 | 4 | 1 | 1 | 1 | 1 | 4 | 1 | 3 | 2 | 2 | 1 | 1 | 1 | 2 | 1 | 1 | 1 | 1 | 1 | 1 | 1 | 1 | 4 | 4 | 2 |
| yeps_YPTB3415 | 7 | 6 | 8 | 7 | 1 | 1 | 8 | 1 | 6 | 7 | 7 | 2 | 1 | 1 | 1 | 7 | 1 | 1 | 7 | 7 | 1 | 1 | 1 | 7 | 1 | 1 | 1 | 1 | 1 | 1 | 1 | 1 | 7 | 7 | 7 |
| yeps_YPTB3417 | 3 | 4 | 3 | 7 | 1 | 1 | 3 | 6 | 4 | 2 | 7 | 1 | 1 | 1 | 1 | 2 | 1 | 2 | 3 | 3 | 1 | 1 | 1 | 3 | 1 | 1 | 1 | 1 | 1 | 1 | 1 | 1 | 7 | 7 | 3 |
| yeps_YPTB3434 | 1 | 1 | 1 | 1 | 1 | 1 | 1 | 1 | 1 | 1 | 1 | 1 | 1 | 1 | 1 | 1 | 1 | 1 | 1 | 1 | 1 | 1 | 1 | 1 | 1 | 1 | 1 | 1 | 1 | 1 | 1 | 1 | 1 | 1 | 1 |
| yeps_YPTB3439 | 5 | 9 | 5 | 12 | 1 | 1 | 5 | 10 | 9 | 12 | 12 | 118 | 5 | 5 | 1 | 12 | 5 | 6 | 5 | 5 | 1 | 1 | 1 | 5 | 5 | 5 | 1 | 1 | 1 | 1 | 1 | 5 | 12 | 12 | 5 |
| yeps_YPTB3440 | 5 | 1 | 5 | 4 | 1 | 1 | 5 | 4 | 1 | 4 | 4 | 3 | 5 | 5 | 1 | 4 | 5 | 1 | 5 | 5 | 1 | 1 | 1 | 5 | 5 | 5 | 1 | 1 | 1 | 1 | 1 | 5 | 4 | 4 | 5 |
| yeps_YPTB3441 | 4 | 4 | 4 | 4 | 1 | 4 | 4 | 6 | 21 | 4 | 4 | 3 | 4 | 4 | 4 | 4 | 4 | 4 | 4 | 4 | 4 | 4 | 4 | 4 | 4 | 4 | 4 | 4 | 4 | 4 | 4 | 4 | 4 | 4 | 4 |
| yeps_YPTB3442 | 1 | 7 | 1 | 9 | 1 | 5 | 1 | 8 | 7 | 9 | 9 | 3 | 1 | 1 | 5 | 9 | 1 | 5 | 1 | 1 | 5 | 5 | 5 | 1 | 1 | 1 | 5 | 5 | 5 | 5 | 5 | 1 | 9 | 9 | 1 |
| yeps_YPTB3443 | 1 | 4 | 1 | 8 | 1 | 4 | 1 | 7 | 4 | 8 | 8 | 3 | 1 | 1 | 4 | 8 | 1 | 4 | 1 | 1 | 4 | 4 | 4 | 1 | 1 | 1 | 4 | 4 | 4 | 4 | 4 | 1 | 8 | 8 | 1 |
| yeps_YPTB3444 | 1 | 6 | 1 | 8 | 1 | 3 | 1 | 2 | 6 | 8 | 8 | 2 | 1 | 1 | 3 | 8 | 1 | 3 | 1 | 1 | 3 | 3 | 3 | 1 | 1 | 1 | 3 | 3 | 3 | 3 | 3 | 1 | 8 | 8 | 1 |
| yeps_YPTB3445 | 1 | 10 | 1 | 16 | 1 | 17 | 1 | 13 | 10 | 15 | 16 | 3 | 8 | 8 | 17 | 15 | 8 | 6 | 1 | 1 | 17 | 17 | 17 | 1 | 8 | 8 | 17 | 17 | 17 | 17 | 17 | 8 | 16 | 16 | 12 |
| yeps_YPTB3446 | 1 | 6 | 1 | 8 | 1 | 1 | 1 | 7 | 6 | 8 | 8 | 1 | 1 | 1 | 1 | 8 | 1 | 1 | 1 | 1 | 1 | 1 | 1 | 1 | 1 | 1 | 1 | 1 | 1 | 1 | 1 | 1 | 8 | 8 | 1 |
| yeps_YPTB3447 | 1 | 6 | 1 | 9 | 1 | 1 | 1 | 7 | 6 | 9 | 9 | 1 | 1 | 1 | 1 | 9 | 1 | 3 | 1 | 1 | 1 | 1 | 1 | 1 | 1 | 1 | 1 | 1 | 1 | 1 | 1 | 1 | 9 | 9 | 1 |
| yeps_YPTB3448 | 1 | 6 | 1 | 10 | 1 | 1 | 1 | 7 | 6 | 9 | 10 | 3 | 1 | 1 | 1 | 9 | 1 | 4 | 1 | 1 | 1 | 1 | 1 | 1 | 1 | 1 | 1 | 1 | 1 | 1 | 1 | 1 | 10 | 10 | 1 |

| | | | | | | | | | | | | | | | | | | | | | | | | | | | | | | | | | | | |
|---|---|---|---|---|---|---|---|---|---|---|---|---|---|---|---|---|---|---|---|---|---|---|---|---|---|---|---|---|---|---|---|---|---|---|---|
| yeps_YPTB3468 | 2 | 3 | 2 | 5 | 1 | 1 | 2 | 3 | 3 | 2 | 5 | 2 | 1 | 29 | 1 | 2 | 1 | 3 | 2 | 2 | 1 | 1 | 1 | 2 | 1 | 1 | 1 | 1 | 1 | 1 | 1 | 1 | 5 | 5 | 2 |
| yeps_YPTB3469 | 4 | 7 | 4 | 10 | 1 | 1 | 4 | 8 | 7 | 10 | 10 | 3 | 1 | 1 | 1 | 10 | 1 | 5 | 4 | 4 | 1 | 1 | 1 | 4 | 1 | 1 | 1 | 1 | 1 | 1 | 1 | 1 | 10 | 10 | 4 |
| yeps_YPTB3472 | 1 | 4 | 1 | 1 | 1 | 1 | 1 | 1 | 4 | 1 | 1 | 3 | 1 | 1 | 1 | 1 | 1 | 1 | 1 | 1 | 1 | 1 | 1 | 1 | 1 | 1 | 1 | 1 | 1 | 1 | 1 | 1 | 1 | 1 | 1 |
| yeps_YPTB3473 | 1 | 1 | 1 | 1 | 1 | 35 | 1 | 5 | 1 | 1 | 1 | 3 | 3 | 3 | 35 | 1 | 3 | 1 | 1 | 1 | 35 | 35 | 35 | 1 | 3 | 3 | 35 | 35 | 35 | 35 | 35 | 3 | 1 | 1 | 1 |
| yeps_YPTB3475 | 1 | 5 | 1 | 3 | 1 | 1 | 1 | 3 | 5 | 3 | 3 | 3 | 3 | 3 | 1 | 3 | 3 | 1 | 1 | 1 | 1 | 1 | 1 | 1 | 3 | 3 | 1 | 1 | 1 | 1 | 1 | 3 | 3 | 3 | 1 |
| yeps_YPTB3477 | 1 | 6 | 1 | 8 | 1 | 1 | 1 | 10 | 6 | 8 | 8 | 3 | 3 | 3 | 1 | 8 | 3 | 1 | 1 | 1 | 1 | 1 | 1 | 1 | 3 | 3 | 1 | 1 | 1 | 1 | 1 | 3 | 8 | 8 | 1 |
| yeps_YPTB3480 | 1 | 1 | 1 | 1 | 1 | 1 | 1 | 4 | 1 | 1 | 1 | 1 | 1 | 1 | 1 | 1 | 1 | 2 | 1 | 1 | 1 | 1 | 1 | 1 | 1 | 1 | 1 | 1 | 1 | 1 | 1 | 1 | 1 | 1 | 1 |
| yeps_YPTB3481 | 1 | 1 | 1 | 1 | 1 | 1 | 1 | 4 | 1 | 31 | 1 | 1 | 1 | 1 | 1 | 1 | 1 | 1 | 1 | 1 | 1 | 1 | 1 | 1 | 1 | 1 | 1 | 1 | 1 | 1 | 1 | 1 | 1 | 1 | 1 |
| yeps_YPTB3482 | 1 | 1 | 1 | 1 | 1 | 1 | 1 | 1 | 1 | 1 | 1 | 1 | 1 | 1 | 1 | 1 | 1 | 1 | 1 | 1 | 1 | 1 | 1 | 1 | 1 | 1 | 1 | 1 | 1 | 1 | 1 | 1 | 1 | 1 | 1 |
| yeps_YPTB3483 | 1 | 5 | 1 | 1 | 1 | 1 | 1 | 1 | 5 | 1 | 1 | 1 | 1 | 1 | 1 | 1 | 1 | 1 | 1 | 1 | 1 | 1 | 1 | 1 | 1 | 1 | 1 | 1 | 1 | 1 | 1 | 1 | 1 | 1 | 1 |
| yeps_YPTB3484 | 1 | 1 | 1 | 1 | 1 | 1 | 1 | 1 | 1 | 1 | 1 | 1 | 1 | 1 | 1 | 1 | 1 | 1 | 1 | 1 | 1 | 1 | 1 | 1 | 1 | 1 | 1 | 1 | 1 | 1 | 1 | 1 | 1 | 1 | 1 |
| yeps_YPTB3486 | 1 | 2 | 1 | 2 | 1 | 1 | 1 | 2 | 2 | 2 | 2 | 1 | 1 | 1 | 1 | 2 | 1 | 2 | 1 | 1 | 1 | 1 | 1 | 1 | 1 | 1 | 1 | 1 | 1 | 1 | 1 | 1 | 2 | 2 | 1 |
| yeps_YPTB3487 | 2 | 2 | 2 | 2 | 1 | 1 | 2 | 2 | 2 | 2 | 2 | 2 | 1 | 1 | 1 | 2 | 1 | 2 | 2 | 2 | 1 | 1 | 1 | 2 | 1 | 1 | 1 | 1 | 1 | 1 | 1 | 1 | 2 | 2 | 2 |
| yeps_YPTB3489 | 3 | 6 | 3 | 3 | 1 | 1 | 3 | 8 | 6 | 3 | 3 | 1 | 1 | 1 | 1 | 3 | 1 | 1 | 3 | 3 | 1 | 1 | 1 | 3 | 1 | 1 | 1 | 1 | 1 | 1 | 1 | 1 | 3 | 3 | 3 |
| yeps_YPTB3490 | 3 | 6 | 3 | 7 | 1 | 1 | 3 | 2 | 6 | 7 | 7 | 2 | 1 | 1 | 1 | 7 | 1 | 2 | 3 | 3 | 1 | 1 | 1 | 3 | 1 | 1 | 1 | 1 | 1 | 1 | 1 | 1 | 7 | 7 | 3 |
| yeps_YPTB3494 | 1 | 1 | 1 | 1 | 1 | 1 | 1 | 1 | 1 | 1 | 1 | 1 | 1 | 1 | 1 | 1 | 1 | 1 | 1 | 1 | 1 | 1 | 1 | 1 | 1 | 1 | 1 | 1 | 1 | 1 | 1 | 1 | 1 | 1 | 1 |
| yeps_YPTB3495 | 3 | 5 | 3 | 1 | 1 | 1 | 3 | 6 | 5 | 1 | 1 | 1 | 1 | 1 | 1 | 1 | 1 | 4 | 3 | 3 | 1 | 1 | 1 | 3 | 1 | 1 | 1 | 1 | 1 | 1 | 1 | 1 | 1 | 1 | 3 |
| yeps_YPTB3496 | 3 | 3 | 3 | 5 | 1 | 1 | 3 | 3 | 3 | 5 | 5 | 1 | 1 | 1 | 1 | 5 | 1 | 4 | 3 | 3 | 1 | 1 | 1 | 3 | 1 | 1 | 1 | 1 | 1 | 1 | 1 | 1 | 5 | 5 | 3 |
| yeps_YPTB3497 | 3 | 7 | 3 | 3 | 1 | 1 | 3 | 4 | 7 | 3 | 3 | 3 | 1 | 1 | 1 | 3 | 1 | 4 | 3 | 3 | 1 | 1 | 1 | 3 | 1 | 1 | 1 | 1 | 1 | 1 | 1 | 1 | 3 | 3 | 3 |
| yeps_YPTB3499 | 2 | 2 | 2 | 2 | 2 | 2 | 2 | 2 | 2 | 2 | 2 | 2 | 2 | 2 | 2 | 2 | 2 | 2 | 2 | 2 | 2 | 2 | 2 | 2 | 2 | 2 | 2 | 2 | 2 | 2 | 2 | 2 | 2 | 2 | 2 |
| yeps_YPTB3500 | 3 | 8 | 3 | 10 | 1 | 1 | 3 | 9 | 8 | 10 | 10 | 2 | 1 | 1 | 1 | 10 | 1 | 4 | 3 | 3 | 1 | 1 | 1 | 3 | 1 | 1 | 1 | 1 | 1 | 1 | 1 | 1 | 10 | 10 | 3 |
| yeps_YPTB3501 | 3 | 3 | 3 | 5 | 1 | 1 | 3 | | 3 | 5 | 5 | 1 | 1 | 1 | 1 | 5 | 1 | 2 | 3 | 3 | 1 | 1 | 1 | 3 | 1 | 1 | 1 | 1 | 1 | 1 | 1 | 1 | 5 | 5 | 3 |
| yeps_YPTB3505 | 2 | 2 | 2 | 3 | 1 | 2 | 2 | 3 | 2 | 3 | 3 | 3 | 1 | 1 | 2 | 3 | 1 | 2 | 2 | 2 | 2 | 2 | 2 | 2 | 1 | 1 | 2 | 2 | 2 | 2 | 2 | 1 | 3 | 3 | 2 |
| yeps_YPTB3509 | 1 | 1 | 1 | 5 | 1 | 1 | 1 | 1 | 1 | 5 | 5 | 1 | 1 | 1 | 1 | 5 | 1 | 3 | 1 | 1 | 1 | 1 | 1 | 1 | 1 | 1 | 1 | 1 | 1 | 1 | 1 | 1 | 5 | 5 | 1 |
| yeps_YPTB3511 | 1 | 2 | 1 | 2 | 1 | 1 | 1 | 5 | 2 | 2 | 2 | 1 | 1 | 1 | 1 | 2 | 1 | 1 | 1 | 1 | 1 | 1 | 1 | 1 | 1 | 1 | 1 | 1 | 1 | 1 | 1 | 1 | 2 | 2 | 1 |
| yeps_YPTB3512 | 1 | 2 | 1 | 2 | 1 | 1 | 1 | 1 | 2 | 2 | 2 | 1 | 1 | 1 | 1 | 2 | 1 | 3 | 1 | 1 | 1 | 1 | 1 | 1 | 1 | 1 | 1 | 1 | 1 | 1 | 1 | 1 | 2 | 2 | 1 |
| yeps_YPTB3513 | 1 | 7 | 1 | 10 | 1 | 1 | 1 | 8 | 7 | 10 | 10 | 3 | 1 | 6 | 1 | 10 | 6 | 4 | 1 | 1 | 1 | 1 | 1 | 1 | 1 | 1 | 1 | 1 | 1 | 1 | 1 | 1 | 10 | 10 | 1 |
| yeps_YPTB3514 | 1 | 1 | 1 | 1 | 1 | 1 | 1 | 1 | 1 | 1 | 1 | 1 | 1 | 1 | 1 | 1 | 1 | 1 | 1 | 1 | 1 | 1 | 1 | 1 | 1 | 1 | 1 | 1 | 1 | 1 | 1 | 1 | 1 | 1 | 1 |
| yeps_YPTB3516 | 2 | 1 | 2 | 1 | 1 | 1 | 2 | 1 | 1 | 1 | 1 | 2 | 1 | 1 | 1 | 1 | 1 | 3 | 2 | 2 | 1 | 1 | 1 | 2 | 1 | 1 | 1 | 1 | 1 | 1 | 1 | 1 | 1 | 1 | 4 |
| yeps_YPTB3517 | 3 | 3 | 3 | 3 | 1 | 3 | 3 | 2 | 3 | 3 | 3 | 3 | 3 | 3 | 3 | 3 | 3 | 3 | 3 | 3 | 3 | 3 | 3 | 3 | 3 | 3 | 3 | 3 | 3 | 3 | 3 | 3 | 3 | 3 | 3 |
| yeps_YPTB3519 | 1 | 4 | 5 | 2 | 1 | 1 | 1 | 2 | 4 | 2 | 2 | 1 | 1 | 1 | 1 | 2 | 1 | 1 | 1 | 1 | 1 | 1 | 1 | 1 | 1 | 1 | 1 | 1 | 1 | 1 | 1 | 1 | 2 | 2 | 1 |
| yeps_YPTB3520 | 1 | 1 | 1 | 1 | 1 | 1 | 1 | 3 | 1 | 1 | 1 | 2 | 1 | 1 | 1 | 1 | 1 | 1 | 1 | 1 | 1 | 1 | 1 | 1 | 1 | 1 | 1 | 1 | 1 | 1 | 1 | 1 | 1 | 1 | 1 |
| yeps_YPTB3521 | 1 | 4 | 1 | 1 | 1 | 1 | 1 | 1 | 4 | 1 | 1 | 2 | 1 | 1 | 1 | 1 | 1 | 3 | 1 | 1 | 1 | 1 | 1 | 1 | 1 | 1 | 1 | 1 | 1 | 1 | 1 | 1 | 1 | 1 | 1 |
| yeps_YPTB3522 | 1 | 2 | 1 | 2 | 1 | 1 | 1 | 1 | 2 | 2 | 2 | 3 | 1 | 1 | 1 | 2 | 1 | 1 | 1 | 1 | 1 | 1 | 1 | 1 | 1 | 1 | 1 | 1 | 1 | 1 | 1 | 1 | 2 | 2 | 1 |
| yeps_YPTB3529 | 3 | 1 | 3 | 1 | 1 | 3 | 3 | 1 | 1 | 1 | 1 | 1 | 1 | 1 | 3 | 1 | 1 | 4 | 3 | 3 | 3 | 3 | 3 | 3 | 1 | 1 | 3 | 3 | 3 | 3 | 3 | 1 | 1 | 1 | 3 |
| yeps_YPTB3530 | 1 | 2 | 1 | 2 | 1 | 1 | 1 | 2 | 2 | 2 | 2 | 3 | 1 | 1 | 1 | 2 | 1 | 2 | 1 | 1 | 1 | 1 | 1 | 1 | 1 | 1 | 1 | 1 | 1 | 1 | 1 | 1 | 2 | 2 | 1 |
| yeps_YPTB3532 | 2 | 2 | 2 | 2 | 1 | 2 | 2 | 4 | 2 | 2 | 2 | 3 | 1 | 1 | 2 | 2 | 1 | 2 | 2 | 2 | 2 | 2 | 2 | 2 | 1 | 1 | 2 | 2 | 2 | 2 | 2 | 1 | 2 | 2 | 2 |
| yeps_YPTB3541 | 3 | 7 | 3 | 9 | 1 | 1 | 3 | 8 | 7 | 9 | 9 | 27 | 1 | 1 | 1 | 9 | 1 | 4 | 3 | 3 | 1 | 1 | 1 | 3 | 1 | 1 | 1 | 1 | 1 | 1 | 1 | 1 | 9 | 9 | 3 |
| yeps_YPTB3542 | 1 | 4 | 1 | 5 | 1 | 1 | 1 | 4 | 4 | 5 | 5 | 3 | 1 | 1 | 1 | 5 | 1 | 3 | 1 | 1 | 1 | 1 | 1 | 1 | 1 | 1 | 1 | 1 | 1 | 1 | 1 | 1 | 5 | 5 | 1 |
| yeps_YPTB3543 | 1 | 1 | 1 | 2 | 1 | 1 | 1 | 1 | 1 | 2 | 2 | 1 | 1 | 1 | 1 | 2 | 1 | 1 | 1 | 1 | 1 | 1 | 1 | 1 | 1 | 1 | 1 | 1 | 1 | 1 | 1 | 1 | 2 | 2 | 1 |
| yeps_YPTB3544 | 1 | 2 | 1 | 2 | 1 | 1 | 1 | 5 | 2 | 2 | 2 | 1 | 1 | 1 | 1 | 2 | 1 | 2 | 1 | 1 | 1 | 1 | 1 | 1 | 1 | 1 | 1 | 1 | 1 | 1 | 1 | 1 | 2 | 2 | 1 |
| yeps_YPTB3546 | 1 | 7 | 1 | 9 | 1 | 3 | 1 | 8 | 7 | 9 | 9 | 1 | 1 | 1 | 3 | 9 | 1 | 3 | 1 | 1 | 3 | 3 | 3 | 1 | 1 | 1 | 3 | 3 | 3 | 3 | 3 | 1 | 9 | 9 | 1 |
| yeps_YPTB3566 | 9 | 8 | 9 | 12 | 1 | 1 | 9 | 8 | 8 | 11 | 12 | 3 | 1 | 1 | 1 | 11 | 1 | 5 | 9 | 9 | 1 | 1 | 1 | 9 | 1 | 1 | 1 | 1 | 1 | 1 | 1 | 1 | 12 | 12 | 9 |
| yeps_YPTB3567 | 10 | 7 | 1 | 9 | 1 | 1 | 1 | 8 | 7 | 9 | 9 | 3 | 1 | 1 | 1 | 9 | 1 | 4 | 1 | 1 | 1 | 1 | 1 | 1 | 1 | 1 | 1 | 1 | 1 | 1 | 1 | 1 | 9 | 9 | 1 |
| yeps_YPTB3571 | 1 | 7 | 1 | 8 | 1 | 1 | 1 | 1 | 7 | 8 | 8 | 1 | 1 | 1 | 1 | 8 | 1 | 4 | 1 | 1 | 1 | 1 | 1 | 1 | 1 | 1 | 1 | 1 | 1 | 1 | 1 | 1 | 8 | 8 | 1 |
| yeps_YPTB3579 | 1 | 3 | 1 | 2 | 1 | 1 | 1 | 5 | 3 | 2 | 2 | 3 | 1 | 1 | 1 | 2 | 1 | 4 | 1 | 1 | 1 | 1 | 1 | 1 | 1 | 1 | 1 | 1 | 1 | 1 | 1 | 1 | 2 | 2 | 1 |
| yeps_YPTB3580 | 9 | 43 | 9 | | 1 | 1 | 9 | 11 | 8 | 12 | 12 | 3 | 1 | 1 | 1 | 12 | 1 | 5 | 9 | 9 | 1 | 1 | 1 | 9 | 1 | 1 | 1 | 1 | 1 | 1 | 1 | 1 | 12 | 12 | 9 |
| yeps_YPTB3581 | 3 | 1 | 3 | 1 | 1 | 1 | 3 | 2 | 1 | 1 | 1 | 2 | 1 | 1 | 1 | 1 | 1 | 1 | 3 | 3 | 1 | 1 | 1 | 3 | 1 | 1 | 1 | 1 | 1 | 1 | 1 | 1 | 1 | 1 | 3 |
| yeps_YPTB3582 | 4 | 6 | 4 | 6 | 1 | 1 | 4 | 7 | 6 | 6 | 6 | 3 | 1 | 1 | 1 | 6 | 1 | 5 | 4 | 4 | 1 | 1 | 1 | 4 | 1 | 1 | 1 | 1 | 1 | 1 | 1 | 1 | 6 | 6 | 4 |
| yeps_YPTB3589 | 1 | 4 | 1 | 7 | 1 | 1 | 1 | 5 | 4 | 7 | 7 | 1 | 1 | 1 | 1 | 7 | 1 | 1 | 1 | 1 | 1 | 1 | 1 | 1 | 1 | 1 | 1 | 1 | 1 | 1 | 1 | 1 | 7 | 7 | 1 |
| yeps_YPTB3590 | 3 | 6 | 3 | 8 | 1 | 1 | 3 | 7 | 6 | 8 | 8 | 1 | 1 | 1 | 1 | 8 | 1 | 1 | 3 | 3 | 1 | 1 | 1 | 3 | 1 | 1 | 1 | 1 | 1 | 1 | 1 | 1 | 8 | 8 | 3 |
| yeps_YPTB3591 | 2 | 1 | 2 | 7 | 1 | 1 | 2 | 5 | 1 | 7 | 7 | 1 | 1 | 1 | 1 | 7 | 1 | 1 | 2 | 2 | 1 | 1 | 1 | 2 | 1 | 1 | 1 | 1 | 1 | 1 | 1 | 1 | 7 | 7 | 2 |
| yeps_YPTB3592 | 1 | 8 | 1 | 10 | 1 | 1 | 1 | 9 | 8 | 10 | 10 | 3 | 1 | 1 | 1 | 10 | 1 | 4 | 1 | 1 | 1 | 1 | 1 | 1 | 1 | 1 | 1 | 1 | 1 | 1 | 1 | 1 | 10 | 10 | 1 |
| yeps_YPTB3593 | 1 | 9 | 1 | 9 | 1 | 1 | 1 | 10 | 9 | 9 | 9 | 3 | 1 | 1 | 1 | 9 | 1 | 5 | 1 | 1 | 1 | 1 | 1 | 1 | 1 | 1 | 1 | 1 | 1 | 1 | 1 | 1 | 9 | 9 | 1 |
| yeps_YPTB3595 | 1 | 3 | 1 | 8 | 1 | 1 | 1 | 1 | 3 | 8 | 8 | 3 | 1 | 1 | 1 | 8 | 1 | 1 | 1 | 1 | 1 | 1 | 1 | 1 | 1 | 1 | 1 | 1 | 1 | 1 | 1 | 1 | 8 | 8 | 1 |
| yeps_YPTB3596 | 4 | 6 | 4 | 10 | 1 | 4 | 4 | 8 | 6 | 10 | 10 | 3 | 1 | 1 | 4 | 10 | 1 | 5 | 4 | 4 | 4 | 4 | 4 | 4 | 1 | 1 | 4 | 4 | 4 | 4 | 4 | 1 | 10 | 10 | 4 |
| yeps_YPTB3597 | 3 | 8 | 3 | 11 | 1 | 3 | 3 | 9 | 8 | 11 | 11 | 1 | 1 | 1 | 3 | 11 | 1 | 4 | 3 | 3 | 3 | 3 | 3 | 3 | 1 | 1 | 3 | 3 | 3 | 3 | 3 | 1 | 11 | 11 | 3 |
| yeps_YPTB3598 | 61 | 9 | 3 | 9 | 1 | 3 | 3 | 10 | 9 | 9 | 9 | 1 | 1 | 1 | 3 | 9 | 1 | 6 | 3 | 3 | 3 | 3 | 3 | 3 | 1 | 1 | 3 | 3 | 3 | 3 | 3 | 1 | 9 | 9 | 3 |
| yeps_YPTB3599 | 4 | 8 | 4 | 10 | 1 | 4 | 4 | 9 | 8 | 10 | 10 | 3 | 1 | 1 | 4 | 10 | 1 | 3 | 4 | 4 | 4 | 4 | 4 | 4 | 1 | 1 | 4 | 4 | 4 | 4 | 4 | 1 | 10 | 10 | 4 |
| yeps_YPTB3603 | 5 | 5 | 5 | 10 | 1 | 31 | 5 | 8 | 5 | 10 | 10 | 3 | 1 | 1 | 31 | 10 | 1 | 5 | 5 | 5 | 31 | 31 | 31 | 5 | 1 | 1 | 31 | 31 | 31 | 31 | 31 | 1 | 10 | 10 | 5 |
| yeps_YPTB3604 | 3 | 5 | 3 | 7 | 1 | 1 | 3 | 6 | 5 | 7 | 7 | 1 | 1 | 1 | 1 | 7 | 1 | 4 | 3 | 3 | 1 | 1 | 1 | 3 | 1 | 1 | 1 | 1 | 1 | 1 | 1 | 1 | 7 | 7 | 3 |
| yeps_YPTB3607 | 4 | 7 | 4 | 10 | 1 | 1 | 4 | 7 | 7 | 10 | 10 | 3 | 1 | 1 | 1 | 10 | 1 | 4 | 4 | 4 | 1 | 1 | 1 | 4 | 1 | 1 | 1 | 1 | 1 | 1 | 1 | 1 | 10 | 10 | 4 |
| yeps_YPTB3608 | 1 | 7 | 1 | 12 | 1 | 1 | 1 | 9 | 8 | 11 | 12 | 3 | 1 | 1 | 1 | 11 | 1 | 4 | 1 | 1 | 1 | 1 | 1 | 1 | 1 | 1 | 1 | 1 | 1 | 1 | 1 | 1 | 12 | 12 | 1 |
| yeps_YPTB3609 | 3 | 1 | 3 | 1 | 1 | 1 | 3 | 1 | 1 | 1 | 1 | 3 | 1 | 1 | 1 | 1 | 1 | 3 | 3 | 3 | 1 | 1 | 1 | 3 | 1 | 1 | 1 | 1 | 1 | 1 | 1 | 1 | 1 | 1 | 3 |
| yeps_YPTB3610 | 5 | 3 | 5 | 3 | 1 | 1 | 5 | 3 | 3 | 3 | 3 | 3 | 1 | 1 | 1 | 3 | 1 | 1 | 5 | 5 | 1 | 1 | 1 | 5 | 1 | 1 | 1 | 1 | 1 | 1 | 1 | 1 | 3 | 3 | 5 |
| yeps_YPTB3611 | 4 | 42 | 4 | 11 | 1 | 1 | 4 | 9 | 8 | 11 | 11 | 3 | 1 | 1 | 1 | 11 | 1 | 5 | 4 | 4 | 1 | 1 | 1 | 4 | 1 | 1 | 1 | 1 | 1 | 1 | 1 | 1 | 11 | 11 | 4 |
| yeps_YPTB3626 | 4 | 9 | 4 | 13 | 1 | 1 | 4 | 10 | 9 | 12 | 13 | 3 | 1 | 1 | 40 | 12 | 1 | 5 | 4 | 4 | 1 | 1 | 1 | 90 | 40 | 91 | 1 | 1 | 1 | 1 | 1 | 40 | 13 | 13 | 92 |
| yeps_YPTB3628 | 1 | 2 | 1 | 5 | 1 | 1 | 1 | 3 | 2 | 4 | 5 | 2 | 1 | 1 | 1 | 4 | 1 | 2 | 1 | 1 | 1 | 1 | 1 | 1 | 1 | 1 | 1 | 1 | 1 | 1 | 1 | 1 | 5 | 5 | 1 |
| yeps_YPTB3629 | 4 | 7 | 4 | 11 | 1 | 1 | 4 | 8 | 7 | 10 | 11 | 3 | 1 | 1 | 1 | 10 | 1 | 5 | 4 | 4 | 1 | 1 | 1 | 4 | 1 | 1 | 1 | 1 | 1 | 1 | 1 | 1 | 11 | 11 | 4 |
| yeps_YPTB3631 | 4 | 7 | 4 | 12 | 1 | 1 | 4 | 10 | 7 | 12 | 12 | 3 | 1 | 1 | 1 | 12 | 1 | 4 | 4 | 4 | 1 | 1 | 1 | 4 | 1 | 1 | 1 | 1 | 1 | 1 | 1 | 1 | 12 | 12 | 4 |
| yeps_YPTB3632 | 4 | 5 | 7 | 4 | 1 | 37 | 7 | 5 | 5 | 4 | 4 | 3 | 1 | 1 | 37 | 4 | 1 | 4 | 4 | 4 | 37 | 37 | 37 | 4 | 1 | 1 | 37 | 37 | 37 | 37 | 37 | 1 | 4 | 4 | 9 |
| yeps_YPTB3633 | 4 | 7 | 4 | 1 | 1 | 8 | 4 | 9 | 7 | 1 | 1 | 3 | 1 | 1 | 8 | 1 | 1 | 4 | 4 | 4 | 8 | 8 | 8 | 4 | 1 | 1 | 8 | 8 | 8 | 8 | 8 | 1 | 1 | 1 | 4 |
| yeps_YPTB3635 | 5 | 8 | 4 | 12 | 1 | 63 | 4 | 10 | 8 | 12 | 12 | 3 | 1 | 1 | 63 | 12 | 1 | 5 | 4 | 4 | 63 | 63 | 63 | 4 | 1 | 1 | 63 | 63 | 63 | 63 | 63 | 1 | 12 | 12 | 5 |
| yeps_YPTB3636 | 19 | 5 | 2 | 7 | 1 | 1 | 2 | 1 | 5 | 1 | 7 | 1 | 1 | 1 | 1 | 1 | 1 | 2 | 2 | 2 | 1 | 1 | 1 | 2 | 1 | 1 | 1 | 1 | 1 | 1 | 1 | 1 | 7 | 7 | 2 |
| yeps_YPTB3637 | 4 | 6 | 4 | 9 | 1 | 1 | 4 | 7 | 6 | 9 | 9 | 3 | 1 | 1 | 1 | 9 | 1 | 4 | 4 | 4 | 1 | 1 | 1 | 4 | 1 | 1 | 1 | 1 | 1 | 1 | 1 | 1 | 9 | 9 | 4 |
| yeps_YPTB3638 | 3 | 5 | 3 | 8 | 1 | 1 | 3 | 7 | 5 | 8 | 8 | 1 | 1 | 1 | 1 | 8 | 1 | 3 | 3 | 3 | 1 | 1 | 1 | 3 | 1 | 1 | 1 | 1 | 1 | 1 | 1 | 1 | 8 | 8 | 3 |
| yeps_YPTB3639 | 4 | 1 | 4 | 8 | 1 | 1 | 4 | 7 | 1 | 8 | 8 | 3 | 1 | 1 | 1 | 8 | 1 | 4 | 4 | 4 | 1 | 1 | 1 | 4 | 1 | 1 | 1 | 1 | 1 | 1 | 1 | 1 | 8 | 8 | 4 |
| yeps_YPTB3642 | 2 | 4 | 2 | 2 | 1 | 1 | 2 | 6 | 4 | 2 | 2 | 1 | 1 | 1 | 1 | 2 | 1 | 2 | 2 | 2 | 1 | 1 | 1 | 2 | 1 | 1 | 1 | 1 | 1 | 1 | 1 | 1 | 2 | 2 | 2 |
| yeps_YPTB3643 | 3 | 2 | 3 | 3 | 1 | 1 | 3 | 6 | 2 | 3 | 3 | 1 | 1 | 1 | 1 | 3 | 1 | 3 | 3 | 3 | 1 | 1 | 1 | 3 | 1 | 1 | 1 | 1 | 1 | 1 | 1 | 1 | 3 | 3 | 3 |
| yeps_YPTB3645 | 4 | 6 | 4 | 9 | 1 | 1 | 4 | 8 | 6 | 9 | 9 | 3 | 1 | 1 | 1 | 9 | 1 | 4 | 4 | 4 | 1 | 1 | 1 | 4 | 1 | 1 | 1 | 1 | 1 | 1 | 1 | 1 | 9 | 9 | 4 |
| yeps_YPTB3646 | 5 | 9 | 5 | 11 | 1 | 1 | 5 | 3 | 9 | 11 | 11 | 3 | 1 | 1 | 1 | 11 | 7 | 5 | 5 | 5 | 1 | 1 | 1 | 5 | 1 | 1 | 1 | 1 | 1 | 1 | 1 | 1 | 11 | 11 | 5 |
| yeps_YPTB3647 | 3 | 2 | 3 | 2 | 1 | 1 | 3 | 2 | 2 | 2 | 2 | 2 | 1 | 1 | 1 | 2 | 1 | 3 | 3 | 3 | 1 | 1 | 1 | 3 | 1 | 1 | 1 | 1 | 1 | 1 | 1 | 1 | 2 | 2 | 3 |
| yeps_YPTB3648 | 2 | 1 | 2 | 5 | 1 | 1 | 2 | 1 | 1 | 5 | 5 | 1 | 1 | 1 | 1 | 5 | 1 | 2 | 2 | 2 | 1 | 1 | 1 | 2 | 1 | 1 | 1 | 1 | 1 | 1 | 1 | 1 | 5 | 5 | 2 |
| yeps_YPTB3650 | 1 | 1 | 1 | 5 | 1 | 1 | 1 | 1 | 1 | 5 | 5 | 3 | 1 | 6 | 1 | 5 | 6 | 4 | 1 | 1 | 1 | 1 | 1 | 1 | 1 | 1 | 1 | 1 | 1 | 1 | 1 | 1 | 5 | 5 | 1 |
| yeps_YPTB3653 | 4 | 8 | 4 | 11 | 1 | 1 | 4 | 9 | 68 | 11 | 11 | 3 | 12 | 3 | 1 | 11 | 3 | 14 | 4 | 4 | 1 | 1 | 1 | 4 | 3 | 3 | 1 | 1 | 1 | 1 | 1 | 3 | 11 | 11 | 4 |
| yeps_YPTB3654 | 1 | 1 | 1 | 3 | 1 | 1 | 1 | 1 | 1 | 3 | 3 | 1 | 1 | 1 | 1 | 3 | 1 | 3 | 1 | 1 | 1 | 1 | 1 | 1 | 1 | 1 | 1 | 1 | 1 | 1 | 1 | 1 | 3 | 3 | 1 |
| yeps_YPTB3661 | 1 | 1 | 1 | 1 | 1 | 1 | 1 | 1 | 1 | 1 | 1 | 1 | 1 | 1 | 1 | 1 | 1 | 1 | 1 | 1 | 1 | 1 | 1 | 1 | 1 | 1 | 1 | 1 | 1 | 1 | 1 | 1 | 1 | 1 | 1 |
| yeps_YPTB3662 | 2 | 3 | 2 | 1 | 1 | 1 | 2 | 2 | 3 | 1 | 1 | 2 | 1 | 1 | 1 | 1 | 1 | 1 | 2 | 2 | 1 | 1 | 1 | 2 | 1 | 1 | 1 | 1 | 1 | 1 | 1 | 1 | 1 | 1 | 2 |
| yeps_YPTB3666 | 1 | 1 | 1 | 1 | 1 | 1 | 1 | 1 | 1 | 1 | 1 | 1 | 1 | 1 | 1 | 1 | 1 | 1 | 1 | 1 | 1 | 1 | 1 | 1 | 1 | 1 | 1 | 1 | 1 | 1 | 1 | 1 | 1 | 1 | 1 |
| yeps_YPTB3669 | 1 | 1 | 1 | 1 | 1 | 1 | 1 | 1 | 1 | 1 | 1 | 1 | 1 | 1 | 1 | 1 | 1 | 1 | 1 | 1 | 1 | 1 | 1 | 1 | 1 | 1 | 1 | 1 | 1 | 1 | 1 | 1 | 1 | 1 | 1 |
| yeps_YPTB3670 | 2 | 2 | 2 | 2 | 2 | 2 | 2 | 2 | 2 | 2 | 2 | 2 | 2 | 2 | 2 | 2 | 2 | 2 | 2 | 2 | 2 | 2 | 2 | 2 | 2 | 2 | 2 | 2 | 2 | 2 | 2 | 2 | 2 | 2 | 2 |
| yeps_YPTB3671 | 1 | 1 | 1 | 1 | 1 | 1 | 1 | 1 | 1 | 1 | 1 | 1 | 1 | 1 | 1 | 1 | 1 | 1 | 1 | 1 | 1 | 1 | 1 | 1 | 1 | 1 | 1 | 1 | 1 | 1 | 1 | 1 | 1 | 1 | 1 |
| yeps_YPTB3672 | 1 | 1 | 1 | 1 | 1 | 1 | 1 | 1 | 1 | 1 | 1 | 1 | 1 | 1 | 1 | 1 | 1 | 1 | 1 | 1 | 1 | 1 | 1 | 1 | 1 | 1 | 1 | 1 | 1 | 1 | 1 | 1 | 1 | 1 | 1 |
| yeps_YPTB3673 | 1 | 1 | 1 | 1 | 1 | 1 | 1 | 1 | 1 | 1 | 1 | 1 | 1 | 1 | 1 | 1 | 1 | 1 | 1 | 1 | 1 | 1 | 1 | 1 | 1 | 1 | 1 | 1 | 1 | 1 | 1 | 1 | 1 | 1 | 1 |
| yeps_YPTB3703 | 1 | 2 | 1 | 1 | 1 | 1 | 1 | 3 | 2 | 1 | 1 | 1 | 1 | 1 | 1 | 1 | 1 | 1 | 1 | 1 | 1 | 1 | 1 | 1 | 1 | 1 | 1 | 1 | 1 | 1 | 1 | 1 | 1 | 1 | 1 |
| yeps_YPTB3707 | 2 | 2 | 2 | 2 | 1 | 1 | 2 | 2 | 2 | 2 | 2 | 1 | 1 | 1 | 1 | 2 | 1 | 2 | 2 | 2 | 1 | 1 | 1 | 2 | 1 | 1 | 1 | 1 | 1 | 1 | 1 | 1 | 2 | 2 | 2 |
| yeps_YPTB3708 | 1 | 3 | 1 | 4 | 1 | 1 | 1 | 1 | 3 | 4 | 4 | 1 | 1 | 1 | 1 | 4 | 1 | 1 | 1 | 1 | 1 | 1 | 1 | 1 | 1 | 1 | 1 | 1 | 1 | 1 | 1 | 1 | 4 | 4 | 1 |
| yeps_YPTB3713 | 2 | 3 | 2 | 1 | 1 | 2 | 2 | 2 | 1 | 1 | 1 | 2 | 2 | 2 | 2 | 1 | 2 | 2 | 2 | 2 | 2 | 2 | 2 | 2 | 2 | 2 | 2 | 2 | 2 | 2 | 2 | 2 | 1 | 1 | 2 |
| yeps_YPTB3715 | 1 | 3 | 1 | 4 | 1 | 1 | 1 | 1 | 3 | 4 | 4 | 1 | 1 | 1 | 1 | 4 | 1 | 1 | 1 | 1 | 1 | 1 | 1 | 1 | 1 | 1 | 1 | 1 | 1 | 1 | 1 | 1 | 4 | 4 | 1 |
| yeps_YPTB3716 | 5 | 5 | 5 | 9 | 1 | 1 | 5 | 7 | 5 | 9 | 9 | 3 | 4 | 4 | 1 | 9 | 4 | 5 | 5 | 5 | 1 | 1 | 1 | 5 | 4 | 4 | 1 | 1 | 1 | 1 | 1 | 4 | 9 | 9 | 10 |
| yeps_YPTB3719 | 2 | 2 | 2 | 2 | 1 | 1 | 2 | 6 | 2 | 2 | 2 | 1 | 1 | 1 | 1 | 2 | 1 | 2 | 2 | 2 | 1 | 1 | 1 | 2 | 1 | 1 | 1 | 1 | 1 | 1 | 1 | 1 | 2 | 2 | 2 |
| yeps_YPTB3720 | 3 | 2 | 3 | 2 | 36 | 1 | 3 | 5 | 2 | 2 | 2 | 1 | 1 | 1 | 1 | 2 | 1 | 3 | 3 | 3 | 1 | 1 | 1 | 3 | 1 | 1 | 1 | 1 | 1 | 1 | 1 | 1 | 2 | 2 | 3 |
| yeps_YPTB3721 | 2 | 3 | 2 | 2 | 1 | 1 | 2 | 5 | 3 | 2 | 2 | 3 | 3 | 3 | 1 | 2 | 3 | 2 | 2 | 2 | 1 | 1 | 1 | 2 | 3 | 3 | 1 | 1 | 1 | 1 | 1 | 3 | 2 | 2 | 2 |
| yeps_YPTB3723 | 8 | 2 | 3 | 7 | 1 | 1 | 3 | 5 | 2 | 6 | 7 | 1 | 1 | 1 | 1 | 6 | 1 | 1 | 3 | 3 | 1 | 1 | 1 | 3 | 1 | 1 | 1 | 1 | 1 | 1 | 1 | 1 | 7 | 7 | 3 |
| yeps_YPTB3724 | 4 | 8 | 4 | 10 | 1 | 4 | 4 | 6 | 8 | 10 | 10 | 3 | 4 | 4 | 4 | 10 | 4 | 1 | 4 | 4 | 4 | 4 | 4 | 4 | 4 | 4 | 4 | 4 | 4 | 4 | 4 | 4 | 10 | 10 | 4 |
| yeps_YPTB3726 | 1 | 1 | 1 | 3 | 1 | 1 | 1 | 1 | 1 | 3 | 3 | 3 | 1 | 1 | 1 | 3 | 1 | 1 | 1 | 1 | 1 | 1 | 1 | 1 | 1 | 1 | 1 | 1 | 1 | 1 | 1 | 1 | 3 | 3 | 1 |
| yeps_YPTB3727 | 1 | 1 | 1 | 1 | 1 | 1 | 1 | 1 | 1 | 1 | 1 | 1 | 1 | 1 | 1 | 1 | 1 | 1 | 1 | 1 | 1 | 1 | 1 | 1 | 1 | 1 | 1 | 1 | 1 | 1 | 1 | 1 | 1 | 1 | 1 |
| yeps_YPTB3728 | 1 | 4 | 1 | 1 | 1 | 3 | 1 | 1 | 4 | 1 | 1 | 1 | 3 | 3 | 3 | 1 | 3 | 1 | 1 | 1 | 3 | 3 | 3 | 1 | 3 | 3 | 3 | 3 | 3 | 3 | 3 | 3 | 1 | 1 | 1 |
| yeps_YPTB3735 | 1 | 6 | 1 | 8 | 1 | 3 | 1 | 1 | 6 | 8 | 8 | 1 | 3 | 3 | 3 | 8 | 3 | 4 | 1 | 1 | 3 | 3 | 3 | 1 | 3 | 3 | 3 | 3 | 3 | 3 | 3 | 3 | 8 | 8 | 1 |
| yeps_YPTB3736 | 4 | 7 | 4 | 9 | 1 | 3 | 4 | 8 | 7 | 9 | 9 | 3 | 3 | 3 | 3 | 9 | 3 | 1 | 4 | 4 | 3 | 3 | 3 | 4 | 3 | 3 | 3 | 3 | 3 | 3 | 3 | 3 | 9 | 9 | 4 |
| yeps_YPTB3737 | 3 | 8 | 3 | 10 | 1 | 4 | 3 | 9 | 8 | 10 | 10 | 3 | 4 | 4 | 4 | 10 | 4 | 1 | 3 | 3 | 4 | 4 | 4 | 3 | 4 | 4 | 4 | 4 | 4 | 4 | 4 | 4 | 10 | 10 | 3 |
| yeps_YPTB3739 | 4 | 5 | 4 | 12 | 1 | 4 | 4 | 10 | 5 | 5 | 12 | 3 | 4 | 4 | 4 | 5 | 4 | 5 | 4 | 4 | 4 | 4 | 4 | 4 | 4 | 4 | 4 | 4 | 4 | 4 | 4 | 4 | 12 | 12 | 4 |
| yeps_YPTB3740 | 1 | 8 | 1 | 11 | 1 | 4 | 1 | 3 | 8 | 10 | 11 | 3 | 4 | 4 | 4 | 10 | 4 | 106 | 1 | 1 | 4 | 4 | 4 | 1 | 4 | 4 | 4 | 4 | 4 | 4 | 4 | 4 | 11 | 11 | 1 |
| yeps_YPTB3741 | 1 | 1 | 1 | 1 | 1 | 1 | 1 | 1 | 1 | 1 | 1 | 1 | 1 | 1 | 1 | 1 | 1 | 1 | 1 | 1 | 1 | 1 | 1 | 1 | 1 | 1 | 1 | 1 | 1 | 1 | 1 | 1 | 1 | 1 | 1 |
| yeps_YPTB3742 | 1 | 5 | 1 | 7 | 1 | 1 | 1 | 6 | 5 | 7 | 7 | 1 | 1 | 1 | 1 | 7 | 1 | 3 | 1 | 1 | 1 | 1 | 1 | 1 | 1 | 1 | 1 | 1 | 1 | 1 | 1 | 1 | 7 | 7 | 1 |
| yeps_YPTB3743 | 1 | 6 | 1 | 9 | 1 | 3 | 1 | 7 | 6 | 9 | 9 | 1 | 3 | 3 | 3 | 9 | 3 | 4 | 1 | 1 | 3 | 3 | 3 | 1 | 3 | 3 | 3 | 3 | 3 | 3 | 3 | 3 | 9 | 9 | 1 |
| yeps_YPTB3744 | 1 | 1 | 1 | 3 | 1 | 3 | 1 | 1 | 1 | 6 | 3 | 3 | 3 | 3 | 3 | 6 | 3 | 4 | 1 | 1 | 3 | 3 | 3 | 1 | 3 | 3 | 3 | 3 | 3 | 3 | 3 | 3 | 3 | 3 | 1 |
| yeps_YPTB3745 | 3 | 6 | 3 | 2 | 1 | 3 | 3 | 7 | 6 | 2 | 2 | 3 | 3 | 3 | 3 | 2 | 3 | 2 | 8 | 8 | 3 | 3 | 3 | 8 | 3 | 3 | 3 | 3 | 3 | 3 | 3 | 3 | 2 | 2 | 3 |
| yeps_YPTB3747 | 3 | 2 | 3 | 4 | 1 | 1 | 3 | 5 | 2 | 4 | 4 | 1 | 1 | 1 | 1 | 4 | 1 | 3 | 3 | 3 | 1 | 1 | 1 | 3 | 1 | 1 | 1 | 1 | 1 | 1 | 1 | 1 | 4 | 4 | 3 |
| yeps_YPTB3748 | 1 | 2 | 1 | 2 | 1 | 1 | 1 | 2 | 2 | 2 | 2 | 1 | 1 | 1 | 1 | 2 | 1 | 2 | 1 | 1 | 1 | 1 | 1 | 1 | 1 | 1 | 1 | 1 | 1 | 1 | 1 | 1 | 2 | 2 | 1 |
| yeps_YPTB3754 | 3 | 5 | 3 | 5 | 1 | 1 | 3 | 1 | | 5 | 5 | 1 | 1 | 1 | 1 | 5 | 1 | 4 | 3 | 3 | 1 | 1 | 1 | 3 | 1 | 1 | 1 | 1 | 1 | 1 | 1 | 1 | 5 | 5 | 3 |
| yeps_YPTB3755 | 3 | 6 | 3 | 5 | 1 | 1 | 3 | 7 | | 5 | 5 | 1 | 1 | 1 | 1 | 5 | 1 | 4 | 3 | 3 | 1 | 1 | 1 | 3 | 1 | 1 | 1 | 1 | 1 | 1 | 1 | 1 | 5 | 5 | 3 |
| yeps_YPTB3757 | 1 | 2 | 1 | 1 | 1 | 1 | 1 | 1 | 2 | 1 | 1 | 1 | 1 | 1 | 1 | 1 | 1 | 1 | 1 | 1 | 1 | 1 | 1 | 1 | 1 | 1 | 1 | 1 | 1 | 1 | 1 | 1 | 1 | 1 | 1 |
| yeps_YPTB3758 | 3 | 8 | 3 | 10 | 1 | 1 | 3 | 9 | 8 | 10 | 10 | 1 | 1 | 1 | 1 | 10 | 1 | 4 | 3 | 3 | 1 | 1 | 1 | 3 | 1 | 1 | 1 | 1 | 1 | 1 | 1 | 1 | 10 | 10 | 3 |
| yeps_YPTB3759 | 1 | 4 | 1 | 5 | 1 | 1 | 1 | 1 | 4 | 1 | 5 | 1 | 1 | 1 | 1 | 1 | 1 | 1 | 1 | 1 | 1 | 1 | 1 | 1 | 1 | 1 | 1 | 1 | 1 | 1 | 1 | 1 | 5 | 5 | 1 |
| yeps_YPTB3761 | 1 | 4 | 1 | 1 | 1 | 1 | 1 | 5 | 4 | 1 | 1 | 1 | 1 | 23 | 1 | 1 | 1 | 3 | 1 | 1 | 1 | 1 | 1 | 1 | 1 | 1 | 1 | 1 | 1 | 1 | 1 | 1 | 1 | 1 | 1 |
| yeps_YPTB3765 | 3 | 4 | 3 | 4 | 1 | 1 | 3 | 4 | 4 | 4 | 4 | 1 | 1 | 1 | 1 | 4 | 1 | 4 | 3 | 3 | 1 | 1 | 1 | 3 | 1 | 1 | 1 | 1 | 1 | 1 | 1 | 1 | 4 | 4 | 3 |
| yeps_YPTB3769 | 1 | 2 | 1 | 2 | 1 | 1 | 1 | 2 | 2 | 2 | 2 | 1 | 1 | 1 | 1 | 2 | 1 | 2 | 1 | 1 | 1 | 1 | 1 | 1 | 1 | 1 | 1 | 1 | 1 | 1 | 1 | 1 | 2 | 2 | 1 |
| yeps_YPTB3770 | 1 | 2 | 1 | 1 | 1 | 1 | 1 | 1 | 2 | 1 | 1 | 1 | 1 | 1 | 1 | 1 | 1 | 1 | 1 | 1 | 1 | 1 | 1 | 1 | 1 | 1 | 1 | 1 | 1 | 1 | 1 | 1 | 1 | 1 | 1 |
| yeps_YPTB3771 | 1 | 4 | 1 | 2 | 1 | 1 | 1 | 5 | 4 | 2 | 2 | 1 | 1 | 1 | 1 | 2 | 1 | 2 | 1 | 1 | 1 | 1 | 1 | 1 | 1 | 1 | 1 | 1 | 1 | 1 | 1 | 1 | 2 | 2 | 1 |
| yeps_YPTB3772 | 1 | 2 | 1 | 3 | 1 | 1 | 1 | 3 | 2 | 3 | 3 | 1 | 1 | 1 | 1 | 3 | 1 | 3 | 1 | 1 | 1 | 1 | 1 | 1 | 1 | 1 | 1 | 1 | 1 | 1 | 1 | 1 | 3 | 3 | 1 |
| yeps_YPTB3773 | 1 | 1 | 1 | 1 | 1 | 1 | 1 | 1 | 1 | 1 | 1 | 1 | 1 | 1 | 1 | 1 | 1 | 1 | 1 | 1 | 1 | 1 | 1 | 1 | 1 | 1 | 1 | 1 | 1 | 1 | 1 | 1 | 1 | 1 | 1 |
| yeps_YPTB3774 | 1 | 93 | 1 | 6 | 1 | 1 | 1 | 2 | 2 | 6 | 6 | 1 | 1 | 1 | 1 | 6 | 1 | 2 | 1 | 1 | 1 | 1 | 1 | 1 | 1 | 1 | 1 | 1 | 1 | 1 | 1 | 1 | 6 | 6 | 1 |
| yeps_YPTB3775 | 4 | 8 | 4 | 2 | 1 | 3 | 4 | 9 | 8 | 10 | 2 | 3 | 3 | 3 | 3 | 10 | 3 | 6 | 4 | 4 | 3 | 3 | 3 | 4 | 3 | 3 | 3 | 3 | 3 | 3 | 3 | 3 | 2 | 2 | 4 |
| yeps_YPTB3776 | 1 | 10 | 1 | 14 | 1 | 4 | 1 | 12 | 10 | 14 | 14 | 3 | 4 | 4 | 4 | 14 | 4 | 8 | 1 | 1 | 4 | 4 | 4 | 1 | 4 | 4 | 4 | 4 | 4 | 4 | 4 | 4 | 14 | 14 | 1 |
| yeps_YPTB3777 | 1 | 1 | 1 | 1 | 1 | 1 | 1 | 4 | 1 | 1 | 1 | 3 | 1 | 1 | 1 | 1 | 1 | 4 | 1 | 1 | 1 | 1 | 1 | 1 | 1 | 1 | 1 | 1 | 1 | 1 | 1 | 1 | 1 | 1 | 1 |
| yeps_YPTB3778 | 1 | 2 | 1 | 2 | 1 | 2 | 1 | 2 | 2 | 2 | 2 | 1 | 2 | 2 | 2 | 2 | 2 | 3 | 1 | 1 | 2 | 2 | 2 | 1 | 2 | 2 | 2 | 2 | 2 | 2 | 2 | 2 | 2 | 2 | 1 |
| yeps_YPTB3783 | 5 | 8 | 5 | 10 | 1 | 1 | 5 | 9 | 8 | 10 | 10 | 3 | 4 | 4 | 1 | 10 | 4 | 6 | 5 | 5 | 1 | 1 | 1 | 5 | 4 | 4 | 1 | 1 | 1 | 1 | 1 | 4 | 10 | 10 | 5 |
| yeps_YPTB3784 | 1 | 7 | 1 | 9 | 1 | 1 | 1 | 8 | 7 | 9 | 9 | 3 | 3 | 3 | 1 | 9 | 3 | 6 | 1 | 1 | 1 | 1 | 1 | 1 | 3 | 3 | 1 | 1 | 1 | 1 | 1 | 3 | 9 | 9 | 1 |
| yeps_YPTB3785 | 1 | 4 | 1 | 3 | 1 | 1 | 1 | 3 | 4 | 6 | 3 | 3 | 1 | 1 | 1 | 6 | 1 | 3 | 1 | 1 | 1 | 1 | 1 | 1 | 1 | 1 | 1 | 1 | 1 | 1 | 1 | 1 | 3 | 3 | 1 |
| yeps_YPTB3788 | 1 | 9 | 1 | 12 | 1 | 1 | 1 | 10 | 9 | 12 | 12 | 1 | 1 | 1 | 1 | 12 | 1 | 6 | 1 | 1 | 1 | 1 | 1 | 1 | 1 | 1 | 1 | 1 | 1 | 1 | 1 | 1 | 12 | 12 | 1 |
| yeps_YPTB3796 | 3 | 6 | 3 | 7 | 1 | 3 | 3 | 3 | 6 | 8 | 7 | 3 | 4 | 4 | 3 | 8 | 4 | 1 | 3 | 3 | 3 | 3 | 3 | 3 | 4 | 4 | 3 | 3 | 3 | 3 | 3 | 4 | 7 | 7 | 3 |
| yeps_YPTB3797 | 1 | 7 | 8 | 12 | 1 | 1 | 1 | 9 | 7 | 11 | 12 | 3 | 4 | 4 | 1 | 11 | 4 | 5 | 1 | 1 | 1 | 1 | 1 | 1 | 4 | 4 | 1 | 1 | 1 | 1 | 1 | 4 | 12 | 12 | 1 |
| yeps_YPTB3798 | 12 | 8 | 9 | 11 | 1 | 1 | 1 | 10 | 8 | 11 | 11 | 3 | 4 | 4 | 1 | 11 | 4 | 5 | 1 | 1 | 1 | 1 | 1 | 1 | 4 | 4 | 1 | 1 | 1 | 1 | 1 | 4 | 11 | 11 | 1 |
| yeps_YPTB3799 | 1 | 2 | 1 | 4 | 1 | 1 | 1 | 4 | 2 | 4 | 4 | 3 | 4 | 4 | 1 | 4 | 4 | 5 | 1 | 1 | 1 | 1 | 1 | 1 | 4 | 4 | 1 | 1 | 1 | 1 | 1 | 4 | 4 | 4 | 1 |
| yeps_YPTB3802 | 1 | 5 | 1 | 6 | 1 | 1 | 1 | 6 | 5 | 6 | 6 | 1 | 1 | 1 | 1 | 6 | 1 | 3 | 1 | 1 | 1 | 1 | 1 | 1 | 1 | 1 | 1 | 1 | 1 | 1 | 1 | 1 | 6 | 6 | 1 |
| yeps_YPTB3803 | 1 | 1 | 1 | 5 | 1 | 1 | 1 | 1 | 1 | 5 | 5 | 1 | 3 | 3 | 1 | 5 | 3 | 2 | 1 | 1 | 1 | 1 | 1 | 1 | 3 | 3 | 1 | 1 | 1 | 1 | 1 | 3 | 5 | 5 | 1 |
| yeps_YPTB3804 | 1 | 5 | 1 | 6 | 1 | 1 | 1 | 5 | 5 | 6 | 6 | 3 | 3 | 3 | 1 | 6 | 3 | 2 | 1 | 1 | 1 | 1 | 1 | 1 | 3 | 3 | 1 | 1 | 1 | 1 | 1 | 3 | 6 | 6 | 1 |
| yeps_YPTB3805 | 1 | 7 | 1 | 12 | 1 | 1 | 1 | 9 | 7 | 11 | 12 | 3 | 4 | 4 | 1 | 11 | 4 | 5 | 1 | 1 | 1 | 1 | 1 | 1 | 4 | 4 | 1 | 1 | 1 | 1 | 1 | 4 | 12 | 12 | 1 |
| yeps_YPTB3806 | 1 | 6 | 1 | 9 | 1 | 1 | 1 | 7 | 6 | 9 | 9 | 3 | 3 | 3 | 1 | 9 | 3 | 12 | 1 | 1 | 1 | 1 | 1 | 1 | 3 | 3 | 1 | 1 | 1 | 1 | 1 | 3 | 9 | 9 | 1 |
| yeps_YPTB3812 | 1 | 3 | 1 | 4 | 1 | 1 | 1 | 3 | 3 | 5 | 4 | 3 | 3 | 3 | 1 | 5 | 3 | 4 | 1 | 1 | 1 | 1 | 1 | 1 | 3 | 3 | 1 | 1 | 1 | 1 | 1 | 3 | 4 | 4 | 1 |
| yeps_YPTB3813 | 1 | 7 | 1 | 9 | 1 | 1 | 1 | 8 | 7 | 9 | 9 | 3 | 4 | 4 | 1 | 9 | 4 | 5 | 1 | 1 | 1 | 1 | 1 | 1 | 4 | 4 | 1 | 1 | 1 | 1 | 1 | 4 | 9 | 9 | 1 |
| yeps_YPTB3814 | 1 | 1 | 1 | 10 | 1 | 1 | 1 | 7 | 1 | 10 | 10 | 1 | 3 | 3 | 1 | 10 | 3 | 11 | 1 | 1 | 1 | 1 | 1 | 1 | 3 | 3 | 1 | 1 | 1 | 1 | 1 | 3 | 10 | 10 | 1 |
| yeps_YPTB3815 | 1 | 1 | 1 | 9 | 1 | 1 | 1 | 1 | 1 | 8 | 9 | 1 | 3 | 3 | 1 | 8 | 3 | 5 | 1 | 1 | 1 | 1 | 1 | 1 | 3 | 3 | 1 | 1 | 1 | 1 | 1 | 3 | 9 | 9 | 1 |
| yeps_YPTB3818 | 1 | 8 | 1 | 5 | 1 | 1 | 1 | 10 | 8 | 5 | 5 | 3 | 4 | 4 | 1 | 5 | 4 | 5 | 1 | 1 | 1 | 1 | 1 | 1 | 4 | 4 | 1 | 1 | 1 | 1 | 1 | 4 | 5 | 5 | 1 |
| yeps_YPTB3820 | 1 | 7 | 1 | 12 | 1 | 1 | 1 | 10 | 8 | 12 | 12 | 3 | 3 | 3 | 1 | 12 | 3 | 4 | 1 | 1 | 1 | 1 | 1 | 1 | 3 | 3 | 1 | 1 | 1 | 1 | 1 | 3 | 12 | 12 | 1 |
| yeps_YPTB3821 | 1 | 7 | 1 | 10 | 1 | 1 | 1 | 8 | 7 | 10 | 10 | 3 | 4 | 4 | 1 | 10 | 4 | 5 | 1 | 1 | 1 | 1 | 1 | 1 | 4 | 4 | 1 | 1 | 1 | 1 | 1 | 4 | 10 | 10 | 1 |
| yeps_YPTB3829 | 1 | 3 | 1 | 3 | 1 | 3 | 1 | 5 | 3 | 3 | 3 | 3 | 3 | 3 | 3 | 3 | 3 | 1 | 1 | 1 | 3 | 3 | 3 | 1 | 3 | 3 | 3 | 3 | 3 | 3 | 3 | 3 | 3 | 3 | 1 |
| yeps_YPTB3830 | 2 | 5 | 2 | 2 | 1 | 3 | 2 | 2 | 5 | 2 | 2 | 2 | 3 | 3 | 3 | 2 | 3 | 2 | 2 | 2 | 3 | 3 | 3 | 2 | 3 | 3 | 3 | 3 | 3 | 3 | 3 | 3 | 2 | 2 | 2 |
| yeps_YPTB3832 | 1 | 8 | 1 | 12 | 1 | 4 | 1 | 9 | 8 | 12 | 12 | 3 | 4 | 4 | 4 | 12 | 4 | 5 | 1 | 1 | 4 | 4 | 4 | 1 | 4 | 4 | 4 | 4 | 4 | 4 | 4 | 4 | 12 | 12 | 1 |
| yeps_YPTB3834 | 4 | 5 | 4 | 8 | 1 | 3 | 4 | 7 | 5 | 8 | 8 | 3 | 3 | 3 | 3 | 8 | 3 | 4 | 4 | 4 | 3 | 3 | 3 | 4 | 3 | 3 | 3 | 3 | 3 | 3 | 3 | 3 | 8 | 8 | 6 |
| yeps_YPTB3838 | 1 | 3 | 1 | 9 | 1 | 4 | 1 | 8 | 3 | 9 | 9 | 3 | 4 | 4 | 4 | 9 | 4 | 5 | 1 | 1 | 4 | 4 | 4 | 1 | 4 | 4 | 4 | 4 | 4 | 4 | 4 | 4 | 9 | 9 | 1 |
| yeps_YPTB3839 | 1 | 7 | 1 | 4 | 1 | 3 | 1 | 8 | 7 | 4 | 4 | 1 | 3 | 3 | 3 | 4 | 3 | 4 | 1 | 1 | 3 | 3 | 3 | 1 | 3 | 3 | 3 | 3 | 3 | 3 | 3 | 3 | 4 | 4 | 1 |
| yeps_YPTB3841 | 1 | 9 | 1 | 1 | 1 | 3 | 1 | 10 | 9 | 11 | 1 | 1 | 3 | 3 | 3 | 11 | 3 | 7 | 1 | 1 | 3 | 3 | 3 | 1 | 3 | 3 | 3 | 3 | 3 | 3 | 3 | 3 | 1 | 1 | 1 |
| yeps_YPTB3842 | 4 | 8 | 4 | 8 | 1 | 1 | 4 | 5 | 8 | 9 | 8 | 3 | 4 | 4 | 1 | 9 | 4 | 5 | 4 | 4 | 1 | 1 | 1 | 4 | 4 | 4 | 1 | 1 | 1 | 1 | 1 | 4 | 8 | 8 | 4 |
| yeps_YPTB3846 | 1 | 4 | 6 | 9 | 1 | 1 | 6 | 7 | 4 | 9 | 9 | 4 | 1 | 1 | 1 | 9 | 1 | 5 | 1 | 1 | 1 | 1 | 1 | 1 | 1 | 1 | 1 | 1 | 1 | 1 | 1 | 1 | 9 | 9 | 1 |

| | | | | | | | | | | | | | | | | | | | | | | | | | | | | | | | | | | | |
|---|---|---|---|---|---|---|---|---|---|---|---|---|---|---|---|---|---|---|---|---|---|---|---|---|---|---|---|---|---|---|---|---|---|---|---|
| yeps_YPTB3848 | 1 | 2 | 1 | 4 | 1 | 1 | 1 | 1 | 2 | 4 | 4 | 2 | 1 | 1 | 1 | 4 | 1 | 3 | 1 | 1 | 1 | 1 | 1 | 1 | 1 | 1 | 1 | 1 | 1 | 1 | 1 | 1 | 4 | 4 | 1 |
| yeps_YPTB3850 | 4 | 8 | 4 | 5 | 1 | 1 | 4 | 5 | 8 | 5 | 5 | 3 | 1 | 1 | 1 | 5 | 1 | 5 | 4 | 4 | 1 | 1 | 1 | 4 | 1 | 1 | 1 | 1 | 1 | 1 | 1 | 1 | 5 | 5 | 4 |
| yeps_YPTB3856 | 4 | 3 | 4 | 8 | 1 | 1 | 4 | 4 | 3 | 8 | 8 | 3 | 1 | 1 | 1 | 8 | 1 | 5 | 4 | 4 | 1 | 1 | 1 | 4 | 1 | 1 | 1 | 1 | 1 | 1 | 1 | 1 | 8 | 8 | 4 |
| yeps_YPTB3857 | 3 | 1 | 3 | 6 | 1 | 1 | 3 | 1 | 1 | 6 | 6 | 2 | 1 | 1 | 1 | 6 | 1 | 4 | 3 | 3 | 1 | 1 | 1 | 3 | 1 | 1 | 1 | 1 | 1 | 1 | 1 | 1 | 6 | 6 | 3 |
| yeps_YPTB3858 | 3 | 6 | 3 | 7 | 1 | 1 | 3 | 3 | 6 | 7 | 7 | 3 | 1 | 1 | 1 | 7 | 1 | 1 | 3 | 3 | 1 | 1 | 1 | 3 | 1 | 1 | 1 | 1 | 1 | 1 | 1 | 1 | 7 | 7 | 3 |
| yeps_YPTB3859 | 4 | 7 | 4 | 3 | 1 | 1 | 4 | 4 | 7 | 3 | 3 | 3 | 1 | 1 | 1 | 3 | 1 | 4 | 4 | 4 | 1 | 1 | 1 | 4 | 1 | 1 | 1 | 1 | 1 | 1 | 1 | 1 | 3 | 3 | 4 |
| yeps_YPTB3888 | 3 | 7 | 3 | 10 | 1 | 1 | 3 | 9 | 7 | 10 | 10 | 1 | 1 | 1 | 1 | 10 | 1 | 4 | 3 | 3 | 1 | 1 | 1 | 3 | 1 | 1 | 1 | 1 | 1 | 1 | 1 | 1 | 10 | 10 | 3 |
| yeps_YPTB3904 | 1 | 4 | 1 | 2 | 1 | 1 | 1 | 1 | 4 | 2 | 2 | 1 | 1 | 1 | 1 | 2 | 1 | 2 | 1 | 1 | 1 | 1 | 1 | 1 | 1 | 1 | 1 | 1 | 1 | 1 | 1 | 1 | 2 | 2 | 1 |
| yeps_YPTB3905 | 1 | 2 | 1 | 1 | 1 | 1 | 1 | 1 | 2 | 1 | 1 | 1 | 1 | 1 | 1 | 1 | 1 | 1 | 1 | 1 | 1 | 1 | 1 | 1 | 1 | 1 | 1 | 1 | 1 | 1 | 1 | 1 | 1 | 1 | 1 |
| yeps_YPTB3906 | 4 | 8 | 4 | 12 | 1 | 1 | 4 | 10 | 8 | 12 | 12 | 3 | 1 | 1 | 1 | 12 | 1 | 5 | 4 | 4 | 1 | 1 | 1 | 4 | 1 | 1 | 1 | 1 | 1 | 1 | 1 | 1 | 12 | 12 | 4 |
| yeps_YPTB3910 | 3 | 6 | 3 | 4 | 1 | 1 | 3 | 7 | 6 | 4 | 4 | 1 | 1 | 1 | 1 | 4 | 1 | 4 | 3 | 3 | 1 | 1 | 1 | 3 | 1 | 1 | 1 | 1 | 1 | 1 | 1 | 1 | 4 | 4 | 3 |
| yeps_YPTB3911 | 2 | 2 | 2 | 5 | 1 | 1 | 2 | 5 | 2 | 5 | 5 | 3 | 1 | 1 | 1 | 5 | 1 | 2 | 2 | 2 | 1 | 1 | 1 | 2 | 1 | 1 | 1 | 1 | 1 | 1 | 1 | 1 | 5 | 5 | 2 |
| yeps_YPTB3912 | 3 | 6 | 3 | 3 | 1 | 1 | 3 | 5 | 6 | 3 | 3 | 3 | 1 | 1 | 1 | 3 | 1 | 3 | 3 | 3 | 1 | 1 | 1 | 3 | 1 | 1 | 1 | 1 | 1 | 1 | 1 | 1 | 3 | 3 | 3 |
| yeps_YPTB3913 | 1 | 1 | 1 | 1 | 1 | 1 | 1 | 7 | 1 | 1 | 1 | 3 | 1 | 1 | 1 | 1 | 1 | 4 | 1 | 8 | 1 | 1 | 1 | 8 | 1 | 1 | 1 | 1 | 1 | 1 | 1 | 1 | 1 | 1 | 1 |
| yeps_YPTB3919 | 4 | 8 | 4 | 11 | 1 | 1 | 4 | 10 | 9 | 11 | 11 | 3 | 1 | 1 | 1 | 11 | 1 | 5 | 4 | 4 | 1 | 1 | 1 | 4 | 1 | 1 | 1 | 1 | 1 | 1 | 1 | 1 | 11 | 11 | 4 |
| yeps_YPTB3922 | 1 | 4 | 1 | 2 | 1 | 1 | 1 | 1 | 4 | 2 | 2 | 1 | 1 | 1 | 1 | 2 | 1 | 2 | 1 | 1 | 1 | 1 | 1 | 1 | 1 | 1 | 1 | 1 | 1 | 1 | 1 | 1 | 2 | 2 | 1 |
| yeps_YPTB3925 | 1 | 2 | 1 | 2 | 1 | 1 | 1 | 3 | 2 | 2 | 2 | 1 | 1 | 1 | 1 | 2 | 1 | 4 | 1 | 1 | 1 | 1 | 1 | 1 | 1 | 1 | 1 | 1 | 1 | 1 | 1 | 1 | 2 | 2 | 1 |
| yeps_YPTB3926 | 4 | 7 | 4 | 5 | 1 | 1 | 4 | 8 | 7 | 5 | 5 | 3 | 1 | 1 | 1 | 5 | 1 | 5 | 4 | 4 | 1 | 1 | 1 | 4 | 1 | 1 | 1 | 1 | 1 | 1 | 1 | 1 | 5 | 5 | 4 |
| yeps_YPTB3929 | 1 | 1 | 1 | 3 | 1 | 1 | 1 | 6 | 1 | 3 | 3 | 2 | 1 | 1 | 1 | 3 | 1 | 3 | 1 | 1 | 1 | 1 | 1 | 1 | 1 | 1 | 1 | 1 | 1 | 1 | 1 | 1 | 3 | 3 | 1 |
| yeps_YPTB3930 | 4 | 4 | 4 | 5 | 1 | 1 | 4 | 5 | 4 | 5 | 5 | 3 | 1 | 1 | 1 | 5 | 1 | 5 | 4 | 4 | 1 | 1 | 1 | 4 | 1 | 1 | 1 | 1 | 1 | 1 | 1 | 1 | 5 | 5 | 4 |
| yeps_YPTB3931 | 2 | 3 | 2 | 5 | 1 | 1 | 2 | 4 | 3 | 5 | 5 | 1 | 1 | 1 | 1 | 5 | 1 | 1 | 2 | 2 | 1 | 1 | 1 | 2 | 1 | 1 | 1 | 1 | 1 | 1 | 1 | 1 | 5 | 5 | 2 |
| yeps_YPTB3941 | 3 | 6 | 3 | 8 | 1 | 1 | 3 | 6 | 6 | 7 | 8 | 1 | 1 | 1 | 1 | 7 | 1 | 4 | 3 | 3 | 1 | 1 | 1 | 3 | 1 | 1 | 1 | 1 | 1 | 1 | 1 | 1 | 8 | 8 | 3 |
| yeps_YPTB3943 | 1 | 7 | 1 | 9 | 1 | 1 | 1 | 1 | 7 | 9 | 9 | 1 | 1 | 1 | 1 | 9 | 1 | 4 | 1 | 1 | 1 | 1 | 1 | 1 | 1 | 1 | 1 | 1 | 1 | 1 | 1 | 1 | 9 | 9 | 1 |
| yeps_YPTB3944 | 1 | 1 | 1 | 1 | 1 | 1 | 1 | 1 | 1 | 1 | 1 | 1 | 1 | 1 | 1 | 1 | 1 | 1 | 1 | 1 | 1 | 1 | 1 | 1 | 1 | 1 | 1 | 1 | 1 | 1 | 1 | 1 | 1 | 1 | 1 |
| yeps_YPTB3948 | 1 | 3 | 1 | 6 | 1 | 1 | 1 | 53 | 54 | 6 | 6 | 1 | 1 | 1 | 1 | 6 | 1 | 2 | 1 | 1 | 1 | 1 | 1 | 1 | 1 | 1 | 1 | 1 | 1 | 1 | 1 | 1 | 6 | 6 | 1 |
| yeps_YPTB3949 | 3 | 3 | 3 | 7 | 1 | 1 | 3 | 8 | 3 | 7 | 7 | 1 | 1 | 1 | 1 | 7 | 1 | 4 | 3 | 3 | 1 | 1 | 1 | 3 | 1 | 1 | 1 | 1 | 1 | 1 | 1 | 1 | 7 | 7 | 3 |
| yeps_YPTB3967 | 3 | 7 | 3 | 7 | 1 | 1 | 3 | 7 | 7 | 7 | 7 | 1 | 1 | 1 | 1 | 7 | 1 | 4 | 3 | 3 | 1 | 1 | 1 | 3 | 1 | 1 | 1 | 1 | 1 | 1 | 1 | 1 | 7 | 7 | 3 |
| yeps_YPTB3968 | 2 | 1 | 2 | 1 | 1 | 1 | 2 | 1 | 1 | 1 | 1 | 1 | 1 | 1 | 1 | 1 | 1 | 1 | 2 | 2 | 1 | 1 | 1 | 2 | 1 | 1 | 1 | 1 | 1 | 1 | 1 | 1 | 1 | 1 | 2 |
| yeps_YPTB3972 | 1 | 1 | 1 | 1 | 1 | 1 | 1 | 1 | 1 | 1 | 1 | 1 | 1 | 1 | 1 | 1 | 1 | 1 | 1 | 1 | 1 | 1 | 1 | 1 | 1 | 1 | 1 | 1 | 1 | 1 | 1 | 1 | 1 | 1 | 1 |
| x_1 | 118 | 118 | 118 | 113 | 1 | 1 | 118 | 116 | 118 | 113 | 113 | 113 | 118 | 118 | 1 | 113 | 118 | 115 | 118 | 118 | 1 | 1 | 1 | 118 | 118 | 118 | 1 | 1 | 1 | 1 | 1 | 118 | 113 | 113 | 118 |
| x_2 | 1 | 1 | 1 | 1 | 1 | 1 | 1 | 1 | 1 | 1 | 1 | 1 | 1 | 1 | 1 | 1 | 1 | 1 | 1 | 1 | 1 | 1 | 1 | 1 | 1 | 1 | 1 | 1 | 1 | 1 | 1 | 1 | 1 | 1 | 1 |
| x_3 | 1 | 1 | 1 | 1 | 1 | 1 | 1 | 1 | 1 | 1 | 1 | 1 | 1 | 1 | 1 | 1 | 1 | 1 | 1 | 1 | 1 | 1 | 1 | 1 | 1 | 1 | 1 | 1 | 1 | 1 | 1 | 1 | 1 | 1 | 1 |
| x_4 | 1 | 1 | 1 | 1 | 1 | 1 | 1 | 1 | 1 | 1 | 1 | 1 | 1 | 1 | 1 | 1 | 1 | 1 | 1 | 1 | 1 | 1 | 1 | 1 | 1 | 1 | 1 | 1 | 1 | 1 | 1 | 1 | 1 | 1 | 1 |
| x_5 | 1 | 1 | 1 | 1 | 1 | 1 | 1 | 1 | 1 | 1 | 1 | 1 | 1 | 1 | 1 | 1 | 1 | 1 | 1 | 1 | 1 | 1 | 1 | 1 | 1 | 1 | 1 | 1 | 1 | 1 | 1 | 1 | 1 | 1 | 1 |
| x_6 | 69 | 1 | 69 | 1 | 1 | 1 | 69 | 1 | 1 | 1 | 1 | 1 | 1 | 1 | 1 | 1 | 1 | 70 | 69 | 69 | 1 | 1 | 1 | 69 | 1 | 1 | 1 | 1 | 1 | 1 | 1 | 1 | 1 | 1 | 69 |
| yeps_x_1 | 2 | 1 | 2 | 3 | 1 | 1 | 2 | 1 | 1 | 3 | 3 | 1 | 2 | 2 | 1 | 3 | 2 | 1 | 2 | 2 | 1 | 1 | 1 | 2 | 2 | 2 | 1 | 1 | 1 | 1 | 1 | 2 | 3 | 3 | 2 |
| yeps_x_10 | 1 | 1 | 1 | 1 | 1 | 1 | 1 | 1 | 1 | 1 | 1 | 1 | 1 | 1 | 1 | 1 | 1 | 1 | 1 | 1 | 1 | 1 | 1 | 1 | 1 | 1 | 1 | 1 | 1 | 1 | 1 | 1 | 1 | 1 | 1 |
| yeps_x_11 | 1 | 1 | 1 | 1 | 1 | 1 | 1 | 1 | 1 | 1 | 1 | 1 | 1 | 1 | 1 | 1 | 1 | 1 | 1 | 1 | 1 | 1 | 1 | 1 | 1 | 1 | 1 | 1 | 1 | 1 | 1 | 1 | 1 | 1 | 1 |
| yeps_x_12 | 4 | 28 | 4 | 8 | 1 | 4 | 4 | 4 | 6 | 8 | 8 | 3 | 1 | 1 | 4 | 8 | 1 | 4 | 4 | 4 | 4 | 4 | 4 | 4 | 1 | 1 | 4 | 4 | 4 | 4 | 4 | 1 | 8 | 8 | 4 |
| yeps_x_13 | 1 | 1 | 1 | 3 | 1 | 1 | 1 | 1 | 1 | 2 | 3 | 1 | 1 | 1 | 1 | 2 | 1 | 2 | 1 | 1 | 1 | 1 | 1 | 1 | 1 | 1 | 1 | 1 | 1 | 1 | 1 | 1 | 3 | 3 | 1 |
| yeps_x_14 | 1 | 1 | 1 | 1 | 1 | 1 | 1 | 1 | 1 | 1 | 1 | 1 | 1 | 1 | 1 | 1 | 1 | 2 | 1 | 1 | 1 | 1 | 1 | 1 | 1 | 1 | 1 | 1 | 1 | 1 | 1 | 1 | 1 | 1 | 1 |
| yeps_x_15 | 1 | | 1 | 1 | 1 | 1 | 1 | 1 | 1 | 1 | 1 | 1 | 1 | 1 | 1 | 1 | 1 | 1 | 1 | 1 | 1 | 1 | 1 | 1 | 1 | 1 | 1 | 1 | 1 | 1 | 1 | 1 | 1 | 1 | 1 |
| yeps_x_16 | 1 | 2 | 1 | 17 | 1 | 1 | 1 | 1 | 2 | 1 | 1 | 1 | 1 | 1 | 1 | 1 | 1 | 1 | 1 | 1 | 1 | 1 | 1 | 1 | 1 | 1 | 1 | 1 | 1 | 1 | 1 | 1 | 1 | 1 | 1 |
| yeps_x_17 | 1 | 2 | 1 | 1 | 1 | 1 | 1 | 1 | 2 | 1 | 1 | 1 | 1 | 1 | 1 | 1 | 1 | 1 | 1 | 1 | 1 | 1 | 1 | 1 | 1 | 1 | 1 | 1 | 1 | 1 | 1 | 1 | 1 | 1 | 1 |
| yeps_x_18 | 1 | 1 | 1 | 4 | 1 | 1 | 1 | 1 | 1 | 4 | 4 | 3 | 1 | 1 | 1 | 4 | 1 | 1 | 1 | 1 | 1 | 1 | 1 | 1 | 1 | 1 | 1 | 1 | 1 | 1 | 1 | 1 | 4 | 4 | 1 |
| yeps_x_19 | 1 | 4 | 1 | 1 | 1 | 1 | 1 | 2 | 2 | 1 | 1 | 2 | 1 | 1 | 1 | 1 | 1 | 1 | 1 | 1 | 1 | 1 | 1 | 1 | 1 | 1 | 1 | 1 | 1 | 1 | 1 | 1 | 1 | 1 | 1 |
| yeps_x_2 | 2 | 1 | 2 | 3 | 1 | 1 | 2 | 1 | 1 | 3 | 3 | 1 | 2 | 2 | 1 | 3 | 2 | 2 | 2 | 2 | 1 | 1 | 1 | 2 | 2 | 2 | 1 | 1 | 1 | 1 | 1 | 2 | 3 | 3 | 2 |
| yeps_x_20 | 1 | 1 | 1 | 1 | 1 | 1 | 1 | 1 | 1 | 1 | 1 | 2 | 1 | 1 | 1 | 1 | 1 | 1 | 1 | 1 | 1 | 1 | 1 | 1 | 1 | 1 | 1 | 1 | 1 | 1 | 1 | 1 | 1 | 1 | 1 |
| yeps_x_21 | 1 | 1 | 1 | 1 | 1 | 1 | 1 | 1 | 1 | 1 | 1 | 1 | 1 | 1 | 1 | 1 | 1 | 1 | 1 | 1 | 1 | 1 | 1 | 1 | 1 | 1 | 1 | 1 | 1 | 1 | 1 | 1 | 1 | 1 | 1 |
| yeps_x_22 | 1 | 1 | 1 | 1 | 1 | 1 | 1 | 2 | 1 | 1 | 1 | 1 | 1 | 1 | 1 | 1 | 1 | 1 | 1 | 1 | 1 | 1 | 1 | 1 | 1 | 1 | 1 | 1 | 1 | 1 | 1 | 1 | 1 | 1 | 1 |
| yeps_x_23 | 1 | 1 | 1 | 1 | 1 | 1 | 1 | 1 | 1 | 1 | 1 | 1 | 1 | 1 | 1 | 1 | 1 | 1 | 1 | 1 | 1 | 1 | 1 | 1 | 1 | 1 | 1 | 1 | 1 | 1 | 1 | 1 | 1 | 1 | 1 |
| yeps_x_24 | 1 | 2 | 1 | 1 | 1 | 1 | 1 | 1 | 2 | 1 | 1 | 1 | 1 | 1 | 1 | 1 | 1 | 1 | 1 | 1 | 1 | 1 | 1 | 1 | 1 | 1 | 1 | 1 | 1 | 1 | 1 | 1 | 1 | 1 | 1 |
| yeps_x_25 | 1 | 1 | 1 | 1 | 1 | 1 | 1 | 1 | 1 | 1 | 1 | 1 | 1 | 1 | 1 | 1 | 1 | 1 | 1 | 1 | 1 | 1 | 1 | 1 | 1 | 1 | 1 | 1 | 1 | 1 | 1 | 1 | 1 | 1 | 1 |
| yeps_x_26 | 1 | 3 | 1 | 2 | 1 | 1 | 1 | 4 | 3 | 2 | 2 | 1 | 1 | 1 | 1 | 2 | 1 | 1 | 1 | 1 | 1 | 1 | 1 | 1 | 1 | 1 | 1 | 1 | 1 | 1 | 1 | 1 | 2 | 2 | 1 |
| yeps_x_27 | 1 | 1 | 1 | 1 | 1 | 1 | 1 | 1 | 1 | 1 | 1 | 1 | 1 | 1 | 1 | 1 | 1 | 1 | 1 | 1 | 1 | 1 | 1 | 1 | 1 | 1 | 1 | 1 | 1 | 1 | 1 | 1 | 1 | 1 | 1 |
| yeps_x_28 | 3 | 1 | 3 | 1 | 1 | 1 | 3 | 1 | 1 | 4 | 1 | 2 | 1 | 1 | 1 | 4 | 1 | 1 | 3 | 3 | 1 | 1 | 1 | 3 | 1 | 1 | 1 | 1 | 1 | 1 | 1 | 1 | 1 | 1 | 3 |
| yeps_x_29 | 1 | 1 | 1 | 1 | 1 | 1 | 1 | 1 | 1 | 1 | 1 | 1 | 1 | 1 | 1 | 1 | 1 | 1 | 1 | 1 | 1 | 1 | 1 | 1 | 1 | 1 | 1 | 1 | 1 | 1 | 1 | 1 | 1 | 1 | 1 |
| yeps_x_3 | 1 | 4 | 1 | 4 | 1 | 1 | 1 | 1 | 4 | 4 | 4 | 1 | 1 | 1 | 1 | 4 | 1 | 3 | 1 | 1 | 1 | 1 | 1 | 1 | 1 | 1 | 1 | 1 | 1 | 1 | 1 | 1 | 4 | 4 | 1 |
| yeps_x_30 | 1 | 1 | 1 | 1 | 1 | 1 | 1 | 2 | 1 | 1 | 1 | 1 | 1 | 1 | 1 | 1 | 1 | 1 | 1 | 1 | | | | 1 | 1 | 1 | | | | | | 1 | 1 | 1 | 1 |
| yeps_x_31 | 1 | 1 | 1 | 1 | 1 | 1 | 1 | 1 | 1 | 1 | 1 | 1 | 1 | 1 | 1 | 1 | 1 | 1 | 1 | 1 | 1 | 1 | 1 | 1 | 1 | 1 | 1 | 1 | 1 | 1 | 1 | 1 | 1 | 1 | 1 |
| yeps_x_32 | 1 | 1 | 1 | 1 | 1 | 1 | 1 | 1 | 1 | 1 | 1 | 1 | 1 | 1 | 1 | 1 | 1 | 1 | 1 | 1 | 1 | 1 | 1 | 1 | 1 | 1 | 1 | 1 | 1 | 1 | 1 | 1 | 1 | 1 | 1 |
| yeps_x_33 | 3 | 1 | 1 | 2 | 1 | 1 | 1 | 2 | 1 | 2 | 2 | 1 | 1 | 1 | 1 | 2 | 1 | 2 | 1 | 1 | 1 | 1 | 1 | 1 | 1 | 1 | 1 | 1 | 1 | 1 | 1 | 1 | 2 | 2 | 1 |
| yeps_x_34 | 2 | 2 | 2 | 2 | 1 | 1 | 2 | 2 | 2 | 2 | 2 | 2 | 1 | 1 | 1 | 2 | 1 | 2 | 2 | 2 | 1 | 1 | 1 | 2 | 1 | 1 | 1 | 1 | 1 | 1 | 1 | 1 | 2 | 2 | 2 |
| yeps_x_35 | 1 | 1 | 1 | 1 | 1 | 1 | 1 | 1 | 1 | 1 | 1 | 1 | 1 | 1 | 1 | 1 | 1 | 1 | 1 | 1 | 1 | 1 | 1 | 1 | 1 | 1 | 1 | 1 | 1 | 1 | 1 | 1 | 1 | 1 | 1 |
| yeps_x_36 | 1 | 1 | 1 | 1 | 1 | 2 | 1 | 2 | 1 | 1 | 1 | 1 | 2 | 2 | 2 | 1 | 2 | 1 | 1 | 1 | 2 | 2 | 2 | 1 | 2 | 2 | 2 | 2 | 2 | 2 | 2 | 2 | 1 | 1 | 1 |
| yeps_x_37 | 1 | 1 | 1 | 1 | 1 | 1 | 1 | 1 | 1 | 1 | 1 | 1 | 1 | 1 | 1 | 1 | 1 | 1 | 1 | 1 | 1 | 1 | 1 | 1 | 1 | 1 | 1 | 1 | 1 | 1 | 1 | 1 | 1 | 1 | 1 |
| yeps_x_38 | 1 | 1 | 1 | 1 | 1 | 1 | 1 | 1 | 1 | 1 | 1 | 1 | 1 | 1 | 1 | 1 | 1 | 1 | 1 | 1 | 1 | 1 | 1 | 1 | 1 | 1 | 1 | 1 | 1 | 1 | 1 | 1 | 1 | 1 | 1 |
| yeps_x_39 | 1 | 1 | 1 | 1 | 1 | 1 | 1 | 3 | 1 | 1 | 1 | 1 | 1 | 1 | 1 | 1 | 1 | 3 | 1 | 1 | 1 | 1 | 1 | 1 | 1 | 1 | 1 | 1 | 1 | 1 | 1 | 1 | 1 | 1 | 1 |
| yeps_x_4 | 1 | 1 | 1 | 1 | 1 | 1 | 1 | 1 | 1 | 1 | 1 | 1 | 1 | 1 | 1 | 1 | 1 | 1 | 1 | 1 | 1 | 1 | 1 | 1 | 1 | 1 | 1 | 1 | 1 | 1 | 1 | 1 | 1 | 1 | 1 |
| yeps_x_40 | 1 | 1 | 1 | 2 | 1 | 1 | 1 | 1 | 1 | 2 | 2 | 1 | 1 | 1 | 1 | 2 | 1 | 1 | 1 | 1 | 1 | 1 | 1 | 1 | 1 | 1 | 1 | 1 | 1 | 1 | 1 | 1 | 2 | 2 | 1 |
| yeps_x_41 | 1 | 1 | 1 | 1 | 1 | 1 | 1 | 1 | 1 | 1 | 1 | 1 | 1 | 1 | 1 | 1 | 1 | 1 | 1 | 1 | 1 | 1 | 1 | 1 | 1 | 1 | 1 | 1 | 1 | 1 | 1 | 1 | 1 | 1 | 1 |
| yeps_x_42 | 1 | 2 | 1 | 3 | 1 | 1 | 1 | 5 | 2 | 3 | 3 | 1 | 1 | 1 | 1 | 3 | 1 | 3 | 1 | 1 | 1 | 1 | 1 | 1 | 1 | 1 | 1 | 1 | 1 | 1 | 1 | 1 | 3 | 3 | 1 |
| yeps_x_43 | 1 | 1 | 1 | 1 | 1 | 1 | 1 | 2 | 1 | 1 | 1 | 1 | 1 | 1 | 1 | 1 | 1 | 1 | 1 | 1 | 1 | 1 | 1 | 1 | 1 | 1 | 1 | 1 | 1 | 1 | 1 | 1 | 1 | 1 | 1 |
| yeps_x_44 | 1 | 1 | 1 | 1 | 1 | 1 | 1 | 2 | 1 | 1 | 1 | 1 | 1 | 1 | 1 | 1 | 1 | 3 | 1 | 1 | 1 | 1 | 1 | 1 | 1 | 1 | 1 | 18 | 1 | 14 | 1 | 1 | 1 | 1 | 1 |
| yeps_x_45 | 1 | 6 | 1 | 6 | 1 | 1 | 1 | 6 | 6 | 6 | 6 | 3 | 3 | 3 | 1 | 6 | 3 | 4 | 1 | 1 | 1 | 1 | 1 | 1 | 3 | 3 | 1 | 1 | 1 | 1 | 1 | 3 | 6 | 6 | 1 |
| yeps_x_46 | 2 | 1 | 2 | 2 | 1 | 1 | 2 | 2 | 1 | 2 | 2 | 2 | 2 | 2 | 1 | 2 | 2 | 2 | 2 | 2 | 1 | 1 | 1 | 2 | 2 | 2 | 1 | 1 | 1 | 1 | 1 | 2 | 2 | 2 | 2 |
| yeps_x_47 | 1 | 1 | 1 | 3 | 1 | 1 | 1 | 1 | 1 | 3 | 3 | 1 | 1 | 1 | 1 | 3 | 1 | 2 | 1 | 1 | 1 | 1 | 1 | 1 | 1 | 1 | 1 | 1 | 1 | 1 | 1 | 1 | 3 | 3 | 1 |
| yeps_x_48 | 2 | 3 | 2 | 1 | 1 | 1 | 2 | 1 | 3 | 1 | 1 | 2 | 2 | 2 | 1 | 1 | 2 | 1 | 2 | 2 | 1 | 1 | 1 | 2 | 2 | 2 | 1 | 1 | 1 | 1 | 1 | 2 | 1 | 1 | 2 |
| yeps_x_49 | 1 | 1 | 1 | 1 | 1 | 1 | 1 | 1 | 1 | 1 | 1 | 1 | 1 | 1 | 1 | 1 | 1 | 1 | 1 | 1 | 1 | 1 | 1 | 1 | 1 | 1 | 1 | 1 | 1 | 1 | 1 | 1 | 1 | 1 | 1 |
| yeps_x_5 | 1 | 1 | 1 | 1 | 1 | 1 | 1 | 1 | 1 | 3 | 1 | 1 | 1 | 1 | 1 | 3 | 1 | 1 | 1 | 1 | 1 | 1 | 1 | 1 | 1 | 1 | 1 | 1 | 1 | 1 | 1 | 1 | 1 | 1 | 1 |
| yeps_x_50 | 1 | 1 | 1 | 1 | 1 | 1 | 1 | 3 | 1 | 3 | 1 | 1 | 1 | 1 | 1 | 3 | 1 | 2 | 1 | 1 | 1 | 1 | 1 | 1 | 1 | 1 | 1 | 1 | 1 | 1 | 1 | 1 | 1 | 1 | 1 |
| yeps_x_51 | 2 | 1 | 2 | 1 | 1 | 1 | 2 | 2 | 1 | 1 | 1 | 1 | 1 | 1 | 1 | 1 | 1 | 1 | 2 | 2 | 1 | 1 | 1 | 2 | 1 | 1 | 1 | 1 | 1 | 1 | 1 | 1 | 1 | 1 | 2 |
| yeps_x_52 | 1 | 2 | 1 | 1 | 1 | 22 | 1 | 3 | 2 | 1 | 1 | 1 | 2 | 2 | 22 | 1 | 2 | 1 | 1 | 1 | 22 | 22 | 22 | 1 | 2 | 2 | 22 | 22 | 22 | 22 | 22 | 2 | 1 | 1 | 1 |
| yeps_x_53 | 1 | 3 | 1 | 1 | 1 | 1 | 1 | 1 | 1 | 1 | 1 | 1 | 1 | 1 | 1 | 1 | 1 | 1 | 1 | 1 | 1 | 1 | 1 | 1 | 1 | 1 | 1 | 1 | 1 | 1 | 1 | 1 | 1 | 1 | 1 |
| yeps_x_54 | 1 | 1 | 1 | 1 | 1 | 1 | 1 | 1 | 1 | 1 | 1 | 1 | 1 | 1 | 1 | 1 | 1 | 1 | 1 | 1 | 1 | 1 | 1 | 1 | 1 | 1 | 1 | 1 | 1 | 1 | 1 | 1 | 1 | 1 | 1 |
| yeps_x_55 | 1 | 1 | 1 | 1 | 1 | 1 | 1 | 1 | 1 | 1 | 1 | 2 | 1 | 1 | 1 | 1 | 1 | 1 | 1 | 1 | 1 | 1 | 1 | 1 | 1 | 1 | 1 | 1 | 1 | 1 | 1 | 1 | 1 | 1 | 1 |
| yeps_x_56 | 1 | 1 | 1 | 1 | 1 | 1 | 1 | 1 | 1 | 1 | 1 | 1 | 1 | 1 | 1 | 1 | 1 | 1 | 1 | 1 | 1 | 1 | 1 | 1 | 1 | 1 | 1 | 1 | 1 | 1 | 1 | 1 | 1 | 1 | 1 |
| yeps_x_57 | 1 | 1 | 1 | 1 | 1 | 1 | 1 | 1 | 1 | 1 | 1 | 1 | 1 | 1 | 1 | 1 | 1 | 1 | 1 | 1 | 1 | 1 | 1 | 1 | 1 | 1 | 1 | 1 | 1 | 1 | 1 | 1 | 1 | 1 | 1 |
| yeps_x_58 | 1 | 1 | 1 | 1 | 1 | 1 | 1 | 1 | 1 | 1 | 1 | 1 | 1 | 1 | 1 | 1 | 1 | 1 | 1 | 1 | 1 | 1 | 1 | 1 | 1 | 1 | 1 | 1 | 1 | 1 | 1 | 1 | 1 | 1 | 1 |
| yeps_x_59 | 1 | 1 | 1 | 2 | 1 | 2 | 1 | 1 | 1 | 2 | 2 | 2 | 2 | 2 | 2 | 2 | 2 | 2 | 1 | 1 | 2 | 2 | 2 | 1 | 2 | 2 | 2 | 2 | 2 | 2 | 2 | 2 | 2 | 2 | 1 |
| yeps_x_6 | 1 | 1 | 1 | 1 | 1 | 1 | 1 | 1 | 1 | 1 | 1 | 1 | 1 | 1 | 1 | 1 | 1 | 1 | 1 | 1 | 1 | 1 | 1 | 1 | 1 | 1 | 1 | 1 | 1 | 1 | 1 | 1 | 1 | 1 | 1 |
| yeps_x_60 | 1 | 1 | 1 | 1 | 1 | 1 | 1 | 1 | 1 | 1 | 1 | 1 | 1 | 1 | 1 | 1 | 1 | 1 | 1 | 1 | 1 | 1 | 1 | 1 | 1 | 1 | 1 | 1 | 1 | 1 | 1 | 1 | 1 | 1 | 1 |
| yeps_x_61 | 1 | 1 | 1 | 1 | 1 | 1 | 1 | | 1 | 1 | 1 | 1 | 1 | 1 | 1 | 1 | 1 | 1 | 1 | 1 | 1 | 1 | 1 | 1 | 1 | 1 | 1 | 1 | 1 | 1 | 1 | 1 | 1 | 1 | 1 |
| yeps_x_62 | 4 | 8 | 4 | 3 | 1 | 1 | 4 | 10 | 8 | 3 | 3 | 3 | 1 | 1 | 1 | 3 | 1 | 5 | 4 | 4 | 1 | 1 | 1 | 4 | 1 | 1 | 1 | 1 | 1 | 1 | 1 | 1 | 3 | 3 | 4 |
| yeps_x_63 | 1 | 1 | 1 | 1 | 1 | 1 | 1 | 1 | 1 | 1 | 1 | 1 | 1 | 1 | 1 | 1 | 1 | 1 | 1 | 1 | 1 | 1 | 1 | 1 | 1 | 1 | 1 | 1 | 1 | 1 | 1 | 1 | 1 | 1 | 1 |
| yeps_x_64 | 11 | 9 | 11 | 15 | 1 | 1 | 11 | 8 | 9 | 14 | 15 | 3 | 1 | 1 | 1 | 14 | 1 | 7 | 4 | 4 | 1 | 1 | 1 | 4 | 1 | 1 | 1 | 1 | 1 | 1 | 1 | 1 | 15 | 15 | 11 |
| yeps_x_65 | 1 | 1 | 1 | 2 | 1 | 1 | 1 | 1 | 1 | 2 | 2 | 1 | 1 | 1 | 1 | 2 | 1 | 2 | 1 | 1 | 1 | 1 | 1 | 1 | 1 | 1 | 1 | 1 | 1 | 1 | 1 | 1 | 2 | 2 | 1 |
| yeps_x_66 | 1 | 1 | 1 | 1 | 1 | 1 | 1 | 1 | 1 | 1 | 1 | 1 | 1 | 1 | 1 | 1 | 1 | 1 | 1 | 1 | 1 | 1 | 1 | 1 | 1 | 1 | 1 | 1 | 1 | 1 | 1 | 1 | 1 | 1 | 1 |
| yeps_x_7 | 1 | 1 | 1 | 1 | 1 | 1 | 1 | 1 | 1 | 1 | 1 | 1 | 1 | 1 | 1 | 1 | 1 | 1 | 1 | 1 | 1 | 1 | 1 | 1 | 1 | 1 | 1 | 1 | 1 | 1 | 1 | 1 | 1 | 1 | 1 |
| yeps_x_8 | 1 | 2 | 1 | 1 | 1 | 1 | 1 | 1 | 2 | 1 | 1 | 1 | 1 | 1 | 1 | 1 | 1 | 1 | 1 | 1 | 1 | 1 | 1 | 1 | 1 | 1 | 1 | 1 | 1 | 1 | 1 | 1 | 1 | 1 | 1 |
| yeps_x_9 | 1 | 1 | 1 | 1 | 1 | 1 | 1 | 1 | 1 | 1 | 1 | 1 | 1 | 1 | 1 | 1 | 1 | 2 | 1 | 1 | 1 | 1 | 1 | 1 | 1 | 1 | 1 | 1 | 1 | 1 | 1 | 1 | 1 | 1 | 1 |
| YPTB0002 | 1 | 1 | 1 | 1 | 1 | 1 | 1 | 72 | 1 | 1 | 1 | 1 | 1 | 1 | 1 | 1 | 1 | 1 | 1 | 1 | 1 | 1 | 1 | 1 | 1 | 1 | 1 | 1 | 1 | 1 | 1 | 1 | 1 | 1 | 1 |
| YPTB0003 | 1 | 1 | 1 | 118 | 1 | 1 | 1 | 1 | 1 | 1 | 118 | 119 | 1 | 1 | 1 | 1 | 1 | 91 | 1 | 1 | 1 | 1 | 1 | 1 | 1 | 1 | 1 | 1 | 1 | 1 | 1 | 1 | 118 | 118 | 1 |
| YPTB0028 | 79 | 79 | 79 | 81 | 1 | 1 | 79 | 79 | 79 | 81 | 81 | 1 | 79 | 79 | 1 | 81 | 79 | 79 | 79 | 79 | 1 | 1 | 1 | 79 | 79 | 79 | 1 | 1 | 1 | 1 | 1 | 79 | 81 | 81 | 79 |
| YPTB0033 | 132 | 135 | 132 | 180 | 1 | 1 | 132 | 136 | 135 | 137 | 180 | 1 | 132 | 132 | 1 | 137 | 132 | 132 | 132 | 132 | 1 | 1 | 1 | 132 | 132 | 132 | 1 | 1 | 1 | 1 | 1 | 132 | 180 | 180 | 132 |
| YPTB0037 | 73 | 73 | 73 | 73 | 1 | 1 | 73 | 73 | 73 | 73 | 73 | 74 | 73 | 73 | 1 | 73 | 73 | 73 | 73 | 73 | 1 | 1 | 1 | 73 | 73 | 73 | 1 | 1 | 1 | 1 | 1 | 73 | 73 | 73 | 73 |
| YPTB0044 | 70 | 70 | 70 | 70 | 1 | 1 | 70 | 70 | 70 | 70 | 70 | 70 | 1 | 1 | 1 | 70 | 1 | 70 | 70 | 70 | 1 | 1 | 1 | 70 | 1 | 1 | 1 | 1 | 1 | 1 | 1 | 1 | 70 | 70 | 70 |
| YPTB0047 | 1 | 1 | 1 | 1 | 1 | 1 | 1 | 1 | 1 | 1 | 1 | 1 | 1 | 1 | 1 | 1 | 1 | 1 | 1 | 1 | 1 | 1 | 1 | 1 | 1 | 1 | 1 | 1 | 1 | 1 | 1 | 1 | 1 | 1 | 1 |
| YPTB0055 | 1 | 1 | 1 | 1 | 1 | 1 | 1 | 1 | 1 | 1 | 1 | 1 | 1 | 1 | 1 | 1 | 1 | 1 | 1 | 1 | 1 | 1 | 1 | 1 | 1 | 1 | 1 | 1 | 1 | 1 | 1 | 1 | 1 | 1 | 1 |
| YPTB0070 | 1 | 1 | 1 | 1 | 1 | 1 | 1 | 1 | 1 | 1 | 1 | 87 | 1 | 1 | 1 | 1 | 1 | 1 | 1 | 1 | 1 | 1 | 1 | 1 | 1 | 1 | 1 | 1 | 1 | 1 | 1 | 1 | 1 | 1 | 1 |
| YPTB0086 | 1 | 126 | 1 | 1 | 1 | 1 | 1 | 126 | 126 | 1 | 1 | 1 | 1 | 1 | 1 | 1 | 1 | 1 | 1 | 1 | 1 | 1 | 1 | 1 | 1 | 1 | 1 | 1 | 1 | 1 | 1 | 1 | 1 | 1 | 1 |
| YPTB0089 | 1 | 1 | 1 | 1 | 1 | 1 | 1 | 1 | 1 | 1 | 1 | 1 | 1 | 1 | 1 | 1 | 1 | 1 | 1 | 1 | 1 | 1 | 1 | 1 | 1 | 1 | 1 | 1 | 1 | 1 | 1 | 1 | 1 | 1 | 1 |
| YPTB0097 | 1 | 110 | 1 | 1 | 1 | 1 | 1 | 110 | 110 | 1 | 1 | 108 | 1 | 1 | 1 | 1 | 1 | 1 | 1 | 1 | 1 | 1 | 1 | 1 | 1 | 1 | 1 | 1 | 1 | 1 | 1 | 1 | 1 | 1 | 1 |
| YPTB0101 | 126 | 129 | 126 | 126 | 1 | 1 | 184 | 131 | 129 | 126 | 126 | 126 | 1 | 1 | 1 | 126 | 1 | 127 | 126 | 126 | 1 | 1 | 1 | 126 | 1 | 1 | 1 | 1 | 1 | 1 | 1 | 1 | 126 | 126 | 126 |
| YPTB0104 | 1 | 39 | 1 | 1 | 1 | 1 | 1 | 39 | 39 | 1 | 1 | 1 | 1 | 1 | 1 | 1 | 1 | 1 | 1 | 1 | 1 | 1 | 1 | 1 | 1 | 1 | 1 | 1 | 1 | 1 | 1 | 1 | 1 | 1 | 1 |
| YPTB0121 | 111 | 1 | 111 | 110 | 1 | 1 | 111 | 114 | 1 | 110 | 110 | 111 | 1 | 1 | 1 | 110 | 1 | 1 | 111 | 111 | 1 | 1 | 1 | 111 | 1 | 1 | 1 | 1 | 1 | 1 | 1 | 1 | 110 | 110 | 111 |
| YPTB0132 | 1 | 1 | 1 | 1 | 1 | 1 | 1 | 1 | 1 | 1 | 1 | 1 | 1 | 1 | 1 | 1 | 1 | 1 | 1 | 1 | 1 | 1 | 1 | 1 | 1 | 1 | 1 | 1 | 1 | 1 | 1 | 1 | 1 | 1 | 1 |
| YPTB0134 | 1 | 124 | 1 | 121 | 1 | 1 | 1 | 123 | 124 | 121 | 121 | 169 | 1 | 1 | 1 | 121 | 1 | 118 | 1 | 1 | 1 | 1 | 1 | 1 | 1 | 1 | 1 | 1 | 1 | 1 | 1 | 1 | 121 | 121 | 1 |
| YPTB0137 | 127 | 131 | 127 | 132 | 1 | 1 | 127 | 134 | 131 | 132 | 132 | 127 | 127 | 127 | 1 | 132 | 127 | 1 | 127 | 127 | 1 | 1 | 1 | 127 | 127 | 127 | 1 | 1 | 1 | 1 | 1 | 127 | 132 | 132 | 127 |
| YPTB0166 | 1 | 1 | 1 | 47 | 1 | 1 | 1 | 1 | 1 | 47 | 47 | 1 | 1 | 1 | 1 | 47 | 1 | 1 | 1 | 1 | 1 | 1 | 1 | 1 | 1 | 1 | 1 | 1 | 1 | 1 | 1 | 1 | 47 | 47 | 1 |
| YPTB0167 | 83 | 1 | 83 | 1 | 1 | 83 | 83 | 1 | 1 | 1 | 1 | 83 | 1 | 1 | 83 | 1 | 1 | 83 | 83 | 83 | 83 | 83 | 83 | 83 | 1 | 1 | 83 | 83 | 83 | 83 | 83 | 1 | 1 | 1 | 83 |
| YPTB0171 | 112 | 112 | 112 | 112 | 1 | 1 | 112 | 112 | 112 | 112 | 112 | 1 | 1 | 1 | 1 | 112 | 1 | 112 | 112 | 112 | 1 | 1 | 1 | 112 | 1 | 1 | 1 | 1 | 1 | 1 | 1 | 1 | 112 | 112 | 112 |
| YPTB0177 | 1 | 112 | 1 | 110 | 1 | 1 | 1 | 114 | 112 | 110 | 110 | 1 | 1 | 1 | 1 | 110 | 1 | 1 | 1 | 1 | 1 | 1 | 1 | 1 | 1 | 1 | 1 | 1 | 1 | 1 | 1 | 1 | 110 | 110 | 1 |
| YPTB0182 | 1 | 107 | 1 | 105 | 1 | 1 | 1 | 108 | 107 | 105 | 105 | 106 | 1 | 1 | 1 | 105 | 1 | 105 | 1 | 1 | 1 | 1 | 1 | 1 | 1 | 1 | 1 | 1 | 1 | 1 | 1 | 1 | 105 | 105 | 1 |
| YPTB0190 | 85 | 89 | 85 | 90 | 1 | 1 | 85 | 88 | 89 | 90 | 90 | 1 | 1 | 1 | 1 | 90 | 1 | 86 | 85 | 85 | 1 | 1 | 1 | 85 | 1 | 1 | 1 | 1 | 1 | 1 | 1 | 1 | 90 | 90 | 85 |
| YPTB0193 | 1 | 89 | 1 | 112 | 1 | 1 | 1 | 1 | 111 | 92 | 112 | 1 | 1 | 1 | 1 | 92 | 1 | 90 | 1 | 1 | 1 | 1 | 1 | 1 | 1 | 1 | 1 | 1 | 1 | 1 | 1 | 1 | 112 | 112 | 1 |
| YPTB0219 | 1 | 1 | 1 | 60 | 1 | 1 | 1 | 1 | 1 | 60 | 60 | 1 | 1 | 1 | 1 | 60 | 1 | 1 | 1 | 1 | 1 | 1 | 1 | 1 | 1 | 1 | 1 | 1 | 1 | 1 | 1 | 1 | 60 | 60 | 1 |
| YPTB0222 | 75 | 1 | 75 | 77 | 1 | 1 | 75 | 79 | 1 | 77 | 77 | 1 | 1 | 1 | 1 | 77 | 1 | 75 | 75 | 75 | 1 | 1 | 1 | 75 | 1 | 1 | 1 | 1 | 1 | 1 | 1 | 1 | 77 | 77 | 75 |
| YPTB0224 | 75 | 75 | 75 | 78 | 1 | 1 | 75 | 80 | 75 | 78 | 78 | 1 | 1 | 1 | 1 | 78 | 1 | 75 | 75 | 75 | 1 | 1 | 1 | 75 | 1 | 1 | 1 | 1 | 1 | 1 | 1 | 1 | 78 | 78 | 75 |
| YPTB0227 | 102 | 105 | 102 | 106 | 1 | 1 | 102 | 102 | 105 | 106 | 106 | 101 | 1 | 1 | 1 | 106 | 1 | 102 | 102 | 102 | 1 | 1 | 1 | 102 | 1 | 1 | 1 | 1 | 1 | 1 | 1 | 1 | 106 | 106 | 102 |
| YPTB0228 | 114 | 117 | 114 | 119 | 1 | 1 | 114 | 118 | 117 | 119 | 119 | 1 | 1 | 1 | 1 | 119 | 1 | 114 | 114 | 114 | 1 | 1 | 1 | 114 | 1 | 1 | 1 | 1 | 1 | 1 | 1 | 1 | 119 | 119 | 114 |
| YPTB0229 | 94 | 98 | 94 | 98 | 1 | 1 | 94 | 99 | 98 | 98 | 98 | 1 | 1 | 1 | 1 | 98 | 1 | 95 | 94 | 94 | 1 | 1 | 1 | 94 | 1 | 1 | 1 | 1 | 1 | 1 | 1 | 1 | 98 | 98 | 94 |
| YPTB0240 | 1 | 1 | 1 | 1 | 1 | 1 | 1 | 1 | 1 | 1 | 1 | 1 | 1 | 1 | 1 | 1 | 1 | 1 | 1 | 1 | 1 | 1 | 1 | 1 | 1 | 1 | 1 | 1 | 1 | 1 | 1 | 1 | 1 | 1 | 1 |
| YPTB0247 | 95 | 98 | 95 | 1 | 1 | 1 | 95 | 102 | 292 | 100 | 1 | 94 | 1 | 1 | 1 | 100 | 1 | 96 | 95 | 95 | 1 | 1 | 1 | 95 | 1 | 1 | 1 | 1 | 1 | 1 | 1 | 1 | 1 | 1 | 95 |
| YPTB0250 | 1 | 1 | 1 | 1 | 1 | 1 | 1 | 1 | 1 | 1 | 1 | 1 | 1 | 1 | 1 | 1 | 1 | 1 | 1 | 1 | 1 | 1 | 1 | 1 | 1 | 1 | 1 | 1 | 1 | 1 | 1 | 1 | 1 | 1 | 1 |
| YPTB0261 | 102 | 105 | 102 | 102 | 1 | 1 | 102 | 105 | 105 | 102 | 102 | 1 | 1 | 1 | 1 | 102 | 1 | 102 | 102 | 102 | 1 | 1 | 1 | 102 | 1 | 1 | 1 | 1 | 1 | 1 | 1 | 1 | 102 | 102 | 102 |
| YPTB0262 | 101 | 101 | 101 | 101 | 1 | 1 | 101 | 101 | 101 | 101 | 101 | 1 | 1 | 1 | 1 | 101 | 1 | 1 | 101 | 101 | 1 | 1 | 1 | 101 | 1 | 1 | 1 | 1 | 1 | 1 | 1 | 1 | 101 | 101 | 101 |
| YPTB0264 | 105 | 1 | 105 | 1 | 1 | 1 | 105 | 104 | 1 | 1 | 1 | 1 | 1 | 1 | 1 | 1 | 1 | 106 | 105 | 105 | 1 | 1 | 1 | 105 | 1 | 1 | 1 | 1 | 1 | 1 | 1 | 1 | 1 | 1 | 105 |
| YPTB0265 | 83 | 83 | 83 | 1 | 1 | 1 | 83 | 83 | 83 | 1 | 1 | 1 | 1 | 1 | 1 | 1 | 1 | 83 | 83 | 83 | 1 | 1 | 1 | 83 | 1 | 1 | 1 | 1 | 1 | 1 | 1 | 1 | 1 | 1 | 83 |
| YPTB0266 | 114 | 115 | 114 | 1 | 1 | 1 | 114 | 114 | 115 | 1 | 1 | 1 | 1 | 1 | 1 | 1 | 1 | 114 | 114 | 114 | 1 | 1 | 1 | 114 | 1 | 1 | 1 | 1 | 1 | 1 | 1 | 1 | 1 | 1 | 114 |
| YPTB0267 | 135 | 491 | 135 | 135 | 1 | 1 | 135 | 133 | 138 | 139 | 135 | 134 | 1 | 1 | 1 | 139 | 1 | 136 | 135 | 135 | 1 | 1 | 1 | 135 | 1 | 1 | 1 | 1 | 1 | 1 | 1 | 1 | 135 | 135 | 135 |
| YPTB0270 | 272 | 91 | 91 | 91 | 1 | 1 | 91 | 91 | 91 | 91 | 91 | 91 | 1 | 1 | 1 | 91 | 1 | 91 | 91 | 91 | 1 | 1 | 1 | 91 | 1 | 1 | 1 | 1 | 1 | 1 | 1 | 1 | 91 | 91 | 91 |
| YPTB0277 | 1 | 1 | 1 | 1 | 1 | 1 | 1 | 1 | 1 | 1 | 1 | 1 | 1 | 1 | 1 | 1 | 1 | 1 | 1 | 1 | 1 | 1 | 1 | 1 | 1 | 1 | 1 | 1 | 1 | 1 | 1 | 1 | 1 | 1 | 1 |
| YPTB0278 | 1 | 1 | 1 | 1 | 1 | 1 | 1 | 1 | 1 | 1 | 1 | 1 | 1 | 67 | 1 | 1 | 67 | 1 | 1 | 1 | 1 | 1 | 1 | 1 | 1 | 1 | 1 | 1 | 1 | 1 | 1 | 1 | 1 | 1 | 1 |
| YPTB0279 | 1 | 1 | 1 | 1 | 1 | 1 | 1 | 1 | 1 | 1 | 1 | 1 | 1 | 1 | 1 | 1 | 1 | 1 | 1 | 1 | 1 | 1 | 1 | 1 | 1 | 1 | 1 | 1 | 1 | 1 | 1 | 1 | 1 | 1 | 1 |
| YPTB0280 | 1 | 1 | 1 | 1 | 1 | 1 | 1 | 1 | 1 | 1 | 1 | 1 | 1 | 1 | 1 | 1 | 1 | 1 | 1 | 1 | 1 | 1 | 1 | 1 | 1 | 1 | 1 | 1 | 1 | 1 | 1 | 1 | 1 | 1 | 1 |
| YPTB0282 | 38 | 38 | 38 | 38 | 1 | 1 | 38 | 1 | 38 | 38 | 38 | 1 | 1 | 1 | 1 | 38 | 1 | 38 | 38 | 38 | 1 | 1 | 1 | 38 | 1 | 1 | 1 | 1 | 1 | 1 | 1 | 1 | 38 | 38 | 38 |
| YPTB0296 | 1 | 1 | 1 | 1 | 1 | 1 | 1 | 1 | 1 | 1 | 1 | 1 | 1 | 1 | 1 | 1 | 1 | 1 | 1 | 1 | 1 | 1 | 1 | 1 | 1 | 1 | 1 | 1 | 1 | 1 | 1 | 1 | 1 | 1 | 1 |
| YPTB0337 | 127 | 1 | 1 | 1 | 1 | 1 | 1 | 1 | 1 | 1 | 1 | 1 | 1 | 1 | 1 | 1 | 1 | 1 | 1 | 1 | 1 | 1 | 1 | 1 | 1 | 1 | 1 | 1 | 1 | 1 | 1 | 1 | 1 | 1 | 1 |
| YPTB0370 | 73 | 74 | 73 | 73 | 1 | 1 | 73 | 75 | 74 | 73 | 73 | 73 | 1 | 1 | 1 | 73 | 1 | 73 | 73 | 73 | 1 | 1 | 1 | 73 | 1 | 1 | 1 | 1 | 1 | 1 | 1 | 1 | 73 | 73 | 73 |
| YPTB0373 | 1 | 1 | 1 | 1 | 1 | 1 | 1 | 1 | 1 | 1 | 1 | 53 | 1 | 1 | 1 | 1 | 1 | 1 | 1 | 1 | 1 | 1 | 1 | 1 | 1 | 1 | 1 | 1 | 1 | 1 | 1 | 1 | 1 | 1 | 1 |
| YPTB0375 | 120 | 123 | 120 | 124 | 1 | 1 | 120 | 126 | 123 | 124 | 124 | 120 | 1 | 1 | 1 | 124 | 1 | 121 | 120 | 120 | 1 | 1 | 1 | 120 | 1 | 1 | 1 | 1 | 1 | 1 | 1 | 1 | 124 | 124 | 120 |
| YPTB0376 | 121 | 123 | 121 | 122 | 1 | 1 | 121 | 126 | 123 | 122 | 122 | 120 | 1 | 1 | 1 | 122 | 1 | 122 | 121 | 121 | 1 | 1 | 1 | 121 | 1 | 1 | 1 | 1 | 1 | 1 | 1 | 1 | 122 | 122 | 121 |
| YPTB0377 | 129 | 130 | 129 | 128 | 1 | 1 | 129 | 128 | 130 | 128 | 128 | 125 | 161 | 1 | 1 | 128 | 1 | 127 | 129 | 129 | 1 | 1 | 1 | 129 | 1 | 1 | 1 | 1 | 1 | 1 | 1 | 1 | 128 | 128 | 129 |
| YPTB0402 | 103 | 106 | 103 | 107 | 1 | 1 | 103 | 109 | 106 | 107 | 107 | 1 | 1 | 1 | 1 | 107 | 1 | 104 | 103 | 103 | 1 | 1 | 1 | 103 | 1 | 1 | 1 | 1 | 1 | 1 | 1 | 1 | 107 | 107 | 103 |
| YPTB0404 | 1 | 31 | 1 | 1 | 1 | 1 | 1 | 1 | 31 | 1 | 1 | 1 | 1 | 1 | 1 | 1 | 1 | 1 | 1 | 1 | 1 | 1 | 1 | 1 | 1 | 1 | 1 | 1 | 1 | 1 | 1 | 1 | 1 | 1 | 1 |
| YPTB0408 | 1 | 75 | 1 | 1 | 1 | 1 | 1 | 78 | 75 | 1 | 1 | 1 | 1 | 1 | 1 | 1 | 1 | 75 | 1 | 1 | 1 | 1 | 1 | 1 | 1 | 1 | 1 | 1 | 1 | 1 | 1 | 1 | 1 | 1 | 1 |
| YPTB0411 | 1 | 1 | 1 | 1 | 1 | 1 | 1 | 1 | 1 | 1 | 1 | 1 | 1 | 1 | 1 | 1 | 1 | 1 | 1 | 1 | 1 | 1 | 1 | 1 | 1 | 1 | 1 | 1 | 1 | 1 | 1 | 1 | 1 | 1 | 1 |
| YPTB0412 | 83 | 86 | 83 | 1 | 1 | 1 | 83 | 83 | 86 | 1 | 1 | 1 | 1 | 1 | 1 | 1 | 1 | 84 | 83 | 83 | 1 | 1 | 1 | 83 | 1 | 1 | 1 | 1 | 1 | 1 | 1 | 1 | 1 | 1 | 83 |
| YPTB0424 | 101 | 101 | 101 | 101 | 1 | 1 | 101 | 1 | 101 | 101 | 101 | 1 | 1 | 1 | 1 | 101 | 1 | 101 | 101 | 101 | 1 | 1 | 1 | 101 | 1 | 1 | 1 | 1 | 1 | 1 | 1 | 1 | 101 | 101 | 101 |
| YPTB0425 | 1 | 1 | 1 | 1 | 1 | 1 | 1 | 1 | 1 | 1 | 1 | 1 | 1 | 1 | 1 | 1 | 1 | 1 | 1 | 1 | 1 | 1 | 1 | 1 | 1 | 1 | 1 | 1 | 1 | 1 | 1 | 1 | 1 | 1 | 1 |
| YPTB0438 | 1 | 1 | 1 | 1 | 1 | 1 | 1 | 1 | 1 | 1 | 1 | 1 | 1 | 1 | 1 | 1 | 1 | 1 | 1 | 1 | 1 | 1 | 1 | 1 | 1 | 1 | 1 | 1 | 1 | 1 | 1 | 1 | 1 | 1 | 1 |
| YPTB0439 | 1 | 1 | 1 | 1 | 1 | 1 | 1 | 1 | 1 | 1 | 1 | 1 | 1 | 1 | 1 | 1 | 1 | 1 | 1 | 1 | 1 | 1 | 1 | 1 | 1 | 1 | 1 | 1 | 1 | 1 | 1 | 1 | 1 | 1 | 1 |
| YPTB0440 | 1 | 1 | 1 | 1 | 1 | 1 | 1 | 1 | 1 | 1 | 1 | 1 | 1 | 1 | 1 | 1 | 1 | 1 | 1 | 1 | 1 | 1 | 1 | 1 | 1 | 1 | 1 | 1 | 1 | 1 | 1 | 1 | 1 | 1 | 1 |
| YPTB0441 | 54 | 54 | 54 | 54 | 1 | 1 | 54 | 54 | 54 | 54 | 54 | 1 | 1 | 1 | 1 | 54 | 1 | 54 | 54 | 54 | 1 | 1 | 1 | 54 | 1 | 1 | 1 | 1 | 1 | 1 | 1 | 1 | 54 | 54 | 54 |
| YPTB0444 | 1 | 79 | 1 | 80 | 1 | 1 | 1 | 78 | 78 | 80 | 80 | 1 | 1 | 1 | 1 | 80 | 1 | 78 | 1 | 1 | 1 | 1 | 1 | 1 | 1 | 1 | 1 | 1 | 1 | 1 | 1 | 1 | 80 | 80 | 1 |

| | | | | | | | | | | | | | | | | | | | | | | | | | | | | | | | | | | | |
|---|---|---|---|---|---|---|---|---|---|---|---|---|---|---|---|---|---|---|---|---|---|---|---|---|---|---|---|---|---|---|---|---|---|---|---|
| YPTB0450 | 1 | 115 | 1 | 120 | 1 | 1 | 1 | 118 | 115 | 1 | 120 | 112 | 118 | 118 | 1 | 1 | 118 | 1 | 1 | 1 | 1 | 1 | 1 | 1 | 118 | 118 | 1 | 1 | 1 | 1 | 1 | 118 | 120 | 120 | 1 |
| YPTB0463 | 105 | 104 | 105 | 1 | 1 | 1 | 105 | 1 | 104 | 1 | 1 | 1 | 1 | 1 | 1 | 1 | 1 | 1 | 105 | 105 | 1 | 1 | 1 | 105 | 1 | 1 | 1 | 1 | 1 | 1 | 1 | 1 | 1 | 1 | 105 |
| YPTB0464 | 1 | 1 | 1 | 1 | 1 | 1 | 1 | 1 | 1 | 1 | 1 | 1 | 1 | 1 | 1 | 1 | 1 | 1 | 1 | 1 | 1 | 1 | 1 | 1 | 1 | 1 | 1 | 1 | 1 | 1 | 1 | 1 | 1 | 1 | 1 |
| YPTB0465 | 1 | 1 | 1 | 1 | 1 | 1 | 1 | 1 | 1 | 1 | 1 | 1 | 1 | 1 | 1 | 1 | 1 | 1 | 1 | 1 | 1 | 1 | 1 | 1 | 1 | 1 | 1 | 1 | 1 | 1 | 1 | 1 | 1 | 1 | 1 |
| YPTB0470 | 119 | 118 | 119 | 119 | 1 | 1 | 119 | 118 | 118 | 119 | 119 | 1 | 1 | 1 | 1 | 119 | 1 | 119 | 119 | 119 | 1 | 1 | 1 | 119 | 1 | 1 | 1 | 1 | 1 | 1 | 1 | 1 | 119 | 119 | 119 |
| YPTB0471 | 1 | 1 | 1 | 1 | 1 | 1 | 1 | 1 | 1 | 1 | 1 | 1 | 1 | 1 | 1 | 1 | 1 | 1 | 1 | 1 | 1 | 1 | 1 | 1 | 1 | 1 | 1 | 1 | 1 | 1 | 1 | 1 | 1 | 1 | 1 |
| YPTB0472 | 1 | 47 | 1 | 47 | 1 | 1 | 1 | 1 | 47 | 47 | 47 | 1 | 1 | 1 | 1 | 47 | 1 | 47 | 1 | 1 | 1 | 1 | 1 | 1 | 1 | 1 | 1 | 1 | 1 | 1 | 1 | 1 | 47 | 47 | 1 |
| YPTB0473 | 40 | 1 | 40 | 41 | 1 | 1 | 40 | 40 | 1 | 41 | 41 | 1 | 1 | 1 | 1 | 41 | 1 | 41 | 40 | 40 | 1 | 1 | 1 | 40 | 1 | 1 | 1 | 1 | 1 | 1 | 1 | 1 | 41 | 41 | 40 |
| YPTB0475 | 89 | 1 | 89 | 1 | 1 | 1 | 89 | 91 | 1 | 1 | 1 | 1 | 1 | 1 | 1 | 1 | 1 | 90 | 89 | 89 | 1 | 1 | 1 | 89 | 1 | 1 | 1 | 1 | 1 | 1 | 1 | 1 | 1 | 1 | 89 |
| YPTB0476 | 102 | 103 | 102 | 101 | 1 | 1 | 102 | 100 | 103 | 101 | 101 | 101 | 1 | 1 | 1 | 101 | 1 | 101 | 102 | 102 | 1 | 1 | 1 | 102 | 1 | 1 | 1 | 1 | 1 | 1 | 1 | 1 | 101 | 101 | 102 |
| YPTB0478 | 1 | 1 | 1 | 1 | 1 | 1 | 1 | 1 | 1 | 1 | 1 | 1 | 1 | 1 | 1 | 1 | 1 | 1 | 1 | 1 | 1 | 1 | 1 | 1 | 1 | 1 | 1 | 1 | 1 | 1 | 1 | 1 | 1 | 1 | 1 |
| YPTB0481 | 1 | 1 | 1 | 1 | 1 | 1 | 1 | 1 | 1 | 1 | 1 | 1 | 1 | 1 | 1 | 1 | 1 | 1 | 1 | 1 | 1 | 1 | 1 | 1 | 1 | 1 | 1 | 1 | 1 | 1 | 1 | 1 | 1 | 1 | 1 |
| YPTB0483 | 1 | 1 | 1 | 1 | 1 | 1 | 1 | 1 | 1 | 1 | 1 | 1 | 1 | 1 | 1 | 1 | 1 | 1 | 1 | 1 | 1 | 1 | 1 | 1 | 1 | 1 | 1 | 1 | 1 | 1 | 1 | 1 | 1 | 1 | 1 |
| YPTB0485 | 1 | 1 | 1 | 1 | 1 | 1 | 1 | 1 | 1 | 1 | 1 | 1 | 1 | 1 | 1 | 1 | 1 | 1 | 1 | 1 | 1 | 1 | 1 | 1 | 1 | 1 | 1 | 1 | 1 | 1 | 1 | 1 | 1 | 1 | 1 |
| YPTB0494 | 105 | 1 | 105 | 105 | 1 | 1 | 105 | 1 | 1 | 105 | 105 | 105 | 1 | 1 | 1 | 105 | 1 | 105 | 105 | 105 | 1 | 1 | 1 | 105 | 1 | 1 | 1 | 1 | 1 | 1 | 1 | 1 | 105 | 105 | 105 |
| YPTB0523 | 120 | 119 | 120 | 119 | 1 | 1 | 120 | 124 | 119 | 119 | 119 | 1 | 1 | 1 | 1 | 119 | 1 | 121 | 358 | 120 | 1 | 1 | 1 | 120 | 1 | 1 | 1 | 1 | 1 | 1 | 1 | 1 | 119 | 119 | 120 |
| YPTB0530 | 1 | 66 | 1 | 68 | 1 | 1 | 1 | 1 | 66 | 68 | 68 | 1 | 1 | 1 | 1 | 68 | 1 | 1 | 1 | 1 | 1 | 1 | 1 | 1 | 1 | 1 | 1 | 1 | 1 | 1 | 1 | 1 | 68 | 68 | 1 |
| YPTB0531 | 1 | 115 | 1 | 1 | 1 | 1 | 1 | 116 | 115 | 1 | 1 | 1 | 1 | 1 | 1 | 1 | 1 | 1 | 1 | 1 | 1 | 1 | 1 | 1 | 1 | 1 | 1 | 1 | 1 | 1 | 1 | 1 | 1 | 1 | 1 |
| YPTB0532 | 1 | 108 | 1 | 1 | 1 | 1 | 1 | 108 | 108 | 1 | 1 | 108 | 108 | 108 | 1 | 1 | 108 | 108 | 1 | 1 | 1 | 1 | 1 | 1 | 108 | 108 | 1 | 1 | 1 | 1 | 1 | 108 | 1 | 1 | 1 |
| YPTB0548 | 1 | 104 | 1 | 105 | 1 | 1 | 1 | 1 | 104 | 105 | 105 | 103 | 1 | 1 | 1 | 105 | 1 | 1 | 1 | 1 | 1 | 1 | 1 | 1 | 1 | 1 | 1 | 1 | 1 | 1 | 1 | 1 | 105 | 105 | 1 |
| YPTB0573 | 73 | 77 | 73 | 76 | 1 | 1 | 73 | 77 | 77 | 76 | 76 | 74 | 1 | 1 | 1 | 76 | 1 | 75 | 73 | 73 | 1 | 1 | 1 | 73 | 1 | 1 | 1 | 1 | 1 | 1 | 1 | 1 | 76 | 76 | 73 |
| YPTB0574 | 80 | 79 | 80 | 1 | 1 | 1 | 80 | 79 | 79 | 1 | 1 | 79 | 1 | 1 | 1 | 1 | 1 | 81 | 80 | 80 | 1 | 1 | 1 | 80 | 1 | 1 | 1 | 1 | 1 | 1 | 1 | 1 | 1 | 1 | 80 |
| YPTB0576 | 1 | 81 | 1 | 1 | 1 | 79 | 1 | 81 | 81 | 1 | 1 | 79 | 79 | 79 | 79 | 1 | 79 | 1 | 1 | 1 | 79 | 79 | 79 | 1 | 79 | 79 | 79 | 79 | 79 | 79 | 79 | 79 | 1 | 1 | 1 |
| YPTB0577 | 1 | 1 | 1 | 30 | 1 | 1 | 1 | 1 | 1 | 30 | 30 | 1 | 1 | 1 | 1 | 30 | 1 | 29 | 1 | 1 | 1 | 1 | 1 | 1 | 1 | 1 | 1 | 1 | 1 | 1 | 1 | 1 | 30 | 30 | 1 |
| YPTB0583 | 133 | 135 | 133 | 133 | 133 | 133 | 133 | 133 | 135 | 133 | 133 | 132 | 133 | 133 | 133 | 133 | 133 | 134 | 133 | 133 | 133 | 133 | 133 | 133 | 133 | 133 | 133 | 133 | 133 | 133 | 133 | 133 | 133 | 133 | 133 |
| YPTB0598 | 1 | 1 | 1 | 108 | 1 | 1 | 1 | 1 | 1 | 82 | 108 | 1 | 1 | 1 | 1 | 82 | 1 | 1 | 1 | 1 | 1 | 1 | 1 | 1 | 1 | 1 | 1 | 1 | 1 | 1 | 1 | 1 | 108 | 108 | 1 |
| YPTB0599 | 1 | 1 | 1 | 90 | 1 | 1 | 1 | 1 | 1 | 91 | 90 | 1 | 1 | 1 | 1 | 91 | 1 | 1 | 1 | 1 | 1 | 1 | 1 | 1 | 1 | 1 | 1 | 1 | 1 | 1 | 1 | 1 | 90 | 90 | 1 |
| YPTB0603 | 97 | 98 | 97 | 99 | 1 | 1 | 97 | 100 | 98 | 99 | 99 | 96 | 1 | 1 | 1 | 99 | 1 | 97 | 97 | 97 | 1 | 1 | 1 | 97 | 1 | 1 | 1 | 1 | 1 | 1 | 1 | 1 | 99 | 99 | 97 |
| YPTB0604 | 422 | 135 | 132 | 136 | 1 | 1 | 132 | 132 | 135 | 136 | 136 | 131 | 1 | 1 | 1 | 136 | 1 | 132 | 178 | 132 | 1 | 1 | 1 | 132 | 1 | 1 | 1 | 1 | 1 | 1 | 1 | 1 | 136 | 136 | 132 |
| YPTB0616 | 93 | 94 | 93 | 94 | 1 | 1 | 93 | 1 | 94 | 94 | 94 | 1 | 1 | 1 | 1 | 94 | 1 | 93 | 93 | 93 | 1 | 1 | 1 | 93 | 1 | 1 | 1 | 1 | 1 | 1 | 1 | 1 | 94 | 94 | 93 |
| YPTB0620 | 1 | 106 | 1 | 106 | 1 | 1 | 1 | 1 | 106 | 106 | 106 | 103 | 1 | 1 | 1 | 106 | 1 | 1 | 1 | 1 | 1 | 1 | 1 | 1 | 1 | 1 | 1 | 1 | 1 | 1 | 1 | 1 | 106 | 106 | 1 |
| YPTB0622 | 152 | 117 | 118 | 117 | 1 | 1 | 118 | 117 | 154 | 117 | 117 | 117 | 1 | 1 | 1 | 117 | 1 | 117 | 118 | 118 | 1 | 1 | 1 | 118 | 1 | 1 | 1 | 1 | 1 | 1 | 1 | 1 | 117 | 117 | 118 |
| YPTB0632 | 1 | 1 | 1 | 1 | 1 | 1 | 1 | 1 | 1 | 1 | 1 | 1 | 1 | 1 | 1 | 1 | 1 | 54 | 1 | 1 | 1 | 1 | 1 | 1 | 1 | 1 | 1 | 1 | 1 | 1 | 1 | 1 | 1 | 1 | 1 |
| YPTB0633 | 86 | 85 | 86 | 1 | 1 | 1 | 86 | 1 | 85 | 1 | 1 | 1 | 1 | 1 | 1 | 1 | 1 | 1 | 86 | 86 | 1 | 1 | 1 | 86 | 1 | 1 | 1 | 1 | 1 | 1 | 1 | 1 | 1 | 1 | 86 |
| YPTB0671 | 124 | 128 | 124 | 129 | 1 | 1 | 124 | 131 | 128 | 129 | 129 | 123 | 1 | 1 | 1 | 129 | 1 | 125 | 124 | 124 | 1 | 1 | 1 | 124 | 1 | 1 | 1 | 1 | 1 | 1 | 1 | 1 | 129 | 129 | 124 |
| YPTB0685 | 1 | 104 | 1 | 105 | 1 | 1 | 1 | 101 | 104 | 105 | 105 | 101 | 1 | 1 | 1 | 105 | 1 | 102 | 1 | 1 | 1 | 1 | 1 | 1 | 1 | 1 | 1 | 1 | 1 | 1 | 1 | 1 | 105 | 105 | 1 |
| YPTB0690 | 107 | 106 | 107 | 107 | 1 | 1 | 107 | 107 | 106 | 107 | 107 | 106 | 1 | 1 | 1 | 107 | 1 | 106 | 107 | 107 | 1 | 1 | 1 | 107 | 1 | 1 | 1 | 1 | 1 | 1 | 1 | 1 | 107 | 107 | 107 |
| YPTB0692 | 1 | 1 | 1 | 1 | 1 | 1 | 1 | 1 | 1 | 1 | 1 | 1 | 1 | 1 | 1 | 1 | 1 | 1 | 1 | 1 | 1 | 1 | 1 | 1 | 1 | 1 | 1 | 1 | 1 | 1 | 1 | 1 | 1 | 1 | 1 |
| YPTB0693 | 92 | 1 | 92 | 92 | 1 | 1 | 92 | 92 | 1 | 92 | 92 | 92 | 1 | 1 | 1 | 92 | 1 | 92 | 92 | 92 | 1 | 1 | 1 | 92 | 1 | 1 | 1 | 1 | 1 | 1 | 1 | 1 | 92 | 92 | 92 |
| YPTB0701 | 81 | 81 | 81 | 81 | 1 | 1 | 81 | 81 | 81 | 81 | 81 | 81 | 1 | 1 | 1 | 81 | 1 | 81 | 81 | 81 | 1 | 1 | 1 | 81 | 1 | 1 | 1 | 1 | 1 | 1 | 1 | 1 | 81 | 81 | 81 |
| YPTB0703 | 102 | 103 | 102 | 102 | 1 | 1 | 102 | 102 | 103 | 102 | 102 | 102 | 1 | 1 | 1 | 102 | 1 | 102 | 102 | 102 | 1 | 1 | 1 | 102 | 1 | 1 | 1 | 1 | 1 | 1 | 1 | 1 | 102 | 102 | 102 |
| YPTB0712 | 1 | 1 | 1 | 1 | 1 | 1 | 1 | 1 | 1 | 1 | 1 | 1 | 1 | 1 | 1 | 1 | 1 | 1 | 1 | 1 | 1 | 1 | 1 | 1 | 1 | 1 | 1 | 1 | 1 | 1 | 1 | 1 | 1 | 1 | 1 |
| YPTB0713 | 133 | 137 | 133 | 139 | 1 | 1 | 133 | 141 | 137 | 139 | 139 | 132 | 1 | 1 | 1 | 139 | 1 | 134 | 133 | 133 | 1 | 1 | 1 | 133 | 1 | 1 | 1 | 1 | 1 | 1 | 1 | 1 | 139 | 139 | 133 |
| YPTB0719 | 91 | 89 | 91 | 93 | 1 | 1 | 91 | 89 | 89 | 93 | 93 | 90 | 1 | 1 | 1 | 93 | 1 | 92 | 91 | 91 | 1 | 1 | 1 | 91 | 1 | 1 | 1 | 1 | 1 | 1 | 1 | 1 | 93 | 93 | 91 |
| YPTB0723 | 1 | 87 | 1 | 1 | 1 | 1 | 1 | 1 | 87 | 1 | 1 | 1 | 1 | 1 | 1 | 1 | 1 | 1 | 1 | 1 | 1 | 1 | 1 | 1 | 1 | 1 | 1 | 1 | 1 | 1 | 1 | 1 | 1 | 1 | 1 |
| YPTB0726 | 96 | 97 | 96 | 99 | 1 | 1 | 96 | 97 | 97 | 99 | 99 | 95 | 1 | 1 | 1 | 99 | 1 | 97 | 96 | 96 | 1 | 1 | 1 | 96 | 1 | 1 | 1 | 1 | 1 | 1 | 1 | 1 | 99 | 99 | 96 |
| YPTB0731 | 71 | 73 | 71 | 72 | 1 | 1 | 71 | 72 | 73 | 72 | 72 | 72 | 1 | 1 | 1 | 72 | 1 | 71 | 71 | 71 | 1 | 1 | 1 | 71 | 1 | 1 | 1 | 1 | 1 | 1 | 1 | 1 | 72 | 72 | 71 |
| YPTB0734 | 1 | 1 | 1 | 1 | 1 | 1 | 1 | 1 | 1 | 1 | 1 | 1 | 1 | 1 | 1 | 1 | 1 | 1 | 1 | 1 | 1 | 1 | 1 | 1 | 1 | 1 | 1 | 1 | 1 | 1 | 1 | 1 | 1 | 1 | 1 |
| YPTB0744 | 1 | 1 | 1 | 1 | 1 | 1 | 1 | 1 | 1 | 1 | 1 | 1 | 1 | 1 | 1 | 1 | 1 | 1 | 1 | 1 | 1 | 1 | 1 | 1 | 1 | 1 | 1 | 1 | 1 | 1 | 1 | 1 | 1 | 1 | 1 |
| YPTB0747 | 92 | 95 | 92 | 95 | 1 | 1 | 92 | 95 | 95 | 95 | 95 | 1 | 1 | 1 | 1 | 95 | 1 | 93 | 92 | 92 | 1 | 1 | 1 | 92 | 1 | 1 | 1 | 1 | 1 | 1 | 1 | 1 | 95 | 95 | 92 |
| YPTB0751 | 180 | 137 | 134 | 139 | 1 | 1 | 134 | 140 | 137 | 139 | 139 | 133 | 1 | 1 | 1 | 139 | 1 | 134 | 134 | 134 | 1 | 1 | 1 | 134 | 1 | 1 | 1 | 1 | 1 | 1 | 1 | 1 | 139 | 139 | 134 |
| YPTB0765 | 1 | 107 | 1 | 107 | 1 | 1 | 1 | 108 | 107 | 107 | 107 | 107 | 1 | 1 | 1 | 107 | 1 | 1 | 1 | 1 | 1 | 1 | 1 | 1 | 1 | 1 | 1 | 1 | 1 | 1 | 1 | 1 | 107 | 107 | 1 |
| YPTB0768 | 1 | 1 | 1 | 1 | 1 | 1 | 1 | 67 | 1 | 1 | 1 | 1 | 1 | 1 | 1 | 1 | 1 | 1 | 1 | 1 | 1 | 1 | 1 | 1 | 1 | 1 | 1 | 1 | 1 | 1 | 1 | 1 | 1 | 1 | 1 |
| YPTB0774 | 1 | 1 | 1 | 1 | 1 | 1 | 1 | 1 | 1 | 1 | 1 | 1 | 1 | 1 | 1 | 1 | 1 | 1 | 1 | 1 | 1 | 1 | 1 | 1 | 1 | 1 | 1 | 1 | 1 | 1 | 1 | 1 | 1 | 1 | 1 |
| YPTB0833 | 113 | 1 | 113 | 117 | 1 | 1 | 113 | 118 | 1 | 117 | 117 | 1 | 1 | 1 | 1 | 117 | 1 | 114 | 113 | 113 | 1 | 1 | 1 | 113 | 1 | 1 | 1 | 1 | 1 | 1 | 1 | 1 | 117 | 117 | 113 |
| YPTB0834 | 1 | 1 | 1 | 1 | 1 | 1 | 1 | 1 | 1 | 1 | 1 | 1 | 1 | 1 | 1 | 1 | 1 | 1 | 1 | 1 | 1 | 1 | 1 | 1 | 1 | 1 | 1 | 1 | 1 | 1 | 1 | 1 | 1 | 1 | 1 |
| YPTB0835 | 1 | 61 | 1 | 61 | 1 | 1 | 1 | 59 | 61 | 61 | 61 | 1 | 1 | 1 | 1 | 61 | 1 | 59 | 1 | 1 | 1 | 1 | 1 | 1 | 1 | 1 | 1 | 1 | 1 | 1 | 1 | 1 | 61 | 61 | 1 |
| YPTB0842 | 1 | 119 | 1 | 120 | 1 | 1 | 1 | 121 | 119 | 120 | 120 | 115 | 1 | 1 | 1 | 120 | 1 | 116 | 1 | 1 | 1 | 1 | 1 | 1 | 1 | 1 | 1 | 1 | 1 | 1 | 1 | 1 | 120 | 120 | 1 |
| YPTB0851 | 1 | 135 | 189 | 136 | 1 | 1 | 1 | 138 | 135 | 136 | 136 | 132 | 132 | 132 | 1 | 136 | 132 | 183 | 1 | 1 | 1 | 1 | 1 | 1 | 132 | 132 | 1 | 1 | 1 | 1 | 1 | 132 | 136 | 136 | 1 |
| YPTB0857 | 1 | 70 | 1 | 1 | 1 | 1 | 1 | 70 | 70 | 1 | 1 | 1 | 1 | 1 | 1 | 1 | 1 | 1 | 1 | 1 | 1 | 1 | 1 | 1 | 1 | 1 | 1 | 1 | 1 | 1 | 1 | 1 | 1 | 1 | 1 |
| YPTB0858 | 1 | 1 | 1 | 77 | 1 | 1 | 1 | 77 | 1 | 77 | 77 | 1 | 1 | 1 | 1 | 77 | 1 | 1 | 1 | 1 | 1 | 1 | 1 | 1 | 1 | 1 | 1 | 1 | 1 | 1 | 1 | 1 | 77 | 77 | 1 |
| YPTB0873 | 121 | 122 | 121 | | 1 | 1 | 121 | 126 | 122 | 124 | 124 | 120 | 1 | 1 | 1 | 124 | 1 | 121 | 121 | 121 | 1 | 1 | 1 | 121 | 1 | 1 | 1 | 1 | 1 | 1 | 1 | 1 | 124 | 124 | 121 |
| YPTB0876 | 1 | 100 | 120 | 101 | 1 | 1 | 120 | 1 | 100 | 101 | 101 | 1 | 1 | 1 | 1 | 101 | 1 | 99 | 1 | 1 | 1 | 1 | 1 | 1 | 1 | 1 | 1 | 1 | 1 | 1 | 1 | 1 | 101 | 101 | 1 |
| YPTB0887 | 1 | 111 | 1 | 145 | 1 | 1 | 1 | 1 | 111 | 112 | 145 | 1 | 1 | 1 | 1 | 112 | 1 | 109 | 1 | 1 | 1 | 1 | 1 | 1 | 1 | 1 | 1 | 1 | 1 | 1 | 1 | 1 | 145 | 145 | 1 |
| YPTB0889 | 91 | 1 | 91 | 91 | 1 | 1 | 91 | 91 | 1 | 91 | 91 | 1 | 1 | 1 | 1 | 91 | 1 | 91 | 91 | 91 | 1 | 1 | 1 | 91 | 1 | 1 | 1 | 1 | 1 | 1 | 1 | 1 | 91 | 91 | 91 |
| YPTB0890 | 79 | 79 | 79 | 81 | 1 | 1 | 79 | 79 | 79 | 81 | 81 | 79 | 1 | 1 | 1 | 81 | 1 | 79 | 79 | 79 | 1 | 1 | 1 | 79 | 1 | 1 | 1 | 1 | 1 | 1 | 1 | 1 | 81 | 81 | 79 |
| YPTB0892 | 108 | 107 | 108 | 110 | 1 | 1 | 108 | 107 | 107 | 110 | 110 | 108 | 139 | 1 | 1 | 110 | 1 | 107 | 108 | 108 | 1 | 1 | 1 | 108 | 1 | 1 | 1 | 1 | 1 | 1 | 1 | 140 | 110 | 110 | 108 |
| YPTB0894 | 1 | 1 | 1 | 1 | 1 | 1 | 1 | 1 | 1 | 1 | 1 | 1 | 1 | 1 | 1 | 1 | 1 | 1 | 1 | 1 | 1 | 1 | 1 | 1 | 1 | 1 | 1 | 1 | 1 | 1 | 1 | 1 | 1 | 1 | 1 |
| YPTB0903 | 1 | 1 | 1 | 1 | 1 | 1 | 1 | 1 | 1 | 1 | 1 | 1 | 1 | 1 | 1 | 1 | 1 | 1 | 1 | 1 | 1 | 1 | 1 | 1 | 1 | 1 | 1 | 1 | 1 | 1 | 1 | 1 | 1 | 1 | 1 |
| YPTB0920 | 109 | 111 | 109 | 147 | 1 | 1 | 109 | 113 | 111 | 109 | 147 | 1 | 1 | 1 | 1 | 109 | 1 | 110 | 109 | 109 | 1 | 1 | 1 | 109 | 1 | 1 | 1 | 1 | 1 | 1 | 1 | 1 | 147 | 147 | 109 |
| YPTB0925 | 67 | 67 | 67 | 69 | 1 | 1 | 67 | 82 | 67 | 69 | 69 | 67 | 1 | 1 | 1 | 69 | 1 | 67 | 67 | 67 | 1 | 1 | 1 | 67 | 1 | 1 | 1 | 1 | 1 | 1 | 1 | 1 | 69 | 69 | 67 |
| YPTB0930 | 340 | 1 | 120 | 123 | 1 | 1 | 120 | 119 | 1 | 123 | 123 | 1 | 1 | 1 | 1 | 123 | 1 | 161 | 412 | 120 | 1 | 1 | 1 | 120 | 1 | 1 | 1 | 1 | 1 | 1 | 1 | 1 | 123 | 123 | 120 |
| YPTB0933 | 1 | 1 | 1 | 1 | 1 | 1 | 1 | 1 | 1 | 1 | 1 | 1 | 1 | 1 | 1 | 1 | 1 | 1 | 1 | 1 | 1 | 1 | 1 | 1 | 150 | 1 | 1 | 1 | 1 | 1 | 1 | 1 | 1 | 1 | 1 |
| YPTB0934 | 126 | 1 | 126 | 1 | 1 | 1 | 126 | 125 | 1 | 1 | 1 | 1 | 1 | 1 | 1 | 1 | 1 | 1 | 126 | 126 | 1 | 1 | 1 | 126 | 1 | 1 | 1 | 1 | 1 | 1 | 1 | 1 | 1 | 1 | 126 |
| YPTB0936 | 1 | 1 | 1 | 1 | 1 | 1 | 1 | 1 | 1 | 1 | 1 | 1 | 1 | 1 | 1 | 1 | 1 | 1 | 1 | 1 | 1 | 1 | 1 | 1 | 1 | 1 | 1 | 1 | 1 | 1 | 1 | 1 | 1 | 1 | 1 |
| YPTB0950 | 1 | 1 | 1 | 1 | 1 | 1 | 1 | 76 | 1 | 1 | 1 | 1 | 1 | 1 | 1 | 1 | 1 | 1 | 1 | 1 | 1 | 1 | 1 | 1 | 1 | 1 | 1 | 1 | 1 | 1 | 1 | 1 | 1 | 1 | 1 |
| YPTB0951 | 1 | 126 | 1 | 125 | 1 | 1 | 1 | 128 | 126 | 125 | 125 | 125 | 1 | 1 | 1 | 125 | 1 | 1 | 1 | 1 | 1 | 1 | 1 | 1 | 1 | 1 | 1 | 1 | 1 | 1 | 1 | 1 | 125 | 125 | 159 |
| YPTB0955 | 1 | 82 | 1 | 1 | 1 | 1 | 1 | 81 | 82 | 1 | 1 | 1 | 1 | 1 | 1 | 1 | 1 | 81 | 1 | 1 | 1 | 1 | 1 | 1 | 1 | 1 | 1 | 1 | 1 | 1 | 1 | 1 | 1 | 1 | 1 |
| YPTB0958 | 1 | 93 | 1 | 93 | 1 | 1 | 1 | 1 | 263 | 93 | 93 | 91 | 1 | 1 | 1 | 93 | 1 | 92 | 1 | 1 | 1 | 1 | 1 | 1 | 1 | 1 | 1 | 1 | 1 | 1 | 1 | 1 | 93 | 93 | 1 |
| YPTB0959 | 1 | 1 | 1 | 1 | 1 | 1 | 1 | 1 | 1 | 1 | 1 | 1 | 1 | 1 | 1 | 1 | 1 | 1 | 1 | 1 | 1 | 1 | 1 | 1 | 1 | 1 | 1 | 1 | 1 | 1 | 1 | 1 | 1 | 1 | 1 |
| YPTB0960 | 100 | 100 | 100 | 102 | 1 | 99 | 100 | 100 | 100 | 102 | 102 | 99 | 99 | 99 | 99 | 102 | 99 | 100 | 100 | 100 | 99 | 99 | 99 | 100 | 99 | 99 | 99 | 99 | 99 | 99 | 99 | 99 | 102 | 102 | 100 |
| YPTB0962 | 1 | 1 | 1 | 1 | 1 | 1 | 1 | 1 | 1 | 1 | 1 | 1 | 1 | 1 | 1 | 1 | 1 | 1 | 1 | 1 | 1 | 1 | 1 | 1 | 1 | 1 | 1 | 1 | 1 | 1 | 1 | 1 | 1 | 1 | 1 |
| YPTB0963 | 129 | 131 | 129 | 130 | 1 | 1 | 129 | 1 | 131 | 130 | 130 | 1 | 1 | 1 | 1 | 130 | 1 | 130 | 129 | 129 | 1 | 1 | 1 | 129 | 1 | 1 | 1 | 1 | 1 | 1 | 1 | 1 | 130 | 130 | 129 |
| YPTB0966 | 1 | 103 | 1 | 1 | 1 | 1 | 1 | 1 | 103 | 1 | 1 | 1 | 1 | 1 | 1 | 1 | 1 | 1 | 1 | 1 | 1 | 1 | 1 | 1 | 1 | 1 | 1 | 1 | 1 | 1 | 1 | 1 | 1 | 1 | 1 |
| YPTB0970 | 1 | 1 | 1 | 1 | 1 | 1 | 1 | 1 | 1 | 1 | 1 | 1 | 1 | 1 | 1 | 1 | 1 | 1 | 1 | 1 | 1 | 1 | 1 | 1 | 1 | 1 | 1 | 1 | 1 | 1 | 1 | 1 | 1 | 1 | 1 |
| YPTB0979 | 46 | 1 | 46 | 1 | 1 | 1 | 46 | 1 | 1 | 1 | 1 | 1 | 1 | 1 | 1 | 1 | 1 | 46 | 46 | 46 | 1 | 1 | 1 | 46 | 1 | 1 | 1 | 1 | 1 | 1 | 1 | 1 | 1 | 1 | 46 |
| YPTB0981 | 1 | 1 | 1 | 1 | 1 | 1 | 1 | 1 | 1 | 1 | 1 | 1 | 1 | 1 | 1 | 1 | 1 | 1 | 1 | 1 | 1 | 1 | 1 | 1 | 1 | 1 | 1 | 1 | 1 | 1 | 1 | 1 | 1 | 1 | 1 |
| YPTB0990 | 1 | 72 | 1 | 72 | 1 | 1 | 1 | 1 | 72 | 72 | 72 | 72 | 1 | 1 | 1 | 72 | 1 | 72 | 1 | 1 | 1 | 1 | 1 | 1 | 1 | 1 | 1 | 1 | 1 | 1 | 1 | 1 | 72 | 72 | 1 |
| YPTB0993 | 48 | 1 | 48 | 48 | 1 | 1 | 48 | 1 | 1 | 48 | 48 | 48 | 1 | 1 | 1 | 48 | 1 | 48 | 48 | 48 | 1 | 1 | 1 | 48 | 1 | 1 | 1 | 1 | 1 | 1 | 1 | 1 | 48 | 48 | 48 |
| YPTB0994 | 96 | 96 | 96 | 97 | 1 | 1 | 96 | 96 | 96 | 97 | 97 | 96 | 1 | 1 | 1 | 97 | 1 | 96 | 96 | 96 | 1 | 1 | 1 | 96 | 1 | 1 | 1 | 1 | 1 | 1 | 1 | 1 | 97 | 97 | 96 |
| YPTB1025 | 1 | 1 | 1 | 1 | 1 | 1 | 1 | 95 | 1 | 1 | 1 | 1 | 1 | 1 | 1 | 1 | 1 | 1 | 1 | 1 | 1 | 1 | 1 | 1 | 1 | 1 | 1 | 1 | 1 | 1 | 1 | 1 | 1 | 1 | 1 |
| YPTB1027 | 316 | 116 | 1 | 116 | 1 | 1 | 1 | 117 | 116 | 116 | 116 | 1 | 1 | 1 | 1 | 116 | 1 | 1 | 1 | 1 | 1 | 1 | 1 | 1 | 1 | 1 | 1 | 1 | 1 | 1 | 1 | 1 | 116 | 116 | 1 |
| YPTB1029 | 95 | 98 | 95 | 95 | 1 | 1 | 95 | 101 | 98 | 99 | 95 | 1 | 1 | 1 | 1 | 99 | 1 | 95 | 95 | 95 | 1 | 1 | 1 | 95 | 1 | 1 | 1 | 1 | 1 | 1 | 1 | 1 | 95 | 95 | 95 |
| YPTB1034 | 1 | 1 | 1 | 71 | 1 | 1 | 1 | 1 | 1 | 71 | 71 | 1 | 1 | 1 | 1 | 71 | 1 | 1 | 1 | 1 | 1 | 1 | 1 | 1 | 1 | 1 | 1 | 1 | 1 | 1 | 1 | 1 | 71 | 71 | 1 |
| YPTB1091 | 1 | 99 | 1 | 98 | 1 | 1 | 1 | 96 | 99 | 98 | 98 | 96 | 96 | 96 | 1 | 98 | 96 | 1 | 1 | 1 | 1 | 1 | 1 | 1 | 96 | 96 | 1 | 1 | 1 | 1 | 1 | 96 | 98 | 98 | 1 |
| YPTB1096 | 1 | 1 | 1 | 1 | 1 | 1 | 1 | 94 | 1 | 1 | 1 | 93 | 93 | 93 | 1 | 1 | 93 | 1 | 1 | 1 | 1 | 1 | 1 | 1 | 93 | 93 | 1 | 1 | 1 | 1 | 1 | 93 | 1 | 1 | 1 |
| YPTB1097 | 126 | 129 | 126 | 131 | 1 | 1 | 126 | 133 | 129 | 131 | 131 | 125 | 1 | 1 | 1 | 131 | 1 | 126 | 126 | 126 | 1 | 1 | 1 | 126 | 1 | 1 | 1 | 1 | 1 | 1 | 1 | 1 | 131 | 131 | 126 |
| YPTB1098 | 1 | 1 | 1 | 1 | 1 | 1 | 1 | 1 | 1 | 75 | 1 | 1 | 1 | 1 | 1 | 75 | 1 | 1 | 1 | 1 | 1 | 1 | 1 | 1 | 1 | 1 | 1 | 1 | 1 | 1 | 1 | 1 | 1 | 1 | 1 |
| YPTB1103 | 1 | 126 | 1 | 128 | 1 | 1 | 1 | 130 | 126 | 128 | 128 | 123 | 123 | 123 | 1 | 128 | 123 | 124 | 1 | 1 | 1 | 1 | 1 | 1 | 123 | 123 | 1 | 1 | 1 | 1 | 1 | 123 | 128 | 128 | 1 |
| YPTB1104 | 1 | 1 | | 1 | 1 | 1 | 1 | 1 | 1 | 74 | 1 | 1 | 1 | 1 | 1 | 74 | 1 | 1 | 1 | 1 | 1 | 1 | 1 | 1 | 1 | 1 | 1 | 1 | 1 | 1 | 1 | 1 | 1 | 1 | 1 |
| YPTB1105 | 1 | 1 | 1 | 1 | 1 | 1 | 1 | 1 | 1 | 1 | 1 | 1 | 1 | 1 | 1 | 1 | 1 | 1 | 1 | 1 | 1 | 1 | 1 | 1 | 1 | 1 | 1 | 1 | 1 | 1 | 1 | 1 | 1 | 1 | 1 |
| YPTB1106 | 1 | 1 | 1 | 84 | 1 | 1 | 1 | 84 | 1 | 84 | 84 | 1 | 1 | 1 | 1 | 84 | 1 | 84 | 1 | 1 | 1 | 1 | 1 | 1 | 1 | 1 | 1 | 1 | 1 | 1 | 1 | 1 | 84 | 84 | 1 |
| YPTB1110 | 1 | 1 | 1 | 89 | 1 | 1 | 1 | 89 | 1 | 89 | 89 | 89 | 89 | 89 | 1 | 89 | 89 | 1 | 1 | 1 | 1 | 1 | 1 | 1 | 89 | 89 | 1 | 1 | 1 | 1 | 1 | 89 | 89 | 89 | 1 |
| YPTB1116 | 92 | 92 | 92 | 1 | 1 | 1 | 92 | 92 | 92 | 1 | 1 | 1 | 1 | 1 | 1 | 1 | 1 | 1 | 92 | 92 | 1 | 1 | 1 | 92 | 1 | 1 | 1 | 1 | 1 | 1 | 1 | 1 | 1 | 1 | 92 |
| YPTB1119 | 1 | 92 | 1 | 118 | 1 | 1 | 1 | 90 | 92 | 1 | 118 | 1 | 1 | 1 | 1 | 1 | 1 | 90 | 1 | 1 | 1 | 1 | 1 | 1 | 1 | 1 | 1 | 1 | 1 | 1 | 1 | 1 | 118 | 118 | 1 |
| YPTB1125 | 1 | 1 | 1 | 1 | 1 | 1 | 1 | 1 | 1 | 1 | 1 | 71 | 1 | 1 | 1 | 1 | 1 | 1 | 1 | 1 | 1 | 1 | 1 | 1 | 1 | 1 | 1 | 1 | 1 | 1 | 1 | 1 | 1 | 1 | 1 |
| YPTB1135 | 1 | 1 | 1 | 1 | 1 | 1 | 1 | 1 | 1 | 1 | 1 | 1 | 1 | 1 | 1 | 1 | 1 | 1 | 1 | 1 | 1 | 1 | 1 | 1 | 1 | 1 | 1 | 1 | 1 | 1 | 1 | 1 | 1 | 1 | 1 |
| YPTB1141 | 1 | 64 | 1 | 64 | 1 | 1 | 1 | 64 | 64 | 64 | 64 | 1 | 1 | 1 | 1 | 64 | 1 | 64 | 1 | 1 | 1 | 1 | 1 | 1 | 1 | 1 | 1 | 1 | 1 | 1 | 1 | 1 | 64 | 64 | 1 |
| YPTB1144 | 1 | 1 | 1 | 46 | 1 | 1 | 1 | 46 | 1 | 46 | 46 | 47 | 1 | 1 | 1 | 46 | 1 | 1 | 1 | 1 | 1 | 1 | 1 | 1 | 1 | 1 | 1 | 1 | 1 | 1 | 1 | 1 | 46 | 46 | 1 |
| YPTB1145 | 1 | 123 | 1 | 122 | 1 | 1 | 1 | 122 | 123 | 122 | 122 | 1 | 1 | 1 | 1 | 122 | 1 | 1 | 1 | 1 | 1 | 1 | 1 | 1 | 1 | 1 | 1 | 1 | 1 | 1 | 1 | 1 | 122 | 122 | 1 |
| YPTB1146 | 1 | 1 | 1 | 1 | 1 | 1 | 1 | 1 | 1 | 1 | 1 | 1 | 1 | 1 | 1 | 1 | 1 | 1 | 1 | 1 | 1 | 1 | 1 | 1 | 1 | 1 | 1 | 1 | 1 | 1 | 1 | 1 | 1 | 1 | 1 |
| YPTB1147 | 178 | 139 | 178 | 135 | 1 | 1 | 178 | 135 | 139 | 135 | 135 | 1 | 1 | 1 | 1 | 135 | 1 | 179 | 178 | 178 | 1 | 1 | 1 | 178 | 1 | 1 | 1 | 1 | 1 | 1 | 1 | 1 | 135 | 135 | 178 |
| YPTB1149 | 108 | 110 | 108 | 107 | 1 | 1 | 108 | 112 | 110 | 107 | 107 | 107 | 1 | 1 | 1 | 107 | 1 | 109 | 108 | 108 | 1 | 1 | 1 | 108 | 1 | 1 | 1 | 1 | 1 | 1 | 1 | 1 | 107 | 107 | 108 |
| YPTB1155 | 1 | 1 | 1 | 48 | 1 | 1 | 1 | 1 | 1 | 48 | 48 | 1 | 1 | 1 | 1 | 48 | 1 | 1 | 1 | 1 | 1 | 1 | 1 | 1 | 1 | 1 | 1 | 1 | 1 | 1 | 1 | 1 | 48 | 48 | 1 |
| YPTB1157 | 64 | 65 | 64 | 64 | 1 | 64 | 64 | 64 | 65 | 64 | 64 | 64 | 64 | 64 | 64 | 64 | 64 | 64 | 64 | 64 | 64 | 64 | 64 | 64 | 64 | 64 | 64 | 64 | 64 | 64 | 64 | 64 | 64 | 64 | 64 |
| YPTB1159 | 1 | 1 | 1 | 1 | 1 | 1 | 1 | 1 | 1 | 1 | 1 | 1 | 120 | 120 | 1 | 1 | 120 | 89 | 1 | 1 | 1 | 1 | 1 | 1 | 120 | 120 | 1 | 1 | 1 | 1 | 1 | 120 | 1 | 1 | 1 |
| YPTB1166 | 100 | 1 | 100 | 1 | 1 | 1 | 100 | 100 | 1 | 1 | 1 | 100 | 1 | 1 | 1 | 1 | 1 | 1 | 100 | 100 | 1 | 1 | 1 | 100 | 1 | 1 | 1 | 1 | 1 | 1 | 1 | 1 | 1 | 1 | 100 |
| YPTB1169 | 112 | 113 | 112 | 113 | 1 | 1 | 112 | 112 | 113 | 113 | 113 | 112 | 146 | 1 | 1 | 113 | 1 | 145 | 112 | 112 | 1 | 1 | 1 | 112 | 1 | 1 | 1 | 1 | 1 | 1 | 1 | 1 | 113 | 113 | 112 |
| YPTB1172 | 104 | 1 | 104 | 107 | 1 | 1 | 104 | 103 | 1 | 107 | 107 | 103 | 1 | 1 | 1 | 107 | 1 | 1 | 104 | 104 | 1 | 1 | 1 | 104 | 1 | 1 | 1 | 1 | 1 | 1 | 1 | 1 | 107 | 107 | 104 |
| YPTB1192 | 62 | 62 | 62 | 1 | 1 | 1 | 62 | 1 | 62 | 1 | 1 | 1 | 1 | 1 | 1 | 1 | 1 | 62 | 62 | 62 | 1 | 1 | 1 | 62 | 1 | 1 | 1 | 1 | 1 | 1 | 1 | 1 | 1 | 1 | 62 |
| YPTB1209 | 150 | 110 | 150 | 110 | 1 | 1 | 150 | 113 | 110 | 110 | 152 | 111 | 114 | 114 | 1 | 110 | 114 | 1 | 150 | 150 | 1 | 1 | 1 | 150 | 151 | 151 | 1 | 1 | 1 | 1 | 1 | 151 | 110 | 110 | 150 |
| YPTB1212 | 109 | 113 | 109 | 114 | 1 | 1 | 109 | 109 | 113 | 114 | 114 | 110 | 114 | 114 | 1 | 114 | 114 | 1 | 109 | 150 | 1 | 1 | 1 | 150 | 114 | 114 | 1 | 1 | 1 | 1 | 1 | 114 | 114 | 114 | 109 |
| YPTB1253 | 1 | 113 | 1 | 115 | 1 | 1 | 1 | 111 | 113 | 115 | 115 | 112 | 1 | 1 | 1 | 115 | 1 | 1 | 1 | 1 | 1 | 1 | 1 | 1 | 1 | 1 | 1 | 1 | 1 | 1 | 1 | 1 | 115 | 115 | 1 |
| YPTB1297 | 1 | 1 | 1 | 35 | 1 | 1 | 1 | 1 | 1 | 34 | 35 | 1 | 1 | 1 | 1 | 34 | 1 | 1 | 1 | 1 | 1 | 1 | 1 | 1 | 1 | 1 | 1 | 1 | 1 | 1 | 1 | 1 | 35 | 35 | 1 |
| YPTB1309 | 1 | 1 | 1 | 1 | 1 | 1 | 1 | 1 | 1 | 1 | 1 | 1 | 1 | 1 | 1 | 1 | 1 | 1 | 1 | 1 | 1 | 1 | 1 | 1 | 1 | 1 | 1 | 1 | 1 | 1 | 1 | 1 | 1 | 1 | 1 |
| YPTB1316 | 1 | 82 | 1 | 81 | 1 | 1 | 1 | 83 | 82 | 81 | 81 | 1 | 1 | 1 | 1 | 81 | 1 | 97 | 1 | 1 | 1 | 1 | 1 | 1 | 1 | 1 | 1 | 1 | 1 | 1 | 1 | 1 | 81 | 81 | 1 |
| YPTB1330 | 356 | 119 | 1 | 121 | 1 | 1 | 1 | 114 | 119 | 121 | 121 | 115 | 1 | 1 | 1 | 121 | 1 | 116 | 1 | 1 | 1 | 1 | 1 | 1 | 1 | 1 | 1 | 1 | 1 | 1 | 1 | 1 | 121 | 121 | 1 |
| YPTB1358 | 1 | 1 | 1 | 1 | 1 | 1 | 1 | 1 | 1 | 1 | 1 | 1 | 1 | 1 | 1 | 1 | 1 | 57 | 1 | 1 | 1 | 1 | 1 | 1 | 1 | 1 | 1 | 1 | 1 | 1 | 1 | 1 | 1 | 1 | 1 |
| YPTB1361 | 105 | 105 | 105 | 105 | 1 | 1 | 105 | 105 | 105 | 105 | 105 | 1 | 1 | 1 | 1 | 105 | 1 | 106 | 105 | 105 | 1 | 1 | 1 | 105 | 1 | 1 | 1 | 1 | 1 | 1 | 1 | 1 | 105 | 105 | 105 |
| YPTB1362 | 1 | 118 | 1 | 1 | 1 | 1 | 1 | 1 | 118 | 1 | 1 | 1 | 1 | 1 | 1 | 1 | 1 | 1 | 1 | 1 | 1 | 1 | 382 | 1 | 1 | 1 | 1 | 1 | 1 | 1 | 1 | 1 | 1 | 1 | 1 |
| YPTB1376 | 94 | 96 | 94 | 1 | 1 | 1 | 94 | 99 | 94 | 1 | 1 | 1 | 1 | 1 | 1 | 1 | 1 | 94 | 94 | 94 | 1 | 1 | 1 | 94 | 1 | 1 | 1 | 1 | 1 | 1 | 1 | 1 | 1 | 1 | 94 |
| YPTB1377 | 1 | 92 | 1 | 1 | 1 | 1 | 1 | 91 | 92 | 1 | 1 | 1 | 1 | 1 | 1 | 1 | 1 | 1 | 1 | 1 | 1 | 1 | 1 | 1 | 1 | 1 | 1 | 1 | 1 | 1 | 1 | 1 | 1 | 1 | 1 |
| YPTB1386 | 1 | 134 | 1 | 136 | 1 | 1 | 1 | 138 | 134 | 136 | 136 | 1 | 1 | 1 | 1 | 136 | 1 | 131 | 1 | 1 | 1 | 1 | 1 | 1 | 1 | 1 | 1 | 1 | 1 | 1 | 1 | 1 | 136 | 136 | 1 |
| YPTB1391 | 132 | 137 | 132 | 132 | 1 | 1 | 132 | 132 | 137 | 132 | 132 | 1 | 1 | 1 | 1 | 132 | 1 | 134 | 132 | 132 | 1 | 1 | 1 | 132 | 1 | 1 | 1 | 1 | 1 | 1 | 1 | 1 | 132 | 132 | 132 |
| YPTB1394 | 1 | 1 | 1 | 1 | 1 | 1 | 1 | 1 | 1 | 1 | 1 | 1 | 1 | 1 | 1 | 1 | 1 | 1 | 1 | 1 | 1 | 1 | 1 | 1 | 1 | 1 | 1 | 1 | 1 | 1 | 1 | 1 | 1 | 1 | 1 |
| YPTB1395 | 1 | 1 | 1 | 1 | 1 | 1 | 1 | 1 | 1 | 1 | 1 | 1 | 1 | 1 | 1 | 1 | 1 | 1 | 1 | 1 | 1 | 1 | 1 | 1 | 1 | 1 | 1 | 1 | 1 | 1 | 1 | 1 | 1 | 1 | 1 |
| YPTB1397 | 127 | 130 | 127 | 131 | 1 | 1 | 127 | 1 | 130 | 131 | 131 | 126 | 1 | 1 | 1 | 131 | 1 | 1 | 127 | 127 | 1 | 1 | 1 | 127 | 1 | 1 | 1 | 1 | 1 | 1 | 1 | 1 | 131 | 131 | 127 |
| YPTB1403 | 115 | 115 | 115 | 116 | 1 | 1 | 115 | 117 | 115 | 116 | 116 | 1 | 1 | 1 | 1 | 116 | 1 | 115 | 115 | 115 | 1 | 1 | 1 | 115 | 1 | 1 | 1 | 1 | 1 | 1 | 1 | 1 | 116 | 116 | 115 |
| YPTB1408 | 1 | 1 | 1 | 125 | 167 | 1 | 1 | 126 | 1 | 125 | 125 | 1 | 1 | 1 | 1 | 125 | 1 | 123 | 1 | 1 | 1 | 1 | 1 | 1 | 1 | 1 | 1 | 1 | 1 | 1 | 1 | 1 | 125 | 125 | 1 |
| YPTB1414 | 1 | 113 | 1 | 113 | 1 | 1 | 1 | 1 | 113 | 113 | 113 | 145 | 115 | 115 | 1 | 113 | 334 | 112 | 1 | 1 | 1 | 1 | 1 | 1 | 115 | 115 | 1 | 1 | 1 | 1 | 1 | 115 | 113 | 113 | 1 |
| YPTB1416 | 1 | 1 | 1 | 68 | 1 | 1 | 1 | 1 | 1 | 68 | 68 | 1 | 68 | 68 | 1 | 68 | 68 | 1 | 1 | 1 | 1 | 1 | 1 | 1 | 68 | 68 | 1 | 1 | 1 | 1 | 1 | 68 | 68 | 68 | 1 |
| YPTB1417 | 1 | 80 | 1 | 1 | 1 | 1 | 1 | 1 | 80 | 1 | 1 | 1 | 82 | 82 | 1 | 1 | 82 | 79 | 1 | 1 | 1 | 1 | 1 | 1 | 82 | 82 | 1 | 1 | 1 | 1 | 1 | 82 | 1 | 1 | 1 |
| YPTB1428 | 1 | 112 | 1 | 1 | 1 | 1 | 1 | 1 | 112 | 1 | 1 | 1 | 1 | 1 | 1 | 1 | 1 | 112 | 1 | 1 | 1 | 1 | 1 | 1 | 1 | 1 | 1 | 1 | 1 | 1 | 1 | 1 | 1 | 1 | 1 |
| YPTB1436 | 107 | 111 | 107 | 112 | 1 | 1 | 107 | 106 | 111 | 112 | 112 | 1 | 1 | 1 | 1 | 112 | 1 | 108 | 107 | 107 | 1 | 1 | 1 | 107 | 1 | 1 | 1 | 1 | 1 | 1 | 1 | 1 | 112 | 112 | 107 |
| YPTB1439 | 1 | 104 | 1 | 100 | 1 | 1 | 1 | 100 | 104 | 100 | 100 | 100 | 100 | 100 | 1 | 100 | 100 | 103 | 1 | 1 | 1 | 1 | 1 | 1 | 100 | 100 | 1 | 1 | 1 | 1 | 1 | 100 | 100 | 100 | 1 |
| YPTB1442 | 1 | 476 | 1 | 130 | 1 | 1 | 1 | 131 | 126 | 130 | 130 | 124 | 124 | 124 | 1 | 130 | 124 | 126 | 1 | 1 | 1 | 1 | 1 | 1 | 124 | 124 | 1 | 1 | 1 | 1 | 1 | 124 | 130 | 130 | 1 |
| YPTB1445 | 1 | 1 | 1 | 1 | 1 | 1 | 1 | 1 | 1 | 1 | 1 | 1 | 1 | 1 | 1 | 1 | 1 | 1 | 1 | 1 | 1 | 1 | 1 | 1 | 1 | 1 | 1 | 1 | 1 | 1 | 1 | 1 | 1 | 1 | 1 |
| YPTB1454 | 1 | 1 | 1 | 1 | 1 | 1 | 1 | 66 | 1 | 1 | 1 | 66 | 66 | 66 | 1 | 1 | 66 | 67 | 1 | 1 | 1 | 1 | 1 | 1 | 66 | 66 | 1 | 1 | 1 | 1 | 1 | 66 | 1 | 1 | 1 |
| YPTB1524 | 101 | 102 | 101 | 108 | 1 | 1 | 101 | 104 | 102 | 101 | 108 | 1 | 1 | 1 | 1 | 101 | 1 | 101 | 126 | 126 | 1 | 1 | 1 | 126 | 1 | 1 | 1 | 1 | 1 | 1 | 1 | 1 | 108 | 108 | 101 |
| YPTB1531 | 130 | 1 | 130 | | 1 | 1 | 130 | 136 | 1 | 134 | | 129 | 137 | 137 | 1 | 134 | 137 | 131 | 130 | 130 | 1 | 1 | 1 | 130 | 137 | 137 | 1 | 1 | 1 | 1 | 1 | 137 | | | 130 |
| YPTB1535 | 126 | 1 | 126 | 1 | 1 | 1 | 126 | 126 | 1 | 1 | 1 | 1 | 1 | 1 | 1 | 1 | 1 | 127 | 126 | 126 | 1 | 1 | 1 | 126 | 1 | 1 | 1 | 1 | 1 | 1 | 1 | 1 | 1 | 1 | 126 |
| YPTB1537 | 1 | 80 | 1 | 80 | 1 | 1 | 1 | 1 | 80 | 80 | 80 | 1 | 1 | 1 | 1 | 80 | 1 | 1 | 1 | 1 | 1 | 1 | 1 | 1 | 1 | 1 | 1 | 1 | 1 | 1 | 1 | 1 | 80 | 80 | 1 |
| YPTB1555 | 80 | 81 | 80 | 80 | 1 | 1 | 80 | 1 | 81 | 80 | 80 | 1 | 1 | 1 | 1 | 80 | 1 | 78 | 80 | 80 | 1 | 1 | 1 | 80 | 1 | 1 | 1 | 1 | 1 | 1 | 1 | 1 | 80 | 80 | 80 |
| YPTB1557 | 104 | 1 | 104 | 1 | 1 | 1 | 104 | 107 | 1 | 1 | 1 | 1 | 1 | 1 | 1 | 1 | 1 | 104 | 104 | 104 | 1 | 1 | 1 | 104 | 1 | 1 | 1 | 1 | 1 | 1 | 1 | 1 | 1 | 1 | 104 |
| YPTB1558 | 1 | 1 | 1 | 1 | 1 | 1 | 1 | 1 | 1 | 1 | 1 | 1 | 1 | 1 | 1 | 1 | 1 | 1 | 1 | 1 | 1 | 1 | 1 | 1 | 114 | 1 | 1 | 1 | 1 | 1 | 1 | 1 | 1 | 1 | 1 |
| YPTB1559 | 121 | 125 | 121 | 126 | 1 | 1 | 121 | 128 | 159 | 126 | 126 | 120 | 1 | 1 | 1 | 126 | 1 | 122 | 121 | 121 | 1 | 1 | 1 | 121 | 1 | 1 | 1 | 1 | 1 | 1 | 1 | 1 | 126 | 126 | 121 |
| YPTB1615 | 62 | 1 | 62 | 1 | 1 | 1 | 62 | 1 | 1 | 1 | 1 | 1 | 1 | 1 | 1 | 1 | 1 | 1 | 62 | 62 | 1 | 1 | 1 | 62 | 1 | 1 | 1 | 1 | 1 | 1 | 1 | 1 | 1 | 1 | 62 |
| YPTB1647 | 124 | 1 | 124 | 126 | 1 | 1 | 124 | 1 | 1 | 126 | 126 | 1 | 1 | 1 | 1 | 126 | 1 | 1 | 124 | 124 | 1 | 1 | 1 | 124 | 1 | 1 | 1 | 1 | 1 | 1 | 1 | 1 | 126 | 126 | 124 |
| YPTB1724 | 123 | 1 | 123 | 124 | 1 | 1 | 123 | 122 | 1 | 126 | 124 | 1 | 1 | 1 | 1 | 126 | 1 | 124 | 123 | 123 | 1 | 1 | 1 | 123 | 1 | 1 | 1 | 1 | 1 | 1 | 1 | 1 | 124 | 124 | 123 |
| YPTB1795 | 100 | 100 | 100 | 100 | 1 | 1 | 100 | 1 | 100 | 100 | 100 | 1 | 100 | 100 | 1 | 100 | 100 | 100 | 100 | 100 | 1 | 1 | 1 | 100 | 100 | 100 | 1 | 1 | 1 | 1 | 1 | 100 | 100 | 100 | 100 |
| YPTB1921 | 1 | 1 | 1 | 1 | 1 | 1 | 1 | 1 | 1 | 1 | 1 | 1 | 1 | 1 | 1 | 1 | 1 | 1 | 1 | 1 | 1 | 1 | 1 | 1 | 1 | 1 | 1 | 1 | 1 | 1 | 1 | 1 | 1 | 1 | 1 |
| YPTB1940 | 1 | 1 | 1 | 1 | 1 | 1 | 1 | 1 | 1 | 1 | 1 | 1 | 1 | 1 | 1 | 1 | 1 | 85 | 1 | 1 | 1 | 1 | 1 | 1 | 1 | 1 | 1 | 1 | 1 | 1 | 1 | 1 | 1 | 1 | 1 |
| YPTB2001 | 1 | 1 | 1 | 65 | 1 | 1 | 1 | 68 | 1 | 65 | 65 | 1 | 1 | 1 | 1 | 65 | 1 | 66 | 1 | 1 | 1 | 1 | 1 | 1 | 1 | 1 | 1 | 1 | 1 | 1 | 1 | 1 | 65 | 65 | 1 |
| YPTB2005 | 1 | 113 | 1 | 115 | 1 | 1 | 1 | 114 | 113 | 115 | 115 | 1 | 114 | 114 | 1 | 115 | 114 | 111 | 1 | 1 | 1 | 1 | 1 | 1 | 145 | 145 | 1 | 1 | 1 | 1 | 1 | 145 | 115 | 115 | 1 |
| YPTB2007 | 1 | 1 | 1 | 1 | 1 | 1 | 1 | 56 | 1 | 1 | 1 | 1 | 1 | 1 | 1 | 1 | 1 | 1 | 1 | 1 | 1 | 1 | 1 | 1 | 1 | 1 | 1 | 1 | 1 | 1 | 1 | 1 | 1 | 1 | 1 |
| YPTB2008 | 1 | 95 | 1 | 93 | 1 | 1 | 1 | 93 | 110 | 93 | 93 | 92 | 92 | 92 | 1 | 93 | 92 | 93 | 1 | 1 | 1 | 1 | 1 | 1 | 92 | 92 | 1 | 1 | 1 | 1 | 1 | 92 | 93 | 93 | 1 |

| | | | | | | | | | | | | | | | | | | | | | | | | | | | | | | | | | | | |
|---|---|---|---|---|---|---|---|---|---|---|---|---|---|---|---|---|---|---|---|---|---|---|---|---|---|---|---|---|---|---|---|---|---|---|---|
| YPTB2009 | 1 | 100 | 1 | 98 | 1 | 1 | 1 | 101 | 100 | 98 | 98 | 1 | 1 | 1 | 1 | 98 | 1 | 121 | 1 | 1 | 1 | 1 | 1 | 1 | 1 | 1 | 1 | 1 | 1 | 1 | 1 | 1 | 98 | 98 | 1 |
| YPTB2019 | 1 | 102 | 1 | 129 | 1 | 1 | 1 | 103 | 102 | 1 | 129 | 1 | 1 | 1 | 1 | 1 | 1 | 1 | 1 | 1 | 1 | 1 | 1 | 1 | 1 | 1 | 1 | 1 | 1 | 1 | 1 | 1 | 129 | 129 | 1 |
| YPTB2023 | 1 | 1 | 1 | 1 | 1 | 1 | 1 | 1 | 1 | 1 | 1 | 1 | 1 | 1 | 1 | 1 | 1 | 1 | 1 | 1 | 1 | 1 | 1 | 1 | 1 | 1 | 1 | 1 | 1 | 1 | 1 | 1 | 1 | 1 | 1 |
| YPTB2032 | 1 | 108 | 1 | 108 | 1 | 1 | 1 | 110 | 108 | 108 | 108 | 1 | 1 | 1 | 1 | 108 | 1 | 108 | 1 | 1 | 1 | 1 | 1 | 1 | 1 | 1 | 1 | 1 | 1 | 1 | 1 | 1 | 108 | 108 | 1 |
| YPTB2038 | 1 | 103 | 1 | 105 | 1 | 1 | 1 | 103 | 103 | 105 | 105 | 1 | 1 | 1 | 1 | 105 | 1 | 1 | 1 | 1 | 1 | 1 | 1 | 1 | 1 | 1 | 1 | 1 | 1 | 1 | 1 | 1 | 105 | 105 | 1 |
| YPTB2042 | 97 | 92 | 97 | 94 | 1 | 1 | 97 | 97 | 92 | 94 | 94 | 1 | 1 | 1 | 1 | 94 | 1 | 94 | 93 | 93 | 1 | 1 | 1 | 93 | 1 | 1 | 1 | 1 | 1 | 1 | 1 | 1 | 94 | 94 | 97 |
| YPTB2045 | 1 | 1 | 1 | 156 | 1 | 1 | 1 | 124 | 1 | 121 | 156 | 1 | 1 | 1 | 1 | 121 | 1 | 121 | 1 | 1 | 1 | 1 | 1 | 1 | 395 | 1 | 1 | 1 | 1 | 1 | 1 | 1 | 156 | 156 | 1 |
| YPTB2047 | 121 | 121 | 121 | 121 | 1 | 122 | 121 | 121 | 121 | 121 | 121 | 122 | 122 | 122 | 122 | 121 | 122 | 121 | 121 | 121 | 122 | 122 | 122 | 121 | 122 | 122 | 122 | 122 | 122 | 122 | 122 | 122 | 121 | 121 | 121 |
| YPTB2053 | 1 | 75 | 1 | 75 | 1 | 1 | 1 | 1 | 75 | 75 | 75 | 1 | 1 | 1 | 1 | 75 | 1 | 1 | 1 | 1 | 1 | 1 | 1 | 1 | 1 | 1 | 1 | 1 | 1 | 1 | 1 | 1 | 75 | 75 | 1 |
| YPTB2059 | 1 | 1 | 1 | 1 | 1 | 1 | 1 | 1 | 1 | 1 | 1 | 1 | 1 | 1 | 1 | 1 | 1 | 1 | 1 | 1 | 1 | 1 | 1 | 1 | 1 | 1 | 1 | 1 | 1 | 1 | 1 | 1 | 1 | 1 | 1 |
| YPTB2060 | 1 | 1 | 1 | 1 | 1 | 1 | 1 | 1 | 1 | 1 | 1 | 1 | 1 | 1 | 1 | 1 | 1 | 1 | 1 | 1 | 1 | 1 | 1 | 1 | 1 | 1 | 1 | 1 | 1 | 1 | 1 | 1 | 1 | 1 | 1 |
| YPTB2064 | 1 | 207 | 1 | 86 | 1 | 1 | 1 | 1 | 86 | 86 | 86 | 1 | 1 | 1 | 1 | 86 | 1 | 86 | 1 | 1 | 1 | 1 | 1 | 1 | 1 | 1 | 1 | 1 | 1 | 1 | 1 | 1 | 86 | 86 | 1 |
| YPTB2083 | 106 | 88 | 85 | 1 | 1 | 1 | 85 | 1 | 88 | 1 | 1 | 1 | 1 | 1 | 1 | 1 | 1 | 1 | 85 | 85 | 1 | 1 | 1 | 85 | 1 | 1 | 1 | 1 | 1 | 1 | 1 | 1 | 1 | 1 | 85 |
| YPTB2084 | 76 | 76 | 76 | 1 | 1 | 1 | 76 | 1 | 76 | 77 | 1 | 1 | 1 | 1 | 1 | 77 | 1 | 1 | 76 | 76 | 1 | 1 | 1 | 76 | 1 | 1 | 1 | 1 | 1 | 1 | 1 | 1 | 1 | 1 | 76 |
| YPTB2089 | 98 | 1 | 98 | 97 | 1 | 1 | 98 | 1 | 1 | 97 | 97 | 97 | 1 | 1 | 1 | 97 | 1 | 99 | 98 | 98 | 1 | 1 | 1 | 98 | 1 | 1 | 1 | 1 | 1 | 1 | 1 | 1 | 97 | 97 | 98 |
| YPTB2090 | 114 | 116 | 114 | 114 | 1 | 1 | 114 | 117 | 116 | 114 | 114 | 113 | 1 | 1 | 1 | 114 | 1 | 113 | 114 | 114 | 1 | 1 | 1 | 114 | 1 | 1 | 1 | 1 | 1 | 1 | 1 | 1 | 114 | 114 | 114 |
| YPTB2116 | 1 | 1 | 1 | 1 | 1 | 1 | 1 | 1 | 1 | 1 | 1 | 1 | 1 | 1 | 1 | 1 | 1 | 1 | 1 | 1 | 1 | 1 | 1 | 1 | 1 | 1 | 1 | 1 | 1 | 1 | 1 | 1 | 1 | 1 | 1 |
| YPTB2125 | 116 | 116 | 116 | 116 | 1 | 1 | 116 | 1 | 116 | 116 | 116 | 115 | 1 | 1 | 1 | 116 | 1 | 1 | 116 | 116 | 1 | 1 | 1 | 116 | 1 | 1 | 1 | 1 | 1 | 1 | 1 | 1 | 116 | 116 | 116 |
| YPTB2126 | 1 | 118 | 1 | 119 | 1 | 1 | 1 | 1 | 118 | 119 | 119 | 118 | 1 | 1 | 1 | 119 | 1 | 1 | 1 | 1 | 1 | 1 | 1 | 1 | 1 | 1 | 1 | 1 | 1 | 1 | 1 | 1 | 119 | 119 | 1 |
| YPTB2137 | 112 | 141 | 112 | 112 | 1 | 1 | 112 | 112 | 141 | 112 | 112 | 1 | 1 | 1 | 1 | 112 | 1 | 113 | 112 | 112 | 1 | 1 | 1 | 112 | 1 | 1 | 1 | 1 | 1 | 1 | 1 | 1 | 112 | 112 | 112 |
| YPTB2139 | 51 | 1 | 51 | 1 | 1 | 1 | 51 | 1 | 1 | 1 | 1 | 1 | 1 | 1 | 1 | 1 | 1 | 1 | 51 | 51 | 1 | 1 | 1 | 51 | 1 | 1 | 1 | 1 | 1 | 1 | 1 | 1 | 1 | 1 | 51 |
| YPTB2144 | 1 | 1 | 1 | 1 | 1 | 1 | 1 | 1 | 1 | 1 | 1 | 1 | 1 | 1 | 1 | 1 | 1 | 1 | 1 | 1 | 1 | 1 | 1 | 1 | 1 | 1 | 1 | 1 | 1 | 1 | 1 | 1 | 1 | 1 | 1 |
| YPTB2146 | 1 | 1 | 1 | 1 | 1 | 1 | 1 | 55 | 1 | 1 | 1 | 1 | 1 | 1 | 1 | 1 | 1 | 1 | 1 | 1 | 1 | 1 | 1 | 1 | 1 | 1 | 1 | 1 | 1 | 1 | 1 | 1 | 1 | 1 | 1 |
| YPTB2147 | 117 | 1 | 117 | 1 | 1 | 1 | 117 | 119 | 352 | 1 | 1 | 1 | 1 | 154 | 1 | 1 | 154 | 117 | 117 | 117 | 1 | 1 | 1 | 117 | 1 | 1 | 1 | 1 | 1 | 1 | 1 | 1 | 1 | 1 | 117 |
| YPTB2156 | 130 | 134 | 170 | 174 | 129 | 129 | 170 | 138 | 134 | 136 | 174 | 129 | 129 | 129 | 129 | 136 | 129 | 131 | 130 | 130 | 129 | 129 | 129 | 130 | 129 | 129 | 129 | 129 | 129 | 129 | 129 | 129 | 174 | 174 | 130 |
| YPTB2160 | 1 | 96 | 1 | 97 | 1 | 1 | 1 | 99 | 96 | 97 | 97 | 93 | 1 | 1 | 1 | 97 | 1 | 1 | 1 | 1 | 1 | 1 | 1 | 1 | 1 | 1 | 1 | 1 | 1 | 1 | 1 | 1 | 97 | 97 | 1 |
| YPTB2167 | 1 | 1 | 1 | 1 | 1 | 1 | 1 | 1 | 1 | 1 | 1 | 1 | 1 | 1 | 1 | 1 | 1 | 1 | 1 | 1 | 1 | 1 | 1 | 1 | 1 | 1 | 1 | 1 | 1 | 1 | 1 | 1 | 1 | 1 | 1 |
| YPTB2261 | 1 | 1 | 1 | 78 | 1 | 1 | 1 | 1 | 1 | 78 | 78 | 1 | 77 | 77 | 1 | 78 | 77 | 1 | 1 | 1 | 1 | 1 | 1 | 1 | 77 | 77 | 1 | 1 | 1 | 1 | 1 | 77 | 78 | 78 | 1 |
| YPTB2263 | 1 | 86 | 1 | 85 | 1 | 1 | 1 | 87 | 86 | 85 | 85 | 84 | 84 | 84 | 1 | 85 | 84 | 1 | 1 | 1 | 1 | 1 | 1 | 1 | 84 | 84 | 1 | 1 | 1 | 1 | 1 | 84 | 85 | 85 | 1 |
| YPTB2266 | 1 | 159 | 1 | 120 | 1 | 1 | 1 | 121 | 159 | 120 | 120 | 115 | 115 | 115 | 1 | 120 | 115 | 117 | 1 | 1 | 1 | 1 | 1 | 1 | 115 | 115 | 1 | 1 | 1 | 1 | 1 | 115 | 120 | 120 | 1 |
| YPTB2273 | 106 | 109 | 106 | 106 | 1 | 1 | 106 | 106 | 109 | 106 | 106 | 107 | 1 | 1 | 1 | 106 | 1 | 141 | 106 | 106 | 1 | 1 | 1 | 106 | 1 | 1 | 1 | 1 | 1 | 1 | 1 | 1 | 106 | 106 | 106 |
| YPTB2274 | 116 | 1 | 116 | 116 | 1 | 1 | 116 | 116 | 1 | 116 | 116 | 116 | 1 | 1 | 1 | 116 | 1 | 117 | 116 | 116 | 1 | 1 | 1 | 116 | 1 | 1 | 1 | 1 | 1 | 1 | 1 | 1 | 116 | 116 | 116 |
| YPTB2275 | 1 | 1 | 1 | 100 | 1 | 1 | 1 | 1 | 1 | 99 | 100 | 98 | 1 | 1 | 1 | 99 | 1 | 1 | 1 | 1 | 1 | 1 | 1 | 1 | 1 | 1 | 1 | 1 | 1 | 1 | 1 | 1 | 100 | 100 | 1 |
| YPTB2283 | 1 | 113 | 1 | 112 | 1 | 1 | 1 | 116 | 113 | 114 | 112 | 1 | 1 | 1 | 1 | 114 | 1 | 110 | 1 | 1 | 1 | 1 | 1 | 1 | 1 | 1 | 1 | 1 | 1 | 1 | 1 | 1 | 112 | 112 | 1 |
| YPTB2312 | 104 | 107 | 104 | 1 | 1 | 1 | 104 | 1 | 107 | 1 | 1 | 1 | 1 | 1 | 1 | 1 | 1 | 105 | 104 | 104 | 1 | 1 | 1 | 104 | 1 | 1 | 1 | 1 | 1 | 1 | 1 | 1 | 1 | 1 | 290 |
| YPTB2322 | 118 | 117 | 118 | 117 | 1 | 1 | 118 | 117 | 117 | 117 | 117 | 1 | 1 | 1 | 1 | 117 | 1 | 118 | 118 | 118 | 1 | 1 | 1 | 118 | 1 | 1 | 1 | 1 | 1 | 1 | 1 | 1 | 117 | 117 | 118 |
| YPTB2323 | 1 | 1 | 1 | 75 | 1 | 1 | 1 | 1 | 1 | 75 | 75 | 1 | 1 | 1 | 1 | 75 | 1 | 1 | 1 | 1 | 1 | 1 | 1 | 1 | 1 | 1 | 1 | 1 | 1 | 1 | 1 | 1 | 75 | 75 | 1 |
| YPTB2324 | 71 | 1 | 86 | 1 | 1 | 1 | 86 | 1 | 1 | 1 | 1 | 1 | 1 | 1 | 1 | 1 | 1 | 71 | 71 | 71 | 1 | 1 | 1 | 71 | 1 | 1 | 1 | 1 | 1 | 1 | 1 | 1 | 1 | 1 | 71 |
| YPTB2336 | 131 | 134 | 131 | 139 | 1 | 1 | 131 | 137 | 134 | 136 | 139 | 1 | 1 | 176 | 1 | 136 | 176 | 177 | 131 | 131 | 1 | 1 | 1 | 131 | 1 | 1 | 1 | 1 | 1 | 1 | 1 | 1 | 139 | 139 | 131 |
| YPTB2337 | 94 | 95 | 94 | 94 | 1 | 1 | 94 | 94 | 95 | 94 | 94 | 1 | 1 | 1 | 1 | 94 | 1 | 94 | 94 | 94 | 1 | 1 | 1 | 94 | 1 | 1 | 1 | 1 | 1 | 1 | 1 | 1 | 94 | 94 | 94 |
| YPTB2339 | 1 | 1 | 1 | 1 | 1 | 1 | 1 | 1 | 1 | 1 | 1 | 1 | 1 | 1 | 1 | 1 | 1 | 1 | 1 | 1 | 1 | 1 | 1 | 1 | 1 | 1 | 1 | 1 | 1 | 1 | 1 | 1 | 1 | 1 | 1 |
| YPTB2340 | 1 | 1 | 1 | 1 | 1 | 1 | 1 | 1 | 1 | 1 | 1 | 1 | 1 | 1 | 1 | 1 | 1 | 1 | 1 | 1 | 1 | 1 | 1 | 1 | 1 | 1 | 1 | 1 | 1 | 1 | 1 | 1 | 1 | 1 | 78 |
| YPTB2341 | 1 | 1 | 1 | 1 | 1 | 1 | 1 | 1 | 1 | 1 | 1 | 1 | 1 | 1 | 1 | 1 | 1 | 1 | 1 | 1 | 1 | 1 | 1 | 1 | 1 | 1 | 1 | 1 | 1 | 1 | 1 | 1 | 1 | 1 | 1 |
| YPTB2353 | 1 | 1 | 1 | 1 | 1 | 1 | 1 | 1 | 1 | 1 | 1 | 79 | 1 | 1 | 1 | 1 | 1 | 79 | 1 | 1 | 1 | 1 | 1 | 1 | 1 | 1 | 1 | 1 | 1 | 1 | 1 | 1 | 1 | 1 | 1 |
| YPTB2354 | 1 | 1 | 1 | 87 | 1 | 1 | 1 | 87 | 1 | 87 | 87 | 87 | 1 | 1 | 1 | 87 | 1 | 87 | 1 | 1 | 1 | 1 | 1 | 1 | 1 | 1 | 1 | 1 | 1 | 1 | 1 | 1 | 87 | 87 | 1 |
| YPTB2389 | 1 | 1 | 1 | 1 | 1 | 1 | 1 | 1 | 1 | 1 | 1 | 1 | 1 | 1 | 1 | 1 | 1 | 1 | 1 | 1 | 1 | 1 | 1 | 1 | 1 | 1 | 1 | 1 | 1 | 1 | 1 | 1 | 1 | 1 | 1 |
| YPTB2396 | 1 | 81 | 1 | 98 | 1 | 1 | 1 | 1 | 81 | 1 | 98 | 1 | 1 | 1 | 1 | 1 | 1 | 1 | 1 | 1 | 1 | 1 | 1 | 1 | 1 | 1 | 1 | 1 | 1 | 1 | 1 | 1 | 98 | 98 | 1 |
| YPTB2397 | 1 | 1 | 1 | 1 | 1 | 1 | 1 | 1 | 1 | 1 | 1 | 1 | 1 | 1 | 1 | 1 | 1 | 1 | 1 | 1 | 1 | 1 | 1 | 1 | 1 | 1 | 1 | 1 | 1 | 1 | 1 | 1 | 1 | 1 | 1 |
| YPTB2427 | 167 | 119 | 167 | 117 | 1 | 1 | 167 | 168 | 119 | 117 | 117 | 114 | 170 | 1 | 1 | 117 | 1 | 125 | 167 | 167 | 1 | 1 | 1 | 167 | 1 | 1 | 1 | 1 | 1 | 1 | 1 | 170 | 117 | 117 | 167 |
| YPTB2432 | 1 | 1 | 1 | 118 | 1 | 1 | 1 | 1 | 1 | 118 | 118 | 1 | 1 | 1 | 1 | 118 | 1 | 116 | 1 | 1 | 1 | 1 | 1 | 1 | 1 | 1 | 1 | 1 | 1 | 1 | 1 | 1 | 118 | 118 | 1 |
| YPTB2442 | 111 | 114 | 111 | 116 | 1 | 111 | 111 | 118 | 141 | 116 | 116 | 110 | 1 | 1 | 111 | 116 | 1 | 111 | 111 | 111 | 111 | 111 | 111 | 111 | 1 | 1 | 111 | 111 | 111 | 111 | 111 | 1 | 116 | 116 | 111 |
| YPTB2447 | 117 | 116 | 113 | 115 | 1 | 1 | 113 | 114 | 116 | 115 | 115 | 113 | 1 | 1 | 1 | 115 | 1 | 114 | 113 | 113 | 1 | 1 | 1 | 113 | 1 | 1 | 1 | 1 | 1 | 1 | 1 | 1 | 115 | 115 | 113 |
| YPTB2448 | 85 | 86 | 85 | 85 | 1 | 1 | 85 | 86 | 86 | 85 | 85 | 85 | 1 | 1 | 1 | 85 | 1 | 86 | 85 | 85 | 1 | 1 | 1 | 85 | 1 | 1 | 1 | 1 | 1 | 1 | 1 | 1 | 85 | 85 | 85 |
| YPTB2467 | 143 | 109 | 1 | 142 | 1 | 1 | 1 | 109 | 109 | 109 | 142 | 106 | 1 | 1 | 1 | 109 | 1 | 109 | 1 | 1 | 1 | 1 | 1 | 1 | 1 | 1 | 1 | 1 | 1 | 1 | 1 | 1 | 142 | 142 | 1 |
| YPTB2470 | 1 | 1 | 1 | 1 | 1 | 1 | 1 | 1 | 1 | 1 | 1 | 18 | 1 | 1 | 1 | 1 | 1 | 1 | 1 | 1 | 1 | 1 | 1 | 1 | 1 | 1 | 1 | 1 | 1 | 1 | 1 | 1 | 1 | 1 | 1 |
| YPTB2471 | 63 | 1 | 63 | 1 | 1 | 1 | 63 | 66 | 1 | 1 | 1 | 1 | 1 | 1 | 1 | 1 | 1 | 1 | 63 | 63 | 1 | 1 | 1 | 63 | 1 | 1 | 1 | 1 | 1 | 1 | 1 | 1 | 1 | 1 | 63 |
| YPTB2474 | 86 | 88 | 86 | 94 | 1 | 1 | 86 | 92 | 88 | 90 | 94 | 1 | 1 | 1 | 1 | 90 | 1 | 87 | 86 | 86 | 1 | 1 | 1 | 86 | 1 | 1 | 1 | 1 | 1 | 1 | 1 | 1 | 94 | 94 | 86 |
| YPTB2475 | 19 | 19 | 19 | 20 | 1 | 1 | 19 | 1 | 19 | 20 | 20 | 1 | 1 | 1 | 1 | 20 | 1 | 20 | 19 | 19 | 1 | 1 | 1 | 19 | 1 | 1 | 1 | 1 | 1 | 1 | 1 | 1 | 20 | 20 | 19 |
| YPTB2549 | 1 | 79 | 100 | 77 | 1 | 1 | 1 | 1 | 79 | 77 | 77 | 1 | 82 | 82 | 1 | 77 | 82 | 77 | 1 | 1 | 1 | 1 | 1 | 1 | 82 | 82 | 1 | 1 | 1 | 1 | 1 | 82 | 77 | 77 | 1 |
| YPTB2557 | 123 | 124 | 123 | 124 | 1 | 382 | 123 | 124 | 124 | 124 | 124 | 122 | 124 | 124 | 382 | 124 | 124 | 124 | 123 | 123 | 382 | 382 | 382 | 123 | 124 | 124 | 382 | 382 | 382 | 382 | 382 | 124 | 124 | 124 | 123 |
| YPTB2558 | 100 | 102 | 100 | 103 | 1 | 100 | 100 | 105 | 102 | 103 | 103 | 99 | 99 | 99 | 100 | 103 | 99 | 1 | 100 | 100 | 100 | 100 | 100 | 100 | 99 | 99 | 100 | 100 | 100 | 100 | 100 | 99 | 103 | 103 | 100 |
| YPTB2577 | 137 | 141 | 137 | 143 | 1 | 1 | 137 | 144 | 184 | 143 | 143 | 1 | 1 | 1 | 1 | 143 | 1 | 187 | 137 | 137 | 1 | 1 | 1 | 137 | 1 | 1 | 1 | 1 | 1 | 1 | 1 | 1 | 143 | 143 | 137 |
| YPTB2578 | 1 | 1 | 1 | 1 | 1 | 1 | 1 | 1 | 1 | 1 | 1 | 1 | 1 | 1 | 1 | 1 | 1 | 1 | 1 | 1 | 1 | 1 | 1 | 1 | 1 | 1 | 1 | 1 | 1 | 1 | 1 | 1 | 1 | 1 | 1 |
| YPTB2580 | 1 | 1 | 1 | 1 | 1 | 1 | 1 | 1 | 1 | 1 | 1 | 1 | 1 | 1 | 1 | 1 | 1 | 1 | 1 | 1 | 1 | 1 | 1 | 1 | 1 | 1 | 1 | 1 | 1 | 1 | 1 | 1 | 1 | 1 | 1 |
| YPTB2581 | 97 | 100 | 97 | 101 | 1 | 1 | 97 | 102 | 100 | 101 | 101 | 1 | 1 | 1 | 1 | 101 | 1 | 97 | 97 | 97 | 1 | 1 | 1 | 97 | 1 | 1 | 1 | 1 | 1 | 1 | 1 | 1 | 101 | 101 | 97 |
| YPTB2585 | 120 | 123 | 120 | 125 | 1 | 1 | 120 | 127 | 123 | 125 | 125 | 119 | 1 | 1 | 1 | 125 | 1 | 120 | 120 | 120 | 1 | 1 | 1 | 120 | 1 | 1 | 1 | 1 | 1 | 1 | 1 | 1 | 125 | 125 | 120 |
| YPTB2594 | 1 | 63 | 1 | 1 | 1 | 1 | 1 | 63 | 63 | 1 | 1 | 63 | 1 | 1 | 1 | 1 | 1 | 1 | 1 | 1 | 1 | 1 | 1 | 1 | 1 | 1 | 1 | 1 | 1 | 1 | 1 | 1 | 1 | 1 | 1 |
| YPTB2595 | 1 | 1 | 1 | 1 | 1 | 1 | 1 | 1 | 1 | 1 | 1 | 1 | 1 | 1 | 1 | 1 | 1 | 1 | 1 | 1 | 1 | 1 | 1 | 1 | 1 | 1 | 1 | 1 | 1 | 1 | 1 | 1 | 1 | 1 | 1 |
| YPTB2597 | 92 | 95 | 92 | 1 | 1 | 1 | 92 | 96 | 95 | 1 | 1 | 93 | 1 | 1 | 1 | 1 | 1 | 1 | 92 | 92 | 1 | 1 | 1 | 92 | 1 | 1 | 1 | 1 | 1 | 1 | 1 | 1 | 1 | 1 | 92 |
| YPTB2600 | 78 | 79 | 78 | 1 | 1 | 1 | 78 | 1 | 79 | 1 | 1 | 1 | 1 | 1 | 1 | 1 | 1 | 1 | 78 | 78 | 1 | 1 | 1 | 78 | 1 | 1 | 1 | 1 | 1 | 1 | 1 | 1 | 1 | 1 | 78 |
| YPTB2601 | 1 | 69 | 1 | 70 | 1 | 1 | 1 | 1 | 69 | 70 | 70 | 1 | 1 | 1 | 1 | 70 | 1 | 1 | 1 | 1 | 1 | 1 | 1 | 1 | 1 | 1 | 1 | 1 | 1 | 1 | 1 | 1 | 70 | 70 | 1 |
| YPTB2602 | 109 | 112 | 109 | 113 | 1 | 1 | 109 | 114 | 112 | 113 | 113 | 1 | 109 | 109 | 1 | 113 | 109 | 110 | 109 | 109 | 1 | 1 | 1 | 109 | 109 | 109 | 1 | 1 | 1 | 1 | 1 | 109 | 113 | 113 | 109 |
| YPTB2609 | 1 | 87 | 1 | 1 | 1 | 1 | 1 | 86 | 87 | 85 | 1 | 1 | 1 | 1 | 1 | 85 | 1 | 1 | 1 | 1 | 1 | 1 | 1 | 1 | 1 | 1 | 1 | 1 | 1 | 1 | 1 | 1 | 1 | 1 | 1 |
| YPTB2612 | 101 | 105 | 101 | 107 | 1 | 1 | 101 | 109 | 105 | 107 | 107 | 1 | 140 | 140 | 1 | 107 | 140 | 103 | 101 | 101 | 1 | 1 | 1 | 101 | 140 | 140 | 1 | 1 | 1 | 1 | 1 | 140 | 107 | 107 | 101 |
| YPTB2613 | 1 | 1 | 1 | 1 | 1 | 1 | 1 | 1 | 1 | 1 | 1 | 1 | 1 | 1 | 1 | 1 | 1 | 1 | 1 | 1 | 1 | 1 | 1 | 1 | 1 | 1 | 1 | 1 | 1 | 1 | 1 | 1 | 1 | 1 | 1 |
| YPTB2617 | 88 | 1 | 88 | 1 | 1 | 1 | 88 | 1 | 205 | 1 | 1 | 1 | 1 | 1 | 1 | 1 | 1 | 1 | 88 | 88 | 1 | 1 | 1 | 88 | 1 | 1 | 1 | 1 | 1 | 1 | 1 | 1 | 1 | 1 | 88 |
| YPTB2630 | 109 | 1 | 109 | 1 | 1 | 1 | 109 | 1 | 1 | 1 | 1 | 1 | 1 | 1 | 1 | 1 | 1 | 1 | 109 | 109 | 1 | 1 | 1 | 109 | 1 | 1 | 1 | 1 | 1 | 1 | 1 | 1 | 1 | 1 | 109 |
| YPTB2632 | 116 | 116 | 116 | 116 | 1 | 1 | 116 | 119 | 116 | 116 | 116 | 116 | 1 | 1 | 1 | 116 | 1 | 117 | 116 | 116 | 1 | 1 | 1 | 116 | 1 | 1 | 1 | 1 | 1 | 1 | 1 | 1 | 116 | 116 | 116 |
| YPTB2637 | 1 | 1 | 1 | 1 | 1 | 1 | 1 | 1 | 1 | 1 | 1 | 106 | 1 | 1 | 1 | 1 | 1 | 1 | 1 | 1 | 1 | 1 | 1 | 1 | 1 | 1 | 1 | 1 | 1 | 1 | 1 | 1 | 1 | 1 | 1 |
| YPTB2640 | 1 | 95 | 1 | 95 | 1 | 1 | 1 | 95 | 95 | 95 | 95 | 96 | 1 | 1 | 1 | 95 | 1 | 95 | 1 | 1 | 1 | 1 | 1 | 1 | 1 | 1 | 1 | 1 | 1 | 1 | 1 | 1 | 95 | 95 | 1 |
| YPTB2714 | 1 | 98 | 1 | 96 | 1 | 1 | 1 | 97 | 98 | 96 | 96 | 1 | 1 | 1 | 1 | 96 | 1 | 97 | 1 | 1 | 1 | 1 | 1 | 1 | 1 | 1 | 1 | 1 | 1 | 1 | 1 | 1 | 96 | 96 | 1 |
| YPTB2715 | 1 | 1 | 1 | 1 | 1 | 1 | 1 | 1 | 1 | 1 | 1 | 1 | 1 | 1 | 1 | 1 | 1 | 1 | 1 | 1 | 1 | 1 | 1 | 1 | 1 | 1 | 1 | 1 | 1 | 1 | 1 | 1 | 1 | 1 | 1 |
| YPTB2716 | 107 | 109 | 107 | 108 | 1 | 1 | 107 | 108 | 140 | 108 | 108 | 106 | 1 | 1 | 1 | 108 | 1 | 108 | 107 | 107 | 1 | 1 | 1 | 107 | 1 | 1 | 1 | 1 | 1 | 1 | 1 | 1 | 108 | 108 | 107 |
| YPTB2761 | 1 | 49 | 1 | 48 | 1 | 1 | 1 | 1 | 49 | 48 | 48 | 48 | 1 | 1 | 1 | 48 | 1 | 1 | 1 | 1 | 1 | 1 | 1 | 1 | 1 | 1 | 1 | 1 | 1 | 1 | 1 | 1 | 48 | 48 | 1 |
| YPTB2775 | 1 | 112 | 1 | 114 | 1 | 1 | 1 | 112 | 112 | 114 | 114 | 109 | 1 | 1 | 1 | 114 | 1 | 108 | 1 | 1 | 1 | 1 | 1 | 1 | 1 | 1 | 1 | 1 | 1 | 1 | 1 | 1 | 114 | 114 | 1 |
| YPTB2779 | 1 | 1 | 1 | 1 | 1 | 1 | 1 | 1 | 1 | 1 | 1 | 78 | 1 | 1 | 1 | 1 | 1 | 1 | 1 | 1 | 1 | 1 | 1 | 1 | 1 | 1 | 1 | 1 | 1 | 1 | 1 | 1 | 1 | 1 | 1 |
| YPTB2781 | 1 | 1 | 1 | 1 | 1 | 1 | 1 | 86 | 1 | 1 | 1 | 1 | 84 | 84 | 1 | 1 | 84 | 1 | 1 | 1 | 1 | 1 | 1 | 1 | 84 | 84 | 1 | 1 | 1 | 1 | 1 | 84 | 1 | 1 | 1 |
| YPTB2785 | 1 | 74 | 75 | 1 | 1 | 1 | 75 | 1 | 74 | 1 | 1 | 1 | 1 | 1 | 1 | 1 | 1 | 74 | 75 | 75 | 1 | 1 | 1 | 75 | 1 | 1 | 1 | 1 | 1 | 1 | 1 | 1 | 1 | 1 | 75 |
| YPTB2789 | 1 | 102 | 1 | 102 | 1 | 1 | 1 | 103 | 102 | 102 | 102 | 102 | 102 | 102 | 1 | 102 | 102 | 1 | 1 | 1 | 1 | 1 | 1 | 1 | 102 | 102 | 1 | 1 | 1 | 1 | 1 | 102 | 102 | 102 | 1 |
| YPTB2794 | 79 | 81 | 79 | 81 | 1 | 1 | 79 | 81 | 81 | 81 | 81 | 79 | 79 | 79 | 1 | 81 | 79 | 1 | 79 | 79 | 1 | 1 | 1 | 79 | 79 | 79 | 1 | 1 | 1 | 1 | 1 | 79 | 81 | 81 | 79 |
| YPTB2797 | 67 | 68 | 67 | 68 | 1 | 1 | 67 | 69 | 68 | 68 | 68 | 1 | 1 | 1 | 1 | 68 | 1 | 1 | 67 | 67 | 1 | 1 | 1 | 67 | 1 | 1 | 1 | 1 | 1 | 1 | 1 | 1 | 68 | 68 | 67 |
| YPTB2799 | 96 | 95 | 96 | 1 | 1 | 1 | 96 | 99 | 95 | 1 | 1 | 96 | 1 | 1 | 1 | 1 | 1 | 1 | 96 | 96 | 1 | 1 | 1 | 96 | 1 | 1 | 1 | 1 | 1 | 1 | 1 | 1 | 1 | 1 | 96 |
| YPTB2819 | 1 | 1 | 1 | 1 | 1 | 125 | 1 | 1 | 1 | 1 | 1 | 1 | 1 | 1 | 125 | 1 | 1 | 1 | 1 | 1 | 125 | 125 | 125 | 1 | 1 | 1 | 125 | 125 | 125 | 125 | 125 | 1 | 1 | 1 | 1 |
| YPTB2838 | 105 | 1 | 105 | 1 | 1 | 1 | 105 | 1 | 1 | 1 | 1 | 105 | 1 | 1 | 1 | 1 | 1 | 105 | 105 | 145 | 1 | 1 | 1 | 105 | 1 | 1 | 1 | 1 | 1 | 1 | 1 | 1 | 1 | 1 | 105 |
| YPTB2853 | 1 | 1 | 1 | 1 | 1 | 1 | 1 | 1 | 1 | 1 | 1 | 1 | 1 | 1 | 1 | 1 | 1 | 1 | 1 | 1 | 1 | 1 | 1 | 1 | 1 | 1 | 1 | 1 | 1 | 1 | 1 | 1 | 1 | 1 | 1 |
| YPTB2857 | 1 | 1 | 1 | 1 | 1 | 1 | 1 | 1 | 1 | 1 | 1 | 1 | 1 | 1 | 1 | 1 | 1 | 1 | 1 | 1 | 1 | 1 | 1 | 1 | 1 | 1 | 1 | 1 | 1 | 1 | 1 | 1 | 1 | 1 | 1 |
| YPTB2859 | 1 | 107 | 1 | 111 | 1 | 1 | 1 | 1 | 107 | 107 | 111 | 106 | 1 | 1 | 1 | 107 | 1 | 107 | 1 | 1 | 1 | 1 | 1 | 1 | 1 | 1 | 1 | 1 | 1 | 1 | 1 | 1 | 111 | 111 | 1 |
| YPTB2869 | 1 | 117 | 1 | 118 | 1 | 1 | 1 | 118 | 117 | 118 | 118 | 115 | 1 | 1 | 1 | 118 | 1 | 1 | 1 | 1 | 1 | 1 | 1 | 1 | 1 | 1 | 1 | 1 | 1 | 1 | 1 | 1 | 118 | 118 | 1 |
| YPTB2872 | 1 | 69 | 1 | 69 | 1 | 1 | 1 | 1 | 69 | 69 | 69 | 1 | 1 | 1 | 1 | 69 | 1 | 1 | 1 | 1 | 1 | 1 | 1 | 1 | 1 | 1 | 1 | 1 | 1 | 1 | 1 | 1 | 69 | 69 | 1 |
| YPTB2873 | 1 | 130 | 1 | 130 | 1 | 1 | 1 | 133 | 175 | 132 | 130 | 1 | 1 | 1 | 1 | 132 | 1 | 127 | 1 | 1 | 1 | 1 | 1 | 1 | 1 | 1 | 1 | 1 | 1 | 1 | 1 | 1 | 130 | 130 | 1 |
| YPTB2885 | 1 | 77 | 1 | 79 | 1 | 1 | 1 | 79 | 77 | 79 | 79 | 1 | 1 | 1 | 1 | 79 | 1 | 1 | 1 | 1 | 1 | 1 | 1 | 1 | 1 | 1 | 1 | 1 | 1 | 1 | 1 | 1 | 79 | 79 | 1 |
| YPTB2886 | 69 | 1 | 69 | 1 | 1 | 1 | 69 | 69 | 1 | 1 | 1 | 69 | 1 | 1 | 1 | 1 | 1 | 1 | 69 | 69 | 1 | 1 | 1 | 69 | 1 | 1 | 1 | 1 | 1 | 1 | 1 | 1 | 1 | 1 | 69 |
| YPTB2887 | 93 | 91 | 93 | 94 | 1 | 1 | 93 | 92 | 91 | 94 | 94 | 92 | 1 | 1 | 1 | 94 | 1 | 94 | 93 | 93 | 1 | 1 | 1 | 93 | 1 | 1 | 1 | 1 | 1 | 1 | 1 | 1 | 94 | 94 | 93 |
| YPTB2889 | 100 | 101 | 100 | 100 | 1 | 1 | 100 | 102 | 101 | 100 | 100 | 100 | 1 | 1 | 1 | 100 | 1 | 100 | 100 | 100 | 1 | 1 | 1 | 100 | 1 | 1 | 1 | 1 | 1 | 1 | 1 | 1 | 100 | 100 | 100 |
| YPTB2890 | 1 | 1 | 59 | 1 | 1 | 1 | 59 | 1 | 1 | 1 | 1 | 1 | 1 | 1 | 1 | 1 | 1 | 1 | 59 | 59 | 1 | 1 | 1 | 59 | 1 | 1 | 1 | 1 | 1 | 1 | 1 | 1 | 1 | 1 | 59 |
| YPTB2895 | 1 | 1 | 1 | 93 | 1 | 1 | 1 | 94 | 1 | 93 | 93 | 1 | 1 | 1 | 1 | 93 | 1 | 1 | 1 | 1 | 1 | 1 | 1 | 1 | 1 | 1 | 1 | 1 | 1 | 1 | 1 | 1 | 93 | 93 | 1 |
| YPTB2897 | 1 | 1 | 1 | 1 | 1 | 1 | 1 | 1 | 1 | 1 | 1 | 1 | 1 | 1 | 1 | 1 | 1 | 1 | 1 | 1 | 1 | 1 | 1 | 1 | 1 | 1 | 1 | 1 | 1 | 1 | 1 | 1 | 1 | 1 | 1 |
| YPTB2915 | 1 | 1 | 1 | 1 | 1 | 1 | 1 | 1 | 1 | 1 | 1 | 1 | 1 | 1 | 1 | 1 | 1 | 1 | 1 | 1 | 1 | 1 | 1 | 1 | 1 | 1 | 1 | 1 | 1 | 1 | 1 | 1 | 1 | 1 | 1 |
| YPTB2929 | 1 | 70 | 1 | 70 | 1 | 1 | 1 | 1 | 70 | 70 | 70 | 68 | 1 | 1 | 1 | 70 | 1 | 68 | 1 | 1 | 1 | 1 | 1 | 1 | 1 | 1 | 1 | 1 | 1 | 1 | 1 | 1 | 70 | 70 | 1 |
| YPTB2940 | 1 | 90 | 1 | 1 | 1 | 1 | 1 | 91 | 90 | 1 | 1 | 91 | 1 | 1 | 1 | 1 | 1 | 1 | 1 | 1 | 1 | 1 | 1 | 1 | 1 | 1 | 1 | 1 | 1 | 1 | 1 | 1 | 1 | 1 | 1 |
| YPTB2942 | 1 | 126 | 1 | 128 | 1 | 436 | 1 | 122 | 126 | 128 | 128 | 122 | 1 | 1 | 436 | 128 | 1 | 123 | 1 | 1 | 436 | 436 | 436 | 1 | 1 | 1 | 436 | 436 | 436 | 436 | 436 | 1 | 128 | 128 | 1 |
| YPTB2965 | 1 | 83 | 1 | 84 | 1 | 1 | 1 | 1 | 83 | 84 | 84 | 1 | 1 | 1 | 1 | 84 | 1 | 83 | 1 | 1 | 1 | 1 | 1 | 1 | 1 | 1 | 1 | 1 | 1 | 1 | 1 | 1 | 84 | 84 | 1 |
| YPTB2966 | 1 | 87 | 1 | 1 | 1 | 1 | 1 | 1 | 87 | 1 | 1 | 1 | 1 | 1 | 1 | 1 | 1 | 1 | 1 | 1 | 1 | 1 | 1 | 1 | 1 | 1 | 1 | 1 | 1 | 1 | 1 | 1 | 1 | 1 | 110 |
| YPTB2967 | 129 | 103 | 101 | 128 | 1 | 1 | 101 | 103 | 103 | 103 | 128 | 1 | 1 | 1 | 1 | 103 | 1 | 130 | 101 | 101 | 1 | 1 | 1 | 101 | 1 | 1 | 1 | 1 | 1 | 1 | 1 | 1 | 128 | 128 | 101 |
| YPTB2973 | 99 | 101 | 99 | 129 | 1 | 1 | 99 | 101 | 101 | 101 | 129 | 1 | 1 | 1 | 1 | 101 | 1 | 1 | 99 | 99 | 1 | 1 | 1 | 99 | 1 | 1 | 1 | 1 | 1 | 1 | 1 | 1 | 129 | 129 | 99 |
| YPTB2974 | 83 | 86 | 83 | 87 | 1 | 1 | 83 | 1 | 86 | 87 | 87 | 82 | 1 | 1 | 1 | 87 | 1 | 84 | 83 | 83 | 1 | 1 | 1 | 83 | 1 | 1 | 1 | 1 | 1 | 1 | 1 | 1 | 87 | 87 | 83 |
| YPTB2975 | 1 | 1 | 1 | 86 | 1 | 1 | 1 | 87 | 1 | 86 | 86 | 86 | 1 | 1 | 1 | 86 | 1 | 87 | 1 | 1 | 1 | 1 | 1 | 1 | 1 | 1 | 1 | 1 | 1 | 1 | 1 | 1 | 86 | 86 | 1 |
| YPTB2977 | 1 | 95 | 1 | 93 | 1 | 1 | 1 | 1 | 95 | 96 | 93 | 1 | 1 | 1 | 1 | 96 | 1 | 93 | 1 | 1 | 1 | 1 | 1 | 1 | 1 | 1 | 1 | 1 | 1 | 1 | 1 | 1 | 93 | 93 | 1 |
| YPTB2978 | 1 | 122 | 1 | 122 | 1 | 1 | 1 | 122 | 122 | 122 | 122 | 122 | 1 | 1 | 1 | 122 | 1 | 122 | 123 | 123 | 1 | 1 | 1 | 123 | 1 | 1 | 1 | 1 | 1 | 1 | 1 | 1 | 122 | 122 | 1 |
| YPTB2987 | 86 | 1 | 86 | 94 | 1 | 1 | 86 | 1 | 1 | 89 | 94 | 85 | 1 | 1 | 1 | 89 | 1 | 87 | 86 | 86 | 1 | 1 | 1 | 86 | 1 | 1 | 1 | 1 | 1 | 1 | 1 | 1 | 94 | 94 | 86 |
| YPTB2993 | 1 | 1 | 1 | 100 | 1 | 1 | 1 | 1 | 1 | 100 | 100 | 1 | 1 | 1 | 1 | 100 | 1 | 98 | 1 | 1 | 1 | 1 | 1 | 1 | 1 | 1 | 1 | 1 | 1 | 1 | 1 | 1 | 100 | 100 | 1 |
| YPTB2994 | 1 | 1 | 1 | 1 | 1 | 1 | 1 | 1 | 1 | 1 | 1 | 1 | 1 | 1 | 1 | 1 | 1 | 1 | 1 | 1 | 1 | 1 | 1 | 1 | 1 | 1 | 1 | 1 | 1 | 1 | 1 | 1 | 1 | 1 | 1 |
| YPTB2997 | 81 | 81 | 81 | 81 | 1 | 1 | 81 | 81 | 81 | 81 | 81 | 82 | 1 | 1 | 1 | 81 | 1 | 81 | 81 | 81 | 1 | 1 | 1 | 81 | 1 | 1 | 1 | 1 | 1 | 1 | 1 | 1 | 81 | 81 | 81 |
| YPTB2998 | 1 | 1 | 1 | 1 | 1 | 1 | 1 | 1 | 1 | 1 | 1 | 1 | 1 | 1 | 1 | 1 | 1 | 72 | 1 | 1 | 1 | 1 | 1 | 1 | 1 | 1 | 1 | 1 | 1 | 1 | 1 | 1 | 1 | 1 | 1 |
| YPTB3003 | 55 | 55 | 55 | 1 | 1 | 1 | 55 | 1 | 55 | 1 | 1 | 1 | 1 | 1 | 1 | 1 | 1 | 55 | 55 | 55 | 1 | 1 | 1 | 55 | 1 | 1 | 1 | 1 | 1 | 1 | 1 | 1 | 1 | 1 | 55 |
| YPTB3012 | 113 | 116 | 113 | 111 | 1 | 1 | 113 | 112 | 111 | 111 | 111 | 112 | 1 | 1 | 1 | 111 | 1 | 113 | 113 | 113 | 1 | 1 | 1 | 113 | 1 | 1 | 1 | 1 | 1 | 1 | 1 | 1 | 111 | 111 | 147 |
| YPTB3017 | 1 | 1 | 1 | 1 | 1 | 1 | 1 | 1 | 1 | 1 | 1 | 1 | 1 | 1 | 1 | 1 | 1 | 84 | 1 | 1 | 1 | 1 | 1 | 1 | 1 | 1 | 1 | 1 | 1 | 1 | 1 | 1 | 1 | 1 | 1 |
| YPTB3023 | 1 | 119 | 1 | 158 | 1 | 1 | 1 | 120 | 119 | 119 | 158 | 116 | 157 | 157 | 1 | 119 | 157 | 117 | 1 | 1 | 1 | 1 | 1 | 1 | 157 | 157 | 1 | 1 | 1 | 1 | 1 | 157 | 158 | 158 | 1 |
| YPTB3033 | 1 | 1 | 1 | 1 | 1 | 1 | 1 | 1 | 1 | 1 | 1 | 1 | 1 | 1 | 1 | 1 | 1 | 98 | 1 | 1 | 1 | 1 | 1 | 1 | 1 | 1 | 1 | 1 | 1 | 1 | 1 | 1 | 1 | 1 | 126 |
| YPTB3034 | 1 | 1 | 1 | 1 | 1 | 1 | 1 | 1 | 1 | 1 | 1 | 1 | 1 | 1 | 1 | 1 | 1 | 1 | 1 | 1 | 1 | 1 | 1 | 1 | 1 | 1 | 1 | 1 | 1 | 1 | 1 | 1 | 1 | 1 | 1 |
| YPTB3035 | 1 | 139 | 1 | 141 | 1 | 1 | 1 | 142 | 139 | 141 | 141 | 1 | 1 | 1 | 1 | 141 | 1 | 136 | 1 | 1 | 1 | 1 | 1 | 1 | 1 | 1 | 1 | 1 | 1 | 1 | 1 | 1 | 141 | 141 | 1 |
| YPTB3036 | 1 | 47 | 1 | 47 | 1 | 1 | 1 | 47 | 47 | 47 | 47 | 1 | 1 | 1 | 1 | 47 | 1 | 47 | 1 | 1 | 1 | 1 | 1 | 1 | 1 | 1 | 1 | 1 | 1 | 1 | 1 | 1 | 47 | 47 | 1 |
| YPTB3042 | 140 | 141 | 140 | 140 | 1 | 1 | 140 | 142 | 141 | 140 | 140 | 1 | 1 | 1 | 1 | 140 | 1 | 138 | 140 | 140 | 1 | 1 | 1 | 140 | 1 | 1 | 1 | 1 | 1 | 1 | 1 | 1 | 140 | 140 | 140 |
| YPTB3165 | 165 | 127 | 165 | 130 | 1 | 1 | 165 | 130 | 127 | 129 | 130 | 122 | 133 | 133 | 1 | 129 | 133 | 124 | 165 | 165 | 1 | 1 | 1 | 165 | 133 | 133 | 1 | 1 | 1 | 1 | 1 | 133 | 130 | 130 | 165 |
| YPTB3167 | 97 | 1 | 97 | 1 | 1 | 1 | 97 | 1 | 1 | 1 | 1 | 97 | 1 | 1 | 1 | 1 | 1 | 1 | 97 | 97 | 1 | 1 | 1 | 97 | 1 | 1 | 1 | 1 | 1 | 1 | 1 | 1 | 1 | 1 | 97 |
| YPTB3181 | 72 | 73 | 72 | 1 | 1 | 1 | 72 | 1 | 73 | 1 | 1 | 1 | 1 | 1 | 1 | 1 | 1 | 1 | 72 | 72 | 1 | 1 | 1 | 72 | 1 | 1 | 1 | 1 | 1 | 1 | 1 | 1 | 1 | 1 | 72 |
| YPTB3182 | 124 | 119 | 116 | 121 | 1 | 1 | 116 | 122 | 119 | 121 | 121 | 115 | 115 | 115 | 1 | 121 | 115 | 1 | 116 | 116 | 1 | 1 | 1 | 116 | 115 | 115 | 1 | 1 | 1 | 1 | 1 | 115 | 121 | 121 | 124 |
| YPTB3196 | 102 | 105 | 102 | 102 | 1 | 1 | 102 | 101 | 105 | 102 | 102 | 101 | 102 | 102 | 1 | 102 | 102 | 1 | 102 | 102 | 1 | 1 | 1 | 102 | 102 | 102 | 1 | 1 | 1 | 1 | 1 | 102 | 102 | 102 | 102 |
| YPTB3197 | 100 | 100 | 100 | 100 | 1 | 1 | 100 | 100 | 100 | 100 | 100 | 100 | 100 | 100 | 1 | 100 | 100 | 1 | 100 | 100 | 1 | 1 | 1 | 100 | 100 | 100 | 1 | 1 | 1 | 1 | 1 | 100 | 100 | 100 | 100 |
| YPTB3198 | 132 | 135 | 132 | 146 | 1 | 1 | 132 | 138 | 135 | 183 | 146 | 131 | 141 | 141 | 1 | 183 | 141 | 1 | 132 | 132 | 1 | 1 | 1 | 132 | 141 | 141 | 1 | 1 | 1 | 1 | 1 | 141 | 146 | 146 | 132 |
| YPTB3205 | 95 | 1 | 95 | 95 | 1 | 1 | 95 | 96 | 1 | 97 | 95 | 1 | 1 | 1 | 1 | 97 | 1 | 95 | 95 | 95 | 1 | 1 | 1 | 95 | 1 | 1 | 1 | 1 | 1 | 1 | 1 | 1 | 95 | 95 | 95 |
| YPTB3210 | 1 | 105 | 1 | 1 | 1 | 1 | 1 | 106 | 105 | 1 | 1 | 1 | 1 | 1 | 1 | 1 | 1 | 1 | 1 | 1 | 1 | 1 | 1 | 1 | 1 | 1 | 1 | 1 | 1 | 1 | 1 | 1 | 1 | 1 | 1 |
| YPTB3215 | 1 | 1 | 95 | 77 | 1 | 1 | 1 | 1 | 1 | 1 | 77 | 70 | 1 | 1 | 1 | 1 | 1 | 1 | 1 | 1 | 1 | 1 | 1 | 1 | 1 | 1 | 1 | 1 | 1 | 1 | 1 | 1 | 77 | 77 | 1 |
| YPTB3217 | 92 | 94 | 92 | 99 | 1 | 1 | 92 | 1 | 94 | 95 | 99 | 91 | 1 | 1 | 1 | 119 | 1 | 1 | 92 | 92 | 1 | 1 | 1 | 92 | 1 | 1 | 1 | 1 | 1 | 1 | 1 | 1 | 99 | 99 | 92 |
| YPTB3218 | 108 | 111 | 108 | 111 | 1 | 1 | 108 | 110 | 111 | 111 | 111 | 107 | 1 | 1 | 1 | 111 | 1 | 1 | 108 | 108 | 1 | 1 | 1 | 108 | 1 | 1 | 1 | 1 | 1 | 1 | 1 | 1 | 111 | 111 | 108 |
| YPTB3222 | 1 | 51 | 1 | 51 | 1 | 1 | 1 | 51 | 51 | 51 | 51 | 1 | 1 | 1 | 1 | 51 | 1 | 51 | 1 | 1 | 1 | 1 | 1 | 1 | 1 | 1 | 1 | 1 | 1 | 1 | 1 | 1 | 51 | 51 | 1 |
| YPTB3225 | 1 | 48 | 1 | 47 | 1 | 1 | 1 | 48 | 48 | 47 | 47 | 1 | 1 | 1 | 1 | 47 | 1 | 47 | 1 | 1 | 1 | 1 | 1 | 1 | 1 | 1 | 1 | 1 | 1 | 1 | 1 | 1 | 47 | 47 | 1 |
| YPTB3230 | 1 | 1 | 1 | 106 | 1 | 1 | 1 | 1 | 1 | 106 | 106 | 1 | 1 | 1 | 1 | 106 | 1 | 1 | 1 | 1 | 1 | 1 | 1 | 1 | 1 | 1 | 1 | 1 | 1 | 1 | 1 | 1 | 106 | 106 | 1 |
| YPTB3313 | 74 | 76 | 74 | 74 | 1 | 1 | 74 | 74 | 76 | 74 | 74 | 1 | 1 | 1 | 1 | 74 | 1 | 75 | 74 | 74 | 1 | 1 | 1 | 74 | 1 | 1 | 1 | 1 | 1 | 1 | 1 | 1 | 74 | 74 | 74 |
| YPTB3319 | 1 | 83 | 1 | 83 | 1 | 1 | 1 | 1 | 83 | 83 | 83 | 1 | 1 | 1 | 1 | 83 | 1 | 83 | 1 | 1 | 1 | 1 | 1 | 1 | 1 | 1 | 1 | 1 | 1 | 1 | 1 | 1 | 83 | 83 | 1 |
| YPTB3320 | 84 | 84 | 84 | 1 | 1 | 1 | 84 | 86 | 84 | 1 | 1 | 83 | 1 | 1 | 1 | 1 | 1 | 84 | 84 | 84 | 1 | 1 | 1 | 84 | 1 | 1 | 1 | 1 | 1 | 1 | 1 | 1 | 1 | 1 | 84 |
| YPTB3339 | 1 | 1 | 1 | 1 | 1 | 1 | 1 | 1 | 1 | 73 | 1 | 1 | 1 | 1 | 1 | 73 | 1 | 1 | 1 | 1 | 1 | 1 | 1 | 1 | 1 | 1 | 1 | 1 | 1 | 1 | 1 | 1 | 1 | 1 | 1 |
| YPTB3340 | 1 | 59 | 1 | 56 | 1 | 1 | 1 | 1 | 59 | 56 | 56 | 1 | 1 | 1 | 1 | 56 | 1 | 56 | 1 | 1 | 1 | 1 | 1 | 1 | 1 | 1 | 1 | 1 | 1 | 1 | 1 | 1 | 56 | 56 | 1 |
| YPTB3347 | 1 | 105 | 1 | 106 | 1 | 1 | 1 | 105 | 105 | 106 | 106 | 1 | 1 | 1 | 1 | 106 | 1 | 106 | 1 | 1 | 1 | 1 | 1 | 1 | 1 | 1 | 1 | 1 | 1 | 1 | 1 | 1 | 106 | 106 | 1 |
| YPTB3352 | 1 | 72 | 1 | 72 | 1 | 1 | 1 | 72 | 72 | 72 | 72 | 1 | 1 | 1 | 1 | 72 | 1 | 72 | 1 | 1 | 1 | 1 | 1 | 1 | 1 | 1 | 1 | 1 | 1 | 1 | 1 | 1 | 72 | 72 | 1 |
| YPTB3354 | 1 | 1 | 1 | 1 | 1 | 1 | 1 | 63 | 1 | 1 | 1 | 1 | 1 | 1 | 1 | 1 | 1 | 1 | 1 | 1 | 1 | 1 | 1 | 1 | 1 | 1 | 1 | 1 | 1 | 1 | 1 | 1 | 1 | 1 | 1 |
| YPTB3357 | 139 | 142 | 139 | 144 | 1 | 1 | 139 | 201 | 142 | 144 | 144 | 138 | 1 | 1 | 1 | 144 | 1 | 140 | 139 | 139 | 1 | 1 | 1 | 139 | 1 | 1 | 1 | 1 | 1 | 1 | 1 | 1 | 144 | 144 | 139 |
| YPTB3395 | 170 | 130 | 170 | 132 | 1 | 1 | 170 | 134 | 130 | 132 | 132 | 1 | 1 | 1 | 1 | 132 | 1 | 127 | 170 | 170 | 1 | 1 | 1 | 170 | 1 | 1 | 1 | 1 | 1 | 1 | 1 | 1 | 132 | 132 | 170 |
| YPTB3396 | 69 | 69 | 69 | 69 | 1 | 1 | 69 | 68 | 69 | 69 | 69 | 1 | 1 | 1 | 1 | 69 | 1 | 69 | 69 | 69 | 1 | 1 | 1 | 69 | 1 | 1 | 1 | 1 | 1 | 1 | 1 | 1 | 69 | 69 | 69 |
| YPTB3398 | 1 | 1 | 1 | 1 | 1 | 1 | 1 | 1 | 1 | 1 | 1 | 1 | 1 | 1 | 1 | 1 | 1 | 1 | 1 | 1 | 1 | 1 | 1 | 1 | 1 | 1 | 1 | 1 | 1 | 1 | 1 | 1 | 1 | 1 | 1 |
| YPTB3407 | 1 | 1 | 1 | 1 | 1 | 1 | 1 | 123 | 1 | 1 | 1 | 1 | 1 | 1 | 1 | 1 | 1 | 121 | 1 | 1 | 1 | 1 | 1 | 1 | 1 | 1 | 1 | 1 | 1 | 1 | 1 | 1 | 1 | 1 | 1 |
| YPTB3413 | 1 | 69 | 1 | 71 | 1 | 1 | 1 | 1 | 69 | 71 | 71 | 1 | 1 | 1 | 1 | 71 | 1 | 1 | 1 | 1 | 1 | 1 | 1 | 1 | 1 | 1 | 1 | 1 | 1 | 1 | 1 | 1 | 71 | 71 | 1 |
| YPTB3416 | 1 | 1 | 1 | 1 | 1 | 1 | 1 | 1 | 1 | 1 | 1 | 1 | 1 | 1 | 1 | 1 | 1 | 1 | 1 | 1 | 1 | 1 | 1 | 1 | 1 | 1 | 1 | 1 | 1 | 1 | 1 | 1 | 1 | 1 | 1 |
| YPTB3476 | 126 | 129 | 126 | 131 | 1 | 126 | 126 | 125 | 133 | 131 | 131 | 125 | 125 | 125 | 126 | 131 | 125 | 126 | 126 | 126 | 126 | 126 | 126 | 126 | 125 | 125 | 126 | 126 | 126 | 126 | 126 | 125 | 131 | 131 | 126 |
| YPTB3478 | 112 | 114 | 112 | 116 | 1 | 1 | 112 | 115 | 160 | 116 | 116 | 1 | 1 | 1 | 1 | 116 | 1 | 112 | 112 | 112 | 1 | 1 | 1 | 112 | 1 | 1 | 1 | 1 | 1 | 1 | 1 | 1 | 116 | 116 | 112 |
| YPTB3479 | 102 | 104 | 102 | 105 | 1 | 1 | 102 | 100 | 104 | 105 | 105 | 101 | 1 | 1 | 1 | 105 | 1 | 103 | 102 | 102 | 1 | 1 | 1 | 102 | 1 | 1 | 1 | 1 | 1 | 1 | 1 | 1 | 105 | 105 | 102 |

| | | | | | | | | | | | | | | | | | | | | | | | | | | | | | | | | | | | |
|---|---|---|---|---|---|---|---|---|---|---|---|---|---|---|---|---|---|---|---|---|---|---|---|---|---|---|---|---|---|---|---|---|---|---|---|
| YPTB3485 | 1 | 1 | 1 | 1 | 1 | 1 | 1 | 1 | 1 | 1 | 1 | 1 | 1 | 1 | 1 | 1 | 1 | 1 | 1 | 1 | 1 | 1 | 1 | 1 | 1 | 1 | 1 | 1 | 1 | 1 | 1 | 1 | 1 | 1 | 1 |
| YPTB3488 | 1 | 1 | 1 | 1 | 1 | 1 | 1 | 1 | 1 | 1 | 1 | 1 | 1 | 1 | 1 | 1 | 92 | 1 | 1 | 1 | 1 | 1 | 1 | 1 | | 217 | 1 | 1 | 1 | 1 | 1 | 1 | 1 | 1 | 1 |
| YPTB3491 | 1 | 101 | 1 | 102 | 1 | 1 | 1 | 1 | 124 | 102 | 102 | 99 | 1 | 1 | 1 | 102 | 1 | 100 | 1 | 1 | 1 | 1 | 1 | 1 | 1 | 1 | 1 | 1 | 1 | 1 | 1 | 1 | 102 | 102 | 1 |
| YPTB3498 | 1 | 84 | 1 | 85 | 1 | 1 | 1 | 83 | 84 | 85 | 85 | 82 | 1 | 1 | 1 | 85 | 1 | 102 | 1 | 1 | 1 | 1 | 1 | 1 | 1 | 1 | 1 | 1 | 1 | 1 | 1 | 1 | 85 | 85 | 1 |
| YPTB3503 | 129 | 132 | 129 | 134 | 1 | 129 | 129 | 127 | 132 | 134 | 134 | 1 | 1 | 1 | 129 | 134 | 1 | 130 | 129 | 129 | 129 | 129 | 129 | 129 | 1 | 1 | 129 | 129 | 129 | 129 | 129 | 1 | 134 | 134 | 129 |
| YPTB3506 | 1 | 1 | 1 | 1 | 1 | 1 | 1 | 1 | 1 | 67 | 1 | 1 | 1 | 1 | 1 | 67 | 1 | 1 | 1 | 1 | 1 | 1 | 1 | 1 | 1 | 1 | 1 | 1 | 1 | 1 | 1 | 1 | 1 | 1 | 1 |
| YPTB3508 | 1 | 1 | 1 | 1 | 1 | 1 | 1 | 1 | 1 | 1 | 1 | 1 | 1 | 1 | 1 | 1 | 1 | 1 | 1 | 1 | 1 | 1 | 1 | 1 | 1 | 1 | 1 | 1 | 1 | 1 | 1 | 1 | 1 | 1 | 1 |
| YPTB3510 | 1 | 1 | 1 | 1 | 1 | 1 | 1 | 53 | 1 | 1 | 1 | 1 | 1 | 1 | 1 | 1 | 1 | 54 | 1 | 1 | 1 | 1 | 1 | 1 | 1 | 1 | 1 | 1 | 1 | 1 | 1 | 1 | 1 | 1 | 1 |
| YPTB3518 | 1 | 1 | 1 | 1 | 1 | 1 | 1 | 1 | 1 | 1 | 1 | 1 | 1 | 1 | 1 | 1 | 1 | 1 | 1 | 1 | 1 | 1 | 1 | 1 | 1 | 1 | 1 | 1 | 1 | 1 | 1 | 1 | 1 | 1 | 1 |
| YPTB3523 | 1 | 72 | 1 | 73 | 1 | 1 | 1 | 72 | 72 | 73 | 73 | 1 | 1 | 1 | 1 | 73 | 1 | 71 | 1 | 1 | 1 | 1 | 1 | 1 | 1 | 1 | 1 | 1 | 1 | 1 | 1 | 1 | 73 | 73 | 1 |
| YPTB3524 | 1 | 1 | 1 | 77 | 1 | 1 | 1 | 77 | 1 | 77 | 77 | 1 | 1 | 1 | 1 | 77 | 1 | 1 | 1 | 1 | 1 | 1 | 1 | 1 | 1 | 1 | 1 | 1 | 1 | 1 | 1 | 1 | 77 | 77 | 1 |
| YPTB3525 | 86 | 86 | 86 | 86 | 1 | 1 | 86 | 86 | 86 | 86 | 86 | 1 | 1 | 1 | 1 | 86 | 1 | 86 | 86 | 86 | 1 | 1 | 1 | 86 | 1 | 1 | 1 | 1 | 1 | 1 | 1 | 1 | 86 | 86 | 86 |
| YPTB3526 | 319 | 114 | 112 | 114 | 1 | 1 | 112 | 116 | 114 | 114 | 114 | 1 | 1 | 1 | 1 | 114 | 1 | 113 | 112 | 112 | 1 | 1 | 1 | 112 | 1 | 1 | 1 | 1 | 1 | 1 | 1 | 1 | 114 | 114 | 112 |
| YPTB3527 | 1 | 1 | 1 | 1 | 1 | 1 | 1 | 1 | 1 | 1 | 1 | 1 | 1 | 1 | 1 | 1 | 1 | 42 | 1 | 1 | 1 | 1 | 1 | 1 | 1 | 1 | 1 | 1 | 1 | 1 | 1 | 1 | 1 | 1 | 1 |
| YPTB3528 | 61 | 62 | 61 | 61 | 1 | 1 | 61 | 1 | 62 | 61 | 61 | 1 | 1 | 1 | 1 | 61 | 1 | 61 | 61 | 61 | 1 | 1 | 1 | 61 | 1 | 1 | 1 | 1 | 1 | 1 | 1 | 1 | 61 | 61 | 61 |
| YPTB3531 | 96 | 94 | 96 | 95 | 1 | 96 | 96 | 101 | 94 | 95 | 95 | 95 | 1 | 1 | 96 | 95 | 1 | 97 | 96 | 96 | 96 | 96 | 96 | 96 | 1 | 1 | 96 | 96 | 96 | 96 | 96 | 1 | 95 | 95 | 96 |
| YPTB3533 | 1 | 1 | 1 | 64 | 1 | 1 | 1 | 64 | 1 | 64 | 64 | 1 | 1 | 1 | 1 | 64 | 1 | 64 | 1 | 1 | 1 | 1 | 1 | 1 | 1 | 1 | 1 | 1 | 1 | 1 | 1 | 1 | 64 | 64 | 1 |
| YPTB3536 | 1 | 120 | 1 | 123 | 1 | 1 | 1 | 125 | 120 | 123 | 123 | 1 | 1 | 1 | 1 | 123 | 1 | 120 | 1 | 1 | 1 | 1 | 1 | 1 | 1 | 1 | 1 | 1 | 1 | 1 | 1 | 1 | 123 | 123 | 1 |
| YPTB3538 | 1 | 72 | 1 | 71 | 1 | 75 | 1 | 1 | 72 | 71 | 71 | 1 | 1 | 1 | 75 | 71 | 1 | 71 | 1 | 1 | 75 | 75 | 75 | 1 | 1 | 1 | 75 | 75 | 75 | 75 | 75 | 1 | 71 | 71 | 1 |
| YPTB3547 | 1 | 114 | 1 | 116 | 1 | 1 | 1 | 117 | 114 | 116 | 116 | 1 | 1 | 1 | 1 | 116 | 1 | 1 | 1 | 1 | 1 | 1 | 1 | 1 | 1 | 1 | 1 | 1 | 1 | 1 | 1 | 1 | 116 | 116 | 1 |
| YPTB3548 | 1 | 1 | 1 | 1 | 1 | 1 | 1 | 1 | 1 | 1 | 1 | 1 | 1 | 1 | 1 | 1 | 1 | 1 | 1 | 1 | 1 | 1 | 1 | 1 | 1 | 1 | 1 | 1 | 1 | 1 | 1 | 1 | 1 | 1 | 1 |
| YPTB3549 | 112 | 115 | 112 | 125 | 1 | 1 | 112 | 119 | 115 | 117 | 125 | 111 | 1 | 1 | 1 | 117 | 1 | 1 | 112 | 112 | 1 | 1 | 1 | 112 | 1 | 1 | 1 | 1 | 1 | 1 | 1 | 1 | 125 | 125 | |
| YPTB3558 | 1 | 131 | 1 | 133 | 1 | 1 | 1 | 135 | 131 | 133 | 133 | 127 | 1 | 1 | 1 | 133 | 1 | 128 | 1 | 1 | 1 | 1 | 1 | 1 | 1 | 1 | 1 | 1 | 1 | 1 | 1 | 1 | 133 | 133 | 1 |
| YPTB3561 | 119 | 122 | 119 | 119 | 1 | 1 | 119 | 123 | 122 | 119 | 119 | 118 | 1 | 1 | 1 | 119 | 1 | 120 | 119 | 119 | 1 | 1 | 1 | 119 | 1 | 1 | 1 | 1 | 1 | 1 | 1 | 1 | 119 | 119 | 119 |
| YPTB3563 | 90 | 1 | 90 | 1 | 1 | 1 | 90 | 1 | 1 | 1 | 1 | 1 | 1 | 1 | 1 | 1 | 1 | 75 | 90 | 90 | 1 | 1 | 1 | 90 | 1 | 1 | 1 | 1 | 1 | 1 | 1 | 1 | 1 | 1 | 90 |
| YPTB3565 | 1 | 95 | 1 | 1 | 1 | 1 | 1 | 92 | 95 | 1 | 1 | 1 | 1 | 1 | 1 | 1 | 1 | 89 | 1 | 1 | 1 | 1 | 1 | 1 | 1 | 1 | 1 | 1 | 1 | 1 | 1 | 1 | 1 | 1 | 1 |
| YPTB3570 | 60 | 61 | 60 | 61 | 1 | 1 | 60 | 62 | 61 | 61 | 61 | 60 | 1 | 1 | 1 | 61 | 1 | 61 | 60 | 60 | 1 | 1 | 1 | 60 | 1 | 1 | 1 | 1 | 1 | 1 | 1 | 1 | 61 | 61 | 60 |
| YPTB3574 | 129 | 133 | 129 | 1 | 1 | 1 | 129 | 136 | 133 | 1 | 1 | 1 | 1 | 1 | 1 | 1 | 1 | 130 | 129 | 129 | 1 | 1 | 1 | 129 | 1 | 1 | 1 | 1 | 1 | 1 | 1 | 1 | 1 | 1 | 129 |
| YPTB3649 | 168 | 123 | 121 | 124 | 1 | 1 | 121 | 126 | 123 | 124 | 124 | 121 | 121 | 121 | 1 | 124 | 121 | 121 | 121 | 121 | 1 | 1 | 1 | 121 | 121 | 121 | 1 | 1 | 1 | 1 | 475 | 121 | 124 | 124 | 121 |
| YPTB3660 | 96 | 1 | 96 | 1 | 1 | 1 | 96 | 1 | 1 | 1 | 1 | 1 | 1 | 1 | 1 | 1 | 1 | 1 | 96 | 96 | 1 | 1 | 1 | 96 | 1 | 1 | 1 | 1 | 1 | 1 | 1 | 1 | 1 | 1 | 96 |
| YPTB3663 | 1 | 63 | 1 | 1 | 1 | 1 | 1 | 1 | 76 | 1 | 1 | 1 | 1 | 1 | 1 | 1 | 1 | 1 | 1 | 1 | 1 | 1 | 1 | 1 | 1 | 1 | 1 | 1 | 1 | 1 | 1 | 1 | 1 | 1 | 1 |
| YPTB3665 | 1 | 1 | 1 | 1 | 1 | 1 | 1 | 1 | 1 | 1 | 1 | 1 | 1 | 1 | 1 | 1 | 1 | 1 | 1 | 1 | 1 | 1 | 1 | 1 | 1 | 1 | 1 | 1 | 1 | 1 | 1 | 1 | 1 | 1 | 1 |
| YPTB3668 | 1 | 1 | 1 | 1 | 1 | 1 | 1 | 1 | 1 | 1 | 1 | 1 | 1 | 1 | 1 | 1 | 1 | 1 | 1 | 1 | 319 | 1 | 1 | 1 | 240 | 1 | 1 | 1 | 1 | 1 | 1 | 1 | 1 | 1 | 366 |
| YPTB3674 | 1 | 1 | 1 | 1 | 1 | 1 | 1 | 1 | 1 | 1 | 1 | 1 | 1 | 1 | 1 | 1 | 1 | 1 | 1 | 1 | 1 | 1 | 1 | 1 | 1 | 1 | 1 | 1 | 1 | 1 | 1 | 1 | 1 | 1 | 1 |
| YPTB3675 | 1 | 1 | 1 | 1 | 1 | 1 | 1 | 1 | 1 | 1 | 1 | 1 | 1 | 1 | 1 | 1 | 1 | 1 | 1 | 1 | 1 | 1 | 1 | 1 | 1 | 1 | 1 | 1 | 1 | 1 | 1 | 1 | 1 | 1 | 1 |
| YPTB3676 | 1 | 1 | 1 | 1 | 1 | 1 | 1 | 1 | 1 | 1 | 1 | 1 | 1 | 1 | 1 | 1 | 1 | 1 | 1 | 1 | 1 | 1 | 1 | 1 | 1 | 1 | 1 | 1 | 1 | 1 | 1 | 1 | 1 | 1 | 1 |
| YPTB3677 | 1 | 1 | 1 | 1 | 1 | 1 | 1 | 1 | 1 | 1 | 1 | 1 | 1 | 1 | 1 | 1 | 1 | 1 | 1 | 1 | 1 | 1 | 1 | 1 | 1 | 1 | 1 | 1 | 1 | 1 | 1 | 1 | 1 | 1 | 1 |
| YPTB3678 | 1 | 60 | 1 | 1 | 1 | 1 | 1 | 1 | 63 | 1 | 1 | 1 | 1 | 1 | 1 | 1 | 1 | 1 | 1 | 1 | 1 | 1 | 1 | 1 | 1 | 1 | 1 | 1 | 1 | 1 | 1 | 1 | 1 | 1 | 1 |
| YPTB3679 | 1 | 28 | 1 | 1 | 1 | 1 | 1 | 1 | 1 | 1 | 1 | 1 | 1 | 1 | 1 | 1 | 1 | 1 | 1 | 1 | 1 | 1 | 1 | 1 | 1 | 1 | 1 | 1 | 1 | 1 | 1 | 1 | 1 | 1 | 1 |
| YPTB3680 | 1 | 1 | 1 | 1 | 1 | 1 | 1 | 1 | 1 | 1 | 1 | 1 | 1 | 1 | 1 | 1 | 1 | 1 | 1 | 1 | 1 | 1 | 1 | 1 | 1 | 1 | 1 | 1 | 1 | 1 | 1 | 1 | 1 | 1 | 1 |
| YPTB3681 | 32 | 33 | 32 | 1 | 1 | 1 | 32 | 32 | 33 | 1 | 1 | 1 | 1 | 1 | 1 | 1 | 1 | 32 | 32 | 32 | 1 | 1 | 1 | 32 | 1 | 1 | 1 | 1 | 1 | 1 | 1 | 1 | 1 | 1 | 32 |
| YPTB3682 | 1 | 1 | 1 | 1 | 1 | 1 | 1 | 1 | 1 | 1 | 1 | 1 | 1 | 1 | 1 | 1 | 1 | 1 | 1 | 1 | 1 | 1 | 1 | 1 | 1 | 1 | 1 | 1 | 1 | 1 | 1 | 1 | 1 | 1 | 1 |
| YPTB3683 | 121 | 1 | 1 | 1 | 1 | 1 | 1 | 1 | 1 | 1 | 1 | 1 | 1 | 1 | 1 | 1 | 1 | 1 | 1 | 1 | 1 | 1 | 1 | 1 | 1 | 1 | 1 | 1 | 1 | 1 | 1 | 1 | 1 | 1 | 1 |
| YPTB3684 | 1 | 1 | 1 | 1 | 1 | 1 | 1 | 1 | 1 | 1 | 1 | 1 | 1 | 1 | 1 | 1 | 1 | 1 | 1 | 1 | 1 | 1 | 1 | 1 | 1 | 1 | 1 | 1 | 1 | 1 | 1 | 1 | 1 | 1 | 1 |
| YPTB3685 | 1 | 1 | 1 | 1 | 1 | 1 | 1 | 1 | 1 | 1 | 1 | 1 | 1 | 1 | 1 | 1 | 1 | 1 | 1 | 1 | 1 | 1 | 1 | 1 | 1 | 1 | 1 | 1 | 1 | 1 | 1 | 1 | 1 | 1 | 1 |
| YPTB3686 | 1 | 1 | 1 | 1 | 1 | 1 | 1 | 1 | 1 | 1 | 1 | 1 | 1 | 1 | 1 | 1 | 1 | 1 | 1 | 1 | 1 | 1 | 1 | 1 | 1 | 1 | 1 | 1 | 1 | 1 | 1 | 1 | 1 | 1 | 1 |
| YPTB3687 | 1 | 1 | 1 | 1 | 1 | 1 | 1 | 1 | 1 | 1 | 1 | 1 | 1 | 1 | 1 | 1 | 1 | 1 | 1 | 1 | 1 | 1 | 1 | 1 | 1 | 1 | 1 | 1 | 1 | 1 | 1 | 1 | 1 | 1 | 1 |
| YPTB3688 | 1 | 1 | 1 | 1 | 1 | 1 | 1 | 1 | 1 | 1 | 1 | 1 | 1 | 1 | 1 | 1 | 1 | 1 | 1 | 1 | 1 | 1 | 1 | 1 | 1 | 1 | 1 | 1 | 1 | 1 | 1 | 1 | 1 | 1 | 1 |
| YPTB3690 | 1 | 1 | 1 | 1 | 1 | 1 | 1 | 1 | 1 | 1 | 1 | 1 | 1 | 1 | 1 | 1 | 1 | 1 | 1 | 1 | 1 | 1 | 1 | 1 | 1 | 1 | 1 | 1 | 1 | 1 | 1 | 1 | 1 | 1 | 1 |
| YPTB3691 | 1 | 1 | 1 | 1 | 1 | 1 | 1 | 1 | 1 | 1 | 1 | 1 | 1 | 1 | 1 | 1 | 1 | 29 | 1 | 1 | 1 | 1 | 1 | 1 | 1 | 1 | 1 | 1 | 1 | 1 | 1 | 1 | 1 | 1 | 1 |
| YPTB3692 | 33 | 1 | 33 | 1 | 1 | 1 | 33 | 1 | 1 | 1 | 1 | 1 | 1 | 1 | 1 | 1 | 1 | 1 | 33 | 33 | 1 | 1 | 1 | 33 | 1 | 1 | 1 | 1 | 1 | 1 | 1 | 1 | 1 | 1 | 33 |
| YPTB3693 | 1 | 1 | 1 | 1 | 1 | 1 | 1 | 1 | 1 | 1 | 1 | 1 | 1 | 1 | 1 | 1 | 1 | 1 | 1 | 1 | 1 | 1 | 1 | 1 | 1 | 1 | 1 | 1 | 1 | 1 | 1 | 1 | 1 | 1 | 1 |
| YPTB3694 | 1 | 1 | 1 | 1 | 1 | 1 | 1 | 1 | 1 | 1 | 1 | 1 | 1 | 1 | 1 | 1 | 1 | 1 | 1 | 1 | 1 | 1 | 1 | 1 | 1 | 1 | 1 | 1 | 1 | 1 | 1 | 1 | 1 | 1 | 1 |
| YPTB3695 | 42 | 1 | 42 | 1 | 1 | 1 | 42 | 1 | 1 | 1 | 1 | 1 | 1 | 1 | 1 | 1 | 1 | 1 | 42 | 42 | 1 | 1 | 1 | 42 | 1 | 1 | 1 | 1 | 1 | 1 | 1 | 1 | 1 | 1 | 42 |
| YPTB3696 | 1 | 1 | 1 | 1 | 1 | 1 | 1 | 1 | 1 | 1 | 1 | 1 | 1 | 1 | 1 | 1 | 1 | 1 | 1 | 1 | 1 | 1 | 1 | 1 | 1 | 1 | 1 | 1 | 1 | 1 | 1 | 1 | 1 | 1 | 1 |
| YPTB3697 | 1 | 1 | 1 | 47 | 1 | 1 | 1 | 1 | 1 | 47 | 47 | 1 | 1 | 1 | 1 | 47 | 1 | 1 | 1 | 1 | 1 | 1 | 1 | 1 | 1 | 1 | 1 | 1 | 1 | 1 | 1 | 1 | 47 | 47 | 1 |
| YPTB3698 | 1 | 47 | 1 | 46 | 1 | 1 | 1 | 1 | 47 | 46 | 46 | 1 | 1 | 1 | 1 | 46 | 1 | 1 | 1 | 1 | 1 | 1 | 1 | 1 | 1 | 1 | 1 | 1 | 1 | 1 | 1 | 1 | 46 | 46 | 1 |
| YPTB3699 | 1 | 1 | 1 | 1 | 1 | 1 | 1 | 1 | 1 | 1 | 1 | 1 | 1 | 1 | 1 | 1 | 1 | 1 | 1 | 1 | 1 | 1 | 1 | 1 | 1 | 1 | 1 | 1 | 1 | 1 | 1 | 1 | 1 | 1 | 1 |
| YPTB3704 | 1 | 1 | 1 | 37 | 1 | 1 | 1 | 37 | 1 | 37 | 37 | 1 | 1 | 1 | 1 | 37 | 1 | 37 | 1 | 1 | 1 | 1 | 1 | 1 | 1 | 1 | 1 | 1 | 1 | 1 | 1 | 1 | 37 | 37 | 1 |
| YPTB3705 | 1 | 1 | 1 | 1 | 1 | 1 | 1 | 1 | 1 | 1 | 1 | 1 | 1 | 1 | 1 | 1 | 1 | 1 | 1 | 1 | 1 | 1 | 1 | 1 | 1 | 1 | 1 | 1 | 1 | 1 | 1 | 1 | 1 | 1 | 1 |
| YPTB3706 | 63 | 1 | 63 | 1 | 1 | 1 | 63 | 1 | 1 | 1 | 1 | 1 | 1 | 1 | 1 | 1 | 1 | 1 | 63 | 63 | 1 | 1 | 1 | 63 | 1 | 1 | 1 | 1 | 1 | 1 | 1 | 1 | 1 | 1 | 63 |
| YPTB3709 | 82 | 1 | 82 | 102 | 1 | 1 | 82 | 1 | 1 | 84 | 102 | 1 | 1 | 1 | 1 | 84 | 1 | 1 | 82 | 82 | 1 | 1 | 1 | 82 | 1 | 1 | 1 | 1 | 1 | 1 | 1 | 1 | 102 | 102 | 82 |
| YPTB3711 | 1 | 1 | 1 | 1 | 1 | 1 | 1 | 1 | 1 | 1 | 1 | 1 | 1 | 1 | 1 | 1 | 1 | 1 | 1 | 1 | 1 | 1 | 1 | 1 | 1 | 1 | 1 | 1 | 1 | 1 | 1 | 1 | 1 | 1 | 1 |
| YPTB3714 | 176 | 124 | 176 | 126 | 1 | 1 | 176 | 119 | 124 | 126 | 126 | 119 | 122 | 122 | 1 | 126 | 122 | 176 | 176 | 176 | 1 | 1 | 1 | 176 | 122 | 122 | 1 | 1 | 1 | 1 | 1 | 122 | 126 | 126 | 176 |
| YPTB3725 | 64 | 65 | 64 | 1 | 1 | 64 | 64 | 1 | 65 | 1 | 1 | 63 | 64 | 64 | 64 | 1 | 64 | 1 | 64 | 64 | 64 | 64 | 64 | 64 | 64 | 64 | 64 | 64 | 64 | 64 | 64 | 64 | 1 | 1 | 64 |
| YPTB3729 | 1 | 53 | 1 | 1 | 1 | 53 | 1 | 53 | 53 | 1 | 1 | 53 | 53 | 53 | 53 | 1 | 53 | 1 | 1 | 1 | 53 | 53 | 53 | 1 | 53 | 53 | 53 | 53 | 53 | 53 | 53 | 53 | 1 | 1 | 1 |
| YPTB3732 | 93 | 95 | 93 | 94 | 1 | 1 | 93 | 1 | 95 | 95 | 94 | 92 | 1 | 1 | 1 | 95 | 1 | 94 | 93 | 93 | 1 | 1 | 1 | 93 | 1 | 1 | 1 | 1 | 1 | 1 | 1 | 1 | 94 | 94 | 93 |
| YPTB3734 | 80 | 81 | 80 | 1 | 1 | 1 | 80 | 1 | 81 | 1 | 1 | 1 | 1 | 1 | 1 | 1 | 1 | 1 | 80 | 80 | 1 | 1 | 1 | 80 | 1 | 1 | 1 | 1 | 1 | 1 | 1 | 1 | 1 | 1 | 80 |
| YPTB3743 | 1 | 124 | 1 | 125 | 1 | 1 | 1 | 126 | 163 | 125 | 125 | 122 | 1 | 1 | 1 | 125 | 1 | 1 | 1 | 1 | 1 | 1 | 1 | 1 | 1 | 1 | 1 | 1 | 1 | 1 | 1 | 1 | 125 | 125 | 1 |
| YPTB3746 | 110 | 1 | 110 | 109 | 1 | 110 | 110 | 109 | 1 | 109 | 109 | 110 | 110 | 110 | 110 | 109 | 110 | 1 | 110 | 110 | 110 | 110 | 110 | 110 | 110 | 110 | 110 | 110 | 110 | 110 | 110 | 110 | 109 | 109 | 110 |
| YPTB3750 | 1 | 47 | 1 | 1 | 1 | 1 | 1 | 1 | 47 | 1 | 1 | 1 | 1 | 1 | 1 | 1 | 1 | 47 | 1 | 1 | 1 | 1 | 1 | 1 | 1 | 1 | 1 | 1 | 1 | 1 | 1 | 1 | 1 | 1 | 1 |
| YPTB3760 | 1 | 1 | 1 | 1 | 1 | 1 | 1 | 1 | 1 | 1 | 1 | 72 | 1 | 1 | 1 | 1 | 1 | 1 | 1 | 1 | 1 | 1 | 1 | 1 | 1 | 1 | 1 | 1 | 1 | 1 | 1 | 1 | 1 | 1 | 1 |
| YPTB3764 | 1 | 1 | 1 | 74 | 1 | 1 | 1 | 1 | 1 | 74 | 74 | 1 | 1 | 1 | 1 | 74 | 1 | 1 | 1 | 1 | 1 | 1 | 1 | 1 | 1 | 1 | 1 | 1 | 1 | 1 | 1 | 1 | 74 | 74 | 1 |
| YPTB3786 | 1 | 141 | 1 | 1 | 1 | 1 | 1 | 137 | 141 | 143 | 1 | 137 | 145 | 145 | 1 | 143 | 145 | 138 | 1 | 1 | 1 | 1 | 1 | 1 | 145 | 145 | 1 | 1 | 1 | 1 | 1 | 145 | 1 | 1 | 177 |
| YPTB3790 | 1 | 1 | 1 | 92 | 1 | 1 | 1 | 1 | 1 | 92 | 92 | 1 | 1 | 1 | 1 | 92 | 1 | 92 | 1 | 1 | 1 | 1 | 1 | 1 | 1 | 1 | 1 | 1 | 1 | 1 | 1 | 1 | 92 | 92 | 1 |
| YPTB3791 | 94 | 96 | 94 | 97 | 1 | 94 | 94 | 99 | 96 | 97 | 97 | 94 | 94 | 94 | 94 | 97 | 94 | 1 | 94 | 94 | 94 | 94 | 94 | 94 | 94 | 94 | 94 | 94 | 94 | 94 | 94 | 94 | 97 | 97 | 94 |
| YPTB3840 | 1 | 117 | 1 | 117 | 1 | 116 | 1 | 117 | 117 | 117 | 117 | 1 | 116 | 116 | 116 | 117 | 116 | 115 | 1 | 1 | 116 | 116 | 116 | 1 | 116 | 116 | 116 | 116 | 116 | 116 | 116 | 116 | 117 | 117 | 1 |
| YPTB3847 | 1 | 1 | 1 | 93 | 1 | 1 | 1 | 1 | 1 | 93 | 93 | 92 | 1 | 1 | 1 | 93 | 1 | 1 | 1 | 1 | 1 | 1 | 1 | 1 | 1 | 1 | 1 | 1 | 1 | 1 | 1 | 1 | 93 | 93 | 1 |
| YPTB3914 | 91 | 94 | 91 | 95 | 1 | 1 | 91 | 96 | 94 | 95 | 95 | 90 | 1 | 1 | 1 | 95 | 1 | 1 | 91 | 91 | 1 | 1 | 1 | 91 | 1 | 1 | 1 | 1 | 1 | 1 | 1 | 1 | 95 | 95 | 91 |
| YPTB3915 | 99 | 99 | 99 | 99 | 99 | 99 | 99 | 99 | 99 | 99 | 99 | 99 | 99 | 99 | 99 | 99 | 99 | 99 | 99 | 99 | 99 | 99 | 99 | 99 | 99 | 99 | 99 | 99 | 99 | 99 | 99 | 99 | 99 | 99 | 99 |
| YPTB3916 | 137 | 141 | 137 | 140 | 1 | 1 | 137 | 192 | 141 | 143 | 140 | 136 | 1 | 1 | 1 | 143 | 1 | 138 | 137 | 137 | 1 | 1 | 1 | 137 | 1 | 1 | 1 | 1 | 1 | 1 | 1 | 1 | 140 | 140 | 137 |
| YPTB3920 | 1 | 81 | 1 | 83 | 1 | 1 | 1 | 1 | 81 | 83 | 83 | 80 | 1 | 1 | 1 | 83 | 1 | 81 | 1 | 1 | 1 | 1 | 1 | 1 | 1 | 1 | 1 | 1 | 1 | 1 | 1 | 1 | 83 | 83 | 1 |
| YPTB3921 | 1 | 61 | 1 | 1 | 1 | 1 | 1 | 1 | 61 | 1 | 1 | 1 | 1 | 1 | 1 | 1 | 1 | 1 | 1 | 1 | 1 | 1 | 1 | 1 | 1 | 1 | 1 | 1 | 1 | 1 | 1 | 1 | 1 | 1 | 1 |
| YPTB3940 | 129 | 133 | 129 | 134 | 1 | 1 | 129 | 136 | 133 | 134 | 134 | 128 | 1 | 1 | 1 | 134 | 1 | 130 | 129 | 129 | 1 | 1 | 1 | 129 | 1 | 1 | 1 | 1 | 1 | 1 | 1 | 1 | 134 | 134 | 129 |
| YPTB3942 | 95 | 98 | 95 | 98 | 1 | 1 | 95 | 98 | 98 | 98 | 98 | 1 | 1 | 1 | 1 | 98 | 1 | 96 | 95 | 95 | 1 | 1 | 1 | 95 | 1 | 1 | 1 | 1 | 1 | 1 | 1 | 1 | 98 | 98 | 95 |
| YPTB3945 | 1 | 1 | 1 | 1 | 1 | 1 | 1 | 1 | 1 | 1 | 1 | 1 | 1 | 1 | 1 | 1 | 1 | 1 | 1 | 1 | 1 | 1 | 1 | 1 | 1 | 1 | 1 | 1 | 1 | 1 | 1 | 1 | 1 | 1 | 1 |
| YPTB3946 | 1 | 1 | 1 | 1 | 1 | 1 | 1 | 1 | 1 | 1 | 1 | 1 | 1 | 1 | 1 | 1 | 1 | 39 | 1 | 1 | 1 | 1 | 1 | 1 | 1 | 1 | 1 | 1 | 1 | 1 | 1 | 1 | 1 | 1 | 1 |
| YPTB3947 | 1 | 29 | 1 | 1 | 1 | 1 | 1 | 1 | 29 | 1 | 1 | 1 | 1 | 1 | 1 | 1 | 1 | 28 | 1 | 1 | 1 | 1 | 1 | 1 | 1 | 1 | 1 | 1 | 1 | 1 | 1 | 1 | 1 | 1 | 1 |
| YPTB3950 | 1 | 1 | 1 | 104 | 1 | 1 | 1 | 99 | 1 | 103 | 104 | 1 | 1 | 133 | 1 | 103 | 133 | 100 | 1 | 1 | 1 | 1 | 1 | 1 | 1 | 1 | 1 | 1 | 1 | 1 | 1 | 1 | 104 | 104 | 1 |
| YPTB3960 | 1 | 1 | 1 | 1 | 1 | 1 | 1 | 1 | 1 | 1 | 1 | 1 | 1 | 1 | 1 | 1 | 1 | 91 | 1 | 1 | 1 | 1 | 1 | 1 | 1 | 1 | 1 | 1 | 1 | 1 | 1 | 1 | 1 | 1 | 1 |
| YPTB3961 | 1 | 97 | 1 | 1 | 1 | 1 | 1 | 1 | 97 | 1 | 1 | 1 | 1 | 1 | 1 | 1 | 1 | 97 | 1 | 125 | 1 | 1 | 1 | 125 | 1 | 1 | 1 | 1 | 1 | 1 | 1 | 1 | 1 | 1 | 1 |
| YPTB3965 | 1 | 118 | 1 | 122 | 1 | 1 | 1 | 123 | 118 | 122 | 122 | 119 | 1 | 1 | 1 | 122 | 1 | 120 | 1 | 1 | 1 | 1 | 1 | 1 | 1 | 1 | 1 | 1 | 1 | 1 | 1 | 1 | 122 | 122 | 1 |
| YPTB3966 | 1 | 1 | 1 | 53 | 1 | 1 | 1 | 1 | 1 | 53 | 53 | 1 | 1 | 1 | 1 | 53 | 1 | 54 | 1 | 1 | 1 | 1 | 1 | 1 | 1 | 1 | 1 | 1 | 1 | 1 | 1 | 1 | 53 | 53 | 1 |
| YPTB3969 | 101 | 101 | 101 | 130 | 1 | 1 | 101 | 101 | 101 | 103 | 130 | 1 | 1 | 1 | 1 | 103 | 1 | 131 | 101 | 101 | 1 | 1 | 1 | 101 | 1 | 1 | 1 | 1 | 1 | 1 | 1 | 1 | 130 | 130 | 101 |
| YPTB3970 | 1 | 1 | 1 | 1 | 1 | 1 | 1 | 1 | 1 | 1 | 1 | 1 | 1 | 1 | 1 | 1 | 1 | 46 | 1 | 1 | 1 | 1 | 1 | 1 | 1 | 1 | 1 | 1 | 1 | 1 | 1 | 1 | 1 | 1 | 1 |
| YPTB3971 | 1 | 1 | 1 | 1 | 1 | 1 | 1 | 1 | 1 | 1 | 1 | 1 | 1 | 1 | 1 | 1 | 1 | 1 | 1 | 1 | 1 | 1 | 1 | 1 | 1 | 1 | 1 | 1 | 1 | 1 | 1 | 1 | 1 | 1 | 1 |
| YPTB3973 | 1 | 1 | 1 | 1 | 1 | 1 | 1 | 1 | 1 | 1 | 1 | 1 | 1 | 1 | 1 | 1 | 1 | 1 | 1 | 1 | 1 | 1 | 1 | 1 | 1 | 1 | 1 | 1 | 1 | 1 | 1 | 1 | 1 | 1 | 1 |
