## Supplementary material for "First description of a *Yersinia pseudotuberculosis* clonal outbreak in France, confirmed using a new core genome multilocus sequence typing method": Table_S3

| cgMLST distance | IP32953 | IP32997 | IP32998 | IP33018 | IP33060 | IP33094 | IP33357 | IP33362 | IP33363 | IP33443 | IP33444 | IP33467 | IP33966 | IP34864 | IP35660 | IP37713 | IP37784 | IP39187 | IP40258 | IP41727 | IP32400 | IP32759 | IP32780 | IP31871 | IP32627 | IP32641 | IP32006 | IP43278 | IP43304 | IP31922 | IP43492 | IP43951 | IP43952 | IP43989 | IP44094 | IP44110 | IP44154 | IP44180 | IP44214 |
|---|---|---|---|---|---|---|---|---|---|---|---|---|---|---|---|---|---|---|---|---|---|---|---|---|---|---|---|---|---|---|---|---|---|---|---|---|---|---|---|
| IP32953 | 0 |
| IP32997 | 105 | 0 |
| IP32998 | 29 | 102 | 0 |
| IP33018 | 105 | 13 | 102 | 0 |
| IP33060 | 31 | 103 | 8 | 103 | 0 |
| IP33094 | 112 | 23 | 109 | 18 | 110 | 0 |
| IP33357 | 114 | 71 | 110 | 71 | 111 | 80 | 0 |
| IP33362 | 31 | 102 | 8 | 102 | 10 | 109 | 110 | 0 |
| IP33363 | 29 | 101 | 6 | 101 | 8 | 108 | 109 | 5 | 0 |
| IP33443 | 114 | 24 | 111 | 19 | 112 | 3 | 81 | 110 | 110 | 0 |
| IP33444 | 109 | 18 | 106 | 17 | 107 | 27 | 75 | 106 | 105 | 28 | 0 |
| IP33467 | 113 | 70 | 109 | 70 | 110 | 79 | 6 | 108 | 108 | 80 | 74 | 0 |
| IP33966 | 31 | 103 | 10 | 103 | 12 | 110 | 111 | 11 | 9 | 112 | 107 | 110 | 0 |
| IP34864 | 29 | 102 | 8 | 102 | 10 | 109 | 110 | 9 | 8 | 111 | 106 | 109 | 6 | 0 |
| IP35660 | 112 | 21 | 109 | 20 | 110 | 30 | 76 | 109 | 108 | 31 | 9 | 75 | 110 | 109 | 0 |
| IP37713 | 115 | 30 | 112 | 29 | 113 | 38 | 80 | 112 | 111 | 39 | 18 | 79 | 113 | 112 | 13 | 0 |
| IP37784 | 35 | 117 | 41 | 117 | 43 | 124 | 125 | 43 | 41 | 126 | 120 | 124 | 43 | 41 | 123 | 123 | 0 |
| IP39187 | 106 | 16 | 104 | 15 | 105 | 25 | 74 | 104 | 103 | 26 | 18 | 73 | 105 | 103 | 21 | 29 | 117 | 0 |
| IP40258 | 113 | 21 | 110 | 20 | 111 | 30 | 78 | 110 | 109 | 31 | 9 | 77 | 111 | 110 | 4 | 13 | 124 | 21 | 0 |
| IP41727 | 105 | 13 | 102 | 12 | 102 | 22 | 70 | 102 | 101 | 23 | 17 | 68 | 103 | 102 | 20 | 28 | 117 | 15 | 20 | 0 |
| IP32400 | 28 | 110 | 34 | 110 | 36 | 117 | 118 | 36 | 34 | 119 | 114 | 117 | 36 | 34 | 117 | 118 | 25 | 112 | 118 | 110 | 0 |
| IP32759 | 33 | 106 | 10 | 106 | 12 | 113 | 113 | 11 | 9 | 114 | 110 | 113 | 14 | 12 | 111 | 116 | 45 | 108 | 114 | 106 | 38 | 0 |
| IP32780 | 107 | 15 | 104 | 14 | 105 | 24 | 73 | 104 | 103 | 25 | 17 | 72 | 105 | 104 | 20 | 29 | 118 | 5 | 20 | 14 | 112 | 108 | 0 |
| IP31871 | 114 | 13 | 111 | 22 | 112 | 32 | 80 | 110 | 110 | 32 | 26 | 79 | 112 | 111 | 29 | 39 | 126 | 25 | 30 | 22 | 119 | 115 | 24 | 0 |
| IP32627 | 116 | 24 | 112 | 23 | 113 | 32 | 82 | 111 | 111 | 33 | 12 | 81 | 113 | 112 | 13 | 22 | 127 | 24 | 13 | 23 | 121 | 116 | 23 | 32 | 0 |
| IP32641 | 105 | 26 | 102 | 26 | 103 | 35 | 71 | 101 | 101 | 34 | 31 | 70 | 103 | 102 | 32 | 43 | 116 | 29 | 34 | 26 | 110 | 106 | 28 | 34 | 35 | 0 |
| IP32006 | 29 | 102 | 6 | 102 | 4 | 109 | 110 | 8 | 6 | 111 | 106 | 109 | 10 | 8 | 109 | 112 | 41 | 104 | 110 | 101 | 34 | 10 | 104 | 111 | 112 | 102 | 0 |
| IP43278 | 28 | 100 | 7 | 100 | 9 | 107 | 108 | 8 | 6 | 109 | 104 | 107 | 8 | 7 | 107 | 109 | 40 | 101 | 108 | 100 | 33 | 11 | 102 | 109 | 110 | 100 | 7 | 0 |
| IP43304 | 145 | 96 | 142 | 96 | 143 | 104 | 109 | 141 | 140 | 104 | 100 | 108 | 143 | 142 | 104 | 111 | 155 | 99 | 104 | 95 | 149 | 146 | 98 | 106 | 106 | 97 | 142 | 140 | 0 |
| IP31922 | 108 | 17 | 105 | 16 | 106 | 25 | 73 | 105 | 104 | 27 | 21 | 72 | 106 | 105 | 24 | 32 | 120 | 19 | 24 | 16 | 113 | 109 | 18 | 26 | 27 | 30 | 105 | 103 | 99 | 0 |
| IP43492 | 147 | 98 | 144 | 98 | 145 | 106 | 111 | 142 | 142 | 105 | 101 | 110 | 145 | 144 | 106 | 113 | 157 | 101 | 106 | 97 | 151 | 148 | 100 | 106 | 107 | 98 | 144 | 142 | 2 | 101 | 0 |
| IP43951 | 146 | 98 | 143 | 98 | 144 | 106 | 110 | 142 | 141 | 106 | 102 | 109 | 143 | 143 | 105 | 113 | 155 | 101 | 106 | 97 | 150 | 147 | 100 | 107 | 108 | 98 | 143 | 141 | 2 | 101 | 4 | 0 |
| IP43952 | 145 | 96 | 142 | 96 | 143 | 104 | 109 | 141 | 140 | 104 | 100 | 108 | 142 | 142 | 104 | 111 | 154 | 99 | 104 | 95 | 149 | 146 | 98 | 106 | 106 | 97 | 142 | 140 | 0 | 99 | 2 | 2 | 0 |
| IP43989 | 146 | 97 | 143 | 97 | 144 | 105 | 110 | 142 | 141 | 105 | 101 | 109 | 143 | 143 | 105 | 112 | 154 | 100 | 105 | 96 | 150 | 147 | 99 | 107 | 107 | 98 | 143 | 141 | 1 | 100 | 3 | 3 | 1 | 0 |
| IP44094 | 145 | 97 | 142 | 97 | 143 | 105 | 109 | 141 | 140 | 105 | 101 | 108 | 142 | 142 | 104 | 112 | 154 | 100 | 105 | 96 | 149 | 146 | 99 | 106 | 107 | 97 | 142 | 140 | 1 | 100 | 3 | 2 | 1 | 2 | 0 |
| IP44110 | 146 | 97 | 143 | 97 | 144 | 105 | 110 | 142 | 139 | 105 | 101 | 109 | 143 | 143 | 105 | 112 | 155 | 100 | 105 | 96 | 150 | 146 | 99 | 107 | 107 | 98 | 143 | 141 | 1 | 100 | 3 | 3 | 1 | 2 | 2 | 0 |
| IP44154 | 145 | 96 | 142 | 96 | 143 | 104 | 109 | 141 | 140 | 104 | 100 | 108 | 142 | 142 | 104 | 111 | 154 | 99 | 104 | 95 | 149 | 146 | 98 | 106 | 106 | 97 | 142 | 140 | 0 | 99 | 2 | 2 | 0 | 1 | 1 | 1 | 0 |
| IP44180 | 146 | 97 | 143 | 97 | 144 | 105 | 110 | 142 | 140 | 105 | 101 | 109 | 143 | 143 | 105 | 112 | 155 | 100 | 105 | 96 | 150 | 146 | 99 | 107 | 107 | 98 | 143 | 141 | 1 | 100 | 3 | 3 | 1 | 2 | 2 | 1 | 1 | 0 |
| IP44214 | 148 | 99 | 145 | 99 | 146 | 107 | 112 | 143 | 143 | 106 | 103 | 111 | 145 | 145 | 107 | 114 | 157 | 102 | 107 | 98 | 152 | 149 | 101 | 108 | 108 | 99 | 145 | 143 | 3 | 102 | 4 | 5 | 3 | 4 | 4 | 4 | 3 | 4 | 0 |

| SNP distance | IP32953 | IP32997 | IP32998 | IP33018 | IP33060 | IP33094 | IP33357 | IP33362 | IP33363 | IP33443 | IP33444 | IP33467 | IP33966 | IP34864 | IP35660 | IP37713 | IP37784 | IP39187 | IP40258 | IP41727 | IP32400 | IP32759 | IP32780 | IP31871 | IP32627 | IP32641 | IP32006 | IP43278 | IP43304 | IP31922 | IP43492 | IP43951 | IP43952 | IP43989 | IP44094 | IP44110 | IP44154 | IP44180 | IP44214 |
|---|---|---|---|---|---|---|---|---|---|---|---|---|---|---|---|---|---|---|---|---|---|---|---|---|---|---|---|---|---|---|---|---|---|---|---|---|---|---|---|
| IP32953 | 0 |
| IP32997 | 243 | 0 |
| IP32998 | 65 | 222 | 0 |
| IP33018 | 245 | 46 | 224 | 0 |
| IP33060 | 69 | 226 | 18 | 228 | 0 |
| IP33094 | 244 | 47 | 223 | 43 | 227 | 0 |
| IP33357 | 267 | 162 | 246 | 164 | 250 | 163 | 0 |
| IP33362 | 69 | 226 | 18 | 228 | 22 | 227 | 250 | 0 |
| IP33363 | 69 | 226 | 18 | 228 | 22 | 227 | 250 | 0 | 0 |
| IP33443 | 244 | 47 | 223 | 43 | 227 | 4 | 163 | 227 | 227 | 0 |
| IP33444 | 243 | 46 | 222 | 48 | 226 | 47 | 162 | 226 | 226 | 47 | 0 |
| IP33467 | 268 | 163 | 247 | 165 | 251 | 164 | 9 | 251 | 251 | 164 | 163 | 0 |
| IP33966 | 71 | 228 | 20 | 230 | 24 | 229 | 252 | 24 | 24 | 229 | 228 | 253 | 0 |
| IP34864 | 71 | 228 | 20 | 230 | 24 | 229 | 252 | 24 | 24 | 229 | 228 | 253 | 12 | 0 |
| IP35660 | 243 | 46 | 222 | 48 | 226 | 47 | 162 | 226 | 226 | 47 | 22 | 163 | 228 | 228 | 0 |
| IP37713 | 242 | 45 | 221 | 47 | 225 | 46 | 161 | 225 | 225 | 46 | 21 | 162 | 227 | 227 | 5 | 0 |
| IP37784 | 85 | 266 | 88 | 268 | 92 | 267 | 290 | 92 | 92 | 267 | 266 | 291 | 94 | 94 | 266 | 265 | 0 |
| IP39187 | 233 | 36 | 212 | 38 | 216 | 37 | 152 | 216 | 216 | 37 | 32 | 153 | 218 | 218 | 32 | 31 | 256 | 0 |
| IP40258 | 245 | 48 | 224 | 50 | 228 | 49 | 164 | 228 | 228 | 49 | 24 | 165 | 230 | 230 | 8 | 7 | 268 | 34 | 0 |
| IP41727 | 244 | 43 | 223 | 49 | 227 | 48 | 163 | 227 | 227 | 48 | 47 | 164 | 229 | 229 | 47 | 46 | 267 | 37 | 49 | 0 |
| IP32400 | 83 | 264 | 86 | 266 | 90 | 265 | 288 | 90 | 90 | 265 | 264 | 289 | 92 | 92 | 264 | 263 | 72 | 254 | 266 | 265 | 0 |
| IP32759 | 71 | 228 | 20 | 230 | 24 | 229 | 252 | 24 | 24 | 229 | 228 | 253 | 26 | 26 | 228 | 227 | 94 | 218 | 230 | 229 | 92 | 0 |
| IP32780 | 235 | 38 | 214 | 40 | 218 | 39 | 154 | 218 | 218 | 39 | 34 | 155 | 220 | 220 | 34 | 33 | 258 | 4 | 36 | 39 | 256 | 220 | 0 |
| IP31871 | 254 | 33 | 233 | 57 | 237 | 58 | 173 | 237 | 237 | 58 | 57 | 174 | 239 | 239 | 57 | 56 | 277 | 47 | 59 | 54 | 275 | 239 | 49 | 0 |
| IP32627 | 248 | 51 | 227 | 53 | 231 | 52 | 167 | 231 | 231 | 52 | 27 | 168 | 233 | 233 | 21 | 20 | 271 | 37 | 23 | 52 | 269 | 233 | 39 | 62 | 0 |
| IP32641 | 425 | 250 | 404 | 252 | 408 | 251 | 344 | 408 | 408 | 251 | 250 | 345 | 410 | 410 | 250 | 249 | 448 | 240 | 252 | 251 | 446 | 410 | 242 | 261 | 255 | 0 |
| IP32006 | 65 | 222 | 14 | 224 | 6 | 223 | 246 | 18 | 18 | 223 | 222 | 247 | 20 | 20 | 222 | 221 | 88 | 212 | 224 | 223 | 86 | 20 | 214 | 233 | 227 | 404 | 0 |
| IP43278 | 69 | 226 | 18 | 228 | 22 | 227 | 250 | 22 | 22 | 227 | 226 | 251 | 22 | 22 | 226 | 225 | 92 | 216 | 228 | 227 | 90 | 24 | 218 | 237 | 231 | 408 | 18 | 0 |
| IP43304 | 723 | 620 | 702 | 622 | 706 | 621 | 644 | 706 | 706 | 621 | 620 | 645 | 706 | 708 | 620 | 619 | 746 | 610 | 622 | 621 | 744 | 708 | 612 | 631 | 625 | 800 | 702 | 706 | 0 |
| IP31922 | 248 | 47 | 227 | 53 | 231 | 52 | 167 | 231 | 231 | 52 | 51 | 168 | 233 | 233 | 51 | 50 | 271 | 41 | 53 | 46 | 269 | 233 | 43 | 58 | 56 | 255 | 227 | 231 | 625 | 0 |
| IP43492 | 724 | 621 | 703 | 623 | 707 | 622 | 645 | 707 | 707 | 622 | 621 | 646 | 707 | 709 | 621 | 620 | 747 | 611 | 623 | 622 | 745 | 709 | 613 | 632 | 626 | 801 | 703 | 707 | 1 | 626 | 0 |
| IP43951 | 723 | 620 | 702 | 622 | 706 | 621 | 644 | 706 | 706 | 621 | 620 | 645 | 706 | 708 | 620 | 619 | 746 | 610 | 622 | 621 | 744 | 708 | 612 | 631 | 625 | 800 | 702 | 706 | 0 | 625 | 1 | 0 |
| IP43952 | 723 | 620 | 702 | 622 | 706 | 621 | 644 | 706 | 706 | 621 | 620 | 645 | 706 | 708 | 620 | 619 | 746 | 610 | 622 | 621 | 744 | 708 | 612 | 631 | 625 | 800 | 702 | 706 | 0 | 625 | 1 | 0 | 0 |
| IP43989 | 723 | 620 | 702 | 622 | 706 | 621 | 644 | 706 | 706 | 621 | 620 | 645 | 706 | 708 | 620 | 619 | 746 | 610 | 622 | 621 | 744 | 708 | 612 | 631 | 625 | 800 | 702 | 706 | 0 | 625 | 1 | 0 | 0 | 0 |
| IP44094 | 723 | 620 | 702 | 622 | 706 | 621 | 644 | 706 | 706 | 621 | 620 | 645 | 706 | 708 | 620 | 619 | 746 | 610 | 622 | 621 | 744 | 708 | 612 | 631 | 625 | 800 | 702 | 706 | 0 | 625 | 1 | 0 | 0 | 0 | 0 |
| IP44110 | 723 | 620 | 702 | 622 | 706 | 621 | 644 | 706 | 706 | 621 | 620 | 645 | 706 | 708 | 620 | 619 | 746 | 610 | 622 | 621 | 744 | 708 | 612 | 631 | 625 | 800 | 702 | 706 | 0 | 625 | 1 | 0 | 0 | 0 | 0 | 0 |
| IP44154 | 723 | 620 | 702 | 622 | 706 | 621 | 644 | 706 | 706 | 621 | 620 | 645 | 706 | 708 | 620 | 619 | 746 | 610 | 622 | 621 | 744 | 708 | 612 | 631 | 625 | 800 | 702 | 706 | 0 | 625 | 1 | 0 | 0 | 0 | 0 | 0 | 0 |
| IP44180 | 723 | 620 | 702 | 622 | 706 | 621 | 644 | 706 | 706 | 621 | 620 | 645 | 706 | 708 | 620 | 619 | 746 | 610 | 622 | 621 | 744 | 708 | 612 | 631 | 625 | 800 | 702 | 706 | 0 | 625 | 1 | 0 | 0 | 0 | 0 | 0 | 0 | 0 |
| IP44214 | 725 | 622 | 704 | 624 | 708 | 623 | 646 | 708 | 708 | 623 | 622 | 647 | 708 | 710 | 622 | 621 | 748 | 612 | 624 | 623 | 746 | 710 | 614 | 633 | 627 | 802 | 704 | 708 | 2 | 627 | 3 | 2 | 2 | 2 | 2 | 2 | 2 | 2 | 0 |
